# Investigating Coronary Artery Disease methylome through targeted bisulfite sequencing

**DOI:** 10.1101/621789

**Authors:** Subhoshree Ghose, Sourav Ghosh, Vinay Singh Tanwar, Priya Tolani, Anju Sharma, Nitin Bhardwaj, KV Shamsudheen, Ankit Verma, Rijith Jayarajan, Sridhar Sivasubbu, Vinod Scaria, Sandeep Seth, Shantanu Sengupta

## Abstract

**Background:** Gene environment interactions leading to epigenetic alterations play pivotal role in the pathogenesis of Coronary Artery Disease (CAD). Altered DNA methylation is one such epigenetic factor that could lead to altered disease etiology. In this study, we comprehensively identified methylation sites in several genes that have been previously associated with young CAD patients.

**Methods:** The study population consisted of 42 healthy controls and 33 young CAD patients (age group < 50 years). We performed targeted bisulfite sequencing of promoter as well as genic regions of several genes in various pathways like cholesterol synthesis and metabolism, endothelial dysfunction, apoptosis, which are implicated in the development of CAD.

**Results:** We observed that the genes like *GALNT2, HMGCR* were hypermethylated in the promoter whereas *LDLR* gene promoter was hypomethylated indicating that intracellular LDL uptake was higher in CAD patients. Although *APOA1* did not show significant change in methylation but *APOC3* and *APOA5* showed variation in methylation in promoter and exonic regions. Glucokinase (*GCK*) and endothelial nitric oxide synthase 3 *(NOS3)* were hyper methylated in the promoter. Genes involved in apoptosis *(BAX/BCL2/AKT2)* and inflammation (*PHACTR1/LCK*) also showed differential methylation between controls and CAD patients.

**Conclusions:** This study is unique because it highlights important gene methylation alterations which might predict the risk of young CAD in Indian population. Large scale studies in different populations would be important for validating our findings and understanding the epigenetic events associated with CAD.

## Introduction

Coronary Artery Disease (CAD), a complex multifactorial disease thought to occur via complex amalgamation between genetic and environmental factors, is manifested via dyslipidemia, obesity, raised blood pressure, aortic constrictions and eventual myocardial infarction. According to a WHO report, it was estimated that between 2000–2012, deaths due to CAD were about 56 million worldwide (Pagidipati and Gaziano 2013). This statistic is expected to alarmingly increase due to sedentary lifestyle and poor nutrition status in developing countries. The burden of CAD is huge worldwide and is identified to be one of the top five causes of death (Gupta, Guptha et al. 2012). A number of modifiable (smoking, diabetes, physical inactivity, hypertriglyceridemia, hypertension, obesity) and non-modifiable risk factors (age, gender, family history) have been attributed to disease pathophysiology (Huma, Tariq et al. 2012). Over the years, genome wide association (GWAS) studies on several populations have identified a significant number of single nucleotide polymorphisms (SNPs) located in genes and their vicinity which might play a role in disease progression. However, these SNPs account only for 10-15% of the disease risk (Consortium 2011; Deloukas, Kanoni et al. 2013) (Yamada, Yasukochi et al. 2018). The largest CAD GWAS study (CARDIoGRAMplusC4D) revealed approximately 46 susceptibility loci and 104 suggestive loci to be significantly associated with the disease (Deloukas, Kanoni et al. 2013) In a recent study, variants in the genes involved in actin remodeling are reported to be associated with LDL-C levels and CAD (Siewert and Voight 2018). Additionally, common variants in the CDKN2B-AS1 region have been reported to influence lipid metabolism and might be associated with CAD in Turkish population (Temel and Ergoren 2019). Exome wide association studies (EWAS) conducted on Japanese population identified 21 genes and 5 new chromosomal regions which were individual determinants of susceptibility towards early onset of CAD (Yamada, Yasukochi et al. 2018). Although a handful of studies have highlighted the importance of genetic variants in influencing lipid parameter but their underlying role in disease manifestation has not been elucidated well.

For complex diseases like CAD, both genetic and environmental factors contribute to the disease etiology where gene-environment interactions are thought to play a major role. Thus, it is not surprising that attempts have been made towards identifying epigenetic modulators especially DNA methylation in the context of CAD. DNA methylation is a stable modification which involves the addition of a methyl group to the fifth carbon position of cytosine base in CpG dinucleotides of the mammalian genome by DNA methyltransferases (DNMTs) and known to regulate gene expression (Vaissiere and Miller 2011). Several risk factors of CAD for example tobacco smoking has been reported to alter DNA methylation status of a few candidate genes involved in atherosclerosis (Steenaard, Ligthart et al. 2015). Moreover, epigenome wide association studies performed in different populations highlight the importance of methylation regulation in cardiovascular disease development (Nakatochi, Ichihara et al. 2017; Yamada, Horibe et al. 2018). A recent study performed on Chinese population affected with Acute Coronary Syndrome (ACS) highlighted some DNA methylation based markers relevant to several pathways like chemotaxis, apoptosis, thrombosis and atherogenic signaling (Li, Zhu et al. 2017). Interestingly, in French Canadian founder population, it was reported that DNA hypomethylation in the promoter region of TNNT1 gene was directly associated with dyslipidemia and the risk for CAD (Guay, Legare et al. 2016). Moreover, leukocyte LINE-1 methylation, a surrogate for global DNA methylation has been identified as a predictor of myocardial infarction and cardiovascular risk in men of Samoan islander population (Cash, McGarvey et al. 2011).

We have earlier shown that global DNA methylation was much higher in CAD patients in India which was also associated with hyperhomocystenemia, an independent risk factor for CAD (Sharma, Kumar et al. 2008; Sharma, Garg et al. 2014). Hyperhomocysteinemia is prevalent in India due to wide spread deficiency of vitamin B_12_, presumably due to adherence to strict vegetarian diet. We also showed low vitamin B_12_ levels were associated with CAD in Indian population (Kumar, Garg et al. 2009; Basak, Tanwar et al. 2016).

Epidemiological reports have documented high incidences of CAD in India which has grown enormously in the past 60 years (Gupta, Mohan et al. 2016). It is anticipated that by 2030, almost 60% of deaths due to CAD will be in India. Notably, CAD occurs at least a decade earlier in Indians than western countries. This was also confirmed in a recent report by. *Chaudhary et al* where it was shown that CAD manifests at an early age (< 40 years) in India. One of the reasons for this would be differences in methylation profile due to unique dietary habits in Indian population. However, to the best of our knowledge there are no comprehensive study in Indian population where alteration of DNA methylation has been looked at specifically in young CAD patients.

Therefore, in the current study, we analyzed the DNA methylation status of several genes that have been implicated in CAD in young individuals (25-50 years) to understand the role of DNA methylation in developing premature CAD.

## Material and Methods

### Study population and biochemical parameter measurement

We performed a cross sectional study to examine the DNA methylation status of promoter and gene body region of several genes associated with CAD in 42 healthy controls and 33 young CAD subjects. CAD patients were recruited from All India Institute of Medical Sciences (AIIMS), New Delhi following coronary angiography and the healthy controls were from the general population. The study was undertaken in accordance with the Principles of the Helsinki Declaration and was approved by the ethical committee of both CSIR-IGIB & AIIMS, New Delhi. All the participants were males between the age group (25-50) years. Participants taking anti-hypertensive or anti diabetic medications were excluded from the study. These healthy individuals were neither having any family history of cardiovascular disease nor reported of any chest pain or obstruction. For the current study, control and CAD patient samples were randomly screened from a group of samples previously collected from National Capital Region (NCR) for a separate study (Sharma, Garg et al. 2014; Basak, Tanwar et al. 2016). Blood samples were collected from all these individuals and were centrifuged at 1200 rpm for 20 minutes at 4°C. High quality genomic DNA was isolated from blood samples using the modified salting out method (Sharma, Garg et al. 2014) from all the 75 subjects and their concentration was estimated using fluorometric quantitation method using Qubit ds DNA BR kit. (Qubit fluorometer 2.0, Invitrogen).

## DNA methylation analysis

### Methylation probe design

We reviewed literature from the year 2003-2013, and retrieved information about the genes which were reported to be associated with CAD in different populations. A total of 48 genes were chosen through literature mining and their promoter and gene body region were selected for biotinylated probe design using Sure Design advanced online tool (https://earray.chem.agilent.com/suredesign). Some of the genes were already reported to have altered methylation in CAD patients in other populations. Mutations or polymorphism in some of these genes have also been reported with CAD earlier. The list of genes is provided in Supplementary Table 1. Repetitive regions were also included in the design for genes like *CDKN2BAS, CDKN2B-AS1, LDLR and APOA1* cluster. The overall probe size was 497.225 kbp corresponding to a total of 8461 probes.

### Sample preparation and targeted bisulfite sequencing

About 3 μg of good quality genomic DNA in 50 µl was fragmented using Covaris (S series S220) to an estimated size of 100-150 bp. Fragmentation size was determined using DNA 1000 Kit (Agilent p/n 5067-1504) according to manufacturer’s instructions. Fragmented DNA was then end repaired using end repair mix followed by AMPure bead based purification. End repaired libraries had an average size of 125-175 bp. Further, end repaired libraries were subjected to dA tailing where a single adenine base “A” was added to the 3’ end of the library and another round of AMPure bead based purification was performed. Methylated adapter ligation was performed on the dA tailed DNA using ligation master mix provided in the kit. Purification after this step was done immediately to avoid self-ligation of adapters. The quality of the end repaired and adapter ligated libraries were checked using DNA 1000 kit in Bioanalyzer (Supplementary fig 3). Following methylated adapter ligation, the size increased to 200-300 bp as expected (Supplementary fig 3). The adapter ligated libraries were then subjected to hybridization with the custom designed RNA bait library (Agilent Technologies) at 65°C for 24 hours followed by capture using streptavidin coated magnetic beads. Hybridized libraries were purified using magnetic bead based purification method. Finally, captured DNA was modified using EZ Gold DNA methylation kit (Zymo research, CA, USA.) After bisulfite conversion, desulfonation was performed and captured bisulfite converted DNA was eluted in nuclease free water. In the final step, all the 75 bisulfite converted libraries were indexed using 75 unique 8 bp Sure Select XT indexes (A01-H012 provided in Agilent Sure Select Methyl Seq Target Enrichment Kit) and PCR amplified for 8 cycles according to the following conditions: 95°C for 2 mins for initial denaturation; 95°C for 30 sec, 60°C for 30 sec, 72°C for 30 sec for amplification; 72°C for 7 min; 4°C hold. The quality of the libraries were then checked in Bioanalyzer using Agilent ds DNA High Sensitivity Kit following manufacturer’s guidelines. Size of the final libraries were between 200-300 bp with unique single indexes (Supplementary fig 3). The indexed libraries were quantified using Qubit High sensitivity DNA kit (Qubit HS). All the indexed libraries were then diluted to 5 nM concentration and combined into a single pool which was used for sequencing.

### Sequencing and downstream analysis

All the 75 libraries were diluted to 5 nM concentration and the indexed libraries were mixed into a pool. The pool was diluted to 15 pM and loaded onto the flow cell. Sequencing was performed in paired end mode (150×2) and run for 150 cycles in HiSeq 2500 (Illumina, CA. USA). Raw targeted bisulfite sequencing reads were initially checked for QC and then filtered through FastQC and low quality reads were removed. Adapter trimming was done by Trimmomatic pipeline to remove adapter contaminations and low quality reads. High quality paired end reads were then aligned against human genome (hg19 genome assembly downloaded from UCSC) using Bismark tool from Babraham bioinformatics. Alignment percentage was calculated considering both uniquely and multiple aligned reads. Further read coverage was calculated using the formula: (Total reads*read length)/targeted capture region. The ‘SAM’ files generated after alignment, were then used as input for methylation extractor component in Bismark tool, which generated genome wide cytosine reports. The cytosine report file gave methylation status of each cytosine in three different sequence context, i.e. CpG, CHG and CHH context where ‘H’ is either of the bases A, T, or C. For further downstream analysis only the CpG methylation was considered from both the strands for all the genes included in the study. A total of 311979 unique C & G coordinates in 139 regions, were subjected to downstream analysis of which 104690 sites were obtained after selecting the read coverage and methylation percentage cut-off (read coverage ≥ 5 and methylation percentage ≥ 20%) (Supplementary fig 2). Out of these, about 6400 coordinates were reported in at least 2 samples (Supplementary Table 3). Unpaired Student’s t-test was performed on these coordinates to identify the differentially methylated CpG sites (p < 0.05). We found 260 such CpG sites, associated with 45 genes to be significantly (p-value < 0.05) differentially methylated between controls and CAD patients (Supplementary Table 4). Further those CpG sites harboring significant differential methylation were mapped back to their respective genomic locations using in-house PERL programme.

### Results

The demographic characteristics of the individuals included in the study are provided in Table 1. The average age of controls and cases were 40.7 and 42.1 years respectively indicating that relatively young individuals were considered for this study.

Targeted bisulfite sequencing of 48 genes (details provided in Supplementary Table 1) were performed in 42 control and 33 Coronary Artery Disease (CAD) patients. The raw sequencing reads were mapped to the reference human genome (hg19) with a median mapping efficiency of approximately 60% in both cases and controls, with more than 30X read coverage (Figure. 1). The raw sequencing read count, alignment percentage and read coverage for all the 75 individuals recruited in the study are shown in Supplementary Table 2.

**Figure 1:**
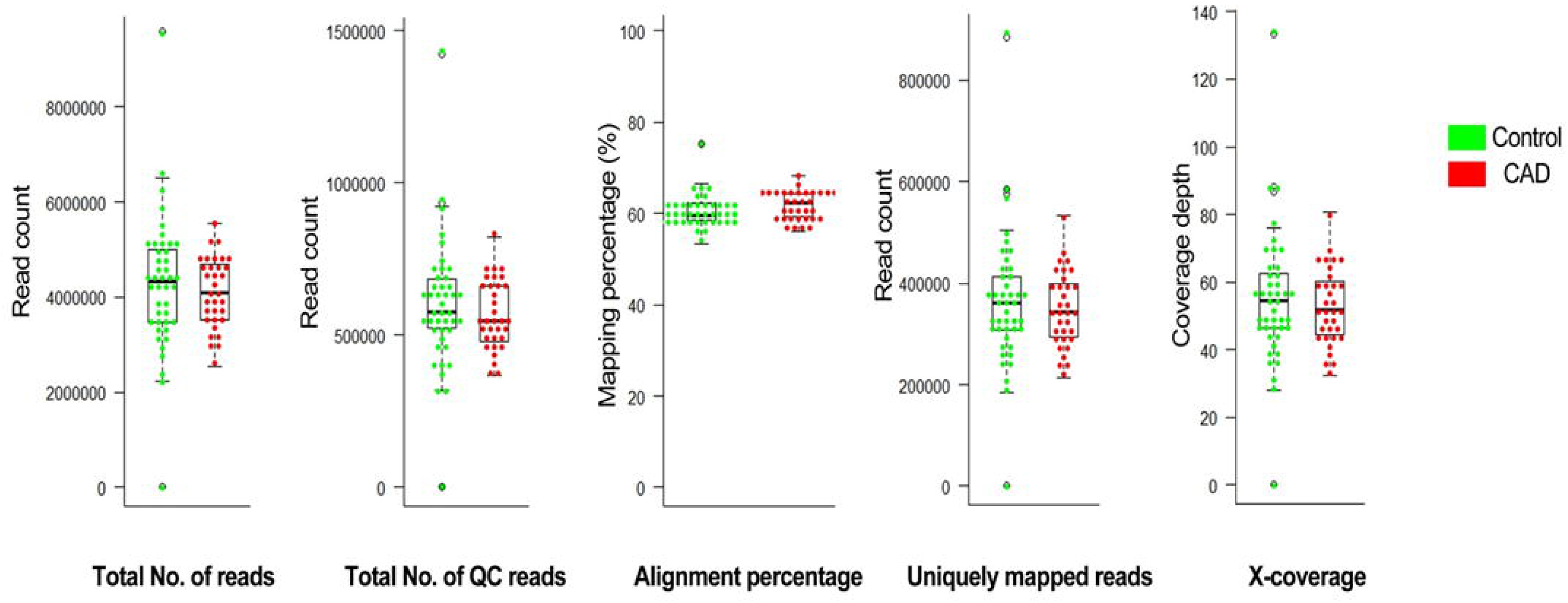
Sequencing statistics (Total no. of reads, QC reads, alignment percentage, and uniquely mapped reads and X coverage) of all the 76 subjects recruited in the study

**Figure 2:**
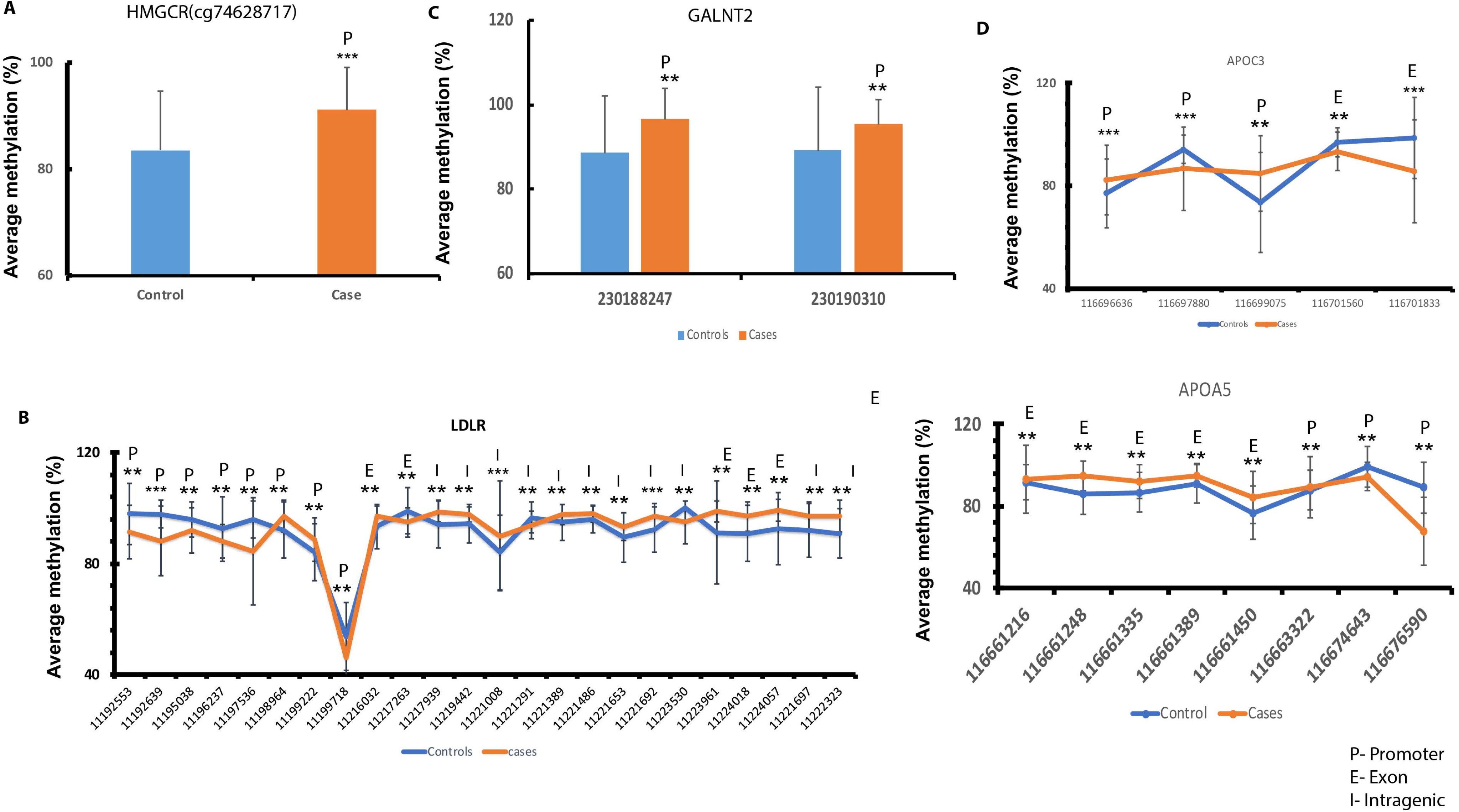
Methylation status of CpG present in the *HMGCR* /GALNT2/APOC3/LDLR/APOA5 gene. Y axis represents average percentage methylation and X axis represents different CpG sites. (P: Promoter; E: Exon; I: Intragenic); ** indicates p value < 0.01; *** indicates p value < 0.001

### Impairment of cholesterol biosynthesis and metabolism in CAD patients

One of the mechanisms proposed for the manifestation of CAD is accumulation of intracellular cholesterol, which could be due to increased synthesis or intracellular import of cholesterol or reduced intracellular cholesterol efflux. The rate limiting step of cholesterol biosynthesis is catalyzed by the enzyme 3-Hydroxy-3-Methylglutaryl-CoA-reductase (*HMGCR*) which exhibited hypermethylation in the promoter region. (Figure 2A) indicating less cholesterol synthesis. This is in agreement with a previous study by Peng et. al. (2014) where they also reported hypermethylation of *HMGCR* promoter region (Peng, Wang et al. 2014). However, we also observed that 24 CpG sites falling in the *LDLR* gene to be significantly differentially methylated in CAD patients. The promoter region of *LDLR* was hypomethylated (average methylation of 87% in controls and 84% in cases, p < 0.05), while the exonic regions were hypermethylated (Figure 2B) in CAD patients (average methylation of 97% in cases and 94% in controls, p<0.05), suggesting increased expression of LDLR. These results indicate lower synthesis of cholesterol but higher intracellular cholesterol uptake in CAD patients.

On the other hand, although there was no significant difference in methylation of the promoter or exonic regions of *APOA1*, an important mediator of cholesterol efflux, we found hypermethylation in the promoter of *GALNT2* (Figure 2C) gene indicating lower expression. A loss of function of *GALNT2* (Khetarpal, Schjoldager et al. 2016) gene has been reported to reduce the activity of phospholipid transfer protein (PLTP) thereby affecting the cholesterol efflux pathway leading to low HDL levels. We indeed found significantly low HDL levels in CAD patients (29.2 mg/dl) as compared to controls (43.5 mg/dl). Besides, one of the targets of *GALNT2* is *APOC3*. Interestingly, the exons of *APOC3* were hypomethylated and 2 of the 3 differentially methylated CpGs in the promoter region were hypermethylated (Figure 2D) suggesting low levels of ApoC3 in CAD patients. Low ApoC3 levels have been reported to be associated with low triglyceride levels which we have also observed in the plasma of CAD patients (Crosby, Peloso et al. 2014). We further observed that promoter of *APOA5* was hypomethylated and exons were hypermethylated which could result in low triglyceride levels in CAD patients (Figure 2E). Consistent with this the levels of triglyceride in the CAD patients were lower (96.9 mg/dl) than controls (142.9 mg/dl).

### Hyperglycemia leading to endothelial dysfunction and inflammation

Elevated blood glucose levels have been an important risk factor for CAD. We found that the promoter region of the gene *GCK* (glucokinse), which plays a major role in converting glucose to glucose 6 phosphate, is hypermethylated (Figure 3A) which could be responsible for lowered expression of GCK and increased blood glucose levels in CAD patients (110 mg/dl) as compared to controls (96 mg/dl) found in this study. Interestingly, the gene activating transcription factor 3 (*ATF3*) which is known to transcriptionally suppress the expression of glucokinase (*GCK*) was observed to be hypomethylated in the 5’UTR region (cg212739706) and hypermethylated in the exon (cg212793952) which could lead to increased ATF expression and low GCK levels. (Figure 3B).

**Figure 3:**
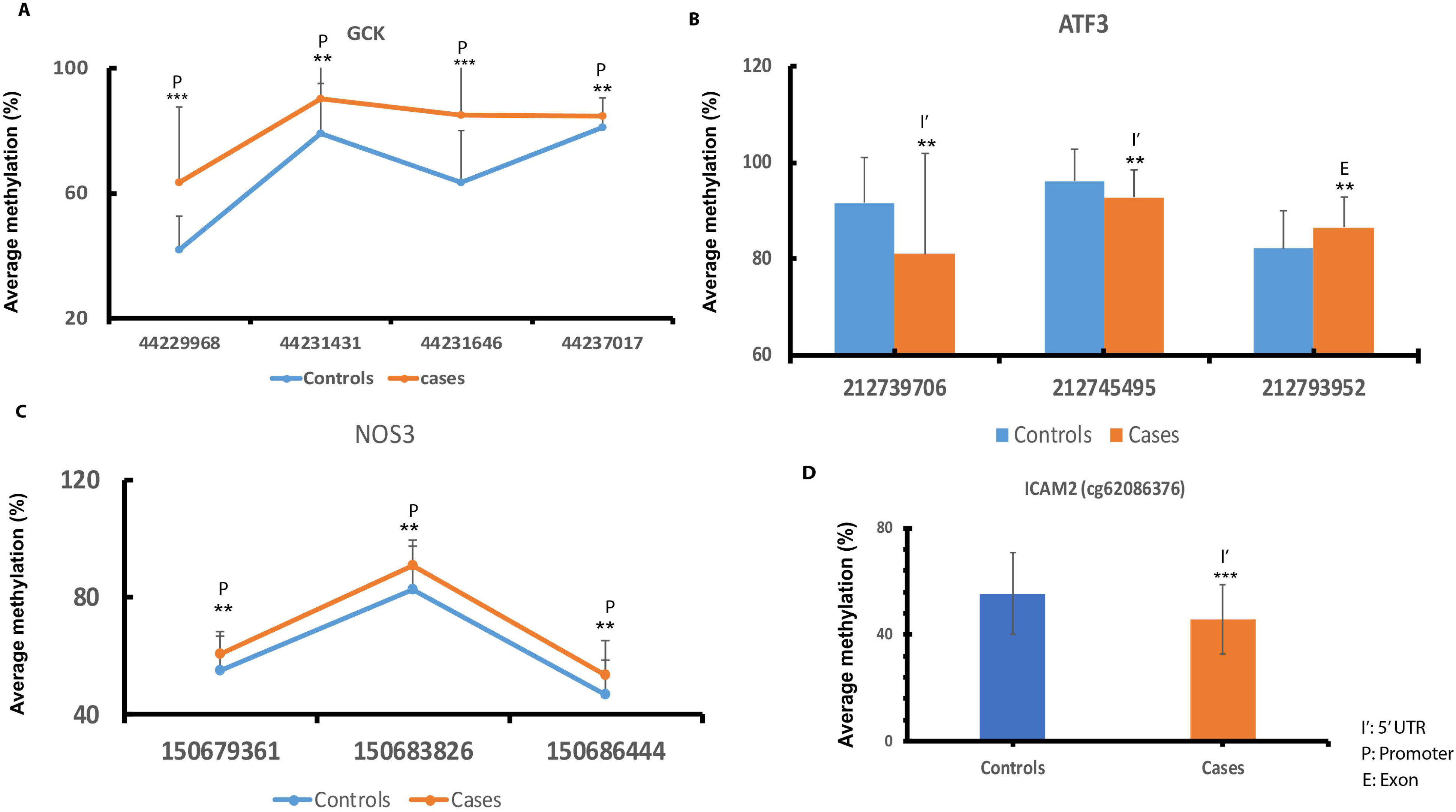
Methylation status of the genes involved in hyperglycemia and vasoconstriction GCK/ATF3/NOS3/ICAM2. P: Promoter, E: Exon; I’: 5’ UTR. ** indicates p value < 0.01; *** indicates p value < 0.001

It is known that endothelial cells exposed to hyperglycemic conditions show decreased NO production, increased levels of adhesion molecules accompanied by inflammation and also increased levels of apoptosis (Funk, Yurdagul et al. 2012). This is also suggestive from our results as we found hypermethylation in the promoter of endothelial nitric oxide synthase (*eNOS*) gene responsible for NO production in the endothelial cells (Figure 3C) which could potentially lead to reduced levels of NO and vasoconstriction in the arteries. We also observed hypomethylation in the 5’ UTR region of cell adhesion molecule *ICAM2* (cg62086376) implying increased adhesion and aggregation processes occurring during atherosclerosis (Figure 3D).

It is also known that endothelial dysfunction is responsible for inducing cell apoptosis via inhibition of NO (van den Oever, Raterman et al. 2010) and increased apoptosis was found in CAD patients (Kaplan and Demircan 2018). This leads us to assess the level of methylation in the genes responsible for causing apoptosis in CAD patients. The pro-apoptotic gene *BAX* promoter (cg49453159) was hypomethylated (Figure A) whereas anti-apoptotic *BCL2* gene promoter (cg60995889) was hypermethylated in CAD patients (Figure 4B). Interestingly, we observed altered methylation in the *LCK* (Src family of tyrosine kinase) gene, where 5’ UTR region was hypomethylated (cg32739493) and exonic region (cg32740607) was hypermethylated (Figure 4C) indicating possible upregulation of the *LCK* gene. Upregulation of LCK is indicative of activated T cell signaling which also leads to activation of pro inflammatory molecule NF-Kβ. Increased expression of LCK is associated with mitochondrial apoptosis(Samraj, Stroh et al. 2006). Besides *AKT2*, a phosphoinositide dependent serine threonine kinase showed hypomethylation in the promoter region (cg40795616, cg40794901) suggesting positive regulation of pro inflammatory factor NF-Kβ (Figure 4D).

**Figure 4:**
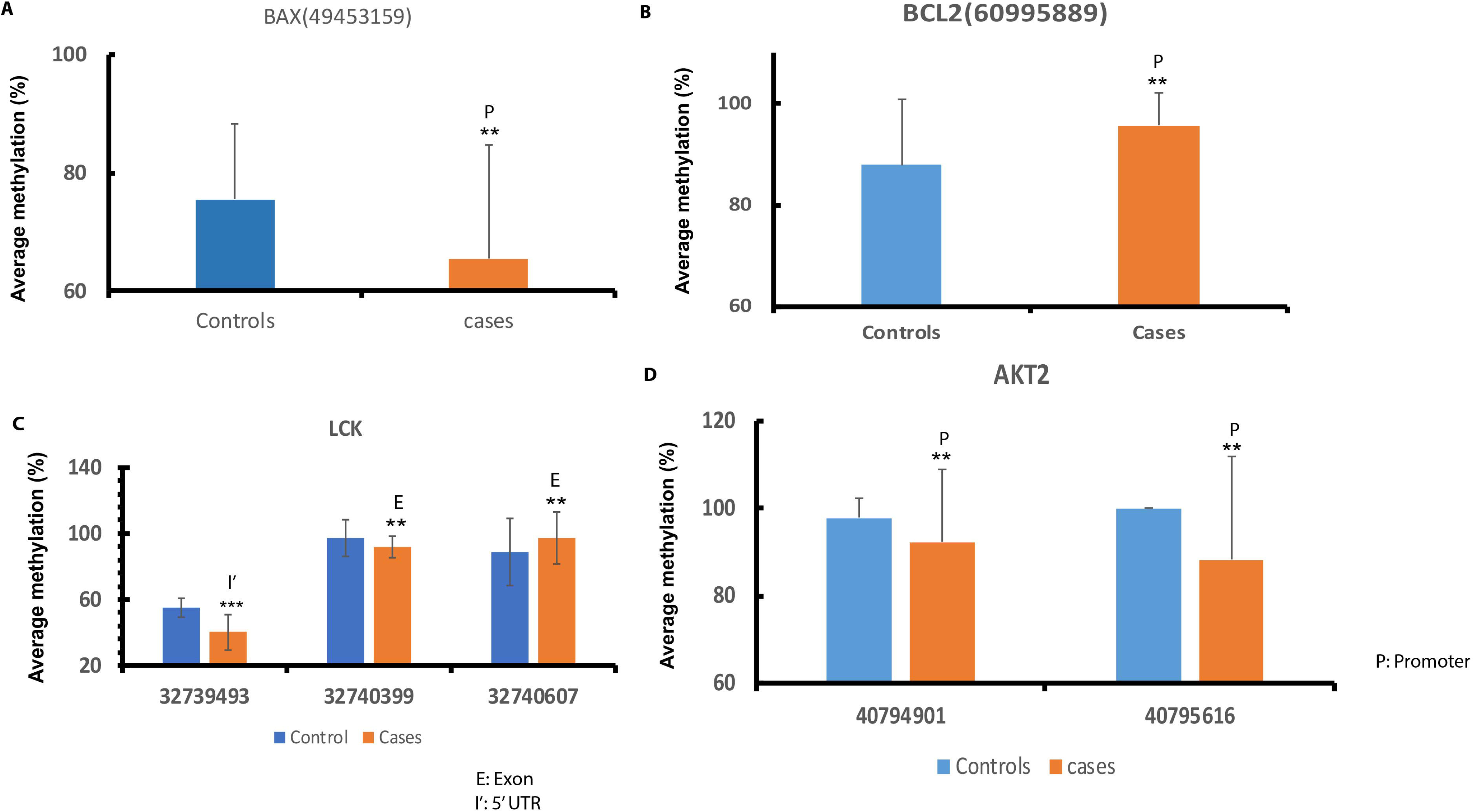
Methylation status of genes involved in apoptosis and inflammation (*BAX/BCL2/LCK/AKT2*). P: Promoter, E: Exon; I’: 5’ UTR, ** indicates p value < 0.01; *** indicates p value < 0.001.

### Genes involved in extracellular matrix remodeling and platelet aggregation were differentially methylated in CAD

We found that phosphatase and actin regulator 1 (*PHACTR1)* gene harbored hypermethylation in the promoter (cg12711037, cg12715861) and hypomethylation in the exon (cg12717069) (Figure 5A) suggestive of lower expression of PHACTR1. Down regulation of *PHACTR1* has been reported to increase expression of matrix-metalloproteinase regulators and pro inflammatory factors (Jarray, Pavoni et al. 2015). Matrix metalloproteinase regulators were altered in acute coronary patients (Liu, Sun et al. 2006) and in this study we observed tissue inhibitor of matrix metalloproteinase 3 (*TIMP3*) to be hypomethylated in the promoter indicating increased expression (Figure 5B).

**Figure 5:**
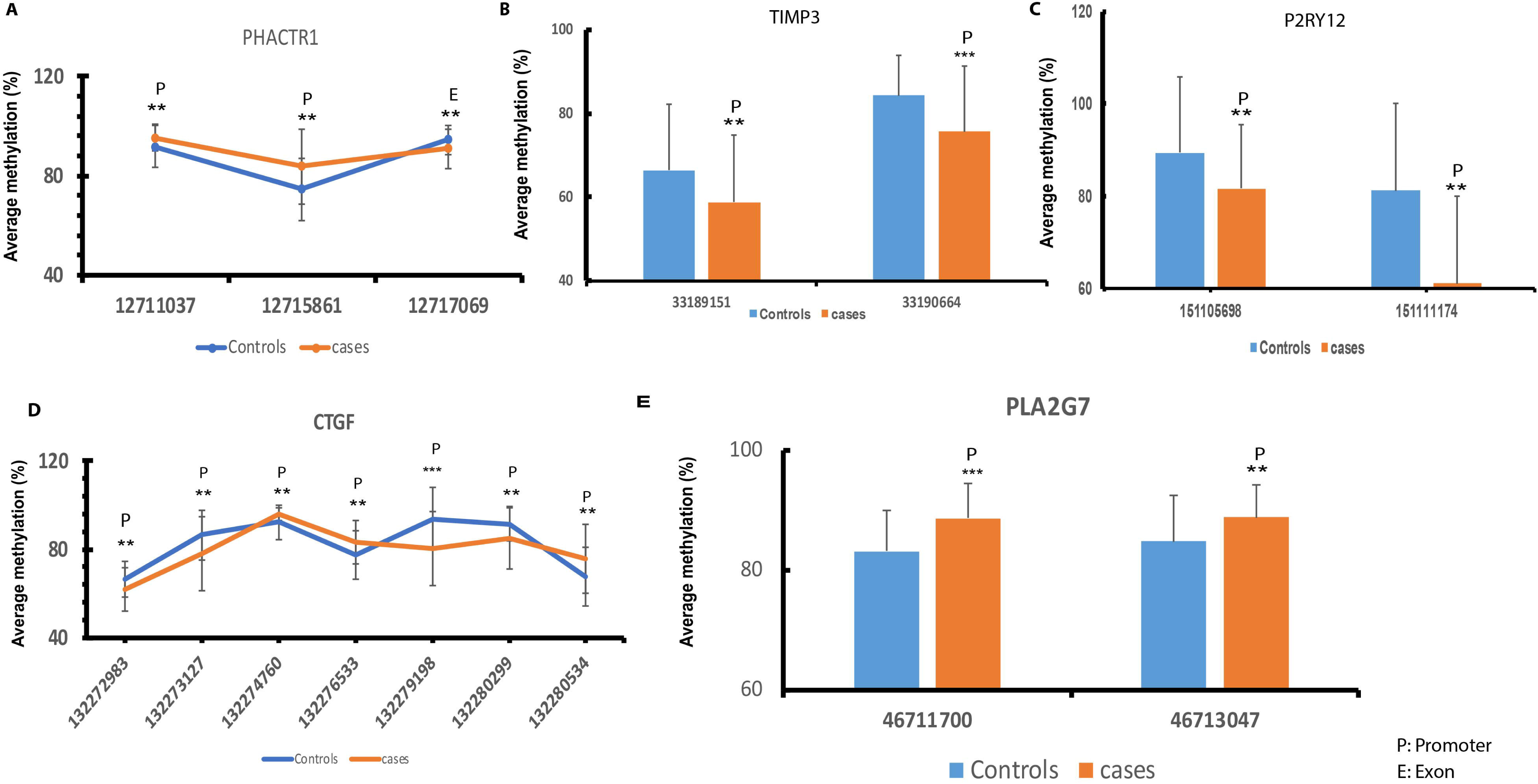
Methylation status of the genes involved in inflammation, extracellular matrix remodeling and platelet aggregation *(PHACTR1/TIMP3/P2RY12/CTGF/PLA2G7).* P: Promoter, E: Exon.** indicates p value < 0.01; *** indicates p value < 0.001.

We probed into the methylation status of the genes causing extracellular matrix turnover of the endothelial wall and intimal angiogenesis. The gene *P2RY12* responsible for platelet activation which encodes for purinergic receptor exhibited hypomethylation in the promoter region (Figure 5C) which could be responsible for platelet aggregation and inflammatory processes associated with atherosclerosis. Interestingly, we observed that the four loci falling into the promoter region of *CTGF* (Connective tissue growth factor) was hypomethylated which could lead to their increased expression and eventually lead to increased angiogenesis (Figure 5D). Promoter of *PLA2G7* (Phospholipase 2G7) known to catalyze degradation of platelet activating factor and abundant in necrotic core of coronary lesions, was found to be hypermethylated indicating lowered expression (Figure 5E).

### Methylation status of CDKN2B-AS1 (ANRIL) loci in CAD

Genome wide association and candidate gene studies have identified that polymorphisms in the gene *CDKN2BAS* (cyclin-dependent kinase inhibitor 2B antisense RNA) is linked to the predisposition towards the risk of coronary artery disease (CAD) (Samani, Erdmann et al. 2007). It is also reported that CDKN2BAS encodes a noncoding RNA (ANRIL) which plays a key role in progression of atherogenesis by modulating pathways like vascular cell proliferation, thrombogenesis and plaque stability (Zhao, Liao et al. 2016). We wanted to investigate the presence of DNA methylation in the regulatory region of cyclin dependent kinase inhibitor 2B anti sense RNA (CDKN2BAS1), polymorphisms of which have been reported with CAD in Han Chinese population. We observed that 5 out of 9 CpG sites falling in the promoter were hypermethylated. Overall promoter percentage methylation in CAD patients were 48.5 whereas in controls it was 47.3 (Figure 6).

**Figure 6:**
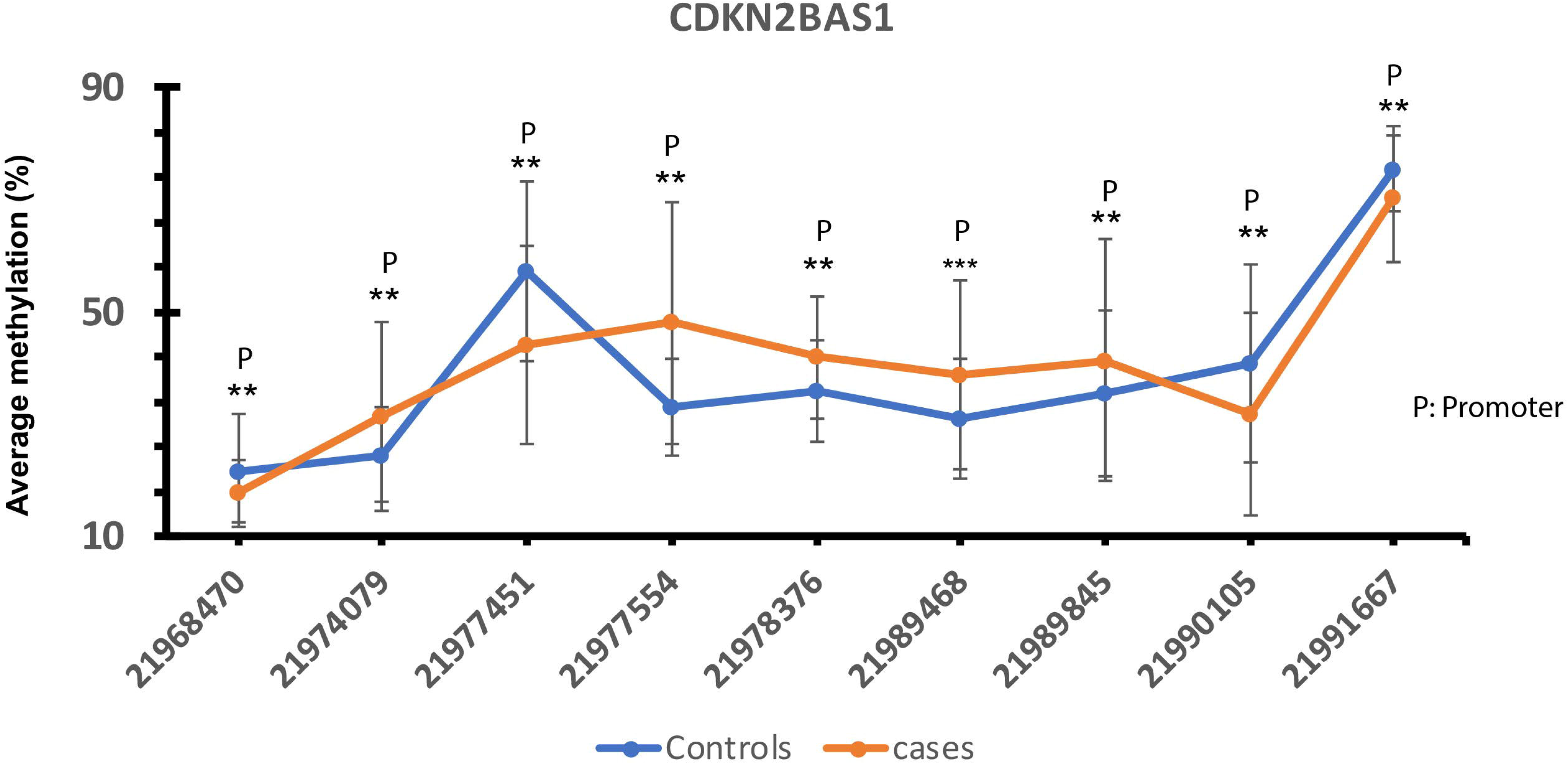
Methylation status of CDKN2B-AS1 gene; P: Promoter.** indicates p value < 0.01; *** indicates p value < 0.001.

## Discussion

In the current study, we interrogated DNA methylation status of candidate genes involved in cholesterol biosynthesis and metabolism, endothelial dysfunction, inflammation and apoptosis in relevance to the pathophysiology in young CAD patients. The genes chosen for the current study were previously reported to be associated with CAD in other populations. Polymorphisms of some of these genes were found to be associated with CAD earlier. Since cholesterol synthesis and metabolism is arguably one of the most important processes involved in the manifestation of CAD, we checked the methylation of a few genes involved in these processes. We observed that cholesterol biosynthetic gene *HMGCR* was hypermethylated in the promoter indicating lower expression. *Peng et al.* reported that *HMGCR* showed hypermethylation in the promoter (Peng, Wang et al. 2014). Lower expression of *HMGCR* could hint at lower cholesterol synthesis. Interestingly, we found promoter hypomethylation and exonic hypermethylation in the *LDLR* gene indicating higher uptake of lipoproteins inside the cell. It is very well established that promoter hypermethylation is associated with loss of transcription (Lim and Maher 2010). We and others have earlier shown that CAD in Indian population is usually associated with low HDL levels (Gupta, Rao et al. 2017). Excess intracellular cholesterol and other lipids are exported out of the cells in the form of HDL. Although, ApoA1, which initiates this process of reverse cholesterol transport did not show any significant alteration in methylation, *GALNT2*, which codes for polypeptide Gal-N-Ac transferase 2, was found to be hypermethylated in the promoter region. *GALNT2* is known to influence O linked oligosaccharide biosynthesis thus modulating HDL-C levels in blood. Interestingly, a recent study by *Sumeet A. Khetarpal* et. al. identified a novel mechanism of regulation of HDL-C metabolism by reducing O-sialylation of APOC3 residues and subsequent reduction of phospholipase transfer protein (PLTP) activity in primates (Khetarpal, Schjoldager et al. 2016). Therefore it could be hypothesized that increased promoter methylation in the *GALNT2* gene could be causing reduced glycosylation of *APOC3* and hence reduce HDL-C levels in blood predisposing the individuals towards CAD. Besides, *GALNT2* gene has also been shown to harbor promoter hypermethylation in CAD patients of Chinese population. We also found that *APOC3* and *APOA5* showed differential methylation in promoter and exonic region, with *APOC3* showing hyper while *APOA5* hypomethylation in the promoter region and vice versa in the exonic region. This suggests that CAD patients could have low APOC3 and high APOA5 levels resulting in low triglyceride levels. This is surprising because high triglyceride levels have long known to be a classical risk factor of CAD (Gotto 1998). Evidences also suggest that increased expression of *APOA5* could cause indirect activation of lipoprotein lipases (LPL) present on the surface of macrophages, endothelial cells and smooth muscle cells which might lead to accumulation of cholesterol esters inside macrophages via inhibition of cholesterol efflux (Ostlund-Lindqvist, Gustafson et al. 1983). This paradox of lower plasma cholesterol and triglyceride levels in Indian patients could probably be due to low vitamin B_12_ levels. We had earlier shown that low vitamin B_12_ is associated with CAD in Indian population. In this study also the levels of vitamin B_12_ were lower in CAD cases than controls (Kumar, Garg et al. 2009). Vitamin B_12_ deficiency has been linked to increased expression of LDLR in hepatocytes (Adaikalakoteswari, Finer et al. 2015). Vitamin B_12_ levels have also been found to directly correlate with HDL levels. Taken together, we hypothesize that vitamin B_12_ deficiency leads to increased cellular influx and decreased efflux of cholesterol. Notably, observations from the landmark INTERHEART study highlighted that South Asians had low triglyceride and LDL-C levels as compared to non-Asians and in acute MI patients triglyceride levels were lower as compared to controls (Karthikeyan, Teo et al. 2009).

Raised blood glucose levels are a known risk factor for CAD and we found hypermethylation in the promoter of *GCK* gene in CAD patients. *GCK* converts glucose to glucose-6-phosphate and the gene promoter (−287 to −158) is known to have a putative binding site for activating transcription factor 3 (ATF3) which ultimately leads to GCK down regulation in pancreatic β cells(Kim, Hwang et al. 2014). 5’UTR hypomethylation in the *ATF* gene suggests that under atherosclerotic conditions, increased expression of ATF3 may potentially mediate its binding to the GCK promoter and hence down regulate its transcription. It is also known that hyperglycemia can augment cardiovascular risk through several signaling pathways, i.e. downregulation of NO production via eNOS phosphorylation, *de novo* synthesis of diacylglycerol (DAG) and upregulation of NF-kβ activity (Funk, Yurdagul et al. 2012). Clinical evidences suggest that hyperglycemia induced endothelial dysfunction is involved in the pathogenesis of atherosclerosis (van den Oever, Raterman et al. 2010). Inflammation has been known to play pivotal role in facilitating atherosclerotic risk independent of serum cholesterol levels (Fioranelli, Bottaccioli et al. 2018). *AKT2* gene, a phosphoinositide dependent serine threonine kinase has been known to induce polarization of macrophages towards M1 state and promote atherosclerosis risk. It is also known that *AKT2* gene in macrophages plays an important role in migration of monocytes and stimulating pro inflammatory response (Rotllan, Chamorro-Jorganes et al. 2015). Besides, activation of *LCK* gene is an indication towards pro-inflammatory state in endothelial cells. We identified promoter hyper methylation in *PHACTR1* gene indicating that expression of this gene could be low in CAD patients. Knockdown of *PHACTR1* gene has been earlier reported to cause impaired vascular development in zebrafish (Jarray, Pavoni et al. 2015). Moreover, knockdown of PHACTR1 has been linked to reduced activity of pro inflammatory cytokine NF-Kβ (Zhang, Jiang et al. 2018).

Collagen deposition and extracellular matrix remodeling in the vessel wall is an integrative component of atherosclerosis. We found promoter hypomethylation in the *CTGF* gene indicating higher expression of CTGF which could also be triggered by pro inflammatory molecules and lead to endothelial apoptosis. Pro and anti-apoptotic genes also showed differential methylation in the promoter. We also checked the methylation status of genes involved in platelet aggregation and angiogenesis. The gene phospholipase A2 (*PLA2G7)* has been reported to be positively correlated to atherosclerotic risk in human studies but in our data but we observed promoter hypermethylation indicating low expression of this gene. However, hyper methylation in the promoter of (*PLA2G7*) gene has previously been reported to be associated with the risk of coronary heart disease (CHD) (Jiang, Zheng et al. 2013). Finally, we investigated methylation levels in the “Chr9p21” region which is known to be associated with coronary artery calcification and atherosclerosis. A recent study by *Shuyu Zhou et. al.* has reported higher methylation in the cyclin dependent kinase inhibitor 2B (*CDKN2B*) gene in peripheral blood of ischemic stroke patients. (Zhou, Zhang et al. 2016). We have observed an opposing trend of methylation in the *CDKN2BAS1* (cyclin dependent kinase inhibitor 2B anti sense1) promoter region. Additionally, *CDKN2B* exonic regions were identified to have significantly increased levels of methylation which is linked to vascular smooth muscle cell proliferation. Our data has provided evidences that DNA methylation has an important role to play in the pathogenesis of CAD which has long term clinical relevance as well. Although our study highlights important methylation alterations in young CAD but they suffer from a few limitations. We did not look at gender specific differences and cell type specific variations of methylation.

## Conclusions

In this study, we observed that in CAD patients *HMGCR* and *GALNT2* genes were hypermethylated in the promoter indicating that cholesterol biosynthesis might be low. Besides, LDLR receptor showed promoter hypomethylation and exonic hypermethylation indicating that intracellular cholesterol uptake might be higher resulting in low plasma LDL-C levels. Low APOC3 and higher APOA5 levels also reinforce our presumptions. Genes involved in endothelial dysfunction and inflammation showed differential methylation in CAD patients. Endothelial *NOS3* gene showed hypermethylation in promoter. These observations point towards the fact that genes involved in processes like cholesterol metabolism, hyperglycemia induced endothelial dysfunction, inflammation, extracellular matrix remodeling and platelet aggregation harbor significant methylation signatures that need further validation in large prospective cohorts. This study is also unique since it has investigated several genes whose methylation status has not been investigated earlier in CAD patients of any other population. This study also highlights the importance of methylation in the *CDKN2A/CDKN2B* cluster whose methylation status has not been very well established. It also forms the basis for further investigation into the functional role of these altered methylated genes in relation to CAD pathogenesis. Our study has investigated genes linked to important pathological processes involved with CAD majorly cholesterol homeostasis, endothelial dysfunction, vasoconstriction, endothelial apoptosis and inflammation. We have also highlighted that DNA methylation at candidate genes belonging to these processes can be interrogated as epigenetic biomarkers of CAD.

## Abbreviations

TG: Triglyceride;
LDL-C: Low density lipoprotein cholesterol;
HDL-C: High density lipoprotein cholesterol;
CAD: Coronary Artery Disease;
LDLR: Low density lipoprotein receptor;
HMGCR: 3-Hydroxy-3-Methylglutaryl-CoA Reductase;
APOA1: Apolipoprotein A1;
APOA4: Apolipoprotein A4;
APOA5: Apolipoprotein A5;
GWAS: Genome wide association study;
EWAS: Epigenome wide association study

## Conflict of interest

The authors report no conflict of interest.

## Author contributions

SSG conceptualized the idea and design of the experiment. SG (Subhoshree) and VST performed the bisulfite experiments. SG (Sourav) & PT (Priya) performed the bisulfite analysis. Downstream analysis was performed by SG (Subhoshree). Initial manuscript was drafted by SG (Subhoshree) with critical inputs from SG (Sourav) and SSG. AS (Anju) helped in performing the experiments and literature survey. SKV, AV, RJ were involved in performing the sequencing experiments. NB (Nitin) helped in collection of CAD samples from AIIMS and also performed the biochemical parameter measurements. SSG, SS (Sridhar) and VS provided critical suggestions to data analysis which helped in improving the manuscript. SS (Sandeep) performed the angiography at AIIMS and supervised the selection of sample cohort in the present study. We certify that all the authors have read and approved the content of the manuscript.

### Acknowledgements

We thank CSIR Network project (BSC0122) for funding. Subhoshree Ghose & Sourav Ghosh received fellowship support from CSIR. VST, PT, AS, NB, SKV, AV, RJ received support from CSIR-IGIB. We thank IGIB sequencing facility for the sequencing experiments. Dr. Sandeep Seth helped with sample collection from AIIMS, New Delhi. The funding agency had no role in study design or interpretation of results.

---

Supplementary figure 1 shows the study design and working methodology

Supplementary figure 2 describes the detailed workflow of Bioinformatic analysis to identify differentially methylated CpG sites

Supplementary figure 3A shown bioanalayzer profile of end repaired library with an average size of 175 bp. Figure 3B shows bioanalyzer profile of adapter ligated library with an average size of 250-280 bp. Figure 3C shows bioanalyzer profile of indexed library with an average size of 300 bp.

