## Supplementary figures and images for "Investigating Coronary Artery Disease methylome through targeted bisulfite sequencing"

### Supplementary figure 1

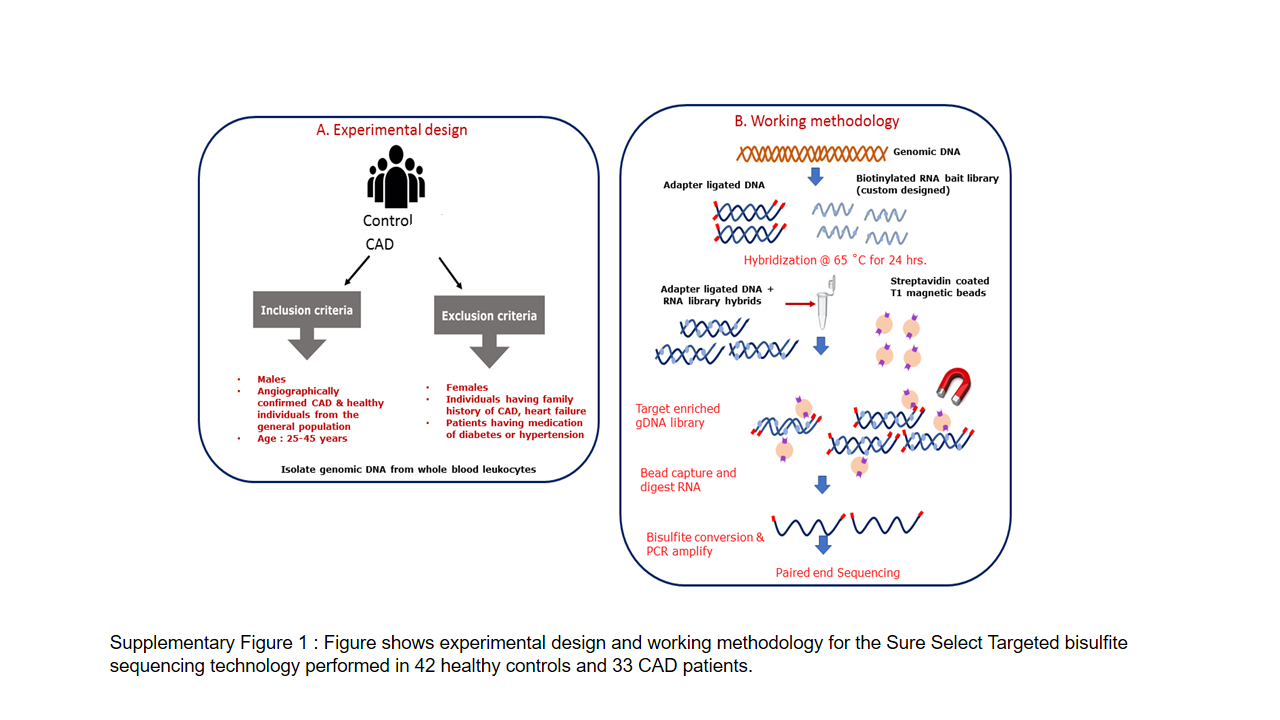

### Supplementary figure 2

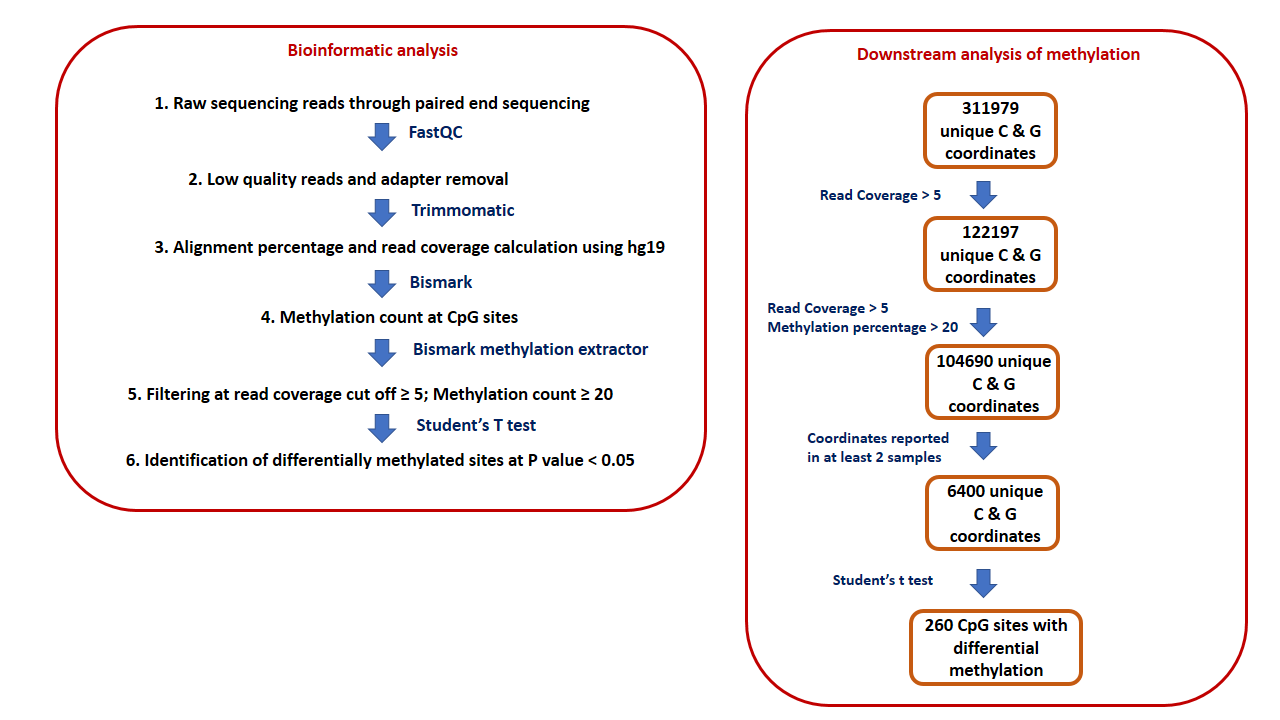

### Supplementary figure 3

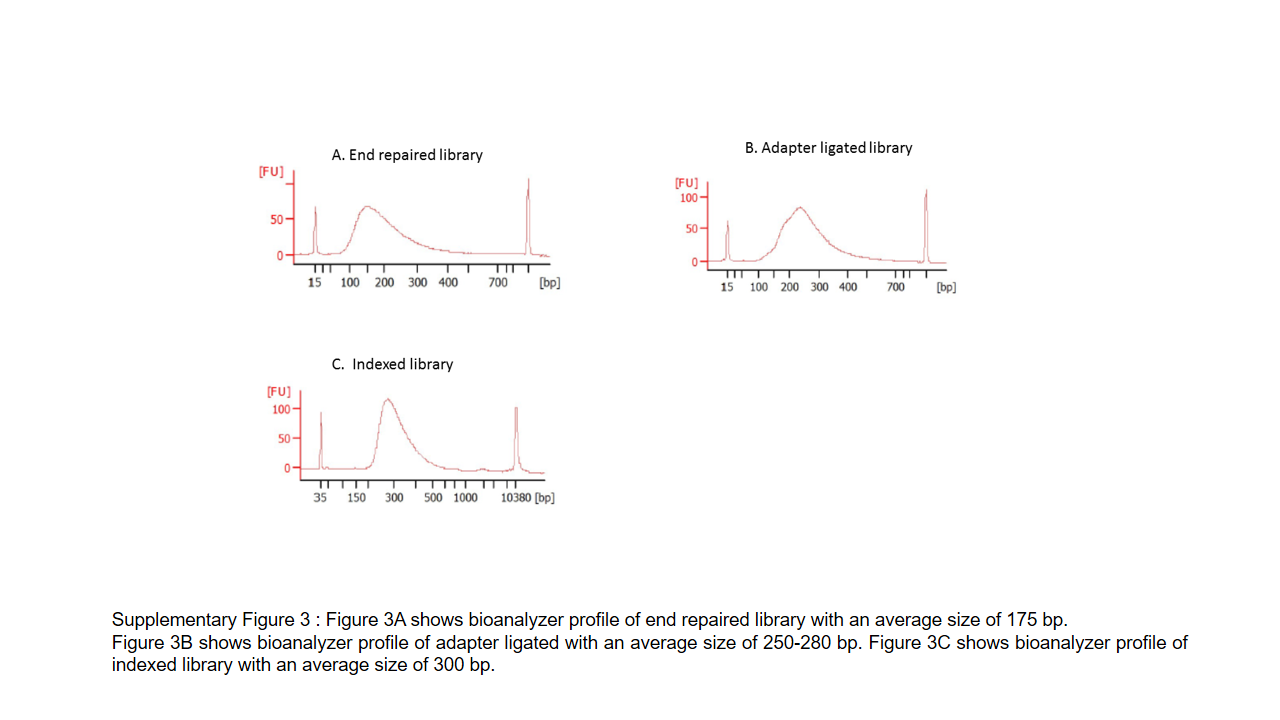
