## Supplementary table 1 for "Investigating Coronary Artery Disease methylome through targeted bisulfite sequencing"

**Supplementary Table 1 shows a list of genes and their coordinates included in the design for biotinylated probe design**

| Gene name | Chromosome | Coordinate |
|---|---|---|
| ABCG1 | chr21 | 43629265-43717855 |
| ADRA2C | chr4 | 3758295-3770253 |
| ALDOA | chr16 | 30054410-30078433 |
| ARHGEF18 | chr19 | 7459498-7537872 |
| ATF3 | chr1 | 212738675-212794119 |
| BAX | chr19 | 49448115-49465556 |
| CDKN1A | chr6 | 36643146-36655453 |
| COL15A1 | chr9 | 101695993-101833074 |
| ESR1 | chr6 | 152001629-152424408 |
| F2RL3 | chr19 | 16989824-17003331 |
| F7 | chr13 | 113750100-113775496 |
| FGF2 | chr4 | 123737861-123819891 |
| GALNT2 | chr1 | 230183534-230418377 |
| HMGCR | chr5 | 74622991-74658427 |
| KLF2 | chr19 | 16425649-16438339 |
| LCK | chr1 | 32706839-32751768 |
| LOXL1 | chr15 | 74208787-74244478 |
| MCPH1 | chr8 | 6254111-6471172 |
| MIA3 | chr1 | 222790920-222841354 |
| NOS3 | chr7 | 150678142-150711707 |
| PHACTR1 | chr6 | 12707740-13276355 |
| TIMP3 | chr22 | 33186800-33259529 |
| TMPO | chr12 | 98899350-98944157 |
| ABCA1 | chr9 | 107542782-107700528 |
| ABCB1 | chr7 | 87133178-87352639 |
| AKT2 | chr19 | 40736223-40801302 |
| ALOX15 | chr17 | 4533712-4554961 |
| AMOTL2 | chr3 | 134074186-134104260 |
| BCL2 | chr18 | 60790077-60996614 |
| CTGF | chr6 | 132269316-132282518 |
| DDAH2 | chr6 | 31694312-31708043 |
| DHFR | chr5 | 79950299-79960801 |
| E2F6 | chr2 | 11584500-11616303 |
| ESR2 | chr14 | 64693429-64815269 |
| FOXP3 | chrx | 49106395-49131289 |
| GCK | chr7 | 44183368-44239023 |
| ICAM2 | chr17 | 62079954-62094283 |
| LPA | chr6 | 11215850-11225123 |
| NR3C1 | chr5 | 142657495-142793254 |
| P2RY12 | chr3 | 151054129-151112602 |
| PECAM1 | chr17 | 62396744-62417083 |
| PLA2G7 | chr6 | 46671551-46713431 |
| TERT | chr5 | 1253286-1305163 |
| TFPI | chr2 | 188343304-188429220 |
| CDKN2BAS | chr9 | 22067093-22104313 |
| LDLR | chr19 | 11190036-11225123 |
| APOA1 | chr11 | 116658412-116726445 |
| CDKN2B-AS1 | chr9 | 21964197-22011962 |
