## Supplementary Table 2 for "Investigating Coronary Artery Disease methylome through targeted bisulfite sequencing"

| | A | B | C | D | E | F | G | H |
|---|---|---|---|---|---|---|---|---|
| 1 | **Supplementary Table 2 shows the values of total sequencing raw read count, filtered reads, alignment percentage and read coverage for all the 42 controls and 33 CAD patients** | | | | | | | |
| 2 | **Sample id** | **Total_Reads** | **QC_Reads** | **Mapping Efficiency** | **Uniquely mapped** | **Coverage** | | |
| 3 | Sample_C007 | 4845993 | 710551 | 61.1 | 433815 | 65.44 | | |
| 4 | Sample_C041 | 3982976 | 517453 | 64.6 | 334221 | 50.41 | | |
| 5 | Sample_C071 | 4832672 | 720244 | 59.6 | 429368 | 64.76 | | |
| 6 | Sample_C074 | 3115474 | 477132 | 56 | 266969 | 40.27 | | |
| 7 | Sample_C080 | 3405863 | 451428 | 64 | 288858 | 43.57 | | |
| 8 | Sample_C081 | 3830517 | 539818 | 64.3 | 346983 | 52.34 | | |
| 9 | Sample_C102 | 3142311 | 426670 | 57.2 | 244216 | 36.84 | | |
| 10 | Sample_C104 | 4400765 | 545300 | 63 | 343520 | 51.82 | | **C= CAD** |
| 11 | Sample_C110 | 2951659 | 410624 | 58.6 | 240723 | 36.31 | | **N= Control** |
| 12 | Sample_C113 | 4756624 | 668190 | 59.8 | 399523 | 60.26 | | |
| 13 | Sample_C115 | 3470167 | 448646 | 68.2 | 306176 | 46.18 | | |
| 14 | Sample_C166 | 3755442 | 525342 | 61.3 | 321879 | 48.55 | | |
| 15 | Sample_C195 | 5174443 | 682283 | 63.7 | 434519 | 65.54 | | |
| 16 | Sample_C196 | 3456721 | 448047 | 65.6 | 293839 | 44.32 | | |
| 17 | Sample_C255 | 4882027 | 711873 | 63.1 | 449178 | 67.75 | | |
| 18 | Sample_C261 | 4625661 | 661648 | 62.9 | 416259 | 62.79 | | |
| 19 | Sample_C267 | 5547516 | 823160 | 64.9 | 534433 | 80.61 | | |
| 20 | Sample_C272 | 4681626 | 697618 | 64.6 | 450358 | 67.93 | | |
| 21 | Sample_C292 | 4797237 | 661330 | 58.7 | 388228 | 58.56 | | |
| 22 | Sample_C303 | 4086339 | 569998 | 64.6 | 368482 | 55.58 | | |
| 23 | Sample_C308 | 4419636 | 557989 | 64.1 | 357874 | 53.98 | | |
| 24 | Sample_C312 | 4371653 | 605563 | 60.3 | 364960 | 55.05 | | |
| 25 | Sample_C316 | 4205317 | 559862 | 58.6 | 328022 | 49.48 | | |
| 26 | Sample_C360 | 5167012 | 702551 | 65.2 | 458199 | 69.11 | | |
| 27 | Sample_C416 | 4540197 | 633873 | 61.3 | 388267 | 58.57 | | |
| 28 | Sample_C429 | 4550426 | 650399 | 59.2 | 385055 | 58.08 | | |
| 29 | Sample_C431 | 3013905 | 386789 | 64.1 | 248058 | 37.42 | | |
| 30 | Sample_C442 | 3618841 | 485885 | 57.6 | 280029 | 42.24 | | |
| 31 | Sample_C448 | 4080738 | 523870 | 56.1 | 293652 | 44.29 | | |
| 32 | Sample_C452 | 3758007 | 534891 | 62.4 | 333994 | 50.38 | | |
| 33 | Sample_C465 | 3801493 | 515846 | 59.5 | 306707 | 46.26 | | |
| 34 | Sample_C495 | 2547824 | 366027 | 58.7 | 214766 | 32.39 | | |
| 35 | Sample_C504 | 3532063 | 479397 | 64.3 | 308158 | 46.48 | | |
| 36 | Sample_N085 | 4254814 | 576852 | 62.7 | 361446 | 54.52 | | |
| 37 | Sample_N088 | 4321650 | 570470 | 64.9 | 370250 | 55.85 | | |
| 38 | Sample_N124 | 4213298 | 524501 | 58.5 | 306929 | 46.3 | | |
| 39 | Sample_N136 | 4623724 | 590417 | 59.1 | 349003 | 52.64 | | |
| 40 | Sample_N137 | 3532138 | 541180 | 58.2 | 314886 | 47.5 | | |
| 41 | Sample_N140 | 4767667 | 671108 | 58 | 389427 | 58.74 | | |
| 42 | Sample_N156 | 5126176 | 716266 | 59.6 | 426971 | 64.4 | | |
| 43 | Sample_N165 | 6508245 | 922777 | 62.4 | 575521 | 86.81 | | |

| | A | B | C | D | E | F | G | H |
|---|---|---|---|---|---|---|---|---|
| 44 | **Sample id** | **Total_Reads** | **QC_Reads** | **Mapping Efficiency** | **Uniquely mapped** | **Coverage** | | |
| 45 | Sample_N167 | 9581749 | 1424208 | 62.1 | 885127 | 133.51 | | |
| 46 | Sample_N172 | 6248166 | 935706 | 62.4 | 584186 | 88.12 | | |
| 47 | Sample_N174 | 5093454 | 699745 | 58.7 | 410948 | 61.99 | | |
| 48 | Sample_N190 | 4963442 | 648112 | 58.6 | 379539 | 57.25 | | |
| 49 | Sample_N191 | 4376907 | 560603 | 58.2 | 326449 | 49.24 | | |
| 50 | Sample_N193 | 5046833 | 553245 | 59 | 326568 | 49.26 | | |
| 51 | Sample_N195 | 4699992 | 668167 | 63.9 | 427029 | 64.41 | | |
| 52 | Sample_N199 | 5290952 | 698308 | 64.9 | 453190 | 68.36 | | |
| 53 | Sample_N200 | 2879353 | 451602 | 59.2 | 267526 | 40.35 | | |
| 54 | Sample_N201 | 4367540 | 619098 | 61.4 | 380080 | 57.33 | | |
| 55 | Sample_N206 | 5174622 | 826014 | 58.8 | 485566 | 73.24 | | |
| 56 | Sample_N208 | 4937043 | 618590 | 58.7 | 362887 | 54.74 | | |
| 57 | Sample_N213 | 5898592 | 804888 | 62.7 | 504275 | 76.06 | | |
| 58 | Sample_N221 | 2751219 | 386621 | 62.5 | 241830 | 36.48 | | |
| 59 | Sample_N223 | 5105496 | 730835 | 63.4 | 463324 | 69.89 | | |
| 60 | Sample_N229 | 4210991 | 629931 | 60.2 | 379356 | 57.22 | | |
| 61 | Sample_N238 | 3842584 | 552650 | 59.7 | 329691 | 49.73 | | |
| 62 | Sample_N246 | 3632109 | 526625 | 58.8 | 309609 | 46.7 | | |
| 63 | Sample_N261 | 4457986 | 724999 | 57.5 | 416877 | 62.88 | | |
| 64 | Sample_N262 | 3522064 | 535785 | 61.9 | 331916 | 50.07 | | |
| 65 | Sample_N267 | 3398228 | 529542 | 53.4 | 283026 | 42.69 | | |
| 66 | Sample_N276 | 3907738 | 558146 | 59 | 329189 | 49.65 | | |
| 67 | Sample_N283 | 3040997 | 390319 | 66.4 | 259042 | 39.07 | | |
| 68 | Sample_N291 | 3339367 | 389567 | 61.3 | 238930 | 36.04 | | |
| 69 | Sample_N296 | 2241147 | 320627 | 57.7 | 185138 | 27.93 | | |
| 70 | Sample_N306 | 3404725 | 483287 | 61.9 | 298935 | 45.09 | | |
| 71 | Sample_N397 | 4530594 | 644269 | 59.9 | 385775 | 58.19 | | |
| 72 | Sample_N591 | 3054342 | 446355 | 57.8 | 257945 | 38.91 | | |
| 73 | Sample_N592 | 4164529 | 637204 | 58.2 | 370748 | 55.92 | | |
| 74 | Sample_N601 | 3223485 | 410129 | 75.3 | 308657 | 46.56 | | |
| 75 | Sample_N604 | 4469450 | 626905 | 58.7 | 368062 | 55.52 | | |
| 76 | Sample_N645 | 5414746 | 750597 | 60.8 | 456727 | 68.89 | | |
| 77 | Sample_N680 | 2328878 | 328530 | 60.2 | 197814 | 29.84 | | |
| 78 | Sample_N776 | 3613768 | 551279 | 55.7 | 307301 | 46.35 | | |
