## Supplementary Table 3 for "Investigating Coronary Artery Disease methylome through targeted bisulfite sequencing"

| Supplementary Table is a list of all the 6400 coordinates which and passed the read coverage and methylation percentage cut off | | | | | | | |
|---|---|---|---|---|---|---|---|
| Gene | Chromosome | Coordinate | No. of Controls | No. of CAD cases | Control methylation count | Case methylation count | p value |
| APOA1 | chr11 | 116686234 | 6 | 4 | 600 | 400 | 6.05E-14 |
| ATF3 | chr1 | 212745296 | 6 | 7 | 600 | 700 | 1.62E-12 |
| KLF2 | chr19 | 16432578 | 10 | 5 | 1000 | 500 | 1.62E-12 |
| LDLR | chr19 | 11222233 | 5 | 6 | 500 | 600 | 1.63E-12 |
| ABCG1 | chr21 | 43631319 | 8 | 3 | 800 | 300 | 1.63E-12 |
| COL15A1 | chr9 | 101696930 | 5 | 6 | 500 | 600 | 1.63E-12 |
| BCL2 | chr18 | 60988746 | 7 | 2 | 700 | 200 | 3.04E-12 |
| LDLR | chr19 | 11194085 | 4 | 5 | 400 | 500 | 3.04E-12 |
| APOA1 | chr11 | 116682998 | 6 | 2 | 600 | 200 | 3.58E-11 |
| APOA1 | chr11 | 116683002 | 6 | 2 | 600 | 200 | 3.58E-11 |
| ATF3 | chr1 | 212745289 | 3 | 4 | 300 | 400 | 9.98E-10 |
| GALNT2 | chr1 | 230188226 | 3 | 4 | 300 | 400 | 9.98E-10 |
| APOA1 | chr11 | 116668291 | 5 | 2 | 500 | 200 | 9.98E-10 |
| APOA1 | chr11 | 116725659 | 4 | 3 | 400 | 300 | 9.98E-10 |
| E2F6 | chr2 | 11608679 | 3 | 4 | 300 | 400 | 9.98E-10 |
| LOXL1 | chr15 | 74210392 | 4 | 2 | 400 | 200 | 3.12E-08 |
| AKT2 | chr19 | 40801015 | 4 | 2 | 400 | 166.6666667 | 3.12E-08 |
| FOXP3 | chrX | 49121911 | 4 | 2 | 400 | 200 | 3.12E-08 |
| FOXP3 | chrX | 49127045 | 2 | 4 | 40 | 80 | 3.12E-08 |
| LDLR | chr19 | 11220413 | 3 | 2 | 300 | 200 | 1.14E-06 |
| F2RL3 | chr19 | 16997237 | 2 | 3 | 40 | 60 | 1.14E-06 |
| BAX | chr19 | 49453884 | 3 | 2 | 300 | 200 | 1.14E-06 |
| APOA1 | chr11 | 116661248 | 31 | 32 | 2660.90125 | 3028.46628 | 0.000135367 |
| PLA2G7 | chr6 | 46711700 | 35 | 32 | 2907.244716 | 2835.20577 | 0.00076233 |
| FOXP3 | chrX | 49121289 | 21 | 18 | 1679.958791 | 1132.435065 | 0.00133892 |
| CTGF | chr6 | 132279198 | 30 | 27 | 2813.331448 | 2168.752184 | 0.00164071 |
| ICAM2 | chr17 | 62086376 | 30 | 33 | 1700.235569 | 1504.472239 | 0.00164379 |
| CDKN1A | chr6 | 36651933 | 31 | 31 | 2987.672346 | 2805.204288 | 0.00166194 |
| F7 | chr13 | 113756669 | 34 | 32 | 2508.381418 | 2579.3089 | 0.00192393 |
| ABCG1 | chr21 | 43629820 | 37 | 32 | 3522.277779 | 2874.058612 | 0.0021751 |
| PHACTR1 | chr6 | 12717192 | 2 | 4 | 200 | 331.4285714 | 0.0022792 |
| NOS3 | chr7 | 150689700 | 26 | 30 | 1769.570013 | 2434.986921 | 0.0024431 |
| GCK | chr7 | 44229968 | 10 | 19 | 419.4540708 | 1207.30464 | 0.0025609 |
| KLF2 | chr19 | 16438034 | 29 | 30 | 972.958874 | 1288.309009 | 0.002597 |
| TFPI | chr2 | 188424480 | 22 | 22 | 2168.293651 | 1981.067821 | 0.0026617 |
| LDLR | chr19 | 11221008 | 27 | 25 | 2269.989557 | 2338.599179 | 0.0028384 |
| F2RL3 | chr19 | 16998127 | 17 | 15 | 1429.184982 | 996.0714286 | 0.0029185 |
| APOA1 | chr11 | 116687627 | 32 | 32 | 2963.553796 | 3090.157093 | 0.0030358 |
| KLF2 | chr19 | 16433304 | 35 | 32 | 3457.780687 | 2977.12297 | 0.0030629 |
| TFPI | chr2 | 188420267 | 39 | 32 | 3004.300012 | 2689.91039 | 0.0030796 |

| | | | | | | | |
|---|---|---|---|---|---|---|---|
| F7 | chr13 | 113755287 | 32 | 31 | 2949.859307 | 3033.300547 | 0.0033371 |
| ADRA2C | chr4 | 3765853 | 18 | 17 | 1330.079365 | 1490.386003 | 0.0033431 |
| ADRA2C | chr4 | 3766721 | 25 | 25 | 1350.293651 | 1672.112724 | 0.0034237 |
| LDLR | chr19 | 11192639 | 22 | 20 | 2151.742424 | 1763.095238 | 0.0035626 |
| GCK | chr7 | 44231646 | 9 | 19 | 572.1753247 | 1612.608225 | 0.0036045 |
| HMGCR | chr5 | 74628717 | 31 | 30 | 2588.22911 | 2730.822788 | 0.0038189 |
| ABCG1 | chr21 | 43717591 | 37 | 32 | 3104.243264 | 2890.112966 | 0.004575 |
| LDLR | chr19 | 11221692 | 31 | 29 | 2851.185835 | 2812.255157 | 0.0045801 |
| CDKN2B-AS1 | chr9 | 22005889 | 25 | 27 | 920.4695867 | 1313.69129 | 0.0048565 |
| CDKN2B-AS1 | chr9 | 21989468 | 24 | 26 | 809.5659048 | 1140.979321 | 0.0049386 |
| LPA | chr6 | 160946112 | 32 | 32 | 3145.479996 | 2953.917403 | 0.0050151 |
| APOA1 | chr11 | 116723854 | 25 | 21 | 739.7567362 | 814.7053852 | 0.0058999 |
| ALOX15 | chr17 | 4553160 | 33 | 33 | 1574.973078 | 1308.38947 | 0.0063216 |
| APOA1 | chr11 | 116697880 | 31 | 30 | 2920.575064 | 2676.844966 | 0.0064352 |
| CDKN2B-AS1 | chr9 | 22009886 | 32 | 30 | 1756.675522 | 1369.526065 | 0.0066222 |
| APOA1 | chr11 | 116701833 | 31 | 32 | 3050.047796 | 2739.742382 | 0.0066394 |
| F7 | chr13 | 113750128 | 33 | 31 | 3204.001114 | 2826.589638 | 0.0066403 |
| KLF2 | chr19 | 16438050 | 33 | 32 | 1347.652719 | 1591.862541 | 0.0067418 |
| LCK | chr1 | 32739493 | 12 | 14 | 782.9691877 | 592.3412698 | 0.0067586 |
| NOS3 | chr7 | 150694867 | 32 | 31 | 2749.596914 | 2861.04145 | 0.0071606 |
| E2F6 | chr2 | 11613547 | 35 | 32 | 3322.792507 | 2931.948827 | 0.0083225 |
| APOA1 | chr11 | 116696636 | 33 | 32 | 2604.42204 | 2694.135944 | 0.0087409 |
| P2RY12 | chr3 | 151111174 | 19 | 11 | 1541.666667 | 672.0634921 | 0.0089032 |
| ADRA2C | chr4 | 3761998 | 26 | 27 | 2479.91009 | 2369.89956 | 0.0093249 |
| TIMP3 | chr22 | 33190664 | 32 | 33 | 2696.616007 | 2495.694229 | 0.009764 |
| TMPO | chr12 | 98900752 | 39 | 32 | 3833.655242 | 3079.942353 | 0.0098957 |
| ABCA1 | chr9 | 107691120 | 15 | 11 | 1402.261905 | 1100 | 0.0099785 |
| PLA2G7 | chr6 | 46713047 | 40 | 32 | 3390.057951 | 2841.381957 | 0.0100352 |
| LDLR | chr19 | 11219442 | 34 | 32 | 3208.279783 | 3129.765784 | 0.0102424 |
| CDKN2B-AS1 | chr9 | 21989845 | 20 | 19 | 709.6549529 | 943.1349206 | 0.0103386 |
| CDKN2B-AS1 | chr9 | 22001063 | 33 | 33 | 2125.792617 | 1811.839638 | 0.0104757 |
| NOS3 | chr7 | 150686444 | 30 | 33 | 1385.473035 | 1764.71109 | 0.011356 |
| LDLR | chr19 | 11197536 | 16 | 16 | 1533.055556 | 1417.380952 | 0.0116574 |
| ESR1 | chr6 | 152010481 | 38 | 32 | 3517.55283 | 3046.173451 | 0.011904 |
| NOS3 | chr7 | 150683826 | 33 | 32 | 2733.188019 | 2912.199969 | 0.0119589 |
| AMOTL2 | chr3 | 134098914 | 35 | 33 | 2860.117139 | 2481.069369 | 0.0120187 |
| TERT | chr5 | 1301170 | 34 | 32 | 3139.173343 | 3076.933869 | 0.0123428 |
| F7 | chr13 | 113756188 | 34 | 33 | 2418.248227 | 2133.7321 | 0.0123448 |
| PECAM1 | chr17 | 62399742 | 14 | 22 | 1342.853535 | 1919.920635 | 0.0124225 |
| GCK | chr7 | 44228831 | 36 | 32 | 3098.395854 | 2896.82119 | 0.012544 |
| ARHGEF18 | chr19 | 7461921 | 33 | 32 | 3189.298143 | 2770.217195 | 0.0127396 |
| LDLR | chr19 | 11217939 | 30 | 29 | 2826.561129 | 2862.804516 | 0.0127805 |
| KLF2 | chr19 | 16437980 | 31 | 28 | 1202.112436 | 1340.635794 | 0.0129163 |
| CDKN2B-AS1 | chr9 | 21968470 | 7 | 6 | 226.9047619 | 141.3071895 | 0.0133153 |
| F2RL3 | chr19 | 16991169 | 27 | 26 | 2226.719033 | 2388.613609 | 0.0134043 |
| ATF3 | chr1 | 212793952 | 37 | 32 | 3039.401717 | 2766.558684 | 0.0134372 |
| LDLR | chr19 | 11221697 | 30 | 29 | 2759.691503 | 2816.666667 | 0.0134572 |

| | | | | | | | |
|---|---|---|---|---|---|---|---|
| CDKN1A | chr6 | 36651959 | 30 | 31 | 2987.487513 | 2964.535595 | 0.0138002 |
| AMOTL2 | chr3 | 134095282 | 32 | 32 | 2722.255284 | 2891.898575 | 0.0143183 |
| FOXP3 | chrX | 49121784 | 16 | 9 | 1416.944444 | 885.7142857 | 0.0143445 |
| F7 | chr13 | 113760587 | 8 | 2 | 510 | 200 | 0.0144414 |
| APOA1 | chr11 | 116708414 | 32 | 30 | 2663.34869 | 2064.316386 | 0.0144582 |
| LDLR | chr19 | 11199718 | 34 | 32 | 1827.30144 | 1506.978573 | 0.0144591 |
| F7 | chr13 | 113750505 | 25 | 24 | 2339.484127 | 2373.333333 | 0.014679 |
| ICAM2 | chr17 | 62087479 | 34 | 32 | 3246.325922 | 3129.406038 | 0.014861 |
| TERT | chr5 | 1299860 | 12 | 17 | 731.0436788 | 772.6679879 | 0.0149007 |
| CTGF | chr6 | 132274760 | 34 | 32 | 3138.349677 | 3068.303241 | 0.0152034 |
| ESR1 | chr6 | 152002099 | 30 | 33 | 2880.262853 | 2940.172458 | 0.0156643 |
| F2RL3 | chr19 | 16998196 | 17 | 9 | 752.2763348 | 253.7301587 | 0.0157371 |
| LPA | chr6 | 11216121 | 32 | 32 | 2973.066835 | 3071.649815 | 0.016035 |
| LDLR | chr19 | 11224018 | 23 | 22 | 2089.569597 | 2133.360071 | 0.0162166 |
| TERT | chr5 | 1299447 | 24 | 27 | 2068.101343 | 2548.134921 | 0.0166429 |
| F2RL3 | chr19 | 16994756 | 32 | 31 | 2798.203341 | 2907.305539 | 0.0168589 |
| ADRA2C | chr4 | 3760991 | 32 | 29 | 2853.639971 | 2766.175691 | 0.0171362 |
| MCPH1 | chr8 | 6470832 | 34 | 31 | 3126.432337 | 2985.708171 | 0.0172374 |
| COL15A1 | chr9 | 101698014 | 34 | 32 | 2895.740938 | 2866.269512 | 0.0172398 |
| APOA1 | chr11 | 116687933 | 32 | 31 | 3033.537113 | 3034.001823 | 0.0172593 |
| CDKN2B-AS1 | chr9 | 21977554 | 13 | 10 | 429.1666667 | 515.4761905 | 0.0172634 |
| ICAM2 | chr17 | 62080221 | 28 | 29 | 2693.326118 | 2879.79798 | 0.0173751 |
| CDKN2B-AS1 | chr9 | 22000964 | 33 | 33 | 2475.752006 | 2130.642638 | 0.0175962 |
| TERT | chr5 | 1296463 | 34 | 33 | 2793.653158 | 2503.444008 | 0.0178199 |
| APOA1 | chr11 | 116691928 | 32 | 32 | 2909.239159 | 3049.794166 | 0.0178604 |
| LDLR | chr19 | 11216032 | 33 | 31 | 3082.045533 | 3012.098074 | 0.0181643 |
| APOA1 | chr11 | 116725861 | 34 | 33 | 1838.142246 | 1550.56936 | 0.0187623 |
| NOS3 | chr7 | 150679361 | 38 | 33 | 2086.237291 | 2000.576065 | 0.0189763 |
| KLF2 | chr19 | 16433445 | 34 | 32 | 3281.516637 | 3004.594136 | 0.0192966 |
| ATF3 | chr1 | 212739706 | 27 | 28 | 2472.885496 | 2265.885226 | 0.0193812 |
| DHFR | chr5 | 79959441 | 20 | 19 | 1825.119048 | 1853.888889 | 0.0194186 |
| ADRA2C | chr4 | 3764755 | 31 | 31 | 1899.559484 | 1592.950393 | 0.019441 |
| CDKN2B-AS1 | chr9 | 21977451 | 29 | 25 | 1657.143918 | 1171.013709 | 0.0195764 |
| APOA1 | chr11 | 116661216 | 31 | 32 | 2840.256015 | 3067.811663 | 0.0195823 |
| ARHGEF18 | chr19 | 7464099 | 33 | 32 | 3039.025467 | 2763.302993 | 0.0197794 |
| COL15A1 | chr9 | 101704337 | 35 | 32 | 3146.366722 | 2995.574006 | 0.0197976 |
| PHACTR1 | chr6 | 12715861 | 25 | 28 | 1865.986455 | 2344.940476 | 0.02086 |
| ADRA2C | chr4 | 3766521 | 27 | 29 | 2386.282096 | 2337.967178 | 0.021116 |
| F2RL3 | chr19 | 16990296 | 30 | 31 | 2825.721092 | 3038.696581 | 0.021201 |
| APOA1 | chr11 | 116721465 | 33 | 32 | 3258.827229 | 2864.249855 | 0.021241 |
| F2RL3 | chr19 | 16998949 | 32 | 33 | 2733.366386 | 2608.380942 | 0.021284 |
| BCL2 | chr18 | 60995889 | 22 | 16 | 1932.832168 | 1531.309524 | 0.021624 |
| CDKN2B-AS1 | chr9 | 21978376 | 36 | 33 | 1317.189357 | 1388.007684 | 0.021637 |
| F7 | chr13 | 113750103 | 32 | 31 | 3067.717763 | 2854.622719 | 0.021658 |
| APOA1 | chr11 | 116725636 | 12 | 9 | 1117.575758 | 900 | 0.021702 |
| TMPO | chr12 | 98906185 | 33 | 32 | 3142.253761 | 2900.953057 | 0.021716 |
| LDLR | chr19 | 11192553 | 31 | 26 | 3034.931041 | 2375.40293 | 0.021903 |

| | | | | | | | |
|---|---|---|---|---|---|---|---|
| TMPO | chr12 | 98900798 | 41 | 32 | 3947.5111 | 3017.745819 | 0.022145 |
| CDKN2B-AS1 | chr9 | 21974079 | 5 | 3 | 130.7936508 | 130 | 0.022356 |
| F7 | chr13 | 113750985 | 30 | 23 | 2697.478355 | 2198.94958 | 0.022797 |
| APOA1 | chr11 | 116704450 | 33 | 33 | 2969.798386 | 2758.633113 | 0.023039 |
| MIA3 | chr1 | 222808651 | 37 | 32 | 3370.595736 | 3006.529588 | 0.023619 |
| LCK | chr1 | 32740607 | 15 | 14 | 1333.607504 | 1357.777778 | 0.023838 |
| FOXP3 | chrX | 49116085 | 24 | 30 | 2105.413059 | 2857.893218 | 0.023857 |
| TERT | chr5 | 1295754 | 27 | 21 | 888.1714169 | 856.9523768 | 0.02406 |
| APOA1 | chr11 | 116690546 | 31 | 33 | 2078.259868 | 2488.141553 | 0.024383 |
| CTGF | chr6 | 132273127 | 30 | 32 | 2591.790324 | 2497.287102 | 0.024401 |
| APOA1 | chr11 | 116714317 | 37 | 32 | 3526.825451 | 2985.376127 | 0.024481 |
| MCPH1 | chr8 | 6470178 | 19 | 23 | 1888.649684 | 2058.441558 | 0.024619 |
| ICAM2 | chr17 | 62091363 | 32 | 32 | 2970.289663 | 3124.852941 | 0.024756 |
| NR3C1 | chr5 | 142788918 | 36 | 32 | 3378.233411 | 2922.490197 | 0.024763 |
| ESR2 | chr14 | 64806245 | 37 | 33 | 3177.031669 | 2643.560749 | 0.025124 |
| ABCB1 | chr7 | 87343538 | 26 | 22 | 2420.627706 | 2179.807692 | 0.025417 |
| FOXP3 | chrX | 49115016 | 13 | 5 | 1055.238095 | 293.3333333 | 0.025441 |
| CDKN2B-AS1 | chr9 | 22005101 | 20 | 13 | 516.2954177 | 466.8903319 | 0.025643 |
| LDLR | chr19 | 11221389 | 34 | 32 | 3224.642383 | 3127.543572 | 0.026705 |
| APOA1 | chr11 | 116661335 | 30 | 32 | 2601.99794 | 2940.67472 | 0.026981 |
| ATF3 | chr1 | 212745495 | 32 | 32 | 3077.167157 | 2967.421722 | 0.027035 |
| LOXL1 | chr15 | 74210831 | 36 | 32 | 3271.388545 | 2998.377202 | 0.027319 |
| LOXL1 | chr15 | 74209559 | 25 | 23 | 2365.436508 | 2273.214286 | 0.027377 |
| CTGF | chr6 | 132280299 | 35 | 31 | 3200.148177 | 2708.951833 | 0.028042 |
| APOA1 | chr11 | 116678334 | 17 | 18 | 1571.190476 | 1757.142857 | 0.028059 |
| LDLR | chr19 | 11224057 | 24 | 18 | 2221.309524 | 1783.333333 | 0.028181 |
| CDKN2B-AS1 | chr9 | 21991667 | 37 | 33 | 2792.539068 | 2318.549655 | 0.028361 |
| FOXP3 | chrX | 49121276 | 21 | 21 | 1745.436785 | 1524.576012 | 0.028478 |
| APOA1 | chr11 | 116708556 | 29 | 30 | 2772.811772 | 2599.416997 | 0.028623 |
| TFPI | chr2 | 188426841 | 24 | 21 | 2076.395467 | 1970.093158 | 0.028664 |
| LCK | chr1 | 32741484 | 25 | 25 | 2211.498779 | 1961.381674 | 0.028778 |
| CDKN2B-AS1 | chr9 | 21990105 | 13 | 18 | 632.089169 | 642.1184371 | 0.028899 |
| LDLR | chr19 | 11221486 | 34 | 32 | 3259.867609 | 3137.146419 | 0.029659 |
| APOA1 | chr11 | 116691961 | 32 | 32 | 2951.202466 | 3088.256473 | 0.030179 |
| APOA1 | chr11 | 116708518 | 31 | 31 | 3021.752315 | 2924.063649 | 0.030359 |
| PECAM1 | chr17 | 62398226 | 36 | 33 | 3089.381442 | 2656.854054 | 0.030424 |
| ABCA1 | chr9 | 107542842 | 34 | 32 | 2828.844758 | 2451.085611 | 0.030518 |
| FGF2 | chr4 | 123741393 | 35 | 32 | 3285.559922 | 2916.456135 | 0.030539 |
| LOXL1 | chr15 | 74218620 | 15 | 19 | 494.8276723 | 890.0968476 | 0.030599 |
| TIMP3 | chr22 | 33189151 | 30 | 30 | 2039.994054 | 1806.386559 | 0.030895 |
| AKT2 | chr19 | 40794901 | 33 | 32 | 3230.521136 | 3041.45447 | 0.030959 |
| APOA1 | chr11 | 116691951 | 32 | 32 | 2959.625112 | 3106.468431 | 0.031043 |
| LOXL1 | chr15 | 74211401 | 35 | 32 | 3174.697564 | 2997.191084 | 0.031402 |
| CTGF | chr6 | 132272983 | 35 | 33 | 2325.489793 | 2037.538089 | 0.03144 |
| ALDOA | chr16 | 30061597 | 23 | 27 | 2184.629815 | 2399.73693 | 0.032084 |
| LOXL1 | chr15 | 74211612 | 14 | 11 | 1253.650794 | 1064.880952 | 0.0323 |
| CDKN1A | chr6 | 36654680 | 32 | 32 | 3071.608865 | 2959.694733 | 0.032456 |

| | | | | | | | |
|---|---|---|---|---|---|---|---|
| NOS3 | chr7 | 150695647 | 23 | 22 | 2249.047619 | 2046.226551 | 0.032844 |
| FOXP3 | chrX | 49130161 | 31 | 31 | 1785.234519 | 2004.693918 | 0.033135 |
| ESR1 | chr6 | 152007888 | 27 | 24 | 2148.986985 | 1585.078059 | 0.033343 |
| TERT | chr5 | 1253550 | 31 | 31 | 2679.064516 | 2879.627458 | 0.033582 |
| FOXP3 | chrX | 49126877 | 7 | 5 | 200.8333333 | 220 | 0.033758 |
| DHFR | chr5 | 79954125 | 6 | 4 | 440.7142857 | 385.7142857 | 0.033943 |
| AMOTL2 | chr3 | 134099761 | 31 | 31 | 3054.586623 | 2800.151876 | 0.034042 |
| ABCG1 | chr21 | 43637024 | 32 | 32 | 2661.951014 | 2836.541639 | 0.034298 |
| LDLR | chr19 | 11217263 | 27 | 31 | 2672.334055 | 2942.353449 | 0.034355 |
| ARHGEF18 | chr19 | 7468074 | 22 | 18 | 2014.761905 | 1759.047619 | 0.034624 |
| FOXP3 | chrX | 49130389 | 29 | 30 | 2507.239403 | 2329.236336 | 0.034699 |
| ADRA2C | chr4 | 3760184 | 35 | 32 | 2794.064131 | 2691.39899 | 0.034985 |
| APOA1 | chr11 | 116705516 | 21 | 24 | 1205.323843 | 1761.111111 | 0.035037 |
| APOA1 | chr11 | 116699075 | 19 | 23 | 1395.766595 | 1953.044733 | 0.035517 |
| APOA1 | chr11 | 116687953 | 32 | 32 | 3021.990188 | 3102.453819 | 0.035637 |
| CTGF | chr6 | 132280534 | 30 | 31 | 2032.789973 | 2348.484965 | 0.035764 |
| KLF2 | chr19 | 16426847 | 7 | 7 | 507.0238095 | 603.3333333 | 0.035802 |
| PHACTR1 | chr6 | 12717341 | 33 | 33 | 3016.100463 | 2839.434731 | 0.035836 |
| APOA1 | chr11 | 116674643 | 28 | 32 | 2778.825439 | 3021.018483 | 0.035998 |
| LPA | chr6 | 160950075 | 32 | 32 | 2930.8083 | 3040.222983 | 0.036044 |
| ABCG1 | chr21 | 43634598 | 28 | 31 | 2320.897034 | 2793.272283 | 0.036222 |
| GCK | chr7 | 44231431 | 14 | 14 | 1106.547619 | 1265 | 0.036557 |
| ARHGEF18 | chr19 | 7461329 | 24 | 22 | 2206.071429 | 2137.142857 | 0.037057 |
| LDLR | chr19 | 11199222 | 32 | 31 | 2670.389903 | 2745.749826 | 0.037064 |
| NOS3 | chr7 | 150705831 | 28 | 27 | 969.4632936 | 1092.170761 | 0.03708 |
| ABCB1 | chr7 | 87342828 | 32 | 31 | 2137.987643 | 2267.592598 | 0.037222 |
| ABCB1 | chr7 | 87350728 | 33 | 32 | 2823.381757 | 2503.065822 | 0.037437 |
| FGF2 | chr4 | 123746991 | 22 | 19 | 2000.125153 | 1834.040404 | 0.037899 |
| CDKN1A | chr6 | 36652302 | 33 | 32 | 3010.647734 | 3047.186878 | 0.038061 |
| APOA1 | chr11 | 116681284 | 23 | 23 | 2258.75 | 2163.883061 | 0.038299 |
| ABCA1 | chr9 | 107691631 | 38 | 32 | 3496.528734 | 2829.008842 | 0.038883 |
| FOXP3 | chrX | 49119927 | 32 | 33 | 2677.842477 | 2545.889789 | 0.038942 |
| LOXL1 | chr15 | 74213839 | 32 | 32 | 2613.61453 | 2448.185977 | 0.0391 |
| GCK | chr7 | 44183776 | 35 | 32 | 3230.062136 | 3035.631651 | 0.039704 |
| ALOX15 | chr17 | 4554325 | 8 | 6 | 676.0606061 | 587.5 | 0.039739 |
| APOA1 | chr11 | 116726424 | 17 | 21 | 528.6580087 | 802.3859963 | 0.039765 |
| ABCB1 | chr7 | 87352198 | 39 | 32 | 3370.126687 | 2857.536804 | 0.039805 |
| E2F6 | chr2 | 11613270 | 16 | 22 | 1471.825397 | 2131.388889 | 0.039821 |
| APOA1 | chr11 | 116692085 | 32 | 32 | 2801.374457 | 2927.095825 | 0.039844 |
| CTGF | chr6 | 132276533 | 33 | 32 | 2560.110867 | 2657.430024 | 0.04004 |
| LDLR | chr19 | 11222323 | 13 | 14 | 1181.428571 | 1359.722222 | 0.040136 |
| FGF2 | chr4 | 123746873 | 33 | 32 | 3115.965202 | 2903.292319 | 0.040155 |
| APOA1 | chr11 | 116661450 | 30 | 31 | 2305.965641 | 2597.882282 | 0.040162 |
| LDLR | chr19 | 11195038 | 33 | 31 | 3161.633989 | 2852.017377 | 0.040355 |
| NOS3 | chr7 | 150708045 | 30 | 33 | 1770.399749 | 2234.542125 | 0.040882 |
| APOA1 | chr11 | 116701560 | 31 | 28 | 3006.767677 | 2615.18759 | 0.041062 |
| LDLR | chr19 | 11223961 | 23 | 21 | 2172.711039 | 2074.603175 | 0.041117 |

| | | | | | | | |
|---|---|---|---|---|---|---|---|
| TERT | chr5 | 1296577 | 32 | 30 | 3125.173395 | 2846.152846 | 0.041422 |
| APOA1 | chr11 | 116708317 | 32 | 32 | 3089.397721 | 2989.079166 | 0.041896 |
| LDLR | chr19 | 11198964 | 23 | 25 | 2117.539683 | 2428.409091 | 0.041985 |
| FOXP3 | chrX | 49106474 | 21 | 23 | 1824.076063 | 1745.952381 | 0.042663 |
| ABCG1 | chr21 | 43635188 | 34 | 32 | 3059.219617 | 3042.813742 | 0.042721 |
| ADRA2C | chr4 | 3762695 | 36 | 32 | 2262.521825 | 2179.978249 | 0.042824 |
| LDLR | chr19 | 11223530 | 11 | 12 | 1100 | 1138.928571 | 0.042998 |
| BAX | chr19 | 49449696 | 31 | 32 | 2905.354111 | 3092.204905 | 0.043163 |
| ADRA2C | chr4 | 3762969 | 28 | 29 | 2392.273005 | 2279.694195 | 0.04326 |
| BAX | chr19 | 49453159 | 25 | 19 | 1886.262626 | 1243.001443 | 0.04328 |
| ABCG1 | chr21 | 43717781 | 37 | 32 | 3544.394002 | 2996.30796 | 0.043396 |
| P2RY12 | chr3 | 151105698 | 31 | 33 | 2771.802538 | 2694.418981 | 0.043457 |
| APOA1 | chr11 | 116692558 | 32 | 32 | 2901.019711 | 3019.623327 | 0.043698 |
| LDLR | chr19 | 11221653 | 32 | 32 | 2862.624706 | 2982.020371 | 0.044098 |
| F2RL3 | chr19 | 16998540 | 2 | 2 | 40 | 61.9047619 | 0.044156 |
| TERT | chr5 | 1295030 | 2 | 2 | 61.9047619 | 40 | 0.044156 |
| LPA | chr6 | 11218999 | 2 | 2 | 40 | 61.9047619 | 0.044156 |
| ARHGEF18 | chr19 | 7463987 | 32 | 32 | 2549.140648 | 2745.909699 | 0.0442 |
| ADRA2C | chr4 | 3760951 | 31 | 28 | 2495.931013 | 2453.137794 | 0.044382 |
| APOA1 | chr11 | 116715660 | 36 | 32 | 3353.911119 | 3056.259274 | 0.044464 |
| APOA1 | chr11 | 116725440 | 34 | 32 | 3336.79097 | 3011.21156 | 0.044464 |
| MCPH1 | chr8 | 6260206 | 12 | 22 | 1200 | 2152.103175 | 0.044521 |
| APOA1 | chr11 | 116708984 | 28 | 27 | 2650.445906 | 2657.478632 | 0.04508 |
| LCK | chr1 | 32740883 | 22 | 26 | 1440.770063 | 1980.06993 | 0.045196 |
| DHFR | chr5 | 79952426 | 26 | 22 | 2291.584603 | 1726.357422 | 0.045456 |
| PECAM1 | chr17 | 62397478 | 32 | 32 | 3124.552596 | 3043.098589 | 0.045539 |
| LCK | chr1 | 32740399 | 26 | 28 | 2518.591921 | 2575.850641 | 0.045604 |
| LPA | chr6 | 11224714 | 2 | 4 | 146.6666667 | 142.1428571 | 0.045681 |
| MCPH1 | chr8 | 6470820 | 33 | 31 | 2985.514881 | 2946.734374 | 0.045939 |
| NOS3 | chr7 | 150695069 | 6 | 4 | 600 | 314.6031746 | 0.045977 |
| LOXL1 | chr15 | 74218777 | 16 | 12 | 406.1105561 | 449.6480643 | 0.046262 |
| MCPH1 | chr8 | 6470110 | 21 | 25 | 2063.214286 | 2337.234432 | 0.046356 |
| APOA1 | chr11 | 116663322 | 34 | 32 | 2985.450879 | 2936.043444 | 0.04636 |
| APOA1 | chr11 | 116724949 | 4 | 3 | 273.8095238 | 283.3333333 | 0.046401 |
| PHACTR1 | chr6 | 12711037 | 33 | 32 | 3027.076802 | 3052.293347 | 0.046483 |
| APOA1 | chr11 | 116676590 | 6 | 4 | 534.7619048 | 270.4761905 | 0.046518 |
| LOXL1 | chr15 | 74218715 | 16 | 17 | 472.056277 | 715.8899434 | 0.046519 |
| ADRA2C | chr4 | 3760295 | 36 | 32 | 3099.450658 | 2872.718022 | 0.04652 |
| AMOTL2 | chr3 | 134097086 | 34 | 32 | 2935.015198 | 2568.723451 | 0.046658 |
| GALNT2 | chr1 | 230188247 | 18 | 15 | 1594.642857 | 1448.333333 | 0.046755 |
| NOS3 | chr7 | 150710683 | 5 | 8 | 158.3333333 | 184.1269841 | 0.046857 |
| AKT2 | chr19 | 40795616 | 13 | 13 | 1300 | 1224.047619 | 0.04686 |
| ADRA2C | chr4 | 3761649 | 34 | 32 | 2961.127215 | 2911.275165 | 0.047017 |
| LDLR | chr19 | 11196237 | 35 | 32 | 3236.037618 | 2814.033387 | 0.047058 |
| GALNT2 | chr1 | 230190310 | 30 | 30 | 2675.920399 | 2858.187923 | 0.047156 |
| APOA1 | chr11 | 116724233 | 33 | 28 | 1905.424417 | 1345.588522 | 0.047204 |
| APOA1 | chr11 | 116661389 | 32 | 32 | 2913.801911 | 3040.75293 | 0.047318 |

| | | | | | | | |
|---|---|---|---|---|---|---|---|
| CDKN1A | chr6 | 36652031 | 29 | 30 | 2663.564214 | 2928.675214 | 0.047403 |
| TFPI | chr2 | 188425287 | 23 | 25 | 2192.247475 | 2245.898268 | 0.047463 |
| PHACTR1 | chr6 | 12717069 | 32 | 32 | 3020.009972 | 2907.285891 | 0.047678 |
| APOA1 | chr11 | 116726158 | 31 | 30 | 2947.512373 | 2744.647595 | 0.047958 |
| ADRA2C | chr4 | 3761372 | 32 | 30 | 2746.269954 | 2738.896761 | 0.048537 |
| LDLR | chr19 | 11221291 | 33 | 32 | 3186.732341 | 3004.793623 | 0.048744 |
| E2F6 | chr2 | 11613685 | 35 | 32 | 3312.092762 | 2937.045275 | 0.048802 |
| FOXP3 | chrX | 49117190 | 31 | 32 | 2571.896673 | 2880.247489 | 0.048879 |
| COL15A1 | chr9 | 101705919 | 7 | 5 | 190.8333333 | 102.2222222 | 0.049277 |
| GCK | chr7 | 44237017 | 38 | 32 | 3078.539383 | 2709.895073 | 0.049821 |
| NOS3 | chr7 | 150695213 | 7 | 14 | 265.2167277 | 806.8470419 | 0.050015 |
| APOA1 | chr11 | 116723031 | 32 | 30 | 1475.991146 | 1198.244767 | 0.050019 |
| NOS3 | chr7 | 150707235 | 20 | 26 | 670.4989455 | 1067.113646 | 0.050042 |
| ARHGEF18 | chr19 | 7469310 | 25 | 26 | 2236.818182 | 2480.753968 | 0.050196 |
| PECAM1 | chr17 | 62399988 | 35 | 32 | 3397.009707 | 3032.529901 | 0.05042 |
| FOXP3 | chrX | 49127038 | 18 | 13 | 1725.833333 | 1120.833333 | 0.050803 |
| F7 | chr13 | 113753782 | 36 | 32 | 3052.959362 | 2852.175183 | 0.051041 |
| AMOTL2 | chr3 | 134074344 | 24 | 26 | 2157.191142 | 2478.103841 | 0.051062 |
| F7 | chr13 | 113751421 | 28 | 29 | 2630.733711 | 2866.666667 | 0.051196 |
| MIA3 | chr1 | 222808763 | 40 | 32 | 3744.927838 | 3075.996275 | 0.051723 |
| ATF3 | chr1 | 212783692 | 37 | 33 | 3324.865509 | 2806.117645 | 0.051776 |
| ADRA2C | chr4 | 3763006 | 7 | 6 | 277.1428571 | 406.1904762 | 0.052458 |
| CTGF | chr6 | 132278753 | 39 | 32 | 3784.248416 | 3043.046461 | 0.052862 |
| CDKN2B-AS1 | chr9 | 21970609 | 26 | 32 | 1502.663635 | 1609.683788 | 0.053023 |
| APOA1 | chr11 | 116662854 | 32 | 32 | 2967.952881 | 3075.773317 | 0.053178 |
| ALOX15 | chr17 | 4545937 | 14 | 21 | 410.493191 | 802.6515151 | 0.053221 |
| MIA3 | chr1 | 222809210 | 33 | 32 | 3129.914329 | 2950.357234 | 0.053298 |
| ABCA1 | chr9 | 107692427 | 34 | 32 | 3084.761671 | 3031.21179 | 0.053456 |
| APOA1 | chr11 | 116704621 | 35 | 32 | 3313.497482 | 2926.81236 | 0.053741 |
| KLF2 | chr19 | 16434561 | 34 | 33 | 1843.165713 | 1590.680185 | 0.05414 |
| ARHGEF18 | chr19 | 7528671 | 6 | 6 | 460 | 585.7142857 | 0.05434 |
| ALOX15 | chr17 | 4550533 | 16 | 17 | 1471.269841 | 1665.512266 | 0.054761 |
| GCK | chr7 | 44234795 | 36 | 32 | 3294.220593 | 3038.915459 | 0.054863 |
| APOA1 | chr11 | 116692228 | 34 | 32 | 3141.45694 | 3040.400757 | 0.055145 |
| BCL2 | chr18 | 60994110 | 25 | 27 | 2313.072809 | 2656.944444 | 0.055197 |
| E2F6 | chr2 | 11614723 | 35 | 33 | 2560.260765 | 2612.216926 | 0.055318 |
| APOA1 | chr11 | 116707791 | 29 | 29 | 2832.155067 | 2639.982071 | 0.055795 |
| CDKN2B-AS1 | chr9 | 21979306 | 29 | 27 | 1769.355274 | 1381.393378 | 0.055921 |
| APOA1 | chr11 | 116680855 | 33 | 32 | 2969.270559 | 2992.144912 | 0.056002 |
| LPA | chr6 | 11220584 | 34 | 32 | 3263.930984 | 3001.630785 | 0.056004 |
| GALNT2 | chr1 | 230190916 | 35 | 32 | 3168.735476 | 3004.487247 | 0.056544 |
| NOS3 | chr7 | 150680528 | 36 | 31 | 1438.439894 | 1395.535346 | 0.056638 |
| KLF2 | chr19 | 16427833 | 32 | 33 | 2685.65899 | 2603.385101 | 0.056695 |
| FOXP3 | chrX | 49127221 | 24 | 24 | 2189.526307 | 2296.897547 | 0.056815 |
| TERT | chr5 | 1296312 | 34 | 33 | 2697.395363 | 2464.689487 | 0.056861 |
| APOA1 | chr11 | 116682390 | 28 | 25 | 2846.568627 | 2405.46176 | 0.056875 |
| FOXP3 | chrX | 49126934 | 12 | 6 | 608.1204906 | 226.9047619 | 0.057219 |

| | | | | | | | |
|---|---|---|---|---|---|---|---|
| P2RY12 | chr3 | 151104745 | 16 | 19 | 1439.642857 | 1529.873737 | 0.057224 |
| APOA1 | chr11 | 116661811 | 8 | 10 | 206.3492063 | 416.6666667 | 0.057496 |
| LDLR | chr19 | 11220878 | 9 | 9 | 880 | 805.7142857 | 0.057563 |
| FGF2 | chr4 | 123740948 | 33 | 32 | 2729.178143 | 2509.885386 | 0.057864 |
| FOXP3 | chrX | 49106836 | 23 | 24 | 2145.775993 | 2102.02381 | 0.057998 |
| LDLR | chr19 | 11215933 | 31 | 30 | 2941.761757 | 2931.902383 | 0.058283 |
| LOXL1 | chr15 | 74214115 | 34 | 33 | 2568.438242 | 2305.511955 | 0.058416 |
| APOA1 | chr11 | 116701309 | 30 | 31 | 2533.707951 | 2815.984301 | 0.058596 |
| PECAM1 | chr17 | 62401401 | 32 | 33 | 1800.562657 | 2089.961278 | 0.058621 |
| COL15A1 | chr9 | 101705288 | 32 | 32 | 3040.028466 | 2934.43574 | 0.058683 |
| ADRA2C | chr4 | 3759892 | 34 | 32 | 2822.450214 | 2783.842877 | 0.059123 |
| ESR1 | chr6 | 152007105 | 8 | 4 | 660 | 251.4285714 | 0.059125 |
| APOA1 | chr11 | 116668970 | 26 | 24 | 2447.487055 | 2361.666667 | 0.0593 |
| CDKN2B-AS1 | chr9 | 21996128 | 8 | 7 | 302.6984127 | 171.3492063 | 0.059675 |
| GCK | chr7 | 44232197 | 33 | 31 | 2987.31691 | 2911.303621 | 0.059702 |
| PHACTR1 | chr6 | 12716671 | 39 | 32 | 3534.731182 | 2824.600799 | 0.059918 |
| GCK | chr7 | 44229820 | 34 | 32 | 2814.788191 | 2772.595104 | 0.060443 |
| LOXL1 | chr15 | 74211520 | 32 | 28 | 2872.559524 | 2317.081007 | 0.060762 |
| LDLR | chr19 | 11196109 | 33 | 30 | 3100.676833 | 2947.151961 | 0.060799 |
| APOA1 | chr11 | 116681899 | 33 | 32 | 2952.560636 | 2706.817519 | 0.060824 |
| AKT2 | chr19 | 40792708 | 39 | 33 | 2812.416854 | 2223.109414 | 0.060898 |
| CDKN2B-AS1 | chr9 | 21973908 | 32 | 26 | 1082.165933 | 1015.131467 | 0.06096 |
| LDLR | chr19 | 11191216 | 32 | 32 | 2637.888823 | 2797.617453 | 0.060997 |
| AMOTL2 | chr3 | 134100622 | 9 | 8 | 732.7777778 | 735.7142857 | 0.061719 |
| F2RL3 | chr19 | 16998820 | 27 | 32 | 2102.690646 | 2165.907539 | 0.061823 |
| BCL2 | chr18 | 60993745 | 35 | 32 | 3042.35621 | 2879.748371 | 0.061837 |
| KLF2 | chr19 | 16437989 | 32 | 31 | 1415.131634 | 1587.13821 | 0.061842 |
| COL15A1 | chr9 | 101705908 | 5 | 11 | 176.6666667 | 247.4603175 | 0.061978 |
| CDKN2B-AS1 | chr9 | 21977420 | 17 | 15 | 473.9845749 | 543.4593185 | 0.062204 |
| TERT | chr5 | 1301062 | 32 | 32 | 2944.100468 | 2837.223826 | 0.062276 |
| HMGCR | chr5 | 74628723 | 29 | 29 | 2576.146465 | 2713.95513 | 0.062641 |
| APOA1 | chr11 | 116686542 | 38 | 32 | 3264.299384 | 2846.689751 | 0.062785 |
| LCK | chr1 | 32741454 | 10 | 5 | 823.7662338 | 483.3333333 | 0.062937 |
| CDKN2B-AS1 | chr9 | 21983169 | 38 | 33 | 2997.689829 | 2426.941191 | 0.063325 |
| ADRA2C | chr4 | 3761000 | 32 | 29 | 2577.072931 | 2535.205447 | 0.063361 |
| FOXP3 | chrX | 49127325 | 14 | 15 | 1371.428571 | 1387.97619 | 0.063635 |
| MIA3 | chr1 | 222792415 | 19 | 19 | 549.1558442 | 700.5460226 | 0.063904 |
| CTGF | chr6 | 132279495 | 34 | 32 | 3090.177707 | 2998.459901 | 0.064143 |
| CDKN2B-AS1 | chr9 | 21964727 | 17 | 18 | 803.3547009 | 661.0335498 | 0.064165 |
| TFPI | chr2 | 188419831 | 37 | 32 | 3290.559255 | 2934.676683 | 0.064183 |
| ESR1 | chr6 | 152005960 | 30 | 31 | 2808.463137 | 2755.761518 | 0.0645 |
| F7 | chr13 | 113750744 | 27 | 26 | 2492.294372 | 2531.269841 | 0.064535 |
| MCPH1 | chr8 | 6471120 | 8 | 7 | 714.7619048 | 683.3333333 | 0.064719 |
| CDKN2B-AS1 | chr9 | 22003005 | 39 | 33 | 3163.311922 | 2552.804087 | 0.064723 |
| LPA | chr6 | 160952436 | 32 | 31 | 2869.079409 | 2898.895743 | 0.064758 |
| LPA | chr6 | 11224679 | 12 | 15 | 848.8383839 | 1270.31746 | 0.064937 |
| LCK | chr1 | 32741597 | 31 | 32 | 2820.589129 | 3006.816043 | 0.065121 |

| | | | | | | | |
|---|---|---|---|---|---|---|---|
| APOA1 | chr11 | 116700484 | 31 | 32 | 2618.687614 | 2477.976307 | 0.065579 |
| PECAM1 | chr17 | 62410571 | 32 | 32 | 2465.538256 | 2734.977736 | 0.065843 |
| ARHGEF18 | chr19 | 7528638 | 11 | 20 | 1087.5 | 1903.051948 | 0.065859 |
| AMOTL2 | chr3 | 134102205 | 9 | 8 | 755.9981685 | 756.6233766 | 0.066105 |
| TIMP3 | chr22 | 33194281 | 35 | 32 | 2494.412636 | 2435.477943 | 0.066299 |
| CDKN2B-AS1 | chr9 | 21977585 | 15 | 11 | 951.5476191 | 547.1428571 | 0.066542 |
| F7 | chr13 | 113758187 | 26 | 21 | 1639.862915 | 1098.603341 | 0.066642 |
| BCL2 | chr18 | 60988765 | 8 | 3 | 717.5 | 226.6666667 | 0.06668 |
| NOS3 | chr7 | 150691037 | 30 | 30 | 2962.846145 | 2838.29823 | 0.066937 |
| ABCG1 | chr21 | 43630293 | 30 | 30 | 2719.072039 | 2838.308775 | 0.067064 |
| NOS3 | chr7 | 150694561 | 27 | 28 | 2381.94636 | 2313.556166 | 0.067118 |
| TMPO | chr12 | 98905173 | 21 | 22 | 1740.790043 | 1995.198413 | 0.067463 |
| TERT | chr5 | 1299657 | 13 | 9 | 1132.664835 | 865.7142857 | 0.067497 |
| PHACTR1 | chr6 | 12716580 | 38 | 32 | 3452.54322 | 2986.927241 | 0.067518 |
| LOXL1 | chr15 | 74209749 | 38 | 32 | 3713.802287 | 3071.380991 | 0.068161 |
| TERT | chr5 | 1295645 | 9 | 11 | 215.6060606 | 322.2727273 | 0.068162 |
| CDKN2B-AS1 | chr9 | 21998263 | 36 | 33 | 2721.953656 | 2339.348807 | 0.068218 |
| TMPO | chr12 | 98906024 | 33 | 30 | 3237.328809 | 2857.476551 | 0.068377 |
| KLF2 | chr19 | 16429197 | 30 | 32 | 2837.396572 | 2901.61132 | 0.068401 |
| ARHGEF18 | chr19 | 7461935 | 33 | 30 | 3149.170889 | 2732.507009 | 0.068541 |
| NOS3 | chr7 | 150689684 | 26 | 30 | 2165.004325 | 2690.45954 | 0.068543 |
| F7 | chr13 | 113752900 | 31 | 32 | 2929.349538 | 3101.362199 | 0.068743 |
| TERT | chr5 | 1299646 | 8 | 7 | 647.5324675 | 660 | 0.069043 |
| F2RL3 | chr19 | 16991840 | 29 | 30 | 2792.797619 | 2967.857143 | 0.069343 |
| BAX | chr19 | 49465310 | 33 | 33 | 2241.155612 | 2059.50891 | 0.069404 |
| APOA1 | chr11 | 116692347 | 33 | 32 | 3143.441011 | 3114.920693 | 0.070025 |
| TERT | chr5 | 1253492 | 28 | 30 | 2345.72956 | 2671.004371 | 0.070163 |
| F7 | chr13 | 113752992 | 33 | 32 | 3190.301289 | 3012.485372 | 0.070242 |
| CDKN2B-AS1 | chr9 | 21990587 | 24 | 28 | 1325.367965 | 1777.451437 | 0.07054 |
| GALNT2 | chr1 | 230190123 | 29 | 32 | 2511.51724 | 2491.517649 | 0.070722 |
| APOA1 | chr11 | 116669033 | 8 | 5 | 711.4285714 | 500 | 0.0708 |
| TIMP3 | chr22 | 33188545 | 35 | 32 | 3167.620372 | 2708.792117 | 0.071379 |
| PECAM1 | chr17 | 62398076 | 38 | 33 | 2827.826697 | 2306.746918 | 0.071575 |
| LOXL1 | chr15 | 74209251 | 33 | 31 | 2934.119197 | 2854.167708 | 0.071799 |
| CDKN2B-AS1 | chr9 | 22006225 | 15 | 15 | 384.1971182 | 516.6775261 | 0.071857 |
| ADRA2C | chr4 | 3760266 | 37 | 32 | 3342.359554 | 2979.711174 | 0.07186 |
| FOXP3 | chrX | 49129683 | 27 | 28 | 1466.987793 | 1287.713585 | 0.071921 |
| P2RY12 | chr3 | 151104241 | 20 | 18 | 1885.714286 | 1777.083333 | 0.071979 |
| NR3C1 | chr5 | 142785502 | 41 | 32 | 3562.505365 | 2678.620614 | 0.072321 |
| F2RL3 | chr19 | 16991504 | 32 | 32 | 2990.607437 | 3090.839931 | 0.072523 |
| CTGF | chr6 | 132277996 | 34 | 32 | 2884.496372 | 2796.062969 | 0.072605 |
| ICAM2 | chr17 | 62079976 | 31 | 30 | 2839.760446 | 2839.7238 | 0.072715 |
| NOS3 | chr7 | 150709684 | 38 | 32 | 1643.703567 | 1526.963813 | 0.073183 |
| APOA1 | chr11 | 116690835 | 32 | 32 | 2867.863699 | 2734.020261 | 0.073379 |
| FGF2 | chr4 | 123745820 | 39 | 33 | 3484.484783 | 2798.152991 | 0.073639 |
| ATF3 | chr1 | 212739022 | 37 | 32 | 3485.769066 | 3095.029153 | 0.074065 |
| CDKN1A | chr6 | 36652043 | 28 | 30 | 2497.924663 | 2829.409757 | 0.07412 |

| | | | | | | | |
|---|---|---|---|---|---|---|---|
| TERT | chr5 | 1298367 | 23 | 27 | 2001.266234 | 2543.147131 | 0.074122 |
| CDKN2B-AS1 | chr9 | 21971614 | 19 | 17 | 576.7621267 | 660.5769231 | 0.074195 |
| FOXP3 | chrX | 49122719 | 21 | 23 | 1674.004607 | 2019.047619 | 0.07452 |
| HMGCR | chr5 | 74630949 | 16 | 11 | 1358.305861 | 1014.761905 | 0.074738 |
| CTGF | chr6 | 132276028 | 33 | 32 | 3034.936698 | 3035.540855 | 0.075658 |
| APOA1 | chr11 | 116692116 | 32 | 32 | 2881.524904 | 2973.939248 | 0.076005 |
| GALNT2 | chr1 | 230185922 | 35 | 32 | 3008.070656 | 2841.555003 | 0.076077 |
| ESR1 | chr6 | 152006643 | 29 | 27 | 2721.763237 | 2630.079365 | 0.076363 |
| ESR1 | chr6 | 152424240 | 38 | 32 | 3596.554306 | 3092.036392 | 0.076636 |
| LDLR | chr19 | 11221090 | 21 | 18 | 1924.91342 | 1768.888889 | 0.077016 |
| NR3C1 | chr5 | 142790450 | 39 | 32 | 3810.652486 | 2928.247906 | 0.077099 |
| ESR2 | chr14 | 64806777 | 41 | 32 | 4042.0525 | 3092.415655 | 0.077117 |
| APOA1 | chr11 | 116663004 | 32 | 30 | 2529.748311 | 2550.902515 | 0.077504 |
| F2RL3 | chr19 | 16990833 | 34 | 32 | 3266.258742 | 3012.478591 | 0.078123 |
| P2RY12 | chr3 | 151102366 | 34 | 32 | 3121.163181 | 2827.689508 | 0.078203 |
| CTGF | chr6 | 132277931 | 31 | 32 | 2588.631538 | 2857.37187 | 0.078585 |
| APOA1 | chr11 | 116699650 | 34 | 33 | 1872.865719 | 2026.93513 | 0.079324 |
| APOA1 | chr11 | 116705456 | 17 | 13 | 1539.79798 | 1079.722222 | 0.079437 |
| CTGF | chr6 | 132280567 | 32 | 32 | 2855.689845 | 2948.177455 | 0.079495 |
| ADRA2C | chr4 | 3768571 | 14 | 15 | 353.4575754 | 500.3589734 | 0.079516 |
| P2RY12 | chr3 | 151054177 | 35 | 32 | 2811.766921 | 2686.914412 | 0.079579 |
| PLA2G7 | chr6 | 46708816 | 35 | 32 | 3274.610957 | 3068.555416 | 0.079761 |
| LDLR | chr19 | 11219840 | 10 | 16 | 882.5 | 1540.909091 | 0.080498 |
| ATF3 | chr1 | 212739113 | 35 | 32 | 2650.675052 | 2651.293457 | 0.080519 |
| DHFR | chr5 | 79952223 | 17 | 24 | 1664.242424 | 2178.095238 | 0.08076 |
| CTGF | chr6 | 132273158 | 25 | 27 | 2386.465022 | 2473.372343 | 0.080856 |
| LCK | chr1 | 32713395 | 33 | 32 | 3126.097043 | 3114.998117 | 0.081159 |
| APOA1 | chr11 | 116661269 | 32 | 32 | 3008.293039 | 2885.222054 | 0.081165 |
| GALNT2 | chr1 | 230184411 | 32 | 32 | 2631.670405 | 2489.106926 | 0.081283 |
| CTGF | chr6 | 132280453 | 27 | 30 | 2166.300269 | 2589.43744 | 0.081336 |
| ADRA2C | chr4 | 3767076 | 12 | 14 | 291.3221418 | 446.2942464 | 0.081497 |
| ABCG1 | chr21 | 43635794 | 35 | 32 | 3278.680221 | 3066.156324 | 0.081739 |
| LDLR | chr19 | 11219926 | 9 | 6 | 846.6666667 | 600 | 0.081979 |
| LCK | chr1 | 32741927 | 25 | 25 | 2188.492063 | 2318.762626 | 0.082041 |
| ESR2 | chr14 | 64807749 | 32 | 32 | 3105.327945 | 3011.746961 | 0.082121 |
| CDKN2B-AS1 | chr9 | 21993475 | 39 | 33 | 1913.008848 | 1489.601683 | 0.082383 |
| APOA1 | chr11 | 116716940 | 32 | 31 | 2882.398127 | 2914.218657 | 0.082416 |
| BAX | chr19 | 49449527 | 32 | 30 | 3227.5 | 2872.979572 | 0.082641 |
| LDLR | chr19 | 11217536 | 17 | 16 | 1571.976912 | 1552.777778 | 0.082701 |
| APOA1 | chr11 | 116692225 | 33 | 32 | 3092.599161 | 3071.013663 | 0.082758 |
| ABCG1 | chr21 | 43632094 | 15 | 20 | 1443.246753 | 1807.857143 | 0.082901 |
| APOA1 | chr11 | 116692506 | 33 | 32 | 3070.324504 | 3049.220919 | 0.082904 |
| NOS3 | chr7 | 150680054 | 33 | 32 | 2268.968703 | 2338.637389 | 0.083155 |
| ESR2 | chr14 | 64811848 | 17 | 17 | 1621.672772 | 1467.326007 | 0.083244 |
| FOXP3 | chrX | 49106442 | 17 | 19 | 1658.492063 | 1754.365079 | 0.083335 |
| CDKN2B-AS1 | chr9 | 21974183 | 16 | 16 | 401.24123 | 525.5350325 | 0.083563 |
| F7 | chr13 | 113750372 | 25 | 24 | 2298.860878 | 2061.135254 | 0.08364 |

| | | | | | | | |
|---|---|---|---|---|---|---|---|
| LDLR | chr19 | 11193171 | 32 | 32 | 2980.224972 | 3079.529282 | 0.083897 |
| APOA1 | chr11 | 116662518 | 12 | 11 | 943.7229437 | 987.7777778 | 0.084064 |
| APOA1 | chr11 | 116660072 | 30 | 30 | 2919.883825 | 2785.034565 | 0.084243 |
| F2RL3 | chr19 | 16990856 | 34 | 32 | 3189.827357 | 3079.159729 | 0.0843 |
| PHACTR1 | chr6 | 12711039 | 33 | 32 | 3044.44676 | 3049.729573 | 0.084364 |
| LDLR | chr19 | 11217834 | 17 | 17 | 1700 | 1662.103175 | 0.084418 |
| P2RY12 | chr3 | 151106556 | 32 | 33 | 2198.15318 | 2404.721764 | 0.084421 |
| TIMP3 | chr22 | 33196884 | 13 | 10 | 1183.333333 | 983.3333333 | 0.084596 |
| FGF2 | chr4 | 123740831 | 36 | 32 | 2993.435239 | 2819.405492 | 0.084616 |
| GALNT2 | chr1 | 230190894 | 35 | 32 | 3200.662892 | 3002.620033 | 0.084636 |
| LDLR | chr19 | 11191618 | 10 | 8 | 956.547619 | 675.4761905 | 0.08484 |
| TERT | chr5 | 1299088 | 28 | 32 | 2307.31352 | 2303.33088 | 0.085378 |
| CDKN2B-AS1 | chr9 | 21996146 | 14 | 11 | 537.2366522 | 325.7142857 | 0.085497 |
| ADRA2C | chr4 | 3759220 | 33 | 30 | 2712.132858 | 2628.343319 | 0.085719 |
| LOXL1 | chr15 | 74209147 | 34 | 32 | 3212.913174 | 3093.358926 | 0.085901 |
| BAX | chr19 | 49449007 | 34 | 32 | 3083.631202 | 2981.215604 | 0.086358 |
| GALNT2 | chr1 | 230190314 | 29 | 32 | 2458.081003 | 2941.9331 | 0.086443 |
| DHFR | chr5 | 79952175 | 32 | 33 | 2974.41522 | 2899.776387 | 0.086484 |
| COL15A1 | chr9 | 101705718 | 34 | 33 | 2655.660251 | 2439.64053 | 0.086562 |
| PHACTR1 | chr6 | 12711842 | 34 | 32 | 2743.009343 | 2697.158251 | 0.08676 |
| NOS3 | chr7 | 150706760 | 23 | 23 | 1390.965399 | 1560.600511 | 0.086775 |
| ADRA2C | chr4 | 3759773 | 32 | 32 | 2460.671254 | 2773.853263 | 0.087056 |
| ADRA2C | chr4 | 3763913 | 32 | 31 | 3014.193935 | 2792.669414 | 0.087078 |
| ESR2 | chr14 | 64806465 | 31 | 30 | 3014.95098 | 2994.736842 | 0.087363 |
| TIMP3 | chr22 | 33193578 | 37 | 32 | 3476.698271 | 3058.705765 | 0.087578 |
| APOA1 | chr11 | 116697788 | 31 | 32 | 2662.382203 | 2596.175207 | 0.087602 |
| PECAM1 | chr17 | 62399312 | 34 | 31 | 2847.294738 | 2731.097528 | 0.087857 |
| LPA | chr6 | 160952215 | 7 | 5 | 637.1428571 | 400.4761905 | 0.087861 |
| P2RY12 | chr3 | 151102837 | 25 | 22 | 733.4774356 | 759.0860859 | 0.088105 |
| AMOTL2 | chr3 | 134103861 | 10 | 4 | 734.6428571 | 351.4285714 | 0.088161 |
| APOA1 | chr11 | 116698401 | 21 | 20 | 1857.849928 | 1880.663781 | 0.088301 |
| ESR1 | chr6 | 152007595 | 11 | 13 | 1054.603175 | 1139.047619 | 0.088876 |
| GCK | chr7 | 44230901 | 36 | 33 | 2198.783764 | 1873.775371 | 0.089018 |
| COL15A1 | chr9 | 101704737 | 28 | 30 | 1782.281572 | 2265.880011 | 0.089637 |
| LDLR | chr19 | 11197559 | 14 | 14 | 1367.5 | 1295.714286 | 0.089838 |
| ICAM2 | chr17 | 62080169 | 33 | 31 | 3190.208408 | 3056.708847 | 0.089844 |
| LCK | chr1 | 32741015 | 28 | 33 | 1908.408593 | 2475.210928 | 0.089898 |
| TERT | chr5 | 1298856 | 26 | 26 | 2304.739011 | 2155.788101 | 0.090077 |
| FOXP3 | chrX | 49115386 | 31 | 31 | 2869.242462 | 3002.812702 | 0.090376 |
| LDLR | chr19 | 11198491 | 17 | 21 | 1684.343434 | 2014.166667 | 0.09038 |
| MCPH1 | chr8 | 6470364 | 32 | 32 | 2851.498257 | 2584.042449 | 0.090682 |
| GCK | chr7 | 44229723 | 32 | 31 | 2655.35578 | 2704.618293 | 0.090684 |
| ATF3 | chr1 | 212748178 | 31 | 32 | 2985.041933 | 2971.755409 | 0.090922 |
| APOA1 | chr11 | 116692625 | 32 | 32 | 2506.201684 | 2656.837647 | 0.091303 |
| F7 | chr13 | 113753723 | 37 | 32 | 3523.12899 | 2985.206907 | 0.091403 |
| AKT2 | chr19 | 40796372 | 35 | 32 | 3411.289077 | 3063.27313 | 0.091502 |
| ALOX15 | chr17 | 4548869 | 36 | 32 | 2634.289331 | 2483.625564 | 0.091757 |

| | | | | | | | |
|---|---|---|---|---|---|---|---|
| FOXP3 | chrX | 49120188 | 30 | 31 | 2061.651645 | 2289.402447 | 0.091821 |
| KLF2 | chr19 | 16430174 | 13 | 9 | 1206.666667 | 885.7142857 | 0.092016 |
| F7 | chr13 | 113752960 | 32 | 32 | 3119.69712 | 2976.945138 | 0.092021 |
| GALNT2 | chr1 | 230184841 | 35 | 32 | 3005.477669 | 2842.247124 | 0.092044 |
| GCK | chr7 | 44237456 | 34 | 32 | 3090.850699 | 2982.508063 | 0.092338 |
| ADRA2C | chr4 | 3760989 | 32 | 29 | 2891.211007 | 2749.067234 | 0.092498 |
| PHACTR1 | chr6 | 12708270 | 32 | 32 | 2385.04944 | 2562.031609 | 0.09262 |
| NOS3 | chr7 | 150683752 | 32 | 31 | 1712.088659 | 1852.276736 | 0.092699 |
| LCK | chr1 | 32716457 | 28 | 31 | 1422.52225 | 1770.713903 | 0.093078 |
| APOA1 | chr11 | 116714753 | 7 | 8 | 700 | 750.8333333 | 0.093164 |
| KLF2 | chr19 | 16437986 | 31 | 30 | 1442.559046 | 1601.77069 | 0.093465 |
| BAX | chr19 | 49450713 | 19 | 28 | 1866.752137 | 2632.267275 | 0.094736 |
| CTGF | chr6 | 132280576 | 33 | 32 | 2509.931548 | 2575.439388 | 0.094959 |
| LCK | chr1 | 32741460 | 15 | 11 | 1323.484848 | 1060 | 0.094995 |
| F7 | chr13 | 113752177 | 30 | 31 | 2652.281387 | 2513.2913 | 0.094997 |
| AMOTL2 | chr3 | 134074448 | 27 | 29 | 2513.264375 | 2806.170696 | 0.095357 |
| AMOTL2 | chr3 | 134095170 | 30 | 31 | 2333.691253 | 2557.859973 | 0.095404 |
| APOA1 | chr11 | 116713675 | 35 | 32 | 3259.811021 | 2901.939095 | 0.095443 |
| CDKN2B-AS1 | chr9 | 21986001 | 38 | 33 | 2410.790836 | 1978.81581 | 0.095779 |
| TMPO | chr12 | 98906222 | 33 | 32 | 3157.227958 | 2795.526885 | 0.095962 |
| APOA1 | chr11 | 116679808 | 34 | 33 | 1977.680024 | 2081.52588 | 0.096361 |
| LDLR | chr19 | 11225023 | 34 | 32 | 3343.727822 | 3062.11194 | 0.096519 |
| APOA1 | chr11 | 116720383 | 37 | 33 | 2411.193846 | 2038.980851 | 0.096642 |
| BAX | chr19 | 49448199 | 34 | 32 | 3116.166104 | 3019.63378 | 0.097016 |
| COL15A1 | chr9 | 101704377 | 32 | 32 | 2511.288962 | 2660.054113 | 0.097075 |
| F2RL3 | chr19 | 16994927 | 22 | 18 | 2001.904762 | 1739.75469 | 0.097944 |
| APOA1 | chr11 | 116704199 | 24 | 28 | 2196.167027 | 2679.55267 | 0.097956 |
| GALNT2 | chr1 | 230191020 | 31 | 32 | 2539.749312 | 2776.679476 | 0.098223 |
| ALOX15 | chr17 | 4552312 | 36 | 33 | 1816.736388 | 1802.622457 | 0.098263 |
| MCPH1 | chr8 | 6470544 | 34 | 32 | 3144.469037 | 3028.700183 | 0.098276 |
| PECAM1 | chr17 | 62400603 | 28 | 32 | 2391.422207 | 2958.880678 | 0.098357 |
| LDLR | chr19 | 11216686 | 5 | 5 | 475 | 406.6666667 | 0.098496 |
| PECAM1 | chr17 | 62410468 | 14 | 18 | 1159.642857 | 1631.067821 | 0.098523 |
| ALOX15 | chr17 | 4547475 | 37 | 32 | 3061.925673 | 2750.843301 | 0.098778 |
| CDKN2B-AS1 | chr9 | 21968510 | 3 | 7 | 61.42857143 | 219.3406593 | 0.098798 |
| MCPH1 | chr8 | 6256628 | 22 | 26 | 1916.070694 | 2418.888889 | 0.098899 |
| ALDOA | chr16 | 30061571 | 30 | 31 | 2748.371263 | 2952.249972 | 0.098978 |
| TERT | chr5 | 1297822 | 15 | 17 | 1338.203463 | 1618.809524 | 0.098996 |
| ESR1 | chr6 | 152002595 | 19 | 25 | 1022.722833 | 1150.498668 | 0.099057 |
| F2RL3 | chr19 | 16990952 | 30 | 30 | 2817.236652 | 2720.501549 | 0.099536 |
| ICAM2 | chr17 | 62087768 | 6 | 3 | 373.3333333 | 250.7142857 | 0.100076 |
| FOXP3 | chrX | 49129231 | 31 | 32 | 1989.071207 | 2230.167478 | 0.100079 |
| CDKN2B-AS1 | chr9 | 21989915 | 6 | 5 | 236.0714286 | 137.2222222 | 0.100363 |
| GCK | chr7 | 44229002 | 35 | 32 | 3105.652585 | 2946.139681 | 0.100501 |
| CDKN2B-AS1 | chr9 | 21968514 | 6 | 7 | 172.4358974 | 165 | 0.100981 |
| APOA1 | chr11 | 116698785 | 35 | 32 | 3001.258197 | 2841.307657 | 0.101062 |
| ADRA2C | chr4 | 3761692 | 35 | 32 | 2786.970144 | 2668.52742 | 0.101101 |

| | | | | | | | |
|---|---|---|---|---|---|---|---|
| PECAM1 | chr17 | 62398282 | 34 | 32 | 2772.372348 | 2487.656583 | 0.101158 |
| GALNT2 | chr1 | 230188418 | 18 | 23 | 1259.776335 | 1881.576479 | 0.101275 |
| ABCG1 | chr21 | 43717642 | 40 | 32 | 3629.85562 | 2981.166077 | 0.101405 |
| TERT | chr5 | 1300423 | 34 | 31 | 2869.540932 | 2745.742566 | 0.101416 |
| FGF2 | chr4 | 123741459 | 35 | 32 | 3148.328744 | 2806.188576 | 0.101777 |
| TERT | chr5 | 1298607 | 27 | 28 | 2101.07206 | 2342.032273 | 0.101938 |
| ADRA2C | chr4 | 3761941 | 24 | 28 | 2070.071595 | 2215.940285 | 0.101978 |
| CDKN2B-AS1 | chr9 | 22007276 | 10 | 9 | 615.2014652 | 435.5952381 | 0.102018 |
| TFPI | chr2 | 188424204 | 3 | 3 | 269.047619 | 231.4285714 | 0.102243 |
| LDLR | chr19 | 11198535 | 28 | 30 | 2856.036839 | 2865.613806 | 0.102322 |
| GCK | chr7 | 44237447 | 33 | 32 | 3070.571693 | 3029.351957 | 0.102344 |
| AMOTL2 | chr3 | 134100882 | 35 | 32 | 3300.013024 | 3094.115525 | 0.102356 |
| BAX | chr19 | 49449689 | 31 | 32 | 2847.216685 | 3029.244515 | 0.102777 |
| AMOTL2 | chr3 | 134102010 | 8 | 9 | 708.021978 | 900 | 0.102935 |
| CDKN2B-AS1 | chr9 | 21977176 | 32 | 33 | 1446.056298 | 1642.224801 | 0.103582 |
| TERT | chr5 | 1299481 | 26 | 28 | 2440.599567 | 2750 | 0.103744 |
| CDKN1A | chr6 | 36654065 | 12 | 9 | 343.6769006 | 386.8073593 | 0.104063 |
| ABCG1 | chr21 | 43634650 | 24 | 28 | 2286.616162 | 2557.25469 | 0.104102 |
| APOA1 | chr11 | 116661491 | 29 | 32 | 2490.789627 | 2891.632145 | 0.104297 |
| KLF2 | chr19 | 16433377 | 32 | 33 | 2886.64776 | 2835.904489 | 0.104502 |
| APOA1 | chr11 | 116721496 | 32 | 32 | 3188.428755 | 3099.935729 | 0.104621 |
| APOA1 | chr11 | 116692314 | 33 | 31 | 3031.778394 | 2926.516994 | 0.104723 |
| TMPO | chr12 | 98906049 | 34 | 32 | 3213.836498 | 3098.0899 | 0.104805 |
| APOA1 | chr11 | 116674211 | 28 | 30 | 2470.596163 | 2791.323693 | 0.104942 |
| FGF2 | chr4 | 123747634 | 18 | 8 | 618.546176 | 380.3968254 | 0.10502 |
| LDLR | chr19 | 11218108 | 26 | 22 | 2600.85213 | 2124.84127 | 0.105043 |
| LCK | chr1 | 32713272 | 32 | 32 | 3048.314475 | 2956.283857 | 0.105617 |
| LDLR | chr19 | 11191299 | 20 | 27 | 1795.045788 | 2187.97619 | 0.105843 |
| LPA | chr6 | 160952386 | 23 | 26 | 1844.563492 | 2247.91904 | 0.106296 |
| LDLR | chr19 | 11216531 | 29 | 25 | 2624.242424 | 2363.021619 | 0.106342 |
| CTGF | chr6 | 132276337 | 33 | 32 | 3001.192584 | 2985.425612 | 0.10642 |
| ATF3 | chr1 | 212742184 | 29 | 26 | 2829.410867 | 2427.709513 | 0.106423 |
| LPA | chr6 | 11217012 | 35 | 32 | 3361.10521 | 3020.80836 | 0.106796 |
| APOA1 | chr11 | 116661185 | 32 | 32 | 2817.082678 | 2957.612321 | 0.107018 |
| APOA1 | chr11 | 116707297 | 18 | 21 | 1708.928571 | 1891.190476 | 0.107265 |
| NOS3 | chr7 | 150691993 | 8 | 9 | 192.2222222 | 332.1825397 | 0.107739 |
| LOXL1 | chr15 | 74214633 | 12 | 14 | 615 | 540.7936508 | 0.108121 |
| NR3C1 | chr5 | 142790715 | 33 | 32 | 3126.968194 | 3093.623407 | 0.10848 |
| ALDOA | chr16 | 30061890 | 20 | 22 | 1731.933622 | 2032.91514 | 0.108545 |
| FOXP3 | chrX | 49128435 | 32 | 32 | 2940.176662 | 3025.417742 | 0.108721 |
| TERT | chr5 | 1295726 | 13 | 10 | 391.6085545 | 377.0192308 | 0.108761 |
| AMOTL2 | chr3 | 134097010 | 35 | 33 | 2499.894379 | 2185.199005 | 0.109003 |
| ARHGEF18 | chr19 | 7466756 | 36 | 32 | 3519.644779 | 3084.520496 | 0.10908 |
| LCK | chr1 | 32741192 | 16 | 13 | 499.0079365 | 521.1255411 | 0.109119 |
| NOS3 | chr7 | 150680387 | 36 | 32 | 2764.650145 | 2565.104641 | 0.109484 |
| NOS3 | chr7 | 150695753 | 29 | 29 | 2487.623273 | 2319.144401 | 0.110384 |
| GCK | chr7 | 44231698 | 30 | 30 | 2382.665974 | 2554.251027 | 0.110423 |

| | | | | | | | |
|---|---|---|---|---|---|---|---|
| CDKN2B-AS1 | chr9 | 21964664 | 16 | 21 | 487.6509602 | 777.2319055 | 0.110481 |
| MCPH1 | chr8 | 6259113 | 35 | 32 | 3243.529414 | 3029.501309 | 0.110558 |
| ABCG1 | chr21 | 43629876 | 36 | 32 | 3302.710226 | 2998.961089 | 0.11058 |
| F2RL3 | chr19 | 16995380 | 16 | 13 | 1468.333333 | 1256.547619 | 0.110638 |
| KLF2 | chr19 | 16433616 | 33 | 32 | 3060.653819 | 3057.055014 | 0.110641 |
| LPA | chr6 | 11216495 | 36 | 32 | 3474.961601 | 3029.090798 | 0.111283 |
| FOXP3 | chrX | 49115206 | 20 | 21 | 1605.47619 | 1827.261905 | 0.111481 |
| NR3C1 | chr5 | 142788777 | 37 | 32 | 3440.839562 | 3055.040851 | 0.111484 |
| APOA1 | chr11 | 116719845 | 34 | 32 | 3177.665346 | 3063.607392 | 0.111525 |
| F7 | chr13 | 113756749 | 30 | 30 | 2497.48022 | 2317.094368 | 0.11208 |
| E2F6 | chr2 | 11613612 | 34 | 32 | 2886.665642 | 2599.995886 | 0.112177 |
| KLF2 | chr19 | 16433618 | 33 | 32 | 3050.909052 | 3047.821604 | 0.112576 |
| APOA1 | chr11 | 116708880 | 9 | 15 | 783.2142857 | 1099.444444 | 0.112741 |
| MCPH1 | chr8 | 6470667 | 32 | 32 | 3131.324473 | 3051.550457 | 0.112804 |
| LDLR | chr19 | 11190651 | 31 | 32 | 2683.496286 | 2559.005072 | 0.112996 |
| CTGF | chr6 | 132282390 | 32 | 32 | 2253.042313 | 2419.878676 | 0.113237 |
| CDKN2B-AS1 | chr9 | 22010312 | 32 | 29 | 1418.400022 | 1133.043547 | 0.11332 |
| F7 | chr13 | 113750160 | 32 | 32 | 3098.560961 | 3006.059755 | 0.113721 |
| APOA1 | chr11 | 116661212 | 31 | 32 | 2821.048675 | 3005.040715 | 0.114038 |
| APOA1 | chr11 | 116687615 | 32 | 32 | 2874.63727 | 2979.086904 | 0.114078 |
| ABCA1 | chr9 | 107690707 | 3 | 3 | 106.6666667 | 86.9047619 | 0.115098 |
| APOA1 | chr11 | 116716943 | 32 | 31 | 2946.152856 | 2765.688429 | 0.115123 |
| APOA1 | chr11 | 116663186 | 32 | 32 | 2605.724365 | 2766.019389 | 0.11514 |
| APOA1 | chr11 | 116692204 | 33 | 32 | 2898.414189 | 2908.434613 | 0.115143 |
| ESR2 | chr14 | 64806870 | 36 | 32 | 3560.825488 | 3115.12487 | 0.11516 |
| APOA1 | chr11 | 116688663 | 10 | 13 | 866.1471862 | 994.7619048 | 0.115202 |
| KLF2 | chr19 | 16438278 | 20 | 17 | 539.5741462 | 574.2470554 | 0.115244 |
| PHACTR1 | chr6 | 12717941 | 39 | 32 | 3488.348467 | 2932.740983 | 0.1153 |
| LCK | chr1 | 32740637 | 15 | 14 | 1466.666667 | 1308.214286 | 0.115356 |
| LDLR | chr19 | 11190984 | 34 | 32 | 3337.801361 | 3055.523671 | 0.115601 |
| ADRA2C | chr4 | 3763426 | 32 | 32 | 2964.920676 | 2851.830434 | 0.115798 |
| LOXL1 | chr15 | 74209503 | 4 | 2 | 340 | 200 | 0.116118 |
| BCL2 | chr18 | 60988367 | 20 | 25 | 534.7153218 | 793.9745539 | 0.11612 |
| CDKN2B-AS1 | chr9 | 21967573 | 21 | 17 | 530.5846583 | 538.3437902 | 0.116221 |
| PLA2G7 | chr6 | 46704232 | 36 | 32 | 3227.859814 | 2934.505827 | 0.116339 |
| LOXL1 | chr15 | 74219172 | 24 | 25 | 943.3848559 | 826.5732858 | 0.116578 |
| NOS3 | chr7 | 150692033 | 24 | 19 | 840.7859121 | 812.0616883 | 0.116596 |
| F2RL3 | chr19 | 16998721 | 17 | 19 | 551.3095238 | 725.6313131 | 0.116676 |
| APOA1 | chr11 | 116708942 | 25 | 22 | 2165.172328 | 2022.45671 | 0.117016 |
| TERT | chr5 | 1299529 | 25 | 30 | 2349.63925 | 2927.777778 | 0.117019 |
| TERT | chr5 | 1298240 | 33 | 32 | 3081.531631 | 3060.009112 | 0.117063 |
| APOA1 | chr11 | 116690235 | 35 | 32 | 3201.687782 | 2993.963797 | 0.117175 |
| NOS3 | chr7 | 150678265 | 32 | 33 | 1052.738116 | 1204.26293 | 0.117515 |
| NOS3 | chr7 | 150684071 | 27 | 31 | 1988.966173 | 2506.200936 | 0.117516 |
| CDKN1A | chr6 | 36652339 | 20 | 26 | 1950.761905 | 2465.442109 | 0.117796 |
| BAX | chr19 | 49465121 | 32 | 31 | 3050.916871 | 3051.211201 | 0.117858 |
| LCK | chr1 | 32740801 | 22 | 21 | 2025.47619 | 1821.663059 | 0.117882 |

| | | | | | | | |
|---|---|---|---|---|---|---|---|
| APOA1 | chr11 | 116708254 | 29 | 27 | 2657.164421 | 2337.009592 | 0.118005 |
| LDLR | chr19 | 11193211 | 34 | 32 | 3290.489537 | 3043.223403 | 0.118236 |
| FOXP3 | chrX | 49121984 | 6 | 6 | 518.3333333 | 421.4285714 | 0.118316 |
| ADRA2C | chr4 | 3766640 | 13 | 13 | 495.4229104 | 624.4805195 | 0.118484 |
| CDKN2B-AS1 | chr9 | 21996744 | 8 | 4 | 411.9047619 | 159.0476191 | 0.118521 |
| ADRA2C | chr4 | 3766535 | 23 | 26 | 2081.566656 | 2218.960623 | 0.118635 |
| ABCG1 | chr21 | 43635088 | 32 | 32 | 3170.372928 | 3058.072945 | 0.11868 |
| F7 | chr13 | 113758669 | 20 | 17 | 1703.333333 | 1550.524476 | 0.119043 |
| ARHGEF18 | chr19 | 7461159 | 9 | 15 | 744.6031746 | 1360.436508 | 0.119402 |
| P2RY12 | chr3 | 151112530 | 34 | 32 | 3278.096601 | 3143.35539 | 0.119717 |
| LDLR | chr19 | 11219829 | 18 | 18 | 1576.825397 | 1691.067821 | 0.119738 |
| F7 | chr13 | 113753008 | 33 | 32 | 3162.114046 | 3006.820493 | 0.119836 |
| APOA1 | chr11 | 116726074 | 34 | 33 | 1793.441076 | 1593.52341 | 0.119956 |
| ABCB1 | chr7 | 87347851 | 37 | 32 | 3218.183313 | 2868.45321 | 0.12006 |
| LOXL1 | chr15 | 74218780 | 20 | 18 | 588.9645497 | 653.9866785 | 0.120155 |
| APOA1 | chr11 | 116705982 | 6 | 7 | 222.5 | 186.9047619 | 0.120259 |
| TERT | chr5 | 1295716 | 20 | 18 | 683.5243188 | 521.9385027 | 0.120316 |
| LOXL1 | chr15 | 74212124 | 32 | 32 | 1409.185965 | 1559.502348 | 0.120782 |
| CDKN2B-AS1 | chr9 | 21993973 | 17 | 18 | 498.1782814 | 627.0315273 | 0.120859 |
| APOA1 | chr11 | 116682396 | 28 | 26 | 2787.042541 | 2469.009324 | 0.121023 |
| F2RL3 | chr19 | 16993466 | 33 | 32 | 3071.194462 | 3037.635954 | 0.121041 |
| MCPH1 | chr8 | 6263119 | 36 | 33 | 1840.086349 | 1846.475168 | 0.12134 |
| PECAM1 | chr17 | 62401957 | 10 | 9 | 934.1666667 | 887.5 | 0.121382 |
| ADRA2C | chr4 | 3764459 | 8 | 10 | 775 | 890 | 0.121384 |
| CDKN2B-AS1 | chr9 | 21997858 | 38 | 33 | 2902.22515 | 2363.702939 | 0.121462 |
| APOA1 | chr11 | 116720122 | 28 | 30 | 2762.519841 | 2736.243952 | 0.121942 |
| AMOTL2 | chr3 | 134100056 | 31 | 32 | 1373.754905 | 1261.308551 | 0.122061 |
| APOA1 | chr11 | 116674031 | 36 | 32 | 3038.482561 | 2600.518621 | 0.122302 |
| APOA1 | chr11 | 116668111 | 11 | 9 | 1058.021978 | 754.2424242 | 0.122401 |
| LDLR | chr19 | 11198141 | 35 | 32 | 3418.462547 | 2995.493058 | 0.122555 |
| APOA1 | chr11 | 116707415 | 11 | 7 | 1083.333333 | 652.2222222 | 0.12292 |
| KLF2 | chr19 | 16429302 | 35 | 32 | 3169.405555 | 2985.703138 | 0.123097 |
| MCPH1 | chr8 | 6260624 | 32 | 31 | 2647.621226 | 2699.989233 | 0.123236 |
| LDLR | chr19 | 11190590 | 17 | 24 | 1616.742424 | 2356.910867 | 0.123361 |
| GCK | chr7 | 44236483 | 4 | 9 | 80 | 297.3520924 | 0.123457 |
| CDKN2B-AS1 | chr9 | 21970928 | 4 | 3 | 93.84615385 | 108.3333333 | 0.12386 |
| APOA1 | chr11 | 116680675 | 13 | 13 | 1249.047619 | 1153.333333 | 0.124218 |
| ABCG1 | chr21 | 43637265 | 32 | 31 | 3091.11588 | 3052.196679 | 0.124423 |
| F7 | chr13 | 113750921 | 32 | 32 | 3143.232791 | 2953.957431 | 0.124523 |
| ALOX15 | chr17 | 4548832 | 33 | 33 | 2008.247353 | 2142.414437 | 0.124657 |
| ABCA1 | chr9 | 107695294 | 38 | 32 | 3312.061341 | 2868.305644 | 0.124697 |
| F2RL3 | chr19 | 16997271 | 28 | 23 | 2672.45671 | 2263.131313 | 0.124863 |
| ABCG1 | chr21 | 43637596 | 20 | 16 | 1885.595238 | 1578.021978 | 0.125103 |
| ARHGEF18 | chr19 | 7537834 | 7 | 7 | 692.8571429 | 643.3333333 | 0.125204 |
| F7 | chr13 | 113753752 | 36 | 32 | 3286.441988 | 3005.793532 | 0.125237 |
| LPA | chr6 | 160945027 | 36 | 33 | 3328.274553 | 2928.989899 | 0.125458 |
| APOA1 | chr11 | 116662358 | 30 | 33 | 2417.244862 | 2520.928697 | 0.125637 |

| | | | | | | | |
|---|---|---|---|---|---|---|---|
| LPA | chr6 | 160950004 | 36 | 32 | 3040.549797 | 2593.821843 | 0.125758 |
| TMPO | chr12 | 98899407 | 31 | 33 | 1441.59768 | 1727.878313 | 0.12576 |
| APOA1 | chr11 | 116709897 | 33 | 33 | 3066.986844 | 2931.244924 | 0.126656 |
| ADRA2C | chr4 | 3766807 | 24 | 22 | 1229.42507 | 1283.470696 | 0.127244 |
| CDKN2B-AS1 | chr9 | 21993583 | 33 | 26 | 1273.109204 | 859.1103545 | 0.127345 |
| NOS3 | chr7 | 150706111 | 32 | 32 | 1722.829408 | 1874.917702 | 0.127665 |
| AMOTL2 | chr3 | 134101778 | 23 | 26 | 2039.509542 | 2468.021978 | 0.127697 |
| LOXL1 | chr15 | 74218767 | 21 | 20 | 623.9714671 | 752.126931 | 0.127764 |
| MCPH1 | chr8 | 6259252 | 34 | 32 | 3091.286131 | 2991.064782 | 0.127864 |
| CDKN1A | chr6 | 36651923 | 14 | 14 | 363.5706547 | 450.4853558 | 0.128296 |
| LCK | chr1 | 32707622 | 17 | 13 | 1437.28022 | 1193.333333 | 0.128556 |
| HMGCR | chr5 | 74625119 | 37 | 32 | 3083.479619 | 2751.500151 | 0.128603 |
| ABCG1 | chr21 | 43634522 | 20 | 26 | 1976.666667 | 2476.980519 | 0.128658 |
| CDKN1A | chr6 | 36654244 | 19 | 18 | 1692.496392 | 1696.969697 | 0.128765 |
| LDLR | chr19 | 11216999 | 10 | 12 | 987.5 | 1130.833333 | 0.12892 |
| APOA1 | chr11 | 116660672 | 32 | 32 | 3002.953049 | 3088.934654 | 0.12896 |
| APOA1 | chr11 | 116691271 | 36 | 33 | 3064.785327 | 2675.148415 | 0.129142 |
| F2RL3 | chr19 | 17000087 | 13 | 9 | 396.8401405 | 358.5409035 | 0.129902 |
| PHACTR1 | chr6 | 12716207 | 32 | 32 | 2393.608467 | 2561.324305 | 0.130063 |
| PECAM1 | chr17 | 62410154 | 33 | 32 | 3005.270706 | 2763.654191 | 0.130242 |
| FGF2 | chr4 | 123744550 | 26 | 26 | 2443.848472 | 2521.780997 | 0.131096 |
| APOA1 | chr11 | 116686681 | 39 | 32 | 3593.144731 | 3001.687008 | 0.131681 |
| LDLR | chr19 | 11216665 | 8 | 9 | 800 | 831.4285714 | 0.132017 |
| LPA | chr6 | 160943049 | 36 | 32 | 3395.921257 | 2960.154403 | 0.132381 |
| NR3C1 | chr5 | 142785811 | 32 | 32 | 3015.920531 | 3122.204486 | 0.1326 |
| APOA1 | chr11 | 116701849 | 31 | 31 | 3106.788475 | 2959.012807 | 0.132962 |
| F2RL3 | chr19 | 16991810 | 26 | 23 | 2257.084305 | 2124.296814 | 0.13316 |
| ARHGEF18 | chr19 | 7537376 | 32 | 32 | 2424.577344 | 2214.813947 | 0.133162 |
| AMOTL2 | chr3 | 134095059 | 28 | 29 | 1949.787117 | 2207.582671 | 0.133339 |
| APOA1 | chr11 | 116697311 | 32 | 32 | 3100.592438 | 3035.979815 | 0.133642 |
| MCPH1 | chr8 | 6470795 | 33 | 31 | 2995.855325 | 2926.533827 | 0.134063 |
| NOS3 | chr7 | 150686325 | 22 | 19 | 1982.380952 | 1586.190476 | 0.134237 |
| F7 | chr13 | 113753484 | 37 | 32 | 3450.516154 | 2921.213735 | 0.134839 |
| ESR1 | chr6 | 152009078 | 33 | 32 | 3093.08948 | 2928.394077 | 0.13498 |
| APOA1 | chr11 | 116697149 | 27 | 30 | 2547.519841 | 2691.797924 | 0.135016 |
| TIMP3 | chr22 | 33194250 | 33 | 33 | 1499.803893 | 1629.08464 | 0.135098 |
| PHACTR1 | chr6 | 12717072 | 32 | 32 | 2840.248439 | 2942.95309 | 0.135336 |
| E2F6 | chr2 | 11612506 | 3 | 2 | 100 | 150 | 0.135342 |
| GCK | chr7 | 44234903 | 34 | 32 | 3028.267982 | 2926.995513 | 0.135585 |
| APOA1 | chr11 | 116663826 | 39 | 32 | 3190.709273 | 2545.536916 | 0.13584 |
| ABCB1 | chr7 | 87348277 | 24 | 24 | 1891.593407 | 1707.13401 | 0.136259 |
| ABCA1 | chr9 | 107691451 | 5 | 5 | 110 | 179.2673993 | 0.136278 |
| CDKN2B-AS1 | chr9 | 22006027 | 11 | 11 | 261.3818861 | 371.020646 | 0.136301 |
| COL15A1 | chr9 | 101696537 | 26 | 28 | 1793.210953 | 2167.242706 | 0.136402 |
| ADRA2C | chr4 | 3763776 | 32 | 32 | 2812.603432 | 2718.85979 | 0.136821 |
| TMPO | chr12 | 98908203 | 19 | 16 | 440.6162468 | 478.4612867 | 0.137016 |
| DHFR | chr5 | 79955781 | 21 | 19 | 541.7627571 | 586.8307717 | 0.137076 |

| | | | | | | | |
|---|---|---|---|---|---|---|---|
| DHFR | chr5 | 79958040 | 2 | 3 | 130 | 280 | 0.138237 |
| F7 | chr13 | 113753244 | 34 | 33 | 2745.95733 | 2523.938766 | 0.138385 |
| ATF3 | chr1 | 212783900 | 28 | 31 | 2599.423654 | 2763.223325 | 0.138481 |
| ABCA1 | chr9 | 107691597 | 38 | 32 | 3251.257627 | 2855.691168 | 0.138483 |
| GCK | chr7 | 44230832 | 33 | 31 | 2941.660467 | 2893.861153 | 0.138523 |
| APOA1 | chr11 | 116695307 | 27 | 28 | 2523.035714 | 2711.944444 | 0.1387 |
| F2RL3 | chr19 | 16990367 | 32 | 32 | 3047.666901 | 3118.712815 | 0.138779 |
| DHFR | chr5 | 79953698 | 34 | 32 | 3266.101147 | 3020.023478 | 0.139 |
| LDLR | chr19 | 11190227 | 3 | 7 | 300 | 627.9761905 | 0.139098 |
| APOA1 | chr11 | 116701028 | 32 | 33 | 2804.515586 | 2732.294496 | 0.139278 |
| F7 | chr13 | 113760223 | 32 | 30 | 2902.116428 | 2821.928174 | 0.139336 |
| BCL2 | chr18 | 60993286 | 32 | 32 | 3006.076878 | 3070.194642 | 0.139343 |
| LDLR | chr19 | 11197407 | 5 | 5 | 485.7142857 | 437.7777778 | 0.139457 |
| APOA1 | chr11 | 116674188 | 27 | 32 | 2435.71578 | 3006.20671 | 0.139479 |
| CDKN2B-AS1 | chr9 | 22011535 | 38 | 32 | 3466.25819 | 2806.620818 | 0.139756 |
| MIA3 | chr1 | 222808706 | 38 | 32 | 3585.240905 | 3078.184614 | 0.139878 |
| F7 | chr13 | 113752972 | 32 | 32 | 3092.472033 | 2987.852598 | 0.140076 |
| PLA2G7 | chr6 | 46712321 | 31 | 32 | 1508.056643 | 1415.931735 | 0.140138 |
| NR3C1 | chr5 | 142786505 | 38 | 32 | 3706.019399 | 3151.006981 | 0.140142 |
| KLF2 | chr19 | 16426775 | 17 | 19 | 1397.904889 | 1386.904762 | 0.140159 |
| APOA1 | chr11 | 116713823 | 34 | 32 | 3282.173262 | 2996.016673 | 0.140484 |
| CTGF | chr6 | 132277934 | 33 | 32 | 2783.574143 | 2825.807101 | 0.140623 |
| LOXL1 | chr15 | 74218826 | 24 | 17 | 738.9307003 | 628.7894136 | 0.14064 |
| LDLR | chr19 | 11197945 | 11 | 16 | 996.119175 | 1570.833333 | 0.140964 |
| TIMP3 | chr22 | 33191569 | 37 | 32 | 3405.479147 | 3015.43893 | 0.14114 |
| CDKN2B-AS1 | chr9 | 21974187 | 12 | 12 | 298.6331969 | 386.7253138 | 0.141381 |
| APOA1 | chr11 | 116709941 | 33 | 32 | 3146.991041 | 2977.856422 | 0.141859 |
| CTGF | chr6 | 132275740 | 32 | 33 | 1006.796201 | 1144.44197 | 0.142083 |
| ESR1 | chr6 | 152005614 | 38 | 32 | 3611.597223 | 3081.723987 | 0.142117 |
| AMOTL2 | chr3 | 134096526 | 25 | 29 | 2270.553401 | 2503.79218 | 0.142517 |
| NOS3 | chr7 | 150706366 | 28 | 26 | 2346.623335 | 2300.47948 | 0.143082 |
| TERT | chr5 | 1301840 | 27 | 28 | 2645.387488 | 2606.219336 | 0.143083 |
| COL15A1 | chr9 | 101703521 | 38 | 32 | 3126.418106 | 2729.993934 | 0.14318 |
| TERT | chr5 | 1301124 | 34 | 32 | 3308.73421 | 3018.813606 | 0.143382 |
| F7 | chr13 | 113752153 | 32 | 31 | 2992.486336 | 2786.187163 | 0.143565 |
| MCPH1 | chr8 | 6256505 | 33 | 32 | 2996.272755 | 2988.350472 | 0.143781 |
| LDLR | chr19 | 11222463 | 31 | 32 | 2932.670158 | 3124.279716 | 0.143795 |
| F2RL3 | chr19 | 16990983 | 26 | 29 | 2520.900129 | 2692.776668 | 0.143965 |
| APOA1 | chr11 | 116661345 | 31 | 32 | 2689.258116 | 2862.718898 | 0.144021 |
| ARHGEF18 | chr19 | 7469098 | 33 | 32 | 3220.028945 | 3070.529548 | 0.144199 |
| ICAM2 | chr17 | 62086890 | 22 | 23 | 891.2003274 | 772.3115773 | 0.144204 |
| LDLR | chr19 | 11191950 | 4 | 4 | 326.6666667 | 286.6666667 | 0.144297 |
| ADRA2C | chr4 | 3760015 | 31 | 33 | 2468.47983 | 2425.891587 | 0.144458 |
| ATF3 | chr1 | 212741715 | 35 | 32 | 3085.382646 | 2892.863928 | 0.144997 |
| ADRA2C | chr4 | 3764637 | 31 | 31 | 2673.899572 | 2812.971016 | 0.145217 |
| NOS3 | chr7 | 150706059 | 33 | 33 | 1947.197525 | 2106.018733 | 0.145444 |
| TIMP3 | chr22 | 33195502 | 38 | 32 | 3558.830227 | 3036.144471 | 0.145561 |

| | | | | | | | |
|---|---|---|---|---|---|---|---|
| LDLR | chr19 | 11199476 | 33 | 32 | 2638.698234 | 2673.319786 | 0.145799 |
| TERT | chr5 | 1298305 | 32 | 32 | 2900.372016 | 3014.312819 | 0.145944 |
| NOS3 | chr7 | 150690936 | 20 | 19 | 1749.794372 | 1766.944444 | 0.145963 |
| APOA1 | chr11 | 116713107 | 34 | 32 | 3359.777612 | 3069.884946 | 0.146136 |
| LOXL1 | chr15 | 74218410 | 33 | 33 | 2872.128344 | 2726.016864 | 0.146201 |
| TIMP3 | chr22 | 33259061 | 20 | 26 | 1788.582529 | 2427.183927 | 0.146278 |
| NOS3 | chr7 | 150710718 | 8 | 7 | 320 | 209.7222222 | 0.146398 |
| BAX | chr19 | 49453114 | 32 | 30 | 2629.542221 | 2591.169562 | 0.146801 |
| LPA | chr6 | 11216709 | 36 | 32 | 3454.621345 | 3011.183906 | 0.146857 |
| LOXL1 | chr15 | 74214297 | 32 | 31 | 2655.795979 | 2675.652364 | 0.146881 |
| CDKN2BAS | chr9 | 22103975 | 36 | 32 | 3328.874492 | 2886.677203 | 0.147681 |
| CDKN2B-AS1 | chr9 | 21980697 | 31 | 31 | 2760.8586 | 2544.504845 | 0.147722 |
| APOA1 | chr11 | 116707100 | 34 | 31 | 3148.068029 | 2950.506007 | 0.147804 |
| APOA1 | chr11 | 116661314 | 31 | 32 | 2504.479622 | 2714.068877 | 0.147823 |
| KLF2 | chr19 | 16437995 | 24 | 30 | 832.2912966 | 1181.59653 | 0.14804 |
| CDKN2B-AS1 | chr9 | 21964438 | 32 | 33 | 1517.427092 | 1712.579113 | 0.148084 |
| ADRA2C | chr4 | 3760642 | 33 | 33 | 2217.670596 | 2077.849508 | 0.148264 |
| CDKN2B-AS1 | chr9 | 21997383 | 32 | 32 | 2699.12988 | 2588.647232 | 0.148299 |
| NR3C1 | chr5 | 142791398 | 29 | 29 | 2769.520202 | 2863.63345 | 0.148577 |
| TIMP3 | chr22 | 33196238 | 33 | 32 | 2962.575727 | 2959.090879 | 0.148762 |
| APOA1 | chr11 | 116692519 | 33 | 32 | 2997.673103 | 2987.241068 | 0.148882 |
| FGF2 | chr4 | 123747226 | 34 | 33 | 2876.804423 | 2667.270926 | 0.149141 |
| CDKN2B-AS1 | chr9 | 22011381 | 38 | 32 | 2776.343477 | 2430.308666 | 0.149438 |
| ALDOA | chr16 | 30060643 | 34 | 32 | 3217.754929 | 2950.751673 | 0.149497 |
| F7 | chr13 | 113758817 | 31 | 33 | 2073.759055 | 2016.027411 | 0.149763 |
| LDLR | chr19 | 11222069 | 31 | 32 | 2631.221718 | 2563.644243 | 0.150021 |
| PECAM1 | chr17 | 62399389 | 22 | 25 | 1959.837662 | 2083.350538 | 0.150042 |
| KLF2 | chr19 | 16438016 | 32 | 32 | 1588.382839 | 1734.610319 | 0.150562 |
| ALDOA | chr16 | 30057611 | 19 | 26 | 1546.43899 | 2259.264763 | 0.150765 |
| BCL2 | chr18 | 60990103 | 9 | 7 | 211.7298053 | 258.7200957 | 0.151115 |
| ARHGEF18 | chr19 | 7461834 | 35 | 32 | 3438.915878 | 3104.768135 | 0.151196 |
| NOS3 | chr7 | 150707679 | 22 | 18 | 593.7578626 | 569.8572408 | 0.151443 |
| KLF2 | chr19 | 16438063 | 26 | 29 | 892.1618969 | 1111.631662 | 0.151662 |
| LDLR | chr19 | 11217942 | 29 | 29 | 2798.145933 | 2864.305556 | 0.151737 |
| APOA1 | chr11 | 116680442 | 32 | 32 | 2548.604899 | 2665.097186 | 0.151922 |
| MIA3 | chr1 | 222810293 | 17 | 15 | 1568.409091 | 1455 | 0.152135 |
| E2F6 | chr2 | 11613071 | 25 | 24 | 2387.972958 | 2360 | 0.152462 |
| AKT2 | chr19 | 40792378 | 29 | 28 | 2711.465216 | 2524.055112 | 0.152743 |
| F7 | chr13 | 113752079 | 32 | 32 | 3010.137451 | 2928.372071 | 0.152879 |
| LPA | chr6 | 11224588 | 16 | 14 | 1160.357143 | 1171.785714 | 0.153016 |
| F2RL3 | chr19 | 16997358 | 31 | 32 | 2730.525148 | 2935.83969 | 0.153176 |
| BAX | chr19 | 49448226 | 37 | 32 | 3058.233033 | 2755.649633 | 0.153404 |
| LDLR | chr19 | 11216634 | 17 | 19 | 1649.95671 | 1771.385281 | 0.153443 |
| F7 | chr13 | 113751051 | 19 | 17 | 1899.603175 | 1611.428571 | 0.153616 |
| LCK | chr1 | 32741652 | 31 | 32 | 2898.265983 | 3056.637524 | 0.153755 |
| ALOX15 | chr17 | 4554100 | 40 | 32 | 3210.55819 | 2643.038393 | 0.153884 |
| KLF2 | chr19 | 16430279 | 34 | 33 | 1872.250786 | 1666.055286 | 0.154076 |

| | | | | | | | |
|---|---|---|---|---|---|---|---|
| LCK | chr1 | 32741543 | 32 | 33 | 3124.349293 | 3047.292149 | 0.154162 |
| CDKN2BAS | chr9 | 22068040 | 14 | 15 | 1186.984127 | 1372.02381 | 0.154199 |
| LDLR | chr19 | 11194157 | 10 | 15 | 906.6666667 | 1465.714286 | 0.154263 |
| LDLR | chr19 | 11193754 | 17 | 19 | 1520.984848 | 1804.444444 | 0.154357 |
| LDLR | chr19 | 11190691 | 35 | 32 | 3315.258563 | 3089.607298 | 0.154358 |
| PHACTR1 | chr6 | 12711477 | 33 | 32 | 2337.984637 | 2397.933094 | 0.154462 |
| F7 | chr13 | 113758142 | 31 | 30 | 2377.203352 | 2135.037793 | 0.154543 |
| MCPH1 | chr8 | 6470839 | 34 | 31 | 3190.160124 | 2982.038943 | 0.154556 |
| FOXP3 | chrX | 49120455 | 18 | 18 | 1709.563492 | 1765.714286 | 0.155116 |
| APOA1 | chr11 | 116680048 | 25 | 25 | 2368.681139 | 2282.460317 | 0.155282 |
| APOA1 | chr11 | 116692688 | 33 | 32 | 2786.643138 | 2817.277616 | 0.155625 |
| LDLR | chr19 | 11197547 | 15 | 15 | 1455.555556 | 1393.333333 | 0.155977 |
| NOS3 | chr7 | 150706607 | 27 | 28 | 2455.985681 | 2406.495449 | 0.156025 |
| TERT | chr5 | 1300643 | 34 | 32 | 2859.337821 | 2794.164457 | 0.15612 |
| FOXP3 | chrX | 49121007 | 28 | 28 | 2243.256263 | 2376.529582 | 0.156221 |
| DHFR | chr5 | 79960306 | 32 | 31 | 3003.187888 | 2850.447941 | 0.156455 |
| BCL2 | chr18 | 60989664 | 30 | 25 | 1062.179349 | 1009.050434 | 0.156721 |
| GCK | chr7 | 44231744 | 2 | 4 | 82.67973856 | 263.0869131 | 0.156795 |
| ABCA1 | chr9 | 107698076 | 12 | 16 | 1106.269841 | 1363.174603 | 0.156804 |
| TIMP3 | chr22 | 33197034 | 34 | 33 | 2134.336896 | 2211.855565 | 0.157025 |
| TIMP3 | chr22 | 33193664 | 34 | 32 | 2539.635327 | 2292.340929 | 0.157099 |
| CDKN2B-AS1 | chr9 | 21966399 | 7 | 6 | 163.8850039 | 261.9047619 | 0.157155 |
| F2RL3 | chr19 | 16990407 | 31 | 32 | 2878.796011 | 3049.912269 | 0.157198 |
| LOXL1 | chr15 | 74211069 | 35 | 32 | 3290.637479 | 3063.141811 | 0.157256 |
| FOXP3 | chrX | 49126920 | 10 | 9 | 383.0952381 | 440.4761905 | 0.157319 |
| NOS3 | chr7 | 150705794 | 35 | 32 | 1739.620468 | 1744.695072 | 0.157435 |
| COL15A1 | chr9 | 101705402 | 30 | 32 | 2717.102339 | 3005.692781 | 0.157478 |
| TERT | chr5 | 1301251 | 34 | 33 | 2310.981384 | 2371.576882 | 0.157559 |
| TERT | chr5 | 1296337 | 34 | 32 | 3125.39789 | 2993.593323 | 0.157657 |
| NOS3 | chr7 | 150707718 | 31 | 31 | 1387.68514 | 1236.523357 | 0.157719 |
| AKT2 | chr19 | 40792065 | 13 | 15 | 1012.020202 | 1334.642857 | 0.157782 |
| APOA1 | chr11 | 116668975 | 26 | 24 | 2459.214952 | 2338.928571 | 0.158456 |
| LCK | chr1 | 32740402 | 28 | 28 | 2573.075105 | 2453.286713 | 0.158462 |
| LDLR | chr19 | 11223321 | 23 | 22 | 2104.361472 | 2105.47619 | 0.158499 |
| FOXP3 | chrX | 49118127 | 12 | 13 | 1156.547619 | 1190 | 0.158541 |
| CDKN2B-AS1 | chr9 | 21968453 | 16 | 13 | 461.9191919 | 495.030525 | 0.158616 |
| NOS3 | chr7 | 150681817 | 38 | 32 | 3180.723225 | 2788.614021 | 0.158798 |
| CDKN2B-AS1 | chr9 | 22009937 | 2 | 2 | 40 | 58.33333333 | 0.158804 |
| LPA | chr6 | 11216231 | 34 | 32 | 3316.024096 | 3078.677911 | 0.158976 |
| APOA1 | chr11 | 116659692 | 32 | 28 | 2984.314125 | 2689.684343 | 0.159043 |
| CDKN2B-AS1 | chr9 | 21977458 | 11 | 16 | 378.0532213 | 447.5108225 | 0.159456 |
| APOA1 | chr11 | 116661885 | 32 | 31 | 2318.835012 | 2380.626624 | 0.159885 |
| TERT | chr5 | 1300308 | 27 | 29 | 2609.532828 | 2664.35824 | 0.159919 |
| FGF2 | chr4 | 123740630 | 5 | 5 | 443.3333333 | 485.7142857 | 0.159961 |
| F2RL3 | chr19 | 16998115 | 15 | 14 | 1040.915751 | 819.1666667 | 0.16002 |
| NOS3 | chr7 | 150683008 | 11 | 10 | 756.8542569 | 568.2326007 | 0.160063 |
| F2RL3 | chr19 | 16995403 | 7 | 5 | 594.7619048 | 483.3333333 | 0.160119 |

| | | | | | | | |
|---|---|---|---|---|---|---|---|
| TERT | chr5 | 1299509 | 27 | 30 | 2559.818841 | 2940.909091 | 0.160216 |
| APOA1 | chr11 | 116706091 | 15 | 10 | 399.6459096 | 356.2606838 | 0.160757 |
| APOA1 | chr11 | 116701216 | 28 | 28 | 1942.083195 | 2138.445166 | 0.160923 |
| APOA1 | chr11 | 116682358 | 32 | 31 | 2944.65876 | 2953.200434 | 0.160977 |
| LCK | chr1 | 32708485 | 32 | 33 | 1967.231683 | 1879.328322 | 0.161242 |
| PECAM1 | chr17 | 62398399 | 31 | 33 | 2824.083103 | 2826.999133 | 0.161383 |
| F2RL3 | chr19 | 16995077 | 10 | 17 | 960.8333333 | 1688.888889 | 0.161424 |
| ICAM2 | chr17 | 62086626 | 32 | 30 | 3010.371404 | 2917.09671 | 0.161521 |
| APOA1 | chr11 | 116690988 | 32 | 32 | 2292.995076 | 2134.598911 | 0.161702 |
| F7 | chr13 | 113750935 | 31 | 29 | 3072.406205 | 2684.615801 | 0.161741 |
| FGF2 | chr4 | 123747558 | 27 | 29 | 1266.828332 | 1220.910204 | 0.161984 |
| APOA1 | chr11 | 116661945 | 33 | 32 | 2870.168506 | 2870.070323 | 0.161995 |
| APOA1 | chr11 | 116661360 | 32 | 32 | 2519.057575 | 2663.489555 | 0.162116 |
| ALDOA | chr16 | 30057140 | 32 | 32 | 2695.375089 | 2555.634746 | 0.162143 |
| ARHGEF18 | chr19 | 7465022 | 36 | 32 | 3344.022663 | 2912.576614 | 0.162544 |
| APOA1 | chr11 | 116707075 | 33 | 33 | 2080.004424 | 1950.101919 | 0.163159 |
| ESR2 | chr14 | 64812846 | 35 | 32 | 2871.505577 | 2711.810521 | 0.16322 |
| LDLR | chr19 | 11217442 | 32 | 30 | 2908.283689 | 2820.350965 | 0.16334 |
| MIA3 | chr1 | 222810211 | 22 | 22 | 2002.086466 | 1806.785714 | 0.163397 |
| ADRA2C | chr4 | 3765969 | 29 | 31 | 2681.368325 | 2958.362082 | 0.163499 |
| F7 | chr13 | 113751557 | 30 | 29 | 2937.198773 | 2643.742846 | 0.163605 |
| F7 | chr13 | 113757071 | 33 | 32 | 3113.023685 | 2941.961864 | 0.163676 |
| AKT2 | chr19 | 40736271 | 32 | 33 | 1765.988757 | 1966.50466 | 0.163678 |
| APOA1 | chr11 | 116714411 | 32 | 32 | 3235.697384 | 3113.814622 | 0.163683 |
| ESR1 | chr6 | 152006559 | 11 | 11 | 1091.666667 | 1046.428571 | 0.163725 |
| ALDOA | chr16 | 30061429 | 32 | 31 | 3005.328283 | 3026.418484 | 0.163735 |
| APOA1 | chr11 | 116710752 | 33 | 32 | 2986.073905 | 2983.328288 | 0.164075 |
| PHACTR1 | chr6 | 12713540 | 32 | 32 | 2889.10476 | 2798.094909 | 0.164361 |
| AKT2 | chr19 | 40798099 | 21 | 21 | 1778.253968 | 1893.293651 | 0.164576 |
| APOA1 | chr11 | 116694447 | 29 | 29 | 2639.836552 | 2731.514319 | 0.16458 |
| GALNT2 | chr1 | 230187901 | 15 | 16 | 1426.547619 | 1435.15873 | 0.164696 |
| APOA1 | chr11 | 116661907 | 32 | 32 | 1977.277818 | 1864.769551 | 0.165162 |
| BCL2 | chr18 | 60988335 | 13 | 11 | 327.8145844 | 383.4887195 | 0.165282 |
| APOA1 | chr11 | 116706047 | 12 | 15 | 477.9545455 | 484.2965368 | 0.165381 |
| CDKN2B-AS1 | chr9 | 21965458 | 13 | 12 | 323.0721278 | 382.9145299 | 0.165496 |
| NOS3 | chr7 | 150707313 | 14 | 11 | 1106.904762 | 992.5 | 0.165697 |
| NOS3 | chr7 | 150689693 | 26 | 31 | 2357.849753 | 2943.884865 | 0.165756 |
| BCL2 | chr18 | 60988257 | 10 | 20 | 285.778813 | 702.5648423 | 0.165784 |
| ALOX15 | chr17 | 4545266 | 6 | 4 | 282.2222222 | 127.5 | 0.165796 |
| LDLR | chr19 | 11217315 | 33 | 32 | 3265.555617 | 3104.82856 | 0.165804 |
| NOS3 | chr7 | 150690672 | 12 | 15 | 1040.865801 | 1386.623377 | 0.16584 |
| LDLR | chr19 | 11223326 | 21 | 21 | 1918.266178 | 2000.833333 | 0.165983 |
| APOA1 | chr11 | 116674582 | 29 | 32 | 1740.651291 | 2091.094739 | 0.166003 |
| CDKN2B-AS1 | chr9 | 21968485 | 4 | 6 | 103.5714286 | 133.015873 | 0.166117 |
| NOS3 | chr7 | 150686593 | 37 | 33 | 2198.996988 | 2062.036648 | 0.166364 |
| LDLR | chr19 | 11217157 | 7 | 13 | 633.3333333 | 1276.388889 | 0.166463 |
| LPA | chr6 | 11218719 | 36 | 32 | 3454.54626 | 3016.180158 | 0.167 |

| | | | | | | | |
|---|---|---|---|---|---|---|---|
| F2RL3 | chr19 | 16990701 | 30 | 32 | 2839.471691 | 3102.103416 | 0.167097 |
| LDLR | chr19 | 11194473 | 30 | 29 | 2921.50216 | 2738.383109 | 0.167121 |
| APOA1 | chr11 | 116674498 | 22 | 21 | 1731.453824 | 1797.9329 | 0.167124 |
| NOS3 | chr7 | 150695523 | 19 | 29 | 1063.03285 | 1844.828505 | 0.167383 |
| F7 | chr13 | 113753282 | 34 | 33 | 2399.486297 | 2185.763388 | 0.167518 |
| ESR2 | chr14 | 64806499 | 4 | 4 | 326.6666667 | 380 | 0.167717 |
| LDLR | chr19 | 11221353 | 33 | 32 | 3147.8224 | 3106.27091 | 0.168039 |
| APOA1 | chr11 | 116709139 | 32 | 32 | 2944.79019 | 3051.36544 | 0.16848 |
| AKT2 | chr19 | 40792602 | 40 | 32 | 3736.441654 | 3035.812832 | 0.168595 |
| LDLR | chr19 | 11193108 | 15 | 13 | 1361.428571 | 1269.047619 | 0.168683 |
| BCL2 | chr18 | 60993080 | 37 | 32 | 3645.128304 | 3093.485304 | 0.16874 |
| LCK | chr1 | 32740912 | 23 | 26 | 1984.858059 | 2080.214508 | 0.168762 |
| APOA1 | chr11 | 116714700 | 24 | 26 | 2330.909091 | 2580 | 0.168938 |
| ADRA2C | chr4 | 3760870 | 4 | 9 | 400 | 860 | 0.16902 |
| AMOTL2 | chr3 | 134094342 | 30 | 33 | 1266.402577 | 1528.186565 | 0.169276 |
| NOS3 | chr7 | 150684620 | 16 | 20 | 1364.274892 | 1818.690476 | 0.169505 |
| APOA1 | chr11 | 116674179 | 28 | 31 | 2563.553391 | 2722.08153 | 0.169537 |
| NR3C1 | chr5 | 142791133 | 19 | 17 | 909.2026813 | 711.3534865 | 0.169736 |
| CDKN2B-AS1 | chr9 | 21996786 | 7 | 5 | 272.0238095 | 153.0555556 | 0.169838 |
| LDLR | chr19 | 11223943 | 24 | 21 | 2295.972222 | 2063.131313 | 0.170159 |
| ABCA1 | chr9 | 107690724 | 5 | 5 | 166.6666667 | 127.2222222 | 0.170345 |
| FOXP3 | chrX | 49126826 | 7 | 4 | 165 | 123.3333333 | 0.170363 |
| CDKN1A | chr6 | 36653580 | 26 | 25 | 1225.568476 | 1019.822677 | 0.170436 |
| LCK | chr1 | 32740718 | 22 | 18 | 2149.673203 | 1706.428571 | 0.170444 |
| ALOX15 | chr17 | 4548997 | 32 | 33 | 2264.760578 | 2186.07167 | 0.170504 |
| FOXP3 | chrX | 49114344 | 32 | 32 | 2526.648641 | 2399.462138 | 0.170644 |
| CDKN2B-AS1 | chr9 | 21967470 | 5 | 8 | 126.0822511 | 313.2539683 | 0.170659 |
| ARHGEF18 | chr19 | 7469114 | 34 | 32 | 3029.695742 | 2934.007504 | 0.170741 |
| KLF2 | chr19 | 16429114 | 31 | 30 | 2915.669814 | 2900.978564 | 0.171301 |
| APOA1 | chr11 | 116692113 | 32 | 32 | 3020.612824 | 3077.569098 | 0.171343 |
| APOA1 | chr11 | 116703147 | 18 | 19 | 578.3716354 | 719.5827287 | 0.171402 |
| LDLR | chr19 | 11224974 | 32 | 32 | 2833.305254 | 2916.324236 | 0.17148 |
| APOA1 | chr11 | 116704001 | 35 | 32 | 3322.359546 | 3081.598887 | 0.171542 |
| BCL2 | chr18 | 60988272 | 16 | 22 | 448.4071011 | 732.2129286 | 0.171956 |
| ALOX15 | chr17 | 4545914 | 29 | 32 | 1302.070241 | 1597.27163 | 0.172104 |
| ABCG1 | chr21 | 43638169 | 8 | 9 | 785.7142857 | 840.9090909 | 0.172297 |
| ABCA1 | chr9 | 107696642 | 3 | 7 | 300 | 666.6666667 | 0.172304 |
| LDLR | chr19 | 11195865 | 5 | 5 | 396.6666667 | 305 | 0.172503 |
| TERT | chr5 | 1298962 | 31 | 31 | 1902.564173 | 1746.384594 | 0.172577 |
| KLF2 | chr19 | 16438117 | 31 | 31 | 1351.761854 | 1532.514756 | 0.172758 |
| TMPO | chr12 | 98901620 | 25 | 25 | 2318.524531 | 2210.121545 | 0.172881 |
| ALOX15 | chr17 | 4547169 | 35 | 33 | 2538.377124 | 2521.563457 | 0.173138 |
| ALOX15 | chr17 | 4533909 | 9 | 11 | 900 | 1065.714286 | 0.173363 |
| ARHGEF18 | chr19 | 7528620 | 12 | 20 | 1153.214286 | 1985.714286 | 0.173421 |
| APOA1 | chr11 | 116714413 | 32 | 32 | 3194.423267 | 3070.242273 | 0.173659 |
| F7 | chr13 | 113752919 | 31 | 32 | 2904.629467 | 3066.059504 | 0.173885 |
| TERT | chr5 | 1300047 | 32 | 33 | 2701.605042 | 2656.725223 | 0.173964 |

| | | | | | | | |
|---|---|---|---|---|---|---|---|
| E2F6 | chr2 | 11615057 | 33 | 32 | 3233.500034 | 3082.474859 | 0.174225 |
| ADRA2C | chr4 | 3758297 | 31 | 31 | 3112.222279 | 2897.531423 | 0.17424 |
| BAX | chr19 | 49454345 | 4 | 5 | 295.7142857 | 270 | 0.174497 |
| F2RL3 | chr19 | 16999113 | 28 | 30 | 2218.498008 | 2211.444071 | 0.174545 |
| F7 | chr13 | 113751490 | 28 | 30 | 2652.072095 | 2741.267401 | 0.174643 |
| ALDOA | chr16 | 30060826 | 32 | 32 | 3020.310523 | 3100.32986 | 0.174964 |
| APOA1 | chr11 | 116705002 | 24 | 24 | 2300.051893 | 2360 | 0.17506 |
| TERT | chr5 | 1295916 | 28 | 30 | 1541.023614 | 1399.38412 | 0.175364 |
| CDKN2B-AS1 | chr9 | 21969344 | 27 | 26 | 1066.976354 | 915.0308473 | 0.175561 |
| F7 | chr13 | 113755434 | 30 | 33 | 2776.553193 | 2906.80406 | 0.175719 |
| ARHGEF18 | chr19 | 7469656 | 35 | 32 | 3223.372039 | 3030.410732 | 0.176404 |
| P2RY12 | chr3 | 151104988 | 8 | 13 | 408.0952381 | 814.5634921 | 0.176875 |
| F7 | chr13 | 113755214 | 6 | 10 | 526.6666667 | 1000 | 0.176883 |
| APOA1 | chr11 | 116710765 | 34 | 32 | 3132.311414 | 3022.06657 | 0.176938 |
| PHACTR1 | chr6 | 12708737 | 35 | 32 | 3021.090612 | 2834.603591 | 0.177123 |
| F2RL3 | chr19 | 16993607 | 31 | 31 | 2489.668284 | 2630.47535 | 0.177395 |
| CTGF | chr6 | 132274005 | 36 | 32 | 3290.626135 | 2868.26275 | 0.178196 |
| APOA1 | chr11 | 116688320 | 32 | 33 | 2598.9608 | 2550.0142 | 0.17824 |
| CDKN2B-AS1 | chr9 | 21976207 | 3 | 6 | 81.9047619 | 133.5714286 | 0.178544 |
| TERT | chr5 | 1296281 | 34 | 33 | 2821.134367 | 2617.722109 | 0.178563 |
| NOS3 | chr7 | 150693966 | 10 | 15 | 300.5663781 | 580.2597403 | 0.178676 |
| LOXL1 | chr15 | 74209141 | 34 | 32 | 3152.148878 | 3051.566362 | 0.178683 |
| P2RY12 | chr3 | 151107792 | 35 | 32 | 3404.528487 | 3036.083147 | 0.178705 |
| CDKN2B-AS1 | chr9 | 21966495 | 10 | 13 | 249.5662233 | 416.2385911 | 0.178863 |
| GCK | chr7 | 44183411 | 37 | 32 | 3491.316609 | 2955.097436 | 0.178963 |
| AKT2 | chr19 | 40795514 | 25 | 30 | 2302.1313 | 2845.739122 | 0.178997 |
| KLF2 | chr19 | 16437958 | 31 | 30 | 1616.217514 | 1744.073382 | 0.179095 |
| ADRA2C | chr4 | 3765676 | 20 | 21 | 1178.773449 | 1086.273449 | 0.179157 |
| CDKN2B-AS1 | chr9 | 21970775 | 32 | 31 | 1192.422318 | 1272.574309 | 0.17948 |
| ESR2 | chr14 | 64807819 | 32 | 32 | 3040.470444 | 3118.413135 | 0.179736 |
| CDKN2B-AS1 | chr9 | 22001078 | 32 | 31 | 1757.773493 | 1537.870724 | 0.179816 |
| APOA1 | chr11 | 116714458 | 33 | 32 | 3381.514727 | 3065.512327 | 0.179979 |
| NOS3 | chr7 | 150688881 | 24 | 28 | 1992.050033 | 2163.560606 | 0.180059 |
| LDLR | chr19 | 11198660 | 37 | 32 | 3327.208258 | 2782.482963 | 0.180076 |
| CTGF | chr6 | 132279465 | 34 | 32 | 2914.976817 | 2832.189698 | 0.180137 |
| ESR1 | chr6 | 152002618 | 28 | 33 | 1826.474205 | 2321.456232 | 0.180184 |
| APOA1 | chr11 | 116717432 | 32 | 33 | 3002.691166 | 2976.701408 | 0.18038 |
| ABCB1 | chr7 | 87343533 | 28 | 26 | 2662.978688 | 2540.313853 | 0.180599 |
| FOXP3 | chrX | 49130541 | 31 | 27 | 2844.11368 | 2573.529901 | 0.180844 |
| F2RL3 | chr19 | 16991286 | 32 | 32 | 2896.789747 | 2990.023072 | 0.180998 |
| HMGCR | chr5 | 74628241 | 37 | 32 | 3226.898149 | 2856.31837 | 0.181003 |
| CDKN2B-AS1 | chr9 | 21989946 | 5 | 5 | 131.9047619 | 195 | 0.181136 |
| NOS3 | chr7 | 150706626 | 28 | 29 | 2659.487057 | 2604.85681 | 0.181358 |
| MCPH1 | chr8 | 6260004 | 12 | 10 | 1014.761905 | 946.6666667 | 0.181622 |
| LDLR | chr19 | 11217661 | 6 | 4 | 600 | 366.6666667 | 0.181695 |
| CTGF | chr6 | 132280086 | 40 | 33 | 2191.072836 | 1704.38221 | 0.18182 |
| TERT | chr5 | 1298345 | 29 | 29 | 2625.345904 | 2712.102065 | 0.182316 |

| | | | | | | | |
|---|---|---|---|---|---|---|---|
| ARHGEF18 | chr19 | 7467732 | 38 | 32 | 3534.493042 | 3025.665238 | 0.182419 |
| LCK | chr1 | 32713614 | 34 | 32 | 3120.900232 | 3008.722365 | 0.182461 |
| GALNT2 | chr1 | 230190780 | 35 | 32 | 2918.829501 | 2731.726068 | 0.182541 |
| NOS3 | chr7 | 150709518 | 20 | 22 | 1864.688645 | 2154.166667 | 0.182562 |
| COL15A1 | chr9 | 101697369 | 34 | 33 | 2242.041769 | 2077.331893 | 0.182619 |
| APOA1 | chr11 | 116711496 | 36 | 32 | 3343.445111 | 3024.508861 | 0.182922 |
| LCK | chr1 | 32740999 | 29 | 30 | 2701.00751 | 2701.28401 | 0.18296 |
| CDKN1A | chr6 | 36653304 | 32 | 33 | 3010.95959 | 2964.748773 | 0.182983 |
| MCPH1 | chr8 | 6470518 | 34 | 32 | 2862.156628 | 2784.418252 | 0.183042 |
| CTGF | chr6 | 132275447 | 5 | 8 | 113.0751042 | 290.5877456 | 0.183081 |
| ABCG1 | chr21 | 43633401 | 32 | 32 | 2713.509202 | 2809.943696 | 0.183118 |
| ADRA2C | chr4 | 3759141 | 32 | 32 | 2318.648179 | 2450.533437 | 0.183318 |
| LCK | chr1 | 32741487 | 25 | 24 | 2371.940837 | 2197.178932 | 0.183539 |
| F7 | chr13 | 113751032 | 22 | 20 | 2211.25 | 1860.036075 | 0.183583 |
| BCL2 | chr18 | 60988268 | 10 | 13 | 254.8214003 | 439.1916417 | 0.183676 |
| APOA1 | chr11 | 116701854 | 30 | 31 | 2942.92812 | 2882.220221 | 0.183697 |
| MCPH1 | chr8 | 6470884 | 30 | 30 | 2843.36784 | 2905.38273 | 0.183944 |
| APOA1 | chr11 | 116716768 | 36 | 32 | 3378.750645 | 2950.079277 | 0.184035 |
| COL15A1 | chr9 | 101705378 | 30 | 32 | 2844.279594 | 2962.810992 | 0.184181 |
| CDKN1A | chr6 | 36653628 | 31 | 29 | 1306.256277 | 1083.230565 | 0.184237 |
| AKT2 | chr19 | 40800291 | 7 | 4 | 700 | 363.3333333 | 0.184305 |
| FOXP3 | chrX | 49117830 | 4 | 6 | 363.3333333 | 600 | 0.184305 |
| CDKN2B-AS1 | chr9 | 21968586 | 5 | 5 | 105.1821862 | 177.6960784 | 0.184382 |
| LPA | chr6 | 11222977 | 34 | 33 | 1687.949481 | 1766.102431 | 0.184536 |
| LDLR | chr19 | 11219058 | 13 | 11 | 851.1904762 | 813.5714286 | 0.184861 |
| ADRA2C | chr4 | 3764681 | 31 | 32 | 2794.707044 | 2992.985821 | 0.185081 |
| CDKN2B-AS1 | chr9 | 21970380 | 25 | 30 | 1101.872389 | 1509.324174 | 0.18512 |
| LDLR | chr19 | 11216959 | 3 | 7 | 300 | 667.5 | 0.185675 |
| PECAM1 | chr17 | 62399377 | 29 | 30 | 2702.064057 | 2666.485557 | 0.185764 |
| PLA2G7 | chr6 | 46671791 | 36 | 32 | 3023.030081 | 2780.42505 | 0.185782 |
| ARHGEF18 | chr19 | 7463748 | 18 | 18 | 511.1759225 | 627.7708174 | 0.185824 |
| BAX | chr19 | 49465364 | 33 | 33 | 2547.021745 | 2666.934797 | 0.185982 |
| APOA1 | chr11 | 116702971 | 35 | 32 | 2977.759117 | 2804.225422 | 0.186096 |
| COL15A1 | chr9 | 101704789 | 29 | 31 | 2738.789384 | 2813.98359 | 0.18618 |
| TMPO | chr12 | 98901096 | 34 | 32 | 3056.325033 | 2943.391279 | 0.186198 |
| APOA1 | chr11 | 116718292 | 35 | 32 | 3318.337568 | 3082.463958 | 0.1863 |
| APOA1 | chr11 | 116719477 | 32 | 32 | 3096.072258 | 3048.005989 | 0.186395 |
| TERT | chr5 | 1301235 | 33 | 32 | 2815.908082 | 2807.86093 | 0.186425 |
| LOXL1 | chr15 | 74218754 | 21 | 23 | 642.8846154 | 828.6156621 | 0.186523 |
| NOS3 | chr7 | 150706620 | 29 | 30 | 2672.074053 | 2614.511322 | 0.186642 |
| LDLR | chr19 | 11217881 | 28 | 29 | 2675.036573 | 2855.496032 | 0.186719 |
| NOS3 | chr7 | 150706830 | 23 | 26 | 2024.259825 | 2123.968254 | 0.186837 |
| CDKN2B-AS1 | chr9 | 21989775 | 10 | 10 | 283.9262372 | 352.8326936 | 0.186863 |
| TERT | chr5 | 1300126 | 33 | 33 | 2226.052394 | 2346.001326 | 0.187165 |
| ALOX15 | chr17 | 4549100 | 9 | 6 | 236.7388167 | 264.9350649 | 0.187196 |
| ADRA2C | chr4 | 3768788 | 18 | 13 | 748.235697 | 629.0628816 | 0.187278 |
| MIA3 | chr1 | 222792406 | 11 | 12 | 291.3492063 | 433.6646687 | 0.187321 |

| | | | | | | | |
|---|---|---|---|---|---|---|---|
| APOA1 | chr11 | 116701779 | 31 | 32 | 2922.422801 | 2925.937658 | 0.187445 |
| P2RY12 | chr3 | 151112522 | 34 | 32 | 3294.458238 | 3148.571711 | 0.187461 |
| APOA1 | chr11 | 116663548 | 32 | 32 | 3051.694289 | 3101.693164 | 0.187559 |
| LPA | chr6 | 11219144 | 30 | 24 | 1036.367041 | 926.9606777 | 0.187624 |
| ALDOA | chr16 | 30060404 | 28 | 32 | 2146.177028 | 2251.52506 | 0.188101 |
| LPA | chr6 | 160943047 | 36 | 32 | 3383.419108 | 3065.649402 | 0.188255 |
| APOA1 | chr11 | 116708330 | 32 | 32 | 3154.488283 | 3053.528444 | 0.188376 |
| GCK | chr7 | 44229320 | 7 | 9 | 415.2380952 | 698.0952381 | 0.188439 |
| GALNT2 | chr1 | 230191681 | 3 | 3 | 255 | 300 | 0.188499 |
| TMPO | chr12 | 98905177 | 16 | 18 | 1291.139971 | 1573.412698 | 0.188644 |
| PHACTR1 | chr6 | 12712125 | 35 | 33 | 2966.837415 | 2679.398723 | 0.188781 |
| NOS3 | chr7 | 150686803 | 32 | 32 | 1559.851965 | 1721.742941 | 0.1888 |
| APOA1 | chr11 | 116677183 | 33 | 31 | 3364.247977 | 3028.030744 | 0.188956 |
| LPA | chr6 | 11219966 | 13 | 19 | 1200.198413 | 1828.409091 | 0.188995 |
| ALOX15 | chr17 | 4552953 | 33 | 33 | 2430.346054 | 2319.8319 | 0.189204 |
| TIMP3 | chr22 | 33195943 | 37 | 32 | 3044.891692 | 2713.50737 | 0.189305 |
| PECAM1 | chr17 | 62399302 | 33 | 31 | 2711.526766 | 2450.20525 | 0.189439 |
| P2RY12 | chr3 | 151112519 | 34 | 32 | 3290.983391 | 3143.088245 | 0.189456 |
| LDLR | chr19 | 11222099 | 31 | 32 | 2864.518308 | 2822.09538 | 0.189495 |
| AKT2 | chr19 | 40794171 | 27 | 30 | 2162.059885 | 2553.210678 | 0.189537 |
| PLA2G7 | chr6 | 46711139 | 34 | 33 | 2734.409174 | 2512.781785 | 0.189742 |
| TIMP3 | chr22 | 33187198 | 34 | 32 | 3120.563233 | 2874.121129 | 0.190336 |
| LDLR | chr19 | 11193763 | 12 | 17 | 1096.865079 | 1635.714286 | 0.190622 |
| LDLR | chr19 | 11194373 | 29 | 30 | 2587.95954 | 2564.572372 | 0.190917 |
| LDLR | chr19 | 11221782 | 5 | 7 | 483.3333333 | 623.3333333 | 0.190937 |
| LPA | chr6 | 11221801 | 34 | 32 | 3178.726263 | 3088.117853 | 0.191216 |
| CTGF | chr6 | 132280889 | 34 | 32 | 3100.016333 | 2995.183494 | 0.191217 |
| TERT | chr5 | 1296051 | 30 | 30 | 2578.367758 | 2450.87149 | 0.191262 |
| FOXP3 | chrX | 49123657 | 16 | 21 | 1534.03505 | 1908.134921 | 0.191496 |
| ESR1 | chr6 | 152006159 | 27 | 30 | 2313.47374 | 2699.087057 | 0.191556 |
| APOA1 | chr11 | 116700143 | 23 | 18 | 1402.44228 | 1300.03663 | 0.191563 |
| ADRA2C | chr4 | 3765118 | 10 | 10 | 279.037629 | 346.9047619 | 0.191604 |
| APOA1 | chr11 | 116699808 | 25 | 25 | 1904.381174 | 1753.650794 | 0.191877 |
| AKT2 | chr19 | 40792016 | 32 | 31 | 2962.828422 | 2950.020786 | 0.192078 |
| COL15A1 | chr9 | 101701632 | 32 | 33 | 2059.623116 | 2001.950395 | 0.192124 |
| GALNT2 | chr1 | 230185645 | 36 | 32 | 3158.661019 | 2874.030531 | 0.192818 |
| ABCG1 | chr21 | 43633344 | 33 | 32 | 2580.688218 | 2645.667933 | 0.192923 |
| NOS3 | chr7 | 150694418 | 30 | 28 | 1266.032072 | 1326.837607 | 0.193016 |
| ARHGEF18 | chr19 | 7469836 | 34 | 32 | 3255.167869 | 2987.137095 | 0.193222 |
| E2F6 | chr2 | 11607910 | 32 | 32 | 2933.714211 | 2777.968812 | 0.193443 |
| APOA1 | chr11 | 116674825 | 24 | 25 | 938.8190318 | 1127.729901 | 0.193556 |
| ALOX15 | chr17 | 4545487 | 8 | 10 | 200.6330117 | 344.4980885 | 0.193819 |
| HMGCR | chr5 | 74624878 | 34 | 32 | 2897.653801 | 2801.750993 | 0.194157 |
| APOA1 | chr11 | 116677371 | 23 | 28 | 1695.162125 | 1734.344577 | 0.194262 |
| NOS3 | chr7 | 150692164 | 14 | 13 | 366.9181549 | 433.7218312 | 0.194521 |
| AKT2 | chr19 | 40795391 | 3 | 4 | 267.5 | 280 | 0.194525 |
| LOXL1 | chr15 | 74244249 | 7 | 9 | 418.7301587 | 446.1904762 | 0.194659 |

| | | | | | | | |
|---|---|---|---|---|---|---|---|
| ADRA2C | chr4 | 3766549 | 16 | 22 | 922.1464646 | 1435.15873 | 0.194761 |
| TERT | chr5 | 1301434 | 23 | 22 | 2077.201108 | 1889.087302 | 0.194801 |
| ATF3 | chr1 | 212743943 | 27 | 28 | 2362.546065 | 2562.45449 | 0.194897 |
| APOA1 | chr11 | 116661665 | 30 | 29 | 2617.420321 | 2383.013653 | 0.195125 |
| APOA1 | chr11 | 116661278 | 32 | 32 | 2995.939438 | 3061.567517 | 0.195284 |
| ABCB1 | chr7 | 87352134 | 39 | 32 | 3416.908151 | 2865.079487 | 0.195464 |
| APOA1 | chr11 | 116677369 | 29 | 30 | 2569.962121 | 2483.015873 | 0.195699 |
| LOXL1 | chr15 | 74215452 | 8 | 6 | 351.0714286 | 349.047619 | 0.195915 |
| F2RL3 | chr19 | 16999025 | 32 | 32 | 2843.166673 | 2932.60911 | 0.195961 |
| CDKN2B-AS1 | chr9 | 21992446 | 37 | 33 | 2596.680279 | 2209.537387 | 0.195985 |
| ABCA1 | chr9 | 107690733 | 7 | 4 | 224.6825397 | 88.57142857 | 0.196462 |
| NOS3 | chr7 | 150686449 | 31 | 33 | 2376.144494 | 2397.103572 | 0.196538 |
| FGF2 | chr4 | 123741986 | 23 | 29 | 2157.850679 | 2607.024379 | 0.196624 |
| E2F6 | chr2 | 11584713 | 30 | 28 | 2763.490953 | 2394.212404 | 0.19668 |
| ALOX15 | chr17 | 4545354 | 12 | 13 | 501.010101 | 667.6984127 | 0.196885 |
| PECAM1 | chr17 | 62399721 | 4 | 9 | 296.547619 | 843.3333333 | 0.197023 |
| CDKN2B-AS1 | chr9 | 21993463 | 38 | 33 | 1995.584961 | 1640.396805 | 0.19712 |
| BCL2 | chr18 | 60993684 | 38 | 32 | 3635.072202 | 3108.209775 | 0.19712 |
| PECAM1 | chr17 | 62399468 | 5 | 4 | 480 | 348.8888889 | 0.197223 |
| TMPO | chr12 | 98904932 | 30 | 28 | 3020.790043 | 2674.404762 | 0.197284 |
| ICAM2 | chr17 | 62080178 | 33 | 31 | 3238.527692 | 3077.516234 | 0.19742 |
| F2RL3 | chr19 | 17000096 | 13 | 10 | 347.4420024 | 344.5881896 | 0.197456 |
| APOA1 | chr11 | 116705969 | 5 | 2 | 235.8333333 | 57.14285714 | 0.197483 |
| LDLR | chr19 | 11224253 | 17 | 16 | 1609.722222 | 1600 | 0.197604 |
| FOXP3 | chrX | 49115543 | 19 | 19 | 1741.043956 | 1810.631313 | 0.197657 |
| ESR2 | chr14 | 64693661 | 32 | 32 | 2910.294025 | 2998.296688 | 0.197721 |
| NOS3 | chr7 | 150711113 | 7 | 8 | 175.7867495 | 306.3682845 | 0.197759 |
| APOA1 | chr11 | 116666249 | 33 | 32 | 3033.692626 | 2870.028388 | 0.197816 |
| COL15A1 | chr9 | 101832947 | 31 | 31 | 2841.042999 | 2958.975212 | 0.197823 |
| APOA1 | chr11 | 116712898 | 34 | 31 | 3311.280604 | 3060.941651 | 0.197839 |
| MCPH1 | chr8 | 6470385 | 33 | 32 | 3167.864665 | 2987.681677 | 0.197898 |
| ABCG1 | chr21 | 43717681 | 36 | 32 | 3228.794416 | 2931.882798 | 0.19804 |
| NOS3 | chr7 | 150708021 | 3 | 4 | 75 | 206.6666667 | 0.198041 |
| NOS3 | chr7 | 150691234 | 36 | 32 | 3118.179817 | 2704.551675 | 0.198101 |
| NOS3 | chr7 | 150690661 | 19 | 23 | 1721.017316 | 2167.142857 | 0.198184 |
| PHACTR1 | chr6 | 12717127 | 28 | 32 | 2604.441367 | 3054.061788 | 0.198203 |
| NOS3 | chr7 | 150680512 | 35 | 33 | 2381.926852 | 2127.083576 | 0.198278 |
| APOA1 | chr11 | 116693168 | 35 | 33 | 3114.97681 | 2823.461513 | 0.198419 |
| PLA2G7 | chr6 | 46712378 | 35 | 32 | 3246.614081 | 2899.59053 | 0.198462 |
| APOA1 | chr11 | 116664040 | 27 | 24 | 1697.160286 | 1726.895231 | 0.198718 |
| APOA1 | chr11 | 116661705 | 29 | 30 | 2516.5967 | 2712.81635 | 0.198936 |
| HMGCR | chr5 | 74623829 | 12 | 19 | 1140 | 1701.626984 | 0.199023 |
| APOA1 | chr11 | 116690115 | 37 | 33 | 3051.991851 | 2602.725756 | 0.199079 |
| LDLR | chr19 | 11192254 | 24 | 20 | 2297.936508 | 1842.063682 | 0.199257 |
| APOA1 | chr11 | 116711561 | 38 | 32 | 3571.691716 | 2959.25816 | 0.199323 |
| APOA1 | chr11 | 116703796 | 32 | 33 | 2985.979163 | 2959.68292 | 0.199443 |
| BAX | chr19 | 49449892 | 27 | 30 | 2413.203121 | 2799.44187 | 0.199659 |

| | | | | | | | |
|---|---|---|---|---|---|---|---|
| CTGF | chr6 | 132279048 | 36 | 33 | 3137.158571 | 2775.143164 | 0.20066 |
| HMGCR | chr5 | 74623767 | 26 | 26 | 2570.631313 | 2409.99889 | 0.20084 |
| F2RL3 | chr19 | 16992944 | 35 | 32 | 3262.355272 | 3035.090125 | 0.20092 |
| LPA | chr6 | 11216518 | 34 | 32 | 3287.095211 | 3058.260287 | 0.20096 |
| KLF2 | chr19 | 16432766 | 33 | 32 | 2900.913683 | 2887.023905 | 0.20108 |
| LDLR | chr19 | 11223983 | 25 | 23 | 2340.958486 | 2217.380952 | 0.2011 |
| APOA1 | chr11 | 116706109 | 11 | 13 | 312.518315 | 449.5199245 | 0.20282 |
| TERT | chr5 | 1298860 | 27 | 26 | 1611.003865 | 1734.235209 | 0.20298 |
| APOA1 | chr11 | 116684287 | 36 | 32 | 2757.749438 | 2548.990809 | 0.20308 |
| BCL2 | chr18 | 60989083 | 27 | 22 | 2084.285714 | 1565.456488 | 0.20312 |
| F7 | chr13 | 113750878 | 32 | 32 | 2936.133132 | 2800.015228 | 0.20372 |
| COL15A1 | chr9 | 101705340 | 31 | 32 | 3006.272941 | 3010.271567 | 0.20392 |
| ADRA2C | chr4 | 3759383 | 35 | 32 | 3186.513261 | 2976.790434 | 0.20398 |
| ADRA2C | chr4 | 3760463 | 37 | 32 | 3469.865912 | 3072.83662 | 0.20414 |
| ABCA1 | chr9 | 107691568 | 3 | 5 | 60 | 212.2222222 | 0.2042 |
| APOA1 | chr11 | 116708902 | 16 | 18 | 1518.333333 | 1641.984127 | 0.20496 |
| TERT | chr5 | 1299067 | 31 | 31 | 2833.99657 | 2710.443641 | 0.20582 |
| TMPO | chr12 | 98899530 | 38 | 32 | 3546.881344 | 2941.566191 | 0.20592 |
| TERT | chr5 | 1296196 | 32 | 32 | 2874.02087 | 2794.045936 | 0.206 |
| ADRA2C | chr4 | 3759717 | 32 | 33 | 2170.480966 | 2395.003933 | 0.20656 |
| CDKN2B-AS1 | chr9 | 21996049 | 13 | 15 | 377.6587302 | 543.1562882 | 0.2066 |
| ABCG1 | chr21 | 43635585 | 33 | 32 | 3040.954389 | 3029.516737 | 0.20668 |
| TIMP3 | chr22 | 33187739 | 36 | 32 | 3048.211392 | 2791.961077 | 0.207 |
| PHACTR1 | chr6 | 12715877 | 26 | 28 | 2452.512536 | 2555.133755 | 0.20712 |
| ABCG1 | chr21 | 43637224 | 27 | 27 | 2632.75264 | 2560.620542 | 0.2072 |
| ADRA2C | chr4 | 3761786 | 30 | 30 | 2115.634779 | 1881.666223 | 0.20722 |
| CDKN2B-AS1 | chr9 | 22006797 | 28 | 26 | 1366.76096 | 1124.618992 | 0.20736 |
| NOS3 | chr7 | 150695765 | 28 | 30 | 2585.93254 | 2677.56549 | 0.20762 |
| PECAM1 | chr17 | 62398846 | 21 | 22 | 1941.984127 | 2100.019523 | 0.20766 |
| APOA1 | chr11 | 116682012 | 36 | 33 | 2618.645156 | 2289.525479 | 0.20772 |
| FOXP3 | chrX | 49117265 | 5 | 5 | 500 | 470 | 0.208 |
| APOA1 | chr11 | 116704389 | 34 | 32 | 3393.356807 | 3006.430509 | 0.20808 |
| LOXL1 | chr15 | 74218167 | 32 | 32 | 2661.694991 | 2749.351332 | 0.20816 |
| CDKN2B-AS1 | chr9 | 21976144 | 17 | 10 | 489.5953443 | 379.7508701 | 0.2082 |
| TERT | chr5 | 1296261 | 35 | 32 | 3222.989921 | 3008.208333 | 0.20876 |
| LCK | chr1 | 32741675 | 30 | 31 | 2625.496618 | 2811.606501 | 0.20932 |
| LDLR | chr19 | 11225018 | 33 | 32 | 3123.158066 | 2959.761582 | 0.2096 |
| BCL2 | chr18 | 60989363 | 31 | 32 | 2415.87268 | 2372.352321 | 0.20968 |
| CDKN2B-AS1 | chr9 | 22007734 | 3 | 2 | 80.55555555 | 40 | 0.2101 |
| ADRA2C | chr4 | 3764783 | 26 | 28 | 1290.155591 | 1227.088934 | 0.2104 |
| PECAM1 | chr17 | 62409047 | 32 | 25 | 1308.924549 | 1156.605319 | 0.21056 |
| APOA1 | chr11 | 116713615 | 33 | 32 | 3002.217351 | 2837.52164 | 0.2106 |
| TERT | chr5 | 1295946 | 31 | 32 | 2247.411135 | 2471.421125 | 0.21076 |
| LOXL1 | chr15 | 74216038 | 33 | 32 | 2821.293783 | 2823.887161 | 0.21174 |
| ALOX15 | chr17 | 4545498 | 11 | 15 | 283.4976963 | 479.6498805 | 0.21188 |
| NOS3 | chr7 | 150710013 | 33 | 32 | 1938.14387 | 2034.823333 | 0.21194 |
| LCK | chr1 | 32751487 | 31 | 31 | 3121.858884 | 3032.079635 | 0.21198 |

| | | | | | | | |
|---|---|---|---|---|---|---|---|
| APOA1 | chr11 | 116697582 | 32 | 31 | 2922.715812 | 2896.917587 | 0.21216 |
| ICAM2 | chr17 | 62093550 | 12 | 18 | 343.0259361 | 627.0310245 | 0.21228 |
| APOA1 | chr11 | 116700304 | 13 | 12 | 372.3979529 | 434.0656566 | 0.21232 |
| CDKN2B-AS1 | chr9 | 21968394 | 9 | 6 | 227.1657754 | 236.9047619 | 0.21262 |
| APOA1 | chr11 | 116698999 | 14 | 20 | 1350.641026 | 1851.111111 | 0.21266 |
| LCK | chr1 | 32714509 | 33 | 33 | 1581.716249 | 1712.129939 | 0.2127 |
| NOS3 | chr7 | 150683004 | 8 | 8 | 409.3398268 | 499.1883117 | 0.2129 |
| LDLR | chr19 | 11191353 | 2 | 8 | 200 | 692.5 | 0.2129 |
| MIA3 | chr1 | 222792554 | 33 | 32 | 1905.447848 | 1723.063682 | 0.21304 |
| AKT2 | chr19 | 40736398 | 32 | 30 | 2219.551111 | 2227.568569 | 0.21346 |
| AKT2 | chr19 | 40800044 | 31 | 27 | 2977.555118 | 2528.002194 | 0.2137 |
| PECAM1 | chr17 | 62398407 | 32 | 32 | 2952.223251 | 2808.565887 | 0.2137 |
| TERT | chr5 | 1295732 | 4 | 6 | 106.3753133 | 141.6666667 | 0.21374 |
| CDKN2B-AS1 | chr9 | 22005129 | 3 | 3 | 76.36363636 | 173.3333333 | 0.21382 |
| F2RL3 | chr19 | 16997461 | 30 | 30 | 2697.057161 | 2595.203824 | 0.21392 |
| LDLR | chr19 | 11215952 | 32 | 32 | 3047.060234 | 2966.677786 | 0.21392 |
| P2RY12 | chr3 | 151109749 | 23 | 17 | 607.9965012 | 529.7386422 | 0.21394 |
| PLA2G7 | chr6 | 46705876 | 31 | 32 | 2870.065259 | 3030.640555 | 0.21422 |
| NOS3 | chr7 | 150706609 | 26 | 29 | 1819.628288 | 1866.834554 | 0.21426 |
| CDKN2B-AS1 | chr9 | 21988688 | 34 | 27 | 1124.226267 | 990.7880832 | 0.21446 |
| NOS3 | chr7 | 150705896 | 20 | 21 | 561.9078371 | 694.501522 | 0.21456 |
| F2RL3 | chr19 | 17000081 | 6 | 5 | 148.3333333 | 226.9047619 | 0.21478 |
| FOXP3 | chrX | 49126858 | 7 | 8 | 182.1428571 | 339.3650794 | 0.21498 |
| ALOX15 | chr17 | 4545788 | 7 | 6 | 173.8888889 | 244.3878928 | 0.2154 |
| APOA1 | chr11 | 116711627 | 37 | 32 | 3496.176483 | 3067.843683 | 0.21548 |
| TERT | chr5 | 1301494 | 32 | 32 | 2879.540246 | 2797.526298 | 0.2155 |
| APOA1 | chr11 | 116693200 | 35 | 32 | 3245.744778 | 3025.540395 | 0.21562 |
| TMPO | chr12 | 98907692 | 39 | 31 | 3146.877862 | 2280.836413 | 0.2157 |
| F7 | chr13 | 113753547 | 38 | 32 | 3409.691604 | 2930.689677 | 0.21576 |
| NOS3 | chr7 | 150687201 | 32 | 29 | 2371.471713 | 2023.801213 | 0.21586 |
| ARHGEF18 | chr19 | 7537863 | 16 | 24 | 1183.705739 | 1993.246753 | 0.2159 |
| CDKN1A | chr6 | 36653609 | 30 | 27 | 1274.139571 | 1014.817114 | 0.21602 |
| GCK | chr7 | 44235035 | 27 | 28 | 2610.811688 | 2754.79798 | 0.21614 |
| NOS3 | chr7 | 150711201 | 7 | 7 | 172.7533578 | 276.6417278 | 0.21618 |
| ADRA2C | chr4 | 3762423 | 31 | 30 | 2432.70279 | 2497.416243 | 0.21632 |
| F2RL3 | chr19 | 16990293 | 30 | 31 | 2844.630862 | 3016.008673 | 0.21634 |
| ARHGEF18 | chr19 | 7466124 | 27 | 30 | 2677.874307 | 2663.874459 | 0.21636 |
| TIMP3 | chr22 | 33190923 | 37 | 32 | 3070.233418 | 2734.218153 | 0.2164 |
| PHACTR1 | chr6 | 12717766 | 35 | 32 | 3245.860523 | 2920.442791 | 0.2164 |
| ABCG1 | chr21 | 43629740 | 38 | 33 | 3313.572086 | 2778.516491 | 0.21646 |
| TERT | chr5 | 1295772 | 28 | 26 | 1060.042162 | 1087.393902 | 0.2168 |
| ESR1 | chr6 | 152008441 | 38 | 32 | 3175.930621 | 2758.750353 | 0.21686 |
| NR3C1 | chr5 | 142789372 | 39 | 32 | 3656.615335 | 3045.083107 | 0.21692 |
| KLF2 | chr19 | 16433257 | 34 | 31 | 3283.81963 | 2942.858342 | 0.21702 |
| CDKN1A | chr6 | 36653771 | 27 | 28 | 1283.040346 | 1185.137967 | 0.2173 |
| AKT2 | chr19 | 40736390 | 32 | 30 | 2516.73768 | 2242.661365 | 0.21738 |
| APOA1 | chr11 | 116698964 | 27 | 26 | 2214.540419 | 1938.463203 | 0.2175 |

| | | | | | | | |
|---|---|---|---|---|---|---|---|
| ABCA1 | chr9 | 107690747 | 12 | 14 | 443.1349206 | 632.9545455 | 0.2175 |
| TERT | chr5 | 1295959 | 32 | 32 | 1965.552421 | 2155.671224 | 0.21762 |
| APOA1 | chr11 | 116706066 | 12 | 13 | 357.057387 | 488.975469 | 0.21762 |
| NR3C1 | chr5 | 142792699 | 34 | 33 | 2267.067143 | 2320.37491 | 0.2177 |
| ESR1 | chr6 | 152004338 | 31 | 32 | 2858.483831 | 2878.912714 | 0.21782 |
| NOS3 | chr7 | 150710384 | 18 | 19 | 545.0430125 | 686.586012 | 0.21802 |
| ADRA2C | chr4 | 3766213 | 34 | 32 | 3165.883133 | 3052.126596 | 0.21808 |
| APOA1 | chr11 | 116726115 | 32 | 32 | 3068.53057 | 3000.40314 | 0.2184 |
| TMPO | chr12 | 98906857 | 40 | 32 | 3376.677295 | 2639.674349 | 0.21842 |
| ABCG1 | chr21 | 43635126 | 33 | 31 | 2528.964914 | 2590.392279 | 0.21864 |
| F7 | chr13 | 113755501 | 14 | 12 | 1061.111111 | 1001.785714 | 0.21872 |
| ADRA2C | chr4 | 3765974 | 29 | 32 | 2674.676084 | 2842.188608 | 0.21894 |
| ICAM2 | chr17 | 62087378 | 37 | 33 | 2870.718749 | 2455.963627 | 0.21906 |
| PLA2G7 | chr6 | 46702940 | 3 | 2 | 73.33333333 | 66.66666666 | 0.2191 |
| CDKN2B-AS1 | chr9 | 21976513 | 3 | 2 | 137.1428571 | 114.2857143 | 0.2191 |
| AKT2 | chr19 | 40800217 | 32 | 31 | 3168.58443 | 3008.4347 | 0.21912 |
| KLF2 | chr19 | 16430699 | 14 | 13 | 902.5649351 | 724.1233766 | 0.2196 |
| F7 | chr13 | 113750710 | 27 | 22 | 2484.850289 | 2108.845599 | 0.21992 |
| CDKN2B-AS1 | chr9 | 21971257 | 11 | 12 | 313.251634 | 441.8864469 | 0.21994 |
| P2RY12 | chr3 | 151107304 | 31 | 32 | 1512.891329 | 1433.841934 | 0.22002 |
| BCL2 | chr18 | 60993279 | 33 | 33 | 2295.439132 | 2190.328283 | 0.2201 |
| ABCA1 | chr9 | 107693716 | 34 | 32 | 2809.424661 | 2742.027025 | 0.2201 |
| GALNT2 | chr1 | 230193999 | 3 | 6 | 243.3333333 | 563.3333333 | 0.22016 |
| NOS3 | chr7 | 150695457 | 21 | 21 | 869.8214286 | 1005.039683 | 0.22018 |
| F7 | chr13 | 113750497 | 23 | 19 | 2150.873016 | 1841.785714 | 0.22026 |
| FOXP3 | chrX | 49126932 | 15 | 6 | 678.2683983 | 222.3015873 | 0.2203 |
| APOA1 | chr11 | 116694931 | 32 | 32 | 3004.576622 | 3066.730271 | 0.2205 |
| GCK | chr7 | 44233049 | 34 | 32 | 3029.065451 | 2929.206958 | 0.22076 |
| LOXL1 | chr15 | 74215202 | 26 | 29 | 1064.207467 | 1312.70757 | 0.22086 |
| F7 | chr13 | 113750717 | 27 | 22 | 2349.847016 | 2024.52381 | 0.22086 |
| ADRA2C | chr4 | 3768582 | 11 | 11 | 304.6428572 | 400.4165856 | 0.22096 |
| TIMP3 | chr22 | 33190685 | 32 | 33 | 2712.089055 | 2676.827188 | 0.22116 |
| APOA1 | chr11 | 116705095 | 17 | 20 | 1664.95098 | 1909.722222 | 0.22116 |
| APOA1 | chr11 | 116673439 | 32 | 29 | 3126.280964 | 2761.121107 | 0.22158 |
| ADRA2C | chr4 | 3764068 | 32 | 32 | 3064.77849 | 3109.536561 | 0.22178 |
| LOXL1 | chr15 | 74212163 | 28 | 22 | 885.8788318 | 775.2909056 | 0.22194 |
| ICAM2 | chr17 | 62080115 | 32 | 32 | 3081.354401 | 3004.476973 | 0.22234 |
| CDKN2B-AS1 | chr9 | 21968436 | 9 | 8 | 236.4285714 | 310.9340659 | 0.22236 |
| CDKN2B-AS1 | chr9 | 22006043 | 11 | 11 | 271.9421102 | 364.4155844 | 0.22288 |
| LDLR | chr19 | 11221368 | 33 | 32 | 3193.889221 | 3033.172883 | 0.22312 |
| F2RL3 | chr19 | 16990943 | 31 | 32 | 2870.224918 | 2873.871973 | 0.22314 |
| LPA | chr6 | 11219394 | 34 | 32 | 2838.075549 | 2737.657285 | 0.22314 |
| PLA2G7 | chr6 | 46710006 | 33 | 32 | 3100.661405 | 2944.371921 | 0.22336 |
| APOA1 | chr11 | 116690529 | 31 | 32 | 2591.295456 | 2785.623761 | 0.22336 |
| PECAM1 | chr17 | 62399703 | 2 | 3 | 200 | 263.3333333 | 0.22342 |
| PECAM1 | chr17 | 62407351 | 7 | 4 | 566.6666667 | 387.5 | 0.22358 |
| APOA1 | chr11 | 116696472 | 35 | 32 | 2906.375701 | 2727.059909 | 0.22366 |

| | | | | | | | |
|---|---|---|---|---|---|---|---|
| ARHGEF18 | chr19 | 7467650 | 36 | 32 | 3449.631841 | 3022.237246 | 0.22366 |
| ICAM2 | chr17 | 62091274 | 30 | 29 | 2464.08811 | 2265.145688 | 0.22388 |
| LPA | chr6 | 11215948 | 32 | 32 | 3205.373737 | 3109.439436 | 0.22406 |
| CDKN2B-AS1 | chr9 | 21973993 | 7 | 13 | 222.2727273 | 346.984127 | 0.2241 |
| MCPH1 | chr8 | 6470120 | 22 | 25 | 2277.873377 | 2409.093407 | 0.22428 |
| TIMP3 | chr22 | 33186917 | 40 | 32 | 3835.79262 | 3038.726927 | 0.2248 |
| NOS3 | chr7 | 150694670 | 3 | 5 | 136.866491 | 304.5839071 | 0.22498 |
| CDKN2B-AS1 | chr9 | 21986404 | 36 | 32 | 3085.673969 | 2675.534301 | 0.2252 |
| KLF2 | chr19 | 16429071 | 25 | 22 | 2371.226551 | 2137.29798 | 0.22532 |
| LCK | chr1 | 32741633 | 31 | 32 | 2746.471653 | 2909.214124 | 0.22538 |
| APOA1 | chr11 | 116667425 | 37 | 33 | 2588.800317 | 2199.182197 | 0.22542 |
| ESR1 | chr6 | 152003453 | 29 | 31 | 2280.024356 | 2529.846322 | 0.22544 |
| ADRA2C | chr4 | 3766930 | 29 | 32 | 1180.624445 | 1431.06772 | 0.22566 |
| HMGCR | chr5 | 74631413 | 31 | 32 | 2852.615129 | 2775.971206 | 0.22578 |
| NOS3 | chr7 | 150695123 | 23 | 23 | 2358.220391 | 2167.380952 | 0.22578 |
| LCK | chr1 | 32707434 | 36 | 33 | 2125.055927 | 1816.53496 | 0.22594 |
| KLF2 | chr19 | 16426990 | 32 | 32 | 1445.407481 | 1312.520176 | 0.22596 |
| LDLR | chr19 | 11218111 | 28 | 22 | 2758.800187 | 2131.873682 | 0.226 |
| APOA1 | chr11 | 116711541 | 36 | 32 | 3146.731618 | 2869.223717 | 0.22604 |
| LDLR | chr19 | 11194822 | 31 | 31 | 1520.649095 | 1409.198342 | 0.22656 |
| NR3C1 | chr5 | 142788716 | 36 | 32 | 3390.679344 | 3080.890558 | 0.22662 |
| KLF2 | chr19 | 16426654 | 32 | 30 | 2897.600062 | 2796.06279 | 0.22664 |
| P2RY12 | chr3 | 151102861 | 35 | 32 | 1438.996033 | 1414.288617 | 0.22676 |
| ICAM2 | chr17 | 62087009 | 17 | 16 | 458.4725271 | 522.368118 | 0.22686 |
| AKT2 | chr19 | 40736565 | 30 | 33 | 2031.431773 | 2100.353304 | 0.2269 |
| LPA | chr6 | 11223960 | 31 | 32 | 2933.619447 | 3110.471224 | 0.22724 |
| F7 | chr13 | 113755385 | 32 | 32 | 3041.756599 | 2994.774258 | 0.22736 |
| ADRA2C | chr4 | 3761721 | 35 | 32 | 3317.448771 | 3067.471875 | 0.22742 |
| GCK | chr7 | 44233555 | 33 | 32 | 2969.555785 | 2775.454024 | 0.22776 |
| ABCG1 | chr21 | 43630546 | 35 | 32 | 3303.582348 | 3079.959063 | 0.22784 |
| CDKN1A | chr6 | 36643237 | 6 | 3 | 540.7142857 | 243.3333333 | 0.22794 |
| CDKN1A | chr6 | 36653034 | 35 | 32 | 3090.150846 | 2872.421435 | 0.22804 |
| F7 | chr13 | 113750829 | 32 | 31 | 3283.465383 | 3006.209207 | 0.22814 |
| F7 | chr13 | 113750753 | 28 | 26 | 2615.469351 | 2503.617216 | 0.22816 |
| GCK | chr7 | 44237131 | 36 | 32 | 3102.685063 | 2699.992955 | 0.22832 |
| LDLR | chr19 | 11218157 | 33 | 31 | 3266.967488 | 3012.351067 | 0.22834 |
| PHACTR1 | chr6 | 12713642 | 32 | 31 | 2153.239669 | 2200.956389 | 0.22838 |
| FGF2 | chr4 | 123744608 | 11 | 12 | 969.6356421 | 1145.714286 | 0.22896 |
| APOA1 | chr11 | 116696335 | 35 | 32 | 2617.794228 | 2470.895705 | 0.22906 |
| KLF2 | chr19 | 16426726 | 29 | 28 | 2573.985486 | 2348.758858 | 0.22922 |
| ADRA2C | chr4 | 3764062 | 32 | 32 | 3102.475905 | 3063.340426 | 0.2294 |
| P2RY12 | chr3 | 151111851 | 37 | 30 | 1256.825655 | 1100.916319 | 0.22952 |
| APOA1 | chr11 | 116680121 | 25 | 26 | 2514.480519 | 2530.138889 | 0.22962 |
| APOA1 | chr11 | 116666744 | 34 | 32 | 3079.135011 | 2969.624476 | 0.2298 |
| CDKN2B-AS1 | chr9 | 21970698 | 19 | 14 | 609.5870439 | 526.0251997 | 0.2302 |
| CDKN1A | chr6 | 36651352 | 28 | 32 | 2532.674858 | 2990.411533 | 0.23032 |
| F7 | chr13 | 113750905 | 32 | 32 | 3154.727505 | 3012.102725 | 0.23048 |

| | | | | | | | |
|---|---|---|---|---|---|---|---|
| LOXL1 | chr15 | 74218819 | 24 | 21 | 815.3471674 | 816.3529101 | 0.23054 |
| F7 | chr13 | 113753314 | 28 | 30 | 2085.952304 | 1985.378737 | 0.231 |
| NOS3 | chr7 | 150691009 | 29 | 29 | 2795.21073 | 2661.544165 | 0.23108 |
| APOA1 | chr11 | 116676695 | 33 | 33 | 1893.079873 | 1999.157932 | 0.23138 |
| APOA1 | chr11 | 116691554 | 32 | 33 | 1986.090793 | 2170.32667 | 0.23148 |
| MCPH1 | chr8 | 6258223 | 32 | 32 | 3057.23163 | 3010.294442 | 0.2317 |
| F2RL3 | chr19 | 16991879 | 31 | 31 | 2907.738095 | 2981.172402 | 0.23172 |
| MIA3 | chr1 | 222815392 | 27 | 31 | 2386.260186 | 2873.332223 | 0.23186 |
| NR3C1 | chr5 | 142788677 | 34 | 32 | 3188.342344 | 3065.150458 | 0.232 |
| ALDOA | chr16 | 30062325 | 36 | 32 | 3322.235287 | 3011.455509 | 0.23224 |
| ARHGEF18 | chr19 | 7537423 | 32 | 33 | 2336.063798 | 2294.134794 | 0.2326 |
| GCK | chr7 | 44230381 | 37 | 32 | 3270.313954 | 2718.921772 | 0.23266 |
| PLA2G7 | chr6 | 46705292 | 4 | 5 | 354.7619048 | 487.5 | 0.23274 |
| APOA1 | chr11 | 116670072 | 37 | 32 | 2928.84091 | 2601.814927 | 0.23298 |
| F7 | chr13 | 113750421 | 19 | 20 | 1446.771284 | 1662.19475 | 0.23308 |
| NOS3 | chr7 | 150708023 | 19 | 22 | 667.6487401 | 896.5295815 | 0.23318 |
| TMPO | chr12 | 98907533 | 37 | 33 | 3080.697974 | 2647.138077 | 0.23334 |
| LDLR | chr19 | 11196305 | 31 | 32 | 2718.084429 | 2709.068652 | 0.23338 |
| ABCA1 | chr9 | 107692081 | 16 | 16 | 751.7103811 | 895.6934405 | 0.23356 |
| PHACTR1 | chr6 | 12711079 | 27 | 30 | 2010.392989 | 2068.254728 | 0.23362 |
| F7 | chr13 | 113751336 | 15 | 14 | 1465.714286 | 1310 | 0.23384 |
| F7 | chr13 | 113752698 | 33 | 32 | 3021.806338 | 3012.542018 | 0.23398 |
| TIMP3 | chr22 | 33193746 | 34 | 32 | 3144.56102 | 2878.834615 | 0.2343 |
| TERT | chr5 | 1253509 | 29 | 31 | 2566.569282 | 2833.424215 | 0.23448 |
| ADRA2C | chr4 | 3766838 | 19 | 20 | 1568.293651 | 1772.664835 | 0.2348 |
| APOA1 | chr11 | 116677973 | 31 | 30 | 2826.801994 | 2638.195951 | 0.23502 |
| LDLR | chr19 | 11221395 | 34 | 32 | 3257.39383 | 3009.098215 | 0.2354 |
| NOS3 | chr7 | 150686970 | 13 | 14 | 1197.532468 | 1350.833333 | 0.2354 |
| CTGF | chr6 | 132277463 | 27 | 25 | 1528.133348 | 1218.099764 | 0.23586 |
| TIMP3 | chr22 | 33186886 | 36 | 32 | 3498.130716 | 3084.149162 | 0.23608 |
| LCK | chr1 | 32740774 | 21 | 18 | 1795.497199 | 1632.936508 | 0.23638 |
| FOXP3 | chrX | 49128948 | 32 | 33 | 1673.14069 | 1845.424939 | 0.23654 |
| ABCA1 | chr9 | 107698876 | 37 | 32 | 3210.822604 | 2837.561366 | 0.23658 |
| NOS3 | chr7 | 150694045 | 7 | 12 | 220 | 512.1428571 | 0.23718 |
| PECAM1 | chr17 | 62401132 | 36 | 32 | 3299.451301 | 2989.708919 | 0.23734 |
| FOXP3 | chrX | 49127860 | 2 | 2 | 40 | 140 | 0.2375 |
| APOA1 | chr11 | 116719710 | 9 | 12 | 740.7539683 | 870.5555555 | 0.23764 |
| MCPH1 | chr8 | 6470315 | 15 | 20 | 1249.318182 | 1789.50938 | 0.2378 |
| LDLR | chr19 | 11193162 | 29 | 32 | 2803.857128 | 3035.651154 | 0.23784 |
| CDKN2B-AS1 | chr9 | 22006078 | 8 | 7 | 207.745098 | 266.6483516 | 0.23802 |
| ABCG1 | chr21 | 43717560 | 36 | 33 | 2659.116403 | 2548.843145 | 0.23818 |
| ARHGEF18 | chr19 | 7461982 | 14 | 5 | 1024.688645 | 426.6666667 | 0.23856 |
| PECAM1 | chr17 | 62398368 | 32 | 32 | 3008.665633 | 2903.425101 | 0.23862 |
| BAX | chr19 | 49449337 | 13 | 13 | 948.3333333 | 1054.642857 | 0.239 |
| NOS3 | chr7 | 150710389 | 14 | 14 | 398.6180486 | 491.9138868 | 0.23912 |
| COL15A1 | chr9 | 101701497 | 36 | 32 | 3395.568968 | 2961.179912 | 0.23928 |
| APOA1 | chr11 | 116716843 | 35 | 32 | 3355.384346 | 3107.284677 | 0.23928 |

| | | | | | | | |
|---|---|---|---|---|---|---|---|
| NOS3 | chr7 | 150694727 | 34 | 33 | 2998.272524 | 2805.62922 | 0.23966 |
| CDKN2B-AS1 | chr9 | 21971051 | 5 | 3 | 107.2222222 | 60 | 0.23976 |
| TERT | chr5 | 1295970 | 30 | 33 | 2425.210623 | 2495.72532 | 0.24002 |
| NOS3 | chr7 | 150711090 | 10 | 9 | 258.5776942 | 321.1915205 | 0.24034 |
| FOXP3 | chrX | 49116550 | 25 | 25 | 2413.194502 | 2298.217159 | 0.24038 |
| LDLR | chr19 | 11198187 | 32 | 33 | 2065.837754 | 1952.841391 | 0.24066 |
| CDKN2B-AS1 | chr9 | 21993448 | 38 | 31 | 1350.366825 | 1182.757859 | 0.241 |
| F7 | chr13 | 113752996 | 33 | 32 | 3194.344959 | 3048.760296 | 0.24118 |
| BCL2 | chr18 | 60991127 | 12 | 12 | 1083.690476 | 1143.333333 | 0.2414 |
| APOA1 | chr11 | 116677175 | 33 | 31 | 3343.534353 | 3003.43312 | 0.24142 |
| ADRA2C | chr4 | 3766681 | 22 | 26 | 1263.540904 | 1666.800977 | 0.2415 |
| TIMP3 | chr22 | 33187578 | 32 | 32 | 3046.851762 | 2971.182962 | 0.2416 |
| BCL2 | chr18 | 60994574 | 38 | 32 | 3606.840172 | 2986.517213 | 0.24204 |
| AKT2 | chr19 | 40800960 | 23 | 24 | 2128.838384 | 2312.380952 | 0.24216 |
| NOS3 | chr7 | 150710411 | 26 | 27 | 969.0572026 | 1133.305975 | 0.24226 |
| LOXL1 | chr15 | 74212404 | 32 | 32 | 1339.452919 | 1467.811808 | 0.2428 |
| TERT | chr5 | 1296377 | 34 | 32 | 3080.038137 | 2841.233004 | 0.243 |
| NOS3 | chr7 | 150683537 | 35 | 33 | 1531.904637 | 1585.637453 | 0.24316 |
| CDKN2B-AS1 | chr9 | 22006223 | 4 | 6 | 92.18570141 | 228.3310841 | 0.24326 |
| CDKN2B-AS1 | chr9 | 21985169 | 38 | 32 | 3116.720894 | 2714.813747 | 0.24336 |
| FGF2 | chr4 | 123742013 | 27 | 31 | 2644.200154 | 2971.653722 | 0.2434 |
| LDLR | chr19 | 11216182 | 20 | 23 | 1940.865385 | 2169.401154 | 0.2434 |
| NOS3 | chr7 | 150680462 | 35 | 33 | 2189.184029 | 2158.080758 | 0.24342 |
| NOS3 | chr7 | 150705982 | 28 | 29 | 996.8082221 | 1153.900141 | 0.2435 |
| APOA1 | chr11 | 116662835 | 32 | 32 | 2966.480384 | 3043.56311 | 0.24356 |
| GALNT2 | chr1 | 230418210 | 22 | 25 | 674.2116029 | 677.2342448 | 0.24358 |
| ADRA2C | chr4 | 3768723 | 29 | 31 | 1455.084154 | 1404.605823 | 0.2437 |
| ESR1 | chr6 | 152009540 | 35 | 32 | 3190.723235 | 2976.408526 | 0.24376 |
| MCPH1 | chr8 | 6470591 | 32 | 32 | 3072.049542 | 2984.092839 | 0.24378 |
| LPA | chr6 | 11224876 | 16 | 17 | 844.469697 | 1018.037518 | 0.24382 |
| FOXP3 | chrX | 49118077 | 30 | 22 | 2921.742424 | 2094.603175 | 0.24384 |
| MCPH1 | chr8 | 6470025 | 14 | 9 | 1342.727273 | 822.2619048 | 0.24438 |
| CTGF | chr6 | 132280293 | 35 | 32 | 3334.61544 | 2990.071491 | 0.24468 |
| FOXP3 | chrX | 49126992 | 20 | 14 | 1256.497114 | 775.2380952 | 0.24468 |
| KLF2 | chr19 | 16430161 | 9 | 4 | 860 | 273.3333333 | 0.2447 |
| ADRA2C | chr4 | 3766875 | 29 | 27 | 2448.512599 | 2387.270507 | 0.2448 |
| ARHGEF18 | chr19 | 7466086 | 30 | 32 | 2960.80359 | 3033.118015 | 0.24498 |
| ADRA2C | chr4 | 3766860 | 26 | 22 | 2012.472089 | 1556.666667 | 0.24502 |
| LDLR | chr19 | 11219824 | 21 | 21 | 1927.97619 | 1811.219336 | 0.24524 |
| APOA1 | chr11 | 116692402 | 34 | 32 | 3201.354858 | 2962.092263 | 0.24552 |
| LDLR | chr19 | 11220596 | 3 | 4 | 300 | 345.7142857 | 0.24574 |
| MCPH1 | chr8 | 6261377 | 29 | 28 | 2563.697969 | 2571.39186 | 0.24578 |
| DHFR | chr5 | 79951646 | 33 | 33 | 2805.343937 | 2702.714025 | 0.24588 |
| ESR2 | chr14 | 64807581 | 2 | 7 | 83.91812865 | 419.0632284 | 0.24604 |
| CDKN2B-AS1 | chr9 | 21964621 | 25 | 26 | 856.687843 | 979.5573445 | 0.24658 |
| PHACTR1 | chr6 | 13276009 | 32 | 33 | 2238.178051 | 2206.372998 | 0.24684 |
| LPA | chr6 | 11221338 | 29 | 31 | 2667.871573 | 2936.435648 | 0.24694 |

| | | | | | | | |
|---|---|---|---|---|---|---|---|
| APOA1 | chr11 | 116702685 | 32 | 31 | 2770.767984 | 2763.070608 | 0.24722 |
| ADRA2C | chr4 | 3761731 | 35 | 32 | 3315.046766 | 3073.132402 | 0.24732 |
| KLF2 | chr19 | 16437844 | 20 | 25 | 815.4105861 | 1139.461143 | 0.24734 |
| LCK | chr1 | 32751472 | 31 | 30 | 2999.434223 | 2803.313187 | 0.24742 |
| FOXP3 | chrX | 49129835 | 30 | 25 | 1012.8614 | 936.5979552 | 0.2476 |
| TERT | chr5 | 1295905 | 31 | 32 | 1902.703735 | 1779.5836 | 0.24792 |
| FGF2 | chr4 | 123745631 | 38 | 32 | 3489.416594 | 2983.531933 | 0.24806 |
| ADRA2C | chr4 | 3767186 | 7 | 7 | 171.244825 | 240.1960784 | 0.24818 |
| LOXL1 | chr15 | 74218880 | 34 | 32 | 1545.928689 | 1326.635218 | 0.2483 |
| APOA1 | chr11 | 116692294 | 33 | 32 | 3205.368404 | 3062.558481 | 0.2483 |
| BCL2 | chr18 | 60994122 | 16 | 18 | 1365.032468 | 1656.190476 | 0.24842 |
| APOA1 | chr11 | 116687837 | 32 | 32 | 2905.25035 | 2976.772396 | 0.2486 |
| APOA1 | chr11 | 116690790 | 30 | 32 | 2666.947608 | 2940.330837 | 0.2487 |
| NOS3 | chr7 | 150710802 | 6 | 3 | 162.5793651 | 106.0714286 | 0.24874 |
| CDKN1A | chr6 | 36652128 | 26 | 28 | 2375.544733 | 2623.848096 | 0.24878 |
| APOA1 | chr11 | 116673838 | 34 | 33 | 2707.084794 | 2716.172378 | 0.24888 |
| LPA | chr6 | 160952668 | 37 | 33 | 2548.592788 | 2364.244552 | 0.2491 |
| NOS3 | chr7 | 150694356 | 6 | 6 | 142.7777778 | 231.1721612 | 0.24922 |
| LDLR | chr19 | 11193484 | 33 | 32 | 3217.684934 | 2986.875275 | 0.24924 |
| ABCA1 | chr9 | 107698642 | 36 | 32 | 3309.67284 | 2990.285964 | 0.2494 |
| CDKN1A | chr6 | 36651475 | 20 | 24 | 1789.756267 | 2275.039683 | 0.2499 |
| F7 | chr13 | 113750316 | 30 | 28 | 2662.149288 | 2567.647353 | 0.24996 |
| LPA | chr6 | 160944423 | 35 | 33 | 2367.726275 | 2334.493349 | 0.2503 |
| NOS3 | chr7 | 150705969 | 7 | 7 | 174.1133377 | 262.1979896 | 0.25036 |
| LOXL1 | chr15 | 74215350 | 3 | 2 | 121.9047619 | 45 | 0.25044 |
| LDLR | chr19 | 11216101 | 28 | 30 | 2618.908694 | 2879.348589 | 0.25088 |
| LDLR | chr19 | 11218078 | 12 | 18 | 1183.333333 | 1718.885281 | 0.25104 |
| APOA1 | chr11 | 116691343 | 34 | 32 | 3072.903345 | 2778.658583 | 0.25134 |
| F7 | chr13 | 113751121 | 4 | 6 | 346.6666667 | 571.4285714 | 0.25184 |
| LPA | chr6 | 11222486 | 40 | 33 | 2627.56061 | 2246.398556 | 0.25196 |
| NOS3 | chr7 | 150692410 | 32 | 31 | 1669.06195 | 1465.172826 | 0.25198 |
| TERT | chr5 | 1298429 | 6 | 7 | 402.8571429 | 570 | 0.25202 |
| ABCA1 | chr9 | 107696435 | 38 | 32 | 3596.754586 | 2926.933199 | 0.25214 |
| APOA1 | chr11 | 116703589 | 36 | 32 | 3160.641071 | 2871.793524 | 0.25222 |
| LCK | chr1 | 32741138 | 14 | 19 | 473.4665335 | 545.1721448 | 0.25222 |
| TMPO | chr12 | 98903274 | 4 | 5 | 320 | 500 | 0.25222 |
| LCK | chr1 | 32740457 | 28 | 32 | 2699.923822 | 3004.696137 | 0.2523 |
| APOA1 | chr11 | 116716840 | 35 | 32 | 3308.813708 | 3067.06698 | 0.25232 |
| LDLR | chr19 | 11216663 | 9 | 10 | 830.5555556 | 980 | 0.25232 |
| F2RL3 | chr19 | 16993519 | 33 | 32 | 3119.207048 | 3073.409424 | 0.25246 |
| ABCG1 | chr21 | 43633092 | 15 | 18 | 1452.103175 | 1780 | 0.25248 |
| KLF2 | chr19 | 16429196 | 7 | 3 | 355.2044514 | 198 | 0.2526 |
| LOXL1 | chr15 | 74219137 | 3 | 2 | 90.83333333 | 133.3333333 | 0.25268 |
| APOA1 | chr11 | 116689610 | 12 | 14 | 1178.888889 | 1330.31746 | 0.25272 |
| NOS3 | chr7 | 150690079 | 8 | 7 | 280 | 340.5769231 | 0.253 |
| LDLR | chr19 | 11216242 | 11 | 10 | 974.047619 | 938.9285714 | 0.25306 |
| LDLR | chr19 | 11190756 | 36 | 32 | 3419.731529 | 3071.748654 | 0.2532 |

| | | | | | | | |
|---|---|---|---|---|---|---|---|
| F7 | chr13 | 113750809 | 31 | 30 | 3052.948718 | 2876.660839 | 0.25324 |
| F2RL3 | chr19 | 16998131 | 15 | 14 | 762.6190476 | 588.0952381 | 0.25338 |
| ESR2 | chr14 | 64807671 | 32 | 32 | 3081.762569 | 3127.541015 | 0.25382 |
| P2RY12 | chr3 | 151054410 | 39 | 32 | 3583.301151 | 2974.212262 | 0.25384 |
| BAX | chr19 | 49449290 | 24 | 24 | 1898.448773 | 2024.163059 | 0.2539 |
| ABCA1 | chr9 | 107694835 | 9 | 3 | 855 | 260 | 0.254 |
| F7 | chr13 | 113756963 | 23 | 27 | 1538.749826 | 1589.077564 | 0.2541 |
| FOXP3 | chrX | 49118324 | 31 | 32 | 2874.423012 | 3041.025472 | 0.25448 |
| PHACTR1 | chr6 | 12717887 | 38 | 32 | 3260.182197 | 2802.640481 | 0.25468 |
| ARHGEF18 | chr19 | 7460675 | 16 | 12 | 414.5939746 | 400.5984215 | 0.25472 |
| CDKN2B-AS1 | chr9 | 21969374 | 28 | 29 | 1033.856592 | 1213.142689 | 0.25484 |
| PHACTR1 | chr6 | 12713951 | 32 | 30 | 2597.424553 | 2540.990398 | 0.25494 |
| ALOX15 | chr17 | 4547614 | 34 | 32 | 3209.372381 | 3075.268917 | 0.25508 |
| CDKN2B-AS1 | chr9 | 21981801 | 9 | 7 | 324.1269841 | 199.047619 | 0.25508 |
| FOXP3 | chrX | 49127194 | 19 | 24 | 1821.355311 | 2237.102065 | 0.25554 |
| CDKN2B-AS1 | chr9 | 21971142 | 26 | 23 | 811.5496078 | 802.0755827 | 0.2556 |
| TERT | chr5 | 1300406 | 32 | 29 | 2815.163989 | 2452.954644 | 0.25624 |
| PLA2G7 | chr6 | 46713412 | 39 | 32 | 3252.854748 | 2732.954571 | 0.25626 |
| APOA1 | chr11 | 116719454 | 33 | 32 | 3228.142773 | 3092.880761 | 0.25646 |
| CDKN2B-AS1 | chr9 | 21964673 | 30 | 30 | 1613.748168 | 1758.654422 | 0.25656 |
| APOA1 | chr11 | 116708600 | 26 | 26 | 2580.992063 | 2472.910335 | 0.25656 |
| F7 | chr13 | 113752605 | 33 | 32 | 2570.888833 | 2351.37025 | 0.25668 |
| CDKN1A | chr6 | 36650733 | 20 | 10 | 533.5438334 | 347.0771428 | 0.25674 |
| APOA1 | chr11 | 116679131 | 39 | 32 | 3714.407112 | 3085.000725 | 0.25678 |
| F7 | chr13 | 113750867 | 32 | 32 | 3241.765138 | 3043.689742 | 0.2568 |
| KLF2 | chr19 | 16429024 | 11 | 10 | 928.8888889 | 934.3253968 | 0.25686 |
| CDKN2B-AS1 | chr9 | 21996151 | 14 | 9 | 435.2272727 | 335.0793651 | 0.25712 |
| APOA1 | chr11 | 116661238 | 31 | 32 | 2870.420127 | 3034.328084 | 0.25738 |
| ABCG1 | chr21 | 43635747 | 34 | 32 | 2998.948008 | 2754.676062 | 0.25756 |
| F2RL3 | chr19 | 16991405 | 30 | 32 | 2777.441218 | 3055.614159 | 0.2576 |
| COL15A1 | chr9 | 101696919 | 2 | 3 | 130 | 300 | 0.25776 |
| NOS3 | chr7 | 150708901 | 5 | 5 | 472.8571429 | 500 | 0.25786 |
| GALNT2 | chr1 | 230188609 | 7 | 4 | 546.7063492 | 363.3333333 | 0.25836 |
| CDKN2B-AS1 | chr9 | 22009870 | 34 | 33 | 2797.284442 | 2577.767323 | 0.25878 |
| PLA2G7 | chr6 | 46704078 | 37 | 32 | 3129.770958 | 2768.287428 | 0.25884 |
| APOA1 | chr11 | 116667086 | 4 | 3 | 371.4285714 | 245.7142857 | 0.25884 |
| CDKN2B-AS1 | chr9 | 21968519 | 2 | 2 | 67.85714286 | 40 | 0.25912 |
| MIA3 | chr1 | 222792372 | 7 | 6 | 164.1025641 | 222.4603175 | 0.25924 |
| HMGCR | chr5 | 74628766 | 5 | 10 | 443.5606061 | 798.7698413 | 0.25928 |
| NOS3 | chr7 | 150683948 | 34 | 33 | 1919.500285 | 1983.161137 | 0.25946 |
| LPA | chr6 | 11221097 | 39 | 32 | 3353.004542 | 2683.397203 | 0.2599 |
| NOS3 | chr7 | 150683561 | 36 | 32 | 2909.577556 | 2667.528287 | 0.25994 |
| PLA2G7 | chr6 | 46703367 | 4 | 3 | 85.55801707 | 167.9761905 | 0.25994 |
| TMPO | chr12 | 98904940 | 30 | 30 | 3019.645355 | 2893.857961 | 0.26014 |
| ADRA2C | chr4 | 3760258 | 37 | 32 | 3444.502102 | 3024.999888 | 0.26034 |
| ABCG1 | chr21 | 43632274 | 9 | 16 | 869.047619 | 1442.380952 | 0.2605 |
| FGF2 | chr4 | 123741421 | 35 | 33 | 2305.729419 | 2087.28364 | 0.26072 |

| | | | | | | | |
|---|---|---|---|---|---|---|---|
| APOA1 | chr11 | 116707064 | 27 | 31 | 917.4597382 | 1150.346761 | 0.26082 |
| MCPH1 | chr8 | 6470967 | 5 | 8 | 444.047619 | 590 | 0.26106 |
| FOXP3 | chrX | 49121301 | 16 | 16 | 907.8342491 | 789.4144452 | 0.26128 |
| BCL2 | chr18 | 60994771 | 36 | 32 | 3310.457531 | 2983.973528 | 0.26146 |
| NOS3 | chr7 | 150711262 | 5 | 2 | 143.3333333 | 42.22222222 | 0.26152 |
| APOA1 | chr11 | 116709102 | 32 | 30 | 2641.908131 | 2594.63253 | 0.26154 |
| APOA1 | chr11 | 116669798 | 17 | 18 | 529.4727203 | 669.6364746 | 0.26166 |
| APOA1 | chr11 | 116687479 | 14 | 12 | 1194.145299 | 1076.706349 | 0.2617 |
| APOA1 | chr11 | 116675264 | 18 | 22 | 1667.609711 | 2121.901154 | 0.26196 |
| NOS3 | chr7 | 150683276 | 7 | 6 | 159.8232323 | 230.8441558 | 0.26252 |
| KLF2 | chr19 | 16426962 | 31 | 32 | 1426.339475 | 1346.98572 | 0.263 |
| APOA1 | chr11 | 116690073 | 33 | 32 | 2678.76391 | 2521.641602 | 0.26316 |
| F7 | chr13 | 113752352 | 36 | 32 | 3272.735667 | 2976.997392 | 0.26324 |
| LDLR | chr19 | 11224354 | 33 | 32 | 3186.775144 | 3140.775746 | 0.26338 |
| ADRA2C | chr4 | 3759830 | 33 | 33 | 2697.032265 | 2580.418075 | 0.26342 |
| LDLR | chr19 | 11220906 | 9 | 10 | 776.6666667 | 938.0952381 | 0.26346 |
| GCK | chr7 | 44230634 | 36 | 32 | 3227.81792 | 2915.572809 | 0.26374 |
| DHFR | chr5 | 79957798 | 34 | 32 | 3192.834587 | 3069.223154 | 0.26376 |
| ALOX15 | chr17 | 4545075 | 9 | 12 | 328.2936508 | 359.1774892 | 0.26382 |
| APOA1 | chr11 | 116704369 | 34 | 32 | 3414.037161 | 2988.566206 | 0.26406 |
| LDLR | chr19 | 11192725 | 27 | 31 | 2451.13101 | 2899.837662 | 0.26442 |
| AMOTL2 | chr3 | 134101472 | 34 | 32 | 2760.93192 | 2694.766948 | 0.26448 |
| ADRA2C | chr4 | 3763828 | 32 | 33 | 2935.494057 | 2909.770736 | 0.26452 |
| F2RL3 | chr19 | 17000202 | 10 | 12 | 579.8412698 | 536.6666667 | 0.26454 |
| CDKN2B-AS1 | chr9 | 21981768 | 21 | 19 | 993.5859973 | 784.1378066 | 0.26478 |
| PECAM1 | chr17 | 62397454 | 33 | 33 | 2758.89668 | 2659.353712 | 0.26498 |
| NOS3 | chr7 | 150690518 | 33 | 32 | 3057.6622 | 2901.994632 | 0.265 |
| BAX | chr19 | 49450955 | 32 | 32 | 3008.779081 | 3050.457831 | 0.26506 |
| CDKN2B-AS1 | chr9 | 21984651 | 35 | 33 | 1995.702472 | 1970.757308 | 0.26512 |
| E2F6 | chr2 | 11584826 | 39 | 32 | 3757.560372 | 3105.855243 | 0.26516 |
| LPA | chr6 | 11224595 | 16 | 15 | 1243.362193 | 1262.261905 | 0.2655 |
| TMPO | chr12 | 98907857 | 33 | 25 | 1072.553121 | 903.440402 | 0.2657 |
| ARHGEF18 | chr19 | 7461399 | 11 | 7 | 1051.428571 | 638.0952381 | 0.26596 |
| ABCA1 | chr9 | 107690995 | 24 | 26 | 1586.618485 | 1539.297078 | 0.26602 |
| ALOX15 | chr17 | 4552039 | 24 | 23 | 769.4606617 | 848.7897963 | 0.26606 |
| PHACTR1 | chr6 | 13276086 | 18 | 16 | 1517.399267 | 1415.137085 | 0.26606 |
| APOA1 | chr11 | 116661135 | 34 | 32 | 2799.063804 | 2734.978926 | 0.26608 |
| NOS3 | chr7 | 150693971 | 6 | 10 | 159.1269841 | 354.3650794 | 0.26608 |
| F2RL3 | chr19 | 16995267 | 36 | 32 | 3318.667654 | 3012.675112 | 0.26614 |
| ARHGEF18 | chr19 | 7461120 | 7 | 14 | 637.5 | 1335 | 0.26644 |
| FOXP3 | chrX | 49116032 | 28 | 29 | 2603.718606 | 2631.529877 | 0.26656 |
| GCK | chr7 | 44232233 | 34 | 31 | 2883.417224 | 2705.047985 | 0.26664 |
| LOXL1 | chr15 | 74218733 | 8 | 9 | 219.71091 | 288.1150794 | 0.26666 |
| GALNT2 | chr1 | 230190735 | 38 | 32 | 3442.097491 | 2844.953886 | 0.26682 |
| ADRA2C | chr4 | 3759259 | 34 | 32 | 3186.881629 | 2938.811101 | 0.26684 |
| NOS3 | chr7 | 150694617 | 32 | 31 | 2531.223537 | 2335.917628 | 0.26684 |
| BCL2 | chr18 | 60990644 | 29 | 29 | 2841.76655 | 2789.497637 | 0.2671 |

| | | | | | | | |
|---|---|---|---|---|---|---|---|
| APOA1 | chr11 | 116687523 | 27 | 27 | 2331.494561 | 2449.976412 | 0.26718 |
| TERT | chr5 | 1253705 | 32 | 32 | 2878.466271 | 2977.202361 | 0.26722 |
| LDLR | chr19 | 11224418 | 35 | 32 | 3322.890657 | 3076.396078 | 0.2673 |
| F7 | chr13 | 113751839 | 32 | 33 | 2222.928256 | 2408.583747 | 0.26742 |
| ALOX15 | chr17 | 4549166 | 35 | 32 | 1493.820905 | 1262.531055 | 0.2675 |
| APOA1 | chr11 | 116671866 | 7 | 2 | 528.5714286 | 200 | 0.26766 |
| APOA1 | chr11 | 116720328 | 35 | 33 | 1522.300339 | 1326.764254 | 0.26768 |
| PECAM1 | chr17 | 62407189 | 8 | 5 | 638.0952381 | 460 | 0.26792 |
| F2RL3 | chr19 | 16990277 | 30 | 30 | 2866.242369 | 2917.041586 | 0.26794 |
| LCK | chr1 | 32741037 | 25 | 31 | 975.3232017 | 1098.729594 | 0.26804 |
| APOA1 | chr11 | 116716288 | 32 | 31 | 2967.927984 | 2949.164645 | 0.2683 |
| AKT2 | chr19 | 40796476 | 32 | 32 | 3164.468864 | 3118.793064 | 0.26836 |
| CDKN2B-AS1 | chr9 | 21970739 | 12 | 10 | 336.0382782 | 360.672884 | 0.26886 |
| COL15A1 | chr9 | 101832613 | 31 | 30 | 2969.026252 | 2928.815629 | 0.26898 |
| TERT | chr5 | 1299061 | 31 | 31 | 3012.620811 | 2918.465356 | 0.26898 |
| ESR2 | chr14 | 64812762 | 34 | 32 | 3268.905001 | 3024.297481 | 0.26916 |
| AKT2 | chr19 | 40799984 | 5 | 5 | 391.4285714 | 440 | 0.26916 |
| NOS3 | chr7 | 150706321 | 31 | 31 | 2876.093059 | 2927.330057 | 0.26934 |
| F7 | chr13 | 113758639 | 8 | 11 | 395.2380952 | 644.2857143 | 0.26942 |
| DHFR | chr5 | 79951587 | 33 | 33 | 2442.22329 | 2333.933839 | 0.26974 |
| APOA1 | chr11 | 116716923 | 30 | 32 | 2821.603348 | 2941.289191 | 0.26976 |
| APOA1 | chr11 | 116725027 | 33 | 33 | 2024.434128 | 1873.900482 | 0.26978 |
| E2F6 | chr2 | 11608275 | 32 | 32 | 2965.24491 | 3030.377578 | 0.26978 |
| ALOX15 | chr17 | 4546394 | 23 | 25 | 1280.891053 | 1534.368132 | 0.26994 |
| LPA | chr6 | 11220764 | 37 | 32 | 3500.755071 | 3088.532828 | 0.27 |
| GALNT2 | chr1 | 230193746 | 31 | 33 | 2166.975063 | 2171.444564 | 0.27016 |
| LDLR | chr19 | 11222334 | 16 | 20 | 1536.388889 | 1967.5 | 0.27038 |
| LPA | chr6 | 160944154 | 12 | 14 | 1058.095238 | 1297.5 | 0.27042 |
| APOA1 | chr11 | 116674499 | 5 | 4 | 115.5555556 | 116.4102564 | 0.2705 |
| TERT | chr5 | 1301422 | 25 | 27 | 2389.972395 | 2473.404373 | 0.2708 |
| TIMP3 | chr22 | 33189229 | 33 | 31 | 2890.622778 | 2778.244465 | 0.27094 |
| LPA | chr6 | 11219939 | 7 | 11 | 660 | 974.6428571 | 0.27106 |
| APOA1 | chr11 | 116701119 | 17 | 18 | 1623.504274 | 1654.206349 | 0.27108 |
| CDKN2B-AS1 | chr9 | 22000838 | 18 | 19 | 1216.906427 | 1178.376623 | 0.2711 |
| E2F6 | chr2 | 11584512 | 31 | 31 | 2908.907302 | 2980.111416 | 0.27124 |
| PHACTR1 | chr6 | 12710305 | 39 | 32 | 3351.323872 | 2796.243369 | 0.27148 |
| ABCA1 | chr9 | 107691557 | 5 | 5 | 108.0769231 | 195.9478022 | 0.2715 |
| TERT | chr5 | 1300697 | 31 | 32 | 2980.194912 | 3024.690302 | 0.27162 |
| LDLR | chr19 | 11218366 | 8 | 7 | 742.5 | 555 | 0.27164 |
| APOA1 | chr11 | 116682606 | 26 | 29 | 2575.892857 | 2748.368298 | 0.27168 |
| ADRA2C | chr4 | 3764802 | 10 | 8 | 266.4656368 | 291.9027756 | 0.27182 |
| MIA3 | chr1 | 222808185 | 34 | 32 | 3217.123901 | 3067.073792 | 0.272 |
| ICAM2 | chr17 | 62084323 | 33 | 31 | 2376.408633 | 2096.027809 | 0.27212 |
| KLF2 | chr19 | 16437855 | 28 | 28 | 1289.053505 | 1434.14474 | 0.27216 |
| APOA1 | chr11 | 116668942 | 31 | 30 | 2954.614469 | 2666.80065 | 0.2722 |
| FOXP3 | chrX | 49106732 | 3 | 2 | 240 | 200 | 0.27222 |
| PHACTR1 | chr6 | 13276066 | 27 | 28 | 2494.044012 | 2449.322169 | 0.27224 |

| | | | | | | | |
|---|---|---|---|---|---|---|---|
| APOA1 | chr11 | 116719861 | 34 | 32 | 3234.005053 | 2996.898441 | 0.2726 |
| APOA1 | chr11 | 116661080 | 34 | 32 | 3250.753122 | 3111.404413 | 0.27262 |
| CDKN2B-AS1 | chr9 | 21991219 | 31 | 33 | 1606.864769 | 1583.021394 | 0.2727 |
| CDKN2B-AS1 | chr9 | 21968462 | 15 | 16 | 455.6962482 | 586.7316017 | 0.27272 |
| ARHGEF18 | chr19 | 7461125 | 7 | 14 | 614.5938375 | 1335 | 0.27282 |
| COL15A1 | chr9 | 101702926 | 32 | 32 | 1463.207094 | 1375.174134 | 0.27284 |
| CDKN2B-AS1 | chr9 | 21965942 | 17 | 11 | 514.6578421 | 385.3361181 | 0.27296 |
| TERT | chr5 | 1253744 | 26 | 29 | 2251.126589 | 2676.699134 | 0.27298 |
| CDKN1A | chr6 | 36645088 | 18 | 12 | 514.5795594 | 410.8230329 | 0.27298 |
| ATF3 | chr1 | 212742181 | 29 | 25 | 2883.850202 | 2374.632035 | 0.27324 |
| BCL2 | chr18 | 60989038 | 5 | 8 | 370.4761905 | 515.2380952 | 0.2735 |
| HMGCR | chr5 | 74624983 | 37 | 32 | 3147.239308 | 2666.154143 | 0.27356 |
| HMGCR | chr5 | 74631844 | 34 | 32 | 3278.452108 | 3024.730085 | 0.27374 |
| F2RL3 | chr19 | 16991418 | 32 | 32 | 3040.992108 | 3109.624506 | 0.27378 |
| APOA1 | chr11 | 116707641 | 31 | 32 | 2809.657041 | 2981.597579 | 0.27418 |
| PLA2G7 | chr6 | 46707600 | 35 | 33 | 3128.878986 | 2859.194548 | 0.27428 |
| LPA | chr6 | 11223649 | 32 | 32 | 3037.692772 | 3089.047918 | 0.27442 |
| BAX | chr19 | 49449503 | 31 | 30 | 3058.987777 | 2872.919164 | 0.2745 |
| APOA1 | chr11 | 116722779 | 33 | 24 | 1255.567553 | 832.4892674 | 0.27464 |
| DHFR | chr5 | 79956308 | 31 | 32 | 2972.946766 | 2946.286569 | 0.27466 |
| F7 | chr13 | 113750643 | 26 | 23 | 2402.81606 | 2051.944444 | 0.27506 |
| APOA1 | chr11 | 116705509 | 26 | 25 | 2147.138156 | 2177.651515 | 0.27508 |
| FOXP3 | chrX | 49130901 | 33 | 31 | 3298.196836 | 2933.524347 | 0.27538 |
| ABCG1 | chr21 | 43633674 | 32 | 32 | 2795.621603 | 2884.253252 | 0.27542 |
| ARHGEF18 | chr19 | 7461385 | 13 | 11 | 1108.095238 | 851.5873016 | 0.27556 |
| F2RL3 | chr19 | 16993398 | 36 | 32 | 3283.721104 | 2970.365232 | 0.27566 |
| KLF2 | chr19 | 16427582 | 36 | 32 | 1629.26619 | 1344.4803 | 0.27588 |
| APOA1 | chr11 | 116710923 | 41 | 32 | 3826.89501 | 3027.094163 | 0.27604 |
| CTGF | chr6 | 132276131 | 33 | 32 | 2957.649077 | 2940.943423 | 0.27608 |
| AKT2 | chr19 | 40800706 | 38 | 32 | 3709.92302 | 3083.986258 | 0.27642 |
| BAX | chr19 | 49454688 | 25 | 26 | 2045.634921 | 1981.818933 | 0.27642 |
| TERT | chr5 | 1294704 | 3 | 5 | 60 | 127.8571429 | 0.2768 |
| CDKN2B-AS1 | chr9 | 22002541 | 4 | 4 | 85 | 123.3333333 | 0.27684 |
| FOXP3 | chrX | 49121206 | 8 | 6 | 604.4444444 | 373.3333333 | 0.277 |
| APOA1 | chr11 | 116702479 | 33 | 32 | 3025.758192 | 2991.123451 | 0.27724 |
| BCL2 | chr18 | 60991131 | 12 | 13 | 1193.333333 | 1263.333333 | 0.27734 |
| P2RY12 | chr3 | 151107261 | 2 | 2 | 50.79365079 | 140 | 0.27736 |
| NOS3 | chr7 | 150682020 | 32 | 32 | 2319.831831 | 2198.70232 | 0.27738 |
| KLF2 | chr19 | 16429473 | 33 | 32 | 3041.619417 | 2858.603147 | 0.27754 |
| CDKN2B-AS1 | chr9 | 21967690 | 10 | 12 | 234.9358213 | 354.4473918 | 0.278 |
| HMGCR | chr5 | 74627469 | 38 | 33 | 1948.320539 | 1770.950454 | 0.27822 |
| HMGCR | chr5 | 74625382 | 33 | 33 | 2677.244941 | 2487.231355 | 0.27846 |
| ABCB1 | chr7 | 87342195 | 21 | 22 | 657.4182381 | 792.8787933 | 0.27852 |
| LDLR | chr19 | 11195859 | 5 | 7 | 450 | 560 | 0.27862 |
| F7 | chr13 | 113750732 | 26 | 27 | 2353.558109 | 2530.945166 | 0.27876 |
| LDLR | chr19 | 11218336 | 14 | 17 | 1282.434428 | 1467.34127 | 0.27888 |
| LDLR | chr19 | 11193137 | 28 | 30 | 2703.908559 | 2838.488456 | 0.2789 |

| | | | | | | | |
|---|---|---|---|---|---|---|---|
| ALDOA | chr16 | 30062228 | 32 | 32 | 2953.657042 | 3011.54061 | 0.27892 |
| DHFR | chr5 | 79953192 | 7 | 16 | 668.8888889 | 1440.47619 | 0.27908 |
| ESR1 | chr6 | 152007546 | 30 | 32 | 2876.488979 | 2966.952077 | 0.27916 |
| LOXL1 | chr15 | 74219214 | 15 | 11 | 414.5923317 | 389.4830024 | 0.2793 |
| MIA3 | chr1 | 222815009 | 32 | 32 | 2619.904875 | 2736.072443 | 0.27972 |
| CDKN2B-AS1 | chr9 | 21993638 | 17 | 19 | 708.2851067 | 690.6364469 | 0.27982 |
| F7 | chr13 | 113751729 | 34 | 32 | 3200.242708 | 3054.024217 | 0.28006 |
| MCPH1 | chr8 | 6470020 | 15 | 10 | 1490 | 965.7142857 | 0.28012 |
| MCPH1 | chr8 | 6261188 | 32 | 32 | 3236.647611 | 3039.418244 | 0.28018 |
| APOA1 | chr11 | 116717289 | 9 | 9 | 880 | 838.3333333 | 0.2803 |
| F7 | chr13 | 113775040 | 29 | 30 | 2618.875291 | 2783.871685 | 0.2804 |
| LCK | chr1 | 32739956 | 28 | 33 | 2062.816657 | 2292.456996 | 0.28044 |
| LDLR | chr19 | 11224005 | 25 | 26 | 2369.68254 | 2525.833333 | 0.28052 |
| ABCA1 | chr9 | 107691559 | 2 | 3 | 48.57142857 | 177.5 | 0.2806 |
| AMOTL2 | chr3 | 134100779 | 36 | 32 | 3017.575216 | 2738.23668 | 0.2808 |
| MIA3 | chr1 | 222841311 | 37 | 32 | 3396.348322 | 2893.60986 | 0.28088 |
| APOA1 | chr11 | 116698409 | 21 | 22 | 2012.380952 | 2154.166667 | 0.28096 |
| LDLR | chr19 | 11191098 | 33 | 32 | 3027.884469 | 2842.063286 | 0.28102 |
| PECAM1 | chr17 | 62407170 | 2 | 2 | 120 | 183.3333333 | 0.28136 |
| APOA1 | chr11 | 116660918 | 35 | 33 | 2636.949152 | 2389.234182 | 0.28172 |
| CDKN1A | chr6 | 36652327 | 32 | 32 | 3059.206006 | 3112.204377 | 0.2818 |
| GCK | chr7 | 44230142 | 36 | 32 | 3027.403894 | 2742.136048 | 0.28234 |
| F2RL3 | chr19 | 16994991 | 9 | 8 | 677.1428571 | 683.3333333 | 0.28242 |
| NOS3 | chr7 | 150708070 | 33 | 33 | 1824.551506 | 1940.748167 | 0.28278 |
| APOA1 | chr11 | 116680822 | 26 | 29 | 2392.276359 | 2753.463481 | 0.28292 |
| APOA1 | chr11 | 116704328 | 34 | 32 | 3468.602022 | 3021.310673 | 0.28302 |
| LOXL1 | chr15 | 74215620 | 23 | 31 | 2043.935008 | 2665.296093 | 0.28304 |
| APOA1 | chr11 | 116668363 | 16 | 13 | 1335.436508 | 1168.214286 | 0.2831 |
| ADRA2C | chr4 | 3768476 | 3 | 4 | 74.76190476 | 198.3333333 | 0.28322 |
| HMGCR | chr5 | 74625665 | 35 | 33 | 3148.667155 | 2876.513045 | 0.2837 |
| ATF3 | chr1 | 212783192 | 9 | 10 | 201.093158 | 307.9630967 | 0.2837 |
| AKT2 | chr19 | 40736559 | 30 | 32 | 2692.299453 | 2800.925537 | 0.28382 |
| E2F6 | chr2 | 11607081 | 34 | 33 | 2077.139451 | 2120.022343 | 0.28408 |
| LDLR | chr19 | 11221179 | 16 | 19 | 1165.528404 | 1176.552892 | 0.28432 |
| CDKN2B-AS1 | chr9 | 22010440 | 39 | 33 | 1900.329843 | 1500.746032 | 0.2849 |
| ESR2 | chr14 | 64809644 | 7 | 7 | 593.3333333 | 646.6666667 | 0.28492 |
| F2RL3 | chr19 | 17000054 | 7 | 6 | 192.947187 | 239.1414141 | 0.28498 |
| TERT | chr5 | 1297646 | 32 | 31 | 2800.668486 | 2636.188516 | 0.28506 |
| APOA1 | chr11 | 116668786 | 31 | 31 | 2650.457804 | 2731.060917 | 0.28512 |
| APOA1 | chr11 | 116683754 | 36 | 32 | 3358.823271 | 3026.77094 | 0.2852 |
| CTGF | chr6 | 132272363 | 3 | 7 | 70 | 260.2380952 | 0.28526 |
| ADRA2C | chr4 | 3758626 | 34 | 33 | 2797.823995 | 2618.722799 | 0.28542 |
| LDLR | chr19 | 11197599 | 8 | 4 | 713.3333333 | 387.5 | 0.2856 |
| PLA2G7 | chr6 | 46672008 | 34 | 33 | 2287.112261 | 2329.811654 | 0.28574 |
| APOA1 | chr11 | 116666147 | 8 | 2 | 626.3095238 | 180 | 0.28574 |
| ADRA2C | chr4 | 3770125 | 31 | 33 | 1939.139435 | 1901.083073 | 0.28578 |
| ALOX15 | chr17 | 4545566 | 35 | 33 | 2212.569467 | 1964.794371 | 0.28592 |

| | | | | | | | |
|---|---|---|---|---|---|---|---|
| F7 | chr13 | 113759210 | 6 | 8 | 352.8571429 | 567.1428571 | 0.28596 |
| ARHGEF18 | chr19 | 7537382 | 32 | 32 | 2072.106369 | 1930.951395 | 0.28634 |
| CDKN2B-AS1 | chr9 | 21996167 | 6 | 13 | 159.2857143 | 453.7262737 | 0.28658 |
| APOA1 | chr11 | 116690012 | 33 | 32 | 2753.930363 | 2596.384763 | 0.2866 |
| PLA2G7 | chr6 | 46706208 | 3 | 2 | 60 | 128.5714286 | 0.2869 |
| CTGF | chr6 | 132281785 | 31 | 31 | 1155.166409 | 1235.625509 | 0.28694 |
| ALDOA | chr16 | 30057820 | 33 | 32 | 2999.44149 | 2856.246952 | 0.28722 |
| BAX | chr19 | 49453816 | 25 | 28 | 2239.016539 | 2598.877614 | 0.2874 |
| CDKN2BAS | chr9 | 22067538 | 33 | 32 | 3037.273702 | 2891.637957 | 0.28782 |
| LPA | chr6 | 160947065 | 38 | 32 | 3613.89202 | 3008.395086 | 0.28808 |
| ABCG1 | chr21 | 43633998 | 26 | 27 | 2479.587269 | 2651.153291 | 0.28864 |
| TERT | chr5 | 1298090 | 34 | 32 | 2817.539378 | 2581.245689 | 0.2888 |
| E2F6 | chr2 | 11614266 | 36 | 32 | 3402.40521 | 2906.807766 | 0.2891 |
| COL15A1 | chr9 | 101702634 | 37 | 33 | 1941.957852 | 1823.232822 | 0.28946 |
| FOXP3 | chrX | 49117338 | 6 | 9 | 569.047619 | 793.9285714 | 0.28964 |
| NOS3 | chr7 | 150692429 | 32 | 33 | 1539.818862 | 1708.507121 | 0.28974 |
| F2RL3 | chr19 | 16994099 | 32 | 32 | 3040.83264 | 3087.20975 | 0.28974 |
| APOA1 | chr11 | 116682548 | 14 | 13 | 1248.571429 | 1209.761905 | 0.28974 |
| KLF2 | chr19 | 16429536 | 18 | 23 | 1629.69475 | 1842.218615 | 0.28976 |
| LDLR | chr19 | 11196505 | 14 | 12 | 1355.090377 | 1129.722222 | 0.28982 |
| APOA1 | chr11 | 116697164 | 30 | 29 | 2807.216811 | 2635.008721 | 0.28994 |
| ATF3 | chr1 | 212738979 | 35 | 32 | 3321.019842 | 3080.642802 | 0.28994 |
| LCK | chr1 | 32741619 | 31 | 32 | 2927.055542 | 2961.78708 | 0.29024 |
| ADRA2C | chr4 | 3762182 | 33 | 33 | 2205.266294 | 2085.094826 | 0.29036 |
| APOA1 | chr11 | 116682562 | 21 | 19 | 1879.95671 | 1618.834776 | 0.29052 |
| TERT | chr5 | 1296743 | 30 | 32 | 2759.529221 | 2856.857019 | 0.29054 |
| KLF2 | chr19 | 16437898 | 24 | 27 | 777.7652281 | 965.6730769 | 0.29054 |
| COL15A1 | chr9 | 101699959 | 26 | 27 | 2320.80416 | 2311.560939 | 0.29056 |
| TMPO | chr12 | 98903378 | 34 | 32 | 3314.377083 | 3149.239874 | 0.29058 |
| COL15A1 | chr9 | 101701040 | 38 | 33 | 3011.960143 | 2516.3581 | 0.29072 |
| APOA1 | chr11 | 116694356 | 26 | 27 | 2202.58547 | 2387.60101 | 0.29112 |
| APOA1 | chr11 | 116668308 | 6 | 2 | 548.8888889 | 200 | 0.29134 |
| CDKN2B-AS1 | chr9 | 21967298 | 3 | 5 | 63.86308909 | 192.3082707 | 0.29142 |
| BAX | chr19 | 49453515 | 26 | 23 | 1690.757576 | 1342.74975 | 0.292 |
| ALOX15 | chr17 | 4545082 | 2 | 3 | 40 | 88.57142857 | 0.29258 |
| FGF2 | chr4 | 123747620 | 23 | 26 | 1538.730436 | 1872.810523 | 0.2928 |
| ALDOA | chr16 | 30060710 | 33 | 32 | 2871.442572 | 2672.96869 | 0.29284 |
| COL15A1 | chr9 | 101697315 | 33 | 33 | 2053.890838 | 1957.834337 | 0.2929 |
| F7 | chr13 | 113753135 | 34 | 32 | 2999.441293 | 2874.819289 | 0.293 |
| COL15A1 | chr9 | 101705792 | 33 | 32 | 3031.763206 | 2876.2795 | 0.29304 |
| MIA3 | chr1 | 222815555 | 38 | 32 | 3668.439666 | 3040.450966 | 0.29316 |
| ADRA2C | chr4 | 3767213 | 10 | 12 | 246.7965368 | 382.1104626 | 0.29316 |
| FOXP3 | chrX | 49122071 | 29 | 32 | 2696.114275 | 2887.133644 | 0.29336 |
| LOXL1 | chr15 | 74211628 | 9 | 9 | 769.047619 | 830.8333333 | 0.29372 |
| ICAM2 | chr17 | 62093907 | 23 | 20 | 629.5867147 | 617.8139983 | 0.29392 |
| PLA2G7 | chr6 | 46709613 | 23 | 26 | 1767.475989 | 2148.259518 | 0.29408 |
| LPA | chr6 | 160943975 | 29 | 28 | 2571.472139 | 2571.738817 | 0.29434 |

| | | | | | | | |
|---|---|---|---|---|---|---|---|
| APOA1 | chr11 | 116720139 | 27 | 28 | 2391.41166 | 2396.966089 | 0.29436 |
| BAX | chr19 | 49449801 | 32 | 31 | 2702.368218 | 2482.269019 | 0.2944 |
| APOA1 | chr11 | 116672970 | 15 | 23 | 1406.563853 | 2223.055556 | 0.29494 |
| NOS3 | chr7 | 150689475 | 7 | 5 | 539.0476191 | 430 | 0.29532 |
| KLF2 | chr19 | 16437998 | 32 | 32 | 1789.352684 | 1919.977562 | 0.29536 |
| CDKN2B-AS1 | chr9 | 21969079 | 8 | 5 | 173.9996511 | 195.2614836 | 0.29544 |
| LPA | chr6 | 11219784 | 34 | 32 | 3278.952289 | 3123.462144 | 0.29548 |
| PHACTR1 | chr6 | 12717016 | 34 | 32 | 2763.78439 | 2671.09976 | 0.29562 |
| TERT | chr5 | 1300807 | 12 | 11 | 1129.047619 | 989.047619 | 0.29562 |
| CDKN1A | chr6 | 36650741 | 22 | 14 | 686.7406591 | 521.286872 | 0.29568 |
| PLA2G7 | chr6 | 46705751 | 8 | 7 | 481.3018926 | 509.2857143 | 0.2961 |
| DHFR | chr5 | 79952986 | 31 | 31 | 2935.504049 | 3021.51076 | 0.29614 |
| MCPH1 | chr8 | 6470790 | 33 | 31 | 3088.701266 | 2831.281424 | 0.29634 |
| APOA1 | chr11 | 116678655 | 9 | 13 | 812.3809524 | 1089.047619 | 0.29638 |
| ADRA2C | chr4 | 3759016 | 37 | 33 | 2561.690875 | 2184.403046 | 0.29644 |
| PECAM1 | chr17 | 62410230 | 2 | 2 | 40 | 133.3333333 | 0.29648 |
| ADRA2C | chr4 | 3762676 | 36 | 32 | 3283.684829 | 2974.294728 | 0.2965 |
| APOA1 | chr11 | 116694115 | 23 | 27 | 1927.605221 | 2139.82351 | 0.29712 |
| F2RL3 | chr19 | 17002951 | 28 | 29 | 2476.647274 | 2638.195569 | 0.29716 |
| ABCA1 | chr9 | 107693937 | 30 | 31 | 3033.520725 | 3066.666667 | 0.2973 |
| F7 | chr13 | 113750521 | 20 | 21 | 1812.22472 | 1977.380952 | 0.29734 |
| BAX | chr19 | 49465530 | 5 | 5 | 113.5714286 | 192.1978022 | 0.2976 |
| ABCA1 | chr9 | 107695574 | 29 | 31 | 2832.226447 | 2958.710317 | 0.29778 |
| TIMP3 | chr22 | 33193439 | 38 | 33 | 2493.406952 | 2086.56952 | 0.29836 |
| CDKN1A | chr6 | 36651840 | 34 | 32 | 3259.112819 | 3026.471238 | 0.29842 |
| CDKN2B-AS1 | chr9 | 21996933 | 31 | 30 | 1507.135977 | 1356.507212 | 0.29852 |
| CDKN2B-AS1 | chr9 | 21968504 | 4 | 8 | 83.65079365 | 184.1269841 | 0.29856 |
| APOA1 | chr11 | 116701503 | 35 | 31 | 3291.542488 | 2839.727832 | 0.29892 |
| AKT2 | chr19 | 40736295 | 32 | 33 | 2075.388473 | 2049.433191 | 0.29898 |
| APOA1 | chr11 | 116699615 | 35 | 32 | 3304.519215 | 3078.230419 | 0.29922 |
| KLF2 | chr19 | 16429542 | 17 | 21 | 1591.064214 | 1815.909091 | 0.29922 |
| LDLR | chr19 | 11193353 | 33 | 32 | 2999.990094 | 2841.12921 | 0.29948 |
| APOA1 | chr11 | 116703799 | 32 | 32 | 3079.218108 | 3039.915774 | 0.29992 |
| ADRA2C | chr4 | 3759616 | 32 | 32 | 2937.162362 | 2862.521223 | 0.30026 |
| ESR1 | chr6 | 152002766 | 5 | 4 | 108.0952381 | 175.9803922 | 0.3006 |
| APOA1 | chr11 | 116678988 | 34 | 32 | 2949.72525 | 2834.17474 | 0.30078 |
| F2RL3 | chr19 | 16991052 | 4 | 4 | 380 | 326.6666667 | 0.30108 |
| CDKN2B-AS1 | chr9 | 21989725 | 27 | 24 | 943.6942665 | 758.0531852 | 0.30118 |
| ABCB1 | chr7 | 87345579 | 33 | 32 | 3002.716631 | 2960.853189 | 0.30126 |
| E2F6 | chr2 | 11611439 | 18 | 20 | 1182.397002 | 1139.379311 | 0.3014 |
| CDKN2B-AS1 | chr9 | 21971058 | 3 | 3 | 62.22222222 | 135.5555556 | 0.30166 |
| CDKN2B-AS1 | chr9 | 21976176 | 14 | 15 | 525.3571429 | 481.8642469 | 0.30234 |
| ADRA2C | chr4 | 3768794 | 14 | 16 | 691.3512949 | 684.6894772 | 0.3025 |
| TERT | chr5 | 1296055 | 30 | 30 | 2378.935801 | 2252.364417 | 0.30272 |
| APOA1 | chr11 | 116699015 | 13 | 15 | 1185.075758 | 1276.904762 | 0.30272 |
| ABCG1 | chr21 | 43632208 | 20 | 25 | 1729.84127 | 2247.621545 | 0.30286 |
| CDKN2B-AS1 | chr9 | 21971037 | 4 | 5 | 85 | 174.7619048 | 0.3031 |

| | | | | | | | |
|---|---|---|---|---|---|---|---|
| APOA1 | chr11 | 116716975 | 33 | 32 | 2913.195815 | 2897.781965 | 0.30312 |
| GCK | chr7 | 44229230 | 32 | 32 | 2865.690947 | 2935.647328 | 0.3032 |
| CDKN2B-AS1 | chr9 | 21970371 | 14 | 19 | 464.3113275 | 726.7825312 | 0.30322 |
| LOXL1 | chr15 | 74216882 | 28 | 31 | 974.6913479 | 1154.03654 | 0.30342 |
| ESR1 | chr6 | 152005590 | 36 | 32 | 3416.625481 | 3067.745972 | 0.30386 |
| CDKN2B-AS1 | chr9 | 22005117 | 3 | 2 | 90.83333333 | 128.5714286 | 0.30388 |
| TIMP3 | chr22 | 33191063 | 34 | 32 | 2483.191784 | 2420.590018 | 0.30398 |
| APOA1 | chr11 | 116712769 | 24 | 20 | 2116.201299 | 1854.764402 | 0.30438 |
| PLA2G7 | chr6 | 46703505 | 19 | 21 | 508.896479 | 625.7044055 | 0.30442 |
| F2RL3 | chr19 | 16991024 | 15 | 11 | 1410.357143 | 1071.428571 | 0.30446 |
| PECAM1 | chr17 | 62398217 | 38 | 33 | 2987.344753 | 2517.022538 | 0.3045 |
| LDLR | chr19 | 11190884 | 34 | 32 | 2911.851653 | 2811.084899 | 0.30458 |
| CDKN2B-AS1 | chr9 | 21996431 | 38 | 33 | 2119.168711 | 1761.564424 | 0.30478 |
| CDKN2B-AS1 | chr9 | 21997025 | 36 | 33 | 1602.069404 | 1555.07107 | 0.3049 |
| NOS3 | chr7 | 150710401 | 22 | 23 | 775.21691 | 904.740807 | 0.30524 |
| AKT2 | chr19 | 40794859 | 35 | 32 | 3216.377416 | 2899.908019 | 0.30574 |
| LOXL1 | chr15 | 74218529 | 32 | 32 | 2906.277067 | 2985.796245 | 0.30584 |
| APOA1 | chr11 | 116698044 | 31 | 30 | 2988.730464 | 2835.992064 | 0.30586 |
| KLF2 | chr19 | 16435997 | 11 | 4 | 260.502065 | 184.3859649 | 0.30632 |
| PLA2G7 | chr6 | 46712398 | 35 | 32 | 3099.196989 | 2897.170327 | 0.30646 |
| CDKN2B-AS1 | chr9 | 21989125 | 24 | 17 | 698.5415435 | 561.1348949 | 0.30654 |
| ADRA2C | chr4 | 3763380 | 33 | 32 | 2984.73939 | 2812.716846 | 0.30662 |
| CDKN2B-AS1 | chr9 | 21965651 | 4 | 3 | 87.22222222 | 160.6060606 | 0.3068 |
| AKT2 | chr19 | 40796249 | 33 | 32 | 3034.509469 | 2868.797478 | 0.30682 |
| CDKN2B-AS1 | chr9 | 21977424 | 12 | 7 | 354.1709572 | 181.8650794 | 0.30692 |
| LCK | chr1 | 32740634 | 14 | 14 | 1229.642857 | 1285.984848 | 0.30696 |
| APOA1 | chr11 | 116692360 | 33 | 32 | 3103.692828 | 3055.848345 | 0.30702 |
| ABCG1 | chr21 | 43636932 | 33 | 32 | 3094.529678 | 3047.73434 | 0.30712 |
| APOA1 | chr11 | 116669429 | 5 | 2 | 211.0903885 | 138.0952381 | 0.30712 |
| APOA1 | chr11 | 116719745 | 24 | 28 | 2284.688645 | 2567.222222 | 0.30716 |
| CDKN1A | chr6 | 36653310 | 33 | 32 | 3235.866159 | 3065.611886 | 0.30724 |
| ABCG1 | chr21 | 43638737 | 5 | 6 | 126.4102564 | 238.0555556 | 0.3073 |
| F7 | chr13 | 113750833 | 32 | 32 | 3067.567988 | 2886.139416 | 0.30734 |
| APOA1 | chr11 | 116721491 | 32 | 32 | 3196.019683 | 3140.168893 | 0.30746 |
| APOA1 | chr11 | 116685840 | 10 | 11 | 686.7261905 | 645 | 0.3075 |
| CDKN2B-AS1 | chr9 | 21977228 | 30 | 33 | 2460.62435 | 2429.41292 | 0.30776 |
| TERT | chr5 | 1300341 | 27 | 27 | 2504.524212 | 2387.82247 | 0.30822 |
| ADRA2C | chr4 | 3767144 | 10 | 7 | 251.961927 | 248.4920635 | 0.30842 |
| APOA1 | chr11 | 116670904 | 35 | 32 | 2787.67031 | 2630.049773 | 0.30848 |
| TIMP3 | chr22 | 33192703 | 13 | 12 | 354.45217 | 404.3214877 | 0.3086 |
| COL15A1 | chr9 | 101832416 | 37 | 32 | 3435.695525 | 3005.749646 | 0.30876 |
| NOS3 | chr7 | 150690909 | 13 | 16 | 1246.666667 | 1575 | 0.30882 |
| APOA1 | chr11 | 116665819 | 2 | 2 | 43.07692308 | 133.3333333 | 0.30886 |
| CDKN2B-AS1 | chr9 | 21993172 | 38 | 32 | 3197.772589 | 2755.537129 | 0.30904 |
| APOA1 | chr11 | 116712038 | 29 | 30 | 2491.263287 | 2671.727276 | 0.30926 |
| LDLR | chr19 | 11221056 | 24 | 20 | 2214.444444 | 1913.131313 | 0.30956 |
| E2F6 | chr2 | 11613068 | 27 | 24 | 2621.915266 | 2364.393939 | 0.30968 |

| | | | | | | | |
|---|---|---|---|---|---|---|---|
| LDLR | chr19 | 11194179 | 14 | 18 | 1360 | 1787.5 | 0.30974 |
| LDLR | chr19 | 11218328 | 19 | 19 | 1798.979266 | 1736.150794 | 0.30994 |
| APOA1 | chr11 | 116662686 | 16 | 19 | 1464.986291 | 1624.354257 | 0.30994 |
| ABCG1 | chr21 | 43630007 | 36 | 32 | 3472.869238 | 3040.909342 | 0.3101 |
| TIMP3 | chr22 | 33195415 | 40 | 33 | 2550.236255 | 2027.591592 | 0.31016 |
| NOS3 | chr7 | 150688887 | 2 | 3 | 40 | 170 | 0.31054 |
| APOA1 | chr11 | 116661444 | 30 | 31 | 2187.838691 | 2365.623487 | 0.31068 |
| LOXL1 | chr15 | 74216981 | 18 | 19 | 887.6907011 | 832.4747475 | 0.31074 |
| CDKN2B-AS1 | chr9 | 21967055 | 32 | 29 | 1225.714675 | 1025.614318 | 0.31082 |
| CDKN2B-AS1 | chr9 | 22000346 | 35 | 33 | 1750.495557 | 1766.50055 | 0.31134 |
| LPA | chr6 | 11217363 | 35 | 32 | 3266.49227 | 2910.599626 | 0.31136 |
| CDKN2B-AS1 | chr9 | 22003179 | 34 | 32 | 2984.857652 | 2861.647672 | 0.31168 |
| F7 | chr13 | 113758307 | 29 | 28 | 2575.212759 | 2565.645107 | 0.31178 |
| CDKN2B-AS1 | chr9 | 21969690 | 6 | 3 | 143.7510443 | 147.7052359 | 0.31184 |
| ESR2 | chr14 | 64806482 | 18 | 15 | 1786.507937 | 1456.547619 | 0.31208 |
| ADRA2C | chr4 | 3762112 | 32 | 31 | 2751.479036 | 2552.36658 | 0.3121 |
| BCL2 | chr18 | 60988255 | 12 | 19 | 387.0812325 | 719.4684701 | 0.31214 |
| F7 | chr13 | 113750742 | 28 | 26 | 2612.886003 | 2487.18254 | 0.31238 |
| TERT | chr5 | 1300762 | 26 | 25 | 2338.311966 | 2143.246753 | 0.31242 |
| BCL2 | chr18 | 60990310 | 4 | 2 | 120 | 40 | 0.3125 |
| CDKN2B-AS1 | chr9 | 22005133 | 2 | 4 | 40 | 198.3333333 | 0.31258 |
| ABCG1 | chr21 | 43634104 | 33 | 31 | 3147.894441 | 3007.43767 | 0.31266 |
| ABCA1 | chr9 | 107693263 | 27 | 25 | 2661.63961 | 2412.81746 | 0.3127 |
| CDKN2B-AS1 | chr9 | 22005569 | 4 | 4 | 86.83716964 | 170.8074534 | 0.31276 |
| HMGCR | chr5 | 74630929 | 15 | 12 | 1169.52381 | 1025.079365 | 0.31286 |
| LCK | chr1 | 32707627 | 12 | 9 | 1111.599512 | 870 | 0.31316 |
| TFPI | chr2 | 188424544 | 19 | 21 | 1796.306194 | 2032.5 | 0.31328 |
| LCK | chr1 | 32713256 | 32 | 32 | 3109.181617 | 3063.110443 | 0.31386 |
| COL15A1 | chr9 | 101702598 | 35 | 32 | 1179.926587 | 1152.884505 | 0.31394 |
| CDKN2B-AS1 | chr9 | 21980590 | 33 | 33 | 2817.827042 | 2719.735981 | 0.31442 |
| TIMP3 | chr22 | 33191603 | 34 | 32 | 3029.077292 | 2913.71474 | 0.3147 |
| APOA1 | chr11 | 116714674 | 26 | 28 | 2486.230159 | 2748.974359 | 0.31474 |
| APOA1 | chr11 | 116702363 | 26 | 27 | 1665.147075 | 1536.582168 | 0.31522 |
| BAX | chr19 | 49465470 | 5 | 4 | 118.7936508 | 182.8700711 | 0.31532 |
| PLA2G7 | chr6 | 46709989 | 33 | 32 | 2668.43417 | 2657.345896 | 0.31536 |
| NOS3 | chr7 | 150690703 | 2 | 6 | 200 | 532.3809524 | 0.31546 |
| F2RL3 | chr19 | 16991886 | 32 | 31 | 3043.55467 | 3001.702742 | 0.31584 |
| FOXP3 | chrX | 49126850 | 10 | 9 | 396.4285714 | 442.7380952 | 0.31608 |
| APOA1 | chr11 | 116692374 | 33 | 32 | 3057.781675 | 3008.97256 | 0.31624 |
| APOA1 | chr11 | 116688377 | 34 | 32 | 3071.872113 | 2964.686463 | 0.31636 |
| CDKN2B-AS1 | chr9 | 21974374 | 5 | 7 | 112.0769231 | 219.3804714 | 0.31648 |
| LOXL1 | chr15 | 74244030 | 18 | 19 | 884.1281233 | 1027.142857 | 0.31652 |
| NOS3 | chr7 | 150686714 | 35 | 32 | 3243.749848 | 2912.758926 | 0.31674 |
| APOA1 | chr11 | 116679733 | 35 | 32 | 2959.295953 | 2762.939676 | 0.31702 |
| FGF2 | chr4 | 123741202 | 40 | 32 | 3839.15222 | 3007.639421 | 0.31708 |
| CDKN1A | chr6 | 36654741 | 32 | 32 | 3026.909555 | 3071.260332 | 0.31712 |
| APOA1 | chr11 | 116704364 | 34 | 32 | 3512.960094 | 3073.156395 | 0.31716 |

| | | | | | | | |
|---|---|---|---|---|---|---|---|
| F2RL3 | chr19 | 16992397 | 24 | 17 | 2209.393939 | 1634.761905 | 0.3172 |
| F7 | chr13 | 113756922 | 31 | 32 | 2893.298471 | 3044.571782 | 0.31724 |
| ARHGEF18 | chr19 | 7469017 | 2 | 2 | 45 | 133.3333333 | 0.3173 |
| BCL2 | chr18 | 60988584 | 32 | 33 | 2090.336093 | 2274.583148 | 0.3175 |
| CDKN2B-AS1 | chr9 | 21971186 | 12 | 13 | 347.5396825 | 416.9099741 | 0.31754 |
| CDKN1A | chr6 | 36652158 | 31 | 30 | 3001.480464 | 2847.267316 | 0.31758 |
| TERT | chr5 | 1299474 | 27 | 29 | 2455.556277 | 2718.695471 | 0.3176 |
| LCK | chr1 | 32741452 | 9 | 3 | 717.9761905 | 263.3333333 | 0.31766 |
| LPA | chr6 | 160952004 | 32 | 33 | 2481.645598 | 2464.295639 | 0.31768 |
| ESR2 | chr14 | 64807610 | 35 | 32 | 3431.492982 | 3102.669282 | 0.31778 |
| KLF2 | chr19 | 16433560 | 33 | 33 | 2700.048609 | 2609.792445 | 0.31786 |
| FOXP3 | chrX | 49127164 | 20 | 23 | 1888.333333 | 2104.801587 | 0.3179 |
| LOXL1 | chr15 | 74218468 | 33 | 32 | 2610.860456 | 2456.094532 | 0.31794 |
| APOA1 | chr11 | 116702373 | 31 | 31 | 2873.152231 | 2932.878477 | 0.3181 |
| F7 | chr13 | 113752461 | 33 | 32 | 3014.516382 | 2987.79658 | 0.31812 |
| CDKN2B-AS1 | chr9 | 21986219 | 40 | 33 | 3526.061722 | 2827.162386 | 0.31828 |
| ESR2 | chr14 | 64693699 | 32 | 32 | 3078.465197 | 3043.849151 | 0.31838 |
| APOA1 | chr11 | 116687947 | 14 | 13 | 368.3402863 | 406.4897465 | 0.31868 |
| GALNT2 | chr1 | 230185640 | 38 | 32 | 3260.909269 | 2801.40532 | 0.31876 |
| APOA1 | chr11 | 116705568 | 7 | 10 | 674.2424242 | 910 | 0.31884 |
| F2RL3 | chr19 | 16990184 | 29 | 33 | 2738.016781 | 3031.696612 | 0.31942 |
| FOXP3 | chrX | 49127035 | 17 | 11 | 1554.642857 | 962.5 | 0.3195 |
| MCPH1 | chr8 | 6259994 | 14 | 15 | 1338.095238 | 1371.547619 | 0.31954 |
| LOXL1 | chr15 | 74215871 | 31 | 32 | 1422.412519 | 1584.408555 | 0.31956 |
| APOA1 | chr11 | 116714629 | 29 | 32 | 2742.238513 | 3072.111152 | 0.31962 |
| GALNT2 | chr1 | 230190909 | 35 | 32 | 3277.203528 | 3039.029596 | 0.31994 |
| NOS3 | chr7 | 150690454 | 32 | 33 | 2034.509939 | 1980.614113 | 0.31998 |
| NOS3 | chr7 | 150685946 | 4 | 5 | 90.79365079 | 186.0714286 | 0.32034 |
| ESR1 | chr6 | 152003002 | 8 | 10 | 188.8721153 | 305.1148376 | 0.32052 |
| COL15A1 | chr9 | 101705492 | 35 | 32 | 3038.614994 | 2704.022491 | 0.32066 |
| PLA2G7 | chr6 | 46710585 | 34 | 32 | 3201.4321 | 2921.515195 | 0.32084 |
| DHFR | chr5 | 79956636 | 35 | 32 | 3376.671453 | 3047.642999 | 0.32104 |
| GALNT2 | chr1 | 230189964 | 29 | 32 | 2239.561272 | 2583.159496 | 0.32132 |
| BCL2 | chr18 | 60994347 | 26 | 26 | 2372.613914 | 2270.602731 | 0.32134 |
| KLF2 | chr19 | 16434843 | 3 | 3 | 238.8888889 | 180 | 0.32138 |
| ATF3 | chr1 | 212779882 | 11 | 8 | 299.1269841 | 271.1507936 | 0.32148 |
| ARHGEF18 | chr19 | 7461114 | 9 | 15 | 721.6558442 | 1295.865801 | 0.3216 |
| CDKN2B-AS1 | chr9 | 21990330 | 16 | 9 | 414.0742655 | 298.5521825 | 0.3216 |
| BCL2 | chr18 | 60988535 | 36 | 33 | 2582.708446 | 2456.767 | 0.32162 |
| E2F6 | chr2 | 11614052 | 34 | 32 | 3213.701801 | 3064.431618 | 0.32182 |
| F7 | chr13 | 113752711 | 33 | 32 | 3189.481163 | 3134.541195 | 0.3219 |
| LPA | chr6 | 11225106 | 2 | 3 | 40 | 165 | 0.32194 |
| TERT | chr5 | 1296728 | 26 | 27 | 2320.502137 | 2309.648823 | 0.32202 |
| LDLR | chr19 | 11215861 | 20 | 20 | 1920.151515 | 1963.333333 | 0.32206 |
| LOXL1 | chr15 | 74215522 | 11 | 11 | 1025.512266 | 1063.333333 | 0.32206 |
| P2RY12 | chr3 | 151104209 | 29 | 32 | 2809.84016 | 3054.034209 | 0.3223 |
| ADRA2C | chr4 | 3761382 | 31 | 31 | 2748.847541 | 2824.094198 | 0.32252 |

| | | | | | | | |
|---|---|---|---|---|---|---|---|
| APOA1 | chr11 | 116701581 | 25 | 27 | 2179.047619 | 2434.904175 | 0.32258 |
| LCK | chr1 | 32741518 | 29 | 30 | 2272.740702 | 2229.689485 | 0.32274 |
| AMOTL2 | chr3 | 134099773 | 26 | 26 | 2382.447275 | 2222.813853 | 0.32292 |
| DHFR | chr5 | 79956441 | 33 | 32 | 3218.931552 | 3069.545682 | 0.32306 |
| PECAM1 | chr17 | 62398676 | 3 | 5 | 63.96135265 | 193.3253589 | 0.32318 |
| ABCA1 | chr9 | 107696501 | 36 | 31 | 3579.531165 | 3008.576817 | 0.32342 |
| TERT | chr5 | 1295675 | 7 | 4 | 212.8924584 | 92.27272727 | 0.32342 |
| APOA1 | chr11 | 116661855 | 33 | 32 | 2030.872031 | 2090.710954 | 0.3236 |
| MCPH1 | chr8 | 6470560 | 32 | 32 | 2984.60312 | 3030.425882 | 0.32374 |
| ESR2 | chr14 | 64814798 | 22 | 24 | 2213.015873 | 2327.678571 | 0.32388 |
| TIMP3 | chr22 | 33192680 | 27 | 22 | 866.0421766 | 644.958758 | 0.32414 |
| PHACTR1 | chr6 | 12709693 | 25 | 23 | 1789.895937 | 1777.150072 | 0.32438 |
| APOA1 | chr11 | 116696996 | 17 | 23 | 1575.584416 | 2193.409091 | 0.32462 |
| LDLR | chr19 | 11192195 | 27 | 26 | 2364.519382 | 2354.363969 | 0.32468 |
| APOA1 | chr11 | 116686183 | 12 | 14 | 1174.242424 | 1330.833333 | 0.3247 |
| CDKN2BAS | chr9 | 22103470 | 33 | 33 | 1569.998869 | 1472.445729 | 0.32484 |
| TIMP3 | chr22 | 33187267 | 35 | 32 | 3303.965214 | 2933.888576 | 0.32492 |
| APOA1 | chr11 | 116680629 | 31 | 30 | 2885.237263 | 2869.383394 | 0.32498 |
| CTGF | chr6 | 132278893 | 31 | 32 | 2905.747503 | 2926.276813 | 0.32528 |
| F7 | chr13 | 113750529 | 4 | 3 | 88.57142857 | 160 | 0.32538 |
| ABCB1 | chr7 | 87133367 | 14 | 14 | 727.0955891 | 630.4833412 | 0.32544 |
| NOS3 | chr7 | 150710526 | 29 | 31 | 1674.281761 | 1918.414865 | 0.32552 |
| LDLR | chr19 | 11220949 | 13 | 18 | 1240.47619 | 1769.047619 | 0.32552 |
| CTGF | chr6 | 132276359 | 34 | 32 | 3200.982076 | 3055.539038 | 0.32558 |
| APOA1 | chr11 | 116677454 | 32 | 31 | 3042.055585 | 2812.965789 | 0.32578 |
| ESR2 | chr14 | 64806585 | 7 | 8 | 615 | 743.3333333 | 0.32582 |
| APOA1 | chr11 | 116703837 | 32 | 32 | 2884.201449 | 2813.808605 | 0.32638 |
| NOS3 | chr7 | 150707415 | 12 | 9 | 592.8571429 | 378.7698413 | 0.3264 |
| CTGF | chr6 | 132272555 | 3 | 2 | 84.44055944 | 110.7142857 | 0.3264 |
| ADRA2C | chr4 | 3759584 | 32 | 33 | 2357.287434 | 2331.185179 | 0.32642 |
| NOS3 | chr7 | 150686270 | 10 | 7 | 516.9047619 | 310 | 0.32662 |
| GCK | chr7 | 44231036 | 38 | 32 | 3124.142449 | 2691.422841 | 0.32698 |
| APOA1 | chr11 | 116695774 | 6 | 5 | 490 | 455 | 0.32724 |
| ADRA2C | chr4 | 3765535 | 10 | 13 | 939.047619 | 1148.571429 | 0.3278 |
| LCK | chr1 | 32741096 | 24 | 24 | 939.0048922 | 848.9978583 | 0.32792 |
| LOXL1 | chr15 | 74219242 | 17 | 17 | 581.7484112 | 668.0471321 | 0.32792 |
| KLF2 | chr19 | 16438187 | 34 | 32 | 2393.529533 | 2360.260399 | 0.3285 |
| PLA2G7 | chr6 | 46709809 | 36 | 32 | 3459.29794 | 3039.117672 | 0.32856 |
| NOS3 | chr7 | 150710074 | 27 | 30 | 1155.237318 | 1388.106879 | 0.329 |
| APOA1 | chr11 | 116688089 | 35 | 32 | 3290.798415 | 3045.075262 | 0.32948 |
| P2RY12 | chr3 | 151105406 | 24 | 19 | 676.1601551 | 605.7096098 | 0.32984 |
| LCK | chr1 | 32740745 | 23 | 19 | 2212.432845 | 1867.380952 | 0.3301 |
| APOA1 | chr11 | 116691769 | 30 | 32 | 2614.449108 | 2672.450196 | 0.33012 |
| APOA1 | chr11 | 116719848 | 35 | 32 | 3342.860744 | 3099.099301 | 0.33032 |
| TMPO | chr12 | 98904929 | 30 | 26 | 2968.557602 | 2480.590521 | 0.33042 |
| BAX | chr19 | 49450783 | 4 | 4 | 225.5952381 | 185.3571429 | 0.33042 |
| TIMP3 | chr22 | 33188966 | 33 | 33 | 2886.162264 | 2688.974487 | 0.33084 |

| | | | | | | | |
|---|---|---|---|---|---|---|---|
| CDKN2B-AS1 | chr9 | 21966908 | 19 | 11 | 494.2183739 | 355.1886405 | 0.331 |
| CDKN2B-AS1 | chr9 | 21966520 | 26 | 27 | 834.6425918 | 946.4622948 | 0.33134 |
| LDLR | chr19 | 11198556 | 33 | 32 | 3312.052843 | 3152.624272 | 0.33136 |
| LDLR | chr19 | 11218350 | 11 | 10 | 1090.909091 | 967.5 | 0.33146 |
| ALDOA | chr16 | 30057701 | 29 | 32 | 1645.470509 | 1697.391618 | 0.33168 |
| CDKN2B-AS1 | chr9 | 21990049 | 18 | 17 | 707.5396825 | 760.9920635 | 0.33168 |
| F7 | chr13 | 113752643 | 32 | 31 | 3077.16954 | 2933.427051 | 0.33204 |
| LDLR | chr19 | 11215960 | 31 | 32 | 2942.043048 | 3089.9255 | 0.33218 |
| ADRA2C | chr4 | 3763652 | 31 | 33 | 2923.354464 | 3022.671046 | 0.33228 |
| CDKN2B-AS1 | chr9 | 21974137 | 11 | 14 | 319.0223665 | 494.5848596 | 0.33228 |
| NOS3 | chr7 | 150709536 | 21 | 24 | 1950.4329 | 2318.650794 | 0.33244 |
| LCK | chr1 | 32739663 | 18 | 20 | 576.8540933 | 716.1549922 | 0.333 |
| PECAM1 | chr17 | 62409802 | 36 | 33 | 1972.970263 | 1718.132423 | 0.33316 |
| LDLR | chr19 | 11221659 | 31 | 32 | 2850.443337 | 2997.276627 | 0.33342 |
| APOA1 | chr11 | 116680133 | 24 | 27 | 2276.310691 | 2495.873016 | 0.33404 |
| CDKN1A | chr6 | 36653413 | 24 | 25 | 885.4825291 | 1047.763642 | 0.33422 |
| AKT2 | chr19 | 40795610 | 14 | 15 | 1400 | 1428.571429 | 0.33428 |
| LOXL1 | chr15 | 74214977 | 31 | 32 | 1561.150648 | 1501.791972 | 0.33452 |
| APOA1 | chr11 | 116692324 | 33 | 31 | 2840.878646 | 2783.906079 | 0.33472 |
| TIMP3 | chr22 | 33190612 | 33 | 31 | 2641.203475 | 2379.809835 | 0.33488 |
| MIA3 | chr1 | 222816714 | 27 | 27 | 2406.165459 | 2488.964646 | 0.33488 |
| ADRA2C | chr4 | 3768519 | 8 | 6 | 208.2862923 | 222.4088656 | 0.33488 |
| FOXP3 | chrX | 49115374 | 30 | 32 | 2764.80039 | 3044.457071 | 0.3351 |
| APOA1 | chr11 | 116692490 | 33 | 32 | 2959.856435 | 2944.316025 | 0.33534 |
| KLF2 | chr19 | 16437992 | 29 | 29 | 1217.44787 | 1321.15838 | 0.33604 |
| FOXP3 | chrX | 49127057 | 18 | 17 | 1743.650794 | 1602.5 | 0.33618 |
| NOS3 | chr7 | 150688674 | 37 | 32 | 3191.503123 | 2816.58464 | 0.3364 |
| APOA1 | chr11 | 116717824 | 32 | 31 | 3036.937799 | 2981.279355 | 0.3369 |
| KLF2 | chr19 | 16433204 | 25 | 22 | 2409.569597 | 2071.831502 | 0.33692 |
| PHACTR1 | chr6 | 13275950 | 12 | 14 | 405.3067827 | 553.8756719 | 0.3371 |
| TERT | chr5 | 1300691 | 33 | 32 | 3150.729015 | 3008.706323 | 0.33722 |
| CDKN2B-AS1 | chr9 | 22006076 | 4 | 6 | 95.19607843 | 223.3838384 | 0.33738 |
| CDKN1A | chr6 | 36648907 | 31 | 30 | 2906.109021 | 2739.881579 | 0.33752 |
| CTGF | chr6 | 132272791 | 28 | 26 | 1662.996032 | 1427.469899 | 0.33756 |
| APOA1 | chr11 | 116718233 | 34 | 32 | 3357.916812 | 3116.498866 | 0.33776 |
| BCL2 | chr18 | 60988474 | 15 | 19 | 533.6977872 | 569.4797045 | 0.33802 |
| LCK | chr1 | 32741116 | 17 | 14 | 527.8143275 | 510.2869026 | 0.33808 |
| FOXP3 | chrX | 49115669 | 25 | 21 | 2052.851095 | 1825.885226 | 0.33832 |
| ADRA2C | chr4 | 3760220 | 37 | 32 | 3101.441071 | 2626.035874 | 0.33854 |
| NOS3 | chr7 | 150706596 | 26 | 25 | 2481.658342 | 2264.051504 | 0.33884 |
| AKT2 | chr19 | 40792595 | 40 | 32 | 3826.066947 | 3089.815852 | 0.33902 |
| MIA3 | chr1 | 222809187 | 35 | 32 | 3265.924249 | 3022.75061 | 0.33922 |
| APOA1 | chr11 | 116693465 | 21 | 23 | 1051.469091 | 1272.326564 | 0.33952 |
| GALNT2 | chr1 | 230193224 | 33 | 32 | 3102.93575 | 2951.144425 | 0.3396 |
| COL15A1 | chr9 | 101697876 | 39 | 32 | 3454.358198 | 2886.958906 | 0.33962 |
| LOXL1 | chr15 | 74244121 | 9 | 6 | 643.9935065 | 375.7142857 | 0.33974 |
| ADRA2C | chr4 | 3765592 | 22 | 27 | 1822.51443 | 2321.428849 | 0.33998 |

| | | | | | | | |
|---|---|---|---|---|---|---|---|
| CDKN2B-AS1 | chr9 | 21966440 | 3 | 8 | 70 | 254.0208648 | 0.34018 |
| APOA1 | chr11 | 116712214 | 32 | 33 | 2359.234858 | 2332.07441 | 0.3402 |
| LPA | chr6 | 160950981 | 17 | 22 | 1477.258297 | 2027.2669 | 0.34022 |
| F2RL3 | chr19 | 16990947 | 31 | 31 | 2821.544184 | 2885.319015 | 0.3403 |
| ADRA2C | chr4 | 3758825 | 38 | 32 | 3067.895665 | 2661.51818 | 0.34042 |
| APOA1 | chr11 | 116700532 | 2 | 2 | 40 | 128.5714286 | 0.34072 |
| ALDOA | chr16 | 30064332 | 2 | 2 | 40 | 128.5714286 | 0.34072 |
| F2RL3 | chr19 | 16998416 | 2 | 2 | 40 | 128.5714286 | 0.34072 |
| AMOTL2 | chr3 | 134098150 | 2 | 2 | 40 | 128.5714286 | 0.34072 |
| FOXP3 | chrX | 49123546 | 2 | 2 | 40 | 128.5714286 | 0.34072 |
| APOA1 | chr11 | 116668994 | 22 | 20 | 2085.2442 | 1934.95671 | 0.34088 |
| ESR1 | chr6 | 152006557 | 11 | 11 | 1100 | 1087.5 | 0.3409 |
| NOS3 | chr7 | 150710807 | 4 | 3 | 109.4444444 | 62.22222222 | 0.3409 |
| APOA1 | chr11 | 116719883 | 34 | 32 | 3220.618538 | 3076.635859 | 0.34096 |
| LPA | chr6 | 160952811 | 36 | 32 | 3012.169452 | 2739.800588 | 0.34116 |
| FOXP3 | chrX | 49122411 | 31 | 30 | 2856.26199 | 2827.597336 | 0.34118 |
| TMPO | chr12 | 98904936 | 30 | 30 | 3059.62371 | 2940.867794 | 0.34142 |
| APOA1 | chr11 | 116698333 | 3 | 3 | 210 | 263.3333333 | 0.34142 |
| F2RL3 | chr19 | 16997331 | 32 | 32 | 2438.091506 | 2531.437583 | 0.34174 |
| ABCG1 | chr21 | 43634402 | 3 | 2 | 251.4285714 | 135.5555556 | 0.3418 |
| TERT | chr5 | 1298927 | 28 | 30 | 1274.512519 | 1478.220075 | 0.3419 |
| APOA1 | chr11 | 116699097 | 23 | 28 | 1742.037789 | 2237.267316 | 0.3421 |
| CDKN1A | chr6 | 36650778 | 6 | 4 | 160.5906593 | 181.8950373 | 0.3421 |
| APOA1 | chr11 | 116659130 | 4 | 4 | 86.2288839 | 168.3805284 | 0.34216 |
| APOA1 | chr11 | 116717749 | 32 | 32 | 3033.446091 | 3070.721121 | 0.34232 |
| KLF2 | chr19 | 16433260 | 34 | 31 | 3269.749591 | 3020.249059 | 0.3425 |
| TERT | chr5 | 1299513 | 26 | 30 | 2410.27417 | 2858.907204 | 0.34264 |
| ABCA1 | chr9 | 107698088 | 8 | 8 | 480.8333333 | 374.7619048 | 0.34274 |
| LPA | chr6 | 11218159 | 40 | 32 | 3713.332461 | 2935.072597 | 0.34288 |
| APOA1 | chr11 | 116718541 | 34 | 32 | 3195.181172 | 2966.663679 | 0.34318 |
| F2RL3 | chr19 | 16990272 | 30 | 30 | 2911.62509 | 2869.586998 | 0.3432 |
| NOS3 | chr7 | 150695676 | 27 | 25 | 2559.751082 | 2313.351648 | 0.34336 |
| AMOTL2 | chr3 | 134104012 | 10 | 3 | 950 | 300 | 0.34344 |
| CDKN2B-AS1 | chr9 | 21967209 | 7 | 10 | 177.2646473 | 331.998002 | 0.34356 |
| FOXP3 | chrX | 49115399 | 31 | 31 | 2809.235328 | 2888.67524 | 0.34408 |
| CDKN2BAS | chr9 | 22103897 | 39 | 32 | 3605.97493 | 3001.653938 | 0.34416 |
| CDKN2B-AS1 | chr9 | 22005892 | 13 | 13 | 359.5761968 | 439.6068551 | 0.34422 |
| APOA1 | chr11 | 116707266 | 22 | 25 | 2190.411255 | 2407.5 | 0.34438 |
| CDKN2B-AS1 | chr9 | 21971133 | 24 | 22 | 851.460928 | 714.4588744 | 0.34502 |
| F7 | chr13 | 113752507 | 30 | 32 | 2825.435727 | 2855.307028 | 0.34506 |
| ARHGEF18 | chr19 | 7461905 | 33 | 32 | 3127.305448 | 2970.414356 | 0.34534 |
| ALOX15 | chr17 | 4550224 | 34 | 32 | 2989.126348 | 2871.627556 | 0.34552 |
| ABCG1 | chr21 | 43630101 | 36 | 33 | 1618.533537 | 1396.714067 | 0.34558 |
| NOS3 | chr7 | 150683282 | 2 | 2 | 40 | 119.5804196 | 0.34568 |
| FOXP3 | chrX | 49131152 | 35 | 32 | 3342.160244 | 3090.2722 | 0.34618 |
| NOS3 | chr7 | 150710304 | 30 | 32 | 1547.90446 | 1551.203905 | 0.34656 |
| ABCG1 | chr21 | 43632053 | 9 | 7 | 880 | 700 | 0.3466 |

| | | | | | | | |
|---|---|---|---|---|---|---|---|
| TIMP3 | chr22 | 33194966 | 9 | 5 | 880 | 500 | 0.3466 |
| ABCG1 | chr21 | 43634695 | 9 | 8 | 790.8333333 | 754.1666667 | 0.34684 |
| ARHGEF18 | chr19 | 7466798 | 36 | 32 | 3483.513675 | 3037.368137 | 0.347 |
| LCK | chr1 | 32740347 | 31 | 31 | 2672.907693 | 2757.607105 | 0.34702 |
| PHACTR1 | chr6 | 12711218 | 37 | 32 | 3022.530891 | 2680.437351 | 0.34706 |
| NOS3 | chr7 | 150693574 | 4 | 3 | 95.55555555 | 60 | 0.34724 |
| GCK | chr7 | 44232647 | 34 | 33 | 2454.111657 | 2476.139606 | 0.3473 |
| LPA | chr6 | 160944263 | 27 | 31 | 2315.722986 | 2552.086364 | 0.34744 |
| CDKN1A | chr6 | 36650745 | 10 | 8 | 294.2225037 | 303.2792087 | 0.34804 |
| AKT2 | chr19 | 40794388 | 35 | 32 | 3202.232803 | 2965.204655 | 0.34812 |
| ABCG1 | chr21 | 43634653 | 3 | 2 | 62.22222222 | 140 | 0.34834 |
| ADRA2C | chr4 | 3768458 | 3 | 2 | 62.22222222 | 140 | 0.34834 |
| LPA | chr6 | 160948111 | 31 | 32 | 2911.21922 | 2955.644266 | 0.34836 |
| APOA1 | chr11 | 116703972 | 34 | 32 | 3089.310964 | 2958.524294 | 0.34848 |
| F2RL3 | chr19 | 16991537 | 32 | 32 | 2379.945336 | 2463.830455 | 0.34892 |
| APOA1 | chr11 | 116669828 | 32 | 29 | 2245.192733 | 1867.37885 | 0.34904 |
| MCPH1 | chr8 | 6470040 | 18 | 12 | 1696.355311 | 1164.123377 | 0.34928 |
| HMGCR | chr5 | 74626355 | 15 | 15 | 1294.31606 | 1352.81746 | 0.34944 |
| ALDOA | chr16 | 30062482 | 32 | 32 | 2944.59217 | 2990.516759 | 0.3495 |
| FOXP3 | chrX | 49129680 | 28 | 28 | 1615.071474 | 1515.902979 | 0.3495 |
| LCK | chr1 | 32740711 | 20 | 19 | 1880.555556 | 1728.051948 | 0.34966 |
| LCK | chr1 | 32741583 | 32 | 32 | 3130.492679 | 3087.323936 | 0.34996 |
| LPA | chr6 | 160949330 | 33 | 32 | 2850.276391 | 2824.101922 | 0.35032 |
| GCK | chr7 | 44234284 | 36 | 33 | 3134.342767 | 2787.803057 | 0.35032 |
| APOA1 | chr11 | 116705130 | 14 | 11 | 1326.666667 | 998.2142857 | 0.3505 |
| GALNT2 | chr1 | 230193439 | 32 | 33 | 2171.853322 | 2142.54833 | 0.35062 |
| LDLR | chr19 | 11219928 | 8 | 6 | 783.3333333 | 600 | 0.35062 |
| ABCA1 | chr9 | 107692157 | 8 | 9 | 783.3333333 | 900 | 0.35062 |
| CDKN1A | chr6 | 36644625 | 33 | 33 | 2424.254791 | 2509.424889 | 0.35074 |
| APOA1 | chr11 | 116713111 | 35 | 32 | 3410.582685 | 3059.596152 | 0.35076 |
| CDKN2BAS | chr9 | 22103513 | 34 | 30 | 1330.902452 | 1100.969998 | 0.35098 |
| APOA1 | chr11 | 116685934 | 5 | 6 | 459.7222222 | 580 | 0.35126 |
| CDKN2B-AS1 | chr9 | 21968455 | 10 | 10 | 276.4790765 | 355.6565656 | 0.35142 |
| F7 | chr13 | 113757079 | 33 | 32 | 3083.597867 | 2947.804575 | 0.3515 |
| CDKN1A | chr6 | 36643691 | 10 | 6 | 724.1666667 | 371.9047619 | 0.3517 |
| COL15A1 | chr9 | 101701081 | 39 | 32 | 3346.290619 | 2822.731795 | 0.35172 |
| LDLR | chr19 | 11196270 | 33 | 32 | 3124.43114 | 2954.901042 | 0.35182 |
| NOS3 | chr7 | 150707811 | 11 | 12 | 274.6559238 | 351.9932396 | 0.35208 |
| CDKN1A | chr6 | 36650301 | 15 | 9 | 701.8922306 | 485.2777778 | 0.3521 |
| APOA1 | chr11 | 116691090 | 33 | 32 | 2809.940381 | 2655.311864 | 0.3522 |
| NOS3 | chr7 | 150707942 | 21 | 21 | 1462.420635 | 1549.383966 | 0.35224 |
| GALNT2 | chr1 | 230189799 | 12 | 13 | 1152.380952 | 1280 | 0.35224 |
| HMGCR | chr5 | 74627234 | 34 | 32 | 3291.956706 | 3130.823596 | 0.35226 |
| TFPI | chr2 | 188343494 | 34 | 32 | 3129.447601 | 3005.497772 | 0.35238 |
| CDKN2B-AS1 | chr9 | 21990171 | 14 | 16 | 637.6556777 | 644.9314574 | 0.35258 |
| CDKN2B-AS1 | chr9 | 22005787 | 7 | 9 | 209.3073593 | 322.8921569 | 0.3526 |
| TERT | chr5 | 1298111 | 36 | 32 | 3280.398841 | 2872.411296 | 0.35274 |

| | | | | | | | |
|---|---|---|---|---|---|---|---|
| FOXP3 | chrX | 49125924 | 2 | 2 | 53.05263158 | 128.7878788 | 0.3528 |
| LOXL1 | chr15 | 74219146 | 7 | 5 | 230.9260915 | 222.4880383 | 0.35286 |
| ESR2 | chr14 | 64806925 | 31 | 29 | 2975.998169 | 2830.161228 | 0.35316 |
| ALDOA | chr16 | 30059669 | 19 | 13 | 1849.807692 | 1219.047619 | 0.35356 |
| ABCA1 | chr9 | 107691566 | 2 | 2 | 41.42857143 | 123.8636364 | 0.35358 |
| APOA1 | chr11 | 116711874 | 7 | 10 | 364.8593074 | 411.1255411 | 0.35374 |
| LCK | chr1 | 32713985 | 2 | 2 | 40 | 120.7843137 | 0.35374 |
| ADRA2C | chr4 | 3761826 | 32 | 33 | 2755.653542 | 2746.693573 | 0.35412 |
| APOA1 | chr11 | 116695461 | 4 | 7 | 309.5238095 | 597.7777778 | 0.3542 |
| CDKN2B-AS1 | chr9 | 21967583 | 15 | 12 | 378.0454411 | 364.9937813 | 0.3543 |
| F2RL3 | chr19 | 16993509 | 33 | 32 | 3075.479934 | 3025.630432 | 0.35442 |
| ARHGEF18 | chr19 | 7468070 | 20 | 14 | 1882.738095 | 1359.047619 | 0.35446 |
| CDKN2B-AS1 | chr9 | 21980685 | 31 | 33 | 2863.396868 | 2939.740856 | 0.35462 |
| NOS3 | chr7 | 150690969 | 23 | 26 | 2346.9315 | 2558.974359 | 0.35508 |
| APOA1 | chr11 | 116704540 | 33 | 32 | 3104.195754 | 3050.737388 | 0.3551 |
| ARHGEF18 | chr19 | 7537795 | 2 | 2 | 86.66666667 | 160 | 0.35516 |
| TFPI | chr2 | 188425278 | 25 | 27 | 2351.996753 | 2467.038517 | 0.35534 |
| AKT2 | chr19 | 40792008 | 32 | 32 | 2840.893281 | 2908.105976 | 0.35566 |
| LDLR | chr19 | 11191513 | 5 | 7 | 485.7142857 | 645.7142857 | 0.35576 |
| AKT2 | chr19 | 40792086 | 4 | 7 | 400 | 666.6666667 | 0.35592 |
| APOA1 | chr11 | 116711891 | 13 | 16 | 1026.906427 | 1143.690476 | 0.35596 |
| CDKN2B-AS1 | chr9 | 21971155 | 16 | 18 | 501.4013764 | 633.2073482 | 0.35598 |
| GCK | chr7 | 44237749 | 32 | 31 | 3039.94281 | 3013.717306 | 0.35656 |
| AMOTL2 | chr3 | 134102002 | 10 | 12 | 883.1652661 | 1119.047619 | 0.35666 |
| APOA1 | chr11 | 116717185 | 32 | 32 | 3085.152928 | 3113.828766 | 0.35684 |
| NOS3 | chr7 | 150706435 | 2 | 3 | 160 | 170 | 0.35754 |
| CDKN1A | chr6 | 36655007 | 33 | 33 | 2535.013871 | 2438.591251 | 0.3576 |
| ADRA2C | chr4 | 3759218 | 33 | 31 | 2584.17243 | 2519.628987 | 0.35786 |
| CDKN2B-AS1 | chr9 | 21967500 | 5 | 5 | 133.8888889 | 211.6483517 | 0.35786 |
| CDKN1A | chr6 | 36650946 | 24 | 21 | 670.7970517 | 652.8649768 | 0.35788 |
| FOXP3 | chrX | 49127004 | 3 | 4 | 60 | 165 | 0.35802 |
| PHACTR1 | chr6 | 12711899 | 34 | 32 | 3211.852496 | 3058.71147 | 0.3585 |
| GCK | chr7 | 44228523 | 23 | 20 | 1721.57039 | 1595.10101 | 0.35852 |
| TERT | chr5 | 1295682 | 12 | 12 | 379.9494949 | 444.505772 | 0.35852 |
| ADRA2C | chr4 | 3765151 | 18 | 14 | 624.9495534 | 549.0589966 | 0.35856 |
| FGF2 | chr4 | 123740122 | 29 | 30 | 2196.646058 | 2190.837684 | 0.3588 |
| CDKN2B-AS1 | chr9 | 21968647 | 6 | 5 | 141.984127 | 181.5360079 | 0.3589 |
| PECAM1 | chr17 | 62409355 | 32 | 29 | 1341.09637 | 1117.820123 | 0.3591 |
| FGF2 | chr4 | 123746885 | 32 | 32 | 2986.252624 | 3038.253013 | 0.35912 |
| F2RL3 | chr19 | 16991333 | 32 | 32 | 3089.208025 | 3033.394712 | 0.35914 |
| APOA1 | chr11 | 116723838 | 19 | 14 | 511.180612 | 449.4160318 | 0.35956 |
| CDKN2B-AS1 | chr9 | 21996208 | 15 | 18 | 421.3077274 | 583.6650987 | 0.3596 |
| APOA1 | chr11 | 116682286 | 36 | 32 | 3502.807566 | 3089.429755 | 0.35992 |
| ESR1 | chr6 | 152004146 | 37 | 32 | 3477.490866 | 2974.514158 | 0.36004 |
| CDKN2B-AS1 | chr9 | 21970749 | 13 | 16 | 358.7361255 | 525.310151 | 0.36004 |
| APOA1 | chr11 | 116709296 | 32 | 32 | 2250.133663 | 2338.619903 | 0.36008 |
| BAX | chr19 | 49465475 | 3 | 3 | 61.42857143 | 149.8452012 | 0.36034 |

| | | | | | | | |
|---|---|---|---|---|---|---|---|
| ABCG1 | chr21 | 43637581 | 2 | 2 | 53.33333333 | 133.3333333 | 0.3604 |
| PHACTR1 | chr6 | 12713997 | 2 | 2 | 53.33333333 | 133.3333333 | 0.3604 |
| F7 | chr13 | 113752693 | 33 | 32 | 3132.773382 | 3080.877045 | 0.36058 |
| APOA1 | chr11 | 116667208 | 33 | 32 | 2929.215721 | 2890.242488 | 0.36082 |
| FGF2 | chr4 | 123747334 | 25 | 25 | 2065.423465 | 2151.433597 | 0.36124 |
| FOXP3 | chrX | 49120166 | 33 | 32 | 2961.474616 | 2936.798116 | 0.36136 |
| ADRA2C | chr4 | 3762479 | 23 | 25 | 2083.042513 | 2348.394024 | 0.36138 |
| ESR2 | chr14 | 64806610 | 2 | 3 | 40 | 160 | 0.36146 |
| AMOTL2 | chr3 | 134100639 | 3 | 2 | 80 | 120 | 0.36146 |
| CDKN1A | chr6 | 36653389 | 32 | 33 | 1794.800757 | 1967.473808 | 0.36174 |
| LDLR | chr19 | 11191777 | 11 | 9 | 704.5054945 | 669.047619 | 0.36188 |
| APOA1 | chr11 | 116682616 | 28 | 27 | 2825 | 2654.393939 | 0.36204 |
| ALOX15 | chr17 | 4545350 | 6 | 9 | 278.3333333 | 333.5714286 | 0.36242 |
| ABCB1 | chr7 | 87133414 | 37 | 32 | 3479.350849 | 2966.023064 | 0.36322 |
| ESR1 | chr6 | 152007779 | 5 | 6 | 500 | 583.3333333 | 0.36322 |
| LDLR | chr19 | 11224235 | 6 | 3 | 533.3333333 | 300 | 0.36322 |
| ATF3 | chr1 | 212745300 | 6 | 9 | 580 | 900 | 0.36322 |
| MCPH1 | chr8 | 6259655 | 6 | 8 | 580 | 800 | 0.36322 |
| MCPH1 | chr8 | 6470660 | 32 | 32 | 3086.693939 | 3028.14114 | 0.36332 |
| ADRA2C | chr4 | 3761193 | 31 | 32 | 2755.900713 | 2932.036857 | 0.36336 |
| ABCG1 | chr21 | 43640107 | 2 | 2 | 45 | 128.5714286 | 0.36348 |
| ABCB1 | chr7 | 87349682 | 2 | 2 | 45 | 128.5714286 | 0.36348 |
| F2RL3 | chr19 | 16998325 | 16 | 10 | 723.6538462 | 391.9047619 | 0.36356 |
| APOA1 | chr11 | 116661489 | 24 | 31 | 1426.526603 | 1715.705035 | 0.36358 |
| F7 | chr13 | 113757953 | 31 | 33 | 2339.124642 | 2589.31225 | 0.36372 |
| ALOX15 | chr17 | 4545374 | 16 | 18 | 723.6263736 | 728.3111333 | 0.3638 |
| FOXP3 | chrX | 49117640 | 33 | 32 | 3083.110365 | 2913.192134 | 0.36424 |
| ABCB1 | chr7 | 87347483 | 32 | 32 | 3009.505588 | 2939.05745 | 0.36478 |
| ADRA2C | chr4 | 3768555 | 8 | 14 | 222.4519633 | 462.687948 | 0.36506 |
| LDLR | chr19 | 11219615 | 34 | 32 | 3097.612836 | 2874.182936 | 0.3652 |
| APOA1 | chr11 | 116697339 | 33 | 32 | 3163.619773 | 3101.36863 | 0.36534 |
| PHACTR1 | chr6 | 12713380 | 37 | 33 | 1896.943998 | 1764.665245 | 0.36542 |
| CDKN2B-AS1 | chr9 | 22003804 | 22 | 26 | 877.0394801 | 928.821742 | 0.3655 |
| FOXP3 | chrX | 49125095 | 33 | 33 | 2070.096256 | 1968.077172 | 0.36558 |
| LDLR | chr19 | 11199425 | 32 | 32 | 2551.768205 | 2626.651402 | 0.36562 |
| CDKN2B-AS1 | chr9 | 21974207 | 10 | 7 | 240.6599318 | 227.974026 | 0.36566 |
| TMPO | chr12 | 98905050 | 37 | 32 | 3664.235082 | 3087.935441 | 0.36598 |
| LDLR | chr19 | 11190238 | 3 | 6 | 280 | 502.3809524 | 0.36604 |
| LDLR | chr19 | 11192504 | 32 | 28 | 2882.709795 | 2456.286924 | 0.36618 |
| NOS3 | chr7 | 150695450 | 21 | 20 | 1502.525253 | 1515.912698 | 0.36628 |
| ARHGEF18 | chr19 | 7469238 | 33 | 32 | 3040.677301 | 2997.620224 | 0.36642 |
| CDKN1A | chr6 | 36653134 | 34 | 32 | 3015.287774 | 2889.997479 | 0.36688 |
| BCL2 | chr18 | 60992886 | 2 | 4 | 185.7142857 | 307.5 | 0.367 |
| BAX | chr19 | 49449607 | 32 | 32 | 2280.456387 | 2372.582328 | 0.36704 |
| CDKN2B-AS1 | chr9 | 21968668 | 12 | 10 | 314.6245318 | 331.9573291 | 0.36718 |
| F2RL3 | chr19 | 16994457 | 13 | 14 | 531.8650794 | 482.481685 | 0.36728 |
| APOA1 | chr11 | 116697082 | 11 | 13 | 876.1904762 | 1118.333333 | 0.36752 |

| | | | | | | | |
|---|---|---|---|---|---|---|---|
| PLA2G7 | chr6 | 46705614 | 12 | 18 | 873.8888889 | 1408.90629 | 0.36806 |
| CDKN2B-AS1 | chr9 | 21990099 | 14 | 14 | 581.5079365 | 509.3362193 | 0.36812 |
| ADRA2C | chr4 | 3768580 | 19 | 22 | 543.6089759 | 692.9635809 | 0.36862 |
| CDKN1A | chr6 | 36652896 | 23 | 25 | 2224.745353 | 2455.277778 | 0.36872 |
| CDKN2B-AS1 | chr9 | 21970440 | 15 | 17 | 459.4641842 | 582.5413475 | 0.36872 |
| LDLR | chr19 | 11190777 | 36 | 32 | 3325.304098 | 2919.832182 | 0.36894 |
| P2RY12 | chr3 | 151106242 | 34 | 32 | 3017.774839 | 2771.913421 | 0.36904 |
| CTGF | chr6 | 132269402 | 40 | 32 | 3509.701448 | 2768.050292 | 0.36928 |
| ATF3 | chr1 | 212742216 | 30 | 31 | 3046.224264 | 3004.653185 | 0.36944 |
| NOS3 | chr7 | 150695176 | 30 | 31 | 2671.888553 | 2610.565575 | 0.36946 |
| F2RL3 | chr19 | 16998135 | 11 | 7 | 347.6984127 | 267.1428571 | 0.3695 |
| ADRA2C | chr4 | 3768575 | 19 | 18 | 555.6883756 | 596.4693674 | 0.36976 |
| CDKN2B-AS1 | chr9 | 21980226 | 36 | 32 | 3401.729587 | 2980.06671 | 0.36986 |
| LCK | chr1 | 32740928 | 22 | 28 | 1827.202812 | 2229.93451 | 0.37018 |
| CDKN2B-AS1 | chr9 | 21990299 | 34 | 32 | 1218.019392 | 1067.341143 | 0.3702 |
| NOS3 | chr7 | 150711105 | 5 | 6 | 128.3838384 | 221.6880342 | 0.37024 |
| F7 | chr13 | 113756313 | 36 | 32 | 2924.276235 | 2546.791411 | 0.37062 |
| NR3C1 | chr5 | 142786406 | 38 | 32 | 3635.352651 | 3091.091008 | 0.37068 |
| FGF2 | chr4 | 123747244 | 34 | 32 | 3175.815111 | 3033.355709 | 0.37086 |
| APOA1 | chr11 | 116660515 | 34 | 32 | 3176.352514 | 2910.302221 | 0.37102 |
| AMOTL2 | chr3 | 134094799 | 37 | 32 | 3479.511368 | 3044.051097 | 0.37112 |
| CDKN2B-AS1 | chr9 | 22005434 | 9 | 4 | 202.6401148 | 161.1627141 | 0.37116 |
| COL15A1 | chr9 | 101705956 | 2 | 5 | 100 | 121.9047619 | 0.3712 |
| TIMP3 | chr22 | 33187651 | 35 | 32 | 3293.055454 | 3045.523747 | 0.37142 |
| LDLR | chr19 | 11224610 | 26 | 30 | 2266.127402 | 2700.572761 | 0.37154 |
| LDLR | chr19 | 11199656 | 34 | 32 | 1355.259731 | 1350.65184 | 0.37164 |
| FOXP3 | chrX | 49117225 | 25 | 26 | 2363.220391 | 2518.841991 | 0.37164 |
| CDKN1A | chr6 | 36651837 | 34 | 32 | 3255.953915 | 3027.042855 | 0.37166 |
| APOA1 | chr11 | 116669309 | 17 | 14 | 1161.428571 | 1025.833333 | 0.37178 |
| ATF3 | chr1 | 212793848 | 2 | 4 | 77.77777777 | 228.8248399 | 0.37214 |
| APOA1 | chr11 | 116664464 | 28 | 30 | 2726.791126 | 2861.487777 | 0.37222 |
| ALOX15 | chr17 | 4546451 | 29 | 31 | 1967.52504 | 1982.088745 | 0.37268 |
| ADRA2C | chr4 | 3758787 | 36 | 32 | 3217.884118 | 2909.042102 | 0.3731 |
| ABCG1 | chr21 | 43630443 | 34 | 33 | 2318.010218 | 2160.854908 | 0.37322 |
| PLA2G7 | chr6 | 46704340 | 32 | 26 | 1538.46086 | 1376.023845 | 0.37326 |
| ALOX15 | chr17 | 4546994 | 19 | 16 | 1552.5222 | 1226.031746 | 0.37334 |
| APOA1 | chr11 | 116711129 | 34 | 32 | 2991.78752 | 2761.524043 | 0.37346 |
| HMGCR | chr5 | 74631770 | 33 | 32 | 3228.903933 | 3096.684234 | 0.3738 |
| CDKN2B-AS1 | chr9 | 22011032 | 20 | 17 | 568.8147919 | 518.1381166 | 0.37382 |
| APOA1 | chr11 | 116718257 | 34 | 32 | 3344.431465 | 3115.699183 | 0.37384 |
| PECAM1 | chr17 | 62399706 | 3 | 3 | 250 | 283.3333333 | 0.3739 |
| PECAM1 | chr17 | 62410251 | 7 | 5 | 700 | 480 | 0.3739 |
| LDLR | chr19 | 11224232 | 5 | 3 | 433.3333333 | 300 | 0.3739 |
| ABCA1 | chr9 | 107693984 | 5 | 5 | 500 | 487.5 | 0.3739 |
| LDLR | chr19 | 11224453 | 36 | 32 | 3433.851334 | 3080.683041 | 0.37438 |
| APOA1 | chr11 | 116704564 | 33 | 32 | 3135.002275 | 3005.062125 | 0.37438 |
| LOXL1 | chr15 | 74215682 | 29 | 30 | 2300.268994 | 2271.738539 | 0.37444 |

| | | | | | | | |
|---|---|---|---|---|---|---|---|
| KLF2 | chr19 | 16433249 | 31 | 30 | 3092.243653 | 2913.65873 | 0.37452 |
| F7 | chr13 | 113750976 | 2 | 2 | 40 | 125 | 0.37464 |
| ALOX15 | chr17 | 4550121 | 2 | 2 | 40 | 125 | 0.37464 |
| NOS3 | chr7 | 150689412 | 2 | 2 | 40 | 125 | 0.37464 |
| CDKN2B-AS1 | chr9 | 21989521 | 15 | 18 | 395.753158 | 553.6389731 | 0.37488 |
| CDKN2B-AS1 | chr9 | 21968592 | 2 | 9 | 40.83333333 | 292.4618438 | 0.37494 |
| ABCA1 | chr9 | 107693931 | 29 | 31 | 2695.067028 | 2741.795977 | 0.37544 |
| CDKN2B-AS1 | chr9 | 22006022 | 3 | 10 | 80.6722689 | 341.1829837 | 0.37544 |
| P2RY12 | chr3 | 151111793 | 37 | 32 | 3573.962733 | 3045.406749 | 0.37548 |
| NOS3 | chr7 | 150691365 | 17 | 16 | 473.9415783 | 509.9039531 | 0.37558 |
| BAX | chr19 | 49454391 | 13 | 13 | 666.7965368 | 591.1111111 | 0.37572 |
| LDLR | chr19 | 11215990 | 31 | 32 | 2964.629438 | 3013.016082 | 0.37582 |
| MIA3 | chr1 | 222792621 | 30 | 32 | 1096.361008 | 1266.336691 | 0.37604 |
| APOA1 | chr11 | 116675100 | 34 | 32 | 2695.911015 | 2445.72505 | 0.3761 |
| ATF3 | chr1 | 212745343 | 28 | 30 | 2924.638695 | 2976.72479 | 0.37614 |
| E2F6 | chr2 | 11611430 | 25 | 29 | 1922.794453 | 2059.82514 | 0.37628 |
| PHACTR1 | chr6 | 12710274 | 38 | 32 | 3570.579405 | 2972.277992 | 0.3763 |
| HMGCR | chr5 | 74628733 | 22 | 25 | 1860.27417 | 2194.821429 | 0.37632 |
| LOXL1 | chr15 | 74215313 | 3 | 5 | 185.8333333 | 260 | 0.37636 |
| ARHGEF18 | chr19 | 7460362 | 5 | 3 | 117.0527634 | 146.1904762 | 0.37636 |
| KLF2 | chr19 | 16435993 | 3 | 3 | 78.46153846 | 153.5714286 | 0.37652 |
| ARHGEF18 | chr19 | 7464035 | 32 | 33 | 2162.231157 | 2340.742282 | 0.37658 |
| ALDOA | chr16 | 30078228 | 26 | 29 | 2115.332632 | 2446.088413 | 0.37672 |
| TMPO | chr12 | 98907356 | 41 | 32 | 3846.193956 | 2974.341717 | 0.37676 |
| CDKN2B-AS1 | chr9 | 22006196 | 10 | 16 | 325.4374266 | 450.1279861 | 0.37676 |
| NOS3 | chr7 | 150678599 | 32 | 32 | 2356.390192 | 2433.426377 | 0.37716 |
| CDKN2B-AS1 | chr9 | 21988080 | 36 | 32 | 2823.959424 | 2569.075667 | 0.37724 |
| CDKN1A | chr6 | 36650930 | 15 | 11 | 371.902072 | 332.7463787 | 0.37756 |
| NOS3 | chr7 | 150682070 | 24 | 25 | 1965.521701 | 2133.784549 | 0.37762 |
| NOS3 | chr7 | 150707755 | 6 | 4 | 152.4314574 | 172.6403743 | 0.37774 |
| ARHGEF18 | chr19 | 7528612 | 12 | 19 | 1160.833333 | 1870.833333 | 0.37804 |
| APOA1 | chr11 | 116692004 | 32 | 32 | 2996.691435 | 2948.713861 | 0.37806 |
| APOA1 | chr11 | 116697533 | 32 | 32 | 2452.234925 | 2544.125331 | 0.37824 |
| ABCA1 | chr9 | 107691567 | 4 | 4 | 93.87057387 | 153.4807105 | 0.37852 |
| NOS3 | chr7 | 150693603 | 5 | 4 | 139.1483517 | 167.2222222 | 0.37862 |
| ESR1 | chr6 | 152003125 | 19 | 17 | 485.952371 | 496.5578841 | 0.37868 |
| GCK | chr7 | 44237498 | 34 | 32 | 3210.427177 | 3056.017011 | 0.37874 |
| CTGF | chr6 | 132272285 | 3 | 2 | 83.88888889 | 120 | 0.37886 |
| LCK | chr1 | 32708244 | 4 | 7 | 95.55555556 | 236.8464869 | 0.37888 |
| NOS3 | chr7 | 150694495 | 27 | 32 | 2345.735196 | 2696.735209 | 0.37894 |
| CDKN2B-AS1 | chr9 | 22005437 | 3 | 5 | 72.06534787 | 186.6170478 | 0.37904 |
| NR3C1 | chr5 | 142789808 | 40 | 33 | 2379.477289 | 2025.164427 | 0.37916 |
| TIMP3 | chr22 | 33188607 | 33 | 32 | 3027.172207 | 2886.001772 | 0.37968 |
| AKT2 | chr19 | 40800022 | 22 | 22 | 2018.888889 | 2070.032468 | 0.3797 |
| F7 | chr13 | 113757875 | 32 | 33 | 2816.685428 | 2817.377629 | 0.37972 |
| LDLR | chr19 | 11219989 | 20 | 24 | 1905.512266 | 2341.428571 | 0.37994 |
| GCK | chr7 | 44236064 | 27 | 29 | 1104.964799 | 1080.396737 | 0.38004 |

| | | | | | | | |
|---|---|---|---|---|---|---|---|
| ESR2 | chr14 | 64806472 | 29 | 25 | 2815.512266 | 2463.141026 | 0.38086 |
| ESR2 | chr14 | 64693598 | 24 | 22 | 2179.062429 | 2051.717033 | 0.3812 |
| PECAM1 | chr17 | 62400576 | 25 | 27 | 1632.65491 | 1601.114223 | 0.38134 |
| LPA | chr6 | 160949907 | 36 | 32 | 3376.180296 | 3029.806891 | 0.38142 |
| LOXL1 | chr15 | 74216953 | 30 | 30 | 1822.931505 | 1941.751086 | 0.38154 |
| PECAM1 | chr17 | 62401696 | 26 | 26 | 2385.20091 | 2437.378732 | 0.38168 |
| CDKN2B-AS1 | chr9 | 21968330 | 4 | 3 | 93.33333333 | 153.3333333 | 0.38172 |
| TIMP3 | chr22 | 33191423 | 38 | 32 | 3458.825514 | 2951.169059 | 0.38176 |
| ICAM2 | chr17 | 62085237 | 29 | 29 | 2717.536075 | 2653.956321 | 0.38182 |
| NOS3 | chr7 | 150710222 | 4 | 9 | 114.6969697 | 334.4047619 | 0.38182 |
| CDKN2B-AS1 | chr9 | 21974181 | 4 | 3 | 132.2431078 | 146.1538462 | 0.38192 |
| ADRA2C | chr4 | 3766322 | 32 | 32 | 2789.46174 | 2720.943569 | 0.38196 |
| APOA1 | chr11 | 116692030 | 31 | 32 | 2875.64679 | 2914.995212 | 0.38226 |
| CDKN2B-AS1 | chr9 | 21996655 | 22 | 23 | 622.3490298 | 727.1451691 | 0.38264 |
| LDLR | chr19 | 11218358 | 11 | 8 | 1078.888889 | 745.7142857 | 0.38266 |
| LDLR | chr19 | 11191293 | 25 | 28 | 2236.881313 | 2426.407204 | 0.38278 |
| CDKN2B-AS1 | chr9 | 21974451 | 2 | 3 | 40 | 146.0526316 | 0.38278 |
| LDLR | chr19 | 11217971 | 29 | 30 | 2717.340743 | 2874.225872 | 0.38288 |
| APOA1 | chr11 | 116697305 | 32 | 32 | 3091.137386 | 3057.989729 | 0.38294 |
| LPA | chr6 | 160952342 | 15 | 17 | 1045.992063 | 1282.142857 | 0.3835 |
| AMOTL2 | chr3 | 134094122 | 11 | 7 | 281.6896389 | 233.0707196 | 0.38352 |
| CDKN2B-AS1 | chr9 | 21970980 | 17 | 16 | 591.4071463 | 505.3439616 | 0.38358 |
| PECAM1 | chr17 | 62401201 | 32 | 32 | 2803.855147 | 2881.136965 | 0.38368 |
| TIMP3 | chr22 | 33188718 | 26 | 27 | 1686.78869 | 1645.73496 | 0.3838 |
| APOA1 | chr11 | 116695295 | 23 | 27 | 2187.380952 | 2518.722944 | 0.3839 |
| ESR1 | chr6 | 152006547 | 10 | 9 | 977.3809524 | 857.1428571 | 0.3841 |
| NOS3 | chr7 | 150691532 | 3 | 3 | 63.65079365 | 136.1616162 | 0.3841 |
| APOA1 | chr11 | 116661330 | 31 | 33 | 2192.654237 | 2416.437842 | 0.38414 |
| NOS3 | chr7 | 150709081 | 37 | 32 | 3214.44211 | 2726.929754 | 0.38422 |
| GALNT2 | chr1 | 230417999 | 17 | 20 | 942.5194916 | 1227.41453 | 0.38438 |
| NOS3 | chr7 | 150709463 | 7 | 6 | 643.4920635 | 580 | 0.38498 |
| APOA1 | chr11 | 116661563 | 16 | 20 | 480.2627846 | 662.7723665 | 0.38502 |
| ABCA1 | chr9 | 107691555 | 7 | 3 | 174.981685 | 155.5555556 | 0.3851 |
| TERT | chr5 | 1297047 | 6 | 9 | 583.3333333 | 844.6428571 | 0.38524 |
| GALNT2 | chr1 | 230186080 | 37 | 32 | 3021.31606 | 2668.472327 | 0.38536 |
| AMOTL2 | chr3 | 134097170 | 32 | 33 | 2548.352653 | 2549.362807 | 0.38556 |
| TIMP3 | chr22 | 33187069 | 38 | 32 | 2748.663209 | 2367.811054 | 0.38574 |
| P2RY12 | chr3 | 151109162 | 37 | 33 | 1961.672077 | 1830.425284 | 0.3861 |
| F7 | chr13 | 113759753 | 25 | 31 | 1909.427661 | 2205.014461 | 0.38648 |
| ABCA1 | chr9 | 107698353 | 18 | 15 | 1491.180223 | 1310 | 0.38648 |
| FOXP3 | chrX | 49114510 | 30 | 31 | 1953.581228 | 2128.26715 | 0.38654 |
| FOXP3 | chrX | 49126956 | 17 | 11 | 1091.962482 | 636.984127 | 0.38656 |
| LDLR | chr19 | 11222344 | 19 | 25 | 1804.603175 | 2280.634921 | 0.38682 |
| CDKN2B-AS1 | chr9 | 22011208 | 7 | 5 | 178.8553781 | 187.6658964 | 0.38694 |
| GCK | chr7 | 44236261 | 2 | 3 | 43.07692308 | 142.7272727 | 0.38714 |
| ESR2 | chr14 | 64805919 | 7 | 7 | 177.2544123 | 237.5 | 0.38718 |
| F2RL3 | chr19 | 16995072 | 6 | 13 | 492.2222222 | 1225.555556 | 0.38718 |

| | | | | | | | |
|---|---|---|---|---|---|---|---|
| CTGF | chr6 | 132279840 | 31 | 27 | 2116.875658 | 1699.728747 | 0.38726 |
| ESR1 | chr6 | 152002912 | 2 | 3 | 40 | 147.3214286 | 0.3873 |
| TERT | chr5 | 1295910 | 30 | 30 | 2180.436143 | 2088.318222 | 0.3874 |
| ESR1 | chr6 | 152007602 | 7 | 11 | 623.6507937 | 908.0952381 | 0.38754 |
| TIMP3 | chr22 | 33191589 | 35 | 32 | 3125.293152 | 2916.043858 | 0.38772 |
| APOA1 | chr11 | 116706234 | 4 | 6 | 109.981685 | 236.3529388 | 0.38772 |
| APOA1 | chr11 | 116699998 | 6 | 5 | 463.5606061 | 340 | 0.38776 |
| TIMP3 | chr22 | 33191471 | 38 | 33 | 2847.926147 | 2384.11198 | 0.388 |
| TERT | chr5 | 1300394 | 28 | 27 | 2571.590237 | 2423.483811 | 0.38816 |
| TIMP3 | chr22 | 33197249 | 14 | 17 | 1181.593407 | 1521.666667 | 0.38824 |
| AKT2 | chr19 | 40792018 | 32 | 32 | 2560.490903 | 2662.900287 | 0.38836 |
| CDKN2B-AS1 | chr9 | 22004698 | 36 | 33 | 2168.001255 | 1908.657324 | 0.38836 |
| PHACTR1 | chr6 | 12716111 | 32 | 33 | 2278.430283 | 2260.978328 | 0.3885 |
| NR3C1 | chr5 | 142785578 | 35 | 32 | 3279.427531 | 2959.922483 | 0.389 |
| LPA | chr6 | 11223915 | 32 | 32 | 3107.5022 | 3134.95671 | 0.389 |
| NOS3 | chr7 | 150695776 | 4 | 2 | 106.3492063 | 65 | 0.389 |
| TIMP3 | chr22 | 33196627 | 33 | 32 | 2863.488099 | 2832.045368 | 0.38904 |
| ALOX15 | chr17 | 4552440 | 32 | 32 | 2192.333928 | 2106.48344 | 0.38906 |
| CDKN2B-AS1 | chr9 | 21988910 | 12 | 9 | 304.7683331 | 289.1646539 | 0.38928 |
| CDKN2B-AS1 | chr9 | 21971157 | 13 | 17 | 383.9255189 | 556.8129093 | 0.38936 |
| HMGCR | chr5 | 74628770 | 4 | 7 | 373.2142857 | 592.2222222 | 0.38964 |
| FGF2 | chr4 | 123741620 | 27 | 28 | 2316.845057 | 2481.690352 | 0.38974 |
| PLA2G7 | chr6 | 46703978 | 2 | 2 | 50 | 128.5714286 | 0.38976 |
| CDKN2B-AS1 | chr9 | 22005420 | 18 | 15 | 484.8984536 | 461.0924171 | 0.38998 |
| APOA1 | chr11 | 116696741 | 36 | 33 | 2853.467779 | 2523.58174 | 0.39 |
| TIMP3 | chr22 | 33191408 | 39 | 32 | 3600.12872 | 2917.468774 | 0.39018 |
| LDLR | chr19 | 11221413 | 33 | 32 | 3171.735518 | 3033.75099 | 0.39026 |
| CDKN2B-AS1 | chr9 | 22006092 | 2 | 5 | 40 | 218.2748538 | 0.39086 |
| LDLR | chr19 | 11197397 | 3 | 4 | 300 | 383.3333333 | 0.391 |
| LPA | chr6 | 160943860 | 4 | 5 | 383.3333333 | 500 | 0.391 |
| LDLR | chr19 | 11197631 | 4 | 3 | 380 | 300 | 0.391 |
| E2F6 | chr2 | 11615687 | 5 | 4 | 500 | 360 | 0.391 |
| AMOTL2 | chr3 | 134102034 | 4 | 4 | 400 | 380 | 0.391 |
| NOS3 | chr7 | 150708903 | 5 | 6 | 488.8888889 | 600 | 0.391 |
| ADRA2C | chr4 | 3761304 | 29 | 32 | 2498.950927 | 2850.330981 | 0.3911 |
| LPA | chr6 | 11217463 | 34 | 33 | 2611.298793 | 2440.023413 | 0.39148 |
| CDKN2B-AS1 | chr9 | 21976106 | 14 | 10 | 370.8563822 | 334.1339869 | 0.3916 |
| E2F6 | chr2 | 11614670 | 34 | 32 | 3055.997617 | 2767.693424 | 0.3918 |
| ABCG1 | chr21 | 43717743 | 38 | 33 | 3423.267808 | 2895.374256 | 0.39194 |
| ATF3 | chr1 | 212741376 | 34 | 32 | 3075.966909 | 2948.717475 | 0.392 |
| AKT2 | chr19 | 40800840 | 35 | 32 | 3331.919971 | 3087.244187 | 0.39208 |
| ESR1 | chr6 | 152005487 | 34 | 33 | 2750.063164 | 2599.456854 | 0.3923 |
| APOA1 | chr11 | 116668919 | 31 | 31 | 2985.146657 | 2901.192451 | 0.3924 |
| LPA | chr6 | 11224767 | 10 | 11 | 639.1269841 | 782.2222222 | 0.39244 |
| F2RL3 | chr19 | 16994186 | 33 | 32 | 3148.271826 | 3010.943614 | 0.39256 |
| ALDOA | chr16 | 30058255 | 32 | 31 | 3114.380814 | 2982.157477 | 0.39306 |
| APOA1 | chr11 | 116696914 | 33 | 32 | 2288.636074 | 2297.401195 | 0.39318 |

| | | | | | | | |
|---|---|---|---|---|---|---|---|
| P2RY12 | chr3 | 151105724 | 30 | 32 | 1993.610297 | 2004.851875 | 0.3932 |
| F7 | chr13 | 113775487 | 31 | 32 | 2812.956111 | 2842.239829 | 0.39352 |
| KLF2 | chr19 | 16437944 | 27 | 30 | 1126.325208 | 1342.99511 | 0.39374 |
| PECAM1 | chr17 | 62401856 | 25 | 27 | 2140.179739 | 2407.49334 | 0.3938 |
| ATF3 | chr1 | 212746380 | 22 | 26 | 2137.140637 | 2557.936508 | 0.39396 |
| F7 | chr13 | 113751103 | 9 | 8 | 866.6666667 | 720 | 0.39402 |
| MIA3 | chr1 | 222792560 | 33 | 32 | 2200.289072 | 2054.110893 | 0.39404 |
| APOA1 | chr11 | 116719020 | 20 | 16 | 1953.353175 | 1513.492063 | 0.39416 |
| PECAM1 | chr17 | 62399152 | 23 | 25 | 2016.231387 | 2259.261156 | 0.3942 |
| NR3C1 | chr5 | 142786449 | 38 | 32 | 3649.022307 | 3102.430435 | 0.39422 |
| LPA | chr6 | 11217843 | 4 | 5 | 105.0793651 | 213.5294118 | 0.3944 |
| ABCA1 | chr9 | 107692896 | 18 | 12 | 1577.432845 | 1107.539683 | 0.39468 |
| LOXL1 | chr15 | 74218735 | 16 | 22 | 516.9387475 | 782.8730991 | 0.39478 |
| PHACTR1 | chr6 | 12717926 | 39 | 32 | 3546.747794 | 2949.730791 | 0.39492 |
| TIMP3 | chr22 | 33195271 | 39 | 32 | 3395.620768 | 2724.617664 | 0.39516 |
| APOA1 | chr11 | 116711396 | 34 | 32 | 2899.287081 | 2798.801613 | 0.39526 |
| LDLR | chr19 | 11217528 | 20 | 15 | 1681.218226 | 1304.166667 | 0.39532 |
| APOA1 | chr11 | 116661516 | 28 | 31 | 2289.138953 | 2622.306044 | 0.39562 |
| APOA1 | chr11 | 116704629 | 35 | 32 | 3385.651701 | 3063.776413 | 0.39572 |
| NOS3 | chr7 | 150695433 | 16 | 11 | 983.7301587 | 736.1904762 | 0.39578 |
| LDLR | chr19 | 11217649 | 7 | 4 | 675 | 369.047619 | 0.3958 |
| CDKN2B-AS1 | chr9 | 21994155 | 12 | 9 | 305.4854242 | 284.9814618 | 0.3959 |
| NOS3 | chr7 | 150706577 | 21 | 22 | 1859.059829 | 1826.017316 | 0.39598 |
| APOA1 | chr11 | 116698665 | 32 | 33 | 2768.086407 | 2770.824275 | 0.39616 |
| APOA1 | chr11 | 116714576 | 30 | 31 | 2661.835751 | 2807.239089 | 0.39622 |
| ALOX15 | chr17 | 4548750 | 32 | 33 | 2531.358892 | 2532.606446 | 0.39634 |
| LDLR | chr19 | 11192082 | 19 | 21 | 1834.123377 | 1979.722222 | 0.39654 |
| APOA1 | chr11 | 116718129 | 28 | 26 | 2565.543312 | 2438.400766 | 0.3966 |
| APOA1 | chr11 | 116691150 | 33 | 32 | 3156.160541 | 3023.306098 | 0.39664 |
| GCK | chr7 | 44229915 | 34 | 32 | 3315.194086 | 3020.440039 | 0.39676 |
| LOXL1 | chr15 | 74214857 | 7 | 11 | 193.5714286 | 378.5714286 | 0.39682 |
| FOXP3 | chrX | 49125616 | 3 | 2 | 106.6666667 | 126.6666667 | 0.39684 |
| LCK | chr1 | 32740942 | 27 | 28 | 2376.769397 | 2529.709873 | 0.39686 |
| FGF2 | chr4 | 123740072 | 34 | 32 | 3082.675419 | 2833.09191 | 0.39688 |
| TERT | chr5 | 1298892 | 30 | 29 | 2773.601265 | 2626.439133 | 0.39702 |
| NOS3 | chr7 | 150708907 | 10 | 9 | 947.5757576 | 820 | 0.39712 |
| TFPI | chr2 | 188425245 | 25 | 27 | 2296.482295 | 2541.123321 | 0.39714 |
| P2RY12 | chr3 | 151102880 | 35 | 31 | 1485.420165 | 1241.821518 | 0.39722 |
| ARHGEF18 | chr19 | 7463783 | 7 | 9 | 180.7877191 | 296.9108947 | 0.39732 |
| FOXP3 | chrX | 49126840 | 7 | 4 | 253.8888889 | 107.2222222 | 0.3974 |
| FOXP3 | chrX | 49126811 | 8 | 3 | 217.1428571 | 160 | 0.39746 |
| ABCG1 | chr21 | 43634716 | 6 | 4 | 563.3333333 | 346.6666667 | 0.39754 |
| CDKN2B-AS1 | chr9 | 21968584 | 3 | 3 | 61.05263158 | 121.8462824 | 0.39756 |
| AMOTL2 | chr3 | 134100452 | 19 | 17 | 1746.302309 | 1480.858586 | 0.39768 |
| F7 | chr13 | 113750930 | 31 | 30 | 2818.019115 | 2611.040955 | 0.39772 |
| CDKN2B-AS1 | chr9 | 22005127 | 3 | 3 | 60 | 160 | 0.39774 |
| APOA1 | chr11 | 116697066 | 3 | 6 | 260 | 420.8333333 | 0.39782 |

| | | | | | | | |
|---|---|---|---|---|---|---|---|
| LOXL1 | chr15 | 74218979 | 26 | 28 | 998.5176307 | 1008.295383 | 0.39792 |
| ALOX15 | chr17 | 4549926 | 3 | 2 | 61.42857143 | 133.3333333 | 0.39798 |
| AKT2 | chr19 | 40792691 | 40 | 32 | 3764.415672 | 3038.260684 | 0.398 |
| LCK | chr1 | 32742037 | 29 | 27 | 2254.312452 | 2189.284882 | 0.39802 |
| PECAM1 | chr17 | 62400560 | 30 | 32 | 2598.942022 | 2669.707529 | 0.39814 |
| CTGF | chr6 | 132272610 | 6 | 5 | 146.7231599 | 175.7070707 | 0.3982 |
| ATF3 | chr1 | 212783896 | 31 | 32 | 2858.109394 | 2890.097331 | 0.39828 |
| LCK | chr1 | 32741891 | 17 | 22 | 1406.699134 | 1929.779942 | 0.39828 |
| CDKN2B-AS1 | chr9 | 21989951 | 6 | 8 | 145.0793651 | 226.9047619 | 0.39848 |
| MCPH1 | chr8 | 6470209 | 17 | 20 | 1654.880952 | 1871.944444 | 0.39862 |
| FGF2 | chr4 | 123819585 | 33 | 32 | 3202.590356 | 3010.819991 | 0.39868 |
| TIMP3 | chr22 | 33189012 | 2 | 2 | 45 | 125 | 0.39868 |
| ICAM2 | chr17 | 62086894 | 26 | 25 | 1154.751331 | 1018.995106 | 0.39874 |
| LDLR | chr19 | 11197902 | 11 | 13 | 969.5436508 | 1226.825397 | 0.39874 |
| APOA1 | chr11 | 116705566 | 25 | 26 | 2186.260504 | 2347.147852 | 0.39886 |
| APOA1 | chr11 | 116682420 | 15 | 11 | 1424.047619 | 1070.833333 | 0.3989 |
| PHACTR1 | chr6 | 12711588 | 34 | 32 | 2883.694322 | 2661.46112 | 0.39892 |
| GALNT2 | chr1 | 230187440 | 38 | 32 | 3638.484905 | 3098.75914 | 0.399 |
| LDLR | chr19 | 11192549 | 30 | 27 | 2829.420996 | 2452.320555 | 0.39912 |
| ESR1 | chr6 | 152002543 | 10 | 10 | 686.1904762 | 765 | 0.39952 |
| APOA1 | chr11 | 116697216 | 32 | 32 | 2842.625931 | 2896.220071 | 0.39954 |
| APOA1 | chr11 | 116706322 | 2 | 2 | 41.42857143 | 133.3333333 | 0.39958 |
| LDLR | chr19 | 11223988 | 25 | 24 | 2367.863248 | 2198.253968 | 0.39966 |
| ESR1 | chr6 | 152007096 | 10 | 9 | 891.547619 | 843.3333333 | 0.39968 |
| CDKN2B-AS1 | chr9 | 22001775 | 28 | 16 | 786.1508391 | 495.873158 | 0.39982 |
| LPA | chr6 | 160944745 | 36 | 32 | 3345.468956 | 2942.553122 | 0.4 |
| CTGF | chr6 | 132281945 | 30 | 29 | 2544.006161 | 2364.369519 | 0.40034 |
| FOXP3 | chrX | 49125696 | 2 | 2 | 54.76190476 | 130 | 0.40038 |
| PHACTR1 | chr6 | 12717097 | 30 | 32 | 2634.705222 | 2733.648753 | 0.4004 |
| MCPH1 | chr8 | 6256457 | 33 | 32 | 2882.642454 | 2840.406923 | 0.40058 |
| E2F6 | chr2 | 11614045 | 34 | 32 | 3098.577637 | 2806.752349 | 0.40064 |
| LDLR | chr19 | 11198943 | 24 | 29 | 2355.984848 | 2813.619714 | 0.40074 |
| ATF3 | chr1 | 212745353 | 31 | 32 | 3141.171159 | 3133.514356 | 0.40082 |
| LOXL1 | chr15 | 74218870 | 28 | 28 | 1030.194898 | 946.173799 | 0.40084 |
| APOA1 | chr11 | 116662013 | 20 | 24 | 606.7593686 | 802.7734949 | 0.40084 |
| F7 | chr13 | 113750705 | 28 | 24 | 2504.587912 | 2207.25469 | 0.4009 |
| ABCG1 | chr21 | 43630410 | 32 | 33 | 2640.578375 | 2643.096514 | 0.40094 |
| LCK | chr1 | 32741202 | 14 | 11 | 446.3095238 | 402.4603175 | 0.40108 |
| ALDOA | chr16 | 30077751 | 2 | 2 | 40 | 122.2222222 | 0.40128 |
| HMGCR | chr5 | 74627145 | 2 | 2 | 40 | 122.2222222 | 0.40128 |
| CTGF | chr6 | 132272329 | 2 | 2 | 40 | 122.2222222 | 0.40128 |
| CDKN2B-AS1 | chr9 | 21973957 | 14 | 4 | 368.0486964 | 167.5213675 | 0.40132 |
| ESR1 | chr6 | 152002206 | 30 | 32 | 2707.464054 | 2831.711516 | 0.40138 |
| CDKN2B-AS1 | chr9 | 21970784 | 19 | 14 | 532.1975086 | 448.3501115 | 0.4015 |
| NOS3 | chr7 | 150678987 | 32 | 32 | 2720.513962 | 2777.082399 | 0.40156 |
| NOS3 | chr7 | 150710302 | 32 | 32 | 2387.931081 | 2305.584488 | 0.40176 |
| NOS3 | chr7 | 150711516 | 17 | 17 | 519.4801685 | 582.0349095 | 0.40186 |

| | | | | | | | |
|---|---|---|---|---|---|---|---|
| LCK | chr1 | 32741220 | 5 | 5 | 161.9047619 | 208.8888889 | 0.40192 |
| LPA | chr6 | 160950882 | 3 | 2 | 75.55555555 | 40 | 0.40196 |
| NOS3 | chr7 | 150679555 | 24 | 22 | 1013.260455 | 1024.475247 | 0.402 |
| TERT | chr5 | 1253671 | 32 | 32 | 2929.01431 | 3004.829293 | 0.40214 |
| ALOX15 | chr17 | 4549872 | 2 | 2 | 42.22222222 | 133.3333333 | 0.40232 |
| ESR1 | chr6 | 152424350 | 34 | 32 | 2942.086771 | 2815.651864 | 0.40272 |
| BAX | chr19 | 49465449 | 7 | 6 | 201.4896215 | 242.5198413 | 0.40276 |
| LCK | chr1 | 32716557 | 31 | 32 | 2022.709205 | 2199.943723 | 0.40304 |
| FGF2 | chr4 | 123742021 | 27 | 31 | 2586.378355 | 2913.737079 | 0.40326 |
| FOXP3 | chrX | 49121056 | 3 | 4 | 248.4848485 | 292.3809524 | 0.40372 |
| ESR1 | chr6 | 152007456 | 35 | 32 | 3372.022028 | 3047.305327 | 0.40394 |
| ADRA2C | chr4 | 3768621 | 22 | 28 | 794.7191185 | 935.4617027 | 0.404 |
| NOS3 | chr7 | 150692061 | 5 | 3 | 151.6666667 | 165.0793651 | 0.40402 |
| GALNT2 | chr1 | 230193574 | 33 | 32 | 1830.182904 | 1677.361785 | 0.40424 |
| LCK | chr1 | 32713241 | 32 | 32 | 3014.498441 | 2972.053011 | 0.40424 |
| LDLR | chr19 | 11216203 | 18 | 16 | 1641.765873 | 1501.388889 | 0.40466 |
| NOS3 | chr7 | 150688166 | 34 | 32 | 2877.178563 | 2777.320428 | 0.4047 |
| F2RL3 | chr19 | 16998718 | 10 | 15 | 291.8253968 | 505.1046176 | 0.4048 |
| FOXP3 | chrX | 49121178 | 11 | 7 | 842.0995671 | 471.6666667 | 0.40484 |
| CDKN2B-AS1 | chr9 | 21967881 | 3 | 3 | 72.24736048 | 150.6060606 | 0.40484 |
| E2F6 | chr2 | 11611966 | 34 | 32 | 3190.417977 | 3053.806292 | 0.40536 |
| CTGF | chr6 | 132278759 | 39 | 32 | 3767.41079 | 3067.27428 | 0.40538 |
| LOXL1 | chr15 | 74210440 | 27 | 29 | 2556.031746 | 2809.304029 | 0.40566 |
| E2F6 | chr2 | 11614337 | 37 | 32 | 3528.79714 | 3010.583532 | 0.40608 |
| LDLR | chr19 | 11224367 | 33 | 32 | 3183.290427 | 3119.994474 | 0.40636 |
| APOA1 | chr11 | 116716921 | 30 | 32 | 2755.383182 | 2994.427503 | 0.40646 |
| APOA1 | chr11 | 116717363 | 26 | 26 | 2295.790043 | 2207.00161 | 0.40658 |
| TERT | chr5 | 1300217 | 31 | 33 | 2225.542394 | 2457.090644 | 0.40708 |
| CDKN1A | chr6 | 36645886 | 2 | 6 | 45 | 250.4906205 | 0.40714 |
| LCK | chr1 | 32714495 | 32 | 33 | 1894.163745 | 2035.139542 | 0.40716 |
| BAX | chr19 | 49465140 | 32 | 32 | 3062.376353 | 3109.634733 | 0.40732 |
| NOS3 | chr7 | 150711058 | 2 | 2 | 50.76923077 | 127.2727273 | 0.40734 |
| LCK | chr1 | 32716817 | 2 | 3 | 42.22222222 | 144.3627451 | 0.40762 |
| ADRA2C | chr4 | 3763568 | 31 | 30 | 2429.056791 | 2424.642931 | 0.40764 |
| HMGCR | chr5 | 74623824 | 15 | 18 | 1381.428571 | 1700.436508 | 0.40764 |
| CDKN2B-AS1 | chr9 | 21977661 | 14 | 10 | 441.9047619 | 368.968254 | 0.40764 |
| CDKN2B-AS1 | chr9 | 21987951 | 16 | 11 | 418.1681463 | 341.6388175 | 0.40776 |
| ABCA1 | chr9 | 107692085 | 25 | 30 | 2389.808802 | 2926.432612 | 0.4079 |
| CDKN2B-AS1 | chr9 | 21990357 | 16 | 11 | 452.8306304 | 365.9704082 | 0.408 |
| APOA1 | chr11 | 116710352 | 39 | 32 | 3430.035559 | 2755.748523 | 0.4081 |
| LDLR | chr19 | 11196231 | 35 | 32 | 2804.345527 | 2491.986021 | 0.40842 |
| FOXP3 | chrX | 49121787 | 16 | 11 | 1490.521978 | 1060 | 0.4087 |
| PLA2G7 | chr6 | 46703124 | 5 | 2 | 110 | 133.3333333 | 0.4087 |
| NOS3 | chr7 | 150678962 | 33 | 32 | 2631.132505 | 2605.696475 | 0.40872 |
| CDKN2B-AS1 | chr9 | 21970802 | 29 | 29 | 1090.822495 | 1018.549212 | 0.40872 |
| GCK | chr7 | 44228636 | 30 | 30 | 2621.449689 | 2694.04762 | 0.40876 |
| F7 | chr13 | 113750925 | 31 | 31 | 3060.592006 | 2942.168803 | 0.40886 |

| | | | | | | | |
|---|---|---|---|---|---|---|---|
| MCPH1 | chr8 | 6260367 | 32 | 32 | 3014.419581 | 2969.289211 | 0.4095 |
| NOS3 | chr7 | 150705760 | 34 | 33 | 2063.997861 | 2105.420673 | 0.40956 |
| MCPH1 | chr8 | 6470761 | 33 | 32 | 3044.721624 | 3007.795271 | 0.41026 |
| CDKN1A | chr6 | 36645335 | 36 | 32 | 3370.124841 | 3031.593796 | 0.4103 |
| APOA1 | chr11 | 116699947 | 3 | 3 | 238.8888889 | 283.3333333 | 0.41072 |
| ADRA2C | chr4 | 3766664 | 20 | 20 | 1771.321195 | 1696.633367 | 0.4111 |
| LCK | chr1 | 32707303 | 11 | 9 | 289.9609155 | 301.4112669 | 0.41124 |
| CDKN2B-AS1 | chr9 | 21979543 | 8 | 7 | 450.4761905 | 320 | 0.41126 |
| F2RL3 | chr19 | 16994412 | 3 | 3 | 65 | 145 | 0.41182 |
| LDLR | chr19 | 11222567 | 32 | 32 | 2983.716461 | 3037.462135 | 0.4121 |
| F7 | chr13 | 113751963 | 34 | 33 | 2739.304727 | 2578.499064 | 0.41224 |
| TIMP3 | chr22 | 33186867 | 36 | 32 | 3360.989201 | 3012.911009 | 0.41232 |
| PLA2G7 | chr6 | 46703133 | 3 | 4 | 65 | 162.2222222 | 0.41238 |
| TERT | chr5 | 1298762 | 4 | 7 | 352.3809524 | 553.3333333 | 0.41268 |
| KLF2 | chr19 | 16437849 | 27 | 26 | 1108.48089 | 1165.871597 | 0.4129 |
| LDLR | chr19 | 11192251 | 26 | 19 | 2427.94511 | 1725.400623 | 0.41352 |
| CDKN2B-AS1 | chr9 | 21968870 | 2 | 3 | 40.28985507 | 137.6221001 | 0.41374 |
| PHACTR1 | chr6 | 12717384 | 33 | 33 | 2628.132619 | 2559.523108 | 0.41394 |
| NOS3 | chr7 | 150686735 | 34 | 32 | 3210.482609 | 2980.639998 | 0.41408 |
| CDKN2B-AS1 | chr9 | 22009471 | 2 | 2 | 44.44444444 | 123.0769231 | 0.4142 |
| APOA1 | chr11 | 116696356 | 35 | 32 | 3010.54159 | 2802.878909 | 0.41432 |
| APOA1 | chr11 | 116661408 | 32 | 32 | 2768.560016 | 2838.277512 | 0.41456 |
| F7 | chr13 | 113760547 | 21 | 21 | 1847.630719 | 1758.809524 | 0.41484 |
| ESR1 | chr6 | 152002033 | 28 | 29 | 2616.095571 | 2775.530303 | 0.41486 |
| LPA | chr6 | 11220557 | 34 | 32 | 3007.11604 | 2927.111887 | 0.41492 |
| CDKN2B-AS1 | chr9 | 22005217 | 18 | 17 | 493.323557 | 516.7195936 | 0.41514 |
| CDKN2B-AS1 | chr9 | 21976490 | 5 | 3 | 196.9047619 | 91.42857143 | 0.41516 |
| ALOX15 | chr17 | 4554187 | 38 | 32 | 3457.027984 | 2878.823972 | 0.41532 |
| LCK | chr1 | 32741224 | 3 | 4 | 95.23809523 | 183.5714286 | 0.41554 |
| GCK | chr7 | 44183433 | 37 | 32 | 2716.333154 | 2487.354967 | 0.41578 |
| LDLR | chr19 | 11197595 | 10 | 5 | 901.6666667 | 483.3333333 | 0.4158 |
| ESR1 | chr6 | 152002461 | 33 | 33 | 2232.779283 | 2325.631251 | 0.41588 |
| PECAM1 | chr17 | 62398713 | 2 | 2 | 40 | 120.6896552 | 0.41602 |
| ATF3 | chr1 | 212780557 | 2 | 2 | 43.07692308 | 122.2222222 | 0.41624 |
| CDKN2B-AS1 | chr9 | 21990196 | 19 | 28 | 688.3704042 | 1109.014524 | 0.41628 |
| P2RY12 | chr3 | 151108101 | 35 | 33 | 2684.889099 | 2456.059693 | 0.4164 |
| F7 | chr13 | 113753435 | 32 | 32 | 3082.741943 | 3022.968404 | 0.41642 |
| LOXL1 | chr15 | 74210924 | 3 | 2 | 93.05194805 | 120 | 0.41646 |
| AMOTL2 | chr3 | 134097633 | 37 | 32 | 3251.525422 | 2773.098406 | 0.41648 |
| LPA | chr6 | 11224158 | 32 | 32 | 2984.392765 | 2929.495664 | 0.4165 |
| BAX | chr19 | 49456810 | 30 | 31 | 2879.822983 | 2935.04237 | 0.4167 |
| ATF3 | chr1 | 212783857 | 32 | 32 | 3015.380023 | 3047.969434 | 0.41686 |
| LOXL1 | chr15 | 74210565 | 33 | 32 | 3085.347988 | 3030.717571 | 0.41704 |
| CDKN2B-AS1 | chr9 | 21975899 | 4 | 3 | 119.4047619 | 162.8571429 | 0.4171 |
| TERT | chr5 | 1296744 | 5 | 9 | 131.3997114 | 284 | 0.41738 |
| TIMP3 | chr22 | 33188779 | 33 | 31 | 2711.752388 | 2473.446788 | 0.41758 |
| ADRA2C | chr4 | 3767248 | 3 | 3 | 85.07936508 | 154.3859649 | 0.41758 |

| | | | | | | | |
|---|---|---|---|---|---|---|---|
| AMOTL2 | chr3 | 134094090 | 19 | 16 | 557.049719 | 528.5304017 | 0.41766 |
| NOS3 | chr7 | 150711018 | 2 | 2 | 45 | 123.0769231 | 0.41774 |
| ICAM2 | chr17 | 62080240 | 25 | 26 | 2435.363248 | 2559.404762 | 0.41778 |
| KLF2 | chr19 | 16433838 | 2 | 2 | 48.57142857 | 125 | 0.41788 |
| BAX | chr19 | 49449766 | 2 | 2 | 48.57142857 | 125 | 0.41788 |
| APOA1 | chr11 | 116672927 | 18 | 28 | 1753.663004 | 2681.666667 | 0.4179 |
| NOS3 | chr7 | 150710985 | 3 | 3 | 81.05263158 | 150.7936508 | 0.41798 |
| APOA1 | chr11 | 116694203 | 10 | 11 | 953.6630037 | 1003.809524 | 0.41812 |
| MIA3 | chr1 | 222816958 | 19 | 25 | 944.5238095 | 1137.911255 | 0.41818 |
| COL15A1 | chr9 | 101832641 | 33 | 32 | 3123.112752 | 3065.720894 | 0.4182 |
| APOA1 | chr11 | 116672184 | 34 | 32 | 3099.636572 | 2956.387196 | 0.41842 |
| ADRA2C | chr4 | 3764982 | 3 | 4 | 73.33333333 | 176.6483517 | 0.41854 |
| P2RY12 | chr3 | 151105615 | 36 | 33 | 3113.7927 | 2776.162525 | 0.41928 |
| CDKN1A | chr6 | 36654848 | 33 | 32 | 2979.469747 | 2942.684132 | 0.41998 |
| ADRA2C | chr4 | 3766477 | 34 | 32 | 2881.95744 | 2795.29664 | 0.42026 |
| NOS3 | chr7 | 150695330 | 32 | 31 | 2621.514405 | 2616.28236 | 0.42026 |
| LOXL1 | chr15 | 74214820 | 15 | 21 | 598.4279609 | 914.1254579 | 0.42028 |
| CDKN2B-AS1 | chr9 | 22005318 | 3 | 2 | 86.02484472 | 106.6666667 | 0.42042 |
| CDKN1A | chr6 | 36651402 | 25 | 27 | 1816.113486 | 2042.878788 | 0.42076 |
| COL15A1 | chr9 | 101701848 | 33 | 32 | 2782.707895 | 2756.974143 | 0.42096 |
| BAX | chr19 | 49450774 | 13 | 16 | 1233.170996 | 1479.603175 | 0.42122 |
| F7 | chr13 | 113760227 | 30 | 25 | 2229.127169 | 1734.368263 | 0.42126 |
| BCL2 | chr18 | 60993449 | 36 | 32 | 3287.149144 | 2952.519557 | 0.42162 |
| E2F6 | chr2 | 11607954 | 33 | 32 | 3161.420048 | 3002.505736 | 0.42164 |
| FOXP3 | chrX | 49127269 | 17 | 22 | 1516.309524 | 2025.595238 | 0.42166 |
| F7 | chr13 | 113750404 | 22 | 19 | 1981.599512 | 1761.837607 | 0.42202 |
| APOA1 | chr11 | 116720951 | 40 | 32 | 3422.044976 | 2705.590442 | 0.42212 |
| ARHGEF18 | chr19 | 7528643 | 11 | 20 | 1087.5 | 1944.242424 | 0.42224 |
| APOA1 | chr11 | 116680131 | 26 | 27 | 2568.845599 | 2612.657204 | 0.42232 |
| FOXP3 | chrX | 49128921 | 33 | 32 | 2413.461935 | 2420.983699 | 0.42234 |
| AMOTL2 | chr3 | 134094925 | 33 | 32 | 3051.521261 | 3007.907707 | 0.42238 |
| ADRA2C | chr4 | 3770090 | 27 | 30 | 1084.210853 | 1304.869575 | 0.42246 |
| F7 | chr13 | 113752175 | 31 | 31 | 2625.505899 | 2527.326701 | 0.42254 |
| ESR2 | chr14 | 64813738 | 13 | 13 | 1063.640526 | 1117.337662 | 0.42254 |
| AKT2 | chr19 | 40796299 | 33 | 32 | 3217.892437 | 3043.080048 | 0.42256 |
| FOXP3 | chrX | 49117271 | 3 | 3 | 300 | 287.5 | 0.42264 |
| APOA1 | chr11 | 116708004 | 2 | 2 | 200 | 166.6666667 | 0.42264 |
| F7 | chr13 | 113751134 | 2 | 2 | 166.6666667 | 200 | 0.42264 |
| LOXL1 | chr15 | 74215603 | 2 | 2 | 43.07692308 | 40 | 0.42264 |
| LCK | chr1 | 32741900 | 2 | 2 | 40 | 120 | 0.42264 |
| ATF3 | chr1 | 212739256 | 2 | 2 | 40 | 120 | 0.42264 |
| ATF3 | chr1 | 212739282 | 2 | 2 | 40 | 80 | 0.42264 |
| MIA3 | chr1 | 222790944 | 2 | 2 | 40 | 110.9090909 | 0.42264 |
| MIA3 | chr1 | 222790952 | 2 | 2 | 40 | 100 | 0.42264 |
| MIA3 | chr1 | 222814691 | 2 | 2 | 40 | 53.33333333 | 0.42264 |
| MIA3 | chr1 | 222816989 | 2 | 2 | 40 | 120 | 0.42264 |
| GALNT2 | chr1 | 230184232 | 2 | 2 | 40 | 120 | 0.42264 |

| | | | | | | | |
|---|---|---|---|---|---|---|---|
| GALNT2 | chr1 | 230188292 | 2 | 2 | 40 | 120 | 0.42264 |
| GALNT2 | chr1 | 230190098 | 2 | 2 | 40 | 120 | 0.42264 |
| APOA1 | chr11 | 116664744 | 2 | 2 | 40 | 120 | 0.42264 |
| APOA1 | chr11 | 116666038 | 2 | 2 | 200 | 160 | 0.42264 |
| APOA1 | chr11 | 116666189 | 2 | 2 | 42.22222222 | 40 | 0.42264 |
| APOA1 | chr11 | 116667078 | 2 | 2 | 180 | 200 | 0.42264 |
| APOA1 | chr11 | 116667699 | 2 | 2 | 40 | 120 | 0.42264 |
| APOA1 | chr11 | 116668936 | 2 | 2 | 40 | 120 | 0.42264 |
| APOA1 | chr11 | 116682375 | 2 | 2 | 40 | 120 | 0.42264 |
| APOA1 | chr11 | 116691709 | 2 | 2 | 40 | 120 | 0.42264 |
| APOA1 | chr11 | 116694563 | 2 | 2 | 40 | 120 | 0.42264 |
| APOA1 | chr11 | 116698413 | 2 | 2 | 40 | 120 | 0.42264 |
| APOA1 | chr11 | 116715915 | 2 | 2 | 40 | 120 | 0.42264 |
| TMPO | chr12 | 98905212 | 2 | 2 | 180 | 200 | 0.42264 |
| ESR2 | chr14 | 64807178 | 2 | 2 | 40 | 53.33333333 | 0.42264 |
| ESR2 | chr14 | 64813189 | 2 | 2 | 40 | 120 | 0.42264 |
| LOXL1 | chr15 | 74216534 | 2 | 2 | 40 | 120 | 0.42264 |
| ALDOA | chr16 | 30060970 | 2 | 2 | 40 | 120 | 0.42264 |
| ALDOA | chr16 | 30064121 | 2 | 2 | 40 | 60 | 0.42264 |
| ALDOA | chr16 | 30076803 | 2 | 2 | 40 | 120 | 0.42264 |
| ALOX15 | chr17 | 4546611 | 2 | 2 | 40 | 45 | 0.42264 |
| ALOX15 | chr17 | 4551816 | 2 | 2 | 40 | 120 | 0.42264 |
| ICAM2 | chr17 | 62084890 | 2 | 2 | 40 | 120 | 0.42264 |
| ICAM2 | chr17 | 62087794 | 2 | 2 | 40 | 120 | 0.42264 |
| PECAM1 | chr17 | 62401254 | 2 | 2 | 40 | 120 | 0.42264 |
| BCL2 | chr18 | 60987967 | 2 | 2 | 60 | 40 | 0.42264 |
| ARHGEF18 | chr19 | 7461112 | 2 | 2 | 40 | 53.33333333 | 0.42264 |
| LDLR | chr19 | 11195173 | 2 | 2 | 180 | 200 | 0.42264 |
| LDLR | chr19 | 11223916 | 2 | 2 | 40 | 120 | 0.42264 |
| KLF2 | chr19 | 16426908 | 2 | 2 | 40 | 120 | 0.42264 |
| BAX | chr19 | 49448388 | 2 | 2 | 40 | 120 | 0.42264 |
| E2F6 | chr2 | 11605919 | 2 | 2 | 40 | 120 | 0.42264 |
| E2F6 | chr2 | 11607499 | 2 | 2 | 40 | 120 | 0.42264 |
| E2F6 | chr2 | 11611502 | 2 | 2 | 40 | 120 | 0.42264 |
| E2F6 | chr2 | 11612044 | 2 | 2 | 40 | 120 | 0.42264 |
| ABCG1 | chr21 | 43637759 | 2 | 2 | 40 | 120 | 0.42264 |
| ABCG1 | chr21 | 43638716 | 2 | 2 | 40 | 120 | 0.42264 |
| TIMP3 | chr22 | 33190502 | 2 | 2 | 40 | 120 | 0.42264 |
| TIMP3 | chr22 | 33259403 | 2 | 2 | 40 | 120 | 0.42264 |
| AMOTL2 | chr3 | 134095545 | 2 | 2 | 40 | 120 | 0.42264 |
| AMOTL2 | chr3 | 134100335 | 2 | 2 | 40 | 105.7142857 | 0.42264 |
| ADRA2C | chr4 | 3762438 | 2 | 2 | 40 | 120 | 0.42264 |
| ADRA2C | chr4 | 3766451 | 2 | 2 | 40 | 120 | 0.42264 |
| ADRA2C | chr4 | 3767337 | 2 | 2 | 40 | 120 | 0.42264 |
| TERT | chr5 | 1299274 | 2 | 2 | 135 | 120 | 0.42264 |
| TERT | chr5 | 1301841 | 2 | 2 | 40 | 120 | 0.42264 |
| HMGCR | chr5 | 74624749 | 2 | 2 | 40 | 120 | 0.42264 |

| | | | | | | | |
|---|---|---|---|---|---|---|---|
| DHFR | chr5 | 79954368 | 2 | 2 | 53.33333333 | 40 | 0.42264 |
| CDKN1A | chr6 | 36654578 | 2 | 2 | 40 | 120 | 0.42264 |
| CTGF | chr6 | 132272228 | 2 | 2 | 40 | 120 | 0.42264 |
| ESR1 | chr6 | 152010736 | 2 | 2 | 40 | 120 | 0.42264 |
| ESR1 | chr6 | 152011732 | 2 | 2 | 40 | 120 | 0.42264 |
| LPA | chr6 | 160951823 | 2 | 2 | 40 | 120 | 0.42264 |
| NOS3 | chr7 | 150684429 | 2 | 2 | 40 | 120 | 0.42264 |
| NOS3 | chr7 | 150689279 | 2 | 2 | 40 | 120 | 0.42264 |
| NOS3 | chr7 | 150709452 | 2 | 2 | 180 | 200 | 0.42264 |
| NOS3 | chr7 | 150710123 | 2 | 2 | 40 | 120 | 0.42264 |
| MCPH1 | chr8 | 6260214 | 2 | 2 | 40 | 120 | 0.42264 |
| MCPH1 | chr8 | 6470957 | 2 | 2 | 40 | 120 | 0.42264 |
| MCPH1 | chr8 | 6471034 | 2 | 2 | 180 | 200 | 0.42264 |
| CDKN2B-AS1 | chr9 | 21966293 | 2 | 2 | 40 | 120 | 0.42264 |
| CDKN2B-AS1 | chr9 | 21971106 | 2 | 2 | 47.27272727 | 40 | 0.42264 |
| CDKN2B-AS1 | chr9 | 21977681 | 2 | 2 | 40 | 120 | 0.42264 |
| CDKN2B-AS1 | chr9 | 21993872 | 2 | 2 | 40 | 120 | 0.42264 |
| ABCA1 | chr9 | 107691513 | 2 | 2 | 40 | 120 | 0.42264 |
| ABCA1 | chr9 | 107697164 | 2 | 2 | 53.33333333 | 40 | 0.42264 |
| FOXP3 | chrX | 49119297 | 2 | 2 | 40 | 120 | 0.42264 |
| FOXP3 | chrX | 49122092 | 2 | 2 | 40 | 112.8571429 | 0.42264 |
| FOXP3 | chrX | 49123680 | 2 | 2 | 40 | 103.3333333 | 0.42264 |
| FOXP3 | chrX | 49127372 | 2 | 2 | 40 | 120 | 0.42264 |
| COL15A1 | chr9 | 101699833 | 30 | 30 | 2573.222823 | 2458.331799 | 0.4227 |
| CDKN2B-AS1 | chr9 | 22006090 | 4 | 2 | 95.29411765 | 133.3333333 | 0.42296 |
| CDKN2B-AS1 | chr9 | 21991699 | 37 | 33 | 2456.066943 | 2120.500827 | 0.42322 |
| LDLR | chr19 | 11224450 | 35 | 32 | 3313.8822 | 3057.941177 | 0.42332 |
| BCL2 | chr18 | 60988286 | 4 | 12 | 96.98335913 | 378.4984358 | 0.42332 |
| FGF2 | chr4 | 123745999 | 25 | 28 | 1747.598162 | 2063.737651 | 0.42338 |
| FOXP3 | chrX | 49131044 | 30 | 31 | 2667.332203 | 2814.120237 | 0.42346 |
| APOA1 | chr11 | 116699663 | 33 | 33 | 2506.67929 | 2591.241037 | 0.42352 |
| LDLR | chr19 | 11191309 | 19 | 18 | 1674.365079 | 1648.571429 | 0.42366 |
| F2RL3 | chr19 | 16992487 | 13 | 15 | 1183.030303 | 1424.047619 | 0.42388 |
| AMOTL2 | chr3 | 134095503 | 31 | 29 | 2706.40762 | 2592.151933 | 0.4239 |
| NOS3 | chr7 | 150682726 | 36 | 32 | 3116.698826 | 2815.397942 | 0.42416 |
| F2RL3 | chr19 | 16993566 | 33 | 32 | 2853.845186 | 2827.346999 | 0.42422 |
| LDLR | chr19 | 11220160 | 29 | 32 | 2841.500722 | 3088.522409 | 0.42434 |
| APOA1 | chr11 | 116674986 | 33 | 33 | 2688.435535 | 2606.170992 | 0.42462 |
| ADRA2C | chr4 | 3767103 | 3 | 4 | 79.76190476 | 182.2222222 | 0.42488 |
| GALNT2 | chr1 | 230191865 | 34 | 32 | 3195.869051 | 3056.918685 | 0.425 |
| NOS3 | chr7 | 150706287 | 33 | 32 | 2396.420031 | 2237.737393 | 0.42512 |
| GCK | chr7 | 44230278 | 37 | 32 | 3011.688689 | 2665.9908 | 0.42518 |
| CDKN2B-AS1 | chr9 | 22007682 | 33 | 31 | 1848.382872 | 1637.743693 | 0.4254 |
| CDKN2B-AS1 | chr9 | 22005985 | 16 | 15 | 433.6430391 | 471.5646686 | 0.42544 |
| AKT2 | chr19 | 40800968 | 16 | 24 | 1510.15873 | 2171.742424 | 0.42566 |
| APOA1 | chr11 | 116685422 | 32 | 33 | 1715.909225 | 1865.8218 | 0.42568 |
| LPA | chr6 | 160946156 | 34 | 32 | 3085.459684 | 2859.965462 | 0.42572 |

| | | | | | | | |
|---|---|---|---|---|---|---|---|
| NOS3 | chr7 | 150711224 | 3 | 4 | 78.57142857 | 174.7619048 | 0.4259 |
| COL15A1 | chr9 | 101704741 | 2 | 2 | 45 | 122.2222222 | 0.4262 |
| APOA1 | chr11 | 116690611 | 34 | 32 | 2944.975065 | 2833.890028 | 0.4263 |
| CTGF | chr6 | 132281493 | 33 | 32 | 2859.634629 | 2717.479718 | 0.42636 |
| APOA1 | chr11 | 116711711 | 32 | 32 | 2778.038217 | 2826.281493 | 0.42636 |
| CDKN2B-AS1 | chr9 | 22005402 | 24 | 22 | 683.1650641 | 683.9835228 | 0.42642 |
| APOA1 | chr11 | 116705097 | 18 | 21 | 1776.190476 | 2037.5 | 0.42654 |
| CDKN2B-AS1 | chr9 | 21969280 | 12 | 15 | 323.3306676 | 471.8236579 | 0.42654 |
| CDKN2B-AS1 | chr9 | 22006119 | 3 | 4 | 75.84415584 | 164.5299145 | 0.42658 |
| AKT2 | chr19 | 40791931 | 37 | 33 | 2972.537892 | 2582.648556 | 0.42666 |
| FOXP3 | chrX | 49128857 | 31 | 33 | 2168.01239 | 2413.558873 | 0.42678 |
| APOA1 | chr11 | 116671421 | 39 | 32 | 3660.004427 | 2977.444244 | 0.427 |
| ABCB1 | chr7 | 87352258 | 38 | 32 | 3418.425522 | 2924.095744 | 0.42706 |
| CDKN2B-AS1 | chr9 | 21969749 | 3 | 2 | 65.2991453 | 130.7692308 | 0.42716 |
| ADRA2C | chr4 | 3765124 | 5 | 8 | 125 | 231.7261905 | 0.42736 |
| APOA1 | chr11 | 116705604 | 19 | 23 | 857.9082029 | 945.0923784 | 0.42748 |
| PECAM1 | chr17 | 62398357 | 32 | 32 | 3152.10807 | 3101.639562 | 0.4277 |
| TERT | chr5 | 1295989 | 27 | 29 | 1219.187381 | 1404.808296 | 0.42806 |
| ARHGEF18 | chr19 | 7466055 | 33 | 32 | 3213.036614 | 3041.907487 | 0.42814 |
| GALNT2 | chr1 | 230190391 | 29 | 30 | 2250.939663 | 2425.052031 | 0.42828 |
| APOA1 | chr11 | 116695807 | 19 | 20 | 1709.563492 | 1849.004329 | 0.42836 |
| APOA1 | chr11 | 116661410 | 32 | 32 | 2919.118424 | 2981.781718 | 0.42842 |
| MIA3 | chr1 | 222808112 | 30 | 32 | 943.5030595 | 1072.240715 | 0.4289 |
| APOA1 | chr11 | 116678414 | 18 | 13 | 1060.639916 | 678.6507937 | 0.4289 |
| LPA | chr6 | 11219498 | 35 | 33 | 2116.408466 | 2056.318976 | 0.429 |
| ESR2 | chr14 | 64805945 | 12 | 8 | 353.0339105 | 288.0372405 | 0.42934 |
| GALNT2 | chr1 | 230193569 | 33 | 33 | 2006.593464 | 1925.546668 | 0.42946 |
| LCK | chr1 | 32707300 | 6 | 10 | 143.0642431 | 296.6298868 | 0.42954 |
| GCK | chr7 | 44231463 | 8 | 6 | 780 | 563.3333333 | 0.42984 |
| APOA1 | chr11 | 116716869 | 25 | 27 | 2352.664597 | 2485.645804 | 0.42992 |
| APOA1 | chr11 | 116708300 | 32 | 31 | 2976.245371 | 2924.705641 | 0.42996 |
| CDKN2B-AS1 | chr9 | 22006155 | 3 | 4 | 85.17316017 | 186.9047619 | 0.43014 |
| NOS3 | chr7 | 150706062 | 30 | 33 | 1442.786115 | 1677.710866 | 0.43024 |
| LDLR | chr19 | 11221689 | 31 | 29 | 2860.903376 | 2726.543873 | 0.43032 |
| E2F6 | chr2 | 11611772 | 29 | 26 | 2781.996435 | 2428.391331 | 0.43084 |
| BAX | chr19 | 49453165 | 23 | 16 | 1910.409035 | 1260.425685 | 0.43086 |
| TERT | chr5 | 1300302 | 30 | 31 | 2882.22445 | 2927.300477 | 0.43106 |
| CDKN2B-AS1 | chr9 | 21992065 | 36 | 33 | 2568.135929 | 2282.378736 | 0.43154 |
| FOXP3 | chrX | 49106867 | 20 | 24 | 1964.242424 | 2324.642857 | 0.43164 |
| F7 | chr13 | 113752435 | 33 | 32 | 2999.859096 | 2949.799792 | 0.4317 |
| GALNT2 | chr1 | 230193804 | 32 | 32 | 2969.199359 | 2913.111059 | 0.43172 |
| MCPH1 | chr8 | 6256556 | 32 | 32 | 2691.930271 | 2756.870433 | 0.43194 |
| AMOTL2 | chr3 | 134100808 | 27 | 22 | 815.4270188 | 616.3989312 | 0.43228 |
| F7 | chr13 | 113775058 | 29 | 30 | 2612.673406 | 2647.291499 | 0.4323 |
| F7 | chr13 | 113751474 | 31 | 31 | 2904.519843 | 2847.661708 | 0.43232 |
| ADRA2C | chr4 | 3764980 | 6 | 5 | 178.5714286 | 214.7727273 | 0.43232 |
| CDKN2B-AS1 | chr9 | 22008804 | 3 | 6 | 68.57142857 | 229.1269841 | 0.4325 |

| | | | | | | | |
|---|---|---|---|---|---|---|---|
| AMOTL2 | chr3 | 134100972 | 32 | 33 | 2966.314856 | 2989.323879 | 0.43254 |
| CDKN2B-AS1 | chr9 | 21967523 | 2 | 3 | 45 | 147.0833333 | 0.43272 |
| BCL2 | chr18 | 60989129 | 2 | 2 | 48.57142857 | 123.5294118 | 0.43274 |
| PHACTR1 | chr6 | 12711001 | 34 | 32 | 3025.078932 | 2895.370311 | 0.43278 |
| FOXP3 | chrX | 49125663 | 3 | 2 | 109.8452012 | 128.5714286 | 0.43278 |
| FOXP3 | chrX | 49125669 | 2 | 2 | 47.77777778 | 123.0769231 | 0.43282 |
| CTGF | chr6 | 132276178 | 32 | 32 | 2658.323609 | 2599.416136 | 0.43284 |
| FOXP3 | chrX | 49131141 | 34 | 32 | 3118.377885 | 2975.950904 | 0.43292 |
| CDKN2B-AS1 | chr9 | 21968704 | 2 | 2 | 45 | 107.4534161 | 0.43308 |
| ADRA2C | chr4 | 3766546 | 17 | 24 | 1276.126241 | 1903.455988 | 0.4333 |
| APOA1 | chr11 | 116685511 | 33 | 33 | 1657.42339 | 1581.711569 | 0.43332 |
| F7 | chr13 | 113750701 | 28 | 24 | 2467.153378 | 2184.7669 | 0.43338 |
| LCK | chr1 | 32740737 | 2 | 3 | 40 | 148.5714286 | 0.43362 |
| PHACTR1 | chr6 | 12708680 | 34 | 32 | 2389.829595 | 2315.502986 | 0.43368 |
| LDLR | chr19 | 11224051 | 26 | 20 | 2497.460317 | 1966.666667 | 0.43368 |
| ADRA2C | chr4 | 3763603 | 31 | 33 | 2630.993591 | 2725.179175 | 0.43374 |
| F7 | chr13 | 113755228 | 10 | 15 | 806.5079365 | 1292.142857 | 0.43386 |
| ABCA1 | chr9 | 107695620 | 3 | 3 | 101.5384615 | 160 | 0.43386 |
| PECAM1 | chr17 | 62400500 | 3 | 4 | 73.57142857 | 168.5714286 | 0.43388 |
| CDKN2B-AS1 | chr9 | 21968682 | 8 | 3 | 185.6592966 | 144.949495 | 0.43404 |
| ADRA2C | chr4 | 3767169 | 3 | 3 | 65 | 132.1052632 | 0.43408 |
| LDLR | chr19 | 11221356 | 33 | 32 | 3138.998233 | 3080.066681 | 0.43414 |
| LDLR | chr19 | 11222331 | 15 | 20 | 1479.166667 | 1945.436508 | 0.4342 |
| CDKN2B-AS1 | chr9 | 22009954 | 3 | 6 | 96.66666666 | 253.8888889 | 0.4344 |
| CDKN2B-AS1 | chr9 | 21993629 | 20 | 23 | 741.9334316 | 779.010545 | 0.43442 |
| ADRA2C | chr4 | 3763593 | 26 | 28 | 1475.354972 | 1718.647889 | 0.43448 |
| TERT | chr5 | 1301204 | 34 | 33 | 2140.49634 | 2152.265831 | 0.43456 |
| CDKN1A | chr6 | 36648839 | 32 | 32 | 2901.274689 | 2815.373412 | 0.43456 |
| TIMP3 | chr22 | 33196992 | 33 | 33 | 1946.082896 | 2045.810351 | 0.4348 |
| ADRA2C | chr4 | 3764919 | 4 | 3 | 113.5714286 | 127.2222222 | 0.43508 |
| BAX | chr19 | 49451077 | 13 | 9 | 1231.623377 | 822.2943723 | 0.43522 |
| ADRA2C | chr4 | 3764891 | 3 | 3 | 87.27272727 | 153.3333333 | 0.43532 |
| KLF2 | chr19 | 16437949 | 30 | 32 | 1496.051115 | 1699.105845 | 0.43544 |
| ESR2 | chr14 | 64806459 | 32 | 30 | 3117.214591 | 2953.333333 | 0.43586 |
| TERT | chr5 | 1300704 | 30 | 31 | 2869.335583 | 2907.133536 | 0.43592 |
| LCK | chr1 | 32713399 | 33 | 32 | 3165.810318 | 3106.678165 | 0.43596 |
| APOA1 | chr11 | 116720090 | 20 | 23 | 1298.804495 | 1352.428576 | 0.43612 |
| CDKN2B-AS1 | chr9 | 21980007 | 37 | 33 | 2898.219162 | 2512.57227 | 0.43622 |
| LOXL1 | chr15 | 74244316 | 29 | 29 | 1313.67549 | 1413.046497 | 0.43642 |
| LDLR | chr19 | 11191927 | 3 | 4 | 240 | 360 | 0.43658 |
| CDKN1A | chr6 | 36644403 | 34 | 32 | 3186.751763 | 3036.699089 | 0.43692 |
| ARHGEF18 | chr19 | 7464894 | 7 | 5 | 680 | 465.7142857 | 0.43696 |
| ADRA2C | chr4 | 3768473 | 2 | 7 | 44.12955466 | 260.6349206 | 0.43714 |
| CDKN2B-AS1 | chr9 | 21965991 | 4 | 3 | 97.5 | 147.7777778 | 0.43718 |
| APOA1 | chr11 | 116695204 | 12 | 11 | 997.6587302 | 857.7777778 | 0.43726 |
| FOXP3 | chrX | 49115028 | 14 | 6 | 1317.142857 | 585.7142857 | 0.43746 |
| APOA1 | chr11 | 116715706 | 38 | 32 | 3701.901119 | 3098.401743 | 0.43772 |

| | | | | | | | |
|---|---|---|---|---|---|---|---|
| F2RL3 | chr19 | 16992144 | 33 | 32 | 3051.435731 | 2995.758059 | 0.43822 |
| AMOTL2 | chr3 | 134096132 | 33 | 33 | 2985.600792 | 3052.489115 | 0.43826 |
| PHACTR1 | chr6 | 12712515 | 19 | 21 | 1149.747475 | 1145.505606 | 0.43836 |
| APOA1 | chr11 | 116700437 | 33 | 32 | 2632.554617 | 2478.614851 | 0.43852 |
| LDLR | chr19 | 11191757 | 10 | 14 | 854.3452381 | 1115.714286 | 0.43856 |
| CDKN2B-AS1 | chr9 | 22006006 | 12 | 11 | 350.4029794 | 386.4778926 | 0.43856 |
| GCK | chr7 | 44236443 | 28 | 33 | 2096.590132 | 2365.799498 | 0.43864 |
| PLA2G7 | chr6 | 46706271 | 11 | 5 | 620.6349206 | 335.2777778 | 0.43876 |
| PLA2G7 | chr6 | 46703496 | 4 | 4 | 105.2052545 | 163.2632633 | 0.43886 |
| LDLR | chr19 | 11194526 | 34 | 32 | 2935.627608 | 2696.432138 | 0.43888 |
| PLA2G7 | chr6 | 46704015 | 33 | 31 | 2589.016793 | 2358.911324 | 0.43896 |
| ATF3 | chr1 | 212783663 | 34 | 32 | 2959.373589 | 2832.25113 | 0.43902 |
| CDKN2B-AS1 | chr9 | 21967389 | 15 | 12 | 440.9691089 | 408.5271673 | 0.43962 |
| LDLR | chr19 | 11218990 | 8 | 12 | 683.3333333 | 918.0952381 | 0.43982 |
| ESR1 | chr6 | 152011103 | 36 | 33 | 2114.509728 | 1874.242143 | 0.43984 |
| ICAM2 | chr17 | 62085197 | 30 | 31 | 2840.246698 | 2987.373737 | 0.43988 |
| LDLR | chr19 | 11216501 | 2 | 2 | 42.22222222 | 128.5714286 | 0.44012 |
| FGF2 | chr4 | 123739122 | 12 | 8 | 1012.178932 | 633.3333333 | 0.44022 |
| NOS3 | chr7 | 150686192 | 4 | 3 | 100.5555556 | 148.5714286 | 0.44024 |
| TFPI | chr2 | 188418807 | 9 | 6 | 249.50333 | 226.0124043 | 0.44028 |
| APOA1 | chr11 | 116694006 | 34 | 32 | 3155.289195 | 3025.157633 | 0.4404 |
| APOA1 | chr11 | 116697812 | 30 | 31 | 2293.556171 | 2288.848619 | 0.44064 |
| ABCG1 | chr21 | 43629989 | 37 | 32 | 3059.419558 | 2694.820871 | 0.4409 |
| LOXL1 | chr15 | 74244101 | 14 | 10 | 1056.269841 | 810.4365079 | 0.44096 |
| F7 | chr13 | 113751564 | 30 | 29 | 2670.681703 | 2476.665304 | 0.4411 |
| LDLR | chr19 | 11219972 | 16 | 17 | 1509.95671 | 1652.380952 | 0.44112 |
| APOA1 | chr11 | 116691407 | 32 | 32 | 3054.788144 | 3000.975501 | 0.44114 |
| ATF3 | chr1 | 212745349 | 31 | 31 | 3120.760284 | 3022.939699 | 0.4413 |
| APOA1 | chr11 | 116676409 | 32 | 32 | 3048.268057 | 3077.358529 | 0.44134 |
| LOXL1 | chr15 | 74216859 | 34 | 33 | 2123.081879 | 2124.983656 | 0.4414 |
| LOXL1 | chr15 | 74216431 | 4 | 3 | 104.4444444 | 150.7936508 | 0.4415 |
| AMOTL2 | chr3 | 134095543 | 2 | 6 | 40 | 231.9047619 | 0.4416 |
| COL15A1 | chr9 | 101699436 | 34 | 33 | 3010.053852 | 2852.214289 | 0.44162 |
| APOA1 | chr11 | 116703529 | 35 | 32 | 3194.775802 | 2879.933147 | 0.44226 |
| CDKN2B-AS1 | chr9 | 22009596 | 3 | 2 | 70.79365079 | 122.8571429 | 0.44236 |
| APOA1 | chr11 | 116671217 | 33 | 32 | 2966.000474 | 2820.699084 | 0.44238 |
| ARHGEF18 | chr19 | 7461704 | 19 | 17 | 1845.714286 | 1683.333333 | 0.44276 |
| BAX | chr19 | 49465339 | 24 | 28 | 802.9851994 | 1009.478826 | 0.44282 |
| ICAM2 | chr17 | 62087500 | 34 | 32 | 3210.34816 | 3056.173607 | 0.44288 |
| F7 | chr13 | 113753614 | 37 | 32 | 3354.276781 | 2945.119735 | 0.44314 |
| COL15A1 | chr9 | 101832886 | 2 | 2 | 60.6060606 | 128.5714286 | 0.44314 |
| P2RY12 | chr3 | 151103169 | 35 | 33 | 2436.389265 | 2380.02421 | 0.4433 |
| LCK | chr1 | 32715485 | 2 | 2 | 43.30357143 | 128.5714286 | 0.44392 |
| PHACTR1 | chr6 | 12713273 | 33 | 32 | 2532.158443 | 2370.539034 | 0.44396 |
| ICAM2 | chr17 | 62080155 | 33 | 32 | 3075.463205 | 3038.561867 | 0.44404 |
| APOA1 | chr11 | 116704360 | 34 | 32 | 3518.906821 | 3135.300279 | 0.4442 |
| LDLR | chr19 | 11217166 | 4 | 10 | 367.5 | 960 | 0.44424 |

| | | | | | | | |
|---|---|---|---|---|---|---|---|
| FOXP3 | chrX | 49126828 | 7 | 6 | 162.2222222 | 162.2222222 | 0.44432 |
| NOS3 | chr7 | 150694593 | 29 | 31 | 1787.209441 | 2014.205073 | 0.44446 |
| ALOX15 | chr17 | 4549954 | 33 | 32 | 3264.342262 | 3036.84494 | 0.44496 |
| APOA1 | chr11 | 116682428 | 12 | 9 | 1162.103175 | 887.5 | 0.44512 |
| TERT | chr5 | 1295700 | 21 | 17 | 777.2077922 | 695.0195638 | 0.44516 |
| F2RL3 | chr19 | 16991338 | 31 | 32 | 2932.30503 | 2961.152591 | 0.44528 |
| F7 | chr13 | 113755281 | 30 | 30 | 2964.615385 | 2942.04472 | 0.44544 |
| LCK | chr1 | 32740783 | 20 | 21 | 1953.492063 | 2023.611111 | 0.4456 |
| CDKN2B-AS1 | chr9 | 21981718 | 22 | 22 | 1510.786436 | 1416.684084 | 0.44562 |
| LDLR | chr19 | 11191729 | 11 | 11 | 968.6904762 | 1015.714286 | 0.44586 |
| PECAM1 | chr17 | 62409285 | 33 | 30 | 1614.796978 | 1382.39785 | 0.446 |
| CDKN2B-AS1 | chr9 | 21966410 | 2 | 2 | 53.33333333 | 125 | 0.44612 |
| TERT | chr5 | 1299597 | 6 | 10 | 537.5757576 | 843.3333333 | 0.4463 |
| ARHGEF18 | chr19 | 7469214 | 33 | 32 | 3174.402532 | 3107.29713 | 0.44638 |
| CDKN2B-AS1 | chr9 | 21979696 | 33 | 33 | 2110.454517 | 2202.132595 | 0.44642 |
| LOXL1 | chr15 | 74215978 | 33 | 32 | 2813.480119 | 2667.442339 | 0.44648 |
| F7 | chr13 | 113752657 | 33 | 30 | 3162.210359 | 2911.064872 | 0.44652 |
| ESR1 | chr6 | 152002470 | 32 | 32 | 2418.988841 | 2490.639383 | 0.44672 |
| CDKN2B-AS1 | chr9 | 21983915 | 6 | 3 | 225.3274063 | 175.8095238 | 0.44686 |
| FGF2 | chr4 | 123739628 | 19 | 16 | 1612.392052 | 1299.365079 | 0.44702 |
| BAX | chr19 | 49448853 | 3 | 8 | 211.4285714 | 627.5 | 0.44732 |
| LPA | chr6 | 11216222 | 35 | 32 | 3350.599773 | 3086.515595 | 0.44734 |
| ABCG1 | chr21 | 43634668 | 17 | 20 | 1547.387057 | 1869.79798 | 0.44736 |
| MCPH1 | chr8 | 6259667 | 19 | 19 | 1863.333333 | 1885.714286 | 0.44774 |
| ADRA2C | chr4 | 3768705 | 31 | 31 | 2100.784493 | 2002.46096 | 0.44776 |
| ICAM2 | chr17 | 62084206 | 19 | 19 | 651.0504886 | 589.9214407 | 0.44784 |
| ADRA2C | chr4 | 3764985 | 4 | 4 | 120 | 184.3406593 | 0.44786 |
| CDKN2B-AS1 | chr9 | 21969012 | 14 | 8 | 373.4629485 | 262.6628395 | 0.4481 |
| LCK | chr1 | 32740822 | 2 | 2 | 45 | 120 | 0.44822 |
| APOA1 | chr11 | 116699123 | 2 | 2 | 45 | 120 | 0.44822 |
| APOA1 | chr11 | 116706078 | 2 | 2 | 45 | 120 | 0.44822 |
| LDLR | chr19 | 11223202 | 2 | 2 | 45 | 120 | 0.44822 |
| ADRA2C | chr4 | 3764482 | 2 | 2 | 45 | 120 | 0.44822 |
| TERT | chr5 | 1253445 | 2 | 2 | 45 | 120 | 0.44822 |
| COL15A1 | chr9 | 101703316 | 2 | 2 | 45 | 120 | 0.44822 |
| TERT | chr5 | 1301481 | 32 | 32 | 2758.530963 | 2812.619363 | 0.44838 |
| TIMP3 | chr22 | 33196328 | 34 | 33 | 2668.405826 | 2530.368535 | 0.44844 |
| APOA1 | chr11 | 116686423 | 33 | 32 | 2739.48408 | 2698.220027 | 0.44864 |
| CDKN2B-AS1 | chr9 | 22005638 | 9 | 3 | 221.3584393 | 143.488529 | 0.44872 |
| CDKN2B-AS1 | chr9 | 22006159 | 2 | 4 | 48.57142857 | 173.2142857 | 0.4489 |
| F2RL3 | chr19 | 16994895 | 30 | 28 | 2778.210874 | 2517.261905 | 0.449 |
| FOXP3 | chrX | 49125897 | 2 | 2 | 47.80952381 | 130 | 0.44912 |
| ALOX15 | chr17 | 4549128 | 21 | 15 | 609.4481325 | 497.5969816 | 0.44922 |
| PLA2G7 | chr6 | 46703512 | 28 | 21 | 829.899255 | 669.4831822 | 0.44932 |
| GCK | chr7 | 44228751 | 36 | 33 | 1977.230279 | 1751.986496 | 0.4494 |
| NOS3 | chr7 | 150684090 | 17 | 21 | 1505.47619 | 1788.279221 | 0.4495 |
| NOS3 | chr7 | 150684671 | 30 | 30 | 2482.679987 | 2410.703574 | 0.4496 |

| | | | | | | | |
|---|---|---|---|---|---|---|---|
| LOXL1 | chr15 | 74219268 | 5 | 5 | 127.5639857 | 188.974195 | 0.44962 |
| LPA | chr6 | 11219555 | 35 | 32 | 3203.901345 | 2892.758687 | 0.44974 |
| NOS3 | chr7 | 150678361 | 35 | 33 | 1819.917819 | 1643.522504 | 0.44988 |
| TERT | chr5 | 1298540 | 13 | 8 | 1153.214286 | 657.2222222 | 0.45006 |
| CDKN2B-AS1 | chr9 | 22009696 | 6 | 4 | 146.2238562 | 148.7907363 | 0.45012 |
| PECAM1 | chr17 | 62398768 | 30 | 32 | 2547.718086 | 2640.041445 | 0.4502 |
| APOA1 | chr11 | 116715735 | 37 | 32 | 3614.528055 | 3103.91779 | 0.45022 |
| LDLR | chr19 | 11199535 | 35 | 32 | 3248.328789 | 2919.417432 | 0.45022 |
| APOA1 | chr11 | 116666256 | 33 | 32 | 3050.448336 | 2998.032013 | 0.45038 |
| NOS3 | chr7 | 150706731 | 27 | 26 | 2424.23818 | 2388.385226 | 0.45072 |
| ALDOA | chr16 | 30060778 | 32 | 33 | 2224.564948 | 2375.869619 | 0.45102 |
| BAX | chr19 | 49449641 | 31 | 32 | 2607.183947 | 2770.540811 | 0.45104 |
| LOXL1 | chr15 | 74218061 | 35 | 31 | 3344.311 | 2933.675597 | 0.45116 |
| APOA1 | chr11 | 116659902 | 29 | 29 | 2558.800821 | 2480.44289 | 0.4513 |
| F7 | chr13 | 113760006 | 28 | 25 | 2171.308969 | 2016.856754 | 0.45138 |
| GALNT2 | chr1 | 230190420 | 30 | 31 | 2768.377783 | 2913.317793 | 0.4514 |
| BCL2 | chr18 | 60994112 | 2 | 2 | 53.33333333 | 113.8888889 | 0.45144 |
| F7 | chr13 | 113757849 | 33 | 33 | 2942.744999 | 2872.64059 | 0.45148 |
| BCL2 | chr18 | 60992370 | 35 | 32 | 3198.163605 | 2967.991178 | 0.4517 |
| NOS3 | chr7 | 150710994 | 6 | 5 | 183.1457431 | 215.8333333 | 0.452 |
| BAX | chr19 | 49465399 | 32 | 33 | 1924.786146 | 1897.810322 | 0.45212 |
| E2F6 | chr2 | 11615192 | 18 | 19 | 1741.666667 | 1789.047619 | 0.45222 |
| APOA1 | chr11 | 116700448 | 33 | 31 | 2439.38031 | 2367.947373 | 0.4523 |
| E2F6 | chr2 | 11614366 | 36 | 32 | 3294.747786 | 2889.990199 | 0.45232 |
| GCK | chr7 | 44233544 | 32 | 32 | 2975.675043 | 3029.935657 | 0.4524 |
| APOA1 | chr11 | 116681658 | 32 | 32 | 3031.145798 | 2934.957847 | 0.45242 |
| LOXL1 | chr15 | 74244299 | 24 | 28 | 1061.711739 | 1315.069204 | 0.45274 |
| F7 | chr13 | 113756678 | 33 | 33 | 2335.898935 | 2267.546473 | 0.4528 |
| CDKN2B-AS1 | chr9 | 22005151 | 2 | 3 | 47.22222222 | 153.3333333 | 0.45286 |
| APOA1 | chr11 | 116690039 | 33 | 32 | 3103.562548 | 3041.292165 | 0.45292 |
| ESR2 | chr14 | 64693606 | 26 | 24 | 2400.744631 | 2255.436508 | 0.45316 |
| CDKN2B-AS1 | chr9 | 21986999 | 3 | 2 | 70 | 60 | 0.45334 |
| CDKN2B-AS1 | chr9 | 21996193 | 29 | 29 | 1648.17698 | 1552.77639 | 0.45364 |
| GALNT2 | chr1 | 230184639 | 39 | 32 | 3134.252823 | 2620.200989 | 0.45368 |
| GALNT2 | chr1 | 230187176 | 26 | 29 | 2280.368707 | 2608.061661 | 0.45372 |
| ADRA2C | chr4 | 3758777 | 34 | 32 | 2944.67121 | 2809.9311 | 0.45378 |
| CDKN2B-AS1 | chr9 | 22009200 | 2 | 2 | 48.57142857 | 121.4285714 | 0.45394 |
| PLA2G7 | chr6 | 46710121 | 32 | 32 | 2131.14716 | 2229.318119 | 0.45414 |
| NOS3 | chr7 | 150710536 | 31 | 31 | 1769.926039 | 1661.350099 | 0.45418 |
| APOA1 | chr11 | 116661393 | 32 | 32 | 2768.666277 | 2844.147145 | 0.45438 |
| BCL2 | chr18 | 60996205 | 38 | 32 | 3756.290073 | 3143.448762 | 0.45466 |
| APOA1 | chr11 | 116717837 | 31 | 31 | 2830.791933 | 2880.171342 | 0.45508 |
| CDKN1A | chr6 | 36651985 | 31 | 31 | 2960.898326 | 3001.697012 | 0.45536 |
| APOA1 | chr11 | 116712996 | 2 | 2 | 45 | 103.3333333 | 0.45544 |
| FOXP3 | chrX | 49125895 | 2 | 2 | 46.31578947 | 120 | 0.45544 |
| CTGF | chr6 | 132272384 | 3 | 3 | 66.42857143 | 140 | 0.45544 |
| F2RL3 | chr19 | 16994246 | 33 | 32 | 3103.923607 | 2972.581354 | 0.45564 |

| | | | | | | | |
|---|---|---|---|---|---|---|---|
| CDKN2B-AS1 | chr9 | 22006607 | 35 | 33 | 2124.165413 | 1936.03321 | 0.45602 |
| LDLR | chr19 | 11216083 | 28 | 30 | 2671.186688 | 2831.336171 | 0.45606 |
| ALDOA | chr16 | 30061407 | 30 | 29 | 2861.277149 | 2818.856797 | 0.45612 |
| TERT | chr5 | 1295966 | 29 | 33 | 1894.719978 | 2039.509518 | 0.45622 |
| FOXP3 | chrX | 49127402 | 21 | 26 | 1947.654392 | 2463.484848 | 0.45644 |
| APOA1 | chr11 | 116683482 | 29 | 30 | 2804.318994 | 2833.439246 | 0.4565 |
| APOA1 | chr11 | 116707514 | 26 | 24 | 2473.579492 | 2239.009324 | 0.45652 |
| KLF2 | chr19 | 16426673 | 33 | 30 | 2988.331239 | 2669.085518 | 0.45696 |
| LDLR | chr19 | 11224381 | 34 | 32 | 3181.340981 | 3023.575923 | 0.457 |
| ADRA2C | chr4 | 3768750 | 17 | 23 | 592.6517926 | 744.119538 | 0.45702 |
| APOA1 | chr11 | 116704662 | 35 | 32 | 2871.713718 | 2708.072739 | 0.45708 |
| KLF2 | chr19 | 16427025 | 34 | 30 | 1462.954065 | 1221.142369 | 0.4573 |
| APOA1 | chr11 | 116697334 | 32 | 32 | 2897.881234 | 2938.568636 | 0.45738 |
| LDLR | chr19 | 11191563 | 11 | 5 | 1017.936508 | 442.2619048 | 0.45744 |
| TIMP3 | chr22 | 33195200 | 32 | 30 | 2387.848293 | 2155.950378 | 0.45754 |
| F7 | chr13 | 113752701 | 33 | 32 | 3165.840188 | 3108.780531 | 0.45754 |
| CDKN2B-AS1 | chr9 | 21993913 | 4 | 4 | 113.5714286 | 165.3781513 | 0.45774 |
| TERT | chr5 | 1300498 | 36 | 32 | 3201.848873 | 2876.977114 | 0.45848 |
| ARHGEF18 | chr19 | 7460681 | 8 | 6 | 198.424801 | 211.7460318 | 0.45852 |
| ARHGEF18 | chr19 | 7467675 | 36 | 32 | 3332.928528 | 2991.828077 | 0.45856 |
| CDKN2B-AS1 | chr9 | 21968050 | 13 | 14 | 343.6180243 | 422.7062533 | 0.4586 |
| HMGCR | chr5 | 74624788 | 2 | 3 | 40 | 145 | 0.45862 |
| TIMP3 | chr22 | 33259146 | 6 | 9 | 467.8571429 | 632.7380952 | 0.45868 |
| KLF2 | chr19 | 16433698 | 35 | 32 | 3237.746821 | 2996.610283 | 0.45876 |
| LPA | chr6 | 160950868 | 27 | 26 | 1918.564426 | 1946.327561 | 0.45882 |
| CDKN2B-AS1 | chr9 | 21969217 | 10 | 11 | 284.7751956 | 379.5848596 | 0.45902 |
| ABCG1 | chr21 | 43629545 | 40 | 32 | 3736.337576 | 3023.923846 | 0.45906 |
| F2RL3 | chr19 | 16989827 | 33 | 32 | 3172.85077 | 3121.738732 | 0.45922 |
| NOS3 | chr7 | 150707764 | 15 | 18 | 427.5739256 | 569.5741536 | 0.45954 |
| MIA3 | chr1 | 222841260 | 37 | 32 | 3433.537104 | 2997.352866 | 0.45956 |
| TERT | chr5 | 1301058 | 33 | 32 | 3014.07531 | 2881.671644 | 0.45978 |
| BAX | chr19 | 49465501 | 4 | 3 | 103.0769231 | 147.2727273 | 0.45996 |
| PECAM1 | chr17 | 62397007 | 19 | 11 | 1705.555556 | 940 | 0.46004 |
| CDKN2B-AS1 | chr9 | 21981802 | 3 | 2 | 60 | 125 | 0.46026 |
| APOA1 | chr11 | 116663516 | 33 | 32 | 3131.498724 | 3063.241985 | 0.46032 |
| DHFR | chr5 | 79959837 | 35 | 32 | 3305.249228 | 3053.925984 | 0.46036 |
| TIMP3 | chr22 | 33196848 | 5 | 3 | 377.1428571 | 260 | 0.46036 |
| ARHGEF18 | chr19 | 7466735 | 37 | 32 | 3606.295377 | 3097.256794 | 0.4604 |
| ADRA2C | chr4 | 3768743 | 15 | 14 | 469.3648019 | 483.466811 | 0.4604 |
| APOA1 | chr11 | 116720744 | 40 | 32 | 3734.348926 | 3014.571298 | 0.46042 |
| FOXP3 | chrX | 49125835 | 2 | 2 | 57.77777778 | 133.3333333 | 0.46044 |
| F2RL3 | chr19 | 16997385 | 31 | 32 | 2817.137177 | 2962.076617 | 0.46064 |
| GALNT2 | chr1 | 230193267 | 31 | 32 | 2790.558002 | 2963.370605 | 0.46066 |
| APOA1 | chr11 | 116723985 | 34 | 33 | 2249.939266 | 2120.125429 | 0.46072 |
| LDLR | chr19 | 11224607 | 2 | 2 | 47.27272727 | 120 | 0.46084 |
| HMGCR | chr5 | 74657944 | 2 | 2 | 47.27272727 | 120 | 0.46084 |
| PLA2G7 | chr6 | 46703316 | 2 | 2 | 47.27272727 | 120 | 0.46084 |

| | | | | | | | |
|---|---|---|---|---|---|---|---|
| CTGF | chr6 | 132272445 | 2 | 2 | 47.27272727 | 120 | 0.46084 |
| ADRA2C | chr4 | 3759780 | 32 | 32 | 3102.053969 | 3066.359086 | 0.46088 |
| F2RL3 | chr19 | 16999093 | 29 | 32 | 2307.359895 | 2446.948231 | 0.46096 |
| LOXL1 | chr15 | 74216471 | 2 | 2 | 57.14285714 | 125 | 0.46108 |
| ARHGEF18 | chr19 | 7464181 | 35 | 32 | 2812.377412 | 2628.180423 | 0.46134 |
| CDKN2B-AS1 | chr9 | 21992334 | 33 | 32 | 1379.099103 | 1262.572172 | 0.46172 |
| F2RL3 | chr19 | 16994271 | 31 | 32 | 2895.261242 | 2927.618573 | 0.4618 |
| LCK | chr1 | 32707389 | 17 | 9 | 437.6018422 | 273.0136696 | 0.46188 |
| NOS3 | chr7 | 150710122 | 2 | 3 | 53.33333333 | 158.3333333 | 0.462 |
| ADRA2C | chr4 | 3768719 | 28 | 32 | 1574.630623 | 1690.520406 | 0.46246 |
| APOA1 | chr11 | 116719942 | 34 | 32 | 3242.239477 | 3089.477516 | 0.46254 |
| GCK | chr7 | 44230108 | 35 | 33 | 2359.442429 | 2158.929505 | 0.46278 |
| APOA1 | chr11 | 116697295 | 32 | 32 | 3044.748305 | 3014.001723 | 0.46292 |
| TERT | chr5 | 1295961 | 31 | 33 | 2092.923359 | 2347.989239 | 0.46362 |
| CDKN2B-AS1 | chr9 | 22005564 | 5 | 5 | 147.7248686 | 192.4565637 | 0.46364 |
| CDKN2B-AS1 | chr9 | 21968458 | 11 | 11 | 325.9653092 | 390.8441558 | 0.46404 |
| NOS3 | chr7 | 150708974 | 30 | 29 | 2918.354701 | 2787.59325 | 0.46458 |
| MCPH1 | chr8 | 6263554 | 5 | 3 | 113.5714286 | 140 | 0.46468 |
| APOA1 | chr11 | 116694066 | 27 | 29 | 2416.125264 | 2522.805112 | 0.4647 |
| E2F6 | chr2 | 11613316 | 13 | 17 | 1236.379552 | 1577.619048 | 0.46476 |
| ADRA2C | chr4 | 3764012 | 32 | 31 | 2637.278482 | 2608.422488 | 0.46494 |
| CDKN2B-AS1 | chr9 | 22006102 | 2 | 5 | 41.73913043 | 193.5714286 | 0.46494 |
| CTGF | chr6 | 132275022 | 28 | 28 | 2179.339795 | 2092.448364 | 0.4651 |
| TERT | chr5 | 1301221 | 33 | 32 | 2917.096644 | 2874.33722 | 0.46516 |
| LCK | chr1 | 32740925 | 22 | 27 | 2035.985224 | 2544.373682 | 0.46528 |
| CDKN2B-AS1 | chr9 | 21992823 | 33 | 32 | 2707.318086 | 2677.480041 | 0.46552 |
| F2RL3 | chr19 | 17000171 | 7 | 4 | 310.6060606 | 200.4761905 | 0.46562 |
| CDKN2B-AS1 | chr9 | 21971063 | 2 | 2 | 41.42857143 | 125 | 0.46568 |
| F7 | chr13 | 113751424 | 28 | 30 | 2664.377289 | 2891.103896 | 0.46636 |
| APOA1 | chr11 | 116692169 | 33 | 32 | 3133.203118 | 3066.396479 | 0.46648 |
| CDKN2B-AS1 | chr9 | 21986445 | 38 | 33 | 2630.78906 | 2351.325658 | 0.46666 |
| F2RL3 | chr19 | 16997264 | 27 | 20 | 2134.345238 | 1481.738817 | 0.46666 |
| BAX | chr19 | 49453121 | 30 | 30 | 2691.264503 | 2752.292462 | 0.46672 |
| F7 | chr13 | 113756814 | 19 | 25 | 1651.215729 | 2105.308303 | 0.46692 |
| APOA1 | chr11 | 116700127 | 27 | 27 | 2558.448773 | 2603.095238 | 0.46706 |
| BAX | chr19 | 49465445 | 11 | 12 | 285.1572039 | 370.0915751 | 0.46732 |
| CDKN2B-AS1 | chr9 | 21968854 | 39 | 33 | 1471.800648 | 1191.745942 | 0.46734 |
| ARHGEF18 | chr19 | 7464952 | 34 | 32 | 3311.105666 | 3087.125985 | 0.46748 |
| CDKN2B-AS1 | chr9 | 21967468 | 3 | 7 | 75.84415584 | 266.6269841 | 0.4676 |
| GALNT2 | chr1 | 230193005 | 31 | 31 | 2305.226806 | 2406.376279 | 0.46818 |
| LCK | chr1 | 32740728 | 22 | 14 | 2091.077534 | 1354.761905 | 0.46828 |
| ESR1 | chr6 | 152007852 | 2 | 2 | 45 | 86.66666667 | 0.46832 |
| AMOTL2 | chr3 | 134095540 | 2 | 2 | 48.57142857 | 120 | 0.46832 |
| NOS3 | chr7 | 150706099 | 32 | 33 | 1784.794404 | 1931.396041 | 0.46834 |
| APOA1 | chr11 | 116690505 | 20 | 21 | 1739.349817 | 1769.542125 | 0.46842 |
| ADRA2C | chr4 | 3761333 | 27 | 31 | 2514.57007 | 2793.060418 | 0.46848 |
| DHFR | chr5 | 79958374 | 22 | 20 | 2169.047619 | 1948.571429 | 0.46856 |

| | | | | | | | |
|---|---|---|---|---|---|---|---|
| F7 | chr13 | 113750653 | 27 | 24 | 2545.169743 | 2309.563492 | 0.46866 |
| FOXP3 | chrX | 49118414 | 25 | 26 | 2311.984127 | 2453.756105 | 0.46878 |
| ABCG1 | chr21 | 43630340 | 2 | 2 | 42.22222222 | 125 | 0.46878 |
| LPA | chr6 | 11219996 | 2 | 2 | 42.22222222 | 125 | 0.46878 |
| LDLR | chr19 | 11199495 | 34 | 32 | 3099.451089 | 2855.903527 | 0.46884 |
| LPA | chr6 | 160946686 | 41 | 32 | 3437.651948 | 2641.135559 | 0.46898 |
| LDLR | chr19 | 11217369 | 32 | 32 | 3072.405516 | 3044.850583 | 0.46904 |
| ESR1 | chr6 | 152002874 | 21 | 14 | 615.3331728 | 469.018759 | 0.46906 |
| NOS3 | chr7 | 150692340 | 5 | 3 | 123.8705739 | 134.4444444 | 0.46908 |
| BCL2 | chr18 | 60988600 | 31 | 32 | 2937.322889 | 3066.910184 | 0.46946 |
| NOS3 | chr7 | 150678556 | 32 | 33 | 2040.107356 | 2186.916166 | 0.46948 |
| NR3C1 | chr5 | 142657679 | 37 | 32 | 3601.225234 | 3097.964067 | 0.46962 |
| MCPH1 | chr8 | 6262739 | 32 | 32 | 3072.910622 | 3113.761298 | 0.46982 |
| TERT | chr5 | 1253589 | 32 | 32 | 2862.354354 | 2920.666479 | 0.46984 |
| LDLR | chr19 | 11224325 | 33 | 32 | 3205.05754 | 3146.746349 | 0.47028 |
| F2RL3 | chr19 | 16995124 | 32 | 30 | 3109.798535 | 2946.32983 | 0.4703 |
| CDKN2B-AS1 | chr9 | 22005647 | 7 | 3 | 176.0434937 | 139.7649847 | 0.47052 |
| ESR1 | chr6 | 152002424 | 33 | 32 | 2968.417671 | 2939.685583 | 0.4707 |
| LCK | chr1 | 32741689 | 28 | 31 | 2465.824592 | 2664.188369 | 0.4708 |
| APOA1 | chr11 | 116661600 | 29 | 29 | 2589.99542 | 2540.792314 | 0.47082 |
| HMGCR | chr5 | 74627144 | 3 | 3 | 60 | 145 | 0.47114 |
| CDKN2B-AS1 | chr9 | 21980033 | 37 | 33 | 1801.069052 | 1671.471942 | 0.47132 |
| ABCG1 | chr21 | 43635099 | 32 | 32 | 3079.812928 | 3034.644277 | 0.47132 |
| KLF2 | chr19 | 16427475 | 35 | 33 | 2524.829876 | 2450.657683 | 0.47146 |
| APOA1 | chr11 | 116689650 | 25 | 28 | 2298.639971 | 2640.246159 | 0.47148 |
| CDKN2B-AS1 | chr9 | 22006138 | 3 | 3 | 73.52941176 | 143.0769231 | 0.47152 |
| APOA1 | chr11 | 116720034 | 33 | 32 | 3142.065067 | 3083.49645 | 0.47162 |
| APOA1 | chr11 | 116699221 | 33 | 32 | 3058.371795 | 2998.23605 | 0.47164 |
| TERT | chr5 | 1299457 | 26 | 29 | 2335.353535 | 2661.688312 | 0.47196 |
| ABCA1 | chr9 | 107691574 | 3 | 3 | 68.07692308 | 127.5 | 0.47232 |
| CTGF | chr6 | 132274539 | 31 | 31 | 2769.465184 | 2816.420638 | 0.47254 |
| NOS3 | chr7 | 150706375 | 23 | 27 | 2068.173077 | 2496.974969 | 0.47254 |
| APOA1 | chr11 | 116676532 | 4 | 5 | 320 | 436.0606061 | 0.47254 |
| CDKN2B-AS1 | chr9 | 21990027 | 8 | 11 | 271.3331113 | 335.3787879 | 0.47258 |
| F7 | chr13 | 113752146 | 32 | 32 | 2892.985224 | 2953.798733 | 0.47292 |
| LOXL1 | chr15 | 74215170 | 21 | 21 | 649.0664945 | 700.524607 | 0.47298 |
| APOA1 | chr11 | 116676473 | 28 | 29 | 2567.157981 | 2714.535012 | 0.47314 |
| E2F6 | chr2 | 11612427 | 3 | 2 | 65 | 125 | 0.47324 |
| KLF2 | chr19 | 16433233 | 2 | 10 | 106.6666667 | 399.4069854 | 0.47356 |
| LDLR | chr19 | 11224280 | 29 | 31 | 2781.168415 | 3027.678571 | 0.47376 |
| ADRA2C | chr4 | 3760698 | 28 | 31 | 2075.144638 | 2378.537851 | 0.4738 |
| CDKN2B-AS1 | chr9 | 22006194 | 15 | 14 | 428.4792009 | 461.0787896 | 0.47402 |
| ABCB1 | chr7 | 87342865 | 27 | 30 | 1170.605889 | 1389.158166 | 0.47406 |
| LDLR | chr19 | 11222366 | 2 | 3 | 53.33333333 | 98.33333333 | 0.4742 |
| ADRA2C | chr4 | 3768437 | 2 | 3 | 40 | 125.5555556 | 0.47442 |
| ADRA2C | chr4 | 3768746 | 24 | 27 | 1104.242979 | 1164.575432 | 0.47452 |
| APOA1 | chr11 | 116673512 | 32 | 29 | 2999.946433 | 2640.909523 | 0.47456 |

| | | | | | | | |
|---|---|---|---|---|---|---|---|
| ALDOA | chr16 | 30078235 | 2 | 2 | 53.33333333 | 122.2222222 | 0.47472 |
| ALDOA | chr16 | 30062216 | 33 | 33 | 2774.25484 | 2706.751704 | 0.47484 |
| ABCA1 | chr9 | 107697728 | 2 | 2 | 50 | 120 | 0.47536 |
| FOXP3 | chrX | 49125647 | 3 | 2 | 109.6240602 | 125 | 0.47552 |
| CDKN2B-AS1 | chr9 | 21996911 | 15 | 17 | 668.8888889 | 681.6233766 | 0.4756 |
| ATF3 | chr1 | 212745345 | 28 | 31 | 2874.979266 | 3086.752137 | 0.47562 |
| FOXP3 | chrX | 49121053 | 3 | 4 | 250 | 380 | 0.47598 |
| NOS3 | chr7 | 150693947 | 10 | 8 | 331.3636364 | 324.3650794 | 0.476 |
| HMGCR | chr5 | 74624619 | 32 | 33 | 2807.926396 | 2828.239084 | 0.47604 |
| APOA1 | chr11 | 116683521 | 2 | 3 | 185.7142857 | 240 | 0.47612 |
| TIMP3 | chr22 | 33190394 | 34 | 32 | 2981.055541 | 2852.641403 | 0.47648 |
| ESR1 | chr6 | 152003072 | 9 | 5 | 273.2707612 | 188.8141528 | 0.47658 |
| FGF2 | chr4 | 123745951 | 2 | 2 | 50 | 120 | 0.4768 |
| CDKN2BAS | chr9 | 22103886 | 38 | 32 | 3561.225074 | 3029.343919 | 0.47688 |
| LPA | chr6 | 160948143 | 30 | 31 | 2919.659119 | 2980.007272 | 0.47692 |
| TERT | chr5 | 1300888 | 11 | 6 | 881.6666667 | 441.8181818 | 0.47724 |
| MCPH1 | chr8 | 6260018 | 6 | 3 | 551.4285714 | 260 | 0.4773 |
| MCPH1 | chr8 | 6263589 | 2 | 2 | 66.66666666 | 128.5714286 | 0.47748 |
| ESR2 | chr14 | 64814347 | 37 | 32 | 3521.60119 | 3027.28135 | 0.47766 |
| CDKN2B-AS1 | chr9 | 21971431 | 7 | 3 | 173.6141054 | 133.3393829 | 0.47768 |
| ARHGEF18 | chr19 | 7466011 | 33 | 32 | 3162.037424 | 3026.614243 | 0.47776 |
| ESR2 | chr14 | 64812934 | 40 | 32 | 3719.793935 | 3019.025924 | 0.47788 |
| F7 | chr13 | 113751018 | 22 | 21 | 2186.547619 | 2021.825397 | 0.47788 |
| E2F6 | chr2 | 11616186 | 38 | 32 | 3389.448644 | 2895.209787 | 0.47804 |
| APOA1 | chr11 | 116686105 | 5 | 6 | 469.047619 | 583.3333333 | 0.47804 |
| ESR1 | chr6 | 152008362 | 33 | 32 | 3131.592302 | 3014.387143 | 0.47812 |
| ESR1 | chr6 | 152008981 | 33 | 31 | 2272.071115 | 2256.894037 | 0.47816 |
| CDKN2B-AS1 | chr9 | 21969248 | 12 | 9 | 312.0054446 | 283.3730159 | 0.47848 |
| CDKN2B-AS1 | chr9 | 21964626 | 22 | 20 | 686.6762978 | 679.7174738 | 0.47886 |
| NOS3 | chr7 | 150706639 | 29 | 30 | 2020.863569 | 1999.805538 | 0.47902 |
| APOA1 | chr11 | 116707073 | 33 | 33 | 2512.547523 | 2435.659771 | 0.47914 |
| LDLR | chr19 | 11220052 | 30 | 31 | 2941.394172 | 2998.978758 | 0.4792 |
| NOS3 | chr7 | 150686728 | 34 | 32 | 3200.568081 | 2952.046809 | 0.47928 |
| LOXL1 | chr15 | 74217341 | 36 | 32 | 3264.322133 | 2865.932416 | 0.47934 |
| APOA1 | chr11 | 116661282 | 32 | 31 | 2817.238227 | 2779.429685 | 0.47934 |
| NOS3 | chr7 | 150695736 | 30 | 29 | 2834.879149 | 2780.050366 | 0.47938 |
| ABCA1 | chr9 | 107693940 | 2 | 2 | 50.79365079 | 120 | 0.47938 |
| CDKN2B-AS1 | chr9 | 21977464 | 17 | 20 | 584.613132 | 752.7869353 | 0.4795 |
| CDKN1A | chr6 | 36645100 | 32 | 33 | 2022.405347 | 2016.036609 | 0.47978 |
| ADRA2C | chr4 | 3764992 | 2 | 3 | 65.45454545 | 169.6969697 | 0.4798 |
| LPA | chr6 | 11219106 | 29 | 28 | 1110.040234 | 1014.688978 | 0.48002 |
| ABCG1 | chr21 | 43630965 | 25 | 23 | 2364.873737 | 2232.777778 | 0.48038 |
| NOS3 | chr7 | 150705848 | 24 | 25 | 745.5572602 | 841.3550876 | 0.4804 |
| APOA1 | chr11 | 116674073 | 33 | 33 | 2057.322078 | 1970.210669 | 0.48058 |
| NOS3 | chr7 | 150691302 | 32 | 33 | 2247.2128 | 2254.796142 | 0.48064 |
| LPA | chr6 | 160952755 | 39 | 32 | 3422.965375 | 2775.966644 | 0.48096 |
| BCL2 | chr18 | 60988466 | 23 | 23 | 680.2384856 | 741.758954 | 0.48104 |

| | | | | | | | |
|---|---|---|---|---|---|---|---|
| CDKN2B-AS1 | chr9 | 22006793 | 21 | 21 | 711.3834776 | 664.9203574 | 0.48114 |
| ALDOA | chr16 | 30062365 | 40 | 32 | 3533.604856 | 2862.431232 | 0.48152 |
| F7 | chr13 | 113775208 | 8 | 6 | 502.5 | 420 | 0.48158 |
| LPA | chr6 | 160942625 | 2 | 2 | 50.79365079 | 117.1428571 | 0.48166 |
| BCL2 | chr18 | 60988590 | 31 | 33 | 1870.391247 | 2066.312104 | 0.48218 |
| F7 | chr13 | 113775436 | 32 | 33 | 2838.803751 | 2865.278788 | 0.48226 |
| CDKN1A | chr6 | 36644902 | 34 | 32 | 3298.822638 | 3071.940842 | 0.48226 |
| LPA | chr6 | 160946754 | 37 | 32 | 3490.427442 | 2990.064865 | 0.48232 |
| LDLR | chr19 | 11216263 | 6 | 4 | 336.3492064 | 281.547619 | 0.48242 |
| GALNT2 | chr1 | 230187458 | 38 | 32 | 3714.894704 | 3089.329368 | 0.4825 |
| FOXP3 | chrX | 49122365 | 32 | 29 | 2894.220071 | 2561.696578 | 0.48262 |
| F2RL3 | chr19 | 17000192 | 8 | 7 | 309.7619048 | 229.4047619 | 0.48272 |
| ADRA2C | chr4 | 3765019 | 3 | 4 | 73.33333333 | 168.0769231 | 0.48288 |
| LCK | chr1 | 32711582 | 5 | 6 | 417.3809524 | 549.047619 | 0.48296 |
| PECAM1 | chr17 | 62398878 | 4 | 4 | 361.1111111 | 330.8333333 | 0.4832 |
| NOS3 | chr7 | 150706066 | 32 | 33 | 1734.447132 | 1694.809436 | 0.48344 |
| COL15A1 | chr9 | 101704325 | 34 | 32 | 3004.398076 | 2856.917611 | 0.48346 |
| TIMP3 | chr22 | 33259066 | 19 | 27 | 1794.047619 | 2500.160118 | 0.4836 |
| ESR1 | chr6 | 152010841 | 32 | 33 | 2992.459207 | 3009.866573 | 0.48362 |
| ABCG1 | chr21 | 43630166 | 33 | 33 | 2000.265511 | 1930.418012 | 0.48368 |
| KLF2 | chr19 | 16427915 | 21 | 19 | 1508.658009 | 1436.756854 | 0.48372 |
| FOXP3 | chrX | 49129714 | 26 | 26 | 1290.264092 | 1364.298259 | 0.4838 |
| NOS3 | chr7 | 150690976 | 2 | 3 | 50 | 131.6483517 | 0.4838 |
| CDKN1A | chr6 | 36651936 | 31 | 32 | 2934.39336 | 2967.522437 | 0.4841 |
| ICAM2 | chr17 | 62080172 | 2 | 6 | 61.9047619 | 268.9285714 | 0.48428 |
| CDKN2B-AS1 | chr9 | 22005396 | 3 | 3 | 74.50166113 | 137.0250896 | 0.48448 |
| F2RL3 | chr19 | 17000072 | 4 | 3 | 100.5769231 | 143.0769231 | 0.48448 |
| FOXP3 | chrX | 49123639 | 16 | 20 | 1512.97619 | 1808.492063 | 0.48466 |
| LPA | chr6 | 11224840 | 3 | 5 | 140 | 268.5714286 | 0.48466 |
| NOS3 | chr7 | 150710245 | 27 | 26 | 1123.256066 | 1159.203248 | 0.48472 |
| FGF2 | chr4 | 123747447 | 18 | 23 | 1468.427406 | 1785.570263 | 0.48484 |
| GCK | chr7 | 44232606 | 24 | 28 | 1444.048166 | 1826.867086 | 0.48488 |
| APOA1 | chr11 | 116700020 | 4 | 4 | 147.2222222 | 113.3333333 | 0.48494 |
| LCK | chr1 | 32707210 | 4 | 4 | 102.7272727 | 151.0147919 | 0.48496 |
| LOXL1 | chr15 | 74218697 | 15 | 20 | 546.871345 | 806.9248003 | 0.48498 |
| AMOTL2 | chr3 | 134095706 | 38 | 32 | 3535.217908 | 3012.032207 | 0.48512 |
| CDKN2B-AS1 | chr9 | 21983786 | 39 | 33 | 2739.384659 | 2257.498547 | 0.48578 |
| GCK | chr7 | 44228746 | 24 | 22 | 707.2161904 | 691.2617184 | 0.48582 |
| PHACTR1 | chr6 | 12709178 | 36 | 33 | 1977.599304 | 1878.210153 | 0.48592 |
| ADRA2C | chr4 | 3768540 | 3 | 2 | 70.19607843 | 118.3333333 | 0.48592 |
| KLF2 | chr19 | 16426656 | 8 | 18 | 371.2698413 | 935.9727772 | 0.48598 |
| CDKN2B-AS1 | chr9 | 21967532 | 9 | 10 | 253.9460784 | 337.6396804 | 0.48598 |
| NOS3 | chr7 | 150707251 | 22 | 27 | 1253.375543 | 1618.437118 | 0.48608 |
| F2RL3 | chr19 | 16998981 | 32 | 33 | 2601.21357 | 2616.556104 | 0.4861 |
| CTGF | chr6 | 132272546 | 2 | 3 | 43.80952381 | 136.6666667 | 0.48614 |
| APOA1 | chr11 | 116674660 | 28 | 32 | 2759.341423 | 3101.951341 | 0.48624 |
| F7 | chr13 | 113775032 | 30 | 30 | 2090.695326 | 2168.97836 | 0.48638 |

| | | | | | | | |
|---|---|---|---|---|---|---|---|
| CDKN2B-AS1 | chr9 | 21989862 | 18 | 23 | 828.968254 | 1175.515873 | 0.48642 |
| CDKN1A | chr6 | 36652077 | 28 | 30 | 2584.52381 | 2831.082251 | 0.48664 |
| APOA1 | chr11 | 116706154 | 24 | 22 | 824.3337108 | 816.6314646 | 0.4867 |
| BAX | chr19 | 49465184 | 32 | 32 | 2183.007136 | 2110.623123 | 0.48698 |
| AMOTL2 | chr3 | 134099340 | 30 | 33 | 1799.963861 | 1886.721874 | 0.48698 |
| CDKN2B-AS1 | chr9 | 21996987 | 35 | 33 | 1836.380952 | 1803.62998 | 0.48704 |
| APOA1 | chr11 | 116716996 | 33 | 32 | 3034.599999 | 2903.352612 | 0.4871 |
| LCK | chr1 | 32708234 | 10 | 13 | 292.2838635 | 441.8108137 | 0.48748 |
| CDKN1A | chr6 | 36650622 | 6 | 6 | 203.2142857 | 176.1904762 | 0.48748 |
| F7 | chr13 | 113751499 | 27 | 28 | 2575.689727 | 2624.969336 | 0.48778 |
| F7 | chr13 | 113775157 | 2 | 3 | 48.57142857 | 150 | 0.48838 |
| PECAM1 | chr17 | 62398840 | 22 | 24 | 2026.522556 | 2254.506152 | 0.4885 |
| LPA | chr6 | 160949887 | 37 | 32 | 3509.645521 | 3015.816183 | 0.48854 |
| GALNT2 | chr1 | 230191043 | 3 | 3 | 160 | 199.2982456 | 0.48868 |
| KLF2 | chr19 | 16427505 | 33 | 33 | 1589.428352 | 1530.901814 | 0.48872 |
| ABCG1 | chr21 | 43632594 | 38 | 33 | 2347.11388 | 2084.655653 | 0.48882 |
| CDKN1A | chr6 | 36650372 | 27 | 29 | 1406.997117 | 1599.525518 | 0.48892 |
| LOXL1 | chr15 | 74215596 | 16 | 22 | 1393.333333 | 1972.489177 | 0.48964 |
| APOA1 | chr11 | 116698917 | 32 | 32 | 3142.518182 | 3055.080375 | 0.48972 |
| ABCG1 | chr21 | 43636522 | 32 | 32 | 2983.828755 | 3018.017213 | 0.48996 |
| NOS3 | chr7 | 150684432 | 12 | 16 | 784.7115385 | 1126.944444 | 0.49002 |
| CDKN2B-AS1 | chr9 | 21976844 | 35 | 33 | 2122.94183 | 1928.271508 | 0.49012 |
| ALDOA | chr16 | 30062955 | 32 | 32 | 3044.262709 | 3074.11999 | 0.49012 |
| CTGF | chr6 | 132274165 | 35 | 32 | 3157.81773 | 2855.998049 | 0.49038 |
| CDKN2B-AS1 | chr9 | 22005440 | 4 | 2 | 84.4568869 | 116.3398693 | 0.49042 |
| CDKN2B-AS1 | chr9 | 21999740 | 2 | 2 | 60 | 125 | 0.49048 |
| CDKN1A | chr6 | 36648991 | 26 | 28 | 2271.533792 | 2377.579643 | 0.4905 |
| APOA1 | chr11 | 116719012 | 15 | 14 | 1313.552754 | 1273.888889 | 0.49066 |
| LDLR | chr19 | 11191841 | 8 | 8 | 734.2424242 | 766.6666667 | 0.49072 |
| BAX | chr19 | 49456215 | 27 | 25 | 2091.490038 | 2012.481962 | 0.49078 |
| APOA1 | chr11 | 116717720 | 33 | 32 | 3071.720248 | 3018.121587 | 0.49086 |
| ABCG1 | chr21 | 43717443 | 2 | 2 | 42.22222222 | 122.2222222 | 0.49116 |
| F7 | chr13 | 113750842 | 32 | 31 | 3230.032657 | 3028.643579 | 0.49132 |
| FOXP3 | chrX | 49120319 | 8 | 9 | 664.1233766 | 795.7142857 | 0.4914 |
| LCK | chr1 | 32740505 | 23 | 26 | 1920.355021 | 2107.348762 | 0.49142 |
| GALNT2 | chr1 | 230193710 | 28 | 25 | 1707.020931 | 1443.157814 | 0.49154 |
| LDLR | chr19 | 11221443 | 34 | 32 | 3311.22249 | 3083.066316 | 0.49206 |
| APOA1 | chr11 | 116680222 | 30 | 31 | 2534.937411 | 2544.783016 | 0.49206 |
| PLA2G7 | chr6 | 46703469 | 21 | 19 | 582.8683626 | 575.4736259 | 0.4922 |
| APOA1 | chr11 | 116723396 | 4 | 2 | 96.06060606 | 117.8571429 | 0.49228 |
| PECAM1 | chr17 | 62398684 | 14 | 12 | 361.0507793 | 357.6370984 | 0.49234 |
| F7 | chr13 | 113760502 | 23 | 26 | 973.9532597 | 1185.087413 | 0.49252 |
| CDKN1A | chr6 | 36653611 | 26 | 22 | 1070.672006 | 838.4099628 | 0.49296 |
| TMPO | chr12 | 98907214 | 39 | 32 | 3729.964954 | 3042.865376 | 0.4931 |
| ATF3 | chr1 | 212779795 | 3 | 2 | 113.3333333 | 125 | 0.49368 |
| LDLR | chr19 | 11224013 | 24 | 26 | 2165.753968 | 2432.5 | 0.49378 |
| ICAM2 | chr17 | 62080148 | 33 | 32 | 3206.50873 | 3069.041819 | 0.494 |

| | | | | | | | |
|---|---|---|---|---|---|---|---|
| LOXL1 | chr15 | 74244445 | 2 | 2 | 53.57142857 | 120 | 0.49404 |
| APOA1 | chr11 | 116661264 | 32 | 33 | 2741.230665 | 2756.311856 | 0.49408 |
| LCK | chr1 | 32712557 | 20 | 25 | 1609.345654 | 1947.104562 | 0.4941 |
| BAX | chr19 | 49456814 | 30 | 29 | 2903.620731 | 2772.073488 | 0.49412 |
| TIMP3 | chr22 | 33190486 | 27 | 26 | 2219.908702 | 2213.485616 | 0.49442 |
| CDKN2B-AS1 | chr9 | 21968763 | 5 | 5 | 126.4371903 | 179.3029001 | 0.49452 |
| CDKN2B-AS1 | chr9 | 22005576 | 20 | 16 | 545.616271 | 486.2043002 | 0.4946 |
| F7 | chr13 | 113752641 | 32 | 31 | 3013.22809 | 2954.770507 | 0.49476 |
| ABCA1 | chr9 | 107542807 | 17 | 15 | 873.6157797 | 835.3391053 | 0.4949 |
| ESR1 | chr6 | 152007591 | 14 | 14 | 1220.47619 | 1265 | 0.49494 |
| NR3C1 | chr5 | 142789685 | 41 | 33 | 2485.996164 | 1948.58815 | 0.49496 |
| TERT | chr5 | 1300296 | 31 | 31 | 2848.67227 | 2897.348675 | 0.49502 |
| TFPI | chr2 | 188424121 | 3 | 2 | 285.7142857 | 200 | 0.49502 |
| LCK | chr1 | 32712353 | 2 | 3 | 40 | 140 | 0.49502 |
| APOA1 | chr11 | 116674391 | 2 | 3 | 40 | 140 | 0.49502 |
| APOA1 | chr11 | 116701105 | 2 | 3 | 40 | 140 | 0.49502 |
| PECAM1 | chr17 | 62408462 | 2 | 3 | 40 | 140 | 0.49502 |
| E2F6 | chr2 | 11612628 | 2 | 3 | 40 | 120 | 0.49502 |
| ABCG1 | chr21 | 43637162 | 2 | 3 | 40 | 125.7142857 | 0.49502 |
| APOA1 | chr11 | 116715923 | 3 | 2 | 68.57142857 | 40 | 0.49502 |
| NOS3 | chr7 | 150685996 | 3 | 2 | 73.33333333 | 40 | 0.49502 |
| NOS3 | chr7 | 150710695 | 2 | 3 | 40 | 67.27272727 | 0.49502 |
| APOA1 | chr11 | 116691666 | 3 | 2 | 62.22222222 | 40 | 0.49502 |
| NOS3 | chr7 | 150706776 | 3 | 2 | 62.22222222 | 40 | 0.49502 |
| GALNT2 | chr1 | 230184267 | 18 | 21 | 1134.365079 | 1421.893939 | 0.49516 |
| CDKN2B-AS1 | chr9 | 21968360 | 3 | 3 | 73.7012987 | 140 | 0.49518 |
| ALOX15 | chr17 | 4547220 | 37 | 32 | 2446.857911 | 2178.882903 | 0.49538 |
| CDKN2B-AS1 | chr9 | 22011076 | 4 | 2 | 84.34044882 | 116.3584637 | 0.49538 |
| FOXP3 | chrX | 49114989 | 12 | 4 | 1103.968254 | 346.6666667 | 0.4954 |
| APOA1 | chr11 | 116705089 | 18 | 20 | 1710.416667 | 1937.188645 | 0.4955 |
| LDLR | chr19 | 11219930 | 8 | 8 | 718.0952381 | 747.5 | 0.49554 |
| NOS3 | chr7 | 150686557 | 35 | 33 | 2358.080955 | 2172.380148 | 0.49568 |
| APOA1 | chr11 | 116678512 | 16 | 15 | 1478.131313 | 1417.380952 | 0.49572 |
| APOA1 | chr11 | 116707223 | 3 | 2 | 65 | 122.2222222 | 0.49578 |
| CTGF | chr6 | 132272334 | 3 | 6 | 74.76190476 | 229.7222222 | 0.49644 |
| LPA | chr6 | 11223909 | 33 | 32 | 3215.740387 | 3137.865613 | 0.49656 |
| NOS3 | chr7 | 150690881 | 6 | 10 | 580 | 923.3333333 | 0.4966 |
| NOS3 | chr7 | 150681977 | 32 | 33 | 2698.622426 | 2717.928177 | 0.49672 |
| TERT | chr5 | 1296105 | 32 | 33 | 2735.387275 | 2741.426013 | 0.49704 |
| ALDOA | chr16 | 30061617 | 18 | 12 | 1712.261905 | 1098.928571 | 0.49706 |
| ESR1 | chr6 | 152002295 | 30 | 32 | 2596.700922 | 2837.461258 | 0.49708 |
| FOXP3 | chrX | 49130593 | 28 | 24 | 2609.482601 | 2273.412698 | 0.4973 |
| LDLR | chr19 | 11192691 | 19 | 22 | 1845.075758 | 2077.29798 | 0.4973 |
| APOA1 | chr11 | 116694559 | 2 | 2 | 53.33333333 | 120 | 0.49748 |
| PHACTR1 | chr6 | 13276352 | 2 | 2 | 53.33333333 | 120 | 0.49748 |
| LOXL1 | chr15 | 74244246 | 7 | 9 | 439.8809524 | 507.3015873 | 0.4975 |
| APOA1 | chr11 | 116703029 | 36 | 32 | 3252.76043 | 2928.953567 | 0.49754 |

| | | | | | | | |
|---|---|---|---|---|---|---|---|
| ESR1 | chr6 | 152008189 | 30 | 32 | 2909.859765 | 3075.7846 | 0.49776 |
| LPA | chr6 | 11218654 | 34 | 32 | 3166.987184 | 3008.564146 | 0.4981 |
| CDKN2B-AS1 | chr9 | 22006199 | 3 | 4 | 89.17748917 | 170 | 0.49816 |
| LPA | chr6 | 160942982 | 36 | 32 | 3443.734621 | 3026.155665 | 0.4982 |
| LDLR | chr19 | 11221235 | 30 | 30 | 2892.197247 | 2923.131689 | 0.49824 |
| TERT | chr5 | 1299576 | 18 | 13 | 1701.275946 | 1256.547619 | 0.4983 |
| LPA | chr6 | 11216604 | 34 | 31 | 3270.727633 | 2957.153671 | 0.49838 |
| ESR1 | chr6 | 152003548 | 34 | 32 | 3107.060851 | 2889.8449 | 0.49844 |
| LDLR | chr19 | 11190180 | 12 | 13 | 1056.269841 | 1092.380952 | 0.4989 |
| CDKN2B-AS1 | chr9 | 21989730 | 14 | 9 | 356.4191967 | 271.8653278 | 0.49894 |
| CDKN2B-AS1 | chr9 | 21977434 | 12 | 10 | 338.5227273 | 315.5555556 | 0.49914 |
| ADRA2C | chr4 | 3767118 | 3 | 2 | 66.42857143 | 109.7222222 | 0.49956 |
| GCK | chr7 | 44232825 | 12 | 17 | 912.2807018 | 1382.857143 | 0.49958 |
| F7 | chr13 | 113759913 | 33 | 29 | 3079.213633 | 2743.41387 | 0.49994 |
| ADRA2C | chr4 | 3759759 | 32 | 32 | 2810.572838 | 2857.851494 | 0.49998 |
| FOXP3 | chrX | 49127284 | 18 | 19 | 1638.650794 | 1777.056277 | 0.49998 |
| LDLR | chr19 | 11220409 | 3 | 3 | 300 | 280 | 0.5 |
| ATF3 | chr1 | 212745580 | 3 | 2 | 300 | 140 | 0.5 |
| APOA1 | chr11 | 116701181 | 3 | 3 | 60 | 140 | 0.5 |
| TMPO | chr12 | 98904883 | 5 | 2 | 500 | 180 | 0.5 |
| LDLR | chr19 | 11196578 | 4 | 2 | 400 | 180 | 0.5 |
| LDLR | chr19 | 11220406 | 3 | 2 | 300 | 180 | 0.5 |
| KLF2 | chr19 | 16432571 | 4 | 2 | 400 | 180 | 0.5 |
| AKT2 | chr19 | 40795535 | 3 | 3 | 60 | 140 | 0.5 |
| AMOTL2 | chr3 | 134094996 | 3 | 2 | 60 | 120 | 0.5 |
| AMOTL2 | chr3 | 134102040 | 3 | 2 | 300 | 180 | 0.5 |
| TERT | chr5 | 1298673 | 3 | 2 | 60 | 120 | 0.5 |
| LPA | chr6 | 160950217 | 3 | 2 | 60 | 120 | 0.5 |
| LOXL1 | chr15 | 74219196 | 17 | 16 | 647.7966893 | 666.4318172 | 0.50002 |
| LDLR | chr19 | 11193081 | 4 | 2 | 363.3333333 | 160 | 0.50014 |
| ABCA1 | chr9 | 107696077 | 2 | 4 | 40 | 145 | 0.50014 |
| NOS3 | chr7 | 150688556 | 38 | 32 | 3396.912767 | 2820.78171 | 0.50022 |
| ESR1 | chr6 | 152002777 | 14 | 13 | 381.9967621 | 406.6291822 | 0.50026 |
| LDLR | chr19 | 11224638 | 13 | 14 | 1158.333333 | 1203.809524 | 0.50038 |
| AKT2 | chr19 | 40800086 | 32 | 32 | 2565.018029 | 2497.103821 | 0.50046 |
| TERT | chr5 | 1299050 | 31 | 31 | 2969.851599 | 2928.450361 | 0.50058 |
| ESR1 | chr6 | 152002856 | 12 | 12 | 328.4992776 | 384.3083565 | 0.50062 |
| TERT | chr5 | 1299698 | 23 | 24 | 2179.60817 | 2224.678932 | 0.50074 |
| CDKN2B-AS1 | chr9 | 21968176 | 9 | 4 | 264.8214286 | 150.0619788 | 0.50074 |
| APOA1 | chr11 | 116662304 | 2 | 2 | 42.26190476 | 109.9415205 | 0.50074 |
| APOA1 | chr11 | 116673603 | 26 | 27 | 2433.214225 | 2576.25 | 0.50124 |
| PHACTR1 | chr6 | 12717874 | 38 | 33 | 2604.336837 | 2313.069305 | 0.5018 |
| PECAM1 | chr17 | 62401970 | 8 | 6 | 717.9365079 | 565.7142857 | 0.50186 |
| CDKN2B-AS1 | chr9 | 22005509 | 8 | 2 | 193.9649123 | 100.8045977 | 0.5019 |
| CDKN2B-AS1 | chr9 | 21974580 | 4 | 3 | 102 | 130.5769231 | 0.50214 |
| ADRA2C | chr4 | 3762312 | 33 | 32 | 2837.660937 | 2794.199659 | 0.50216 |
| CDKN2B-AS1 | chr9 | 21990278 | 14 | 10 | 373.588966 | 310.9607891 | 0.50216 |

| | | | | | | | |
|---|---|---|---|---|---|---|---|
| CDKN2B-AS1 | chr9 | 21990318 | 18 | 10 | 484.657738 | 306.1098807 | 0.5022 |
| CDKN2B-AS1 | chr9 | 21971024 | 3 | 3 | 84.44444444 | 120 | 0.5023 |
| ESR2 | chr14 | 64809000 | 26 | 27 | 2014.604069 | 2173.730159 | 0.50244 |
| NOS3 | chr7 | 150706268 | 33 | 32 | 2873.996276 | 2737.117256 | 0.50254 |
| NOS3 | chr7 | 150707044 | 31 | 33 | 2711.212969 | 2820.376358 | 0.50272 |
| FOXP3 | chrX | 49127091 | 18 | 16 | 1501.742424 | 1268.928571 | 0.5028 |
| MCPH1 | chr8 | 6470230 | 14 | 15 | 1432.282913 | 1480 | 0.50324 |
| TMPO | chr12 | 98903392 | 36 | 32 | 3466.655143 | 3102.3376 | 0.50328 |
| ARHGEF18 | chr19 | 7469590 | 31 | 32 | 2985.335361 | 3106.419858 | 0.50346 |
| NOS3 | chr7 | 150692387 | 32 | 29 | 2562.844297 | 2387.777702 | 0.5036 |
| TERT | chr5 | 1296422 | 35 | 32 | 3070.379271 | 2766.857998 | 0.50362 |
| FOXP3 | chrX | 49106612 | 4 | 5 | 363.3333333 | 420 | 0.50362 |
| P2RY12 | chr3 | 151105443 | 34 | 27 | 1338.660251 | 1108.841897 | 0.50388 |
| APOA1 | chr11 | 116670792 | 38 | 32 | 3476.016323 | 2870.9721 | 0.504 |
| F2RL3 | chr19 | 16990332 | 32 | 32 | 3004.372674 | 3049.098402 | 0.50402 |
| NOS3 | chr7 | 150689835 | 20 | 24 | 1737.63986 | 2006.150794 | 0.50406 |
| LPA | chr6 | 11224993 | 31 | 31 | 2337.770038 | 2419.861294 | 0.50454 |
| ABCA1 | chr9 | 107696191 | 38 | 33 | 2792.393761 | 2376.216194 | 0.50466 |
| TERT | chr5 | 1296127 | 32 | 33 | 2359.828065 | 2504.583104 | 0.50482 |
| GALNT2 | chr1 | 230193513 | 32 | 33 | 2164.049368 | 2298.024813 | 0.50498 |
| ESR1 | chr6 | 152002746 | 11 | 11 | 300.1943155 | 352.4964838 | 0.50506 |
| TERT | chr5 | 1298966 | 32 | 30 | 2651.221993 | 2407.643236 | 0.50514 |
| CDKN1A | chr6 | 36643727 | 12 | 10 | 1067.472527 | 929.3650794 | 0.50514 |
| TERT | chr5 | 1296187 | 32 | 32 | 2911.767688 | 2947.437483 | 0.50516 |
| KLF2 | chr19 | 16432772 | 34 | 33 | 3091.372537 | 3067.171124 | 0.50526 |
| F7 | chr13 | 113758488 | 32 | 32 | 2977.82299 | 3025.561495 | 0.50556 |
| CDKN2B-AS1 | chr9 | 21971412 | 2 | 2 | 41.21212121 | 110.625 | 0.50556 |
| FOXP3 | chrX | 49126818 | 8 | 4 | 253.3333333 | 160 | 0.50586 |
| ABCA1 | chr9 | 107696093 | 19 | 19 | 692.670173 | 751.3239538 | 0.5061 |
| LCK | chr1 | 32740717 | 3 | 2 | 68.57142857 | 122.2222222 | 0.5061 |
| GCK | chr7 | 44228820 | 35 | 32 | 3180.170263 | 2947.449464 | 0.50622 |
| F7 | chr13 | 113751091 | 11 | 9 | 958.4126984 | 816.547619 | 0.50626 |
| APOA1 | chr11 | 116662274 | 5 | 2 | 114.3703704 | 112.5448029 | 0.50628 |
| ADRA2C | chr4 | 3760325 | 36 | 32 | 2438.794708 | 2220.036756 | 0.5065 |
| CDKN2B-AS1 | chr9 | 21966533 | 17 | 18 | 526.9872831 | 600.3846154 | 0.5065 |
| NR3C1 | chr5 | 142790973 | 7 | 9 | 573.8888889 | 780.3571429 | 0.5065 |
| ABCA1 | chr9 | 107693935 | 26 | 28 | 1973.109084 | 1991.413898 | 0.50668 |
| ALDOA | chr16 | 30062388 | 40 | 32 | 3529.749519 | 2860.228168 | 0.50692 |
| APOA1 | chr11 | 116707770 | 31 | 30 | 3060.531968 | 2902.307692 | 0.50692 |
| ADRA2C | chr4 | 3763757 | 33 | 33 | 2236.085361 | 2311.101428 | 0.50742 |
| NOS3 | chr7 | 150694556 | 28 | 30 | 2083.723884 | 2305.905483 | 0.5075 |
| LPA | chr6 | 160943827 | 10 | 11 | 899.476912 | 947.2619048 | 0.50766 |
| ADRA2C | chr4 | 3766458 | 34 | 33 | 2440.636542 | 2440.142845 | 0.50776 |
| LOXL1 | chr15 | 74218894 | 31 | 27 | 1238.765611 | 1012.848751 | 0.508 |
| ALDOA | chr16 | 30078133 | 2 | 3 | 43.80952381 | 138.5185185 | 0.50818 |
| APOA1 | chr11 | 116670506 | 33 | 32 | 2736.804379 | 2717.031746 | 0.50832 |
| ABCA1 | chr9 | 107690763 | 13 | 15 | 574.047619 | 724.4444445 | 0.50844 |

| | | | | | | | |
|---|---|---|---|---|---|---|---|
| PHACTR1 | chr6 | 12714865 | 33 | 32 | 3000.730865 | 2951.097163 | 0.5086 |
| DHFR | chr5 | 79955780 | 33 | 32 | 3048.996491 | 2917.990305 | 0.50878 |
| CDKN2B-AS1 | chr9 | 21970731 | 7 | 8 | 190.2932035 | 267.8969269 | 0.50884 |
| GCK | chr7 | 44233109 | 33 | 32 | 2889.606252 | 2850.12816 | 0.50894 |
| TIMP3 | chr22 | 33195667 | 33 | 32 | 3172.193225 | 3015.064077 | 0.50906 |
| F7 | chr13 | 113752208 | 2 | 2 | 42.22222222 | 120 | 0.50908 |
| F7 | chr13 | 113758166 | 2 | 2 | 42.22222222 | 120 | 0.50908 |
| PECAM1 | chr17 | 62401726 | 2 | 2 | 42.22222222 | 120 | 0.50908 |
| LDLR | chr19 | 11196028 | 2 | 2 | 42.22222222 | 120 | 0.50908 |
| LDLR | chr19 | 11198500 | 2 | 2 | 42.22222222 | 120 | 0.50908 |
| BAX | chr19 | 49456524 | 2 | 2 | 42.22222222 | 120 | 0.50908 |
| AMOTL2 | chr3 | 134097304 | 2 | 2 | 42.22222222 | 120 | 0.50908 |
| FOXP3 | chrX | 49115550 | 2 | 2 | 42.22222222 | 120 | 0.50908 |
| FOXP3 | chrX | 49127941 | 2 | 2 | 42.22222222 | 120 | 0.50908 |
| TERT | chr5 | 1296233 | 34 | 33 | 3034.690998 | 2887.902694 | 0.50922 |
| NOS3 | chr7 | 150705893 | 14 | 17 | 414.5258592 | 556.2478491 | 0.50936 |
| COL15A1 | chr9 | 101700479 | 34 | 32 | 3214.153235 | 3056.808985 | 0.50938 |
| ADRA2C | chr4 | 3758329 | 32 | 32 | 2036.237737 | 1957.144996 | 0.50942 |
| ALOX15 | chr17 | 4549514 | 32 | 33 | 2815.204469 | 2836.431186 | 0.5098 |
| ABCG1 | chr21 | 43633746 | 2 | 2 | 42.22222222 | 114.1176471 | 0.50984 |
| LOXL1 | chr15 | 74219207 | 32 | 30 | 1448.156212 | 1273.109367 | 0.51002 |
| ESR1 | chr6 | 152003123 | 6 | 8 | 150.9465036 | 247.0138024 | 0.51016 |
| ICAM2 | chr17 | 62080144 | 33 | 32 | 3204.248202 | 3074.815904 | 0.51024 |
| F2RL3 | chr19 | 16990992 | 23 | 25 | 2089.383394 | 2315.121545 | 0.51024 |
| E2F6 | chr2 | 11607107 | 37 | 33 | 2423.019003 | 2228.112309 | 0.51026 |
| GALNT2 | chr1 | 230185315 | 37 | 32 | 3300.05114 | 2889.858231 | 0.51036 |
| ADRA2C | chr4 | 3766656 | 19 | 19 | 1537.004662 | 1470.764791 | 0.51038 |
| PLA2G7 | chr6 | 46706158 | 29 | 28 | 2362.725379 | 2216.906149 | 0.51044 |
| GALNT2 | chr1 | 230183622 | 34 | 32 | 3104.727522 | 2948.66995 | 0.51064 |
| ESR1 | chr6 | 152011200 | 39 | 32 | 3551.349508 | 2882.992351 | 0.51078 |
| PLA2G7 | chr6 | 46707523 | 34 | 32 | 2844.921016 | 2729.565458 | 0.5109 |
| NOS3 | chr7 | 150705872 | 33 | 33 | 1794.564671 | 1716.32998 | 0.5112 |
| CDKN1A | chr6 | 36646127 | 9 | 12 | 252.6872798 | 383.7302238 | 0.51148 |
| FOXP3 | chrX | 49119260 | 2 | 2 | 53.33333333 | 108.8888889 | 0.5115 |
| ADRA2C | chr4 | 3761346 | 24 | 25 | 1745.855584 | 1713.486545 | 0.51152 |
| LCK | chr1 | 32716051 | 3 | 5 | 78.33333333 | 186.3415893 | 0.51184 |
| APOA1 | chr11 | 116671442 | 39 | 32 | 3773.718929 | 3109.945791 | 0.51206 |
| APOA1 | chr11 | 116675401 | 30 | 31 | 2758.792125 | 2888.165768 | 0.51206 |
| MCPH1 | chr8 | 6470052 | 17 | 17 | 1650.32967 | 1623.690476 | 0.51212 |
| NOS3 | chr7 | 150695705 | 27 | 28 | 2591.850649 | 2646.155789 | 0.51216 |
| CTGF | chr6 | 132272356 | 2 | 2 | 42.22222222 | 100 | 0.51222 |
| CDKN2B-AS1 | chr9 | 21971043 | 2 | 2 | 42.22222222 | 100 | 0.51222 |
| NOS3 | chr7 | 150710618 | 9 | 8 | 265.9090909 | 288.7201687 | 0.51226 |
| APOA1 | chr11 | 116692261 | 34 | 32 | 3223.519825 | 3003.280816 | 0.51254 |
| APOA1 | chr11 | 116662503 | 14 | 20 | 813.0952381 | 1256.269841 | 0.51272 |
| MIA3 | chr1 | 222817152 | 2 | 2 | 43.07692308 | 120 | 0.51272 |
| APOA1 | chr11 | 116694353 | 2 | 2 | 43.07692308 | 120 | 0.51272 |

| | | | | | | | |
|---|---|---|---|---|---|---|---|
| HMGCR | chr5 | 74628451 | 2 | 2 | 43.07692308 | 120 | 0.51272 |
| LPA | chr6 | 11224390 | 2 | 2 | 43.07692308 | 120 | 0.51272 |
| COL15A1 | chr9 | 101702053 | 2 | 2 | 43.07692308 | 120 | 0.51272 |
| COL15A1 | chr9 | 101704884 | 2 | 2 | 43.07692308 | 120 | 0.51272 |
| F7 | chr13 | 113750133 | 17 | 24 | 928.217681 | 1425.135976 | 0.5128 |
| ALOX15 | chr17 | 4552959 | 33 | 33 | 2512.417899 | 2463.548407 | 0.51286 |
| GALNT2 | chr1 | 230191774 | 17 | 8 | 1344.484127 | 669.8809524 | 0.51304 |
| TMPO | chr12 | 98908170 | 35 | 33 | 1544.029896 | 1404.07138 | 0.51308 |
| FOXP3 | chrX | 49127123 | 18 | 22 | 1705.726496 | 2119.722222 | 0.51324 |
| CDKN2B-AS1 | chr9 | 21996008 | 2 | 2 | 52.99145299 | 103.3333333 | 0.51338 |
| MCPH1 | chr8 | 6469995 | 8 | 3 | 760 | 266.6666667 | 0.51352 |
| APOA1 | chr11 | 116692129 | 33 | 32 | 3012.837796 | 2955.589819 | 0.51366 |
| MIA3 | chr1 | 222792359 | 9 | 9 | 266.6659072 | 318.8639792 | 0.51384 |
| LDLR | chr19 | 11190599 | 2 | 3 | 42.22222222 | 130.9090909 | 0.51404 |
| CDKN2B-AS1 | chr9 | 21975682 | 5 | 2 | 119.3756244 | 122.2222222 | 0.51404 |
| CDKN2B-AS1 | chr9 | 21993498 | 40 | 33 | 2395.00952 | 1924.998144 | 0.51412 |
| GCK | chr7 | 44237650 | 33 | 32 | 2658.215939 | 2626.136517 | 0.51436 |
| ABCG1 | chr21 | 43629437 | 34 | 32 | 2749.132063 | 2542.182497 | 0.51446 |
| F7 | chr13 | 113750737 | 27 | 26 | 2503.611111 | 2461.428571 | 0.5145 |
| APOA1 | chr11 | 116674238 | 29 | 30 | 2739.781145 | 2868.905498 | 0.51488 |
| FOXP3 | chrX | 49125974 | 2 | 2 | 65 | 125 | 0.51492 |
| CTGF | chr6 | 132278861 | 32 | 32 | 2988.214176 | 2925.806318 | 0.5153 |
| CDKN2B-AS1 | chr9 | 21993603 | 27 | 20 | 893.9663021 | 619.5155172 | 0.51542 |
| ADRA2C | chr4 | 3759165 | 32 | 32 | 2701.608941 | 2778.244518 | 0.51546 |
| LPA | chr6 | 11221856 | 36 | 32 | 2834.243482 | 2639.342509 | 0.51594 |
| E2F6 | chr2 | 11613872 | 26 | 28 | 2290.165368 | 2517.724061 | 0.51626 |
| CDKN2BAS | chr9 | 22098457 | 32 | 32 | 2749.819689 | 2801.422833 | 0.51638 |
| FGF2 | chr4 | 123743987 | 35 | 32 | 3258.567346 | 2949.606992 | 0.5164 |
| NOS3 | chr7 | 150692178 | 19 | 20 | 659.3365108 | 749.5922512 | 0.51676 |
| ADRA2C | chr4 | 3765335 | 9 | 12 | 628.452381 | 770.6759907 | 0.51692 |
| PECAM1 | chr17 | 62398535 | 2 | 3 | 45 | 142.2222222 | 0.51698 |
| CDKN2B-AS1 | chr9 | 21971190 | 11 | 11 | 337.8944911 | 308.7490287 | 0.51704 |
| FGF2 | chr4 | 123739112 | 17 | 16 | 1458.769841 | 1427.103175 | 0.51706 |
| F7 | chr13 | 113750595 | 26 | 28 | 2509.88272 | 2664.707792 | 0.51722 |
| ALOX15 | chr17 | 4547886 | 6 | 7 | 517.5 | 643.3333333 | 0.51744 |
| HMGCR | chr5 | 74624589 | 35 | 32 | 2849.616876 | 2652.537255 | 0.51774 |
| TFPI | chr2 | 188419934 | 37 | 32 | 3389.795301 | 2902.880834 | 0.5182 |
| CDKN2B-AS1 | chr9 | 21974466 | 3 | 2 | 66.05263158 | 112.8571429 | 0.5184 |
| ADRA2C | chr4 | 3764557 | 31 | 32 | 2725.496189 | 2875.893488 | 0.51842 |
| APOA1 | chr11 | 116718522 | 7 | 13 | 301.7735535 | 604.3478905 | 0.51854 |
| NOS3 | chr7 | 150711165 | 12 | 12 | 366.938196 | 421.4500996 | 0.51862 |
| ADRA2C | chr4 | 3763836 | 33 | 33 | 2908.520529 | 2838.598976 | 0.51866 |
| NOS3 | chr7 | 150695139 | 27 | 28 | 2584.02417 | 2585.159008 | 0.51882 |
| NOS3 | chr7 | 150706128 | 32 | 33 | 1826.586761 | 1964.926236 | 0.51908 |
| CDKN2B-AS1 | chr9 | 22005367 | 3 | 4 | 83.39335733 | 161.3748423 | 0.51912 |
| APOA1 | chr11 | 116687545 | 2 | 2 | 55 | 107.5 | 0.51912 |
| CDKN2B-AS1 | chr9 | 21967370 | 4 | 6 | 100.5555556 | 202.296798 | 0.51914 |

| | | | | | | | |
|---|---|---|---|---|---|---|---|
| TIMP3 | chr22 | 33192380 | 35 | 32 | 3386.257618 | 3077.607612 | 0.51924 |
| LOXL1 | chr15 | 74215049 | 29 | 24 | 986.2772125 | 868.5885273 | 0.51928 |
| F7 | chr13 | 113750392 | 22 | 21 | 1936.030473 | 1799.458042 | 0.51944 |
| APOA1 | chr11 | 116704339 | 34 | 32 | 3448.65094 | 3118.64983 | 0.51944 |
| FOXP3 | chrX | 49121718 | 8 | 6 | 743.2142857 | 534.1666667 | 0.5195 |
| MIA3 | chr1 | 222791614 | 2 | 2 | 60 | 122.2222222 | 0.51952 |
| LCK | chr1 | 32713425 | 33 | 32 | 3110.050177 | 3055.363935 | 0.51968 |
| PECAM1 | chr17 | 62401606 | 32 | 32 | 2000.712554 | 2064.876508 | 0.51972 |
| APOA1 | chr11 | 116699052 | 15 | 17 | 1367.387057 | 1586.060606 | 0.5198 |
| AMOTL2 | chr3 | 134095542 | 3 | 2 | 67.22222222 | 120 | 0.51996 |
| AMOTL2 | chr3 | 134095544 | 3 | 2 | 67.22222222 | 120 | 0.51996 |
| HMGCR | chr5 | 74627583 | 39 | 33 | 2886.602637 | 2497.053778 | 0.51998 |
| LDLR | chr19 | 11222129 | 31 | 32 | 2842.131618 | 2859.702967 | 0.51998 |
| ESR1 | chr6 | 152010376 | 38 | 32 | 3342.48596 | 2741.864626 | 0.5201 |
| ADRA2C | chr4 | 3766209 | 34 | 32 | 2776.722161 | 2658.542461 | 0.52022 |
| ESR2 | chr14 | 64693809 | 35 | 33 | 2643.28066 | 2431.158867 | 0.5203 |
| APOA1 | chr11 | 116668619 | 32 | 32 | 3125.892546 | 3101.627566 | 0.52046 |
| LCK | chr1 | 32712780 | 32 | 31 | 2852.463545 | 2804.64468 | 0.52054 |
| BAX | chr19 | 49454643 | 31 | 30 | 2626.977931 | 2486.067244 | 0.52072 |
| LCK | chr1 | 32740385 | 26 | 28 | 2369.836543 | 2595.73478 | 0.52102 |
| CTGF | chr6 | 132281865 | 28 | 30 | 1775.543288 | 2010.417794 | 0.52104 |
| GCK | chr7 | 44230056 | 32 | 32 | 2733.505385 | 2776.631613 | 0.5211 |
| APOA1 | chr11 | 116665861 | 12 | 20 | 1084.464286 | 1751.904762 | 0.52128 |
| DHFR | chr5 | 79954158 | 25 | 22 | 2000.057457 | 1700.642136 | 0.5213 |
| F2RL3 | chr19 | 16995385 | 14 | 11 | 1177.525946 | 975.8333333 | 0.52136 |
| AMOTL2 | chr3 | 134094236 | 34 | 33 | 2230.180424 | 2111.19098 | 0.52184 |
| TIMP3 | chr22 | 33196103 | 33 | 33 | 2870.022623 | 2813.356332 | 0.52186 |
| GALNT2 | chr1 | 230418324 | 32 | 32 | 2140.477353 | 2079.124871 | 0.52188 |
| NOS3 | chr7 | 150679863 | 35 | 33 | 1784.621003 | 1743.405746 | 0.52202 |
| LCK | chr1 | 32740671 | 21 | 19 | 2002.5 | 1783.690476 | 0.52234 |
| F7 | chr13 | 113751070 | 13 | 14 | 1290.151515 | 1351.666667 | 0.52238 |
| DHFR | chr5 | 79960175 | 34 | 32 | 2754.601879 | 2708.991913 | 0.5225 |
| ESR1 | chr6 | 152002540 | 14 | 15 | 755.515873 | 879.9603175 | 0.52264 |
| LDLR | chr19 | 11192061 | 13 | 7 | 1217.777778 | 680 | 0.52264 |
| APOA1 | chr11 | 116687900 | 32 | 32 | 2896.902253 | 2845.308525 | 0.52268 |
| APOA1 | chr11 | 116706057 | 12 | 15 | 480.4256854 | 544.1919192 | 0.52286 |
| PECAM1 | chr17 | 62396973 | 30 | 29 | 2826.054968 | 2768.969103 | 0.52328 |
| ADRA2C | chr4 | 3767166 | 8 | 16 | 221.969697 | 496.7856975 | 0.52364 |
| GCK | chr7 | 44230320 | 38 | 32 | 3108.762588 | 2655.476508 | 0.52378 |
| APOA1 | chr11 | 116716794 | 37 | 33 | 3212.05408 | 2811.128776 | 0.52398 |
| APOA1 | chr11 | 116716215 | 31 | 30 | 2810.446039 | 2653.218856 | 0.52406 |
| GCK | chr7 | 44232261 | 33 | 32 | 2820.365083 | 2677.620023 | 0.52412 |
| LDLR | chr19 | 11190945 | 36 | 32 | 3372.511914 | 3024.46132 | 0.5244 |
| FOXP3 | chrX | 49127200 | 19 | 24 | 1826.550117 | 2270.27417 | 0.52444 |
| LPA | chr6 | 160942889 | 37 | 32 | 3304.900217 | 2896.105169 | 0.52446 |
| ESR1 | chr6 | 152008611 | 38 | 32 | 3394.879882 | 2826.339068 | 0.52476 |
| ALOX15 | chr17 | 4533904 | 15 | 13 | 1429.047619 | 1210.714286 | 0.5252 |

| | | | | | | | |
|---|---|---|---|---|---|---|---|
| NOS3 | chr7 | 150710114 | 14 | 15 | 473.9105339 | 566.6378066 | 0.52544 |
| APOA1 | chr11 | 116706303 | 9 | 7 | 251.7568543 | 245.1616162 | 0.52544 |
| CDKN2BAS | chr9 | 22067829 | 19 | 13 | 956.8731997 | 599.0972108 | 0.5255 |
| FOXP3 | chrX | 49117921 | 23 | 26 | 2028.914141 | 2356.707737 | 0.52552 |
| CDKN2B-AS1 | chr9 | 21975678 | 3 | 2 | 75.66666667 | 122.2222222 | 0.5256 |
| APOA1 | chr11 | 116701045 | 31 | 33 | 2743.061511 | 2854.972702 | 0.52564 |
| ABCG1 | chr21 | 43636472 | 32 | 31 | 2929.329207 | 2873.297274 | 0.52566 |
| NOS3 | chr7 | 150695261 | 33 | 32 | 2971.818191 | 2803.108854 | 0.5259 |
| APOA1 | chr11 | 116706037 | 12 | 15 | 451.468254 | 625.9523809 | 0.526 |
| CTGF | chr6 | 132275640 | 3 | 2 | 71.64835165 | 107.5396825 | 0.52604 |
| CDKN2B-AS1 | chr9 | 21974464 | 9 | 5 | 240.4446586 | 180.1455734 | 0.52606 |
| ABCG1 | chr21 | 43630128 | 33 | 31 | 1328.06948 | 1182.58516 | 0.52678 |
| LPA | chr6 | 11224454 | 27 | 32 | 1973.900658 | 2424.848599 | 0.52682 |
| PECAM1 | chr17 | 62398521 | 2 | 5 | 60 | 233.6507937 | 0.5272 |
| LCK | chr1 | 32740758 | 23 | 17 | 2188.075397 | 1641.901154 | 0.52756 |
| CDKN2B-AS1 | chr9 | 21971139 | 10 | 6 | 303.3333333 | 164.2202242 | 0.5278 |
| F7 | chr13 | 113760302 | 21 | 22 | 1659.334416 | 1796.788212 | 0.52788 |
| AMOTL2 | chr3 | 134094100 | 27 | 28 | 845.2520158 | 929.5762652 | 0.52796 |
| LDLR | chr19 | 11224442 | 35 | 32 | 3380.875249 | 3108.931546 | 0.52808 |
| ALDOA | chr16 | 30060654 | 19 | 21 | 1010.802154 | 1208.727888 | 0.5281 |
| ARHGEF18 | chr19 | 7467865 | 7 | 7 | 292.8663004 | 337.0250664 | 0.52812 |
| GALNT2 | chr1 | 230191263 | 33 | 32 | 3125.753614 | 3002.209944 | 0.52828 |
| LCK | chr1 | 32740034 | 33 | 32 | 2844.249203 | 2809.66799 | 0.52852 |
| ABCG1 | chr21 | 43638675 | 24 | 26 | 2276.428571 | 2412.261905 | 0.5286 |
| F7 | chr13 | 113753766 | 35 | 32 | 3000.555838 | 2784.958718 | 0.5287 |
| TERT | chr5 | 1295760 | 15 | 11 | 418.3689657 | 351.8330987 | 0.5287 |
| NOS3 | chr7 | 150711206 | 6 | 7 | 198.452381 | 283.961039 | 0.5288 |
| LPA | chr6 | 160946999 | 36 | 32 | 3385.856667 | 3040.585688 | 0.52886 |
| F7 | chr13 | 113750807 | 31 | 29 | 3028.185991 | 2789.488012 | 0.5289 |
| KLF2 | chr19 | 16429237 | 33 | 32 | 2996.73015 | 2871.866275 | 0.52954 |
| APOA1 | chr11 | 116688161 | 35 | 32 | 3363.439108 | 3054.23742 | 0.52978 |
| F2RL3 | chr19 | 16997292 | 30 | 31 | 2703.436841 | 2846.825397 | 0.53004 |
| NOS3 | chr7 | 150695345 | 2 | 2 | 47.22222222 | 120 | 0.53026 |
| CDKN1A | chr6 | 36645812 | 6 | 6 | 393.8095238 | 431.9047619 | 0.53034 |
| APOA1 | chr11 | 116687555 | 30 | 33 | 2506.09508 | 2690.662337 | 0.53058 |
| F2RL3 | chr19 | 16992496 | 9 | 12 | 820 | 1129.242424 | 0.53062 |
| F7 | chr13 | 113752682 | 33 | 32 | 3193.580888 | 3067.76929 | 0.53064 |
| LOXL1 | chr15 | 74217662 | 32 | 32 | 3045.080406 | 3017.47296 | 0.53066 |
| APOA1 | chr11 | 116681576 | 22 | 26 | 1942.113213 | 2239.86791 | 0.53072 |
| LPA | chr6 | 11216211 | 32 | 32 | 3047.483271 | 3065.690068 | 0.53074 |
| PECAM1 | chr17 | 62397972 | 39 | 32 | 3191.84718 | 2588.836608 | 0.5308 |
| ESR1 | chr6 | 152002200 | 31 | 32 | 2965.485548 | 3029.022007 | 0.53094 |
| APOA1 | chr11 | 116701149 | 21 | 15 | 2045.670996 | 1437.936508 | 0.53096 |
| LDLR | chr19 | 11195067 | 31 | 30 | 2652.276298 | 2489.393939 | 0.53104 |
| COL15A1 | chr9 | 101705161 | 33 | 33 | 2302.413239 | 2367.25599 | 0.5311 |
| ALOX15 | chr17 | 4550208 | 34 | 32 | 3149.748077 | 2935.303812 | 0.53146 |
| NOS3 | chr7 | 150692124 | 9 | 8 | 229.4164676 | 247.4579832 | 0.53146 |

| | | | | | | | |
|---|---|---|---|---|---|---|---|
| BCL2 | chr18 | 60988282 | 26 | 30 | 908.3388321 | 1103.945527 | 0.53184 |
| MCPH1 | chr8 | 6470114 | 21 | 26 | 2029.13961 | 2539.047619 | 0.53202 |
| FOXP3 | chrX | 49127053 | 18 | 15 | 1673.170996 | 1356.428571 | 0.53216 |
| ESR1 | chr6 | 152003785 | 37 | 32 | 3376.081923 | 2897.281364 | 0.53234 |
| TIMP3 | chr22 | 33195138 | 31 | 32 | 2650.282657 | 2806.139351 | 0.53294 |
| APOA1 | chr11 | 116700050 | 20 | 21 | 1101.925636 | 1216.749084 | 0.53304 |
| F2RL3 | chr19 | 16997402 | 30 | 32 | 2691.365483 | 2927.818073 | 0.53308 |
| LCK | chr1 | 32740891 | 21 | 26 | 1840.753968 | 2343.345405 | 0.53312 |
| APOA1 | chr11 | 116661604 | 29 | 28 | 2781.883117 | 2719.747942 | 0.53324 |
| LCK | chr1 | 32708250 | 9 | 10 | 270.7089107 | 353.967699 | 0.53328 |
| CDKN2B-AS1 | chr9 | 21989594 | 30 | 27 | 1115.468758 | 950.4775066 | 0.53338 |
| LDLR | chr19 | 11217534 | 2 | 3 | 45 | 140 | 0.5338 |
| GCK | chr7 | 44236488 | 2 | 3 | 45 | 140 | 0.5338 |
| ABCA1 | chr9 | 107690765 | 2 | 3 | 45 | 140 | 0.5338 |
| FOXP3 | chrX | 49115826 | 16 | 8 | 1552.380952 | 757.5 | 0.53396 |
| FOXP3 | chrX | 49116345 | 29 | 29 | 2674.460539 | 2620.257618 | 0.53404 |
| NOS3 | chr7 | 150710633 | 7 | 8 | 197.1825397 | 277.3999856 | 0.53422 |
| ESR1 | chr6 | 152012049 | 26 | 17 | 796.0311177 | 562.518308 | 0.53434 |
| FOXP3 | chrX | 49127116 | 20 | 21 | 1403.538961 | 1543.809524 | 0.53444 |
| NOS3 | chr7 | 150707245 | 15 | 20 | 515.0596022 | 745.1406926 | 0.53446 |
| APOA1 | chr11 | 116663479 | 34 | 32 | 2834.365937 | 2715.863774 | 0.53454 |
| LPA | chr6 | 160950765 | 11 | 17 | 949.7186147 | 1518.333333 | 0.53456 |
| ABCB1 | chr7 | 87345189 | 8 | 8 | 679.8809524 | 711.4285714 | 0.53458 |
| E2F6 | chr2 | 11584851 | 40 | 32 | 3749.393358 | 2980.747925 | 0.53474 |
| CDKN2B-AS1 | chr9 | 21998121 | 40 | 33 | 2968.372254 | 2395.227995 | 0.5348 |
| TIMP3 | chr22 | 33196384 | 35 | 31 | 2341.011369 | 2143.049003 | 0.53496 |
| PLA2G7 | chr6 | 46708139 | 32 | 32 | 2605.737214 | 2561.653057 | 0.53518 |
| LCK | chr1 | 32741128 | 19 | 18 | 608.0060952 | 529.6658119 | 0.53522 |
| ADRA2C | chr4 | 3760440 | 36 | 32 | 3296.333848 | 2963.299593 | 0.53554 |
| MCPH1 | chr8 | 6261231 | 32 | 32 | 3145.475721 | 3107.090215 | 0.53556 |
| APOA1 | chr11 | 116704314 | 33 | 32 | 3272.479768 | 3068.437384 | 0.53596 |
| P2RY12 | chr3 | 151102704 | 30 | 27 | 1054.354353 | 1004.464077 | 0.536 |
| PLA2G7 | chr6 | 46710918 | 40 | 32 | 3375.121129 | 2733.126521 | 0.53602 |
| AMOTL2 | chr3 | 134100344 | 20 | 20 | 1546.287879 | 1617.496948 | 0.5361 |
| ICAM2 | chr17 | 62080191 | 33 | 31 | 3192.644876 | 2971.634477 | 0.5362 |
| CDKN2B-AS1 | chr9 | 21970988 | 7 | 6 | 211.9444444 | 233.0769231 | 0.53638 |
| APOA1 | chr11 | 116671908 | 9 | 8 | 746.8253968 | 620 | 0.5364 |
| LCK | chr1 | 32742054 | 28 | 24 | 2545.040252 | 2223.686869 | 0.53672 |
| CDKN2B-AS1 | chr9 | 21976536 | 2 | 3 | 135 | 227.5 | 0.53672 |
| LDLR | chr19 | 11216642 | 16 | 18 | 1478.931069 | 1620.198413 | 0.53676 |
| APOA1 | chr11 | 116661949 | 31 | 33 | 1375.590705 | 1522.432704 | 0.53678 |
| ALOX15 | chr17 | 4545701 | 4 | 2 | 160 | 62.85714286 | 0.53696 |
| NOS3 | chr7 | 150693653 | 2 | 4 | 45 | 168.5714286 | 0.53704 |
| NOS3 | chr7 | 150684116 | 4 | 8 | 348.8888889 | 740 | 0.53732 |
| APOA1 | chr11 | 116704866 | 34 | 32 | 3001.722534 | 2867.934367 | 0.53738 |
| ABCB1 | chr7 | 87345429 | 32 | 32 | 2089.124631 | 2155.458929 | 0.53746 |
| KLF2 | chr19 | 16435990 | 7 | 2 | 185.3174603 | 122.2222222 | 0.53748 |

| | | | | | | | |
|---|---|---|---|---|---|---|---|
| APOA1 | chr11 | 116661280 | 32 | 32 | 2522.690099 | 2587.185051 | 0.5375 |
| FOXP3 | chrX | 49127103 | 19 | 18 | 1742.449495 | 1686.190476 | 0.53752 |
| COL15A1 | chr9 | 101705939 | 11 | 5 | 347.4603175 | 137.1428571 | 0.53756 |
| PHACTR1 | chr6 | 13276250 | 31 | 31 | 1946.341332 | 1851.071597 | 0.53762 |
| BCL2 | chr18 | 60994662 | 38 | 32 | 3414.027933 | 2898.405936 | 0.53766 |
| P2RY12 | chr3 | 151107263 | 3 | 3 | 80.55555555 | 140 | 0.5377 |
| APOA1 | chr11 | 116704215 | 29 | 30 | 2674.930821 | 2731.082904 | 0.53778 |
| GCK | chr7 | 44232358 | 36 | 32 | 2895.526819 | 2617.713915 | 0.53784 |
| TERT | chr5 | 1301707 | 37 | 32 | 3195.96904 | 2820.666343 | 0.5379 |
| LCK | chr1 | 32741476 | 24 | 23 | 2019.922577 | 1996.309524 | 0.53816 |
| F7 | chr13 | 113750311 | 31 | 29 | 2495.661949 | 2404.609753 | 0.53842 |
| CTGF | chr6 | 132274782 | 34 | 32 | 2962.81961 | 2745.107961 | 0.53852 |
| ADRA2C | chr4 | 3766351 | 33 | 32 | 2985.064528 | 2936.623295 | 0.53856 |
| ABCA1 | chr9 | 107690754 | 16 | 16 | 765.8730159 | 832.6406927 | 0.5386 |
| APOA1 | chr11 | 116697118 | 12 | 22 | 777.5487013 | 1286.861472 | 0.53864 |
| LPA | chr6 | 11216837 | 39 | 32 | 3744.859299 | 3051.402778 | 0.53868 |
| TERT | chr5 | 1296760 | 24 | 18 | 1272.011133 | 1034.668395 | 0.5387 |
| APOA1 | chr11 | 116709383 | 33 | 32 | 2823.451457 | 2781.988022 | 0.5388 |
| ADRA2C | chr4 | 3764651 | 31 | 32 | 2894.596279 | 3037.171335 | 0.53946 |
| CDKN2B-AS1 | chr9 | 22006303 | 4 | 2 | 100.4437413 | 106.0439561 | 0.53966 |
| AKT2 | chr19 | 40792614 | 41 | 32 | 3901.950908 | 3030.518281 | 0.53988 |
| F2RL3 | chr19 | 16990346 | 31 | 32 | 2586.663926 | 2559.715321 | 0.54012 |
| APOA1 | chr11 | 116679673 | 36 | 32 | 2985.081885 | 2697.494391 | 0.54014 |
| CDKN2B-AS1 | chr9 | 21968305 | 16 | 15 | 446.6702308 | 458.7745886 | 0.54078 |
| CDKN2B-AS1 | chr9 | 21969875 | 11 | 15 | 310.8760684 | 474.4614209 | 0.54134 |
| CDKN2B-AS1 | chr9 | 21996780 | 4 | 4 | 101.5873016 | 113.5714286 | 0.54136 |
| PECAM1 | chr17 | 62400146 | 4 | 2 | 380 | 200 | 0.54146 |
| LDLR | chr19 | 11223611 | 4 | 2 | 387.5 | 200 | 0.54146 |
| LDLR | chr19 | 11223858 | 4 | 2 | 380 | 200 | 0.54146 |
| BAX | chr19 | 49457187 | 2 | 4 | 200 | 380 | 0.54146 |
| E2F6 | chr2 | 11613243 | 2 | 4 | 40 | 160 | 0.54146 |
| ABCG1 | chr21 | 43631314 | 4 | 2 | 387.5 | 200 | 0.54146 |
| CDKN2B-AS1 | chr9 | 22008845 | 2 | 4 | 40 | 148.8888889 | 0.54146 |
| LDLR | chr19 | 11223602 | 4 | 2 | 385.7142857 | 200 | 0.54146 |
| LDLR | chr19 | 11216095 | 28 | 30 | 2613.484848 | 2837.259317 | 0.54172 |
| LPA | chr6 | 160949528 | 38 | 32 | 3426.399546 | 2905.947164 | 0.54226 |
| FOXP3 | chrX | 49130961 | 33 | 31 | 3146.878419 | 2910.25377 | 0.54228 |
| APOA1 | chr11 | 116725625 | 2 | 2 | 60 | 120 | 0.5425 |
| ARHGEF18 | chr19 | 7466475 | 2 | 2 | 60 | 120 | 0.5425 |
| TIMP3 | chr22 | 33192967 | 2 | 2 | 60 | 120 | 0.5425 |
| AMOTL2 | chr3 | 134098182 | 2 | 2 | 60 | 120 | 0.5425 |
| FOXP3 | chrX | 49126941 | 12 | 9 | 616.8253968 | 407.4603175 | 0.54252 |
| MCPH1 | chr8 | 6260458 | 18 | 18 | 1384.198954 | 1424.897834 | 0.54256 |
| AMOTL2 | chr3 | 134097937 | 27 | 30 | 2184.688813 | 2481.585041 | 0.5426 |
| ABCB1 | chr7 | 87343509 | 32 | 30 | 2753.920671 | 2625.848186 | 0.5427 |
| CDKN2B-AS1 | chr9 | 21996056 | 11 | 14 | 381.547619 | 539.97114 | 0.54312 |
| ALOX15 | chr17 | 4548217 | 12 | 7 | 998.8888889 | 617.1428571 | 0.5432 |

| | | | | | | | |
|---|---|---|---|---|---|---|---|
| ADRA2C | chr4 | 3768708 | 28 | 29 | 1345.501443 | 1469.521595 | 0.5433 |
| AKT2 | chr19 | 40794981 | 24 | 29 | 2168.131596 | 2557.447275 | 0.54348 |
| ADRA2C | chr4 | 3765047 | 9 | 6 | 275.1113021 | 232.8282828 | 0.5436 |
| CDKN2B-AS1 | chr9 | 21976041 | 22 | 26 | 735.3131787 | 821.0027496 | 0.54368 |
| ALOX15 | chr17 | 4549900 | 25 | 24 | 2333.888889 | 2282.568543 | 0.54374 |
| APOA1 | chr11 | 116690502 | 2 | 2 | 61.9047619 | 120 | 0.54384 |
| ABCA1 | chr9 | 107695540 | 31 | 29 | 2194.787874 | 2179.950289 | 0.54394 |
| COL15A1 | chr9 | 101696973 | 22 | 27 | 2100 | 2622.099567 | 0.54402 |
| CDKN2B-AS1 | chr9 | 21970965 | 9 | 7 | 232.8282828 | 213.0349063 | 0.5442 |
| APOA1 | chr11 | 116713587 | 31 | 32 | 2903.595329 | 2945.459776 | 0.54422 |
| PLA2G7 | chr6 | 46703131 | 4 | 3 | 107.8571429 | 140 | 0.5443 |
| KLF2 | chr19 | 16437878 | 23 | 29 | 1029.710252 | 1219.62399 | 0.54434 |
| ESR1 | chr6 | 152008032 | 34 | 32 | 3151.401926 | 2994.796787 | 0.5445 |
| LCK | chr1 | 32741961 | 29 | 24 | 2681.295094 | 2174.88234 | 0.54454 |
| ADRA2C | chr4 | 3764957 | 6 | 4 | 160.1298701 | 93.33333333 | 0.54474 |
| TFPI | chr2 | 188425123 | 3 | 4 | 275 | 343.3333333 | 0.54474 |
| NR3C1 | chr5 | 142787929 | 33 | 32 | 2913.876796 | 2868.810665 | 0.54478 |
| AMOTL2 | chr3 | 134094173 | 26 | 19 | 752.9819982 | 595.2651969 | 0.545 |
| APOA1 | chr11 | 116665080 | 29 | 28 | 1986.686079 | 1804.013574 | 0.54504 |
| GALNT2 | chr1 | 230186778 | 17 | 12 | 729.5246038 | 567.6757458 | 0.54506 |
| NOS3 | chr7 | 150711220 | 2 | 4 | 48.57142857 | 173.3333333 | 0.54508 |
| CTGF | chr6 | 132276495 | 34 | 33 | 2135.837719 | 2143.368976 | 0.54598 |
| F7 | chr13 | 113760043 | 7 | 4 | 407.9761905 | 273.3333333 | 0.5461 |
| TERT | chr5 | 1299650 | 10 | 8 | 905.7142857 | 745.7142857 | 0.54616 |
| APOA1 | chr11 | 116679777 | 35 | 32 | 2957.030779 | 2745.449414 | 0.54626 |
| AMOTL2 | chr3 | 134094602 | 2 | 3 | 48.57142857 | 142.2222222 | 0.54644 |
| CDKN2B-AS1 | chr9 | 22002171 | 21 | 18 | 567.811676 | 526.8427333 | 0.54648 |
| ICAM2 | chr17 | 62084074 | 33 | 33 | 1985.019959 | 1916.119846 | 0.54696 |
| CDKN2B-AS1 | chr9 | 22005123 | 3 | 3 | 81.9047619 | 140 | 0.54702 |
| F7 | chr13 | 113751083 | 12 | 10 | 1172.380952 | 951.4285714 | 0.54706 |
| ESR1 | chr6 | 152002589 | 20 | 26 | 1273.287546 | 1750.328283 | 0.5471 |
| CDKN2B-AS1 | chr9 | 22011082 | 9 | 2 | 220.0058815 | 105 | 0.5471 |
| TERT | chr5 | 1300439 | 33 | 33 | 2227.5396 | 2143.783041 | 0.5477 |
| P2RY12 | chr3 | 151104780 | 32 | 32 | 2725.680482 | 2785.336053 | 0.5477 |
| LOXL1 | chr15 | 74216963 | 27 | 30 | 2066.984241 | 2218.561439 | 0.54786 |
| TFPI | chr2 | 188424525 | 19 | 16 | 1788.160266 | 1526.762266 | 0.54814 |
| TMPO | chr12 | 98901355 | 37 | 32 | 3566.961265 | 3071.763994 | 0.54854 |
| LDLR | chr19 | 11218360 | 11 | 7 | 1002.857143 | 590.8333333 | 0.54862 |
| LPA | chr6 | 11222001 | 35 | 32 | 3363.085002 | 3057.44532 | 0.54892 |
| ESR1 | chr6 | 152002505 | 8 | 9 | 197.8082247 | 267.5030525 | 0.54902 |
| GCK | chr7 | 44237676 | 33 | 32 | 2744.931731 | 2706.975932 | 0.54912 |
| APOA1 | chr11 | 116705514 | 27 | 28 | 2461.418129 | 2501.638084 | 0.54934 |
| AMOTL2 | chr3 | 134102215 | 15 | 12 | 1443.214286 | 1133.214286 | 0.54948 |
| F7 | chr13 | 113755406 | 32 | 33 | 2913.101601 | 3056.894807 | 0.5497 |
| TERT | chr5 | 1299538 | 25 | 28 | 2192.083195 | 2501.904762 | 0.54978 |
| LCK | chr1 | 32741148 | 18 | 20 | 675.5517019 | 695.6747337 | 0.55016 |
| LPA | chr6 | 11224401 | 25 | 27 | 2340.35964 | 2572.667333 | 0.55034 |

| | | | | | | | |
|---|---|---|---|---|---|---|---|
| PLA2G7 | chr6 | 46707628 | 34 | 32 | 3262.585116 | 3091.674146 | 0.5504 |
| TERT | chr5 | 1298574 | 23 | 23 | 2054.79798 | 2105.627706 | 0.55044 |
| ATF3 | chr1 | 212739139 | 34 | 32 | 3031.363873 | 2811.929185 | 0.55054 |
| F7 | chr13 | 113755353 | 33 | 32 | 2716.405725 | 2678.355104 | 0.55066 |
| F7 | chr13 | 113750444 | 20 | 23 | 1778.055556 | 2099.325397 | 0.55074 |
| ALOX15 | chr17 | 4547909 | 16 | 19 | 1262.247475 | 1436.349206 | 0.55076 |
| APOA1 | chr11 | 116661036 | 34 | 32 | 2981.501868 | 2844.832076 | 0.55084 |
| ABCA1 | chr9 | 107691630 | 10 | 8 | 874.6356421 | 668.3333333 | 0.55098 |
| HMGCR | chr5 | 74626039 | 36 | 32 | 3468.384832 | 3061.492045 | 0.55116 |
| CDKN2B-AS1 | chr9 | 21988621 | 23 | 29 | 1271.574271 | 1690.649342 | 0.55116 |
| APOA1 | chr11 | 116668442 | 32 | 32 | 2994.982696 | 2965.64871 | 0.55122 |
| AKT2 | chr19 | 40798802 | 32 | 32 | 3034.280222 | 2990.241621 | 0.55132 |
| CDKN2B-AS1 | chr9 | 21970694 | 4 | 3 | 110 | 135.1785714 | 0.55132 |
| ALOX15 | chr17 | 4549905 | 26 | 25 | 2450.111833 | 2388.845599 | 0.55188 |
| ABCG1 | chr21 | 43630820 | 8 | 6 | 615.0915751 | 434.4444445 | 0.5519 |
| TERT | chr5 | 1298744 | 4 | 2 | 102.8571429 | 120 | 0.55204 |
| P2RY12 | chr3 | 151103844 | 29 | 25 | 2371.678925 | 2100.30969 | 0.55242 |
| APOA1 | chr11 | 116659865 | 27 | 25 | 2471.817072 | 2328.928571 | 0.55284 |
| GCK | chr7 | 44232133 | 34 | 31 | 3006.928117 | 2681.999242 | 0.55288 |
| PECAM1 | chr17 | 62409928 | 35 | 33 | 1640.830299 | 1475.35439 | 0.55334 |
| HMGCR | chr5 | 74631872 | 35 | 32 | 3046.248206 | 2730.959833 | 0.55344 |
| APOA1 | chr11 | 116671313 | 38 | 32 | 3342.101309 | 2838.264273 | 0.55346 |
| LDLR | chr19 | 11194843 | 32 | 33 | 2689.704437 | 2715.558679 | 0.55384 |
| LCK | chr1 | 32716144 | 14 | 14 | 383.5292304 | 409.420224 | 0.5541 |
| TERT | chr5 | 1301131 | 34 | 32 | 3318.873116 | 3090.454262 | 0.55412 |
| F7 | chr13 | 113756528 | 35 | 32 | 3231.643826 | 2976.493634 | 0.55414 |
| LPA | chr6 | 160945327 | 39 | 32 | 3611.176056 | 2985.986293 | 0.55426 |
| CDKN2B-AS1 | chr9 | 21987512 | 30 | 33 | 1246.057771 | 1427.031925 | 0.55436 |
| CDKN2B-AS1 | chr9 | 22011109 | 7 | 2 | 184.9932612 | 115.8263305 | 0.55438 |
| FOXP3 | chrX | 49125874 | 2 | 2 | 85.29411765 | 133.3333333 | 0.55452 |
| GCK | chr7 | 44231814 | 33 | 32 | 2983.708504 | 2941.582118 | 0.55466 |
| MCPH1 | chr8 | 6470725 | 34 | 32 | 3213.268577 | 3054.557032 | 0.55476 |
| LDLR | chr19 | 11224134 | 12 | 14 | 1064.126984 | 1295.714286 | 0.55486 |
| ESR2 | chr14 | 64809928 | 36 | 32 | 3334.424398 | 2995.937349 | 0.55502 |
| APOA1 | chr11 | 116675441 | 33 | 33 | 2929.952367 | 2982.339229 | 0.55508 |
| ADRA2C | chr4 | 3765186 | 22 | 19 | 1163.064553 | 929.7106403 | 0.55526 |
| LDLR | chr19 | 11223315 | 24 | 21 | 2266.147186 | 2013.412698 | 0.55528 |
| CDKN2B-AS1 | chr9 | 21990057 | 15 | 14 | 535.5708181 | 456.7171717 | 0.5553 |
| AKT2 | chr19 | 40800952 | 28 | 28 | 2664.242424 | 2703.226496 | 0.55544 |
| PLA2G7 | chr6 | 46703555 | 20 | 17 | 536.9442568 | 498.8500781 | 0.55552 |
| ESR1 | chr6 | 152006044 | 33 | 32 | 3112.733791 | 3050.610299 | 0.55556 |
| LDLR | chr19 | 11218208 | 33 | 32 | 3204.773639 | 3132.220432 | 0.5556 |
| APOA1 | chr11 | 116706042 | 11 | 18 | 471.4568764 | 696.2371615 | 0.5559 |
| COL15A1 | chr9 | 101699971 | 29 | 30 | 2697.292877 | 2744.16137 | 0.5563 |
| APOA1 | chr11 | 116689648 | 23 | 28 | 2151.020924 | 2673.974359 | 0.55644 |
| LCK | chr1 | 32707241 | 16 | 13 | 441.9146282 | 404.8614309 | 0.55676 |
| ALOX15 | chr17 | 4547395 | 35 | 32 | 3200.766324 | 2900.430503 | 0.55706 |

| | | | | | | | |
|---|---|---|---|---|---|---|---|
| F7 | chr13 | 113752904 | 31 | 32 | 2865.309837 | 2914.840715 | 0.55712 |
| ATF3 | chr1 | 212743930 | 27 | 27 | 2556.572455 | 2600 | 0.55722 |
| MIA3 | chr1 | 222792587 | 32 | 31 | 1489.488342 | 1384.131505 | 0.55758 |
| P2RY12 | chr3 | 151106039 | 32 | 32 | 1164.71087 | 1220.650583 | 0.5576 |
| LPA | chr6 | 160947460 | 37 | 32 | 3174.168223 | 2780.736882 | 0.55762 |
| LDLR | chr19 | 11221137 | 19 | 20 | 1760.979494 | 1894.166667 | 0.5577 |
| PLA2G7 | chr6 | 46703539 | 28 | 23 | 800.9673736 | 694.6750301 | 0.55784 |
| ALOX15 | chr17 | 4550285 | 36 | 32 | 3350.786299 | 2940.699935 | 0.55808 |
| BCL2 | chr18 | 60995885 | 20 | 14 | 1901.984127 | 1284.047619 | 0.55814 |
| CDKN2B-AS1 | chr9 | 21967941 | 30 | 26 | 1019.927635 | 932.7939628 | 0.5582 |
| DHFR | chr5 | 79957689 | 36 | 32 | 3206.287957 | 2927.650513 | 0.55828 |
| ARHGEF18 | chr19 | 7462076 | 25 | 20 | 2080.937951 | 1614.642857 | 0.55836 |
| CDKN1A | chr6 | 36643506 | 28 | 24 | 2731.666667 | 2317.951128 | 0.55852 |
| DHFR | chr5 | 79957738 | 31 | 24 | 2232.874566 | 1823.532071 | 0.55866 |
| PECAM1 | chr17 | 62410105 | 35 | 31 | 1351.019751 | 1246.011274 | 0.55894 |
| GCK | chr7 | 44234910 | 34 | 32 | 2865.961914 | 2650.068645 | 0.55894 |
| TERT | chr5 | 1298266 | 32 | 32 | 3056.148877 | 3084.381294 | 0.55914 |
| APOA1 | chr11 | 116696681 | 24 | 23 | 1251.824364 | 1284.859612 | 0.55914 |
| TIMP3 | chr22 | 33259492 | 32 | 32 | 2938.674498 | 2872.353038 | 0.55936 |
| NOS3 | chr7 | 150692566 | 26 | 25 | 1091.751582 | 992.2186147 | 0.55996 |
| ABCA1 | chr9 | 107699094 | 37 | 32 | 3136.699138 | 2740.936792 | 0.5602 |
| APOA1 | chr11 | 116659603 | 9 | 6 | 327.4747475 | 248.4126984 | 0.56028 |
| F7 | chr13 | 113752684 | 33 | 32 | 3215.682431 | 3137.189725 | 0.56036 |
| LDLR | chr19 | 11217356 | 33 | 32 | 3139.582011 | 3018.996875 | 0.56042 |
| APOA1 | chr11 | 116688224 | 36 | 32 | 3331.460804 | 2992.750442 | 0.56058 |
| F7 | chr13 | 113758341 | 31 | 31 | 2640.573871 | 2573.51101 | 0.56106 |
| AMOTL2 | chr3 | 134094803 | 37 | 32 | 3311.854208 | 2839.306595 | 0.56118 |
| APOA1 | chr11 | 116690915 | 32 | 32 | 2773.462869 | 2726.983445 | 0.56132 |
| LDLR | chr19 | 11197932 | 11 | 16 | 1017.460317 | 1526.547619 | 0.56132 |
| NOS3 | chr7 | 150710573 | 2 | 6 | 45 | 212.2727273 | 0.56178 |
| TMPO | chr12 | 98906749 | 37 | 32 | 2983.362079 | 2616.374972 | 0.56202 |
| ADRA2C | chr4 | 3763941 | 32 | 30 | 2890.842944 | 2664.48495 | 0.56208 |
| NOS3 | chr7 | 150707830 | 20 | 22 | 602.0611301 | 706.8921521 | 0.56208 |
| NOS3 | chr7 | 150710028 | 32 | 33 | 1663.336408 | 1781.026143 | 0.5621 |
| DHFR | chr5 | 79960269 | 29 | 24 | 2209.726244 | 1729.684891 | 0.56242 |
| NOS3 | chr7 | 150695664 | 25 | 21 | 2272.968245 | 1944.68254 | 0.56246 |
| TERT | chr5 | 1253617 | 32 | 33 | 2770.587147 | 2916.085227 | 0.56248 |
| ABCG1 | chr21 | 43636822 | 36 | 32 | 3433.924118 | 3072.218018 | 0.5625 |
| NR3C1 | chr5 | 142785456 | 36 | 33 | 1877.964806 | 1670.894964 | 0.56262 |
| ARHGEF18 | chr19 | 7465990 | 35 | 32 | 3280.305594 | 2976.530126 | 0.56262 |
| CDKN2B-AS1 | chr9 | 21971016 | 4 | 4 | 118.0555556 | 169.0118577 | 0.56274 |
| TERT | chr5 | 1300567 | 37 | 32 | 3287.975132 | 2875.21278 | 0.56296 |
| APOA1 | chr11 | 116691155 | 33 | 32 | 3096.447013 | 3027.548648 | 0.56344 |
| CDKN2B-AS1 | chr9 | 21988935 | 33 | 23 | 955.490509 | 705.786574 | 0.56346 |
| ESR1 | chr6 | 152007586 | 15 | 21 | 1074.480519 | 1591.497114 | 0.5635 |
| NOS3 | chr7 | 150694183 | 2 | 3 | 48.57142857 | 140 | 0.56354 |
| CDKN2B-AS1 | chr9 | 22008830 | 2 | 3 | 48.57142857 | 140 | 0.56354 |

| | | | | | | | |
|---|---|---|---|---|---|---|---|
| CDKN2B-AS1 | chr9 | 21968857 | 33 | 26 | 947.7112092 | 707.8080102 | 0.56366 |
| CDKN2B-AS1 | chr9 | 21976149 | 16 | 15 | 530.9158605 | 543.9131375 | 0.56372 |
| CDKN2B-AS1 | chr9 | 21965446 | 4 | 2 | 106.8170426 | 114.1176471 | 0.56378 |
| ALOX15 | chr17 | 4549938 | 33 | 32 | 3147.925614 | 2964.72604 | 0.56382 |
| F7 | chr13 | 113751054 | 17 | 17 | 1690.357143 | 1646.428571 | 0.56386 |
| KLF2 | chr19 | 16430537 | 7 | 5 | 189.4426009 | 174.6924603 | 0.56386 |
| F7 | chr13 | 113755271 | 28 | 29 | 2664.365079 | 2705.518109 | 0.56426 |
| FOXP3 | chrX | 49127333 | 13 | 15 | 1219.761905 | 1433.055556 | 0.5644 |
| ABCG1 | chr21 | 43632608 | 38 | 33 | 2230.134036 | 1984.86778 | 0.56466 |
| ESR2 | chr14 | 64813073 | 35 | 33 | 2111.230375 | 2037.765554 | 0.5647 |
| TERT | chr5 | 1296736 | 30 | 29 | 2713.269051 | 2590.53188 | 0.565 |
| FGF2 | chr4 | 123743035 | 37 | 32 | 3462.400728 | 3017.072798 | 0.56522 |
| LDLR | chr19 | 11224397 | 35 | 32 | 3349.872554 | 3081.434131 | 0.56524 |
| MCPH1 | chr8 | 6470129 | 23 | 25 | 2332.694805 | 2467.460317 | 0.5655 |
| DHFR | chr5 | 79952913 | 31 | 31 | 2972.052544 | 3004.328994 | 0.56552 |
| LPA | chr6 | 160944073 | 29 | 29 | 2712.970779 | 2752.458791 | 0.56568 |
| CDKN2B-AS1 | chr9 | 21974051 | 3 | 3 | 67.22222222 | 80 | 0.56624 |
| ABCG1 | chr21 | 43629488 | 37 | 33 | 2575.010703 | 2249.813911 | 0.56646 |
| CDKN2B-AS1 | chr9 | 22005614 | 3 | 2 | 108.9795918 | 111.5552326 | 0.56646 |
| NOS3 | chr7 | 150694017 | 10 | 11 | 318.0952381 | 397.4603175 | 0.56658 |
| TIMP3 | chr22 | 33187717 | 35 | 32 | 3329.582419 | 3072.524114 | 0.5667 |
| AKT2 | chr19 | 40794892 | 34 | 32 | 3283.220436 | 3070.282839 | 0.56678 |
| LPA | chr6 | 11221240 | 36 | 32 | 2861.627355 | 2586.206284 | 0.56684 |
| APOA1 | chr11 | 116684688 | 31 | 32 | 2791.767636 | 2942.059433 | 0.56696 |
| KLF2 | chr19 | 16432722 | 34 | 32 | 3010.909222 | 2869.578974 | 0.56716 |
| CDKN2B-AS1 | chr9 | 22004917 | 29 | 26 | 883.2730943 | 835.7792041 | 0.56724 |
| LDLR | chr19 | 11196467 | 17 | 24 | 1557.621545 | 2139.516595 | 0.56732 |
| APOA1 | chr11 | 116691858 | 32 | 32 | 2949.345467 | 2979.13426 | 0.56734 |
| TERT | chr5 | 1299096 | 31 | 31 | 2783.122171 | 2747.743337 | 0.56738 |
| ARHGEF18 | chr19 | 7460724 | 15 | 15 | 461.9666383 | 504.1970103 | 0.56742 |
| CDKN2B-AS1 | chr9 | 21976104 | 3 | 5 | 89.76190477 | 205.8859441 | 0.5675 |
| APOA1 | chr11 | 116661095 | 35 | 32 | 3111.419174 | 2883.777462 | 0.56804 |
| NOS3 | chr7 | 150710437 | 26 | 32 | 1706.031746 | 2170.10545 | 0.56832 |
| APOA1 | chr11 | 116702333 | 28 | 30 | 2320.220058 | 2543.438228 | 0.56838 |
| E2F6 | chr2 | 11613912 | 20 | 24 | 1932.020202 | 2286.269841 | 0.56844 |
| CTGF | chr6 | 132272852 | 7 | 3 | 204.1819292 | 133.0623306 | 0.56864 |
| CDKN2B-AS1 | chr9 | 21980725 | 24 | 26 | 2191.72619 | 2419.761905 | 0.56866 |
| FGF2 | chr4 | 123747294 | 31 | 31 | 2892.009061 | 2929.314209 | 0.56884 |
| CDKN2B-AS1 | chr9 | 21990363 | 18 | 16 | 514.2842206 | 493.1287203 | 0.56892 |
| BCL2 | chr18 | 60994314 | 10 | 7 | 889.047619 | 646.6666667 | 0.56892 |
| FOXP3 | chrX | 49120342 | 22 | 18 | 1884.278499 | 1594.603175 | 0.56894 |
| TERT | chr5 | 1298570 | 20 | 20 | 1766.153291 | 1809.599567 | 0.56898 |
| TFPI | chr2 | 188424180 | 3 | 3 | 160.4761905 | 197.1428571 | 0.56916 |
| CDKN2B-AS1 | chr9 | 21974191 | 24 | 23 | 718.6508857 | 736.0553615 | 0.56924 |
| CDKN2B-AS1 | chr9 | 21976481 | 3 | 2 | 108.5714286 | 88.57142857 | 0.56926 |
| ADRA2C | chr4 | 3768528 | 13 | 10 | 421.8174551 | 358.4235209 | 0.5695 |
| ABCG1 | chr21 | 43629952 | 36 | 32 | 3419.299343 | 3060.076658 | 0.56972 |

| | | | | | | | |
|---|---|---|---|---|---|---|---|
| F7 | chr13 | 113756725 | 31 | 33 | 2175.612027 | 2394.273381 | 0.56982 |
| F2RL3 | chr19 | 16998279 | 17 | 17 | 772.2942554 | 712.5213675 | 0.56988 |
| FOXP3 | chrX | 49129041 | 18 | 12 | 516.3031327 | 372.9571623 | 0.56988 |
| NR3C1 | chr5 | 142790700 | 32 | 32 | 3045.128517 | 3066.64394 | 0.5699 |
| ARHGEF18 | chr19 | 7469577 | 31 | 31 | 2971.925585 | 2995.506361 | 0.57052 |
| AKT2 | chr19 | 40798055 | 33 | 31 | 3059.896014 | 2900.164928 | 0.57056 |
| ESR1 | chr6 | 152009426 | 37 | 32 | 3239.11778 | 2761.248747 | 0.57072 |
| CDKN2B-AS1 | chr9 | 21966006 | 8 | 13 | 205.0213675 | 380.4259706 | 0.57076 |
| LDLR | chr19 | 11217933 | 30 | 29 | 2844.833386 | 2719.669148 | 0.57102 |
| TERT | chr5 | 1295825 | 31 | 28 | 1423.15199 | 1341.033233 | 0.57116 |
| NOS3 | chr7 | 150692049 | 18 | 18 | 748.1395014 | 686.8967874 | 0.57122 |
| FOXP3 | chrX | 49117244 | 20 | 23 | 1966.666667 | 2223.333333 | 0.57138 |
| F7 | chr13 | 113759635 | 32 | 31 | 2851.710173 | 2722.895272 | 0.57158 |
| GALNT2 | chr1 | 230189882 | 32 | 32 | 3059.798263 | 3023.802659 | 0.57166 |
| F7 | chr13 | 113757183 | 28 | 28 | 2617.741017 | 2580.850641 | 0.57172 |
| ESR1 | chr6 | 152011666 | 7 | 2 | 203.5714286 | 113.8888889 | 0.572 |
| LOXL1 | chr15 | 74218587 | 30 | 30 | 1377.068174 | 1438.350714 | 0.57218 |
| ABCG1 | chr21 | 43633279 | 37 | 32 | 3487.500782 | 2996.870322 | 0.57234 |
| F2RL3 | chr19 | 17002922 | 27 | 29 | 2154.472021 | 2383.94678 | 0.57236 |
| LDLR | chr19 | 11219885 | 5 | 11 | 472.2222222 | 987.5 | 0.57256 |
| GALNT2 | chr1 | 230187734 | 32 | 32 | 2899.587302 | 2841.549937 | 0.57262 |
| AMOTL2 | chr3 | 134097779 | 34 | 32 | 3058.082085 | 2846.342068 | 0.57296 |
| LOXL1 | chr15 | 74218974 | 26 | 25 | 969.0959096 | 886.1949259 | 0.57338 |
| NOS3 | chr7 | 150678915 | 32 | 32 | 2666.571434 | 2624.480677 | 0.57366 |
| ABCB1 | chr7 | 87342302 | 37 | 33 | 1925.418727 | 1667.826208 | 0.57368 |
| P2RY12 | chr3 | 151102379 | 33 | 32 | 2932.853862 | 2808.075607 | 0.57368 |
| COL15A1 | chr9 | 101696645 | 27 | 25 | 2049.505217 | 1838.192641 | 0.574 |
| LDLR | chr19 | 11222295 | 9 | 9 | 794.1666667 | 762.3809524 | 0.57412 |
| AMOTL2 | chr3 | 134094538 | 33 | 32 | 2884.940118 | 2841.190915 | 0.5743 |
| LOXL1 | chr15 | 74211407 | 34 | 32 | 3212.588668 | 3046.898948 | 0.5743 |
| APOA1 | chr11 | 116678000 | 22 | 22 | 1620.528075 | 1522.474747 | 0.57444 |
| CTGF | chr6 | 132273156 | 26 | 27 | 2481.755135 | 2542.000798 | 0.57448 |
| APOA1 | chr11 | 116693893 | 33 | 33 | 2448.132028 | 2516.121862 | 0.5745 |
| APOA1 | chr11 | 116692156 | 33 | 32 | 2892.30193 | 2837.777756 | 0.57476 |
| LCK | chr1 | 32740917 | 23 | 25 | 2155.714286 | 2374.965729 | 0.57504 |
| AMOTL2 | chr3 | 134100721 | 35 | 32 | 2535.183489 | 2360.166838 | 0.57512 |
| APOA1 | chr11 | 116691055 | 32 | 31 | 2439.230825 | 2287.692724 | 0.5752 |
| APOA1 | chr11 | 116675287 | 16 | 16 | 1369.440837 | 1415.151515 | 0.57532 |
| GALNT2 | chr1 | 230187632 | 31 | 32 | 2977.283319 | 3036.571009 | 0.57554 |
| LPA | chr6 | 160945093 | 37 | 32 | 3303.713022 | 2834.279307 | 0.57558 |
| LDLR | chr19 | 11195029 | 8 | 7 | 418.4782609 | 325.6977226 | 0.5756 |
| ADRA2C | chr4 | 3761056 | 31 | 30 | 2119.196661 | 2117.648879 | 0.57604 |
| ARHGEF18 | chr19 | 7460671 | 34 | 32 | 1459.226788 | 1323.778848 | 0.57614 |
| APOA1 | chr11 | 116668298 | 5 | 2 | 475 | 200 | 0.57614 |
| NOS3 | chr7 | 150707226 | 26 | 27 | 1394.219869 | 1512.004793 | 0.57616 |
| ESR1 | chr6 | 152011399 | 36 | 33 | 1964.344525 | 1746.16953 | 0.57658 |
| TERT | chr5 | 1299828 | 31 | 30 | 2434.72636 | 2297.848009 | 0.5767 |

| | | | | | | | |
|---|---|---|---|---|---|---|---|
| APOA1 | chr11 | 116661659 | 4 | 3 | 119.6969697 | 142.2222222 | 0.57676 |
| APOA1 | chr11 | 116706289 | 11 | 6 | 307.4173659 | 215.5581261 | 0.57742 |
| ADRA2C | chr4 | 3765601 | 22 | 27 | 1812.330447 | 2280.380175 | 0.57746 |
| ADRA2C | chr4 | 3765807 | 7 | 6 | 553.3333333 | 430 | 0.57754 |
| APOA1 | chr11 | 116661668 | 30 | 31 | 2349.78805 | 2509.71159 | 0.57768 |
| ARHGEF18 | chr19 | 7528676 | 5 | 4 | 363.3333333 | 321.6666667 | 0.57788 |
| ADRA2C | chr4 | 3766270 | 33 | 33 | 2691.418912 | 2626.294307 | 0.5781 |
| COL15A1 | chr9 | 101698371 | 30 | 30 | 2120.876553 | 2192.688635 | 0.57834 |
| TIMP3 | chr22 | 33195688 | 34 | 32 | 3044.522045 | 2899.536142 | 0.57858 |
| LPA | chr6 | 11220303 | 37 | 32 | 3192.592329 | 2788.625518 | 0.57894 |
| ALOX15 | chr17 | 4534094 | 31 | 30 | 2948.337912 | 2804.901961 | 0.57918 |
| NOS3 | chr7 | 150689873 | 19 | 20 | 1262.380952 | 1265.342713 | 0.57934 |
| FOXP3 | chrX | 49120887 | 35 | 32 | 3210.196568 | 2961.019296 | 0.57936 |
| TFPI | chr2 | 188425345 | 8 | 2 | 666.3888889 | 183.3333333 | 0.57946 |
| PHACTR1 | chr6 | 12711562 | 34 | 33 | 3028.094654 | 2891.178893 | 0.57972 |
| ABCA1 | chr9 | 107691882 | 34 | 32 | 3093.448504 | 2943.387803 | 0.57998 |
| ABCG1 | chr21 | 43632504 | 36 | 32 | 3325.656133 | 2977.767564 | 0.5801 |
| FOXP3 | chrX | 49126922 | 13 | 3 | 574.6536796 | 111.4285714 | 0.58016 |
| F7 | chr13 | 113755283 | 30 | 30 | 2919.781011 | 2938.667911 | 0.58024 |
| CDKN1A | chr6 | 36652134 | 28 | 29 | 2686.561217 | 2748.577534 | 0.58028 |
| F7 | chr13 | 113758392 | 30 | 32 | 2816.127761 | 2949.535544 | 0.58042 |
| LOXL1 | chr15 | 74218843 | 26 | 22 | 780.2380007 | 701.8265762 | 0.58048 |
| BCL2 | chr18 | 60989802 | 12 | 10 | 558.5714286 | 422.1212121 | 0.58066 |
| HMGCR | chr5 | 74631328 | 31 | 29 | 2740.790253 | 2526.193031 | 0.5808 |
| F7 | chr13 | 113752576 | 32 | 32 | 3009.299271 | 2980.70431 | 0.58084 |
| NR3C1 | chr5 | 142657577 | 37 | 32 | 3344.543257 | 2868.318577 | 0.58124 |
| CDKN1A | chr6 | 36651972 | 31 | 32 | 2997.499641 | 3067.118298 | 0.5815 |
| TERT | chr5 | 1295649 | 2 | 6 | 47.27272727 | 163.3333333 | 0.58154 |
| NOS3 | chr7 | 150693978 | 11 | 13 | 330.8136308 | 439.6933622 | 0.58158 |
| CDKN1A | chr6 | 36652612 | 28 | 28 | 2556.909757 | 2512.38345 | 0.5816 |
| FOXP3 | chrX | 49120380 | 24 | 21 | 2242.423132 | 1930.051893 | 0.58174 |
| KLF2 | chr19 | 16432725 | 34 | 32 | 3148.709225 | 2992.098646 | 0.58182 |
| CDKN2B-AS1 | chr9 | 21995897 | 2 | 5 | 56.81818182 | 214.0789474 | 0.582 |
| CDKN2B-AS1 | chr9 | 21971109 | 3 | 2 | 69.49494949 | 53.33333333 | 0.582 |
| FOXP3 | chrX | 49128168 | 32 | 32 | 2849.604128 | 2798.914796 | 0.5823 |
| GCK | chr7 | 44228639 | 31 | 32 | 2686.529978 | 2721.141034 | 0.5826 |
| MCPH1 | chr8 | 6470899 | 29 | 30 | 2621.379731 | 2760.219062 | 0.5826 |
| CDKN2B-AS1 | chr9 | 22009939 | 6 | 2 | 186.1904762 | 53.33333333 | 0.58272 |
| APOA1 | chr11 | 116688727 | 16 | 17 | 1472.261905 | 1525.793651 | 0.58278 |
| FOXP3 | chrX | 49127352 | 15 | 12 | 1389.559885 | 1134.603175 | 0.58294 |
| FOXP3 | chrX | 49117807 | 18 | 18 | 1454.444444 | 1509.087302 | 0.58316 |
| BCL2 | chr18 | 60989090 | 28 | 22 | 2534.31606 | 2029.155844 | 0.58392 |
| P2RY12 | chr3 | 151108412 | 39 | 32 | 3578.148252 | 2962.921305 | 0.58396 |
| MCPH1 | chr8 | 6258186 | 33 | 32 | 3070.535954 | 3001.202027 | 0.584 |
| NOS3 | chr7 | 150705767 | 33 | 33 | 1810.420644 | 1875.86294 | 0.58408 |
| BCL2 | chr18 | 60993303 | 34 | 32 | 3106.062168 | 2949.908166 | 0.58434 |
| BAX | chr19 | 49465468 | 9 | 12 | 260.5764562 | 385.2543652 | 0.58468 |

| | | | | | | | |
|---|---|---|---|---|---|---|---|
| NOS3 | chr7 | 150706203 | 33 | 32 | 2597.076404 | 2570.594909 | 0.58496 |
| TERT | chr5 | 1296491 | 34 | 32 | 2963.4771 | 2822.385229 | 0.58506 |
| ADRA2C | chr4 | 3764150 | 33 | 32 | 3023.373825 | 2892.689362 | 0.58544 |
| CDKN2B-AS1 | chr9 | 21966758 | 8 | 10 | 214.8378694 | 304.8677136 | 0.58544 |
| F2RL3 | chr19 | 16989881 | 34 | 32 | 3253.887509 | 3088.195338 | 0.58576 |
| APOA1 | chr11 | 116697309 | 32 | 32 | 3091.510338 | 3072.199033 | 0.58606 |
| LPA | chr6 | 160944300 | 32 | 32 | 2372.656689 | 2431.586608 | 0.58612 |
| LDLR | chr19 | 11215976 | 32 | 32 | 3104.570897 | 3126.949727 | 0.5862 |
| TERT | chr5 | 1298645 | 26 | 30 | 1793.424518 | 1956.21781 | 0.58626 |
| LOXL1 | chr15 | 74215318 | 3 | 4 | 176.6666667 | 200 | 0.58654 |
| BAX | chr19 | 49465492 | 5 | 2 | 168.115942 | 116.3157895 | 0.58656 |
| FOXP3 | chrX | 49126927 | 10 | 2 | 392.2366522 | 65.07936508 | 0.58676 |
| TERT | chr5 | 1253780 | 13 | 17 | 1252.307692 | 1669.047619 | 0.58696 |
| LPA | chr6 | 11219559 | 35 | 33 | 2426.118373 | 2333.025721 | 0.58718 |
| LOXL1 | chr15 | 74219049 | 29 | 27 | 1129.906354 | 997.7042883 | 0.58724 |
| APOA1 | chr11 | 116675944 | 34 | 32 | 3101.777819 | 2947.846009 | 0.58726 |
| NOS3 | chr7 | 150711192 | 6 | 6 | 167.2039072 | 213.5714286 | 0.58732 |
| CDKN1A | chr6 | 36652083 | 26 | 30 | 2473.290043 | 2807.775558 | 0.58736 |
| APOA1 | chr11 | 116680883 | 35 | 32 | 3185.035575 | 2878.603321 | 0.58758 |
| NOS3 | chr7 | 150695535 | 22 | 28 | 1332.403167 | 1772.579643 | 0.58782 |
| LOXL1 | chr15 | 74217125 | 4 | 6 | 271.9047619 | 457.1428571 | 0.58782 |
| FOXP3 | chrX | 49130039 | 4 | 3 | 118.3579812 | 137.6043557 | 0.58794 |
| APOA1 | chr11 | 116662312 | 2 | 6 | 44.86215539 | 203.1447964 | 0.58828 |
| ABCG1 | chr21 | 43717649 | 40 | 32 | 3520.838453 | 2847.997846 | 0.5885 |
| TERT | chr5 | 1300801 | 13 | 13 | 1029.52381 | 1077.777778 | 0.58864 |
| F7 | chr13 | 113750189 | 32 | 32 | 3044.532431 | 2983.116771 | 0.58902 |
| LCK | chr1 | 32741200 | 10 | 12 | 340.9290383 | 378.6111111 | 0.58982 |
| APOA1 | chr11 | 116703897 | 33 | 32 | 2977.04922 | 2855.063469 | 0.58984 |
| TERT | chr5 | 1301512 | 33 | 32 | 2696.981451 | 2653.575042 | 0.59004 |
| F7 | chr13 | 113750605 | 26 | 27 | 2433.021978 | 2477.247475 | 0.59004 |
| ABCA1 | chr9 | 107693293 | 10 | 9 | 899.047619 | 833.3333333 | 0.59018 |
| APOA1 | chr11 | 116659429 | 35 | 33 | 1631.43337 | 1494.661996 | 0.59034 |
| NOS3 | chr7 | 150711125 | 7 | 9 | 201.3168449 | 300.9173502 | 0.59036 |
| ABCG1 | chr21 | 43637710 | 18 | 23 | 1731.547619 | 2185.375874 | 0.59086 |
| TIMP3 | chr22 | 33188833 | 35 | 33 | 2402.00995 | 2217.429717 | 0.59132 |
| PECAM1 | chr17 | 62410239 | 9 | 9 | 864.2857143 | 877.5 | 0.59146 |
| ADRA2C | chr4 | 3764620 | 32 | 31 | 2520.70802 | 2493.427776 | 0.59174 |
| TERT | chr5 | 1300310 | 14 | 15 | 1223.489623 | 1356.666667 | 0.59174 |
| PHACTR1 | chr6 | 12710664 | 33 | 33 | 2309.360375 | 2360.501058 | 0.59184 |
| ADRA2C | chr4 | 3764234 | 32 | 32 | 3006.657522 | 2936.910026 | 0.59194 |
| NOS3 | chr7 | 150690992 | 27 | 29 | 2469.924432 | 2600.273126 | 0.592 |
| ARHGEF18 | chr19 | 7464191 | 35 | 33 | 2468.215785 | 2250.266574 | 0.59212 |
| CDKN2B-AS1 | chr9 | 21991146 | 28 | 24 | 839.814227 | 763.1177038 | 0.59222 |
| ABCB1 | chr7 | 87345209 | 7 | 4 | 663.3333333 | 360 | 0.59234 |
| COL15A1 | chr9 | 101704767 | 29 | 32 | 2664.353936 | 2975.260645 | 0.5925 |
| TMPO | chr12 | 98900739 | 37 | 32 | 3569.851817 | 3068.173665 | 0.5927 |
| FGF2 | chr4 | 123740008 | 36 | 32 | 3438.106092 | 3037.457843 | 0.59274 |

| | | | | | | | |
|---|---|---|---|---|---|---|---|
| PLA2G7 | chr6 | 46713225 | 30 | 30 | 2779.01951 | 2738.675214 | 0.59314 |
| GCK | chr7 | 44228688 | 28 | 32 | 1108.690788 | 1225.46895 | 0.59364 |
| APOA1 | chr11 | 116698624 | 32 | 33 | 2747.429954 | 2890.01174 | 0.594 |
| APOA1 | chr11 | 116697390 | 33 | 33 | 2452.633168 | 2390.858295 | 0.59404 |
| LCK | chr1 | 32707632 | 10 | 8 | 912.2222222 | 703.3333333 | 0.5942 |
| F7 | chr13 | 113752713 | 33 | 32 | 3190.31691 | 3116.948926 | 0.59436 |
| ATF3 | chr1 | 212741471 | 33 | 33 | 2063.008531 | 2112.115698 | 0.59448 |
| TERT | chr5 | 1253739 | 28 | 31 | 2616.592898 | 2946.040349 | 0.59496 |
| PLA2G7 | chr6 | 46703719 | 35 | 33 | 2230.59957 | 2151.96023 | 0.59538 |
| LPA | chr6 | 11219364 | 33 | 33 | 2693.236338 | 2641.529748 | 0.59544 |
| APOA1 | chr11 | 116686122 | 8 | 8 | 743.3333333 | 763.3333333 | 0.59552 |
| GCK | chr7 | 44229937 | 33 | 32 | 3088.908026 | 2960.894072 | 0.5957 |
| CDKN2BAS | chr9 | 22096335 | 35 | 32 | 3053.448245 | 2765.423573 | 0.59584 |
| CDKN2B-AS1 | chr9 | 21996245 | 27 | 27 | 964.3015048 | 1007.024746 | 0.59592 |
| CDKN2B-AS1 | chr9 | 21990065 | 11 | 12 | 328.4160284 | 400.5266955 | 0.59634 |
| CDKN2B-AS1 | chr9 | 21992794 | 33 | 32 | 2737.34173 | 2694.806897 | 0.59646 |
| APOA1 | chr11 | 116676893 | 33 | 32 | 2411.971575 | 2386.820281 | 0.59672 |
| APOA1 | chr11 | 116669797 | 31 | 24 | 2728.92544 | 2070.886932 | 0.5968 |
| KLF2 | chr19 | 16430314 | 33 | 32 | 1351.886012 | 1265.168543 | 0.59694 |
| NOS3 | chr7 | 150706013 | 31 | 31 | 1234.914292 | 1293.830295 | 0.59736 |
| F7 | chr13 | 113751795 | 33 | 32 | 2994.248789 | 2932.298998 | 0.59768 |
| CDKN2B-AS1 | chr9 | 21993501 | 40 | 33 | 2339.576325 | 1892.495135 | 0.59794 |
| CDKN1A | chr6 | 36652020 | 29 | 30 | 2719.069644 | 2853.375348 | 0.59796 |
| NR3C1 | chr5 | 142791483 | 24 | 22 | 2263.963078 | 2030.32967 | 0.5987 |
| F2RL3 | chr19 | 17000078 | 6 | 7 | 180.4102564 | 251.2851038 | 0.59898 |
| APOA1 | chr11 | 116675389 | 30 | 29 | 2559.725534 | 2429.837613 | 0.59908 |
| ALDOA | chr16 | 30060334 | 12 | 9 | 1170.833333 | 850 | 0.59912 |
| COL15A1 | chr9 | 101833055 | 31 | 29 | 2935.573593 | 2770.551589 | 0.5992 |
| ARHGEF18 | chr19 | 7469195 | 33 | 32 | 3084.674848 | 3017.574779 | 0.59938 |
| ALOX15 | chr17 | 4546990 | 8 | 7 | 336.2452107 | 335.5555556 | 0.6 |
| ABCG1 | chr21 | 43637484 | 35 | 32 | 3309.658199 | 3049.178421 | 0.60034 |
| APOA1 | chr11 | 116688072 | 34 | 32 | 3188.716669 | 3026.604022 | 0.60054 |
| FOXP3 | chrX | 49123711 | 10 | 10 | 965.7142857 | 980 | 0.60056 |
| GALNT2 | chr1 | 230418310 | 29 | 30 | 1058.607457 | 1140.596237 | 0.60066 |
| F7 | chr13 | 113751453 | 31 | 31 | 2734.883964 | 2773.524675 | 0.6007 |
| FOXP3 | chrX | 49129047 | 33 | 33 | 1726.069386 | 1668.442541 | 0.6008 |
| ICAM2 | chr17 | 62084360 | 29 | 30 | 1800.058952 | 1803.339468 | 0.6009 |
| BAX | chr19 | 49456830 | 29 | 28 | 2742.019092 | 2679.949495 | 0.60104 |
| CDKN2B-AS1 | chr9 | 21996368 | 37 | 31 | 1332.0317 | 1072.531635 | 0.60124 |
| NOS3 | chr7 | 150707728 | 2 | 4 | 53.33333333 | 162.782863 | 0.60124 |
| APOA1 | chr11 | 116702790 | 33 | 32 | 2586.762352 | 2468.533361 | 0.60128 |
| APOA1 | chr11 | 116664798 | 32 | 32 | 2977.715131 | 2953.559257 | 0.6014 |
| NOS3 | chr7 | 150710357 | 30 | 26 | 1292.60928 | 1176.03122 | 0.60142 |
| CDKN2B-AS1 | chr9 | 21966141 | 8 | 9 | 227.2840354 | 292.4404762 | 0.60142 |
| TERT | chr5 | 1253656 | 32 | 33 | 2733.250632 | 2877.837086 | 0.60188 |
| MCPH1 | chr8 | 6256422 | 32 | 32 | 2911.286909 | 2941.525994 | 0.60244 |
| ATF3 | chr1 | 212742686 | 36 | 32 | 2938.188436 | 2650.247987 | 0.60248 |

| | | | | | | | |
|---|---|---|---|---|---|---|---|
| DHFR | chr5 | 79953247 | 30 | 32 | 2820.064009 | 2957.741852 | 0.60272 |
| ALDOA | chr16 | 30060961 | 30 | 30 | 2747.179487 | 2704.959207 | 0.60308 |
| DHFR | chr5 | 79956775 | 6 | 2 | 566.6666667 | 200 | 0.60364 |
| GCK | chr7 | 44229489 | 32 | 33 | 2135.967247 | 2276.841138 | 0.60368 |
| ADRA2C | chr4 | 3764970 | 4 | 6 | 109.4444444 | 147.1428571 | 0.60386 |
| CDKN2B-AS1 | chr9 | 21968707 | 8 | 8 | 204.8408299 | 237.3581652 | 0.60386 |
| FOXP3 | chrX | 49121378 | 20 | 20 | 984.2712843 | 931.3763687 | 0.60402 |
| AKT2 | chr19 | 40800998 | 10 | 7 | 950.8333333 | 680 | 0.60434 |
| ESR1 | chr6 | 152003212 | 31 | 28 | 1073.355984 | 921.6492213 | 0.60442 |
| BAX | chr19 | 49465087 | 32 | 32 | 3012.399635 | 3057.343767 | 0.60448 |
| LDLR | chr19 | 11192701 | 22 | 23 | 2006.024531 | 2132.460317 | 0.60486 |
| LDLR | chr19 | 11194727 | 22 | 24 | 748.9286528 | 771.3796691 | 0.6049 |
| ESR1 | chr6 | 152011415 | 34 | 28 | 1483.702892 | 1172.032234 | 0.60494 |
| ATF3 | chr1 | 212742784 | 33 | 31 | 2882.35877 | 2752.120916 | 0.6051 |
| CDKN2B-AS1 | chr9 | 21977655 | 4 | 6 | 115.8333333 | 218.3333333 | 0.60552 |
| ABCG1 | chr21 | 43631000 | 2 | 3 | 53.33333333 | 140 | 0.60558 |
| FOXP3 | chrX | 49115042 | 14 | 9 | 1112.142857 | 750 | 0.6056 |
| ADRA2C | chr4 | 3761173 | 31 | 32 | 2852.640663 | 2891.893064 | 0.60562 |
| ABCG1 | chr21 | 43717483 | 32 | 32 | 2486.122724 | 2547.412809 | 0.60564 |
| FOXP3 | chrX | 49131196 | 34 | 32 | 2707.369366 | 2605.669415 | 0.60584 |
| COL15A1 | chr9 | 101696935 | 6 | 8 | 554.2424242 | 760 | 0.606 |
| HMGCR | chr5 | 74625929 | 32 | 32 | 2906.000292 | 2940.630176 | 0.60626 |
| APOA1 | chr11 | 116660149 | 3 | 3 | 88.57142857 | 127.5 | 0.60628 |
| KLF2 | chr19 | 16434523 | 35 | 32 | 3217.814096 | 2914.239359 | 0.60654 |
| F7 | chr13 | 113759771 | 31 | 31 | 2715.637369 | 2678.265875 | 0.60656 |
| CDKN2B-AS1 | chr9 | 22000827 | 4 | 3 | 97.5 | 85 | 0.60692 |
| APOA1 | chr11 | 116717439 | 32 | 32 | 2980.988024 | 3024.909066 | 0.60696 |
| APOA1 | chr11 | 116680846 | 31 | 32 | 2847.205122 | 2895.31521 | 0.60698 |
| CDKN2B-AS1 | chr9 | 21989449 | 21 | 25 | 708.3159708 | 889.2217866 | 0.60714 |
| ADRA2C | chr4 | 3767073 | 31 | 33 | 1668.699284 | 1718.673181 | 0.60786 |
| P2RY12 | chr3 | 151108195 | 32 | 32 | 2616.287993 | 2658.970999 | 0.60794 |
| LOXL1 | chr15 | 74209990 | 32 | 32 | 2937.212977 | 2967.130193 | 0.60806 |
| CDKN2B-AS1 | chr9 | 22006025 | 2 | 8 | 50.47619048 | 286.5401265 | 0.60866 |
| TMPO | chr12 | 98906524 | 24 | 22 | 2343.333333 | 2170.833333 | 0.60872 |
| F7 | chr13 | 113755374 | 32 | 32 | 3019.626408 | 2993.684317 | 0.6088 |
| FOXP3 | chrX | 49125571 | 4 | 2 | 133.3552632 | 117.5213675 | 0.60888 |
| KLF2 | chr19 | 16436148 | 3 | 3 | 96.66666667 | 80.55555555 | 0.6089 |
| NOS3 | chr7 | 150710072 | 27 | 29 | 1204.721553 | 1354.710996 | 0.6093 |
| TIMP3 | chr22 | 33186845 | 37 | 32 | 3491.600353 | 3041.00653 | 0.60938 |
| NOS3 | chr7 | 150708310 | 35 | 33 | 1865.049579 | 1703.153389 | 0.60968 |
| ESR2 | chr14 | 64807577 | 36 | 32 | 3497.182802 | 3083.129992 | 0.60972 |
| APOA1 | chr11 | 116720223 | 32 | 32 | 2940.253611 | 2907.813785 | 0.60986 |
| CTGF | chr6 | 132279347 | 15 | 15 | 1267.892108 | 1304.718615 | 0.60986 |
| FOXP3 | chrX | 49127095 | 20 | 16 | 1849.953102 | 1450.079365 | 0.60996 |
| NOS3 | chr7 | 150695425 | 13 | 8 | 974.4444445 | 558.0952381 | 0.61022 |
| APOA1 | chr11 | 116718264 | 34 | 32 | 3301.230743 | 3080.634974 | 0.61082 |
| CDKN1A | chr6 | 36644951 | 34 | 33 | 1797.608171 | 1690.015286 | 0.61086 |

| | | | | | | | |
|---|---|---|---|---|---|---|---|
| APOA1 | chr11 | 116678812 | 30 | 29 | 2934.086747 | 2813.284314 | 0.61094 |
| APOA1 | chr11 | 116691971 | 31 | 32 | 2955.35849 | 3020.80436 | 0.61102 |
| TERT | chr5 | 1299057 | 31 | 31 | 2936.533759 | 2900.713094 | 0.61108 |
| CDKN2B-AS1 | chr9 | 21990053 | 20 | 23 | 927.492785 | 1126.002886 | 0.61108 |
| BAX | chr19 | 49451093 | 3 | 3 | 262.5 | 240 | 0.6111 |
| COL15A1 | chr9 | 101701450 | 36 | 32 | 2697.058082 | 2347.317934 | 0.61126 |
| LDLR | chr19 | 11215918 | 31 | 30 | 2902.219364 | 2779.034674 | 0.61166 |
| F2RL3 | chr19 | 16997257 | 18 | 16 | 1697.619048 | 1544.642857 | 0.61224 |
| ADRA2C | chr4 | 3760005 | 31 | 33 | 2508.333129 | 2725.779082 | 0.6123 |
| F7 | chr13 | 113759909 | 33 | 30 | 2562.984188 | 2373.832666 | 0.61232 |
| CDKN1A | chr6 | 36654955 | 31 | 32 | 2765.440357 | 2789.034158 | 0.61232 |
| APOA1 | chr11 | 116667180 | 32 | 32 | 2983.277191 | 2952.933111 | 0.6126 |
| NOS3 | chr7 | 150690132 | 6 | 7 | 480 | 601.4285714 | 0.61288 |
| ABCG1 | chr21 | 43637473 | 37 | 32 | 3460.678855 | 3013.970062 | 0.61296 |
| ABCG1 | chr21 | 43636362 | 25 | 27 | 1711.381674 | 1789.308396 | 0.613 |
| FOXP3 | chrX | 49123231 | 4 | 4 | 340 | 363.3333333 | 0.6137 |
| ALOX15 | chr17 | 4549485 | 32 | 33 | 2024.883363 | 2142.069841 | 0.61384 |
| NOS3 | chr7 | 150695707 | 29 | 28 | 2775.497836 | 2707.342657 | 0.61394 |
| BCL2 | chr18 | 60996272 | 38 | 32 | 3642.868607 | 3052.042958 | 0.61398 |
| APOA1 | chr11 | 116680801 | 19 | 17 | 1772.651515 | 1543.650794 | 0.61402 |
| APOA1 | chr11 | 116695770 | 3 | 5 | 263.3333333 | 460 | 0.61408 |
| NOS3 | chr7 | 150678749 | 33 | 32 | 2646.544828 | 2606.530441 | 0.61424 |
| TERT | chr5 | 1300107 | 33 | 33 | 2587.065543 | 2635.61249 | 0.61436 |
| AKT2 | chr19 | 40796481 | 31 | 32 | 2928.791419 | 3049.852025 | 0.61452 |
| FGF2 | chr4 | 123747255 | 33 | 32 | 2748.260568 | 2623.427859 | 0.61458 |
| LCK | chr1 | 32741034 | 27 | 30 | 1027.32637 | 1207.428142 | 0.6146 |
| TIMP3 | chr22 | 33195165 | 2 | 4 | 48.57142857 | 162.2222222 | 0.61466 |
| APOA1 | chr11 | 116690124 | 30 | 26 | 1552.671356 | 1274.741318 | 0.61478 |
| LDLR | chr19 | 11218352 | 5 | 2 | 166.6666667 | 105 | 0.6151 |
| ALOX15 | chr17 | 4549910 | 29 | 27 | 2597.554113 | 2460.307225 | 0.6153 |
| NOS3 | chr7 | 150707095 | 34 | 32 | 3164.454466 | 3006.055214 | 0.6157 |
| APOA1 | chr11 | 116673587 | 27 | 28 | 2243.231899 | 2389.047619 | 0.6161 |
| CDKN2B-AS1 | chr9 | 22005770 | 27 | 25 | 962.7730028 | 842.3010489 | 0.61612 |
| FOXP3 | chrX | 49114934 | 6 | 5 | 353.3333333 | 268.5714286 | 0.61622 |
| PHACTR1 | chr6 | 12716655 | 39 | 33 | 2458.870271 | 2119.758709 | 0.61674 |
| NR3C1 | chr5 | 142786230 | 36 | 32 | 3505.476631 | 3103.038251 | 0.6175 |
| NOS3 | chr7 | 150688109 | 26 | 30 | 2226.084934 | 2463.524531 | 0.6176 |
| CDKN2BAS | chr9 | 22067558 | 32 | 32 | 2980.685665 | 2958.635794 | 0.61764 |
| APOA1 | chr11 | 116704697 | 37 | 32 | 3398.707217 | 2898.87502 | 0.61768 |
| APOA1 | chr11 | 116705601 | 23 | 25 | 1929.457764 | 2153.04447 | 0.61772 |
| ADRA2C | chr4 | 3766328 | 32 | 31 | 2771.322267 | 2645.510365 | 0.61792 |
| PLA2G7 | chr6 | 46707692 | 33 | 32 | 3077.039349 | 2955.128099 | 0.61794 |
| LOXL1 | chr15 | 74216605 | 19 | 17 | 709.537839 | 595.5504948 | 0.61794 |
| TERT | chr5 | 1301659 | 41 | 32 | 3500.649218 | 2755.035144 | 0.61828 |
| APOA1 | chr11 | 116719905 | 34 | 32 | 3276.793459 | 3104.276047 | 0.6187 |
| APOA1 | chr11 | 116669959 | 37 | 32 | 3369.377379 | 2933.922155 | 0.6188 |
| TIMP3 | chr22 | 33190773 | 34 | 32 | 2801.556862 | 2673.581232 | 0.6188 |

| | | | | | | | |
|---|---|---|---|---|---|---|---|
| APOA1 | chr11 | 116713524 | 27 | 22 | 2506.85251 | 2001.765734 | 0.61906 |
| LPA | chr6 | 160944079 | 29 | 29 | 2727.054155 | 2764.404762 | 0.6191 |
| PHACTR1 | chr6 | 12710382 | 38 | 33 | 2016.423661 | 1803.031532 | 0.61928 |
| APOA1 | chr11 | 116692311 | 33 | 31 | 3122.053158 | 2953.944123 | 0.61936 |
| TIMP3 | chr22 | 33187417 | 35 | 32 | 3280.467849 | 2973.224617 | 0.61966 |
| APOA1 | chr11 | 116681691 | 34 | 32 | 3092.402536 | 2879.705044 | 0.6197 |
| F7 | chr13 | 113758321 | 29 | 30 | 2758.455727 | 2826.142624 | 0.61976 |
| NOS3 | chr7 | 150706623 | 28 | 30 | 2221.674951 | 2318.628871 | 0.61978 |
| LDLR | chr19 | 11191367 | 2 | 8 | 180 | 664.6428571 | 0.61988 |
| AMOTL2 | chr3 | 134094145 | 13 | 10 | 348.4171384 | 307.6354843 | 0.61994 |
| APOA1 | chr11 | 116673299 | 24 | 25 | 2179.13961 | 2305.2442 | 0.62004 |
| LPA | chr6 | 11218145 | 40 | 32 | 3436.189708 | 2778.13401 | 0.62026 |
| LOXL1 | chr15 | 74244076 | 8 | 13 | 357.3092267 | 530.8477634 | 0.62058 |
| GALNT2 | chr1 | 230193689 | 28 | 26 | 1965.394759 | 1888.164058 | 0.62086 |
| APOA1 | chr11 | 116714244 | 36 | 32 | 3380.418498 | 2987.586338 | 0.62112 |
| APOA1 | chr11 | 116692269 | 33 | 32 | 3191.742467 | 3076.837065 | 0.62114 |
| ATF3 | chr1 | 212741186 | 33 | 32 | 3012.019199 | 2946.887989 | 0.62114 |
| ADRA2C | chr4 | 3761581 | 26 | 30 | 2264.990773 | 2669.642718 | 0.62122 |
| AKT2 | chr19 | 40800771 | 37 | 32 | 3153.431507 | 2752.94653 | 0.62126 |
| CDKN2B-AS1 | chr9 | 21970915 | 4 | 5 | 122.5108225 | 185.4153767 | 0.6216 |
| FGF2 | chr4 | 123743528 | 35 | 32 | 2945.172817 | 2615.31128 | 0.62166 |
| F7 | chr13 | 113751057 | 16 | 16 | 1588.333333 | 1563.888889 | 0.62214 |
| KLF2 | chr19 | 16432781 | 33 | 32 | 3080.809607 | 3015.470608 | 0.6223 |
| CDKN2B-AS1 | chr9 | 21971160 | 8 | 11 | 255.2625153 | 324.2324342 | 0.6226 |
| LCK | chr1 | 32708341 | 2 | 5 | 60.45454545 | 215.8439231 | 0.6226 |
| F7 | chr13 | 113757126 | 35 | 33 | 2801.782155 | 2593.651295 | 0.62302 |
| ICAM2 | chr17 | 62093817 | 35 | 26 | 1231.637437 | 944.6222947 | 0.62306 |
| FOXP3 | chrX | 49125139 | 32 | 33 | 2140.490491 | 2153.559696 | 0.62308 |
| TMPO | chr12 | 98905964 | 10 | 7 | 954.7619048 | 683.3333333 | 0.62328 |
| APOA1 | chr11 | 116701621 | 7 | 8 | 555.7142857 | 669.5238095 | 0.6235 |
| TERT | chr5 | 1295714 | 13 | 16 | 406.8667444 | 472.4075189 | 0.62368 |
| F7 | chr13 | 113752551 | 32 | 32 | 2958.363977 | 2873.158427 | 0.62372 |
| ICAM2 | chr17 | 62084115 | 33 | 32 | 2740.169213 | 2607.116576 | 0.62376 |
| BCL2 | chr18 | 60995873 | 10 | 9 | 886.3888889 | 818.0952381 | 0.62378 |
| ALOX15 | chr17 | 4545438 | 32 | 33 | 1718.24247 | 1716.88749 | 0.6238 |
| ALOX15 | chr17 | 4551661 | 21 | 21 | 1074.50938 | 1117.972583 | 0.62392 |
| ADRA2C | chr4 | 3768690 | 31 | 32 | 2306.045711 | 2308.259992 | 0.6242 |
| AKT2 | chr19 | 40799035 | 7 | 8 | 509.0476191 | 538.3549784 | 0.6242 |
| APOA1 | chr11 | 116680449 | 27 | 30 | 2110.954881 | 2247.470414 | 0.62442 |
| NR3C1 | chr5 | 142791387 | 32 | 31 | 3019.292754 | 2958.840059 | 0.62442 |
| ADRA2C | chr4 | 3760206 | 37 | 32 | 3481.840305 | 3031.85927 | 0.62458 |
| F2RL3 | chr19 | 16999195 | 3 | 7 | 283.3333333 | 626.6666667 | 0.62476 |
| ALDOA | chr16 | 30078077 | 10 | 9 | 260.8736433 | 261.4494479 | 0.62486 |
| P2RY12 | chr3 | 151105776 | 10 | 8 | 341.2759171 | 250.7142857 | 0.62494 |
| CDKN2B-AS1 | chr9 | 22001938 | 33 | 31 | 1277.585774 | 1247.594056 | 0.62502 |
| ALOX15 | chr17 | 4554309 | 23 | 20 | 2255.651154 | 1919.047619 | 0.62514 |
| P2RY12 | chr3 | 151107738 | 36 | 32 | 3384.648907 | 3022.414483 | 0.62522 |

| | | | | | | | |
|---|---|---|---|---|---|---|---|
| LDLR | chr19 | 11216124 | 28 | 31 | 2623.383208 | 2939.220143 | 0.62524 |
| ABCA1 | chr9 | 107698613 | 34 | 32 | 2861.272327 | 2605.172574 | 0.6256 |
| E2F6 | chr2 | 11614372 | 36 | 32 | 3276.331155 | 2892.055328 | 0.6258 |
| ESR1 | chr6 | 152424266 | 36 | 32 | 3365.905895 | 3010.095467 | 0.6263 |
| APOA1 | chr11 | 116725011 | 17 | 24 | 637.6683502 | 848.8021374 | 0.62674 |
| ABCA1 | chr9 | 107692515 | 34 | 33 | 2735.964388 | 2599.267554 | 0.62678 |
| PLA2G7 | chr6 | 46710397 | 32 | 32 | 2924.926001 | 2955.107794 | 0.62694 |
| CDKN2B-AS1 | chr9 | 22000829 | 3 | 3 | 68.57142857 | 80 | 0.6272 |
| GCK | chr7 | 44232540 | 37 | 32 | 3570.435938 | 3060.629257 | 0.62722 |
| E2F6 | chr2 | 11611796 | 23 | 26 | 2240.796391 | 2552.147852 | 0.6274 |
| LCK | chr1 | 32741933 | 26 | 24 | 2421.150794 | 2262.936508 | 0.62752 |
| F7 | chr13 | 113751014 | 22 | 21 | 2165.330087 | 2019.642857 | 0.62754 |
| APOA1 | chr11 | 116726167 | 31 | 29 | 2861.480464 | 2706.610987 | 0.62762 |
| LDLR | chr19 | 11197965 | 16 | 15 | 1514.529915 | 1443.888889 | 0.62778 |
| PECAM1 | chr17 | 62399960 | 36 | 32 | 3172.165785 | 2844.419335 | 0.6281 |
| GALNT2 | chr1 | 230193750 | 32 | 33 | 2258.426532 | 2256.412002 | 0.62812 |
| NR3C1 | chr5 | 142789348 | 39 | 32 | 3665.56454 | 2990.943708 | 0.62822 |
| CDKN2B-AS1 | chr9 | 21970314 | 7 | 8 | 263.3333333 | 270 | 0.62822 |
| PECAM1 | chr17 | 62396931 | 31 | 32 | 2843.159994 | 2963.154272 | 0.62888 |
| LPA | chr6 | 11218487 | 35 | 32 | 3237.649914 | 2927.916763 | 0.62894 |
| AKT2 | chr19 | 40791877 | 22 | 22 | 590.4474783 | 619.0576802 | 0.629 |
| LDLR | chr19 | 11224914 | 31 | 32 | 3058.493055 | 3136.938264 | 0.6291 |
| ESR1 | chr6 | 152010088 | 34 | 33 | 2980.270692 | 2852.141547 | 0.62928 |
| CDKN2B-AS1 | chr9 | 22007337 | 30 | 31 | 1953.051671 | 2074.208233 | 0.62932 |
| PHACTR1 | chr6 | 13276102 | 2 | 3 | 60 | 145 | 0.62934 |
| FGF2 | chr4 | 123747297 | 31 | 31 | 2802.779341 | 2767.498133 | 0.62978 |
| F7 | chr13 | 113775145 | 25 | 28 | 2041.126235 | 2347.649295 | 0.6299 |
| GCK | chr7 | 44183643 | 35 | 32 | 3096.50534 | 2863.237675 | 0.62992 |
| APOA1 | chr11 | 116661297 | 32 | 31 | 2795.904579 | 2742.255794 | 0.63 |
| APOA1 | chr11 | 116691955 | 31 | 32 | 2778.654514 | 2910.595382 | 0.63034 |
| F7 | chr13 | 113752540 | 32 | 32 | 3102.866121 | 3012.11007 | 0.63058 |
| ALOX15 | chr17 | 4547413 | 35 | 33 | 3119.493103 | 2899.848721 | 0.63064 |
| APOA1 | chr11 | 116720400 | 30 | 27 | 947.5134129 | 887.9251968 | 0.63076 |
| ESR1 | chr6 | 152001997 | 25 | 26 | 2411.04798 | 2477.02381 | 0.63094 |
| COL15A1 | chr9 | 101700590 | 38 | 32 | 3338.688991 | 2838.702421 | 0.63096 |
| PECAM1 | chr17 | 62408378 | 34 | 32 | 2684.191276 | 2486.475967 | 0.63136 |
| MIA3 | chr1 | 222809407 | 24 | 26 | 1871.486569 | 2104.859307 | 0.63138 |
| E2F6 | chr2 | 11611661 | 27 | 29 | 2561.290043 | 2796.901154 | 0.63158 |
| LDLR | chr19 | 11220036 | 29 | 31 | 2809.64591 | 2972.892108 | 0.63164 |
| APOA1 | chr11 | 116681640 | 32 | 32 | 3000.765853 | 3053.780042 | 0.63166 |
| CTGF | chr6 | 132272352 | 2 | 4 | 55 | 153.5714286 | 0.63174 |
| CTGF | chr6 | 132276797 | 35 | 32 | 3080.049665 | 2844.297559 | 0.63182 |
| E2F6 | chr2 | 11608332 | 32 | 32 | 3001.070881 | 3025.563393 | 0.63186 |
| F7 | chr13 | 113750500 | 26 | 25 | 2506.742424 | 2431.269841 | 0.63188 |
| COL15A1 | chr9 | 101705734 | 33 | 32 | 2957.258032 | 2892.85801 | 0.63228 |
| APOA1 | chr11 | 116688539 | 34 | 32 | 3197.67677 | 3037.999755 | 0.63234 |
| PLA2G7 | chr6 | 46705365 | 26 | 26 | 1099.505984 | 1153.135476 | 0.63252 |

| | | | | | | | |
|---|---|---|---|---|---|---|---|
| LPA | chr6 | 160949113 | 34 | 33 | 2386.992859 | 2257.27186 | 0.6327 |
| ICAM2 | chr17 | 62084343 | 31 | 30 | 1857.914298 | 1743.726253 | 0.63282 |
| COL15A1 | chr9 | 101696979 | 23 | 28 | 2253.571429 | 2724.632035 | 0.63284 |
| GALNT2 | chr1 | 230186010 | 21 | 12 | 1020.915794 | 628.701422 | 0.63334 |
| COL15A1 | chr9 | 101703951 | 19 | 13 | 1678.095238 | 1105.357143 | 0.63356 |
| P2RY12 | chr3 | 151107890 | 35 | 32 | 3173.787813 | 2863.35615 | 0.63422 |
| APOA1 | chr11 | 116699681 | 32 | 33 | 1959.267977 | 2084.307772 | 0.63426 |
| CDKN1A | chr6 | 36652242 | 33 | 32 | 2887.343116 | 2834.197323 | 0.63428 |
| ADRA2C | chr4 | 3768666 | 30 | 32 | 1147.650381 | 1171.318902 | 0.63512 |
| TIMP3 | chr22 | 33188325 | 35 | 32 | 3284.843922 | 2974.387428 | 0.63518 |
| CDKN2B-AS1 | chr9 | 22005755 | 26 | 25 | 820.179236 | 817.0141794 | 0.63538 |
| FGF2 | chr4 | 123747249 | 33 | 32 | 3118.711408 | 3000.020953 | 0.6354 |
| NOS3 | chr7 | 150705778 | 35 | 33 | 2229.862699 | 2164.79658 | 0.63572 |
| LDLR | chr19 | 11219854 | 7 | 14 | 685.7142857 | 1350 | 0.63574 |
| NOS3 | chr7 | 150687471 | 36 | 32 | 2998.105411 | 2700.038791 | 0.63582 |
| TERT | chr5 | 1295852 | 32 | 33 | 1823.584838 | 1937.905792 | 0.63582 |
| CDKN2B-AS1 | chr9 | 21968126 | 29 | 27 | 969.3215016 | 869.4775363 | 0.63602 |
| BAX | chr19 | 49454708 | 13 | 19 | 1159.047619 | 1732.178932 | 0.63606 |
| ABCA1 | chr9 | 107698295 | 33 | 33 | 2578.706874 | 2634.518808 | 0.63618 |
| APOA1 | chr11 | 116712759 | 18 | 15 | 1540.663781 | 1317.936508 | 0.63626 |
| ADRA2C | chr4 | 3761565 | 22 | 28 | 1846.685907 | 2410.484099 | 0.63642 |
| KLF2 | chr19 | 16437873 | 25 | 27 | 988.3230964 | 1011.557381 | 0.63648 |
| TMPO | chr12 | 98903420 | 37 | 31 | 3254.64665 | 2791.668553 | 0.63668 |
| CDKN2B-AS1 | chr9 | 21981748 | 20 | 21 | 1168.896659 | 1170.287539 | 0.6367 |
| APOA1 | chr11 | 116685587 | 33 | 33 | 1629.757613 | 1570.842686 | 0.63696 |
| ARHGEF18 | chr19 | 7463844 | 9 | 11 | 231.4146027 | 313.5088736 | 0.6372 |
| ADRA2C | chr4 | 3763137 | 27 | 27 | 2049.261268 | 2106.510712 | 0.63724 |
| ADRA2C | chr4 | 3766420 | 35 | 32 | 2834.558145 | 2640.023475 | 0.63768 |
| FOXP3 | chrX | 49127110 | 20 | 18 | 1579.592352 | 1384.047619 | 0.63772 |
| LDLR | chr19 | 11216275 | 4 | 4 | 290 | 313.3333333 | 0.63794 |
| ATF3 | chr1 | 212742065 | 34 | 32 | 3139.392986 | 2978.558593 | 0.63806 |
| APOA1 | chr11 | 116711634 | 38 | 32 | 3621.908011 | 3068.36351 | 0.63828 |
| CDKN2B-AS1 | chr9 | 21988682 | 24 | 21 | 734.339602 | 676.8253668 | 0.63836 |
| ADRA2C | chr4 | 3764216 | 19 | 14 | 1097.276635 | 754.428093 | 0.63842 |
| E2F6 | chr2 | 11584875 | 40 | 32 | 3866.084785 | 3104.217444 | 0.6385 |
| CDKN2B-AS1 | chr9 | 21973528 | 25 | 24 | 738.5553759 | 740.3934438 | 0.63852 |
| APOA1 | chr11 | 116718430 | 34 | 32 | 3029.435311 | 2821.861464 | 0.63918 |
| ATF3 | chr1 | 212742587 | 30 | 30 | 2802.352092 | 2769.775542 | 0.6395 |
| F7 | chr13 | 113751431 | 30 | 32 | 2904.889971 | 3120.597736 | 0.63958 |
| ATF3 | chr1 | 212746339 | 14 | 15 | 1245.147908 | 1365.595238 | 0.63964 |
| LPA | chr6 | 160948254 | 14 | 16 | 1262.261905 | 1408.809524 | 0.63996 |
| ALOX15 | chr17 | 4534071 | 21 | 20 | 1752.301587 | 1626.349206 | 0.64004 |
| F7 | chr13 | 113750819 | 32 | 30 | 3156.973859 | 2909.208847 | 0.6401 |
| NOS3 | chr7 | 150693657 | 6 | 10 | 178.0555556 | 337.1428571 | 0.64042 |
| F7 | chr13 | 113751107 | 7 | 7 | 613.3333333 | 636.547619 | 0.64046 |
| AMOTL2 | chr3 | 134096596 | 5 | 5 | 457.9365079 | 442.2619048 | 0.64058 |
| GALNT2 | chr1 | 230193087 | 34 | 32 | 2662.209798 | 2459.062452 | 0.64064 |

| | | | | | | | |
|---|---|---|---|---|---|---|---|
| CDKN2B-AS1 | chr9 | 21967651 | 2 | 3 | 60 | 138.0706288 | 0.64064 |
| ADRA2C | chr4 | 3762285 | 33 | 33 | 2910.066076 | 2866.244726 | 0.64072 |
| GCK | chr7 | 44230724 | 33 | 32 | 2275.595418 | 2252.148563 | 0.6408 |
| CDKN2B-AS1 | chr9 | 22011030 | 12 | 12 | 322.8240978 | 346.2530284 | 0.6408 |
| F7 | chr13 | 113755432 | 31 | 32 | 2802.638391 | 2855.531968 | 0.6411 |
| TERT | chr5 | 1300694 | 33 | 32 | 3037.987994 | 2971.350675 | 0.6412 |
| LDLR | chr19 | 11194256 | 31 | 30 | 2805.301284 | 2667.040043 | 0.64126 |
| CDKN2B-AS1 | chr9 | 22011092 | 14 | 11 | 451.6946223 | 323.2490952 | 0.64126 |
| MCPH1 | chr8 | 6260934 | 28 | 32 | 1771.011952 | 1931.812955 | 0.64134 |
| AKT2 | chr19 | 40798792 | 31 | 32 | 2923.087462 | 2966.862112 | 0.64136 |
| CDKN2B-AS1 | chr9 | 22001688 | 36 | 32 | 3162.843004 | 2784.49113 | 0.64168 |
| F7 | chr13 | 113750524 | 20 | 22 | 1840.119048 | 2073.174603 | 0.6418 |
| F7 | chr13 | 113750585 | 22 | 25 | 1671.111111 | 1978.424076 | 0.642 |
| BAX | chr19 | 49465385 | 31 | 32 | 1544.927881 | 1534.534619 | 0.6421 |
| LPA | chr6 | 11216473 | 35 | 31 | 3388.119315 | 2988.560435 | 0.64254 |
| TERT | chr5 | 1253383 | 26 | 30 | 2241.928192 | 2533.54798 | 0.64298 |
| APOA1 | chr11 | 116726410 | 17 | 21 | 530.0876803 | 702.7886656 | 0.64336 |
| F7 | chr13 | 113755437 | 29 | 32 | 2572.804022 | 2791.376956 | 0.64356 |
| FOXP3 | chrX | 49125657 | 2 | 2 | 83.80952381 | 121.6783217 | 0.64356 |
| ESR1 | chr6 | 152424341 | 34 | 32 | 3157.445233 | 2994.595825 | 0.6437 |
| LPA | chr6 | 11216780 | 40 | 32 | 3798.747448 | 3020.708233 | 0.64374 |
| TERT | chr5 | 1299660 | 14 | 11 | 1301.269841 | 1002.261905 | 0.64382 |
| CDKN2B-AS1 | chr9 | 22006055 | 4 | 6 | 122.4269006 | 230.0649351 | 0.64396 |
| NOS3 | chr7 | 150707058 | 31 | 32 | 2478.534434 | 2598.83219 | 0.64418 |
| CDKN1A | chr6 | 36643494 | 24 | 20 | 2330.767762 | 1956.666667 | 0.6445 |
| NR3C1 | chr5 | 142790271 | 37 | 33 | 2276.653287 | 1997.382757 | 0.64452 |
| NOS3 | chr7 | 150679043 | 13 | 13 | 586.9102871 | 617.1938638 | 0.6449 |
| MIA3 | chr1 | 222792449 | 4 | 4 | 123.5209235 | 163.0769231 | 0.6449 |
| LOXL1 | chr15 | 74244312 | 27 | 30 | 1385.44689 | 1482.849677 | 0.64498 |
| NOS3 | chr7 | 150695715 | 29 | 28 | 2784.285904 | 2658.174603 | 0.64514 |
| ESR1 | chr6 | 152007462 | 34 | 32 | 3250.190309 | 3084.007642 | 0.6452 |
| NOS3 | chr7 | 150710713 | 5 | 2 | 168.3333333 | 58.33333333 | 0.6452 |
| GALNT2 | chr1 | 230190320 | 30 | 31 | 2786.793712 | 2913.772771 | 0.64524 |
| LOXL1 | chr15 | 74218268 | 21 | 25 | 1765.790043 | 2146.607143 | 0.64526 |
| APOA1 | chr11 | 116720249 | 34 | 33 | 1992.408759 | 1977.234951 | 0.64536 |
| AKT2 | chr19 | 40800068 | 32 | 32 | 2550.615733 | 2595.603329 | 0.64538 |
| NR3C1 | chr5 | 142786486 | 38 | 32 | 3658.369126 | 3067.659742 | 0.64564 |
| NOS3 | chr7 | 150706003 | 29 | 30 | 1063.814254 | 1150.226165 | 0.6457 |
| ESR1 | chr6 | 152002499 | 31 | 32 | 2542.579141 | 2664.745965 | 0.64572 |
| FGF2 | chr4 | 123746820 | 35 | 32 | 3131.827181 | 2838.068832 | 0.64578 |
| APOA1 | chr11 | 116662413 | 27 | 31 | 1528.43832 | 1828.752498 | 0.64578 |
| APOA1 | chr11 | 116690507 | 7 | 4 | 201.4393939 | 141.2987013 | 0.64584 |
| ABCG1 | chr21 | 43636979 | 33 | 32 | 3051.736527 | 2987.215612 | 0.64592 |
| APOA1 | chr11 | 116661339 | 31 | 33 | 2365.109415 | 2567.647345 | 0.64596 |
| NOS3 | chr7 | 150692395 | 32 | 31 | 2537.913802 | 2409.878032 | 0.64632 |
| APOA1 | chr11 | 116718652 | 33 | 32 | 3226.86404 | 3110.746441 | 0.6469 |
| NR3C1 | chr5 | 142785960 | 14 | 13 | 964.978355 | 858.3333333 | 0.64698 |

| | | | | | | | |
|---|---|---|---|---|---|---|---|
| KLF2 | chr19 | 16434677 | 29 | 31 | 1703.238901 | 1759.351789 | 0.64712 |
| APOA1 | chr11 | 116712771 | 24 | 21 | 2121.997724 | 1904.123377 | 0.64712 |
| LPA | chr6 | 160947120 | 35 | 32 | 3333.640411 | 3033.823172 | 0.64738 |
| BCL2 | chr18 | 60996180 | 38 | 32 | 3291.022728 | 2796.332313 | 0.64744 |
| CDKN2B-AS1 | chr9 | 21971039 | 13 | 6 | 333.8071895 | 173.1457431 | 0.6476 |
| TERT | chr5 | 1296447 | 35 | 32 | 2910.239996 | 2627.133169 | 0.64776 |
| LPA | chr6 | 11221344 | 28 | 30 | 2525.827686 | 2743.194722 | 0.64782 |
| APOA1 | chr11 | 116717148 | 32 | 32 | 3091.527109 | 3106.555628 | 0.64788 |
| ABCB1 | chr7 | 87345891 | 39 | 32 | 3650.005424 | 2977.535917 | 0.648 |
| APOA1 | chr11 | 116661363 | 33 | 32 | 2920.598691 | 2795.716163 | 0.64832 |
| BCL2 | chr18 | 60994747 | 36 | 32 | 3361.625583 | 2972.063683 | 0.6485 |
| CTGF | chr6 | 132278483 | 36 | 32 | 2644.110072 | 2279.087806 | 0.6487 |
| ESR1 | chr6 | 152011656 | 3 | 5 | 80 | 185 | 0.64924 |
| ABCA1 | chr9 | 107693842 | 38 | 32 | 3529.214029 | 2995.278268 | 0.64936 |
| GCK | chr7 | 44229556 | 32 | 32 | 3078.474923 | 3060.579487 | 0.64966 |
| APOA1 | chr11 | 116692222 | 33 | 33 | 3011.144716 | 2970.097364 | 0.64984 |
| CDKN2B-AS1 | chr9 | 21970463 | 18 | 18 | 588.8473507 | 553.264827 | 0.64996 |
| CDKN2B-AS1 | chr9 | 21996274 | 27 | 25 | 881.6685562 | 851.6462881 | 0.6501 |
| APOA1 | chr11 | 116699357 | 8 | 2 | 282.8571429 | 120 | 0.65074 |
| BAX | chr19 | 49465059 | 30 | 30 | 2830.568182 | 2868.336623 | 0.65076 |
| CDKN1A | chr6 | 36643854 | 31 | 23 | 2534.933122 | 1919.490232 | 0.65082 |
| NOS3 | chr7 | 150692222 | 9 | 13 | 262.252886 | 417.6337785 | 0.65118 |
| ARHGEF18 | chr19 | 7469506 | 18 | 25 | 1730 | 2438.484848 | 0.6513 |
| DHFR | chr5 | 79956767 | 8 | 7 | 688.8888889 | 560 | 0.65154 |
| ALDOA | chr16 | 30061354 | 25 | 24 | 2315.313853 | 2260.932401 | 0.65158 |
| FGF2 | chr4 | 123745565 | 34 | 32 | 3220.744429 | 3012.199926 | 0.65162 |
| CDKN1A | chr6 | 36650467 | 32 | 28 | 1571.154555 | 1336.497604 | 0.65166 |
| APOA1 | chr11 | 116690256 | 33 | 32 | 3137.489228 | 3059.161772 | 0.65198 |
| PECAM1 | chr17 | 62407117 | 12 | 11 | 1114.736842 | 994.7619048 | 0.65252 |
| LDLR | chr19 | 11220911 | 10 | 11 | 930 | 983.3333333 | 0.65286 |
| CDKN1A | chr6 | 36651994 | 31 | 31 | 2973.66987 | 3001.59628 | 0.65292 |
| ABCA1 | chr9 | 107692954 | 32 | 32 | 2857.647674 | 2893.032825 | 0.65302 |
| ATF3 | chr1 | 212741008 | 34 | 32 | 2903.088124 | 2783.9904 | 0.6532 |
| LDLR | chr19 | 11221716 | 27 | 27 | 2403.175991 | 2444.490232 | 0.65326 |
| NOS3 | chr7 | 150710386 | 12 | 15 | 428.8609146 | 578.3590103 | 0.65344 |
| CDKN1A | chr6 | 36652117 | 26 | 30 | 2450.375458 | 2854.197608 | 0.6539 |
| ALDOA | chr16 | 30060974 | 26 | 28 | 2463.054446 | 2678.774836 | 0.65394 |
| LPA | chr6 | 11216362 | 33 | 32 | 3091.291139 | 3020.493052 | 0.65406 |
| FOXP3 | chrX | 49127041 | 18 | 13 | 1589.880952 | 1176.428571 | 0.6541 |
| CDKN2B-AS1 | chr9 | 21974035 | 7 | 11 | 228.5714286 | 327.2727273 | 0.65422 |
| APOA1 | chr11 | 116661291 | 32 | 31 | 3018.032832 | 2903.449393 | 0.65454 |
| NOS3 | chr7 | 150682948 | 17 | 16 | 1064.307359 | 1052.380952 | 0.65494 |
| F7 | chr13 | 113756928 | 31 | 32 | 2895.905357 | 2961.508859 | 0.65538 |
| NOS3 | chr7 | 150695619 | 20 | 19 | 1697.698413 | 1661.904762 | 0.6556 |
| CDKN2B-AS1 | chr9 | 21974171 | 9 | 8 | 267.8679654 | 270.4040404 | 0.65562 |
| KLF2 | chr19 | 16437869 | 22 | 28 | 936.5409104 | 1137.384944 | 0.65564 |
| ADRA2C | chr4 | 3759197 | 32 | 32 | 2799.137676 | 2753.127869 | 0.65566 |

| | | | | | | | |
|---|---|---|---|---|---|---|---|
| F7 | chr13 | 113750566 | 26 | 25 | 2246.538462 | 2106.028139 | 0.65612 |
| ESR1 | chr6 | 152006654 | 31 | 31 | 2932.172185 | 2955.219421 | 0.6562 |
| ADRA2C | chr4 | 3759203 | 32 | 33 | 2639.48362 | 2663.219686 | 0.65628 |
| CTGF | chr6 | 132276073 | 33 | 32 | 2395.418662 | 2275.840303 | 0.65634 |
| ARHGEF18 | chr19 | 7469490 | 8 | 11 | 769.047619 | 1072.222222 | 0.65636 |
| CDKN2B-AS1 | chr9 | 22005226 | 5 | 6 | 139.9350649 | 199.9098124 | 0.6564 |
| ICAM2 | chr17 | 62084040 | 31 | 32 | 2303.905181 | 2427.930661 | 0.65646 |
| APOA1 | chr11 | 116689986 | 33 | 32 | 3059.527109 | 2988.682059 | 0.65672 |
| MIA3 | chr1 | 222792345 | 17 | 18 | 524.8654854 | 596.7113438 | 0.65704 |
| CTGF | chr6 | 132277083 | 32 | 31 | 2769.692209 | 2644.490002 | 0.6573 |
| NOS3 | chr7 | 150708912 | 13 | 12 | 1225.075758 | 1112.380952 | 0.65746 |
| F2RL3 | chr19 | 16998268 | 14 | 10 | 489.2919799 | 319.2857143 | 0.65748 |
| ALOX15 | chr17 | 4552943 | 32 | 33 | 2597.888394 | 2633.384424 | 0.65752 |
| F2RL3 | chr19 | 16998974 | 32 | 31 | 2903.78314 | 2788.90096 | 0.65754 |
| MCPH1 | chr8 | 6262871 | 36 | 32 | 3154.22589 | 2783.634366 | 0.65766 |
| F2RL3 | chr19 | 16990490 | 29 | 31 | 2793.568197 | 2950.847127 | 0.65782 |
| ADRA2C | chr4 | 3762043 | 27 | 30 | 2177.215729 | 2471.630781 | 0.65828 |
| LDLR | chr19 | 11220668 | 13 | 16 | 1240.714286 | 1546.666667 | 0.65836 |
| ICAM2 | chr17 | 62084349 | 30 | 29 | 1414.237138 | 1314.26957 | 0.65848 |
| GCK | chr7 | 44234861 | 5 | 8 | 227.1997304 | 402.9798224 | 0.6588 |
| ICAM2 | chr17 | 62087088 | 24 | 18 | 737.5262726 | 585.5882026 | 0.65896 |
| DHFR | chr5 | 79951639 | 34 | 32 | 2959.953781 | 2814.201677 | 0.659 |
| TERT | chr5 | 1300379 | 29 | 25 | 2291.750262 | 1927.540112 | 0.65902 |
| PECAM1 | chr17 | 62401032 | 34 | 32 | 3087.581345 | 2881.995542 | 0.65908 |
| NOS3 | chr7 | 150693990 | 10 | 11 | 299.8443224 | 366.0927961 | 0.6591 |
| APOA1 | chr11 | 116707401 | 15 | 10 | 1345.555556 | 862.5396825 | 0.65916 |
| CDKN1A | chr6 | 36645648 | 4 | 10 | 189.5238095 | 436.468254 | 0.65924 |
| CDKN1A | chr6 | 36643466 | 13 | 12 | 1071.742424 | 1040 | 0.65962 |
| LCK | chr1 | 32708530 | 21 | 25 | 1691.31746 | 1961.723554 | 0.65968 |
| APOA1 | chr11 | 116685304 | 33 | 33 | 2259.438079 | 2318.667703 | 0.65996 |
| COL15A1 | chr9 | 101703431 | 34 | 32 | 3216.857336 | 3002.042634 | 0.6601 |
| BAX | chr19 | 49449869 | 30 | 29 | 2812.342519 | 2692.813174 | 0.66012 |
| NOS3 | chr7 | 150686169 | 17 | 10 | 611.9530908 | 398.3882784 | 0.66038 |
| P2RY12 | chr3 | 151110762 | 36 | 32 | 3394.505524 | 2993.6575 | 0.66046 |
| CDKN2B-AS1 | chr9 | 21974015 | 4 | 4 | 97.14285714 | 105.5769231 | 0.6608 |
| APOA1 | chr11 | 116697168 | 30 | 29 | 2795.670996 | 2671.459482 | 0.66082 |
| GALNT2 | chr1 | 230418169 | 7 | 8 | 188.6456384 | 248.5807742 | 0.66082 |
| PECAM1 | chr17 | 62398427 | 30 | 30 | 1251.539428 | 1197.790452 | 0.6609 |
| GALNT2 | chr1 | 230185513 | 16 | 8 | 781.463455 | 362.4969397 | 0.66112 |
| ADRA2C | chr4 | 3759996 | 33 | 33 | 2760.955046 | 2707.347743 | 0.66184 |
| APOA1 | chr11 | 116712731 | 6 | 3 | 449.1666667 | 207.1428571 | 0.6623 |
| COL15A1 | chr9 | 101704474 | 31 | 32 | 2900.854454 | 3018.500418 | 0.66238 |
| PLA2G7 | chr6 | 46713336 | 37 | 32 | 3039.612381 | 2601.734486 | 0.6628 |
| CDKN2B-AS1 | chr9 | 21966279 | 5 | 3 | 157.5541126 | 124.4444444 | 0.6628 |
| ABCG1 | chr21 | 43717733 | 39 | 33 | 2568.771346 | 2207.11765 | 0.66308 |
| NOS3 | chr7 | 150692296 | 6 | 11 | 171.6133866 | 351.1312518 | 0.66322 |
| ICAM2 | chr17 | 62084218 | 19 | 16 | 640.0308902 | 570.9341463 | 0.66326 |

| | | | | | | | |
|---|---|---|---|---|---|---|---|
| APOA1 | chr11 | 116702671 | 32 | 31 | 2805.452393 | 2682.18466 | 0.66344 |
| F2RL3 | chr19 | 16990822 | 34 | 32 | 3289.335117 | 3108.628314 | 0.66366 |
| APOA1 | chr11 | 116701111 | 23 | 24 | 2228.214286 | 2343.214286 | 0.6639 |
| TIMP3 | chr22 | 33193714 | 34 | 32 | 3165.964815 | 3001.803017 | 0.66394 |
| TMPO | chr12 | 98899733 | 35 | 32 | 3287.239415 | 2977.545549 | 0.66396 |
| LDLR | chr19 | 11220433 | 4 | 3 | 363.3333333 | 283.3333333 | 0.66442 |
| ADRA2C | chr4 | 3768617 | 16 | 19 | 490.8984594 | 550.0729478 | 0.6645 |
| CDKN2B-AS1 | chr9 | 21990091 | 11 | 10 | 356.001221 | 301.031746 | 0.66476 |
| APOA1 | chr11 | 116659138 | 5 | 10 | 124.5962733 | 292.7014214 | 0.66476 |
| BCL2 | chr18 | 60988675 | 18 | 16 | 694.1984127 | 577.6587301 | 0.66494 |
| CDKN2B-AS1 | chr9 | 21967988 | 32 | 28 | 1220.83854 | 1032.781131 | 0.66518 |
| LPA | chr6 | 11216176 | 33 | 32 | 3159.475323 | 3042.337209 | 0.66522 |
| LOXL1 | chr15 | 74218418 | 34 | 32 | 3113.040666 | 2901.719683 | 0.66554 |
| ARHGEF18 | chr19 | 7528627 | 12 | 20 | 1126.547619 | 1905.714286 | 0.6656 |
| LOXL1 | chr15 | 74211117 | 34 | 32 | 3269.978394 | 3092.383031 | 0.6659 |
| APOA1 | chr11 | 116712846 | 34 | 31 | 3223.110273 | 2957.918318 | 0.66598 |
| FGF2 | chr4 | 123741608 | 30 | 30 | 2349.860344 | 2392.854603 | 0.66604 |
| ADRA2C | chr4 | 3768608 | 28 | 29 | 898.8035233 | 899.4522284 | 0.66608 |
| CDKN2B-AS1 | chr9 | 21989805 | 12 | 14 | 360.3663004 | 457.1101121 | 0.66618 |
| LOXL1 | chr15 | 74213871 | 23 | 20 | 1366.539036 | 1265.466712 | 0.66622 |
| PECAM1 | chr17 | 62396971 | 30 | 28 | 2839.917027 | 2676.923077 | 0.66646 |
| F7 | chr13 | 113757172 | 30 | 29 | 2660.663934 | 2541.466046 | 0.66668 |
| TIMP3 | chr22 | 33190339 | 34 | 32 | 2593.047529 | 2408.408404 | 0.66696 |
| GALNT2 | chr1 | 230193883 | 23 | 26 | 1819.876983 | 2006.067821 | 0.6671 |
| MCPH1 | chr8 | 6260870 | 33 | 33 | 2467.863833 | 2510.533195 | 0.66746 |
| LDLR | chr19 | 11191004 | 35 | 32 | 3244.040352 | 2986.458353 | 0.6676 |
| LCK | chr1 | 32715449 | 33 | 32 | 1819.409485 | 1714.455821 | 0.66772 |
| E2F6 | chr2 | 11607489 | 19 | 18 | 1423.525919 | 1300.238095 | 0.66776 |
| F7 | chr13 | 113758676 | 21 | 23 | 1978.412698 | 2193.470696 | 0.66812 |
| F7 | chr13 | 113756835 | 24 | 27 | 1903.821734 | 2090.916583 | 0.66822 |
| APOA1 | chr11 | 116692258 | 34 | 32 | 3280.607788 | 3069.072382 | 0.6683 |
| APOA1 | chr11 | 116693354 | 32 | 32 | 2418.055695 | 2505.994959 | 0.66832 |
| NOS3 | chr7 | 150705956 | 22 | 28 | 791.0610801 | 1051.921313 | 0.66838 |
| ABCB1 | chr7 | 87343314 | 2 | 3 | 60 | 140 | 0.6684 |
| FOXP3 | chrX | 49122424 | 31 | 31 | 2913.465235 | 2890.341509 | 0.66854 |
| APOA1 | chr11 | 116699196 | 33 | 32 | 3103.146379 | 3034.866851 | 0.66864 |
| MIA3 | chr1 | 222814807 | 25 | 23 | 2136.993538 | 1998.843158 | 0.66872 |
| F7 | chr13 | 113759894 | 33 | 30 | 2900.346484 | 2662.801441 | 0.66878 |
| P2RY12 | chr3 | 151105633 | 36 | 33 | 2471.533312 | 2223.924487 | 0.6688 |
| KLF2 | chr19 | 16430385 | 34 | 33 | 1964.5072 | 1863.167527 | 0.66916 |
| CDKN2B-AS1 | chr9 | 21996550 | 38 | 33 | 2054.45132 | 1748.917821 | 0.6692 |
| CDKN2B-AS1 | chr9 | 21969251 | 10 | 12 | 292.2062733 | 390.4761905 | 0.66948 |
| APOA1 | chr11 | 116697570 | 32 | 32 | 2150.57451 | 2104.23115 | 0.66968 |
| TERT | chr5 | 1297491 | 34 | 32 | 2976.371412 | 2827.30862 | 0.6697 |
| APOA1 | chr11 | 116693247 | 35 | 32 | 3132.981962 | 2886.596754 | 0.6699 |
| BAX | chr19 | 49465131 | 32 | 32 | 2997.819942 | 2953.446296 | 0.66994 |
| LCK | chr1 | 32742001 | 28 | 26 | 2478.293651 | 2335.47619 | 0.6702 |

| | | | | | | | |
|---|---|---|---|---|---|---|---|
| P2RY12 | chr3 | 151111765 | 37 | 33 | 2275.222916 | 2001.870135 | 0.67042 |
| E2F6 | chr2 | 11607877 | 4 | 4 | 366.6666667 | 383.3333333 | 0.67042 |
| CDKN2B-AS1 | chr9 | 21966325 | 11 | 10 | 346.007326 | 353.3044733 | 0.67052 |
| ADRA2C | chr4 | 3762136 | 32 | 31 | 2669.126346 | 2625.545028 | 0.67062 |
| COL15A1 | chr9 | 101705250 | 32 | 32 | 3061.197912 | 3034.299844 | 0.67074 |
| CDKN2B-AS1 | chr9 | 21973120 | 12 | 18 | 337.3018648 | 539.3216922 | 0.67074 |
| F7 | chr13 | 113751351 | 20 | 18 | 1800.47619 | 1655.147908 | 0.67082 |
| ESR2 | chr14 | 64807531 | 35 | 33 | 3268.125793 | 3124.06748 | 0.6709 |
| FOXP3 | chrX | 49121228 | 17 | 16 | 1419.166667 | 1369.880952 | 0.6709 |
| E2F6 | chr2 | 11607397 | 30 | 31 | 2338.712312 | 2472.631965 | 0.67096 |
| TERT | chr5 | 1299501 | 26 | 30 | 2477.185037 | 2891.183261 | 0.671 |
| FOXP3 | chrX | 49118134 | 7 | 8 | 669.047619 | 743.3333333 | 0.67108 |
| ALDOA | chr16 | 30057850 | 34 | 32 | 3141.342085 | 2979.727683 | 0.67148 |
| TERT | chr5 | 1298238 | 33 | 32 | 3074.530874 | 2998.021704 | 0.67154 |
| PLA2G7 | chr6 | 46708903 | 36 | 32 | 3004.06429 | 2692.212527 | 0.67162 |
| LDLR | chr19 | 11217581 | 17 | 17 | 1445.242272 | 1485.912698 | 0.67182 |
| F7 | chr13 | 113750556 | 25 | 24 | 2350.865801 | 2218.498168 | 0.6722 |
| E2F6 | chr2 | 11609390 | 35 | 32 | 3100.392677 | 2867.706917 | 0.67236 |
| ESR1 | chr6 | 152005596 | 36 | 32 | 3445.295861 | 3050.034275 | 0.6724 |
| TERT | chr5 | 1295686 | 5 | 6 | 115 | 145.5555556 | 0.67244 |
| ALDOA | chr16 | 30078191 | 29 | 32 | 1399.593211 | 1595.4529 | 0.6726 |
| TIMP3 | chr22 | 33192858 | 22 | 27 | 703.1762693 | 900.6929431 | 0.67284 |
| FOXP3 | chrX | 49115103 | 3 | 4 | 285.7142857 | 369.047619 | 0.67294 |
| FOXP3 | chrX | 49114964 | 10 | 5 | 888.0952381 | 460.7142857 | 0.67308 |
| LDLR | chr19 | 11224283 | 31 | 31 | 2909.574592 | 2945.197303 | 0.67326 |
| ESR1 | chr6 | 152005882 | 10 | 5 | 855.4166667 | 445.7142857 | 0.67334 |
| APOA1 | chr11 | 116676850 | 37 | 32 | 3373.646752 | 2939.099179 | 0.67378 |
| CDKN2B-AS1 | chr9 | 21968506 | 3 | 3 | 80.71428571 | 73.57142857 | 0.67424 |
| ARHGEF18 | chr19 | 7469827 | 34 | 32 | 3321.590026 | 3106.78344 | 0.67428 |
| TIMP3 | chr22 | 33196112 | 33 | 32 | 3084.259358 | 2967.248172 | 0.6743 |
| F7 | chr13 | 113753102 | 34 | 32 | 3217.585973 | 3010.565093 | 0.6744 |
| APOA1 | chr11 | 116686129 | 8 | 9 | 747.1428571 | 857.1428571 | 0.67454 |
| F2RL3 | chr19 | 16992033 | 33 | 32 | 2993.582606 | 2877.40558 | 0.67464 |
| AKT2 | chr19 | 40796466 | 32 | 32 | 3124.234403 | 3145.085456 | 0.6748 |
| ESR2 | chr14 | 64809652 | 10 | 10 | 884.7619048 | 912.6984127 | 0.6749 |
| FOXP3 | chrX | 49118375 | 30 | 30 | 2883.75439 | 2864.397547 | 0.67518 |
| ADRA2C | chr4 | 3768501 | 6 | 5 | 175.5555556 | 174.040404 | 0.67518 |
| ABCA1 | chr9 | 107696635 | 5 | 11 | 491.6666667 | 1065.714286 | 0.67524 |
| CDKN1A | chr6 | 36654546 | 19 | 21 | 1844.252137 | 2014.444444 | 0.67528 |
| APOA1 | chr11 | 116676403 | 32 | 32 | 3055.235003 | 3070.615877 | 0.67544 |
| ATF3 | chr1 | 212739171 | 33 | 32 | 2726.510947 | 2610.489093 | 0.67556 |
| APOA1 | chr11 | 116714937 | 17 | 18 | 1598.214286 | 1728.055556 | 0.67556 |
| APOA1 | chr11 | 116674243 | 28 | 30 | 2500.862258 | 2709.717762 | 0.67564 |
| NOS3 | chr7 | 150690699 | 3 | 6 | 283.3333333 | 549.047619 | 0.6758 |
| LCK | chr1 | 32712917 | 30 | 31 | 2810.546259 | 2875.928179 | 0.67634 |
| NOS3 | chr7 | 150688767 | 34 | 33 | 2691.62707 | 2571.280229 | 0.6764 |
| APOA1 | chr11 | 116661183 | 32 | 32 | 3017.401441 | 3042.605671 | 0.67646 |

| | | | | | | | |
|---|---|---|---|---|---|---|---|
| LDLR | chr19 | 11220006 | 24 | 28 | 2274.754183 | 2604.572095 | 0.67654 |
| FGF2 | chr4 | 123742217 | 5 | 4 | 217.3015873 | 194.047619 | 0.67658 |
| ICAM2 | chr17 | 62086839 | 32 | 29 | 1423.606724 | 1336.198851 | 0.67756 |
| GALNT2 | chr1 | 230186065 | 37 | 32 | 3391.062288 | 2952.509603 | 0.678 |
| CDKN1A | chr6 | 36653382 | 25 | 30 | 1035.066667 | 1288.097752 | 0.6782 |
| COL15A1 | chr9 | 101697761 | 21 | 12 | 597.9210244 | 363.7485178 | 0.67832 |
| LDLR | chr19 | 11223936 | 23 | 21 | 2233.165266 | 2020.512266 | 0.67838 |
| ESR1 | chr6 | 152011723 | 2 | 7 | 73.01587301 | 302.8571429 | 0.67848 |
| E2F6 | chr2 | 11613480 | 34 | 32 | 3149.717527 | 2983.19869 | 0.67878 |
| NR3C1 | chr5 | 142788801 | 36 | 32 | 3302.422338 | 2954.333503 | 0.6793 |
| AKT2 | chr19 | 40800006 | 10 | 15 | 889.781746 | 1365.436508 | 0.67934 |
| MIA3 | chr1 | 222809447 | 2 | 3 | 61.9047619 | 140 | 0.67946 |
| LDLR | chr19 | 11196094 | 33 | 29 | 3109.426995 | 2761.125042 | 0.67948 |
| F2RL3 | chr19 | 16990559 | 31 | 30 | 2877.283117 | 2817.07709 | 0.67962 |
| PECAM1 | chr17 | 62408868 | 13 | 14 | 574.3506494 | 667.5793651 | 0.67964 |
| AMOTL2 | chr3 | 134095414 | 34 | 32 | 3055.126646 | 2830.644509 | 0.67968 |
| LOXL1 | chr15 | 74218200 | 31 | 32 | 2853.000022 | 2972.743437 | 0.67986 |
| CDKN2B-AS1 | chr9 | 21964756 | 12 | 14 | 443.8095238 | 559.8809524 | 0.68028 |
| LPA | chr6 | 11223209 | 31 | 31 | 2353.616087 | 2403.36118 | 0.68036 |
| APOA1 | chr11 | 116670183 | 18 | 21 | 806.885323 | 996.0452912 | 0.68044 |
| BCL2 | chr18 | 60988430 | 12 | 9 | 363.330927 | 302.2114442 | 0.68076 |
| CDKN1A | chr6 | 36654687 | 32 | 32 | 3067.404141 | 3035.984788 | 0.68116 |
| NOS3 | chr7 | 150690119 | 13 | 14 | 947.5 | 1060.15873 | 0.68126 |
| ALDOA | chr16 | 30060886 | 32 | 32 | 3133.131313 | 3147.899323 | 0.68142 |
| KLF2 | chr19 | 16432759 | 34 | 32 | 3001.493928 | 2852.813377 | 0.6815 |
| CDKN2B-AS1 | chr9 | 21967396 | 31 | 32 | 1449.111822 | 1449.935258 | 0.68168 |
| APOA1 | chr11 | 116661130 | 34 | 33 | 2864.053584 | 2733.092297 | 0.68184 |
| P2RY12 | chr3 | 151108714 | 38 | 32 | 3255.362627 | 2771.158422 | 0.6823 |
| FOXP3 | chrX | 49116462 | 2 | 4 | 53.57142857 | 160 | 0.68254 |
| NOS3 | chr7 | 150690506 | 32 | 32 | 2956.08853 | 2979.504365 | 0.68264 |
| APOA1 | chr11 | 116691984 | 31 | 32 | 2936.058484 | 3053.920401 | 0.6829 |
| F7 | chr13 | 113758700 | 26 | 28 | 2247.307342 | 2464.020147 | 0.683 |
| NOS3 | chr7 | 150682507 | 13 | 11 | 1183.333333 | 1032.222222 | 0.68316 |
| LPA | chr6 | 160944151 | 11 | 14 | 1008.888889 | 1252.5 | 0.68322 |
| CDKN2B-AS1 | chr9 | 21997429 | 31 | 30 | 2234.34723 | 2097.167555 | 0.68334 |
| P2RY12 | chr3 | 151106019 | 3 | 6 | 92.58974359 | 229.6405229 | 0.68338 |
| LOXL1 | chr15 | 74219043 | 19 | 16 | 633.0037953 | 501.5611884 | 0.68342 |
| APOA1 | chr11 | 116694934 | 32 | 32 | 3004.640386 | 2972.726656 | 0.6838 |
| ADRA2C | chr4 | 3768558 | 13 | 8 | 350.7239252 | 246.5079365 | 0.68386 |
| ATF3 | chr1 | 212738678 | 32 | 31 | 3035.58931 | 2914.590941 | 0.6846 |
| AKT2 | chr19 | 40797817 | 38 | 32 | 3516.068849 | 2946.039798 | 0.68466 |
| ESR1 | chr6 | 152002406 | 33 | 32 | 2402.49821 | 2294.727944 | 0.68526 |
| F7 | chr13 | 113752892 | 32 | 33 | 2825.62543 | 2875.016284 | 0.68528 |
| FOXP3 | chrX | 49127361 | 17 | 15 | 1429.456654 | 1290.714286 | 0.68558 |
| ADRA2C | chr4 | 3761145 | 26 | 30 | 2094.275827 | 2455.02164 | 0.68562 |
| FOXP3 | chrX | 49127367 | 17 | 16 | 1497.489177 | 1438.524531 | 0.68602 |
| KLF2 | chr19 | 16429013 | 3 | 7 | 283.3333333 | 640 | 0.68632 |

| | | | | | | | |
|---|---|---|---|---|---|---|---|
| MCPH1 | chr8 | 6470875 | 30 | 30 | 2813.268706 | 2835.868055 | 0.68644 |
| LDLR | chr19 | 11220868 | 7 | 9 | 680 | 885.7142857 | 0.68674 |
| FOXP3 | chrX | 49127168 | 19 | 24 | 1813.290043 | 2263.131313 | 0.68684 |
| FOXP3 | chrX | 49129969 | 19 | 26 | 644.9327179 | 842.2338133 | 0.68706 |
| F7 | chr13 | 113756891 | 30 | 32 | 2703.9005 | 2920.621184 | 0.6871 |
| ADRA2C | chr4 | 3760448 | 37 | 32 | 3508.183051 | 3053.075451 | 0.6877 |
| CDKN2B-AS1 | chr9 | 21988978 | 29 | 25 | 845.9294565 | 756.569053 | 0.68806 |
| FGF2 | chr4 | 123740011 | 36 | 32 | 3452.621595 | 3055.250115 | 0.6881 |
| E2F6 | chr2 | 11607896 | 22 | 17 | 1369.972527 | 1005.353535 | 0.68842 |
| CDKN2B-AS1 | chr9 | 21984021 | 34 | 33 | 2073.797378 | 2048.974394 | 0.68862 |
| NOS3 | chr7 | 150706355 | 31 | 30 | 2586.318002 | 2545.66838 | 0.68868 |
| PLA2G7 | chr6 | 46703863 | 31 | 33 | 1700.793161 | 1759.993751 | 0.68896 |
| GCK | chr7 | 44232534 | 37 | 32 | 3487.046893 | 3028.270056 | 0.68924 |
| CDKN2B-AS1 | chr9 | 21993928 | 2 | 4 | 52.99145299 | 153.75 | 0.68944 |
| GCK | chr7 | 44237190 | 35 | 32 | 2918.998432 | 2644.773534 | 0.6903 |
| FOXP3 | chrX | 49117253 | 15 | 16 | 1463.333333 | 1540 | 0.69032 |
| APOA1 | chr11 | 116695078 | 4 | 5 | 308.3333333 | 416.6666667 | 0.6904 |
| ALOX15 | chr17 | 4553111 | 35 | 33 | 2578.790855 | 2395.581814 | 0.69048 |
| NOS3 | chr7 | 150710426 | 25 | 28 | 1136.995828 | 1322.273976 | 0.69068 |
| APOA1 | chr11 | 116679711 | 35 | 31 | 2869.215251 | 2462.956865 | 0.6908 |
| CDKN2B-AS1 | chr9 | 21966783 | 35 | 33 | 2013.768957 | 1859.958488 | 0.69094 |
| APOA1 | chr11 | 116662969 | 33 | 32 | 2903.522804 | 2851.295259 | 0.69094 |
| AMOTL2 | chr3 | 134102265 | 32 | 31 | 2745.48654 | 2622.561376 | 0.69098 |
| MIA3 | chr1 | 222815171 | 31 | 32 | 2987.378442 | 3113.650855 | 0.69126 |
| ADRA2C | chr4 | 3761166 | 31 | 32 | 2619.564466 | 2752.779205 | 0.69144 |
| ATF3 | chr1 | 212742646 | 36 | 33 | 2872.550747 | 2667.459401 | 0.6921 |
| CTGF | chr6 | 132274264 | 32 | 32 | 2793.479776 | 2726.201262 | 0.69228 |
| ATF3 | chr1 | 212746991 | 17 | 10 | 1656.547619 | 983.3333333 | 0.69234 |
| F7 | chr13 | 113751439 | 30 | 32 | 2691.322011 | 2900.778491 | 0.69236 |
| APOA1 | chr11 | 116694373 | 29 | 30 | 2758.201433 | 2827.2114 | 0.6924 |
| GALNT2 | chr1 | 230189809 | 15 | 22 | 1463.095238 | 2160.31746 | 0.6926 |
| ALOX15 | chr17 | 4545390 | 29 | 27 | 1914.488983 | 1726.687055 | 0.6927 |
| LDLR | chr19 | 11221435 | 34 | 32 | 3260.177471 | 3083.878729 | 0.69292 |
| FGF2 | chr4 | 123747552 | 22 | 18 | 732.4483722 | 631.8491476 | 0.6937 |
| APOA1 | chr11 | 116707384 | 17 | 18 | 1536.825397 | 1588.885281 | 0.69372 |
| NOS3 | chr7 | 150708968 | 29 | 28 | 2797.813645 | 2716.245686 | 0.69376 |
| KLF2 | chr19 | 16438136 | 33 | 32 | 2022.715326 | 2009.71987 | 0.69402 |
| ADRA2C | chr4 | 3758685 | 34 | 33 | 1834.43206 | 1743.704774 | 0.69404 |
| MCPH1 | chr8 | 6470218 | 17 | 16 | 1673.253968 | 1543.492063 | 0.69404 |
| APOA1 | chr11 | 116662644 | 5 | 2 | 408.8095238 | 175 | 0.6943 |
| F7 | chr13 | 113775150 | 24 | 28 | 2200 | 2524.756077 | 0.69448 |
| TIMP3 | chr22 | 33193908 | 34 | 32 | 2886.830735 | 2678.563786 | 0.69476 |
| APOA1 | chr11 | 116702346 | 2 | 9 | 80 | 388.0298786 | 0.69478 |
| APOA1 | chr11 | 116701456 | 34 | 31 | 3104.420273 | 2793.599859 | 0.69492 |
| LPA | chr6 | 160951780 | 12 | 14 | 996.6269841 | 1129.87013 | 0.69496 |
| ALDOA | chr16 | 30061016 | 19 | 20 | 1709.206349 | 1827.02381 | 0.695 |
| CDKN2B-AS1 | chr9 | 22010340 | 35 | 32 | 3272.790979 | 3018.394405 | 0.69512 |

| | | | | | | | |
|---|---|---|---|---|---|---|---|
| F7 | chr13 | 113759544 | 29 | 28 | 2421.834879 | 2376.552614 | 0.69516 |
| ICAM2 | chr17 | 62085533 | 6 | 9 | 447.0238095 | 633.6904762 | 0.69518 |
| ARHGEF18 | chr19 | 7537624 | 7 | 4 | 680 | 380 | 0.69528 |
| APOA1 | chr11 | 116693863 | 32 | 32 | 2990.303306 | 3010.062436 | 0.69548 |
| ALDOA | chr16 | 30061977 | 33 | 33 | 2570.567857 | 2607.050024 | 0.69566 |
| ABCA1 | chr9 | 107696497 | 36 | 31 | 3404.303221 | 2901.88918 | 0.6958 |
| LDLR | chr19 | 11196394 | 25 | 24 | 2241.733087 | 2178.472222 | 0.69586 |
| FOXP3 | chrX | 49117333 | 5 | 5 | 460 | 446.6666667 | 0.69602 |
| CDKN2B-AS1 | chr9 | 22006229 | 7 | 10 | 221.017316 | 346.025605 | 0.69612 |
| LDLR | chr19 | 11194314 | 31 | 31 | 2972.74087 | 2995.597042 | 0.69624 |
| F7 | chr13 | 113760286 | 22 | 26 | 1237.872544 | 1412.691589 | 0.69636 |
| LDLR | chr19 | 11216129 | 28 | 31 | 2655.041723 | 2965.989844 | 0.69638 |
| PHACTR1 | chr6 | 12711173 | 35 | 32 | 2905.501546 | 2679.752446 | 0.6964 |
| GALNT2 | chr1 | 230193196 | 32 | 32 | 2688.167654 | 2656.777146 | 0.6966 |
| APOA1 | chr11 | 116712443 | 31 | 26 | 2902.258297 | 2395.936476 | 0.69684 |
| NOS3 | chr7 | 150681190 | 3 | 7 | 83.47338935 | 241.5257516 | 0.6971 |
| APOA1 | chr11 | 116700488 | 31 | 31 | 2782.195998 | 2821.182397 | 0.69714 |
| PLA2G7 | chr6 | 46707727 | 24 | 31 | 2046.415545 | 2685.736902 | 0.6978 |
| ARHGEF18 | chr19 | 7537482 | 33 | 32 | 3200.280887 | 3084.916922 | 0.69784 |
| APOA1 | chr11 | 116720419 | 37 | 33 | 1946.526692 | 1708.862017 | 0.69786 |
| AMOTL2 | chr3 | 134102032 | 4 | 6 | 383.3333333 | 585.7142857 | 0.69804 |
| ESR1 | chr6 | 152007558 | 27 | 33 | 2481.305557 | 2986.005917 | 0.69838 |
| CDKN2B-AS1 | chr9 | 21967154 | 27 | 30 | 981.8192232 | 1121.477663 | 0.69846 |
| P2RY12 | chr3 | 151111309 | 16 | 11 | 1147.361111 | 822.2619048 | 0.69854 |
| AMOTL2 | chr3 | 134094559 | 32 | 32 | 2732.586063 | 2767.491184 | 0.69882 |
| ARHGEF18 | chr19 | 7467586 | 36 | 32 | 3238.996901 | 2898.571746 | 0.69886 |
| PLA2G7 | chr6 | 46708316 | 39 | 33 | 2773.291493 | 2377.516533 | 0.69936 |
| NOS3 | chr7 | 150706087 | 29 | 31 | 1122.691192 | 1242.375687 | 0.69938 |
| APOA1 | chr11 | 116664915 | 38 | 32 | 3580.75251 | 3032.025713 | 0.6995 |
| LOXL1 | chr15 | 74217221 | 33 | 32 | 2949.651284 | 2837.56563 | 0.69998 |
| ATF3 | chr1 | 212741799 | 33 | 32 | 2930.803577 | 2869.813718 | 0.70028 |
| F7 | chr13 | 113758242 | 23 | 24 | 2031.557332 | 2087.302697 | 0.70058 |
| CDKN2B-AS1 | chr9 | 21996836 | 8 | 7 | 384.2207792 | 311.6666667 | 0.70066 |
| AKT2 | chr19 | 40794783 | 33 | 33 | 2961.746056 | 2925.594708 | 0.70096 |
| ADRA2C | chr4 | 3766501 | 31 | 31 | 2731.510483 | 2765.343976 | 0.70108 |
| E2F6 | chr2 | 11607206 | 34 | 33 | 2857.686418 | 2733.834445 | 0.70122 |
| TFPI | chr2 | 188343362 | 34 | 33 | 2902.435129 | 2853.137522 | 0.70148 |
| ESR1 | chr6 | 152002832 | 13 | 11 | 345.9027504 | 321.8664084 | 0.7015 |
| DHFR | chr5 | 79956188 | 3 | 4 | 120 | 186.6666667 | 0.70154 |
| NOS3 | chr7 | 150710552 | 9 | 6 | 311.6277874 | 231.8841126 | 0.70158 |
| ADRA2C | chr4 | 3758771 | 35 | 32 | 3162.263499 | 2907.990513 | 0.70178 |
| AMOTL2 | chr3 | 134074282 | 2 | 2 | 73.33333333 | 100 | 0.70186 |
| APOA1 | chr11 | 116684808 | 27 | 27 | 1122.628446 | 1087.897889 | 0.70198 |
| TERT | chr5 | 1300895 | 17 | 14 | 1532.420635 | 1227.619048 | 0.70206 |
| GCK | chr7 | 44183531 | 35 | 32 | 2901.746267 | 2677.492775 | 0.70242 |
| ADRA2C | chr4 | 3759667 | 32 | 32 | 2879.243204 | 2903.950094 | 0.70242 |
| MCPH1 | chr8 | 6263126 | 37 | 33 | 2576.548103 | 2334.34983 | 0.70248 |

| | | | | | | | |
|---|---|---|---|---|---|---|---|
| APOA1 | chr11 | 116685391 | 30 | 31 | 2266.299987 | 2278.062198 | 0.70284 |
| LCK | chr1 | 32708276 | 16 | 19 | 508.3835073 | 636.4208468 | 0.70284 |
| LOXL1 | chr15 | 74215085 | 19 | 16 | 551.140802 | 488.1108847 | 0.70296 |
| APOA1 | chr11 | 116671918 | 9 | 9 | 812.6984127 | 786.1904762 | 0.7031 |
| CDKN1A | chr6 | 36652067 | 28 | 30 | 2612.272409 | 2834.292986 | 0.70312 |
| APOA1 | chr11 | 116711050 | 37 | 32 | 3434.3071 | 2952.851913 | 0.70318 |
| NOS3 | chr7 | 150690676 | 11 | 14 | 1025.670996 | 1280 | 0.70326 |
| F2RL3 | chr19 | 16995086 | 19 | 21 | 1655 | 1764.529915 | 0.70362 |
| APOA1 | chr11 | 116693416 | 36 | 32 | 2890.196359 | 2533.780405 | 0.704 |
| ABCB1 | chr7 | 87349785 | 30 | 28 | 2761.669638 | 2546.858256 | 0.70408 |
| AKT2 | chr19 | 40794277 | 36 | 32 | 3441.30274 | 3045.614795 | 0.70438 |
| ADRA2C | chr4 | 3765937 | 28 | 27 | 2534.182762 | 2466.471225 | 0.70438 |
| F7 | chr13 | 113750939 | 30 | 28 | 2812.64468 | 2596.298701 | 0.7044 |
| FGF2 | chr4 | 123742007 | 28 | 31 | 2777.391083 | 3054.998196 | 0.70442 |
| APOA1 | chr11 | 116719926 | 34 | 32 | 3216.208649 | 3007.45425 | 0.70444 |
| ADRA2C | chr4 | 3766151 | 33 | 33 | 2866.032462 | 2826.265696 | 0.70448 |
| LPA | chr6 | 160949284 | 32 | 33 | 2233.507749 | 2338.499544 | 0.70458 |
| ADRA2C | chr4 | 3760692 | 30 | 31 | 2460.395269 | 2506.396983 | 0.70458 |
| APOA1 | chr11 | 116694752 | 38 | 32 | 3697.785695 | 3086.279582 | 0.70468 |
| TERT | chr5 | 1295919 | 29 | 32 | 1470.112662 | 1682.25646 | 0.70478 |
| APOA1 | chr11 | 116661569 | 29 | 30 | 2794.465312 | 2912.541358 | 0.7054 |
| CDKN2B-AS1 | chr9 | 22008441 | 13 | 12 | 351.3057428 | 352.1640225 | 0.70568 |
| GCK | chr7 | 44237616 | 33 | 32 | 3118.856133 | 3042.468502 | 0.70572 |
| DHFR | chr5 | 79954174 | 27 | 24 | 2200.200216 | 1918.956044 | 0.7059 |
| LCK | chr1 | 32741797 | 11 | 7 | 917.6984127 | 600.4761905 | 0.7062 |
| FOXP3 | chrX | 49127044 | 19 | 15 | 1790.833333 | 1392.142857 | 0.7064 |
| ALDOA | chr16 | 30061433 | 31 | 31 | 2815.85252 | 2839.322927 | 0.70646 |
| LOXL1 | chr15 | 74216030 | 33 | 33 | 1823.744357 | 1859.905327 | 0.7065 |
| APOA1 | chr11 | 116699524 | 37 | 33 | 3020.883716 | 2727.968601 | 0.70656 |
| APOA1 | chr11 | 116694328 | 21 | 20 | 1540.608395 | 1435.903541 | 0.70748 |
| LDLR | chr19 | 11215999 | 31 | 31 | 2869.860062 | 2889.357505 | 0.70762 |
| APOA1 | chr11 | 116721459 | 33 | 32 | 3258.860418 | 3149.431076 | 0.70768 |
| LPA | chr6 | 11217074 | 8 | 9 | 338.4623823 | 414.1268664 | 0.70784 |
| FOXP3 | chrX | 49127189 | 18 | 26 | 1680.40293 | 2462.532468 | 0.7079 |
| APOA1 | chr11 | 116684029 | 29 | 28 | 2029.922951 | 1898.298231 | 0.70798 |
| TIMP3 | chr22 | 33187010 | 38 | 32 | 3222.819624 | 2733.094329 | 0.70848 |
| ADRA2C | chr4 | 3765038 | 10 | 7 | 274.8697054 | 184.1553545 | 0.7088 |
| GCK | chr7 | 44229210 | 33 | 32 | 2697.763325 | 2648.950308 | 0.70892 |
| PHACTR1 | chr6 | 12712109 | 35 | 32 | 3315.976076 | 3049.368085 | 0.70926 |
| P2RY12 | chr3 | 151112568 | 26 | 25 | 1819.842677 | 1822.629176 | 0.70968 |
| ALOX15 | chr17 | 4545356 | 7 | 10 | 260.6926407 | 401.1904762 | 0.70974 |
| FOXP3 | chrX | 49115240 | 23 | 26 | 1956.016484 | 2250.025253 | 0.7098 |
| TERT | chr5 | 1295509 | 5 | 10 | 168.5714286 | 375.4761905 | 0.7099 |
| LDLR | chr19 | 11222534 | 32 | 32 | 3040.010871 | 3061.959089 | 0.70998 |
| F7 | chr13 | 113751365 | 25 | 21 | 2373.409091 | 1968.527029 | 0.71016 |
| FOXP3 | chrX | 49119938 | 33 | 32 | 3064.886215 | 2951.220475 | 0.7105 |
| ALDOA | chr16 | 30061013 | 18 | 21 | 1715.108225 | 2019.444444 | 0.71092 |

| | | | | | | | |
|---|---|---|---|---|---|---|---|
| AMOTL2 | chr3 | 134099217 | 40 | 32 | 1695.338957 | 1389.693955 | 0.71136 |
| F2RL3 | chr19 | 16999145 | 22 | 27 | 1921.882936 | 2314.559885 | 0.71158 |
| PECAM1 | chr17 | 62401737 | 24 | 29 | 2035.769249 | 2417.782773 | 0.71246 |
| CDKN2B-AS1 | chr9 | 21968838 | 38 | 32 | 1415.227078 | 1166.66891 | 0.71254 |
| LOXL1 | chr15 | 74212327 | 33 | 32 | 3159.96404 | 3079.021494 | 0.7126 |
| CDKN2B-AS1 | chr9 | 21967384 | 6 | 4 | 196.127451 | 163.0769231 | 0.71282 |
| LPA | chr6 | 160948436 | 36 | 32 | 2973.854484 | 2618.752847 | 0.71306 |
| LDLR | chr19 | 11216013 | 32 | 31 | 3038.860682 | 2961.399573 | 0.71328 |
| KLF2 | chr19 | 16437865 | 24 | 28 | 1116.958086 | 1353.63272 | 0.71328 |
| APOA1 | chr11 | 116689954 | 33 | 32 | 3092.612408 | 3025.495146 | 0.71332 |
| ABCA1 | chr9 | 107698086 | 18 | 22 | 1626.150794 | 2020.541505 | 0.7135 |
| F7 | chr13 | 113758450 | 32 | 32 | 2342.604754 | 2377.771974 | 0.71352 |
| BAX | chr19 | 49449020 | 34 | 32 | 3245.059101 | 3039.641209 | 0.71394 |
| ALOX15 | chr17 | 4545386 | 26 | 24 | 1415.970696 | 1348.326118 | 0.71398 |
| APOA1 | chr11 | 116684563 | 30 | 31 | 2213.033263 | 2210.598617 | 0.71424 |
| APOA1 | chr11 | 116720179 | 21 | 29 | 1636.370851 | 2205.555556 | 0.71426 |
| NOS3 | chr7 | 150710350 | 31 | 31 | 1839.958996 | 1791.431221 | 0.71444 |
| NOS3 | chr7 | 150683100 | 24 | 21 | 2253.006895 | 1949.402046 | 0.71446 |
| GCK | chr7 | 44237431 | 35 | 32 | 2977.697166 | 2749.440967 | 0.71476 |
| TIMP3 | chr22 | 33195517 | 37 | 32 | 3380.759122 | 2940.992221 | 0.71478 |
| FOXP3 | chrX | 49120284 | 24 | 26 | 1850.104618 | 2050.847763 | 0.7149 |
| APOA1 | chr11 | 116705503 | 25 | 25 | 2251.984127 | 2283.220391 | 0.71504 |
| ATF3 | chr1 | 212739150 | 34 | 32 | 2902.647217 | 2696.244377 | 0.71522 |
| MCPH1 | chr8 | 6471144 | 6 | 5 | 533.2142857 | 460 | 0.71522 |
| LPA | chr6 | 11216878 | 39 | 32 | 3746.634743 | 3083.87029 | 0.71562 |
| HMGCR | chr5 | 74623854 | 5 | 10 | 392.3809524 | 822.8571429 | 0.7157 |
| CDKN2B-AS1 | chr9 | 22006036 | 17 | 19 | 533.0184766 | 626.4316239 | 0.71572 |
| PECAM1 | chr17 | 62408480 | 24 | 24 | 1021.648235 | 1063.115912 | 0.71598 |
| TERT | chr5 | 1298259 | 33 | 32 | 3135.213399 | 3022.340131 | 0.71624 |
| APOA1 | chr11 | 116717118 | 35 | 32 | 3296.787805 | 3029.35548 | 0.71626 |
| NOS3 | chr7 | 150711159 | 16 | 12 | 490.1561914 | 397.8282828 | 0.7167 |
| COL15A1 | chr9 | 101699603 | 32 | 32 | 2697.256447 | 2728.0585 | 0.71674 |
| F7 | chr13 | 113752476 | 33 | 32 | 2844.705264 | 2726.108651 | 0.71686 |
| APOA1 | chr11 | 116703493 | 35 | 32 | 3117.940022 | 2874.756913 | 0.71694 |
| NOS3 | chr7 | 150683937 | 34 | 33 | 2106.674901 | 2005.466163 | 0.7171 |
| APOA1 | chr11 | 116724126 | 28 | 28 | 1172.797272 | 1206.615283 | 0.71712 |
| APOA1 | chr11 | 116723716 | 27 | 29 | 1195.043387 | 1325.132913 | 0.7173 |
| CDKN2B-AS1 | chr9 | 21969342 | 24 | 24 | 877.4092926 | 840.9848808 | 0.71752 |
| ABCB1 | chr7 | 87351323 | 34 | 32 | 3268.920817 | 3066.116379 | 0.71766 |
| PLA2G7 | chr6 | 46707055 | 34 | 32 | 2757.326576 | 2573.881705 | 0.71804 |
| BAX | chr19 | 49465099 | 32 | 33 | 2467.823877 | 2586.475633 | 0.71804 |
| F2RL3 | chr19 | 16991644 | 13 | 23 | 1150.972222 | 2083.170996 | 0.71822 |
| MCPH1 | chr8 | 6260014 | 7 | 5 | 649.047619 | 451.4285714 | 0.71824 |
| APOA1 | chr11 | 116690566 | 34 | 33 | 1801.873054 | 1785.464622 | 0.7183 |
| F7 | chr13 | 113752655 | 33 | 30 | 3163.688862 | 2856.701863 | 0.71918 |
| ATF3 | chr1 | 212742900 | 27 | 30 | 2437.018398 | 2673.669299 | 0.71918 |
| GCK | chr7 | 44232918 | 32 | 32 | 2771.108278 | 2714.855877 | 0.71942 |

| | | | | | | | |
|---|---|---|---|---|---|---|---|
| MCPH1 | chr8 | 6470336 | 25 | 27 | 2223.708791 | 2329.053724 | 0.71952 |
| CDKN2B-AS1 | chr9 | 21993387 | 38 | 33 | 2444.295787 | 2100.313744 | 0.71974 |
| DHFR | chr5 | 79958367 | 24 | 22 | 2065.972222 | 1937.650369 | 0.71976 |
| F7 | chr13 | 113751042 | 21 | 17 | 2066.903652 | 1647.619048 | 0.71988 |
| ESR1 | chr6 | 152003403 | 34 | 33 | 1766.992259 | 1752.704555 | 0.72008 |
| APOA1 | chr11 | 116692645 | 33 | 32 | 3048.735493 | 2933.885755 | 0.72078 |
| PECAM1 | chr17 | 62409682 | 30 | 27 | 1222.83795 | 1067.775547 | 0.72086 |
| ALDOA | chr16 | 30078186 | 30 | 30 | 1376.58743 | 1338.611932 | 0.7211 |
| ADRA2C | chr4 | 3762528 | 20 | 26 | 1186.975524 | 1618.7389 | 0.72114 |
| ABCG1 | chr21 | 43629407 | 33 | 32 | 3025.955978 | 2914.622019 | 0.72144 |
| F2RL3 | chr19 | 17003189 | 21 | 26 | 1845.28416 | 2250.879121 | 0.72188 |
| COL15A1 | chr9 | 101696963 | 18 | 22 | 1704.047619 | 2104.603175 | 0.72204 |
| CDKN2B-AS1 | chr9 | 21979603 | 6 | 2 | 174.1666667 | 53.33333333 | 0.72236 |
| ATF3 | chr1 | 212742073 | 32 | 32 | 2936.103852 | 2915.07767 | 0.72258 |
| F2RL3 | chr19 | 16991041 | 10 | 7 | 887.5 | 643.3333333 | 0.72292 |
| LDLR | chr19 | 11195861 | 5 | 6 | 370.9090909 | 467.5 | 0.72292 |
| TIMP3 | chr22 | 33192057 | 32 | 32 | 1217.700335 | 1256.10062 | 0.72344 |
| ESR1 | chr6 | 152002028 | 27 | 28 | 2565.899541 | 2636.933304 | 0.72368 |
| GCK | chr7 | 44228544 | 25 | 24 | 1893.791486 | 1769.581807 | 0.72374 |
| ALOX15 | chr17 | 4553297 | 16 | 17 | 926.1219336 | 1028.333333 | 0.72374 |
| F7 | chr13 | 113758234 | 25 | 21 | 2105.255717 | 1738.310856 | 0.72406 |
| ADRA2C | chr4 | 3761457 | 29 | 29 | 2451.022311 | 2486.720143 | 0.7241 |
| PECAM1 | chr17 | 62409738 | 34 | 32 | 2894.369639 | 2749.10907 | 0.72448 |
| APOA1 | chr11 | 116680419 | 24 | 28 | 1385.101704 | 1554.630732 | 0.7246 |
| NOS3 | chr7 | 150693983 | 4 | 9 | 117.294686 | 307.2222222 | 0.7247 |
| FGF2 | chr4 | 123747542 | 23 | 27 | 870.4598854 | 992.2682486 | 0.72496 |
| NOS3 | chr7 | 150710242 | 27 | 28 | 1134.449374 | 1211.718253 | 0.72524 |
| APOA1 | chr11 | 116699165 | 33 | 32 | 3166.117762 | 3028.297235 | 0.72528 |
| APOA1 | chr11 | 116700557 | 27 | 30 | 2270.627167 | 2475.664336 | 0.72548 |
| F7 | chr13 | 113751543 | 30 | 28 | 2581.318959 | 2365.126956 | 0.72564 |
| APOA1 | chr11 | 116678013 | 21 | 21 | 1961.784604 | 1982.41342 | 0.72572 |
| LOXL1 | chr15 | 74218700 | 18 | 20 | 834.526862 | 966.3492063 | 0.72608 |
| F7 | chr13 | 113751492 | 28 | 29 | 2591.682674 | 2708.151154 | 0.7262 |
| NOS3 | chr7 | 150684128 | 2 | 3 | 183.3333333 | 263.3333333 | 0.72634 |
| ADRA2C | chr4 | 3758391 | 32 | 32 | 2957.870174 | 2936.898487 | 0.72668 |
| NOS3 | chr7 | 150706020 | 32 | 33 | 1682.953607 | 1775.281988 | 0.72672 |
| TERT | chr5 | 1295855 | 32 | 33 | 2455.67601 | 2501.392779 | 0.72684 |
| ABCG1 | chr21 | 43634560 | 25 | 28 | 2432.391457 | 2747.380952 | 0.7269 |
| ADRA2C | chr4 | 3763068 | 5 | 6 | 436.6233766 | 500 | 0.72714 |
| AMOTL2 | chr3 | 134101185 | 33 | 32 | 2876.002408 | 2811.299319 | 0.72738 |
| APOA1 | chr11 | 116725573 | 31 | 30 | 3039.386736 | 2926.145857 | 0.72768 |
| GCK | chr7 | 44233215 | 36 | 32 | 3325.355176 | 2941.172186 | 0.72774 |
| TERT | chr5 | 1299822 | 31 | 30 | 2873.696291 | 2758.477217 | 0.72784 |
| NOS3 | chr7 | 150710413 | 24 | 23 | 1020.559516 | 1011.728737 | 0.72788 |
| PECAM1 | chr17 | 62401694 | 26 | 27 | 2429.251027 | 2500.276668 | 0.7283 |
| CDKN2B-AS1 | chr9 | 21968147 | 15 | 16 | 495.1848141 | 496.3748713 | 0.72832 |
| CDKN2B-AS1 | chr9 | 22009720 | 12 | 11 | 321.4734777 | 317.3511648 | 0.72858 |

| | | | | | | | |
|---|---|---|---|---|---|---|---|
| F7 | chr13 | 113758134 | 31 | 32 | 2035.053149 | 2152.735043 | 0.72864 |
| ICAM2 | chr17 | 62083948 | 25 | 26 | 1396.336164 | 1412.109557 | 0.7291 |
| DHFR | chr5 | 79957261 | 35 | 32 | 3328.486708 | 3056.1894 | 0.7292 |
| ADRA2C | chr4 | 3768638 | 18 | 19 | 530.3115725 | 539.0344246 | 0.7297 |
| ADRA2C | chr4 | 3762850 | 35 | 33 | 2559.776998 | 2381.820384 | 0.72994 |
| COL15A1 | chr9 | 101702290 | 34 | 33 | 2201.198849 | 2170.329208 | 0.73002 |
| AMOTL2 | chr3 | 134095392 | 33 | 32 | 3136.062953 | 3028.177789 | 0.73016 |
| CTGF | chr6 | 132280561 | 32 | 31 | 2944.271092 | 2871.309696 | 0.73016 |
| TFPI | chr2 | 188419717 | 20 | 20 | 538.5731303 | 561.2830227 | 0.73016 |
| ESR1 | chr6 | 152002859 | 20 | 15 | 673.8698179 | 534.9476557 | 0.73042 |
| P2RY12 | chr3 | 151104035 | 34 | 32 | 3050.318039 | 2891.708473 | 0.7305 |
| NR3C1 | chr5 | 142657622 | 7 | 8 | 353.0515732 | 432.8306878 | 0.73072 |
| APOA1 | chr11 | 116716985 | 33 | 32 | 3209.866915 | 3096.822726 | 0.73078 |
| AKT2 | chr19 | 40798540 | 32 | 32 | 3048.170711 | 3033.590758 | 0.7309 |
| TERT | chr5 | 1297628 | 31 | 31 | 2723.408157 | 2699.996854 | 0.73098 |
| PECAM1 | chr17 | 62409236 | 31 | 24 | 1547.100896 | 1238.697104 | 0.73108 |
| APOA1 | chr11 | 116709736 | 32 | 32 | 2937.968433 | 2900.731105 | 0.73124 |
| APOA1 | chr11 | 116699397 | 23 | 22 | 837.1389787 | 769.9810643 | 0.73128 |
| LCK | chr1 | 32707266 | 20 | 17 | 653.2880739 | 529.14798 | 0.7315 |
| F7 | chr13 | 113753041 | 33 | 32 | 3173.567615 | 3061.73874 | 0.73156 |
| FGF2 | chr4 | 123741097 | 37 | 32 | 3335.642804 | 2899.873414 | 0.73158 |
| LPA | chr6 | 11218291 | 38 | 32 | 3679.336319 | 3106.394161 | 0.73164 |
| APOA1 | chr11 | 116716810 | 36 | 32 | 3487.647904 | 3109.153266 | 0.73182 |
| GALNT2 | chr1 | 230193668 | 26 | 27 | 1468.838764 | 1569.63543 | 0.73188 |
| APOA1 | chr11 | 116691128 | 33 | 32 | 3060.838502 | 2983.500707 | 0.73192 |
| CDKN1A | chr6 | 36654802 | 33 | 32 | 2671.882146 | 2624.055412 | 0.73206 |
| MIA3 | chr1 | 222811525 | 28 | 24 | 2519.425574 | 2187.525632 | 0.73228 |
| F7 | chr13 | 113760101 | 20 | 16 | 1599.868604 | 1248.441558 | 0.7323 |
| APOA1 | chr11 | 116674664 | 29 | 32 | 2832.804217 | 3098.412698 | 0.7328 |
| APOA1 | chr11 | 116692176 | 33 | 32 | 3121.7719 | 3014.81104 | 0.73302 |
| ESR1 | chr6 | 152008514 | 39 | 32 | 3163.731733 | 2575.538132 | 0.7334 |
| BAX | chr19 | 49465094 | 32 | 32 | 2978.388246 | 3007.6123 | 0.73362 |
| APOA1 | chr11 | 116703351 | 32 | 32 | 2511.922543 | 2538.41079 | 0.73374 |
| LCK | chr1 | 32713404 | 32 | 32 | 3023.445717 | 3043.493708 | 0.73382 |
| LPA | chr6 | 11221340 | 28 | 30 | 2470.362744 | 2591.63624 | 0.7342 |
| TFPI | chr2 | 188424643 | 9 | 3 | 822.5 | 283.3333333 | 0.73428 |
| GALNT2 | chr1 | 230191197 | 34 | 32 | 3211.898511 | 3005.714454 | 0.73444 |
| CTGF | chr6 | 132276769 | 33 | 32 | 2764.279574 | 2654.987891 | 0.73468 |
| APOA1 | chr11 | 116661111 | 34 | 32 | 3082.897261 | 2878.2613 | 0.73484 |
| CDKN2B-AS1 | chr9 | 21967932 | 32 | 27 | 1176.855523 | 962.0871932 | 0.73514 |
| APOA1 | chr11 | 116692440 | 35 | 32 | 3104.709902 | 2861.946449 | 0.7354 |
| TERT | chr5 | 1299983 | 32 | 33 | 2613.592627 | 2661.279768 | 0.73564 |
| APOA1 | chr11 | 116692195 | 33 | 32 | 2985.626806 | 2913.872661 | 0.73568 |
| LPA | chr6 | 11220554 | 34 | 32 | 3191.646668 | 3022.566107 | 0.73588 |
| COL15A1 | chr9 | 101704466 | 30 | 32 | 2789.970012 | 2993.630567 | 0.73616 |
| CDKN2B-AS1 | chr9 | 22008026 | 3 | 5 | 235.130719 | 402.579259 | 0.73626 |
| LOXL1 | chr15 | 74218884 | 27 | 25 | 889.9599999 | 794.0365985 | 0.73678 |

| | | | | | | | |
|---|---|---|---|---|---|---|---|
| LCK | chr1 | 32740552 | 15 | 19 | 1367.625153 | 1704.484127 | 0.73678 |
| ICAM2 | chr17 | 62085467 | 30 | 31 | 2829.845925 | 2901.240111 | 0.7368 |
| MCPH1 | chr8 | 6470873 | 30 | 30 | 2828.18572 | 2810.319257 | 0.73682 |
| LOXL1 | chr15 | 74216845 | 8 | 9 | 221.6477273 | 272.9339392 | 0.737 |
| NOS3 | chr7 | 150688849 | 30 | 31 | 2340.284347 | 2374.15626 | 0.73702 |
| F7 | chr13 | 113758367 | 30 | 30 | 2801.50457 | 2779.569602 | 0.73726 |
| F7 | chr13 | 113750759 | 27 | 27 | 2319.579587 | 2349.790765 | 0.7375 |
| BCL2 | chr18 | 60989069 | 20 | 19 | 1216.857864 | 1190.833333 | 0.73752 |
| ALDOA | chr16 | 30063148 | 33 | 33 | 1498.872342 | 1465.265881 | 0.7377 |
| F7 | chr13 | 113750441 | 3 | 5 | 186.6666667 | 288.8888889 | 0.73784 |
| PECAM1 | chr17 | 62399778 | 30 | 32 | 2539.745451 | 2675.792788 | 0.738 |
| ALOX15 | chr17 | 4534083 | 29 | 25 | 2604.437229 | 2270.270563 | 0.73838 |
| ESR2 | chr14 | 64806624 | 29 | 28 | 2618.851704 | 2482.923563 | 0.73838 |
| E2F6 | chr2 | 11616273 | 35 | 32 | 2892.759729 | 2625.06927 | 0.73852 |
| FOXP3 | chrX | 49114546 | 17 | 11 | 574.2272348 | 392.0634921 | 0.73854 |
| F2RL3 | chr19 | 16990514 | 28 | 30 | 2694.16514 | 2852.293956 | 0.73858 |
| FOXP3 | chrX | 49127089 | 18 | 15 | 1624.642857 | 1331.984127 | 0.73912 |
| LPA | chr6 | 160945954 | 31 | 27 | 2830.075902 | 2488.803696 | 0.73948 |
| CTGF | chr6 | 132280132 | 41 | 33 | 2973.774098 | 2364.865166 | 0.7395 |
| LPA | chr6 | 11222805 | 31 | 33 | 1890.373108 | 1976.262329 | 0.73966 |
| APOA1 | chr11 | 116693536 | 34 | 32 | 3087.227024 | 2887.803209 | 0.73972 |
| APOA1 | chr11 | 116690249 | 34 | 32 | 3230.540196 | 3027.045614 | 0.74002 |
| MCPH1 | chr8 | 6257953 | 32 | 32 | 3103.744827 | 3090.269031 | 0.7404 |
| AKT2 | chr19 | 40800856 | 35 | 32 | 3380.764767 | 3102.25232 | 0.74042 |
| LCK | chr1 | 32716507 | 26 | 27 | 1009.741322 | 1077.850074 | 0.74052 |
| NOS3 | chr7 | 150692137 | 23 | 25 | 905.2322418 | 942.1051799 | 0.74052 |
| TERT | chr5 | 1296156 | 34 | 33 | 2820.414128 | 2705.395717 | 0.7406 |
| APOA1 | chr11 | 116660973 | 34 | 32 | 3137.999538 | 2934.888306 | 0.74066 |
| P2RY12 | chr3 | 151107472 | 37 | 32 | 2531.450716 | 2233.280007 | 0.74116 |
| LCK | chr1 | 32713137 | 32 | 31 | 2993.466528 | 2879.209729 | 0.74116 |
| NOS3 | chr7 | 150706175 | 32 | 32 | 2548.550098 | 2517.556747 | 0.74124 |
| F7 | chr13 | 113758738 | 31 | 29 | 2995.094073 | 2786.737892 | 0.74128 |
| ABCB1 | chr7 | 87352350 | 3 | 6 | 132.1052632 | 297.8571429 | 0.7413 |
| APOA1 | chr11 | 116674716 | 24 | 29 | 1459.43297 | 1722.579365 | 0.74152 |
| ALDOA | chr16 | 30063046 | 37 | 32 | 3476.250697 | 3018.529819 | 0.7422 |
| LPA | chr6 | 160942747 | 39 | 32 | 3687.843355 | 3016.249378 | 0.7426 |
| GALNT2 | chr1 | 230186060 | 37 | 32 | 3467.637635 | 3010.214318 | 0.74268 |
| ABCG1 | chr21 | 43636741 | 36 | 32 | 3338.669648 | 2950.062859 | 0.74288 |
| LCK | chr1 | 32741530 | 30 | 31 | 2678.232612 | 2791.851406 | 0.74308 |
| LOXL1 | chr15 | 74216568 | 16 | 18 | 535.3744565 | 578.1940223 | 0.74326 |
| APOA1 | chr11 | 116698323 | 5 | 7 | 420 | 623.3333333 | 0.74336 |
| CDKN2B-AS1 | chr9 | 21967707 | 35 | 31 | 1280.770764 | 1158.494534 | 0.74364 |
| LPA | chr6 | 11224786 | 6 | 8 | 245.3571429 | 303.1168831 | 0.74396 |
| GCK | chr7 | 44230839 | 33 | 31 | 2931.874075 | 2778.901906 | 0.744 |
| PLA2G7 | chr6 | 46712959 | 38 | 32 | 3225.020571 | 2733.400848 | 0.74404 |
| F2RL3 | chr19 | 16994981 | 10 | 7 | 940.7142857 | 645.8333333 | 0.74416 |
| CDKN2B-AS1 | chr9 | 22008075 | 34 | 32 | 1444.770038 | 1388.967131 | 0.74422 |

| | | | | | | | |
|---|---|---|---|---|---|---|---|
| LPA | chr6 | 160949185 | 30 | 31 | 2426.930112 | 2471.914905 | 0.74444 |
| FOXP3 | chrX | 49115701 | 25 | 21 | 2326.092437 | 1932.579365 | 0.74452 |
| ESR1 | chr6 | 152007101 | 9 | 6 | 830 | 540 | 0.74472 |
| PHACTR1 | chr6 | 12707873 | 35 | 32 | 2421.54433 | 2246.765058 | 0.745 |
| ESR2 | chr14 | 64809225 | 32 | 32 | 3090.488632 | 3107.727562 | 0.74532 |
| AKT2 | chr19 | 40791835 | 32 | 32 | 1086.245637 | 1061.714714 | 0.74562 |
| ABCA1 | chr9 | 107695353 | 20 | 17 | 1077.193146 | 953.7308743 | 0.74632 |
| LDLR | chr19 | 11224068 | 21 | 17 | 1989.285714 | 1631.222944 | 0.74664 |
| PECAM1 | chr17 | 62398361 | 32 | 32 | 3085.043763 | 3063.429592 | 0.74668 |
| LDLR | chr19 | 11221080 | 21 | 20 | 2004.603175 | 1933.055556 | 0.74704 |
| APOA1 | chr11 | 116718111 | 19 | 20 | 1644.047619 | 1761.379552 | 0.74724 |
| APOA1 | chr11 | 116684466 | 37 | 32 | 3442.438088 | 2965.154088 | 0.74732 |
| ALDOA | chr16 | 30060371 | 25 | 28 | 2321.937229 | 2624.152514 | 0.7475 |
| ADRA2C | chr4 | 3763812 | 33 | 33 | 2725.395274 | 2687.491212 | 0.7481 |
| CDKN2B-AS1 | chr9 | 21982509 | 34 | 33 | 2579.374164 | 2531.872364 | 0.74812 |
| LDLR | chr19 | 11224931 | 28 | 32 | 2270.588688 | 2530.473677 | 0.74816 |
| F7 | chr13 | 113751516 | 29 | 25 | 2780.211733 | 2380.849341 | 0.7482 |
| TIMP3 | chr22 | 33191803 | 40 | 32 | 3779.454133 | 3012.93665 | 0.74832 |
| PHACTR1 | chr6 | 12718031 | 38 | 32 | 3577.775887 | 3002.060632 | 0.74866 |
| ABCG1 | chr21 | 43637323 | 33 | 32 | 3206.37049 | 3096.022337 | 0.74866 |
| APOA1 | chr11 | 116688366 | 34 | 32 | 3184.557006 | 2974.692971 | 0.74904 |
| MCPH1 | chr8 | 6260592 | 10 | 11 | 473.3085542 | 555.9992806 | 0.74948 |
| CDKN1A | chr6 | 36654284 | 6 | 6 | 460.3571429 | 483.3333333 | 0.74994 |
| NOS3 | chr7 | 150695700 | 27 | 28 | 2617.179952 | 2730.439005 | 0.74998 |
| F2RL3 | chr19 | 17002948 | 27 | 29 | 2047.205775 | 2231.632118 | 0.75006 |
| ESR2 | chr14 | 64808379 | 17 | 22 | 1554.059829 | 1982.25469 | 0.7502 |
| CDKN2B-AS1 | chr9 | 21966553 | 32 | 33 | 2257.332671 | 2366.822284 | 0.75038 |
| GCK | chr7 | 44229893 | 34 | 32 | 2903.730932 | 2700.340645 | 0.7511 |
| NOS3 | chr7 | 150689889 | 16 | 16 | 826.7640693 | 799.3253968 | 0.75158 |
| ADRA2C | chr4 | 3761278 | 30 | 31 | 2636.257071 | 2745.933527 | 0.75174 |
| CDKN2B-AS1 | chr9 | 22011094 | 14 | 7 | 413.2649031 | 224.956182 | 0.7521 |
| BCL2 | chr18 | 60990023 | 19 | 13 | 1067.46915 | 697.5066407 | 0.75216 |
| F7 | chr13 | 113751845 | 32 | 32 | 2723.631637 | 2699.449425 | 0.7525 |
| F7 | chr13 | 113775163 | 21 | 25 | 1937.777778 | 2340.27417 | 0.7529 |
| GCK | chr7 | 44234804 | 36 | 32 | 3416.610606 | 3052.758011 | 0.753 |
| APOA1 | chr11 | 116692331 | 33 | 31 | 3060.416256 | 2889.861932 | 0.75304 |
| LPA | chr6 | 160944248 | 25 | 25 | 2060.584971 | 2022.882395 | 0.75326 |
| APOA1 | chr11 | 116713691 | 35 | 32 | 3306.557953 | 3034.685508 | 0.75338 |
| MIA3 | chr1 | 222841215 | 37 | 32 | 3563.847279 | 3072.530855 | 0.75346 |
| AMOTL2 | chr3 | 134102221 | 18 | 14 | 1631.547619 | 1253.214286 | 0.75348 |
| LPA | chr6 | 160943685 | 30 | 28 | 2873.190629 | 2695.135281 | 0.75362 |
| ARHGEF18 | chr19 | 7469806 | 35 | 32 | 3412.05126 | 3104.737274 | 0.75372 |
| PECAM1 | chr17 | 62400575 | 14 | 17 | 1170.6676 | 1382.267696 | 0.75372 |
| APOA1 | chr11 | 116711473 | 35 | 32 | 3231.150251 | 2971.067257 | 0.7538 |
| LDLR | chr19 | 11220164 | 29 | 32 | 2773.511586 | 3040.782053 | 0.75396 |
| LDLR | chr19 | 11218199 | 33 | 32 | 3199.321236 | 3121.506562 | 0.754 |
| TMPO | chr12 | 98906041 | 33 | 32 | 3213.397632 | 3126.785624 | 0.75426 |

| | | | | | | | |
|---|---|---|---|---|---|---|---|
| FGF2 | chr4 | 123743977 | 35 | 32 | 3309.743408 | 3013.231934 | 0.75454 |
| CDKN2B-AS1 | chr9 | 21989864 | 16 | 18 | 714.0079365 | 837.3015873 | 0.75476 |
| F7 | chr13 | 113750852 | 31 | 32 | 2933.414903 | 3011.05927 | 0.75502 |
| APOA1 | chr11 | 116692389 | 33 | 32 | 3108.845434 | 3029.123404 | 0.75502 |
| LPA | chr6 | 11216941 | 38 | 32 | 3632.613146 | 3067.751874 | 0.75522 |
| MCPH1 | chr8 | 6470878 | 30 | 30 | 2839.867856 | 2855.961268 | 0.75564 |
| CDKN2B-AS1 | chr9 | 21981674 | 18 | 14 | 695.0108225 | 568.7012987 | 0.75572 |
| COL15A1 | chr9 | 101699709 | 33 | 33 | 2372.963737 | 2334.84327 | 0.75602 |
| F2RL3 | chr19 | 16990306 | 32 | 31 | 2807.823109 | 2763.894635 | 0.75612 |
| APOA1 | chr11 | 116698167 | 7 | 7 | 531.1111111 | 548.9285714 | 0.75626 |
| F7 | chr13 | 113758041 | 31 | 33 | 2317.20476 | 2432.618059 | 0.75628 |
| AMOTL2 | chr3 | 134100876 | 35 | 33 | 3139.972573 | 2988.60456 | 0.75632 |
| GALNT2 | chr1 | 230190991 | 33 | 32 | 3088.785737 | 2976.09428 | 0.75656 |
| LDLR | chr19 | 11216975 | 6 | 10 | 587.5 | 968.8888889 | 0.75698 |
| HMGCR | chr5 | 74631168 | 41 | 32 | 3984.05324 | 3098.842194 | 0.75724 |
| HMGCR | chr5 | 74623601 | 6 | 2 | 543.3333333 | 173.2142857 | 0.7576 |
| HMGCR | chr5 | 74631897 | 39 | 33 | 2032.389746 | 1751.823808 | 0.75772 |
| APOA1 | chr11 | 116691864 | 32 | 32 | 2999.262445 | 3016.512089 | 0.75796 |
| ABCG1 | chr21 | 43637525 | 31 | 32 | 2913.854216 | 2990.193777 | 0.75802 |
| CDKN2B-AS1 | chr9 | 21964696 | 26 | 28 | 1139.244396 | 1259.28911 | 0.75802 |
| MCPH1 | chr8 | 6471088 | 6 | 7 | 580 | 666.6666667 | 0.75854 |
| TERT | chr5 | 1301191 | 34 | 32 | 3124.130129 | 2959.887579 | 0.75872 |
| F2RL3 | chr19 | 16994089 | 22 | 28 | 1503.341737 | 1977.659006 | 0.75888 |
| LCK | chr1 | 32711449 | 34 | 32 | 3142.684517 | 2968.856139 | 0.75896 |
| LPA | chr6 | 11219629 | 33 | 32 | 2894.429543 | 2820.40985 | 0.75914 |
| LOXL1 | chr15 | 74211618 | 12 | 10 | 1108.888889 | 943.3333333 | 0.75916 |
| ADRA2C | chr4 | 3766630 | 9 | 13 | 542.5 | 815.3787879 | 0.75932 |
| APOA1 | chr11 | 116712775 | 25 | 25 | 2204.241453 | 2171.011799 | 0.75962 |
| APOA1 | chr11 | 116718995 | 10 | 11 | 901.4285714 | 975.8333333 | 0.75974 |
| MIA3 | chr1 | 222815047 | 33 | 32 | 3121.130378 | 3054.167518 | 0.76044 |
| ICAM2 | chr17 | 62086287 | 31 | 33 | 1909.373151 | 2068.332479 | 0.76078 |
| MIA3 | chr1 | 222816858 | 8 | 11 | 521.1904762 | 694.0476191 | 0.76088 |
| LDLR | chr19 | 11220672 | 13 | 18 | 1266.666667 | 1765.714286 | 0.76094 |
| LDLR | chr19 | 11190087 | 17 | 23 | 1610.238095 | 2197.575758 | 0.76114 |
| APOA1 | chr11 | 116682290 | 36 | 32 | 2961.817524 | 2611.105264 | 0.76124 |
| LPA | chr6 | 11216280 | 32 | 32 | 2858.836407 | 2813.542701 | 0.76144 |
| ALDOA | chr16 | 30057810 | 33 | 32 | 2727.354305 | 2664.72471 | 0.7616 |
| NR3C1 | chr5 | 142790684 | 32 | 33 | 2975.383441 | 3042.866677 | 0.76186 |
| APOA1 | chr11 | 116709265 | 34 | 33 | 2429.745214 | 2389.85097 | 0.76204 |
| LCK | chr1 | 32707582 | 27 | 26 | 2473.468615 | 2353.127706 | 0.7623 |
| COL15A1 | chr9 | 101698737 | 35 | 32 | 3204.932706 | 2914.578625 | 0.76232 |
| KLF2 | chr19 | 16437867 | 20 | 24 | 754.8256446 | 873.7018545 | 0.7624 |
| P2RY12 | chr3 | 151105707 | 29 | 32 | 2612.291815 | 2858.2771 | 0.76254 |
| LDLR | chr19 | 11192771 | 29 | 32 | 2798.619714 | 3105.416667 | 0.76256 |
| ALDOA | chr16 | 30061581 | 27 | 31 | 2427.207107 | 2755.378151 | 0.76264 |
| ESR1 | chr6 | 152003020 | 21 | 17 | 598.593217 | 507.0383912 | 0.76264 |
| GALNT2 | chr1 | 230191555 | 12 | 12 | 920.1515152 | 951.1111111 | 0.76316 |

| | | | | | | | |
|---|---|---|---|---|---|---|---|
| ESR2 | chr14 | 64812322 | 39 | 32 | 3359.845619 | 2742.768422 | 0.76346 |
| CDKN2B-AS1 | chr9 | 21978673 | 11 | 9 | 314.4018008 | 277.8457142 | 0.76352 |
| LOXL1 | chr15 | 74219184 | 17 | 20 | 672.6519167 | 755.6114096 | 0.76446 |
| APOA1 | chr11 | 116699590 | 33 | 32 | 3039.525071 | 2967.84347 | 0.7648 |
| CDKN2B-AS1 | chr9 | 22011346 | 33 | 32 | 2947.123653 | 2844.540983 | 0.76494 |
| LOXL1 | chr15 | 74218904 | 28 | 17 | 913.3149444 | 573.7129509 | 0.76614 |
| ADRA2C | chr4 | 3759490 | 31 | 32 | 2942.206117 | 3020.868147 | 0.7663 |
| APOA1 | chr11 | 116696420 | 35 | 32 | 3260.237573 | 2966.184252 | 0.76646 |
| TERT | chr5 | 1295606 | 2 | 4 | 58.33333333 | 128.5714286 | 0.7667 |
| TIMP3 | chr22 | 33193550 | 37 | 32 | 3464.647693 | 3006.027538 | 0.76682 |
| CDKN1A | chr6 | 36643510 | 28 | 25 | 2562.945527 | 2307.733845 | 0.76682 |
| FOXP3 | chrX | 49122211 | 35 | 32 | 3254.510144 | 2990.466242 | 0.76704 |
| ESR2 | chr14 | 64810000 | 36 | 32 | 3248.16007 | 2908.262691 | 0.76716 |
| LPA | chr6 | 160946987 | 36 | 32 | 3378.213518 | 2990.842381 | 0.76724 |
| ESR2 | chr14 | 64813327 | 33 | 32 | 3114.162511 | 3008.478579 | 0.76818 |
| F7 | chr13 | 113755477 | 21 | 28 | 1850.649351 | 2503.412698 | 0.76826 |
| ADRA2C | chr4 | 3761250 | 30 | 33 | 1941.918524 | 2101.500126 | 0.76846 |
| GCK | chr7 | 44234235 | 34 | 33 | 2700.877883 | 2596.393857 | 0.76862 |
| KLF2 | chr19 | 16426978 | 33 | 30 | 1398.275884 | 1240.519568 | 0.76896 |
| F7 | chr13 | 113758718 | 29 | 29 | 2545.804853 | 2520.300865 | 0.76934 |
| CDKN2B-AS1 | chr9 | 21989605 | 13 | 9 | 425.4017023 | 270.8670741 | 0.7695 |
| CDKN2B-AS1 | chr9 | 21989308 | 26 | 19 | 872.5347837 | 615.7884856 | 0.76962 |
| APOA1 | chr11 | 116661857 | 32 | 32 | 1745.938627 | 1712.588058 | 0.76966 |
| ALOX15 | chr17 | 4552869 | 27 | 21 | 2362.77417 | 1814.036797 | 0.76986 |
| ATF3 | chr1 | 212743583 | 32 | 32 | 2837.110808 | 2816.613486 | 0.77016 |
| KLF2 | chr19 | 16436131 | 3 | 4 | 70 | 99.72222222 | 0.77056 |
| APOA1 | chr11 | 116674433 | 18 | 14 | 1047.123016 | 847.6190476 | 0.77112 |
| LDLR | chr19 | 11218189 | 33 | 32 | 3197.345605 | 3115.191259 | 0.77116 |
| DHFR | chr5 | 79953582 | 17 | 21 | 1010.53156 | 1299.200301 | 0.77136 |
| P2RY12 | chr3 | 151105480 | 33 | 33 | 1780.920768 | 1809.967531 | 0.77146 |
| APOA1 | chr11 | 116672965 | 17 | 25 | 1598.885281 | 2374.047619 | 0.7715 |
| KLF2 | chr19 | 16437887 | 17 | 23 | 617.1176828 | 804.8302351 | 0.77168 |
| APOA1 | chr11 | 116707737 | 31 | 31 | 2996.679848 | 2979.505197 | 0.77204 |
| FOXP3 | chrX | 49127023 | 20 | 11 | 1913.333333 | 1061.428571 | 0.77218 |
| LDLR | chr19 | 11220938 | 13 | 15 | 1265.714286 | 1449.047619 | 0.77238 |
| COL15A1 | chr9 | 101706281 | 3 | 3 | 68.57142857 | 65.75163398 | 0.77242 |
| NOS3 | chr7 | 150705811 | 32 | 33 | 1475.396545 | 1550.931367 | 0.77272 |
| CDKN2B-AS1 | chr9 | 21970711 | 2 | 3 | 71.66666667 | 138.0952381 | 0.77274 |
| LCK | chr1 | 32716212 | 26 | 24 | 928.8974832 | 835.3419077 | 0.77312 |
| TFPI | chr2 | 188425221 | 24 | 27 | 2183.174603 | 2478.000611 | 0.77318 |
| GALNT2 | chr1 | 230187460 | 38 | 32 | 3689.091182 | 3091.680532 | 0.7736 |
| TERT | chr5 | 1297978 | 7 | 6 | 600 | 488.8888889 | 0.77364 |
| PECAM1 | chr17 | 62400137 | 6 | 5 | 571.4285714 | 466.6666667 | 0.77364 |
| F2RL3 | chr19 | 16994937 | 19 | 16 | 1786.879787 | 1520.006105 | 0.77368 |
| GCK | chr7 | 44232890 | 28 | 30 | 2397.753068 | 2596.776557 | 0.77404 |
| GALNT2 | chr1 | 230187303 | 32 | 31 | 2955.482431 | 2879.435309 | 0.7742 |
| TERT | chr5 | 1253690 | 32 | 32 | 2999.461154 | 3020.184997 | 0.77494 |

| | | | | | | | |
|---|---|---|---|---|---|---|---|
| F7 | chr13 | 113751064 | 13 | 13 | 1251.388889 | 1263.333333 | 0.77494 |
| NOS3 | chr7 | 150693935 | 7 | 13 | 240 | 477.0238095 | 0.77496 |
| KLF2 | chr19 | 16437896 | 29 | 29 | 1370.323358 | 1406.072962 | 0.77526 |
| NOS3 | chr7 | 150711138 | 18 | 21 | 684.4203264 | 766.0892105 | 0.7755 |
| NOS3 | chr7 | 150695629 | 19 | 20 | 1677.135642 | 1783.524531 | 0.7766 |
| KLF2 | chr19 | 16427723 | 2 | 5 | 100.4545455 | 279.3604651 | 0.7766 |
| NOS3 | chr7 | 150710282 | 29 | 31 | 1466.819106 | 1601.637162 | 0.77662 |
| APOA1 | chr11 | 116696952 | 32 | 30 | 3102.595048 | 2898.056249 | 0.7767 |
| KLF2 | chr19 | 16433157 | 6 | 6 | 433.8095238 | 450 | 0.7767 |
| LPA | chr6 | 11221930 | 39 | 32 | 3718.256261 | 3059.230224 | 0.77692 |
| NOS3 | chr7 | 150694702 | 33 | 32 | 3042.266869 | 2962.027897 | 0.77732 |
| APOA1 | chr11 | 116691994 | 32 | 32 | 3033.656585 | 3019.912904 | 0.77758 |
| CDKN2B-AS1 | chr9 | 22007558 | 18 | 16 | 489.17773 | 453.0395266 | 0.7776 |
| E2F6 | chr2 | 11614038 | 34 | 32 | 2910.073795 | 2756.219315 | 0.77764 |
| NOS3 | chr7 | 150687420 | 37 | 33 | 2834.933222 | 2553.056303 | 0.7777 |
| TERT | chr5 | 1300269 | 31 | 32 | 2487.714657 | 2605.58866 | 0.7782 |
| LCK | chr1 | 32742014 | 29 | 26 | 2406.214838 | 2135.1554 | 0.7782 |
| BAX | chr19 | 49449842 | 7 | 8 | 402.5595238 | 487.6624462 | 0.77866 |
| TERT | chr5 | 1295798 | 33 | 30 | 1361.058937 | 1263.503649 | 0.7788 |
| MIA3 | chr1 | 222809058 | 38 | 32 | 3653.161474 | 3069.666716 | 0.7789 |
| NOS3 | chr7 | 150710654 | 3 | 2 | 78.33333333 | 48.57142857 | 0.77908 |
| APOA1 | chr11 | 116696067 | 32 | 32 | 2746.44755 | 2719.348018 | 0.77912 |
| ABCA1 | chr9 | 107693359 | 3 | 5 | 287.5 | 451.4285714 | 0.7798 |
| CDKN2B-AS1 | chr9 | 21970782 | 30 | 29 | 1036.823334 | 1026.507165 | 0.77982 |
| CDKN2B-AS1 | chr9 | 21971422 | 8 | 6 | 232.6717632 | 194.8534504 | 0.78022 |
| LDLR | chr19 | 11216225 | 12 | 13 | 1138.888889 | 1218.333333 | 0.78078 |
| LDLR | chr19 | 11217341 | 33 | 32 | 3215.121152 | 3105.551583 | 0.78082 |
| FOXP3 | chrX | 49128552 | 34 | 32 | 3164.011818 | 2993.700358 | 0.78098 |
| LDLR | chr19 | 11217473 | 33 | 29 | 3044.615873 | 2692.250348 | 0.78112 |
| P2RY12 | chr3 | 151106158 | 31 | 25 | 2156.149291 | 1692.292661 | 0.78118 |
| PHACTR1 | chr6 | 12711300 | 37 | 33 | 2500.32857 | 2203.018553 | 0.78138 |
| E2F6 | chr2 | 11607901 | 29 | 24 | 2594.039849 | 2122.499445 | 0.7814 |
| APOA1 | chr11 | 116697193 | 32 | 32 | 2908.495028 | 2930.191022 | 0.78164 |
| NR3C1 | chr5 | 142785359 | 33 | 32 | 2862.738302 | 2760.442312 | 0.78188 |
| KLF2 | chr19 | 16434626 | 32 | 32 | 2629.753784 | 2604.753361 | 0.78194 |
| APOA1 | chr11 | 116707210 | 29 | 31 | 1334.598804 | 1463.280729 | 0.78196 |
| FOXP3 | chrX | 49120142 | 34 | 32 | 3173.116601 | 3001.041711 | 0.78202 |
| PECAM1 | chr17 | 62409296 | 34 | 31 | 1609.830186 | 1434.237511 | 0.78232 |
| CDKN2B-AS1 | chr9 | 21980730 | 20 | 21 | 1879.047619 | 1953.333333 | 0.7824 |
| CDKN2B-AS1 | chr9 | 22000178 | 37 | 33 | 2068.537449 | 1820.939261 | 0.78244 |
| TIMP3 | chr22 | 33196504 | 32 | 32 | 2856.24832 | 2822.906715 | 0.78274 |
| NOS3 | chr7 | 150706297 | 33 | 31 | 3069.856007 | 2897.875835 | 0.78276 |
| TFPI | chr2 | 188426874 | 18 | 10 | 1655.47619 | 932.3809524 | 0.78298 |
| LCK | chr1 | 32741141 | 19 | 17 | 604.022089 | 563.4543819 | 0.78324 |
| F2RL3 | chr19 | 16993527 | 33 | 32 | 3168.76973 | 3083.67218 | 0.7835 |
| LPA | chr6 | 160948309 | 29 | 29 | 2649.718615 | 2626.226551 | 0.78352 |
| CTGF | chr6 | 132272349 | 2 | 4 | 65 | 165 | 0.78354 |

| | | | | | | | |
|---|---|---|---|---|---|---|---|
| LPA | chr6 | 11221301 | 33 | 32 | 2893.999369 | 2787.535571 | 0.78376 |
| ARHGEF18 | chr19 | 7469207 | 34 | 32 | 3250.317711 | 3046.237682 | 0.78388 |
| ADRA2C | chr4 | 3766258 | 33 | 32 | 2983.171519 | 2869.000944 | 0.78388 |
| GCK | chr7 | 44237823 | 29 | 29 | 2524.98484 | 2492.118731 | 0.7842 |
| ABCA1 | chr9 | 107693408 | 24 | 23 | 2103.639971 | 2043.409091 | 0.78422 |
| APOA1 | chr11 | 116704195 | 25 | 27 | 2367.371615 | 2571.302309 | 0.7844 |
| KLF2 | chr19 | 16429104 | 31 | 29 | 2884.530493 | 2715.263633 | 0.78508 |
| AMOTL2 | chr3 | 134074440 | 27 | 29 | 2537.561703 | 2741.826564 | 0.78558 |
| APOA1 | chr11 | 116679587 | 35 | 32 | 2232.444099 | 2065.56025 | 0.78596 |
| AMOTL2 | chr3 | 134097042 | 34 | 32 | 1641.450976 | 1518.637506 | 0.78622 |
| APOA1 | chr11 | 116705644 | 15 | 14 | 570.515873 | 556.8253968 | 0.78622 |
| TIMP3 | chr22 | 33187665 | 35 | 32 | 3299.761638 | 3027.516401 | 0.78636 |
| DHFR | chr5 | 79959023 | 32 | 32 | 2870.847358 | 2848.326094 | 0.78648 |
| FOXP3 | chrX | 49116584 | 21 | 23 | 1973.968254 | 2142.819958 | 0.7865 |
| LCK | chr1 | 32707297 | 15 | 15 | 412.1989559 | 431.2465531 | 0.78694 |
| GCK | chr7 | 44228655 | 30 | 32 | 2581.087755 | 2774.071705 | 0.78704 |
| FOXP3 | chrX | 49126838 | 11 | 5 | 408.8888889 | 170 | 0.7871 |
| ALOX15 | chr17 | 4533860 | 31 | 27 | 2858.126971 | 2507.920568 | 0.78742 |
| FOXP3 | chrX | 49123779 | 4 | 5 | 346.6666667 | 445.7142857 | 0.78742 |
| ABCG1 | chr21 | 43634708 | 6 | 6 | 554.7619048 | 563.3333333 | 0.7875 |
| AKT2 | chr19 | 40800691 | 37 | 32 | 3495.886475 | 3012.536536 | 0.78752 |
| LPA | chr6 | 160944306 | 32 | 32 | 2377.265687 | 2350.846944 | 0.78754 |
| PHACTR1 | chr6 | 12714071 | 36 | 33 | 2548.606442 | 2306.861191 | 0.78758 |
| F7 | chr13 | 113751395 | 28 | 26 | 2593.844489 | 2389.52381 | 0.7879 |
| CDKN2B-AS1 | chr9 | 21979676 | 32 | 30 | 1609.903681 | 1480.891619 | 0.78826 |
| NOS3 | chr7 | 150690136 | 3 | 6 | 240 | 503.3333333 | 0.78832 |
| LPA | chr6 | 11221416 | 2 | 3 | 183.3333333 | 283.3333333 | 0.78878 |
| GCK | chr7 | 44232524 | 39 | 32 | 3779.697374 | 3106.590547 | 0.78908 |
| BAX | chr19 | 49456424 | 4 | 6 | 351.4285714 | 539.047619 | 0.78922 |
| LCK | chr1 | 32740563 | 15 | 18 | 1203.686869 | 1470.833333 | 0.78942 |
| GCK | chr7 | 44230013 | 32 | 33 | 2285.509681 | 2384.737886 | 0.78948 |
| CDKN1A | chr6 | 36648794 | 33 | 32 | 2654.653547 | 2552.623978 | 0.7895 |
| APOA1 | chr11 | 116724177 | 30 | 32 | 2040.405814 | 2219.385337 | 0.78968 |
| PLA2G7 | chr6 | 46707580 | 31 | 29 | 1864.162157 | 1697.021461 | 0.78996 |
| APOA1 | chr11 | 116719913 | 34 | 32 | 3325.814064 | 3122.543534 | 0.78998 |
| ADRA2C | chr4 | 3761793 | 33 | 32 | 2979.736986 | 2873.702523 | 0.79012 |
| APOA1 | chr11 | 116681403 | 9 | 4 | 828.8888889 | 380 | 0.79054 |
| ADRA2C | chr4 | 3761215 | 32 | 33 | 2777.08037 | 2833.937421 | 0.79064 |
| CDKN1A | chr6 | 36651917 | 31 | 32 | 2924.09753 | 3031.847762 | 0.79068 |
| PLA2G7 | chr6 | 46709235 | 37 | 32 | 3503.316642 | 3021.359993 | 0.79086 |
| ADRA2C | chr4 | 3762510 | 23 | 26 | 1910.802823 | 2126.778499 | 0.79092 |
| BAX | chr19 | 49465285 | 32 | 32 | 2290.415456 | 2261.918422 | 0.79098 |
| APOA1 | chr11 | 116662559 | 5 | 4 | 451.4285714 | 371.4285714 | 0.791 |
| APOA1 | chr11 | 116663104 | 26 | 29 | 1641.093127 | 1862.199285 | 0.7911 |
| LOXL1 | chr15 | 74217647 | 32 | 32 | 3052.93902 | 3062.39776 | 0.79116 |
| AMOTL2 | chr3 | 134102269 | 32 | 32 | 2858.59015 | 2838.127317 | 0.7914 |
| PECAM1 | chr17 | 62408924 | 28 | 29 | 1207.616798 | 1285.534731 | 0.7914 |

| | | | | | | | |
|---|---|---|---|---|---|---|---|
| F7 | chr13 | 113758384 | 31 | 32 | 2859.453792 | 2979.083875 | 0.7915 |
| PECAM1 | chr17 | 62408908 | 25 | 27 | 1188.726869 | 1316.648142 | 0.79194 |
| FGF2 | chr4 | 123742913 | 15 | 17 | 1374.801587 | 1572.178932 | 0.79214 |
| F7 | chr13 | 113760313 | 18 | 18 | 1363.445166 | 1387.142857 | 0.79226 |
| ABCA1 | chr9 | 107690792 | 10 | 13 | 425.2092352 | 529.9206349 | 0.79266 |
| E2F6 | chr2 | 11613212 | 14 | 22 | 1286.547619 | 2045.436508 | 0.79268 |
| CDKN2B-AS1 | chr9 | 21989890 | 14 | 15 | 602.5396825 | 674.6626984 | 0.79294 |
| APOA1 | chr11 | 116694342 | 25 | 22 | 2088.716284 | 1817.166167 | 0.79304 |
| APOA1 | chr11 | 116661044 | 33 | 32 | 3130.449979 | 3026.303978 | 0.7931 |
| CDKN1A | chr6 | 36653048 | 35 | 32 | 3361.038845 | 3066.003588 | 0.79328 |
| CDKN1A | chr6 | 36651903 | 31 | 32 | 2846.772833 | 2956.141397 | 0.79334 |
| CDKN2B-AS1 | chr9 | 21970967 | 15 | 15 | 474.0068995 | 454.6535628 | 0.79364 |
| LCK | chr1 | 32741131 | 17 | 16 | 577.8632478 | 564.5970575 | 0.7943 |
| APOA1 | chr11 | 116713878 | 34 | 32 | 3181.032792 | 2981.584391 | 0.79454 |
| CDKN2BAS | chr9 | 22103517 | 33 | 33 | 1579.75295 | 1556.034039 | 0.79464 |
| APOA1 | chr11 | 116691861 | 32 | 32 | 2938.515945 | 2924.007715 | 0.79466 |
| BAX | chr19 | 49456536 | 30 | 30 | 1430.468567 | 1399.426128 | 0.79474 |
| CDKN2B-AS1 | chr9 | 21996333 | 34 | 30 | 1164.367301 | 1051.181207 | 0.79504 |
| CDKN2B-AS1 | chr9 | 21990108 | 14 | 14 | 571.1577965 | 546.953602 | 0.79556 |
| P2RY12 | chr3 | 151105583 | 35 | 32 | 3204.695809 | 2914.883037 | 0.79588 |
| APOA1 | chr11 | 116706373 | 4 | 2 | 95.55555555 | 45 | 0.79598 |
| NOS3 | chr7 | 150692473 | 31 | 32 | 1313.086451 | 1328.656409 | 0.796 |
| GALNT2 | chr1 | 230191234 | 35 | 32 | 2939.753364 | 2706.523907 | 0.79618 |
| LCK | chr1 | 32713067 | 32 | 32 | 3045.91426 | 3033.545336 | 0.79628 |
| NOS3 | chr7 | 150694826 | 33 | 32 | 2912.78134 | 2840.059453 | 0.7965 |
| GALNT2 | chr1 | 230185351 | 35 | 32 | 3292.280358 | 3000.287059 | 0.79654 |
| FOXP3 | chrX | 49117261 | 9 | 8 | 883.3333333 | 790 | 0.79664 |
| ESR1 | chr6 | 152011220 | 39 | 32 | 3615.848312 | 2958.412719 | 0.79668 |
| CDKN2B-AS1 | chr9 | 21970434 | 16 | 16 | 506.8193582 | 527.3271783 | 0.7967 |
| LOXL1 | chr15 | 74215830 | 32 | 32 | 3045.89206 | 3059.115093 | 0.79674 |
| BAX | chr19 | 49465143 | 32 | 33 | 2664.492881 | 2776.153848 | 0.79704 |
| PECAM1 | chr17 | 62397542 | 31 | 33 | 2184.855035 | 2296.746455 | 0.79714 |
| APOA1 | chr11 | 116693213 | 34 | 32 | 2928.433474 | 2733.305361 | 0.7972 |
| COL15A1 | chr9 | 101698133 | 34 | 32 | 3029.632912 | 2865.947851 | 0.79722 |
| CDKN2B-AS1 | chr9 | 21992459 | 37 | 33 | 1818.283332 | 1598.26092 | 0.79742 |
| AKT2 | chr19 | 40792012 | 33 | 33 | 2779.331385 | 2755.790292 | 0.79746 |
| NOS3 | chr7 | 150707216 | 28 | 30 | 1971.955874 | 2080.271562 | 0.79762 |
| LOXL1 | chr15 | 74210568 | 33 | 32 | 3123.562191 | 3015.649641 | 0.79768 |
| APOA1 | chr11 | 116704246 | 32 | 32 | 2758.071131 | 2737.445121 | 0.7977 |
| GCK | chr7 | 44228840 | 37 | 32 | 3480.83618 | 3001.466126 | 0.7978 |
| BAX | chr19 | 49457725 | 19 | 20 | 893.7487512 | 973.6904762 | 0.7978 |
| F2RL3 | chr19 | 16990162 | 30 | 32 | 2843.293299 | 3046.239451 | 0.79786 |
| F7 | chr13 | 113751414 | 27 | 27 | 2593.764014 | 2610.727931 | 0.79796 |
| LPA | chr6 | 11219517 | 34 | 32 | 2917.955912 | 2734.716659 | 0.79816 |
| AMOTL2 | chr3 | 134094045 | 27 | 28 | 976.8467355 | 985.1249346 | 0.79856 |
| CDKN1A | chr6 | 36650771 | 15 | 11 | 409.4187841 | 319.4698679 | 0.79904 |
| FOXP3 | chrX | 49117080 | 32 | 32 | 3004.196974 | 3020.660578 | 0.79958 |

| | | | | | | | |
|---|---|---|---|---|---|---|---|
| CDKN1A | chr6 | 36650546 | 19 | 12 | 1125.873016 | 684.9350649 | 0.79974 |
| FGF2 | chr4 | 123743049 | 39 | 32 | 3615.135664 | 2976.25328 | 0.79982 |
| AMOTL2 | chr3 | 134074480 | 27 | 29 | 2039.389908 | 2158.577395 | 0.79986 |
| FOXP3 | chrX | 49127048 | 20 | 13 | 1858.095238 | 1194.880952 | 0.80022 |
| LPA | chr6 | 11218347 | 38 | 32 | 3653.685486 | 3084.411731 | 0.80032 |
| LDLR | chr19 | 11190752 | 36 | 32 | 3397.073448 | 3009.965449 | 0.80038 |
| NOS3 | chr7 | 150707089 | 33 | 32 | 2907.981707 | 2838.433236 | 0.80038 |
| CTGF | chr6 | 132269645 | 34 | 32 | 2826.475238 | 2646.70398 | 0.8006 |
| FGF2 | chr4 | 123739107 | 22 | 22 | 1917.215007 | 1893.203463 | 0.80066 |
| CDKN2B-AS1 | chr9 | 21997078 | 37 | 33 | 1633.518185 | 1432.487759 | 0.80068 |
| ADRA2C | chr4 | 3762500 | 23 | 26 | 1934.197037 | 2158.286436 | 0.8007 |
| APOA1 | chr11 | 116692779 | 37 | 32 | 3337.88586 | 2900.353327 | 0.8009 |
| ABCG1 | chr21 | 43638345 | 35 | 32 | 3326.431763 | 3055.787738 | 0.80114 |
| E2F6 | chr2 | 11615171 | 31 | 29 | 3038.745967 | 2795.075758 | 0.80124 |
| PHACTR1 | chr6 | 12710989 | 34 | 33 | 2547.760372 | 2449.951516 | 0.80136 |
| ADRA2C | chr4 | 3769987 | 22 | 20 | 1367.16256 | 1271.338384 | 0.80144 |
| APOA1 | chr11 | 116660296 | 34 | 32 | 3160.085027 | 2986.506875 | 0.8018 |
| F7 | chr13 | 113750783 | 29 | 29 | 2707.394966 | 2689.739427 | 0.80194 |
| AMOTL2 | chr3 | 134094048 | 29 | 25 | 1011.368417 | 895.7309346 | 0.80198 |
| MIA3 | chr1 | 222809193 | 34 | 32 | 3186.846248 | 3009.272931 | 0.80216 |
| PLA2G7 | chr6 | 46707617 | 35 | 32 | 3210.91392 | 2923.222116 | 0.80254 |
| NOS3 | chr7 | 150711133 | 14 | 16 | 430.5149013 | 467.9609348 | 0.80254 |
| PECAM1 | chr17 | 62409384 | 33 | 29 | 1372.154774 | 1180.240571 | 0.80296 |
| CDKN2B-AS1 | chr9 | 21971179 | 30 | 31 | 1447.113174 | 1462.843 | 0.80318 |
| FGF2 | chr4 | 123744105 | 32 | 33 | 2567.892266 | 2621.586351 | 0.80348 |
| LDLR | chr19 | 11192633 | 24 | 22 | 2222.274531 | 2018.834776 | 0.8038 |
| FGF2 | chr4 | 123747610 | 18 | 23 | 742.992008 | 926.2717838 | 0.80386 |
| MIA3 | chr1 | 222808781 | 39 | 32 | 3688.406683 | 3034.190186 | 0.804 |
| APOA1 | chr11 | 116691776 | 31 | 32 | 2896.985294 | 2972.789966 | 0.80416 |
| LDLR | chr19 | 11197567 | 14 | 13 | 1253.055556 | 1180.47619 | 0.80416 |
| PECAM1 | chr17 | 62399203 | 33 | 31 | 2800.123734 | 2602.906849 | 0.80424 |
| GCK | chr7 | 44237529 | 32 | 32 | 2944.126252 | 2957.838726 | 0.80426 |
| APOA1 | chr11 | 116692027 | 31 | 32 | 2903.472661 | 2983.292263 | 0.8043 |
| CDKN2B-AS1 | chr9 | 21996224 | 22 | 25 | 736.1882079 | 858.6483981 | 0.80432 |
| ATF3 | chr1 | 212742962 | 7 | 2 | 610 | 166.6666667 | 0.80436 |
| ARHGEF18 | chr19 | 7528664 | 8 | 13 | 757.1428571 | 1216.547619 | 0.80496 |
| KLF2 | chr19 | 16437892 | 19 | 22 | 674.9077601 | 758.8901621 | 0.8052 |
| LCK | chr1 | 32707181 | 17 | 13 | 558.9103054 | 447.5054414 | 0.8058 |
| CDKN2B-AS1 | chr9 | 21979075 | 32 | 31 | 1924.192337 | 1833.631777 | 0.80592 |
| CDKN2B-AS1 | chr9 | 21989567 | 21 | 16 | 620.0346754 | 491.4159624 | 0.8062 |
| APOA1 | chr11 | 116690150 | 35 | 32 | 3218.871567 | 2930.719769 | 0.80638 |
| LDLR | chr19 | 11190988 | 34 | 32 | 3253.626879 | 3073.652465 | 0.80638 |
| DHFR | chr5 | 79951537 | 34 | 33 | 2423.401534 | 2376.579328 | 0.8065 |
| TERT | chr5 | 1299404 | 14 | 14 | 1249.117549 | 1262.738095 | 0.80662 |
| GALNT2 | chr1 | 230193853 | 30 | 30 | 2602.233294 | 2623.730047 | 0.80666 |
| CDKN2B-AS1 | chr9 | 22005969 | 12 | 10 | 382.8327228 | 335.4358387 | 0.80678 |
| NOS3 | chr7 | 150707256 | 23 | 24 | 1824.671717 | 1929.966422 | 0.8068 |

| | | | | | | | |
|---|---|---|---|---|---|---|---|
| CTGF | chr6 | 132281608 | 13 | 21 | 374.8365266 | 623.4673797 | 0.80704 |
| APOA1 | chr11 | 116669652 | 39 | 32 | 3773.783887 | 3102.064678 | 0.80708 |
| CDKN1A | chr6 | 36644997 | 35 | 31 | 2855.228597 | 2545.928994 | 0.80734 |
| F2RL3 | chr19 | 16994229 | 24 | 29 | 1616.530205 | 2000.374628 | 0.8074 |
| DHFR | chr5 | 79955843 | 32 | 30 | 3091.855187 | 2908.172861 | 0.80754 |
| F2RL3 | chr19 | 17003229 | 6 | 5 | 434.6428571 | 340 | 0.80764 |
| BAX | chr19 | 49454937 | 29 | 31 | 2560.282145 | 2760.112583 | 0.80814 |
| FOXP3 | chrX | 49115798 | 15 | 15 | 1171.850649 | 1149.246032 | 0.80898 |
| APOA1 | chr11 | 116714708 | 21 | 23 | 2039.464286 | 2246.284965 | 0.80916 |
| FOXP3 | chrX | 49117093 | 33 | 32 | 3135.268287 | 3056.435784 | 0.80928 |
| F7 | chr13 | 113753802 | 35 | 32 | 3229.110572 | 2942.570889 | 0.80936 |
| LPA | chr6 | 11220567 | 34 | 32 | 3165.275565 | 2991.107443 | 0.80966 |
| BCL2 | chr18 | 60994360 | 32 | 28 | 2799.180428 | 2469.289599 | 0.81016 |
| ABCA1 | chr9 | 107690847 | 11 | 3 | 504.9206349 | 128.5714286 | 0.81028 |
| ALOX15 | chr17 | 4533869 | 31 | 26 | 2990.133478 | 2494.932845 | 0.81046 |
| F7 | chr13 | 113755487 | 21 | 26 | 1544.484127 | 1949.285714 | 0.81062 |
| F7 | chr13 | 113756107 | 34 | 33 | 2326.930142 | 2234.861855 | 0.81068 |
| LDLR | chr19 | 11190095 | 17 | 23 | 1660.238095 | 2255.277778 | 0.8108 |
| BAX | chr19 | 49465332 | 33 | 33 | 2349.007965 | 2377.655468 | 0.81082 |
| AKT2 | chr19 | 40736309 | 31 | 33 | 1937.663907 | 2085.511868 | 0.81084 |
| CDKN2B-AS1 | chr9 | 21966540 | 30 | 33 | 1599.643209 | 1735.937486 | 0.81128 |
| APOA1 | chr11 | 116715935 | 16 | 10 | 1483.216783 | 937.1428571 | 0.81136 |
| LCK | chr1 | 32716378 | 33 | 31 | 1809.834835 | 1726.703768 | 0.81178 |
| APOA1 | chr11 | 116708225 | 16 | 19 | 1462.09596 | 1718.061661 | 0.81188 |
| F7 | chr13 | 113759852 | 32 | 28 | 2363.709208 | 2043.553977 | 0.81204 |
| F7 | chr13 | 113750638 | 25 | 24 | 2363.134921 | 2281.746032 | 0.81208 |
| FOXP3 | chrX | 49120474 | 12 | 15 | 1147.532468 | 1424.040404 | 0.81248 |
| APOA1 | chr11 | 116692675 | 31 | 32 | 2799.963176 | 2907.112067 | 0.81274 |
| LOXL1 | chr15 | 74214301 | 31 | 31 | 2430.660357 | 2408.115006 | 0.81292 |
| LOXL1 | chr15 | 74215888 | 32 | 33 | 2008.706555 | 2100.745328 | 0.81312 |
| ABCB1 | chr7 | 87133567 | 37 | 32 | 3518.566144 | 3052.163096 | 0.81314 |
| TIMP3 | chr22 | 33196425 | 34 | 32 | 2863.730716 | 2709.084113 | 0.81344 |
| FOXP3 | chrX | 49121418 | 15 | 13 | 613.0555556 | 512.51443 | 0.8136 |
| LCK | chr1 | 32742074 | 21 | 20 | 1871.63961 | 1798.722944 | 0.81374 |
| CDKN2B-AS1 | chr9 | 21985398 | 40 | 33 | 1973.665607 | 1608.751893 | 0.81412 |
| P2RY12 | chr3 | 151107004 | 37 | 32 | 2697.305391 | 2311.102925 | 0.81418 |
| APOA1 | chr11 | 116720136 | 27 | 28 | 2578.187229 | 2685.219421 | 0.81422 |
| APOA1 | chr11 | 116685127 | 33 | 32 | 2584.30717 | 2484.605281 | 0.8143 |
| F7 | chr13 | 113753716 | 36 | 32 | 3344.708125 | 2961.995136 | 0.8144 |
| ICAM2 | chr17 | 62086278 | 27 | 32 | 1184.493939 | 1377.286575 | 0.8147 |
| APOA1 | chr11 | 116698334 | 4 | 4 | 157.1428571 | 166.6666667 | 0.81492 |
| CDKN2B-AS1 | chr9 | 21989616 | 15 | 10 | 415.9372259 | 293.4085186 | 0.81508 |
| PHACTR1 | chr6 | 12711015 | 34 | 32 | 3091.866405 | 2926.325259 | 0.81532 |
| CDKN1A | chr6 | 36651875 | 31 | 32 | 2993.606006 | 3081.772492 | 0.81542 |
| AMOTL2 | chr3 | 134103881 | 15 | 10 | 1224.722222 | 797.7777778 | 0.81552 |
| LDLR | chr19 | 11195034 | 34 | 31 | 3214.57396 | 2914.197332 | 0.81556 |
| APOA1 | chr11 | 116689790 | 34 | 32 | 2341.399847 | 2181.123213 | 0.81572 |

| | | | | | | | |
|---|---|---|---|---|---|---|---|
| TMPO | chr12 | 98908148 | 36 | 33 | 2845.64594 | 2590.87585 | 0.81592 |
| ADRA2C | chr4 | 3763692 | 32 | 32 | 2943.889188 | 2957.602675 | 0.816 |
| APOA1 | chr11 | 116711189 | 11 | 14 | 312.0070651 | 413.7932295 | 0.816 |
| ESR1 | chr6 | 152010253 | 14 | 12 | 703.4108323 | 581.750817 | 0.81618 |
| NOS3 | chr7 | 150709877 | 37 | 32 | 3520.751875 | 3054.977763 | 0.81632 |
| CDKN2B-AS1 | chr9 | 21991452 | 36 | 33 | 2505.730747 | 2316.858508 | 0.81644 |
| PHACTR1 | chr6 | 12713899 | 32 | 32 | 2851.610726 | 2829.159313 | 0.8169 |
| NOS3 | chr7 | 150711010 | 4 | 5 | 142.2222222 | 200.7142857 | 0.81716 |
| P2RY12 | chr3 | 151107869 | 34 | 32 | 2547.958765 | 2419.659303 | 0.8172 |
| ABCG1 | chr21 | 43630172 | 29 | 22 | 1149.282105 | 850.735406 | 0.81732 |
| ABCA1 | chr9 | 107691551 | 3 | 5 | 97.27272727 | 181.9775542 | 0.81746 |
| ESR1 | chr6 | 152006061 | 33 | 32 | 3126.983425 | 3041.132271 | 0.8179 |
| ADRA2C | chr4 | 3766395 | 33 | 31 | 2485.949544 | 2314.071135 | 0.818 |
| FGF2 | chr4 | 123744235 | 17 | 21 | 1465.032468 | 1786.865079 | 0.81808 |
| BAX | chr19 | 49456623 | 32 | 32 | 2462.064003 | 2435.968509 | 0.81812 |
| APOA1 | chr11 | 116690830 | 32 | 32 | 2756.256693 | 2775.7901 | 0.81818 |
| ABCG1 | chr21 | 43638596 | 16 | 14 | 1356.904762 | 1168.003108 | 0.8183 |
| ABCA1 | chr9 | 107696063 | 14 | 17 | 546.1471861 | 684.2460317 | 0.81854 |
| PHACTR1 | chr6 | 12717138 | 27 | 31 | 2592.874544 | 2964.71604 | 0.8188 |
| F2RL3 | chr19 | 16991414 | 31 | 32 | 2960.073198 | 3071.03171 | 0.8192 |
| ABCA1 | chr9 | 107690743 | 13 | 13 | 524.484127 | 545.487013 | 0.81938 |
| FOXP3 | chrX | 49127250 | 19 | 24 | 1795.716783 | 2282.81746 | 0.81976 |
| ATF3 | chr1 | 212779858 | 8 | 9 | 287.4814075 | 305.3174603 | 0.81984 |
| F2RL3 | chr19 | 16990995 | 23 | 23 | 2118.448773 | 2102.61544 | 0.8199 |
| APOA1 | chr11 | 116691844 | 32 | 32 | 2967.982122 | 2980.41051 | 0.82012 |
| APOA1 | chr11 | 116692719 | 33 | 32 | 2997.636533 | 2917.5093 | 0.82032 |
| GCK | chr7 | 44232208 | 33 | 31 | 3184.776524 | 3001.953117 | 0.82048 |
| ICAM2 | chr17 | 62086423 | 23 | 24 | 763.3506526 | 814.5855752 | 0.82054 |
| APOA1 | chr11 | 116674229 | 29 | 31 | 2713.754995 | 2913.081934 | 0.82058 |
| PLA2G7 | chr6 | 46706324 | 10 | 13 | 817.9365079 | 1038.611111 | 0.82084 |
| AMOTL2 | chr3 | 134096536 | 21 | 27 | 1618.808151 | 2049.931319 | 0.821 |
| ADRA2C | chr4 | 3762224 | 32 | 32 | 2960.439773 | 2972.784775 | 0.82104 |
| TERT | chr5 | 1295738 | 22 | 21 | 735.6111002 | 685.1919549 | 0.82118 |
| CDKN2B-AS1 | chr9 | 21968132 | 29 | 31 | 930.9119045 | 975.9211709 | 0.82132 |
| AMOTL2 | chr3 | 134101221 | 23 | 28 | 1426.605697 | 1777.090507 | 0.82132 |
| TERT | chr5 | 1299390 | 10 | 11 | 911.3852814 | 989.2063492 | 0.82132 |
| F2RL3 | chr19 | 16990451 | 29 | 30 | 2824.042556 | 2911.550117 | 0.82166 |
| FOXP3 | chrX | 49116290 | 30 | 31 | 2795.431887 | 2873.78423 | 0.82168 |
| MCPH1 | chr8 | 6470942 | 15 | 23 | 1039.199346 | 1555.357143 | 0.82168 |
| ARHGEF18 | chr19 | 7468107 | 30 | 29 | 2777.513845 | 2698.03276 | 0.82176 |
| CTGF | chr6 | 132272744 | 4 | 4 | 130 | 121.9047619 | 0.82176 |
| APOA1 | chr11 | 116705997 | 8 | 10 | 216.6666667 | 282.4603175 | 0.82178 |
| ESR1 | chr6 | 152424275 | 36 | 32 | 3448.914879 | 3059.171181 | 0.82208 |
| APOA1 | chr11 | 116707212 | 27 | 28 | 1145.322782 | 1164.854519 | 0.82232 |
| LDLR | chr19 | 11217629 | 10 | 9 | 902.5518926 | 820.4761905 | 0.82238 |
| CDKN2B-AS1 | chr9 | 21996217 | 21 | 20 | 722.6124022 | 707.458207 | 0.82242 |
| BAX | chr19 | 49455024 | 35 | 32 | 3128.532922 | 2871.245207 | 0.82282 |

| | | | | | | | |
|---|---|---|---|---|---|---|---|
| FOXP3 | chrX | 49116238 | 30 | 29 | 2803.336936 | 2720.694877 | 0.8229 |
| F2RL3 | chr19 | 16998247 | 24 | 16 | 972.105665 | 631.9552669 | 0.82296 |
| GCK | chr7 | 44230868 | 33 | 32 | 2751.784423 | 2652.679643 | 0.82354 |
| CTGF | chr6 | 132282312 | 19 | 15 | 1035.766101 | 791.6704973 | 0.82356 |
| CDKN2B-AS1 | chr9 | 22008003 | 32 | 31 | 1275.404433 | 1256.402821 | 0.82372 |
| CDKN2B-AS1 | chr9 | 21969256 | 23 | 28 | 845.271743 | 1008.505654 | 0.82426 |
| FOXP3 | chrX | 49127105 | 20 | 17 | 1835.468975 | 1570.833333 | 0.82446 |
| GCK | chr7 | 44228569 | 28 | 28 | 2202.332251 | 2175.115775 | 0.82454 |
| F7 | chr13 | 113750613 | 26 | 26 | 2443.549784 | 2455.138571 | 0.82456 |
| ADRA2C | chr4 | 3759515 | 32 | 32 | 3006.220892 | 3017.989796 | 0.8246 |
| APOA1 | chr11 | 116701548 | 32 | 29 | 2991.894407 | 2726.763285 | 0.82478 |
| P2RY12 | chr3 | 151111795 | 37 | 32 | 3361.741831 | 2919.463943 | 0.82478 |
| NOS3 | chr7 | 150688483 | 34 | 32 | 3108.702213 | 2913.345747 | 0.82492 |
| F7 | chr13 | 113756117 | 34 | 32 | 2840.887373 | 2657.487812 | 0.82496 |
| BAX | chr19 | 49455150 | 33 | 32 | 2741.330356 | 2641.219831 | 0.82504 |
| FOXP3 | chrX | 49120452 | 19 | 16 | 1756.315629 | 1493.192918 | 0.82506 |
| AMOTL2 | chr3 | 134094055 | 32 | 33 | 1400.640945 | 1421.602002 | 0.82508 |
| PHACTR1 | chr6 | 12717711 | 35 | 33 | 2634.316928 | 2505.275372 | 0.82514 |
| CDKN1A | chr6 | 36653148 | 34 | 33 | 2290.092969 | 2206.287257 | 0.82548 |
| LDLR | chr19 | 11219567 | 35 | 33 | 2890.707141 | 2705.447641 | 0.8256 |
| APOA1 | chr11 | 116689549 | 3 | 4 | 280 | 365.7142857 | 0.8256 |
| LOXL1 | chr15 | 74218983 | 24 | 23 | 794.9698995 | 776.8850357 | 0.82566 |
| APOA1 | chr11 | 116695819 | 23 | 25 | 2069.539309 | 2266.337274 | 0.8257 |
| MIA3 | chr1 | 222816708 | 27 | 29 | 2445.706654 | 2610.395993 | 0.82574 |
| LPA | chr6 | 11224503 | 25 | 31 | 1812.265512 | 2274.462759 | 0.82596 |
| NOS3 | chr7 | 150710485 | 28 | 31 | 2028.542967 | 2271.969615 | 0.82608 |
| CDKN2B-AS1 | chr9 | 21987353 | 31 | 31 | 1220.1794 | 1199.244714 | 0.82626 |
| CTGF | chr6 | 132276106 | 33 | 32 | 2776.540321 | 2674.041384 | 0.82632 |
| CDKN2B-AS1 | chr9 | 21989200 | 36 | 32 | 1376.768611 | 1205.93221 | 0.82648 |
| ESR1 | chr6 | 152002693 | 31 | 32 | 1314.232892 | 1336.124525 | 0.82652 |
| LPA | chr6 | 160944778 | 35 | 32 | 3246.008119 | 2974.805251 | 0.82656 |
| APOA1 | chr11 | 116680489 | 34 | 32 | 3225.128083 | 3026.057656 | 0.82658 |
| CTGF | chr6 | 132275999 | 33 | 32 | 2963.225184 | 2859.913034 | 0.8268 |
| NOS3 | chr7 | 150711004 | 3 | 3 | 113.3333333 | 131.1111111 | 0.8275 |
| ALOX15 | chr17 | 4550074 | 31 | 32 | 2319.25244 | 2431.162303 | 0.82762 |
| APOA1 | chr11 | 116708590 | 27 | 26 | 2569.928658 | 2460.795342 | 0.8278 |
| ESR2 | chr14 | 64806970 | 4 | 6 | 385.7142857 | 572.2222222 | 0.82782 |
| LDLR | chr19 | 11218332 | 15 | 20 | 1387.129913 | 1829.996392 | 0.82822 |
| LCK | chr1 | 32741207 | 10 | 11 | 381.3095238 | 400.9704185 | 0.8284 |
| FOXP3 | chrX | 49118314 | 30 | 32 | 2836.999365 | 3012.702869 | 0.82852 |
| LDLR | chr19 | 11192830 | 4 | 8 | 203.5714286 | 375.2777778 | 0.82858 |
| CDKN2B-AS1 | chr9 | 21986063 | 38 | 33 | 2979.762342 | 2570.399085 | 0.82878 |
| NOS3 | chr7 | 150710447 | 22 | 25 | 811.0025482 | 942.2566322 | 0.82886 |
| TERT | chr5 | 1296077 | 30 | 32 | 1944.651763 | 2046.410885 | 0.82902 |
| APOA1 | chr11 | 116692241 | 34 | 32 | 3197.480444 | 3017.626316 | 0.82918 |
| LDLR | chr19 | 11221096 | 20 | 18 | 1894.047619 | 1719.365079 | 0.82944 |
| LCK | chr1 | 32741020 | 25 | 32 | 1280.312748 | 1610.821114 | 0.82952 |

| | | | | | | | |
|---|---|---|---|---|---|---|---|
| ESR1 | chr6 | 152005559 | 35 | 32 | 3384.944143 | 3089.385056 | 0.82964 |
| ALOX15 | chr17 | 4534096 | 31 | 30 | 2776.794244 | 2709.078242 | 0.82968 |
| CDKN2B-AS1 | chr9 | 21993154 | 37 | 32 | 3501.739008 | 3035.55195 | 0.83008 |
| LPA | chr6 | 160948527 | 35 | 32 | 2975.829013 | 2732.713689 | 0.83026 |
| GCK | chr7 | 44229360 | 13 | 16 | 1040.714286 | 1306.547619 | 0.83034 |
| APOA1 | chr11 | 116701354 | 31 | 32 | 2938.33441 | 3022.452686 | 0.83054 |
| F2RL3 | chr19 | 16992375 | 22 | 26 | 1966.020924 | 2298.857809 | 0.83108 |
| APOA1 | chr11 | 116711083 | 35 | 33 | 2553.095969 | 2427.756239 | 0.83132 |
| COL15A1 | chr9 | 101832878 | 28 | 21 | 2465.703463 | 1866.738817 | 0.83162 |
| LDLR | chr19 | 11221410 | 33 | 32 | 3154.940029 | 3069.996115 | 0.83166 |
| NOS3 | chr7 | 150692198 | 23 | 24 | 777.3035577 | 828.4668692 | 0.83176 |
| APOA1 | chr11 | 116676108 | 38 | 32 | 3189.482177 | 2697.784401 | 0.83182 |
| APOA1 | chr11 | 116701337 | 32 | 32 | 3068.417277 | 3078.591036 | 0.83192 |
| BCL2 | chr18 | 60995100 | 39 | 32 | 3761.64541 | 3080.23272 | 0.832 |
| E2F6 | chr2 | 11611980 | 34 | 33 | 3123.418424 | 3051.483385 | 0.8323 |
| APOA1 | chr11 | 116695362 | 28 | 27 | 2680.749251 | 2595.194805 | 0.83254 |
| ADRA2C | chr4 | 3766840 | 19 | 18 | 1697.5 | 1586.904762 | 0.83268 |
| CDKN1A | chr6 | 36653545 | 10 | 7 | 381.1904762 | 251.9047619 | 0.83288 |
| APOA1 | chr11 | 116661679 | 29 | 30 | 1887.346265 | 1979.726779 | 0.83306 |
| ABCA1 | chr9 | 107690932 | 26 | 27 | 2265.368814 | 2370.859973 | 0.83316 |
| APOA1 | chr11 | 116693095 | 26 | 26 | 1447.040575 | 1481.330915 | 0.83326 |
| ABCA1 | chr9 | 107690798 | 10 | 16 | 456.7712843 | 751.8650794 | 0.8334 |
| CTGF | chr6 | 132280361 | 28 | 28 | 2394.113537 | 2373.020225 | 0.8335 |
| AKT2 | chr19 | 40792993 | 36 | 33 | 3024.968953 | 2756.13136 | 0.8335 |
| FOXP3 | chrX | 49117118 | 33 | 33 | 2704.668589 | 2726.978486 | 0.8337 |
| P2RY12 | chr3 | 151103697 | 32 | 32 | 1315.476649 | 1289.636013 | 0.83378 |
| F7 | chr13 | 113750672 | 28 | 27 | 2485.721338 | 2412.707431 | 0.83388 |
| APOA1 | chr11 | 116696399 | 35 | 33 | 2658.532816 | 2488.327319 | 0.83448 |
| TIMP3 | chr22 | 33193849 | 32 | 33 | 1595.665736 | 1620.205902 | 0.83462 |
| PLA2G7 | chr6 | 46713364 | 39 | 32 | 3160.943459 | 2606.363978 | 0.83494 |
| APOA1 | chr11 | 116678783 | 28 | 29 | 2642.595974 | 2724.412918 | 0.835 |
| COL15A1 | chr9 | 101699464 | 33 | 32 | 2881.193029 | 2780.584798 | 0.83576 |
| LCK | chr1 | 32714623 | 15 | 20 | 466.1743485 | 606.6333009 | 0.8358 |
| F7 | chr13 | 113751000 | 29 | 23 | 2700.4329 | 2161.925411 | 0.83592 |
| CDKN2B-AS1 | chr9 | 21968488 | 8 | 12 | 260 | 403.7079587 | 0.83598 |
| TERT | chr5 | 1253761 | 22 | 26 | 1995.344198 | 2382.993395 | 0.83602 |
| CDKN2B-AS1 | chr9 | 21970767 | 32 | 30 | 1263.807206 | 1162.038031 | 0.83612 |
| ABCA1 | chr9 | 107693185 | 33 | 32 | 3023.286585 | 2918.103123 | 0.83668 |
| APOA1 | chr11 | 116707817 | 27 | 25 | 2501.945669 | 2287.410192 | 0.83668 |
| PHACTR1 | chr6 | 12712894 | 34 | 33 | 2394.577181 | 2347.974234 | 0.83682 |
| E2F6 | chr2 | 11613152 | 3 | 3 | 194.1558442 | 183.3333333 | 0.83708 |
| FGF2 | chr4 | 123745760 | 38 | 32 | 3448.651784 | 2913.577715 | 0.8372 |
| E2F6 | chr2 | 11614602 | 35 | 32 | 3078.515867 | 2824.668167 | 0.83772 |
| ESR1 | chr6 | 152007627 | 2 | 2 | 180 | 185.7142857 | 0.83778 |
| FOXP3 | chrX | 49119237 | 29 | 30 | 2095.536112 | 2138.020846 | 0.83794 |
| LDLR | chr19 | 11222420 | 28 | 31 | 2600.552813 | 2864.732937 | 0.83796 |
| APOA1 | chr11 | 116688150 | 35 | 32 | 3261.217054 | 2970.109063 | 0.83802 |

| | | | | | | | |
|---|---|---|---|---|---|---|---|
| LCK | chr1 | 32740731 | 22 | 15 | 2131.428571 | 1446.428571 | 0.8382 |
| CDKN1A | chr6 | 36653477 | 6 | 7 | 264.7619048 | 296.6666667 | 0.83824 |
| APOA1 | chr11 | 116669302 | 16 | 12 | 1090.119048 | 833.3333333 | 0.83836 |
| APOA1 | chr11 | 116723235 | 27 | 30 | 907.5092928 | 991.25417 | 0.83858 |
| F7 | chr13 | 113775393 | 35 | 32 | 2965.052352 | 2693.058025 | 0.83902 |
| KLF2 | chr19 | 16430448 | 2 | 4 | 66.17647059 | 153.5784905 | 0.8393 |
| F7 | chr13 | 113750786 | 30 | 29 | 2697.984099 | 2624.358142 | 0.83948 |
| NOS3 | chr7 | 150707964 | 6 | 9 | 269.0476191 | 422.1428571 | 0.83966 |
| ABCG1 | chr21 | 43630973 | 25 | 23 | 2388.167388 | 2213.333333 | 0.83968 |
| ARHGEF18 | chr19 | 7461161 | 10 | 14 | 697.5649351 | 1003.650794 | 0.83998 |
| TMPO | chr12 | 98944103 | 2 | 3 | 138.0995475 | 186.2717471 | 0.84006 |
| APOA1 | chr11 | 116661629 | 30 | 30 | 2025.241915 | 2052.582695 | 0.84038 |
| APOA1 | chr11 | 116684082 | 34 | 33 | 2996.273319 | 2881.081821 | 0.84038 |
| CDKN2B-AS1 | chr9 | 21993654 | 14 | 13 | 501.1616161 | 451.8833944 | 0.8404 |
| TMPO | chr12 | 98903357 | 34 | 31 | 3326.041477 | 3025.700459 | 0.84072 |
| ATF3 | chr1 | 212741827 | 31 | 32 | 2765.64003 | 2834.664894 | 0.84092 |
| MCPH1 | chr8 | 6258265 | 34 | 32 | 3265.329182 | 3064.204678 | 0.84094 |
| ADRA2C | chr4 | 3764165 | 33 | 33 | 2652.246905 | 2634.236851 | 0.84118 |
| LCK | chr1 | 32740738 | 22 | 18 | 2141.904762 | 1741.428571 | 0.84124 |
| FOXP3 | chrX | 49126909 | 14 | 10 | 612.3593074 | 421.1904762 | 0.8415 |
| LOXL1 | chr15 | 74217732 | 32 | 32 | 3037.959401 | 3030.03103 | 0.8418 |
| GCK | chr7 | 44233568 | 31 | 32 | 2534.576461 | 2595.066369 | 0.8419 |
| APOA1 | chr11 | 116685912 | 8 | 7 | 785.7142857 | 683.3333333 | 0.84216 |
| KLF2 | chr19 | 16437889 | 25 | 25 | 1037.490187 | 1057.032853 | 0.84234 |
| CDKN1A | chr6 | 36653159 | 34 | 33 | 2107.762398 | 2062.692809 | 0.84238 |
| ABCG1 | chr21 | 43637289 | 34 | 32 | 3289.024468 | 3087.807642 | 0.84248 |
| E2F6 | chr2 | 11612970 | 34 | 32 | 3270.120419 | 3070.845685 | 0.84252 |
| LDLR | chr19 | 11216626 | 18 | 23 | 1635.736624 | 2073.802309 | 0.84284 |
| P2RY12 | chr3 | 151110931 | 5 | 6 | 233.2142857 | 301.4285714 | 0.843 |
| CDKN1A | chr6 | 36651889 | 31 | 32 | 2909.522603 | 2993.960067 | 0.84304 |
| E2F6 | chr2 | 11613936 | 16 | 14 | 1530 | 1346.666667 | 0.8433 |
| GCK | chr7 | 44229543 | 32 | 32 | 3074.551787 | 3067.226573 | 0.84354 |
| BCL2 | chr18 | 60990594 | 32 | 33 | 2189.010574 | 2276.44384 | 0.84356 |
| NOS3 | chr7 | 150710084 | 23 | 28 | 1005.278997 | 1199.578575 | 0.84362 |
| F2RL3 | chr19 | 16997255 | 17 | 15 | 1550.833333 | 1380.793651 | 0.84376 |
| LOXL1 | chr15 | 74218217 | 28 | 33 | 2635.4928 | 3086.689155 | 0.84378 |
| F7 | chr13 | 113775325 | 33 | 31 | 3006.591394 | 2833.766331 | 0.84426 |
| NOS3 | chr7 | 150695652 | 22 | 21 | 2120.039683 | 2031.547619 | 0.84442 |
| CDKN2BAS | chr9 | 22098618 | 28 | 30 | 1802.253726 | 1973.495528 | 0.84446 |
| LPA | chr6 | 160942848 | 38 | 32 | 3488.790201 | 2927.499611 | 0.84468 |
| ESR1 | chr6 | 152423971 | 33 | 32 | 3044.060784 | 2937.614867 | 0.845 |
| ARHGEF18 | chr19 | 7461683 | 8 | 7 | 750 | 645.7142857 | 0.84518 |
| LOXL1 | chr15 | 74216694 | 34 | 33 | 2676.718483 | 2615.62941 | 0.8452 |
| BAX | chr19 | 49456858 | 27 | 26 | 2458.348041 | 2384.371184 | 0.84542 |
| CDKN2B-AS1 | chr9 | 21969847 | 13 | 14 | 452.7489177 | 470.5663781 | 0.84542 |
| LDLR | chr19 | 11222212 | 3 | 4 | 280 | 380 | 0.84568 |
| LDLR | chr19 | 11190252 | 3 | 4 | 260 | 340 | 0.84568 |

| | | | | | | | |
|---|---|---|---|---|---|---|---|
| ALDOA | chr16 | 30061639 | 4 | 2 | 349.2307692 | 180 | 0.84614 |
| BCL2 | chr18 | 60989305 | 26 | 28 | 2257.893218 | 2407.240815 | 0.84626 |
| CDKN2B-AS1 | chr9 | 21989066 | 27 | 28 | 873.0686079 | 888.8450767 | 0.84648 |
| TIMP3 | chr22 | 33187289 | 34 | 32 | 3278.794384 | 3094.209746 | 0.84708 |
| NR3C1 | chr5 | 142791017 | 28 | 30 | 2593.923631 | 2759.652847 | 0.84718 |
| FOXP3 | chrX | 49131083 | 33 | 32 | 2907.922332 | 2838.067998 | 0.84726 |
| CDKN2B-AS1 | chr9 | 21989777 | 10 | 11 | 343.2189542 | 365.0421508 | 0.8473 |
| CDKN2B-AS1 | chr9 | 21990539 | 14 | 15 | 468.7700345 | 483.7048298 | 0.8483 |
| LPA | chr6 | 160948170 | 29 | 27 | 2512.197919 | 2354.089799 | 0.84834 |
| F7 | chr13 | 113755472 | 21 | 31 | 1679.764957 | 2505.768398 | 0.84848 |
| NOS3 | chr7 | 150711461 | 5 | 2 | 154.981685 | 58.33333333 | 0.84848 |
| LPA | chr6 | 11223277 | 32 | 31 | 1915.631773 | 1879.661123 | 0.84904 |
| APOA1 | chr11 | 116675246 | 22 | 21 | 2123.333333 | 2036.547619 | 0.8493 |
| CDKN2BAS | chr9 | 22067554 | 15 | 13 | 846.5869362 | 757.3252922 | 0.84962 |
| NOS3 | chr7 | 150683812 | 33 | 33 | 2473.874171 | 2497.682746 | 0.84968 |
| ABCA1 | chr9 | 107691065 | 31 | 32 | 2908.802882 | 3013.399815 | 0.85004 |
| NOS3 | chr7 | 150706119 | 33 | 33 | 2189.860355 | 2213.033752 | 0.85008 |
| AMOTL2 | chr3 | 134101391 | 32 | 32 | 2870.187405 | 2880.83228 | 0.8501 |
| HMGCR | chr5 | 74623833 | 11 | 17 | 1038.928571 | 1590 | 0.8501 |
| PLA2G7 | chr6 | 46708190 | 36 | 32 | 3404.799029 | 3018.494264 | 0.85022 |
| F7 | chr13 | 113751097 | 9 | 8 | 885.7142857 | 783.3333333 | 0.8504 |
| APOA1 | chr11 | 116698431 | 18 | 20 | 1695.521978 | 1872.380952 | 0.85052 |
| BCL2 | chr18 | 60993408 | 36 | 32 | 3250.115996 | 2881.223273 | 0.85062 |
| E2F6 | chr2 | 11615048 | 34 | 32 | 3269.609734 | 3068.178963 | 0.85084 |
| LDLR | chr19 | 11219549 | 34 | 32 | 3211.135089 | 3029.798359 | 0.85102 |
| ABCG1 | chr21 | 43717701 | 37 | 32 | 3373.772523 | 2909.493899 | 0.85124 |
| APOA1 | chr11 | 116692162 | 33 | 32 | 3165.604669 | 3075.650865 | 0.85188 |
| CDKN2B-AS1 | chr9 | 21996078 | 5 | 6 | 160 | 201.9047619 | 0.85196 |
| CDKN2B-AS1 | chr9 | 21990465 | 26 | 27 | 862.4688333 | 910.6882455 | 0.85204 |
| CDKN2B-AS1 | chr9 | 21989853 | 19 | 21 | 966.5444359 | 1042.712843 | 0.85204 |
| LOXL1 | chr15 | 74214755 | 20 | 19 | 1358.917749 | 1267.049062 | 0.85212 |
| ATF3 | chr1 | 212745333 | 24 | 29 | 2309.257044 | 2776.903652 | 0.85226 |
| FOXP3 | chrX | 49126944 | 11 | 9 | 420.8333333 | 333.015873 | 0.85236 |
| ABCG1 | chr21 | 43636356 | 24 | 24 | 1967.450344 | 1984.163059 | 0.85238 |
| LOXL1 | chr15 | 74210944 | 34 | 32 | 3175.457212 | 2979.819783 | 0.85252 |
| GALNT2 | chr1 | 230193672 | 25 | 26 | 1510.286103 | 1595.402979 | 0.85258 |
| DHFR | chr5 | 79953683 | 17 | 13 | 998.6071075 | 744.5041193 | 0.85258 |
| GCK | chr7 | 44231779 | 30 | 30 | 1679.270604 | 1706.649214 | 0.85324 |
| DHFR | chr5 | 79952008 | 37 | 32 | 3517.084179 | 3034.577055 | 0.85348 |
| ALDOA | chr16 | 30057291 | 6 | 7 | 506.6666667 | 577.7777778 | 0.85372 |
| APOA1 | chr11 | 116707862 | 14 | 13 | 1333.736264 | 1230 | 0.85388 |
| CDKN2BAS | chr9 | 22096028 | 32 | 33 | 2464.398669 | 2558.820642 | 0.85444 |
| CTGF | chr6 | 132276060 | 33 | 32 | 3104.063047 | 3021.284841 | 0.85478 |
| FGF2 | chr4 | 123744444 | 10 | 7 | 927.5 | 640 | 0.85492 |
| AMOTL2 | chr3 | 134095971 | 30 | 32 | 2615.874084 | 2776.53193 | 0.85508 |
| F7 | chr13 | 113752835 | 32 | 33 | 2401.663381 | 2496.109762 | 0.85556 |
| DHFR | chr5 | 79955632 | 31 | 30 | 2883.835602 | 2774.527099 | 0.8556 |

| | | | | | | | |
|---|---|---|---|---|---|---|---|
| FOXP3 | chrX | 49127031 | 18 | 10 | 1734.166667 | 958.3333333 | 0.85584 |
| APOA1 | chr11 | 116680384 | 32 | 33 | 2842.546912 | 2914.217887 | 0.85628 |
| APOA1 | chr11 | 116688381 | 34 | 33 | 2629.342598 | 2570.544415 | 0.85644 |
| CDKN2B-AS1 | chr9 | 21989835 | 21 | 23 | 891.998557 | 995.5555556 | 0.8565 |
| BAX | chr19 | 49453859 | 3 | 9 | 283.3333333 | 860 | 0.85656 |
| MCPH1 | chr8 | 6470227 | 15 | 16 | 1463.611111 | 1543.055556 | 0.85668 |
| ABCG1 | chr21 | 43636928 | 33 | 32 | 3123.267808 | 3036.801433 | 0.85684 |
| TERT | chr5 | 1299708 | 24 | 25 | 2063.596959 | 2133.707959 | 0.85686 |
| ADRA2C | chr4 | 3759246 | 34 | 32 | 3000.883678 | 2812.151772 | 0.85702 |
| CDKN2B-AS1 | chr9 | 22009670 | 25 | 21 | 701.2471139 | 602.1910419 | 0.85718 |
| APOA1 | chr11 | 116669257 | 5 | 3 | 256.6666667 | 150 | 0.85744 |
| LPA | chr6 | 11220701 | 37 | 32 | 3522.21859 | 3039.956529 | 0.85826 |
| APOA1 | chr11 | 116722761 | 14 | 10 | 433.981544 | 322.8236597 | 0.85826 |
| LDLR | chr19 | 11192269 | 19 | 17 | 1847.619048 | 1642.5 | 0.85848 |
| LCK | chr1 | 32740880 | 19 | 26 | 1129.867311 | 1522.215007 | 0.8585 |
| NR3C1 | chr5 | 142789017 | 37 | 32 | 3427.422164 | 2969.923691 | 0.85868 |
| TERT | chr5 | 1295591 | 11 | 8 | 334.6031746 | 250.5555556 | 0.85872 |
| LDLR | chr19 | 11192066 | 13 | 11 | 1270.833333 | 1080 | 0.8588 |
| ALDOA | chr16 | 30057480 | 5 | 5 | 337.1428571 | 321.4285714 | 0.85902 |
| APOA1 | chr11 | 116680386 | 32 | 32 | 3014.974203 | 3005.181789 | 0.85924 |
| BCL2 | chr18 | 60994559 | 38 | 32 | 3634.789143 | 3071.79592 | 0.85964 |
| CDKN2B-AS1 | chr9 | 21996698 | 25 | 26 | 1106.757502 | 1170.946872 | 0.85986 |
| FOXP3 | chrX | 49118531 | 2 | 2 | 160 | 166.6666667 | 0.85998 |
| LDLR | chr19 | 11194134 | 6 | 9 | 570.9090909 | 863.3333333 | 0.86022 |
| GCK | chr7 | 44233467 | 32 | 32 | 2733.177314 | 2759.053014 | 0.8603 |
| APOA1 | chr11 | 116701851 | 26 | 29 | 1768.153069 | 1929.611507 | 0.8605 |
| APOA1 | chr11 | 116701165 | 17 | 16 | 1579.615385 | 1496.547619 | 0.86092 |
| COL15A1 | chr9 | 101705978 | 4 | 3 | 141.9047619 | 100 | 0.86108 |
| GALNT2 | chr1 | 230193558 | 31 | 33 | 1851.543333 | 1989.553064 | 0.86112 |
| NOS3 | chr7 | 150695777 | 28 | 29 | 2505.357143 | 2582.0227 | 0.8612 |
| CDKN2B-AS1 | chr9 | 21968861 | 35 | 29 | 1059.18884 | 865.9770225 | 0.86144 |
| NOS3 | chr7 | 150705948 | 26 | 28 | 1004.74122 | 1098.024317 | 0.86168 |
| KLF2 | chr19 | 16433642 | 35 | 32 | 3180.820277 | 2919.436395 | 0.862 |
| AKT2 | chr19 | 40798831 | 32 | 32 | 2974.570491 | 2985.736967 | 0.86206 |
| APOA1 | chr11 | 116670738 | 29 | 27 | 1936.108246 | 1770.789503 | 0.86232 |
| ALDOA | chr16 | 30057669 | 24 | 27 | 2140.150242 | 2420.828996 | 0.86236 |
| APOA1 | chr11 | 116718488 | 33 | 32 | 2855.944021 | 2758.02525 | 0.86242 |
| APOA1 | chr11 | 116724951 | 5 | 5 | 409.1666667 | 400 | 0.86262 |
| ABCG1 | chr21 | 43637311 | 35 | 32 | 3426.921037 | 3126.270892 | 0.86268 |
| ADRA2C | chr4 | 3766850 | 18 | 19 | 1516.508908 | 1618.170996 | 0.86292 |
| APOA1 | chr11 | 116699936 | 3 | 3 | 231.7460318 | 243.3333333 | 0.86296 |
| NOS3 | chr7 | 150705937 | 20 | 23 | 658.7551646 | 740.249758 | 0.86306 |
| LPA | chr6 | 11219959 | 11 | 16 | 1039.393939 | 1521.428571 | 0.86308 |
| APOA1 | chr11 | 116681748 | 33 | 33 | 2357.349615 | 2340.313394 | 0.86324 |
| APOA1 | chr11 | 116672069 | 32 | 32 | 2710.748886 | 2721.893216 | 0.86332 |
| DHFR | chr5 | 79957099 | 26 | 25 | 1653.742116 | 1618.00812 | 0.86376 |
| APOA1 | chr11 | 116724218 | 32 | 33 | 1984.284553 | 2020.004856 | 0.86394 |

| | | | | | | | |
|---|---|---|---|---|---|---|---|
| GALNT2 | chr1 | 230188481 | 16 | 15 | 1497.087176 | 1396.536797 | 0.86402 |
| P2RY12 | chr3 | 151112544 | 34 | 32 | 3284.745254 | 3098.749719 | 0.8644 |
| APOA1 | chr11 | 116661695 | 29 | 31 | 2070.799814 | 2233.717704 | 0.86454 |
| BCL2 | chr18 | 60996452 | 33 | 31 | 3164.719393 | 2978.276258 | 0.86464 |
| FOXP3 | chrX | 49124432 | 28 | 24 | 2548.106296 | 2196.008159 | 0.86556 |
| F7 | chr13 | 113758678 | 24 | 23 | 2284.06746 | 2200.357143 | 0.86576 |
| FOXP3 | chrX | 49126891 | 3 | 5 | 130 | 197.1428571 | 0.86594 |
| BAX | chr19 | 49465421 | 17 | 24 | 607.7771983 | 874.1515204 | 0.86636 |
| F7 | chr13 | 113750978 | 29 | 24 | 2538.54768 | 2077.196333 | 0.86668 |
| F2RL3 | chr19 | 16990469 | 27 | 31 | 2592.599512 | 2960.985599 | 0.86678 |
| APOA1 | chr11 | 116686874 | 32 | 32 | 2942.671523 | 2954.986799 | 0.86692 |
| LDLR | chr19 | 11217944 | 29 | 29 | 2797.10928 | 2806.300187 | 0.867 |
| DHFR | chr5 | 79953676 | 35 | 32 | 3294.385915 | 3019.688005 | 0.86706 |
| MIA3 | chr1 | 222814921 | 29 | 29 | 2439.759464 | 2455.935037 | 0.86714 |
| F7 | chr13 | 113756293 | 36 | 32 | 3124.102827 | 2785.705262 | 0.86732 |
| CDKN2B-AS1 | chr9 | 22002716 | 38 | 33 | 2425.128483 | 2121.127901 | 0.86738 |
| F7 | chr13 | 113751619 | 33 | 32 | 2890.774289 | 2814.883923 | 0.86764 |
| APOA1 | chr11 | 116686139 | 9 | 10 | 854.1666667 | 955 | 0.86768 |
| APOA1 | chr11 | 116669926 | 38 | 32 | 3189.360018 | 2694.804779 | 0.86792 |
| ADRA2C | chr4 | 3766844 | 17 | 17 | 1354.68254 | 1370.4329 | 0.86894 |
| FGF2 | chr4 | 123740222 | 29 | 27 | 2482.768343 | 2328.495949 | 0.86898 |
| FGF2 | chr4 | 123742581 | 26 | 27 | 2389.341631 | 2469.507349 | 0.86904 |
| ICAM2 | chr17 | 62085828 | 34 | 33 | 2310.912541 | 2270.015268 | 0.86912 |
| FOXP3 | chrX | 49118609 | 12 | 19 | 1128.097736 | 1776.547619 | 0.86924 |
| PECAM1 | chr17 | 62401868 | 21 | 22 | 1845.448718 | 1914.199134 | 0.86942 |
| CDKN2BAS | chr9 | 22104207 | 38 | 32 | 3275.091047 | 2748.560215 | 0.8695 |
| COL15A1 | chr9 | 101697376 | 35 | 33 | 2765.033542 | 2592.636507 | 0.86958 |
| CDKN2B-AS1 | chr9 | 21970963 | 9 | 11 | 293.1796412 | 345.9693815 | 0.86972 |
| LOXL1 | chr15 | 74213939 | 29 | 31 | 1831.185437 | 1935.338217 | 0.86982 |
| ABCA1 | chr9 | 107691326 | 32 | 32 | 2920.539289 | 2931.303565 | 0.8701 |
| LDLR | chr19 | 11191989 | 3 | 2 | 260 | 180 | 0.87028 |
| COL15A1 | chr9 | 101703926 | 3 | 2 | 260 | 180 | 0.87028 |
| ALOX15 | chr17 | 4552036 | 29 | 29 | 1117.073802 | 1097.850319 | 0.87048 |
| ABCA1 | chr9 | 107698328 | 28 | 29 | 2209.87093 | 2321.942414 | 0.8707 |
| F7 | chr13 | 113753879 | 37 | 32 | 3398.359519 | 2946.57407 | 0.87078 |
| CDKN2B-AS1 | chr9 | 21993645 | 16 | 13 | 535.7264957 | 445.6967787 | 0.87088 |
| LOXL1 | chr15 | 74244278 | 6 | 5 | 443.992674 | 360.1190476 | 0.87098 |
| FOXP3 | chrX | 49123414 | 32 | 32 | 3007.619449 | 2998.920087 | 0.87104 |
| HMGCR | chr5 | 74623252 | 36 | 32 | 3074.965245 | 2722.058479 | 0.87126 |
| TERT | chr5 | 1298006 | 23 | 20 | 2052.891649 | 1798.203463 | 0.87128 |
| NOS3 | chr7 | 150694028 | 6 | 7 | 207.4603175 | 255.4761905 | 0.87136 |
| LCK | chr1 | 32713275 | 32 | 32 | 2772.816997 | 2760.405614 | 0.87144 |
| PLA2G7 | chr6 | 46706546 | 33 | 32 | 2772.595888 | 2678.156473 | 0.87148 |
| APOA1 | chr11 | 116676441 | 31 | 31 | 2871.003495 | 2882.005726 | 0.8716 |
| F7 | chr13 | 113759558 | 30 | 29 | 2400.451242 | 2333.548577 | 0.87198 |
| TERT | chr5 | 1301467 | 31 | 30 | 2535.382386 | 2438.994976 | 0.872 |
| LCK | chr1 | 32740044 | 33 | 32 | 3046.670331 | 2942.567949 | 0.87206 |

| | | | | | | | |
|---|---|---|---|---|---|---|---|
| F7 | chr13 | 113750995 | 30 | 23 | 2814.723361 | 2140.316726 | 0.87268 |
| LDLR | chr19 | 11196544 | 10 | 4 | 938.4090909 | 380 | 0.87416 |
| LDLR | chr19 | 11224131 | 12 | 16 | 1079.404762 | 1460 | 0.87484 |
| GCK | chr7 | 44229005 | 35 | 32 | 3134.11119 | 2855.92478 | 0.8749 |
| F2RL3 | chr19 | 16998668 | 14 | 16 | 1117.437022 | 1293.531746 | 0.87498 |
| ADRA2C | chr4 | 3759442 | 33 | 33 | 1995.23763 | 1980.696444 | 0.875 |
| APOA1 | chr11 | 116686839 | 32 | 33 | 2429.004017 | 2518.8182 | 0.87522 |
| ALDOA | chr16 | 30062086 | 34 | 32 | 2977.53359 | 2810.550041 | 0.87524 |
| LPA | chr6 | 160948952 | 34 | 33 | 2868.608141 | 2797.853725 | 0.87526 |
| DHFR | chr5 | 79951974 | 36 | 32 | 3407.580022 | 3022.428123 | 0.87536 |
| APOA1 | chr11 | 116664172 | 5 | 3 | 110 | 65 | 0.87536 |
| FOXP3 | chrX | 49121221 | 17 | 12 | 1451.269841 | 1012.222222 | 0.87598 |
| TERT | chr5 | 1298976 | 32 | 33 | 1839.70108 | 1877.950815 | 0.87622 |
| LCK | chr1 | 32739981 | 30 | 33 | 1732.165188 | 1887.954606 | 0.8763 |
| NOS3 | chr7 | 150689881 | 12 | 16 | 467.1139971 | 608.2683983 | 0.8763 |
| MCPH1 | chr8 | 6470381 | 33 | 32 | 2975.560433 | 2872.085975 | 0.87636 |
| LDLR | chr19 | 11194888 | 32 | 32 | 2815.850186 | 2801.295791 | 0.8764 |
| PLA2G7 | chr6 | 46706038 | 34 | 32 | 3229.44851 | 3033.550969 | 0.87652 |
| F7 | chr13 | 113750487 | 25 | 26 | 2283.101343 | 2386.073926 | 0.87658 |
| TIMP3 | chr22 | 33195343 | 41 | 33 | 2913.266003 | 2333.179631 | 0.87698 |
| TMPO | chr12 | 98907342 | 40 | 32 | 3695.081324 | 2951.231073 | 0.87736 |
| ABCB1 | chr7 | 87351216 | 32 | 32 | 2755.37306 | 2745.775576 | 0.87772 |
| ABCG1 | chr21 | 43632149 | 16 | 20 | 1218.949384 | 1540.652958 | 0.87774 |
| ESR2 | chr14 | 64814793 | 24 | 25 | 2300.270147 | 2411.014402 | 0.87824 |
| NOS3 | chr7 | 150684108 | 7 | 13 | 594.4444445 | 1117.380952 | 0.87824 |
| KLF2 | chr19 | 16426573 | 14 | 17 | 1173.650794 | 1445.555556 | 0.87862 |
| APOA1 | chr11 | 116716431 | 37 | 32 | 3317.489789 | 2876.406845 | 0.87886 |
| APOA1 | chr11 | 116661042 | 33 | 32 | 3172.712383 | 3082.772566 | 0.8789 |
| TIMP3 | chr22 | 33187582 | 32 | 32 | 2957.635265 | 2965.771262 | 0.87898 |
| CDKN2B-AS1 | chr9 | 21984591 | 36 | 32 | 3143.209526 | 2801.36857 | 0.87918 |
| ABCG1 | chr21 | 43636907 | 34 | 32 | 3206.010792 | 3025.398697 | 0.8792 |
| CDKN2B-AS1 | chr9 | 21989821 | 15 | 20 | 594.4444444 | 810.3496503 | 0.87934 |
| TERT | chr5 | 1299757 | 20 | 29 | 1278.085901 | 1816.946754 | 0.87946 |
| ABCG1 | chr21 | 43633218 | 35 | 32 | 3300.202494 | 3010.58784 | 0.8796 |
| TIMP3 | chr22 | 33187494 | 34 | 33 | 3046.28253 | 2943.421783 | 0.8796 |
| LDLR | chr19 | 11223908 | 20 | 20 | 1923.333333 | 1929.880952 | 0.8797 |
| ICAM2 | chr17 | 62085933 | 13 | 12 | 1245.238095 | 1155.714286 | 0.87994 |
| LPA | chr6 | 160945476 | 36 | 32 | 3297.208995 | 2925.306044 | 0.87998 |
| GCK | chr7 | 44229558 | 32 | 32 | 3016.052519 | 3008.17188 | 0.8804 |
| BAX | chr19 | 49450628 | 13 | 24 | 1223.214286 | 2268.812021 | 0.88042 |
| ABCA1 | chr9 | 107690745 | 9 | 12 | 313.4126984 | 425.8333333 | 0.8807 |
| LOXL1 | chr15 | 74244344 | 19 | 14 | 748.0502896 | 538.8258408 | 0.88072 |
| TERT | chr5 | 1300899 | 23 | 21 | 1847.896825 | 1668.928571 | 0.88086 |
| CDKN2B-AS1 | chr9 | 21988774 | 39 | 33 | 2044.112144 | 1716.571619 | 0.88088 |
| F2RL3 | chr19 | 17000231 | 8 | 9 | 316.7857143 | 369.6825397 | 0.88098 |
| LCK | chr1 | 32715428 | 30 | 31 | 1284.762982 | 1343.52689 | 0.88146 |
| ESR1 | chr6 | 152002878 | 22 | 20 | 729.7829475 | 675.911347 | 0.88164 |

| | | | | | | | |
|---|---|---|---|---|---|---|---|
| LCK | chr1 | 32741152 | 21 | 22 | 779.1206016 | 800.8238545 | 0.88226 |
| TERT | chr5 | 1300820 | 8 | 8 | 653.3333333 | 666.6666667 | 0.8827 |
| GALNT2 | chr1 | 230188031 | 34 | 32 | 3214.527306 | 3015.187572 | 0.88272 |
| ABCA1 | chr9 | 107690774 | 16 | 17 | 1000.577201 | 1081.150794 | 0.88284 |
| LCK | chr1 | 32712237 | 8 | 6 | 387.5 | 301.9047619 | 0.8829 |
| LPA | chr6 | 11216215 | 34 | 32 | 3269.2669 | 3081.824712 | 0.88342 |
| CTGF | chr6 | 132277919 | 27 | 29 | 1778.069847 | 1935.942946 | 0.88354 |
| CDKN2B-AS1 | chr9 | 21968388 | 2 | 6 | 62.22222222 | 205.5555556 | 0.88368 |
| LDLR | chr19 | 11194911 | 34 | 32 | 2964.045217 | 2799.518786 | 0.884 |
| ALOX15 | chr17 | 4547307 | 37 | 33 | 2775.413976 | 2462.752427 | 0.88456 |
| FOXP3 | chrX | 49130289 | 30 | 29 | 1880.88797 | 1797.423623 | 0.8848 |
| PHACTR1 | chr6 | 12716137 | 32 | 33 | 2524.494222 | 2586.713139 | 0.88486 |
| LCK | chr1 | 32739752 | 26 | 29 | 912.2695884 | 1028.478671 | 0.88492 |
| CDKN2BAS | chr9 | 22103812 | 26 | 31 | 1777.642366 | 2091.490549 | 0.88546 |
| ABCG1 | chr21 | 43635417 | 19 | 19 | 1778.637849 | 1769.87013 | 0.88548 |
| F2RL3 | chr19 | 16994742 | 4 | 14 | 206.6666667 | 697.9069541 | 0.88548 |
| APOA1 | chr11 | 116693076 | 35 | 32 | 2620.852987 | 2384.818256 | 0.88554 |
| LCK | chr1 | 32713188 | 31 | 32 | 3008.005999 | 3112.33446 | 0.88562 |
| CDKN1A | chr6 | 36645500 | 33 | 32 | 2361.187571 | 2278.980488 | 0.88584 |
| E2F6 | chr2 | 11611897 | 28 | 28 | 2114.663356 | 2130.882898 | 0.8859 |
| PLA2G7 | chr6 | 46705358 | 25 | 25 | 2182.162698 | 2193.690476 | 0.88622 |
| APOA1 | chr11 | 116707519 | 29 | 27 | 2749.752747 | 2570.353535 | 0.88624 |
| CDKN2B-AS1 | chr9 | 21978980 | 23 | 23 | 797.5969711 | 809.4562112 | 0.88658 |
| CDKN2B-AS1 | chr9 | 21980775 | 2 | 3 | 140 | 215 | 0.88672 |
| ALDOA | chr16 | 30062262 | 33 | 32 | 3070.46607 | 2963.83137 | 0.88674 |
| ADRA2C | chr4 | 3759226 | 32 | 32 | 1793.048371 | 1808.73366 | 0.88684 |
| CDKN2B-AS1 | chr9 | 22008314 | 12 | 11 | 350.232233 | 330.263659 | 0.88696 |
| F7 | chr13 | 113756218 | 34 | 33 | 1598.037923 | 1539.477273 | 0.88698 |
| ABCA1 | chr9 | 107699196 | 12 | 12 | 847.4682223 | 835.7239409 | 0.88716 |
| ALOX15 | chr17 | 4548776 | 33 | 32 | 3009.821289 | 2912.992486 | 0.88736 |
| BCL2 | chr18 | 60991143 | 9 | 7 | 850.3296703 | 666.6666667 | 0.88738 |
| FGF2 | chr4 | 123741401 | 35 | 32 | 2851.314406 | 2615.180217 | 0.88742 |
| ABCA1 | chr9 | 107690771 | 17 | 15 | 1052.12482 | 912.020202 | 0.88766 |
| AMOTL2 | chr3 | 134098667 | 2 | 3 | 160 | 232.3809524 | 0.88806 |
| ABCG1 | chr21 | 43637496 | 33 | 32 | 3110.605357 | 3009.946348 | 0.88854 |
| FOXP3 | chrX | 49116231 | 29 | 29 | 2611.246032 | 2600.077839 | 0.88862 |
| APOA1 | chr11 | 116679014 | 35 | 32 | 2523.935705 | 2318.408421 | 0.88884 |
| PECAM1 | chr17 | 62398344 | 32 | 32 | 2996.208125 | 3002.316243 | 0.88904 |
| CTGF | chr6 | 132273902 | 38 | 32 | 3612.280143 | 3036.603572 | 0.88948 |
| COL15A1 | chr9 | 101699737 | 32 | 33 | 1931.800618 | 1971.744324 | 0.88988 |
| F7 | chr13 | 113759220 | 3 | 3 | 243.3333333 | 250 | 0.88992 |
| ESR1 | chr6 | 152002104 | 31 | 32 | 2956.316048 | 3059.786911 | 0.89 |
| BAX | chr19 | 49454676 | 29 | 28 | 2636.782107 | 2557.589869 | 0.89008 |
| LDLR | chr19 | 11217665 | 5 | 3 | 471.6666667 | 285.7142857 | 0.89032 |
| GALNT2 | chr1 | 230187253 | 31 | 27 | 3032.561336 | 2633.183997 | 0.89054 |
| F7 | chr13 | 113757758 | 34 | 33 | 2574.955869 | 2484.258935 | 0.89058 |
| LDLR | chr19 | 11216089 | 28 | 30 | 2565.758963 | 2737.912878 | 0.89062 |

| | | | | | | | |
|---|---|---|---|---|---|---|---|
| CDKN1A | chr6 | 36648835 | 32 | 33 | 2594.904309 | 2660.068663 | 0.8913 |
| KLF2 | chr19 | 16426675 | 33 | 30 | 3009.460577 | 2725.572863 | 0.8914 |
| CTGF | chr6 | 132279650 | 38 | 32 | 3161.882278 | 2654.299229 | 0.8915 |
| F7 | chr13 | 113775254 | 26 | 28 | 1961.329745 | 2096.948899 | 0.89166 |
| APOA1 | chr11 | 116697345 | 24 | 25 | 1607.733056 | 1695.047614 | 0.89204 |
| F2RL3 | chr19 | 16998796 | 27 | 31 | 1750.184544 | 2027.88073 | 0.8925 |
| F7 | chr13 | 113753082 | 34 | 32 | 3276.556174 | 3088.199466 | 0.8926 |
| APOA1 | chr11 | 116672325 | 34 | 32 | 3260.350968 | 3062.193398 | 0.8926 |
| LCK | chr1 | 32711529 | 33 | 32 | 2880.1225 | 2781.1945 | 0.89278 |
| ESR2 | chr14 | 64810458 | 35 | 32 | 3277.196537 | 3000.920809 | 0.89294 |
| F2RL3 | chr19 | 16990816 | 34 | 32 | 3256.527802 | 3069.11133 | 0.893 |
| F7 | chr13 | 113751410 | 27 | 26 | 2503.268398 | 2420.481659 | 0.89306 |
| TERT | chr5 | 1253593 | 32 | 32 | 2936.657831 | 2927.215153 | 0.89332 |
| CDKN2B-AS1 | chr9 | 22010691 | 32 | 32 | 2664.051575 | 2677.888897 | 0.89336 |
| CTGF | chr6 | 132281522 | 33 | 29 | 2357.857838 | 2060.993905 | 0.89344 |
| TERT | chr5 | 1300231 | 31 | 32 | 2594.718481 | 2689.532553 | 0.89356 |
| ABCB1 | chr7 | 87348875 | 29 | 31 | 2678.706675 | 2874.629121 | 0.89356 |
| AMOTL2 | chr3 | 134074279 | 15 | 13 | 1392.380952 | 1194.615385 | 0.89366 |
| F7 | chr13 | 113751346 | 19 | 18 | 1669.960317 | 1570.634921 | 0.89378 |
| ARHGEF18 | chr19 | 7468270 | 31 | 32 | 2279.946405 | 2337.42121 | 0.89388 |
| PHACTR1 | chr6 | 12717790 | 35 | 32 | 2861.712123 | 2608.093425 | 0.89454 |
| ADRA2C | chr4 | 3764649 | 31 | 31 | 2923.22263 | 2913.044307 | 0.89458 |
| GCK | chr7 | 44183806 | 35 | 32 | 3399.882721 | 3111.877492 | 0.89476 |
| NOS3 | chr7 | 150695400 | 24 | 18 | 1672.783883 | 1268.968254 | 0.89478 |
| COL15A1 | chr9 | 101699271 | 36 | 32 | 3433.104377 | 3046.698072 | 0.89494 |
| TIMP3 | chr22 | 33192396 | 35 | 32 | 3354.686703 | 3063.275748 | 0.89502 |
| NOS3 | chr7 | 150694062 | 7 | 2 | 336.6666667 | 93.33333333 | 0.89506 |
| FGF2 | chr4 | 123740029 | 35 | 32 | 3255.393348 | 2969.790077 | 0.89538 |
| LOXL1 | chr15 | 74209739 | 38 | 32 | 3685.148641 | 3099.21974 | 0.89544 |
| NOS3 | chr7 | 150707758 | 2 | 3 | 77.5 | 127.3024361 | 0.89544 |
| ABCB1 | chr7 | 87351940 | 35 | 33 | 2838.453571 | 2663.14593 | 0.89566 |
| APOA1 | chr11 | 116692287 | 32 | 32 | 3090.247763 | 3096.388839 | 0.89602 |
| GALNT2 | chr1 | 230418375 | 7 | 9 | 567.7777778 | 738.4848485 | 0.89602 |
| ALDOA | chr16 | 30060471 | 33 | 33 | 2652.121803 | 2666.918277 | 0.89626 |
| TERT | chr5 | 1295666 | 6 | 8 | 162.7777778 | 221.5628816 | 0.89674 |
| FOXP3 | chrX | 49125810 | 2 | 2 | 107.1428571 | 120 | 0.89678 |
| TERT | chr5 | 1295800 | 30 | 29 | 1161.64864 | 1133.112964 | 0.89718 |
| P2RY12 | chr3 | 151104217 | 26 | 30 | 1984.065102 | 2317.416056 | 0.89736 |
| NOS3 | chr7 | 150678186 | 33 | 32 | 2699.392568 | 2607.983787 | 0.89748 |
| APOA1 | chr11 | 116680031 | 23 | 16 | 2052.278048 | 1418.650794 | 0.89754 |
| KLF2 | chr19 | 16433189 | 18 | 11 | 1753.492063 | 1067.857143 | 0.898 |
| LPA | chr6 | 11224220 | 34 | 32 | 3230.17546 | 3034.390045 | 0.8983 |
| TIMP3 | chr22 | 33197061 | 35 | 33 | 1866.439241 | 1748.028658 | 0.89844 |
| APOA1 | chr11 | 116663840 | 39 | 32 | 3676.037292 | 3020.853956 | 0.89846 |
| APOA1 | chr11 | 116684488 | 37 | 32 | 3455.710963 | 2983.639958 | 0.89876 |
| PHACTR1 | chr6 | 12708732 | 35 | 32 | 3144.967776 | 2880.700516 | 0.89892 |
| F2RL3 | chr19 | 16992455 | 16 | 16 | 1580 | 1583.333333 | 0.89898 |

| | | | | | | | |
|---|---|---|---|---|---|---|---|
| DHFR | chr5 | 79956266 | 31 | 30 | 2953.184201 | 2840.792541 | 0.89906 |
| NOS3 | chr7 | 150687388 | 37 | 32 | 2802.189094 | 2433.289658 | 0.89916 |
| LDLR | chr19 | 11194649 | 35 | 33 | 2162.6154 | 2049.443366 | 0.89924 |
| LDLR | chr19 | 11222134 | 29 | 30 | 2514.621906 | 2613.782802 | 0.8994 |
| LDLR | chr19 | 11215905 | 28 | 28 | 2550.40182 | 2540.062993 | 0.8994 |
| ESR1 | chr6 | 152006563 | 12 | 12 | 1132.307692 | 1138.492063 | 0.8994 |
| FOXP3 | chrX | 49127011 | 20 | 13 | 1477.261905 | 949.5959596 | 0.89956 |
| APOA1 | chr11 | 116661301 | 32 | 32 | 2995.360602 | 2987.847762 | 0.89974 |
| NOS3 | chr7 | 150683050 | 12 | 15 | 586.6498779 | 742.8684211 | 0.89994 |
| APOA1 | chr11 | 116667171 | 32 | 31 | 2963.214998 | 2861.23947 | 0.90014 |
| BCL2 | chr18 | 60989046 | 7 | 9 | 594.7619048 | 756.4285714 | 0.90022 |
| NOS3 | chr7 | 150678792 | 33 | 32 | 2699.714884 | 2627.874642 | 0.90028 |
| APOA1 | chr11 | 116673356 | 30 | 31 | 2539.204846 | 2611.165779 | 0.90048 |
| NOS3 | chr7 | 150695591 | 13 | 20 | 929.6320346 | 1410.284993 | 0.90052 |
| AKT2 | chr19 | 40794896 | 33 | 32 | 3156.484589 | 3065.448932 | 0.90102 |
| MIA3 | chr1 | 222816937 | 19 | 25 | 1493.160173 | 1945.386003 | 0.9013 |
| APOA1 | chr11 | 116702641 | 32 | 31 | 2875.859064 | 2794.544764 | 0.90152 |
| FOXP3 | chrX | 49121240 | 19 | 17 | 1679.242424 | 1490 | 0.90168 |
| ALDOA | chr16 | 30058320 | 30 | 31 | 2518.177222 | 2586.548705 | 0.90216 |
| PECAM1 | chr17 | 62400777 | 21 | 18 | 1899.520202 | 1637.178932 | 0.90216 |
| GCK | chr7 | 44233448 | 32 | 32 | 2599.801397 | 2608.904897 | 0.90226 |
| NOS3 | chr7 | 150691432 | 27 | 31 | 994.2859713 | 1154.391971 | 0.9023 |
| LDLR | chr19 | 11192516 | 31 | 25 | 2693.776772 | 2181.308789 | 0.90258 |
| APOA1 | chr11 | 116661001 | 34 | 32 | 3210.381233 | 3016.776842 | 0.90276 |
| APOA1 | chr11 | 116693860 | 33 | 32 | 3151.060875 | 3061.049788 | 0.90288 |
| BAX | chr19 | 49454601 | 33 | 31 | 2734.926579 | 2579.710018 | 0.9029 |
| LPA | chr6 | 11216548 | 35 | 32 | 3405.363816 | 3116.708407 | 0.90302 |
| ARHGEF18 | chr19 | 7537511 | 31 | 32 | 2863.681454 | 2964.51266 | 0.90312 |
| ICAM2 | chr17 | 62080249 | 21 | 22 | 1977.619048 | 2078.809524 | 0.90404 |
| CDKN2B-AS1 | chr9 | 21993118 | 39 | 33 | 3056.879756 | 2596.120709 | 0.90408 |
| CDKN2B-AS1 | chr9 | 21980734 | 18 | 17 | 1695.714286 | 1608.214286 | 0.90408 |
| TIMP3 | chr22 | 33188552 | 34 | 32 | 3060.187287 | 2885.576501 | 0.90418 |
| F7 | chr13 | 113753071 | 34 | 32 | 3315.48786 | 3117.308398 | 0.90422 |
| ALDOA | chr16 | 30060869 | 32 | 32 | 2872.217562 | 2881.135695 | 0.90438 |
| AKT2 | chr19 | 40795339 | 19 | 19 | 1809.126984 | 1819.166667 | 0.90446 |
| ADRA2C | chr4 | 3759478 | 33 | 32 | 3068.750067 | 2967.773551 | 0.90494 |
| ADRA2C | chr4 | 3762758 | 16 | 20 | 476.4311545 | 584.7807699 | 0.90514 |
| ALOX15 | chr17 | 4545384 | 23 | 20 | 957.1664446 | 844.2540792 | 0.9052 |
| E2F6 | chr2 | 11608213 | 34 | 32 | 3072.944493 | 2899.809975 | 0.90528 |
| TERT | chr5 | 1299414 | 18 | 17 | 1500.079365 | 1426.071429 | 0.90564 |
| APOA1 | chr11 | 116661892 | 32 | 33 | 2440.378717 | 2528.574688 | 0.90624 |
| BCL2 | chr18 | 60988274 | 19 | 27 | 689.4343405 | 992.0017945 | 0.9063 |
| CDKN2B-AS1 | chr9 | 21988096 | 36 | 33 | 1797.089275 | 1637.357923 | 0.90634 |
| APOA1 | chr11 | 116668627 | 32 | 32 | 3126.851359 | 3123.279422 | 0.90638 |
| CTGF | chr6 | 132277334 | 16 | 14 | 771.1338386 | 687.361364 | 0.90638 |
| ADRA2C | chr4 | 3762274 | 34 | 32 | 3153.455629 | 2974.1092 | 0.90678 |
| LDLR | chr19 | 11221456 | 28 | 28 | 1889.282921 | 1867.135418 | 0.90704 |

| | | | | | | | |
|---|---|---|---|---|---|---|---|
| FOXP3 | chrX | 49114371 | 32 | 32 | 2229.591136 | 2218.804417 | 0.9071 |
| ESR2 | chr14 | 64806937 | 20 | 23 | 1945 | 2241.269841 | 0.90712 |
| ALOX15 | chr17 | 4545388 | 28 | 24 | 1614.476357 | 1370.3774 | 0.90742 |
| CDKN2B-AS1 | chr9 | 21964752 | 15 | 17 | 1084.603175 | 1241.071429 | 0.9075 |
| CDKN1A | chr6 | 36654979 | 31 | 32 | 2692.622043 | 2771.163823 | 0.9078 |
| TIMP3 | chr22 | 33188261 | 33 | 32 | 2428.059755 | 2344.315606 | 0.90796 |
| LPA | chr6 | 11215984 | 25 | 25 | 2422.38345 | 2416.686508 | 0.9081 |
| FOXP3 | chrX | 49126866 | 13 | 12 | 624.5238095 | 565.9704185 | 0.90834 |
| E2F6 | chr2 | 11612963 | 34 | 32 | 3284.056608 | 3086.434989 | 0.90838 |
| ESR2 | chr14 | 64806159 | 37 | 33 | 1854.669862 | 1643.833888 | 0.90844 |
| APOA1 | chr11 | 116707281 | 18 | 26 | 1713.214286 | 2481.421356 | 0.90862 |
| APOA1 | chr11 | 116703640 | 18 | 19 | 1686.873119 | 1786.178571 | 0.90914 |
| BCL2 | chr18 | 60995216 | 36 | 32 | 3307.405476 | 2934.810385 | 0.90956 |
| F2RL3 | chr19 | 16999136 | 24 | 30 | 2010.477023 | 2527.813853 | 0.90962 |
| PECAM1 | chr17 | 62401213 | 31 | 29 | 2686.541748 | 2522.270151 | 0.90982 |
| F2RL3 | chr19 | 16998072 | 2 | 2 | 183.3333333 | 180 | 0.90984 |
| PLA2G7 | chr6 | 46709841 | 35 | 32 | 3226.13701 | 2954.960161 | 0.90988 |
| FOXP3 | chrX | 49123437 | 31 | 32 | 2984.548525 | 3073.75927 | 0.90994 |
| LCK | chr1 | 32741766 | 13 | 12 | 1190.119048 | 1092.261905 | 0.91002 |
| NOS3 | chr7 | 150694383 | 13 | 13 | 428.8971168 | 436.9369519 | 0.91008 |
| ALOX15 | chr17 | 4549203 | 35 | 33 | 2838.817114 | 2690.117404 | 0.91022 |
| CDKN2B-AS1 | chr9 | 21966435 | 12 | 19 | 445.7683983 | 693.9498247 | 0.91028 |
| ALOX15 | chr17 | 4552894 | 30 | 32 | 2459.731535 | 2610.244547 | 0.91048 |
| F7 | chr13 | 113753016 | 32 | 32 | 3096.077613 | 3100.191485 | 0.9105 |
| GCK | chr7 | 44237303 | 35 | 32 | 3288.462449 | 3002.613754 | 0.9111 |
| F2RL3 | chr19 | 16994188 | 32 | 32 | 3138.6147 | 3135.076711 | 0.9112 |
| TERT | chr5 | 1301385 | 30 | 33 | 1906.061107 | 2110.236776 | 0.91132 |
| APOA1 | chr11 | 116700831 | 13 | 13 | 982.8174603 | 993.2142857 | 0.91144 |
| APOA1 | chr11 | 116697773 | 31 | 31 | 2587.580245 | 2596.065687 | 0.91168 |
| ALOX15 | chr17 | 4547979 | 32 | 33 | 2760.499505 | 2858.293456 | 0.91178 |
| APOA1 | chr11 | 116710168 | 16 | 17 | 941.0939242 | 985.3735154 | 0.91188 |
| BCL2 | chr18 | 60988598 | 31 | 33 | 2596.395296 | 2775.729406 | 0.91204 |
| HMGCR | chr5 | 74627138 | 28 | 28 | 2330.72976 | 2321.078023 | 0.9121 |
| ESR1 | chr6 | 152011178 | 38 | 32 | 3199.000228 | 2688.785309 | 0.91212 |
| F7 | chr13 | 113750944 | 30 | 28 | 2726.782107 | 2554.123377 | 0.91214 |
| CDKN1A | chr6 | 36651974 | 31 | 32 | 2986.274615 | 3077.551174 | 0.91238 |
| F2RL3 | chr19 | 16995280 | 33 | 32 | 3132.051743 | 3041.60671 | 0.91254 |
| TERT | chr5 | 1298512 | 2 | 6 | 177.7777778 | 540 | 0.91254 |
| LOXL1 | chr15 | 74215418 | 10 | 11 | 455 | 489.7619048 | 0.91256 |
| LDLR | chr19 | 11220915 | 11 | 9 | 1063.333333 | 866.6666667 | 0.91266 |
| CDKN2B-AS1 | chr9 | 22002737 | 37 | 33 | 1815.493981 | 1627.076134 | 0.9128 |
| APOA1 | chr11 | 116678717 | 19 | 22 | 1653.150183 | 1926.211705 | 0.91304 |
| LPA | chr6 | 11219136 | 32 | 27 | 1243.115569 | 1058.814066 | 0.91344 |
| ESR1 | chr6 | 152004261 | 33 | 32 | 2958.793367 | 2874.931249 | 0.91364 |
| GALNT2 | chr1 | 230190570 | 36 | 32 | 3309.05403 | 2936.782724 | 0.91384 |
| AMOTL2 | chr3 | 134094894 | 35 | 33 | 2714.860494 | 2579.81615 | 0.9142 |
| LDLR | chr19 | 11220439 | 4 | 3 | 369.047619 | 274.6031746 | 0.91438 |

| | | | | | | | |
|---|---|---|---|---|---|---|---|
| ESR2 | chr14 | 64806643 | 32 | 31 | 3009.634496 | 2927.254653 | 0.91442 |
| NOS3 | chr7 | 150710524 | 29 | 31 | 1597.302214 | 1693.186735 | 0.9146 |
| ADRA2C | chr4 | 3768584 | 12 | 16 | 365.5588179 | 479.9447082 | 0.91466 |
| ADRA2C | chr4 | 3761162 | 30 | 32 | 2732.467562 | 2903.937215 | 0.91474 |
| LDLR | chr19 | 11222678 | 27 | 32 | 1933.06689 | 2304.914946 | 0.91478 |
| ADRA2C | chr4 | 3762337 | 33 | 32 | 3063.729785 | 2964.560298 | 0.91492 |
| CTGF | chr6 | 132272944 | 27 | 28 | 906.073197 | 929.6878546 | 0.91498 |
| LOXL1 | chr15 | 74215784 | 31 | 32 | 2294.650069 | 2379.757543 | 0.91522 |
| LDLR | chr19 | 11217255 | 24 | 31 | 2353.409091 | 3035.308025 | 0.91544 |
| NOS3 | chr7 | 150710066 | 30 | 32 | 1356.315606 | 1460.123964 | 0.9159 |
| APOA1 | chr11 | 116718759 | 22 | 24 | 2091.345599 | 2276.246392 | 0.9159 |
| ARHGEF18 | chr19 | 7537497 | 32 | 32 | 3019.394981 | 3013.871263 | 0.91592 |
| CDKN2B-AS1 | chr9 | 22006218 | 28 | 32 | 1162.351982 | 1317.798221 | 0.91626 |
| APOA1 | chr11 | 116665516 | 28 | 32 | 2347.207074 | 2668.330003 | 0.91636 |
| P2RY12 | chr3 | 151107378 | 34 | 32 | 3040.889764 | 2867.471752 | 0.91638 |
| CDKN2B-AS1 | chr9 | 21969849 | 7 | 11 | 200.2380952 | 306.5873016 | 0.91642 |
| FOXP3 | chrX | 49106814 | 22 | 23 | 2112.806638 | 2203.736264 | 0.91648 |
| PECAM1 | chr17 | 62407287 | 15 | 13 | 1430.555556 | 1234.603175 | 0.91654 |
| ABCA1 | chr9 | 107693947 | 28 | 28 | 2714.933955 | 2707.20567 | 0.9173 |
| LPA | chr6 | 11219099 | 30 | 28 | 1178.025294 | 1090.423989 | 0.91732 |
| ALOX15 | chr17 | 4553258 | 24 | 25 | 1085.087853 | 1116.055056 | 0.91758 |
| NOS3 | chr7 | 150710116 | 12 | 13 | 502.3953824 | 553.5714286 | 0.91796 |
| NOS3 | chr7 | 150687569 | 34 | 32 | 3162.578564 | 2970.465538 | 0.91798 |
| E2F6 | chr2 | 11607092 | 35 | 33 | 1828.48456 | 1713.626453 | 0.91844 |
| ADRA2C | chr4 | 3770160 | 32 | 33 | 1902.825532 | 1950.277477 | 0.91844 |
| ESR2 | chr14 | 64809252 | 32 | 32 | 3104.224037 | 3099.976551 | 0.91852 |
| GCK | chr7 | 44237632 | 33 | 32 | 2997.131019 | 2900.763022 | 0.91856 |
| PHACTR1 | chr6 | 12717776 | 35 | 32 | 3260.921379 | 2986.086996 | 0.91858 |
| NOS3 | chr7 | 150709186 | 34 | 30 | 1047.83889 | 932.0603107 | 0.9192 |
| APOA1 | chr11 | 116707169 | 30 | 33 | 1758.122763 | 1921.313647 | 0.91936 |
| KLF2 | chr19 | 16429076 | 25 | 22 | 2253.170996 | 1973.863636 | 0.9197 |
| CDKN2BAS | chr9 | 22067118 | 25 | 21 | 1979.949495 | 1673.272283 | 0.91992 |
| ESR1 | chr6 | 152002518 | 27 | 28 | 2019.29732 | 2106.346534 | 0.92006 |
| LDLR | chr19 | 11221755 | 19 | 17 | 1693.556166 | 1521.288156 | 0.9201 |
| FOXP3 | chrX | 49127489 | 31 | 32 | 2499.858007 | 2569.407034 | 0.92014 |
| PLA2G7 | chr6 | 46713021 | 40 | 33 | 2899.453035 | 2383.822786 | 0.92024 |
| FOXP3 | chrX | 49131131 | 33 | 32 | 3133.668082 | 3044.673585 | 0.92042 |
| TERT | chr5 | 1298327 | 31 | 32 | 2749.547789 | 2845.941575 | 0.92042 |
| TIMP3 | chr22 | 33194446 | 31 | 32 | 2124.073655 | 2178.907397 | 0.9206 |
| CDKN2B-AS1 | chr9 | 21966565 | 26 | 28 | 991.9626577 | 1058.277944 | 0.9209 |
| TERT | chr5 | 1298672 | 31 | 31 | 2904.45671 | 2910.460583 | 0.92096 |
| TERT | chr5 | 1294723 | 6 | 3 | 153.8888889 | 78.33333333 | 0.92112 |
| LPA | chr6 | 160951507 | 7 | 13 | 642.1031746 | 1185.595238 | 0.92122 |
| TERT | chr5 | 1295708 | 19 | 23 | 669.5176388 | 818.4504303 | 0.9214 |
| FOXP3 | chrX | 49126139 | 2 | 4 | 86.36363636 | 160 | 0.9214 |
| LOXL1 | chr15 | 74219236 | 19 | 15 | 737.6182394 | 590.0226408 | 0.92162 |
| TIMP3 | chr22 | 33190874 | 38 | 32 | 3613.723102 | 3039.049978 | 0.9217 |

| | | | | | | | |
|---|---|---|---|---|---|---|---|
| FOXP3 | chrX | 49116873 | 10 | 15 | 940.3174603 | 1404.166667 | 0.9217 |
| F7 | chr13 | 113753323 | 28 | 33 | 2282.431166 | 2678.161962 | 0.92172 |
| LOXL1 | chr15 | 74215773 | 28 | 33 | 1616.574247 | 1892.584989 | 0.9219 |
| AKT2 | chr19 | 40798699 | 33 | 32 | 2658.252744 | 2586.153009 | 0.9222 |
| BCL2 | chr18 | 60992455 | 32 | 32 | 3105.374576 | 3101.458223 | 0.9222 |
| NOS3 | chr7 | 150691143 | 33 | 33 | 2489.235489 | 2480.451954 | 0.92232 |
| F7 | chr13 | 113760346 | 7 | 6 | 517.5324675 | 450.7142857 | 0.92236 |
| CDKN2B-AS1 | chr9 | 21977387 | 20 | 23 | 777.8314659 | 904.3939394 | 0.92244 |
| ABCG1 | chr21 | 43634663 | 18 | 24 | 1679.95671 | 2230 | 0.92252 |
| BAX | chr19 | 49449573 | 32 | 31 | 3029.801717 | 2941.18044 | 0.92256 |
| ABCB1 | chr7 | 87352289 | 36 | 32 | 3300.405993 | 2939.362965 | 0.92256 |
| FOXP3 | chrX | 49119856 | 12 | 17 | 806.6666667 | 1156.071429 | 0.92256 |
| F7 | chr13 | 113757707 | 33 | 33 | 2522.099877 | 2513.363915 | 0.92258 |
| F7 | chr13 | 113751316 | 5 | 5 | 463.3333333 | 460 | 0.9228 |
| GCK | chr7 | 44229009 | 35 | 32 | 3254.064359 | 2970.32633 | 0.92294 |
| CDKN1A | chr6 | 36652296 | 33 | 32 | 3009.488501 | 2924.925115 | 0.9233 |
| LDLR | chr19 | 11219624 | 34 | 32 | 3206.133392 | 3014.108799 | 0.92332 |
| P2RY12 | chr3 | 151102323 | 34 | 32 | 2889.934919 | 2714.728682 | 0.92372 |
| CDKN1A | chr6 | 36653548 | 12 | 14 | 458.6967418 | 526.9552669 | 0.9241 |
| GCK | chr7 | 44230177 | 36 | 32 | 3272.084841 | 2912.676623 | 0.92424 |
| COL15A1 | chr9 | 101703777 | 4 | 3 | 338.9285714 | 251.4285714 | 0.92454 |
| ESR2 | chr14 | 64806848 | 36 | 32 | 3544.072411 | 3146.442695 | 0.92488 |
| ARHGEF18 | chr19 | 7460794 | 17 | 18 | 837.7342755 | 898.6868687 | 0.92496 |
| FOXP3 | chrX | 49115057 | 11 | 11 | 922.1031746 | 931.4285714 | 0.92498 |
| F7 | chr13 | 113752095 | 32 | 32 | 2918.4356 | 2926.857691 | 0.92504 |
| ESR2 | chr14 | 64807190 | 26 | 24 | 2377.758126 | 2204.361472 | 0.9252 |
| F2RL3 | chr19 | 16994752 | 32 | 31 | 3010.498257 | 2923.193658 | 0.92524 |
| ICAM2 | chr17 | 62080079 | 33 | 32 | 3066.667995 | 2967.59702 | 0.92526 |
| LPA | chr6 | 160944113 | 23 | 23 | 2218.888889 | 2214.166667 | 0.92534 |
| CDKN2B-AS1 | chr9 | 22006181 | 15 | 17 | 467.103189 | 539.4906556 | 0.92552 |
| P2RY12 | chr3 | 151109980 | 32 | 32 | 2864.827135 | 2857.158261 | 0.9256 |
| ARHGEF18 | chr19 | 7468124 | 33 | 32 | 3052.028864 | 2964.861577 | 0.9261 |
| GCK | chr7 | 44237296 | 35 | 33 | 2681.043188 | 2519.909912 | 0.92618 |
| ABCB1 | chr7 | 87133635 | 37 | 32 | 3534.677004 | 3050.937294 | 0.92634 |
| CDKN2B-AS1 | chr9 | 21967596 | 14 | 17 | 388.9149584 | 465.9046671 | 0.92688 |
| TIMP3 | chr22 | 33195922 | 38 | 32 | 3510.110767 | 2959.856833 | 0.92698 |
| TERT | chr5 | 1296008 | 28 | 32 | 2013.2089 | 2313.063296 | 0.9271 |
| APOA1 | chr11 | 116726399 | 9 | 9 | 273.5411255 | 280.2925697 | 0.92764 |
| MCPH1 | chr8 | 6256530 | 33 | 33 | 2712.092127 | 2703.211865 | 0.92766 |
| AMOTL2 | chr3 | 134096390 | 32 | 33 | 2628.513713 | 2701.399327 | 0.92786 |
| E2F6 | chr2 | 11615731 | 20 | 19 | 1589.126984 | 1517.97619 | 0.9279 |
| BCL2 | chr18 | 60989552 | 32 | 32 | 1324.587249 | 1316.237243 | 0.92826 |
| FOXP3 | chrX | 49126815 | 2 | 2 | 93.33333333 | 100 | 0.92838 |
| ADRA2C | chr4 | 3768656 | 23 | 27 | 837.6211663 | 974.9848645 | 0.9286 |
| ADRA2C | chr4 | 3762081 | 29 | 28 | 2318.829422 | 2229.621453 | 0.92872 |
| LDLR | chr19 | 11197859 | 7 | 9 | 668.8888889 | 863.3333333 | 0.9288 |
| PECAM1 | chr17 | 62408142 | 31 | 31 | 2634.980627 | 2641.812512 | 0.92896 |

| | | | | | | | |
|---|---|---|---|---|---|---|---|
| COL15A1 | chr9 | 101705884 | 18 | 11 | 574.6903097 | 358.0135387 | 0.92912 |
| TIMP3 | chr22 | 33187350 | 35 | 32 | 3080.122572 | 2821.921818 | 0.9292 |
| ARHGEF18 | chr19 | 7460712 | 18 | 24 | 581.1454789 | 782.7837749 | 0.92946 |
| MCPH1 | chr8 | 6470917 | 19 | 29 | 1767.018634 | 2705.08436 | 0.9299 |
| TFPI | chr2 | 188424747 | 5 | 2 | 445.7142857 | 180 | 0.92998 |
| BCL2 | chr18 | 60988515 | 34 | 33 | 1504.720445 | 1451.792381 | 0.9301 |
| LPA | chr6 | 160945060 | 38 | 32 | 3682.048277 | 3097.418579 | 0.93012 |
| NOS3 | chr7 | 150686787 | 32 | 33 | 2636.561327 | 2709.473171 | 0.9302 |
| NOS3 | chr7 | 150694403 | 28 | 31 | 1400.631085 | 1539.079162 | 0.93034 |
| LDLR | chr19 | 11224157 | 6 | 8 | 528.5714286 | 713.3333333 | 0.93038 |
| F7 | chr13 | 113750200 | 32 | 32 | 3049.802809 | 3040.258239 | 0.93042 |
| TIMP3 | chr22 | 33192544 | 30 | 33 | 2060.24413 | 2251.761669 | 0.93048 |
| KLF2 | chr19 | 16430210 | 2 | 8 | 105.1282051 | 406.3059163 | 0.93072 |
| TERT | chr5 | 1300209 | 32 | 33 | 1976.889215 | 2028.155594 | 0.93074 |
| E2F6 | chr2 | 11612054 | 19 | 17 | 1836.547619 | 1639.79798 | 0.93074 |
| ADRA2C | chr4 | 3768778 | 19 | 19 | 751.7063492 | 743.0294705 | 0.9308 |
| LPA | chr6 | 11223897 | 33 | 32 | 3184.017341 | 3090.715148 | 0.93088 |
| E2F6 | chr2 | 11584561 | 32 | 32 | 2818.843779 | 2813.445415 | 0.93092 |
| LDLR | chr19 | 11191624 | 11 | 7 | 997.9761905 | 630.8333333 | 0.93092 |
| CTGF | chr6 | 132276372 | 35 | 32 | 3286.472142 | 3008.255507 | 0.93156 |
| F2RL3 | chr19 | 16997515 | 10 | 4 | 797.1428571 | 323.2142857 | 0.93158 |
| CDKN2B-AS1 | chr9 | 21967891 | 17 | 18 | 490.4295386 | 525.5435188 | 0.93164 |
| ESR1 | chr6 | 152005542 | 35 | 32 | 3326.736445 | 3038.398222 | 0.9317 |
| ABCA1 | chr9 | 107690720 | 11 | 12 | 461.5873016 | 511.5223665 | 0.93178 |
| TERT | chr5 | 1300291 | 31 | 31 | 2667.004387 | 2659.798829 | 0.93182 |
| BAX | chr19 | 49453863 | 3 | 8 | 283.3333333 | 760 | 0.93192 |
| PECAM1 | chr17 | 62400139 | 6 | 4 | 571.4285714 | 383.3333333 | 0.93242 |
| ALOX15 | chr17 | 4551493 | 25 | 24 | 1027.791445 | 979.0567765 | 0.93254 |
| PECAM1 | chr17 | 62410445 | 7 | 4 | 594.1666667 | 343.3333333 | 0.93254 |
| APOA1 | chr11 | 116698946 | 27 | 29 | 2431.518949 | 2618.246198 | 0.93264 |
| ARHGEF18 | chr19 | 7465898 | 37 | 32 | 3240.765693 | 2799.053173 | 0.93292 |
| KLF2 | chr19 | 16427015 | 31 | 30 | 1179.632557 | 1150.213781 | 0.93318 |
| CDKN2B-AS1 | chr9 | 21996286 | 35 | 33 | 1557.697481 | 1476.640308 | 0.93326 |
| F2RL3 | chr19 | 16990647 | 25 | 27 | 2322.418831 | 2500.716783 | 0.93332 |
| COL15A1 | chr9 | 101705968 | 2 | 5 | 62.85714286 | 153.3333333 | 0.93354 |
| KLF2 | chr19 | 16432671 | 32 | 32 | 2787.628237 | 2781.737478 | 0.9336 |
| ADRA2C | chr4 | 3759195 | 31 | 32 | 2428.063852 | 2513.869217 | 0.93366 |
| LPA | chr6 | 160944477 | 35 | 32 | 2921.947673 | 2666.124861 | 0.93376 |
| F2RL3 | chr19 | 16994897 | 30 | 27 | 2824.280706 | 2536.831502 | 0.9339 |
| APOA1 | chr11 | 116694172 | 17 | 18 | 1414.184982 | 1504.206349 | 0.93398 |
| NR3C1 | chr5 | 142785934 | 22 | 21 | 2090.595238 | 1990.555556 | 0.93408 |
| LOXL1 | chr15 | 74212042 | 19 | 24 | 707.9900019 | 887.6562367 | 0.93422 |
| FOXP3 | chrX | 49119954 | 33 | 32 | 3023.211225 | 2927.765967 | 0.93442 |
| F7 | chr13 | 113758753 | 32 | 32 | 2947.145632 | 2938.217778 | 0.93466 |
| PHACTR1 | chr6 | 12717057 | 33 | 32 | 3083.003705 | 2994.129195 | 0.93472 |
| CTGF | chr6 | 132273258 | 23 | 23 | 1424.826097 | 1436.147741 | 0.93474 |
| PECAM1 | chr17 | 62407210 | 12 | 13 | 1080.833333 | 1176.666667 | 0.9348 |

| | | | | | | | |
|---|---|---|---|---|---|---|---|
| CDKN2BAS | chr9 | 22098908 | 34 | 32 | 3045.154555 | 2870.297955 | 0.935 |
| PHACTR1 | chr6 | 12717736 | 34 | 32 | 3054.36579 | 2871.051055 | 0.93536 |
| F7 | chr13 | 113758964 | 27 | 27 | 2210.379065 | 2202.07709 | 0.93542 |
| AMOTL2 | chr3 | 134097685 | 38 | 33 | 2529.765948 | 2190.006362 | 0.93564 |
| ADRA2C | chr4 | 3760888 | 13 | 16 | 1228.021978 | 1515.079365 | 0.93564 |
| NOS3 | chr7 | 150681579 | 34 | 32 | 3155.052537 | 2966.194142 | 0.9357 |
| LOXL1 | chr15 | 74211115 | 35 | 32 | 3295.90752 | 3010.149972 | 0.93576 |
| APOA1 | chr11 | 116699689 | 34 | 32 | 2696.569944 | 2545.177991 | 0.93592 |
| PECAM1 | chr17 | 62398786 | 28 | 29 | 1968.353175 | 2027.254909 | 0.93608 |
| PECAM1 | chr17 | 62410115 | 30 | 28 | 1079.991437 | 1000.77953 | 0.93616 |
| LDLR | chr19 | 11219075 | 11 | 11 | 720.8333333 | 712.0238095 | 0.93618 |
| AKT2 | chr19 | 40800037 | 31 | 26 | 2290.562394 | 1911.693862 | 0.9363 |
| FOXP3 | chrX | 49130017 | 8 | 6 | 249.4303786 | 191.9503439 | 0.93658 |
| CDKN2B-AS1 | chr9 | 21984029 | 34 | 33 | 2541.834241 | 2459.31074 | 0.93668 |
| CDKN2B-AS1 | chr9 | 21977490 | 11 | 10 | 353.9285714 | 317.7777778 | 0.937 |
| NOS3 | chr7 | 150706569 | 19 | 20 | 1805.277778 | 1909.206349 | 0.93716 |
| LDLR | chr19 | 11216146 | 27 | 29 | 2549.848485 | 2733.870215 | 0.93736 |
| BAX | chr19 | 49449580 | 32 | 32 | 2794.435594 | 2787.377535 | 0.93758 |
| FOXP3 | chrX | 49128199 | 32 | 30 | 2917.722381 | 2729.331779 | 0.93774 |
| LDLR | chr19 | 11215876 | 24 | 24 | 2345.310245 | 2342.222222 | 0.93814 |
| TERT | chr5 | 1295706 | 15 | 19 | 489.7063186 | 614.6801809 | 0.93872 |
| APOA1 | chr11 | 116706052 | 17 | 17 | 636.7490842 | 630.8080808 | 0.93884 |
| AKT2 | chr19 | 40736404 | 32 | 31 | 2131.285107 | 2055.62984 | 0.93892 |
| F7 | chr13 | 113751748 | 32 | 32 | 2731.603408 | 2736.395427 | 0.93896 |
| APOA1 | chr11 | 116663701 | 32 | 32 | 3056.08067 | 3059.94398 | 0.93904 |
| LOXL1 | chr15 | 74219225 | 20 | 14 | 766.4325698 | 542.8451228 | 0.93918 |
| LOXL1 | chr15 | 74218554 | 33 | 33 | 2156.402148 | 2147.86192 | 0.93946 |
| LCK | chr1 | 32740740 | 21 | 18 | 1945.039683 | 1673.095238 | 0.93964 |
| APOA1 | chr11 | 116703393 | 33 | 32 | 2972.989009 | 2886.342663 | 0.9397 |
| FGF2 | chr4 | 123747253 | 33 | 32 | 2980.228046 | 2884.988499 | 0.93978 |
| APOA1 | chr11 | 116662801 | 31 | 32 | 2504.577966 | 2592.308766 | 0.9399 |
| LPA | chr6 | 160948723 | 36 | 32 | 2778.307249 | 2465.748257 | 0.94044 |
| APOA1 | chr11 | 116684685 | 32 | 32 | 2781.009731 | 2789.016637 | 0.94082 |
| APOA1 | chr11 | 116661518 | 29 | 31 | 2633.76089 | 2808.210956 | 0.94092 |
| LDLR | chr19 | 11215925 | 31 | 30 | 3046.061912 | 2945.116607 | 0.94096 |
| APOA1 | chr11 | 116679526 | 34 | 32 | 2931.133447 | 2753.361529 | 0.94098 |
| LOXL1 | chr15 | 74209723 | 38 | 32 | 3669.837756 | 3088.371621 | 0.94118 |
| ESR2 | chr14 | 64807157 | 4 | 4 | 335.5555556 | 332.7777778 | 0.9414 |
| ESR1 | chr6 | 152003052 | 4 | 7 | 141.9978802 | 241.2448553 | 0.9417 |
| LDLR | chr19 | 11193116 | 24 | 22 | 2320.079365 | 2129.761905 | 0.94264 |
| F7 | chr13 | 113775124 | 29 | 31 | 2522.525635 | 2703.008576 | 0.94352 |
| MCPH1 | chr8 | 6470493 | 34 | 32 | 3023.61588 | 2841.993325 | 0.94382 |
| ESR2 | chr14 | 64806689 | 37 | 32 | 3497.783218 | 3021.399901 | 0.94382 |
| LOXL1 | chr15 | 74212306 | 33 | 32 | 3065.016597 | 2976.903305 | 0.94384 |
| LDLR | chr19 | 11221323 | 33 | 32 | 3185.113577 | 3091.650215 | 0.94392 |
| F7 | chr13 | 113756539 | 34 | 33 | 2599.747427 | 2536.956987 | 0.9444 |
| AMOTL2 | chr3 | 134094133 | 33 | 32 | 1380.007667 | 1344.705606 | 0.94444 |

| | | | | | | | |
|---|---|---|---|---|---|---|---|
| NOS3 | chr7 | 150683001 | 8 | 9 | 500.3751804 | 568.8816739 | 0.94448 |
| LDLR | chr19 | 11198067 | 32 | 31 | 3132.521225 | 3040.392488 | 0.94508 |
| LPA | chr6 | 160945050 | 38 | 32 | 3624.970382 | 3049.892401 | 0.9452 |
| ABCG1 | chr21 | 43637157 | 30 | 31 | 2772.2312 | 2859.001124 | 0.94562 |
| LDLR | chr19 | 11196538 | 11 | 4 | 1057.45098 | 385.7142857 | 0.94588 |
| ABCA1 | chr9 | 107691596 | 24 | 22 | 2133.349567 | 1950.515873 | 0.94624 |
| LCK | chr1 | 32741672 | 30 | 32 | 2437.842925 | 2607.738163 | 0.94626 |
| APOA1 | chr11 | 116718980 | 4 | 5 | 355 | 440 | 0.94634 |
| COL15A1 | chr9 | 101705436 | 14 | 13 | 712.3000489 | 652.6254267 | 0.94646 |
| APOA1 | chr11 | 116685339 | 33 | 33 | 2407.749331 | 2399.099866 | 0.9465 |
| TERT | chr5 | 1299907 | 31 | 32 | 2651.18593 | 2743.526586 | 0.94654 |
| AKT2 | chr19 | 40736506 | 32 | 32 | 2521.560766 | 2527.761105 | 0.94658 |
| ADRA2C | chr4 | 3759146 | 32 | 32 | 2608.216861 | 2600.829015 | 0.9467 |
| F2RL3 | chr19 | 16999175 | 10 | 17 | 834.1666667 | 1409.047619 | 0.94712 |
| TIMP3 | chr22 | 33190824 | 37 | 32 | 3172.510831 | 2748.16731 | 0.94742 |
| F7 | chr13 | 113751494 | 28 | 28 | 2677.58547 | 2672.714646 | 0.94746 |
| ADRA2C | chr4 | 3762519 | 25 | 26 | 2043.120326 | 2133.587575 | 0.94752 |
| GCK | chr7 | 44235002 | 31 | 32 | 3008.725212 | 3103.33593 | 0.94802 |
| LPA | chr6 | 160943055 | 35 | 32 | 3311.031394 | 3029.682181 | 0.9482 |
| APOA1 | chr11 | 116724173 | 30 | 30 | 1399.397447 | 1391.669719 | 0.9482 |
| AKT2 | chr19 | 40736428 | 31 | 31 | 1376.143366 | 1381.866817 | 0.94834 |
| F2RL3 | chr19 | 16991864 | 32 | 31 | 3076.429918 | 2983.988095 | 0.94852 |
| CTGF | chr6 | 132274156 | 36 | 32 | 3345.435191 | 2975.777988 | 0.9486 |
| LDLR | chr19 | 11216175 | 25 | 25 | 2448.816796 | 2443.845599 | 0.94876 |
| BAX | chr19 | 49448341 | 33 | 33 | 2904.616429 | 2911.198751 | 0.9488 |
| GCK | chr7 | 44228650 | 30 | 33 | 2154.724689 | 2363.599226 | 0.9489 |
| LDLR | chr19 | 11222002 | 31 | 31 | 2751.833082 | 2757.134182 | 0.94896 |
| APOA1 | chr11 | 116695823 | 25 | 26 | 2242.72049 | 2327.514528 | 0.949 |
| BCL2 | chr18 | 60989550 | 33 | 32 | 1508.917636 | 1456.357545 | 0.94928 |
| APOA1 | chr11 | 116659523 | 13 | 6 | 1180 | 547.5 | 0.94962 |
| LOXL1 | chr15 | 74218718 | 17 | 19 | 619.9350649 | 686.9280719 | 0.9497 |
| BCL2 | chr18 | 60994512 | 38 | 32 | 3608.6742 | 3040.908544 | 0.94978 |
| ALOX15 | chr17 | 4553312 | 4 | 8 | 146.6666667 | 289.4047619 | 0.94988 |
| NOS3 | chr7 | 150706363 | 28 | 27 | 2368.211957 | 2277.076703 | 0.94998 |
| ICAM2 | chr17 | 62085259 | 31 | 29 | 2892.710623 | 2702.068487 | 0.95026 |
| CDKN2B-AS1 | chr9 | 21980687 | 31 | 33 | 2728.00877 | 2913.082546 | 0.95026 |
| ARHGEF18 | chr19 | 7465066 | 36 | 32 | 3439.002169 | 3054.155127 | 0.95054 |
| LDLR | chr19 | 11219011 | 11 | 12 | 564.1666667 | 608.4126984 | 0.95062 |
| KLF2 | chr19 | 16430583 | 17 | 17 | 566.8457808 | 572.1836376 | 0.95084 |
| TMPO | chr12 | 98901037 | 34 | 32 | 3138.33372 | 2956.485908 | 0.95116 |
| TIMP3 | chr22 | 33193709 | 35 | 33 | 2620.852793 | 2465.481297 | 0.9514 |
| BAX | chr19 | 49448185 | 33 | 32 | 3093.634425 | 2996.609095 | 0.9514 |
| F7 | chr13 | 113752526 | 31 | 33 | 2816.193722 | 2992.001256 | 0.95152 |
| F7 | chr13 | 113775021 | 29 | 30 | 2203.860495 | 2285.759122 | 0.9517 |
| LOXL1 | chr15 | 74218830 | 24 | 22 | 783.1394642 | 713.2648352 | 0.95204 |
| LDLR | chr19 | 11191799 | 11 | 9 | 602.3593074 | 497.5 | 0.9521 |
| LDLR | chr19 | 11221073 | 22 | 19 | 2104.410867 | 1813.888889 | 0.95214 |

| | | | | | | | |
|---|---|---|---|---|---|---|---|
| TIMP3 | chr22 | 33259396 | 27 | 29 | 2369.058275 | 2537.293099 | 0.95222 |
| F7 | chr13 | 113760510 | 30 | 29 | 1675.586562 | 1628.202353 | 0.95234 |
| CTGF | chr6 | 132276374 | 35 | 32 | 3328.781193 | 3041.258841 | 0.95246 |
| LDLR | chr19 | 11190635 | 31 | 32 | 2849.59991 | 2945.507285 | 0.9525 |
| CDKN2B-AS1 | chr9 | 21969438 | 17 | 14 | 628.2423433 | 521.8559219 | 0.95252 |
| CDKN2BAS | chr9 | 22067542 | 15 | 13 | 835.5404812 | 731.7201949 | 0.9534 |
| ICAM2 | chr17 | 62084068 | 29 | 30 | 1136.351228 | 1170.112096 | 0.95344 |
| CDKN1A | chr6 | 36651944 | 31 | 31 | 2926.621016 | 2929.381528 | 0.95348 |
| FOXP3 | chrX | 49117977 | 32 | 31 | 2878.221843 | 2783.832769 | 0.95364 |
| APOA1 | chr11 | 116715794 | 34 | 32 | 3220.087405 | 3033.170045 | 0.95402 |
| CDKN2B-AS1 | chr9 | 21990225 | 21 | 21 | 678.9553844 | 683.6336501 | 0.95402 |
| BCL2 | chr18 | 60989319 | 28 | 30 | 2471.727043 | 2652.925367 | 0.95408 |
| APOA1 | chr11 | 116669790 | 31 | 32 | 2912.653351 | 3003.367768 | 0.9541 |
| CDKN1A | chr6 | 36652014 | 31 | 30 | 2932.507405 | 2841.726052 | 0.9542 |
| APOA1 | chr11 | 116694766 | 37 | 32 | 3520.693119 | 3040.571074 | 0.95434 |
| LPA | chr6 | 11221200 | 38 | 32 | 3199.702295 | 2690.953923 | 0.95438 |
| TMPO | chr12 | 98907828 | 27 | 21 | 897.5593486 | 693.9659268 | 0.95452 |
| ALDOA | chr16 | 30078393 | 12 | 11 | 386.6017316 | 357.7777778 | 0.9546 |
| APOA1 | chr11 | 116711064 | 35 | 32 | 3067.717669 | 2801.767189 | 0.95466 |
| CDKN2B-AS1 | chr9 | 21996521 | 16 | 14 | 454.8981705 | 402.6449679 | 0.95472 |
| LPA | chr6 | 11216595 | 36 | 32 | 3508.849721 | 3117.484665 | 0.95474 |
| NR3C1 | chr5 | 142792831 | 32 | 32 | 3045.406559 | 3049.446145 | 0.95482 |
| NOS3 | chr7 | 150687503 | 35 | 32 | 3112.942287 | 2842.475237 | 0.95488 |
| CDKN1A | chr6 | 36652305 | 32 | 32 | 3063.41542 | 3060.03121 | 0.9549 |
| MIA3 | chr1 | 222815968 | 36 | 32 | 3205.183195 | 2852.115962 | 0.95496 |
| PECAM1 | chr17 | 62398094 | 37 | 32 | 1445.789607 | 1245.200856 | 0.95502 |
| APOA1 | chr11 | 116663805 | 38 | 32 | 3331.633811 | 2802.979299 | 0.95502 |
| BCL2 | chr18 | 60992502 | 31 | 30 | 2468.375069 | 2382.269675 | 0.95506 |
| HMGCR | chr5 | 74623354 | 32 | 32 | 2896.515154 | 2893.109381 | 0.95522 |
| CDKN1A | chr6 | 36651328 | 29 | 32 | 2605.096545 | 2870.403778 | 0.9561 |
| F7 | chr13 | 113750385 | 23 | 22 | 2149.964115 | 2052.896825 | 0.95612 |
| PHACTR1 | chr6 | 12714871 | 33 | 32 | 2842.863082 | 2753.046517 | 0.95634 |
| GCK | chr7 | 44234738 | 34 | 32 | 2448.615743 | 2314.024217 | 0.9564 |
| PHACTR1 | chr6 | 12712998 | 6 | 6 | 206.9047619 | 210 | 0.95662 |
| PLA2G7 | chr6 | 46703671 | 24 | 27 | 735.7700238 | 832.1401011 | 0.95668 |
| F2RL3 | chr19 | 16993024 | 33 | 33 | 2906.958524 | 2912.687064 | 0.9572 |
| AKT2 | chr19 | 40800726 | 37 | 32 | 3297.013213 | 2848.061721 | 0.958 |
| CDKN2BAS | chr9 | 22098573 | 26 | 30 | 1786.646425 | 2072.916987 | 0.95806 |
| TERT | chr5 | 1295651 | 4 | 13 | 109.1774892 | 357.5396825 | 0.95816 |
| LDLR | chr19 | 11191297 | 22 | 26 | 1968.021978 | 2320.17094 | 0.95842 |
| APOA1 | chr11 | 116661547 | 30 | 32 | 2254.696585 | 2412.097723 | 0.95844 |
| ESR1 | chr6 | 152011511 | 22 | 19 | 633.6962065 | 551.3055297 | 0.95876 |
| MIA3 | chr1 | 222792386 | 3 | 6 | 110 | 213.8705739 | 0.95876 |
| TERT | chr5 | 1295762 | 27 | 29 | 1056.748405 | 1130.487891 | 0.95914 |
| BCL2 | chr18 | 60989843 | 28 | 29 | 1079.727677 | 1123.798562 | 0.95914 |
| APOA1 | chr11 | 116678986 | 34 | 32 | 3179.118551 | 2989.734368 | 0.95936 |
| AKT2 | chr19 | 40791843 | 24 | 26 | 705.1910607 | 760.1404541 | 0.95948 |

| | | | | | | | |
|---|---|---|---|---|---|---|---|
| F2RL3 | chr19 | 16998963 | 32 | 32 | 2930.800889 | 2934.402916 | 0.95984 |
| ALDOA | chr16 | 30057712 | 30 | 31 | 2479.922897 | 2567.826428 | 0.96 |
| ABCG1 | chr21 | 43636936 | 33 | 32 | 3172.897038 | 3073.899355 | 0.96012 |
| LPA | chr6 | 11216726 | 36 | 32 | 2907.297168 | 2593.111396 | 0.96036 |
| F2RL3 | chr19 | 16993325 | 34 | 32 | 3351.121087 | 3155.584536 | 0.96038 |
| CDKN2B-AS1 | chr9 | 21973983 | 3 | 10 | 93.07692308 | 305.5982906 | 0.96062 |
| APOA1 | chr11 | 116662407 | 28 | 31 | 2381.399798 | 2631.143195 | 0.96072 |
| APOA1 | chr11 | 116690579 | 25 | 25 | 1378.316167 | 1385.004671 | 0.96074 |
| ICAM2 | chr17 | 62084912 | 30 | 32 | 2754.133351 | 2941.488523 | 0.96076 |
| MCPH1 | chr8 | 6470328 | 20 | 26 | 1806.453824 | 2352.681624 | 0.96106 |
| APOA1 | chr11 | 116663743 | 33 | 32 | 3102.635328 | 3005.590267 | 0.9611 |
| F2RL3 | chr19 | 16990973 | 28 | 29 | 2506.312092 | 2588.615551 | 0.96118 |
| CDKN2BAS | chr9 | 22096457 | 36 | 32 | 3469.397471 | 3082.481864 | 0.96124 |
| BAX | chr19 | 49456476 | 25 | 27 | 2041.592866 | 2209.79576 | 0.96128 |
| FOXP3 | chrX | 49116394 | 22 | 21 | 1882.780112 | 1801.093074 | 0.96134 |
| ADRA2C | chr4 | 3764035 | 32 | 32 | 2894.178797 | 2897.439726 | 0.96168 |
| MIA3 | chr1 | 222816658 | 32 | 31 | 2989.388881 | 2898.531511 | 0.96178 |
| MIA3 | chr1 | 222792590 | 29 | 31 | 1227.160937 | 1307.33796 | 0.96196 |
| TIMP3 | chr22 | 33191140 | 37 | 33 | 2598.641906 | 2313.382401 | 0.9623 |
| PECAM1 | chr17 | 62400643 | 28 | 30 | 2511.691079 | 2686.488475 | 0.96232 |
| E2F6 | chr2 | 11613049 | 31 | 30 | 2770.683683 | 2678.15073 | 0.96298 |
| CDKN2B-AS1 | chr9 | 21989796 | 9 | 13 | 272.3179272 | 390.5588856 | 0.96308 |
| CDKN2B-AS1 | chr9 | 21970957 | 9 | 12 | 251.314127 | 338.1875314 | 0.96348 |
| NOS3 | chr7 | 150688516 | 36 | 32 | 3287.990032 | 2920.050557 | 0.96364 |
| TIMP3 | chr22 | 33196610 | 33 | 33 | 2541.556193 | 2545.921582 | 0.96368 |
| LDLR | chr19 | 11221271 | 33 | 32 | 3188.911867 | 3094.094126 | 0.9638 |
| APOA1 | chr11 | 116693345 | 34 | 32 | 3219.637842 | 3027.285183 | 0.96382 |
| NOS3 | chr7 | 150706582 | 21 | 23 | 1688.296703 | 1854.664502 | 0.96398 |
| DHFR | chr5 | 79956708 | 26 | 28 | 2344.074853 | 2521.142052 | 0.96438 |
| LOXL1 | chr15 | 74218965 | 22 | 21 | 740.0789847 | 702.7464481 | 0.96448 |
| CDKN2B-AS1 | chr9 | 21967490 | 16 | 20 | 519.0740996 | 644.8080013 | 0.9648 |
| LOXL1 | chr15 | 74219065 | 31 | 29 | 1249.293232 | 1163.763866 | 0.96486 |
| ESR1 | chr6 | 152003194 | 19 | 18 | 560.7145539 | 528.3166524 | 0.96544 |
| TFPI | chr2 | 188420196 | 10 | 11 | 523.2390771 | 579.3146152 | 0.96556 |
| PECAM1 | chr17 | 62399904 | 36 | 33 | 1679.322525 | 1542.560627 | 0.9657 |
| TMPO | chr12 | 98901069 | 35 | 32 | 3351.801509 | 3065.918645 | 0.96578 |
| APOA1 | chr11 | 116662393 | 30 | 32 | 2056.250684 | 2188.26011 | 0.966 |
| LDLR | chr19 | 11223911 | 20 | 20 | 1929.880952 | 1928.214286 | 0.96622 |
| LDLR | chr19 | 11223288 | 24 | 24 | 2332.380952 | 2330.641026 | 0.9664 |
| NR3C1 | chr5 | 142791812 | 5 | 2 | 395.5555556 | 160 | 0.9665 |
| FOXP3 | chrX | 49122581 | 34 | 32 | 3310.828705 | 3119.057939 | 0.96678 |
| BCL2 | chr18 | 60992423 | 32 | 32 | 3084.504468 | 3082.816675 | 0.96756 |
| APOA1 | chr11 | 116699101 | 25 | 29 | 2108.506608 | 2450.339318 | 0.96806 |
| ALDOA | chr16 | 30058312 | 32 | 31 | 2930.506429 | 2834.814964 | 0.96856 |
| CDKN1A | chr6 | 36646033 | 12 | 17 | 346.4588763 | 494.2596929 | 0.96864 |
| FOXP3 | chrX | 49117274 | 3 | 2 | 280 | 187.5 | 0.96872 |
| APOA1 | chr11 | 116687084 | 16 | 16 | 1507.103175 | 1504.880952 | 0.96882 |

| | | | | | | | |
|---|---|---|---|---|---|---|---|
| CTGF | chr6 | 132269587 | 38 | 32 | 3326.881473 | 2803.634212 | 0.96916 |
| CDKN2B-AS1 | chr9 | 21989866 | 14 | 18 | 628.8690476 | 804.3849206 | 0.96916 |
| CDKN1A | chr6 | 36651810 | 34 | 32 | 3158.249561 | 2970.541608 | 0.96954 |
| ESR2 | chr14 | 64810082 | 32 | 32 | 3027.81975 | 3026.435709 | 0.96966 |
| NOS3 | chr7 | 150682950 | 14 | 14 | 1122.142857 | 1125.714286 | 0.96968 |
| PHACTR1 | chr6 | 12716260 | 32 | 32 | 2865.657682 | 2868.517071 | 0.96978 |
| GALNT2 | chr1 | 230190670 | 39 | 32 | 3729.462262 | 3058.890714 | 0.97018 |
| CDKN2B-AS1 | chr9 | 21996691 | 21 | 21 | 742.2003091 | 745.5331025 | 0.97018 |
| CDKN1A | chr6 | 36651297 | 29 | 31 | 2590.384409 | 2772.125807 | 0.97028 |
| LDLR | chr19 | 11221268 | 33 | 32 | 3162.690879 | 3065.08189 | 0.97036 |
| APOA1 | chr11 | 116666261 | 33 | 32 | 3096.631542 | 3000.926929 | 0.9705 |
| APOA1 | chr11 | 116689825 | 33 | 33 | 2090.021148 | 2093.640462 | 0.9705 |
| TFPI | chr2 | 188420569 | 29 | 30 | 2736.813844 | 2828.468856 | 0.9705 |
| APOA1 | chr11 | 116695172 | 4 | 3 | 326.4285714 | 245.7142857 | 0.97072 |
| APOA1 | chr11 | 116718988 | 6 | 5 | 554.7619048 | 463.3333333 | 0.9711 |
| APOA1 | chr11 | 116692358 | 33 | 32 | 3162.625285 | 3065.500504 | 0.97112 |
| BAX | chr19 | 49453492 | 14 | 8 | 801.5079365 | 460.8333333 | 0.97126 |
| KLF2 | chr19 | 16426721 | 30 | 30 | 2547.792485 | 2543.424368 | 0.97136 |
| ESR2 | chr14 | 64807679 | 32 | 32 | 3103.359276 | 3105.217474 | 0.97142 |
| BCL2 | chr18 | 60992144 | 36 | 32 | 3166.900052 | 2812.998623 | 0.97164 |
| LDLR | chr19 | 11221335 | 33 | 32 | 3090.911568 | 2995.46278 | 0.97164 |
| LDLR | chr19 | 11222731 | 9 | 16 | 819.7619048 | 1454.444444 | 0.97168 |
| F7 | chr13 | 113753038 | 33 | 32 | 3037.047464 | 2943.028896 | 0.97174 |
| CDKN2B-AS1 | chr9 | 21980741 | 11 | 10 | 830 | 751.4285714 | 0.97174 |
| F7 | chr13 | 113760429 | 24 | 25 | 1999.671717 | 2079.473304 | 0.97186 |
| F7 | chr13 | 113755521 | 4 | 3 | 325.7142857 | 243.3333333 | 0.9721 |
| PHACTR1 | chr6 | 12712421 | 31 | 33 | 1938.523227 | 2067.909112 | 0.97264 |
| CDKN2B-AS1 | chr9 | 21969468 | 14 | 13 | 523.3424908 | 483.4920635 | 0.97266 |
| LOXL1 | chr15 | 74217904 | 33 | 32 | 3157.994621 | 3061.013931 | 0.97276 |
| ABCG1 | chr21 | 43638706 | 18 | 10 | 643.7193362 | 359.7330447 | 0.97296 |
| F2RL3 | chr19 | 16992276 | 32 | 31 | 2573.712796 | 2490.086513 | 0.97298 |
| COL15A1 | chr9 | 101705311 | 32 | 33 | 2853.011269 | 2938.59004 | 0.973 |
| MCPH1 | chr8 | 6470140 | 23 | 27 | 2269.95098 | 2660.707071 | 0.97304 |
| F2RL3 | chr19 | 16994692 | 30 | 32 | 2732.868429 | 2918.688036 | 0.97332 |
| TERT | chr5 | 1298385 | 17 | 21 | 1361.655844 | 1678.715729 | 0.97354 |
| PHACTR1 | chr6 | 12711774 | 36 | 32 | 3013.802017 | 2681.6069 | 0.97366 |
| ESR1 | chr6 | 152006025 | 32 | 31 | 2950.472045 | 2860.167969 | 0.97378 |
| AMOTL2 | chr3 | 134095850 | 33 | 32 | 2798.018359 | 2710.268798 | 0.97388 |
| ESR2 | chr14 | 64807696 | 32 | 31 | 3068.246574 | 2970.855769 | 0.97412 |
| TERT | chr5 | 1301026 | 34 | 32 | 2719.092994 | 2561.802766 | 0.97424 |
| CDKN2B-AS1 | chr9 | 21986843 | 35 | 33 | 2130.021049 | 2011.434835 | 0.97444 |
| MCPH1 | chr8 | 6470717 | 34 | 32 | 3248.96454 | 3059.331609 | 0.97452 |
| ADRA2C | chr4 | 3767226 | 2 | 4 | 87.27272727 | 178.5714286 | 0.97452 |
| CDKN1A | chr6 | 36653719 | 32 | 33 | 1839.66271 | 1893.745068 | 0.97468 |
| LOXL1 | chr15 | 74215804 | 33 | 33 | 1990.513659 | 1987.146276 | 0.97526 |
| LDLR | chr19 | 11217303 | 33 | 31 | 3192.998401 | 3001.351198 | 0.97528 |
| LDLR | chr19 | 11219638 | 33 | 32 | 2752.415343 | 2667.140303 | 0.97556 |

| | | | | | | | |
|---|---|---|---|---|---|---|---|
| ABCG1 | chr21 | 43635767 | 33 | 32 | 3152.960098 | 3058.323145 | 0.97562 |
| E2F6 | chr2 | 11607465 | 24 | 22 | 1907.044818 | 1751.095408 | 0.9758 |
| LDLR | chr19 | 11217863 | 27 | 28 | 2626.474359 | 2722.258297 | 0.97588 |
| NOS3 | chr7 | 150690125 | 9 | 10 | 502.8571429 | 556.1904762 | 0.976 |
| FOXP3 | chrX | 49128649 | 32 | 32 | 2820.568724 | 2818.623895 | 0.97628 |
| ABCG1 | chr21 | 43635173 | 17 | 23 | 980.3632207 | 1321.276464 | 0.97676 |
| APOA1 | chr11 | 116723195 | 23 | 22 | 758.4838789 | 728.0984879 | 0.97678 |
| FOXP3 | chrX | 49116879 | 13 | 15 | 1233.333333 | 1425 | 0.97702 |
| FGF2 | chr4 | 123738376 | 27 | 30 | 1786.877392 | 1989.954579 | 0.97726 |
| F2RL3 | chr19 | 16997374 | 31 | 33 | 2129.101606 | 2269.698231 | 0.97742 |
| CDKN2B-AS1 | chr9 | 21969861 | 8 | 13 | 259.4261294 | 424.3323343 | 0.9775 |
| PLA2G7 | chr6 | 46707290 | 36 | 32 | 3138.535216 | 2787.839222 | 0.97764 |
| NOS3 | chr7 | 150692305 | 30 | 32 | 1312.90617 | 1403.546438 | 0.9778 |
| LPA | chr6 | 11218832 | 34 | 32 | 3294.341989 | 3098.864609 | 0.97786 |
| FGF2 | chr4 | 123738862 | 34 | 32 | 2904.841852 | 2732.384982 | 0.9779 |
| LOXL1 | chr15 | 74214246 | 39 | 33 | 2062.467343 | 1743.248385 | 0.97862 |
| LOXL1 | chr15 | 74208853 | 35 | 32 | 3245.716558 | 2966.485824 | 0.9787 |
| PECAM1 | chr17 | 62407260 | 17 | 17 | 1641.666667 | 1640.15873 | 0.9789 |
| F7 | chr13 | 113759694 | 31 | 31 | 2286.386296 | 2283.798523 | 0.97904 |
| PECAM1 | chr17 | 62399734 | 10 | 17 | 910.3030303 | 1549.880952 | 0.97926 |
| CDKN1A | chr6 | 36652125 | 28 | 29 | 2654.52381 | 2750.951646 | 0.97946 |
| LCK | chr1 | 32712756 | 32 | 31 | 3070.010376 | 2975.409005 | 0.9796 |
| PLA2G7 | chr6 | 46710856 | 36 | 32 | 2349.272865 | 2085.84704 | 0.98028 |
| ESR2 | chr14 | 64809895 | 35 | 32 | 3358.246283 | 3071.164855 | 0.98082 |
| TERT | chr5 | 1297489 | 27 | 24 | 1437.661521 | 1281.389787 | 0.98086 |
| AMOTL2 | chr3 | 134095834 | 33 | 32 | 2775.96775 | 2689.607058 | 0.9809 |
| APOA1 | chr11 | 116707351 | 20 | 19 | 1696.272339 | 1613.170996 | 0.98124 |
| BAX | chr19 | 49451083 | 8 | 7 | 720.3174603 | 629.1666667 | 0.98152 |
| APOA1 | chr11 | 116666319 | 36 | 32 | 3246.268937 | 2884.439484 | 0.98158 |
| FOXP3 | chrX | 49129520 | 9 | 8 | 545.5555556 | 486.8253968 | 0.98172 |
| HMGCR | chr5 | 74624308 | 31 | 32 | 2969.160709 | 3065.938706 | 0.98178 |
| LCK | chr1 | 32741535 | 32 | 32 | 3043.631723 | 3042.046365 | 0.9818 |
| AKT2 | chr19 | 40794243 | 32 | 32 | 2834.690912 | 2833.272876 | 0.9818 |
| NOS3 | chr7 | 150679622 | 7 | 9 | 276.4285714 | 357.3412699 | 0.98198 |
| LPA | chr6 | 160945168 | 21 | 23 | 622.346923 | 683.3038022 | 0.9822 |
| FOXP3 | chrX | 49118041 | 33 | 27 | 2988.678209 | 2446.954908 | 0.98248 |
| APOA1 | chr11 | 116664423 | 32 | 32 | 3151.968212 | 3153.202181 | 0.98256 |
| APOA1 | chr11 | 116701445 | 34 | 31 | 2726.819788 | 2484.021972 | 0.98262 |
| FOXP3 | chrX | 49128823 | 31 | 31 | 2781.75201 | 2783.116467 | 0.983 |
| DHFR | chr5 | 79958384 | 12 | 10 | 1147.222222 | 956.6666667 | 0.983 |
| APOA1 | chr11 | 116675698 | 35 | 32 | 2928.831182 | 2679.054242 | 0.98318 |
| APOA1 | chr11 | 116667117 | 19 | 23 | 1648.203463 | 1993.629426 | 0.98328 |
| NOS3 | chr7 | 150679767 | 31 | 29 | 2836.501056 | 2651.950035 | 0.9833 |
| FOXP3 | chrX | 49118383 | 31 | 29 | 2944.337802 | 2755.633811 | 0.98334 |
| LDLR | chr19 | 11195139 | 9 | 3 | 735.2380952 | 245.7142857 | 0.9834 |
| CDKN2B-AS1 | chr9 | 21990253 | 30 | 26 | 958.7854112 | 832.3607863 | 0.9838 |
| APOA1 | chr11 | 116667156 | 29 | 31 | 2575.439079 | 2755.204379 | 0.98384 |

| | | | | | | | |
|---|---|---|---|---|---|---|---|
| E2F6 | chr2 | 11608430 | 18 | 18 | 1279.246375 | 1276.143024 | 0.98398 |
| CDKN2B-AS1 | chr9 | 21990046 | 15 | 12 | 538.0952381 | 431.8922744 | 0.984 |
| CDKN1A | chr6 | 36644220 | 33 | 33 | 2203.633756 | 2206.377281 | 0.98456 |
| ESR1 | chr6 | 152009575 | 33 | 32 | 2963.620784 | 2872.709726 | 0.98472 |
| AMOTL2 | chr3 | 134098695 | 12 | 12 | 938.3910534 | 936.504329 | 0.98504 |
| LCK | chr1 | 32740365 | 27 | 30 | 2477.992914 | 2751.90042 | 0.98518 |
| ADRA2C | chr4 | 3766879 | 30 | 28 | 2790.20091 | 2602.609711 | 0.98524 |
| CDKN2B-AS1 | chr9 | 21987473 | 15 | 15 | 448.1313131 | 449.1663892 | 0.98524 |
| CDKN2B-AS1 | chr9 | 22009250 | 3 | 3 | 75.55555555 | 75.84415584 | 0.98532 |
| ABCA1 | chr9 | 107693975 | 10 | 15 | 967.5 | 1450 | 0.98536 |
| ALDOA | chr16 | 30078162 | 21 | 25 | 687.5314092 | 820.1395701 | 0.98562 |
| CDKN1A | chr6 | 36654542 | 19 | 20 | 1798.888889 | 1894.559885 | 0.98566 |
| ADRA2C | chr4 | 3762647 | 36 | 32 | 3058.810645 | 2720.06296 | 0.98592 |
| TFPI | chr2 | 188343466 | 34 | 32 | 2830.564441 | 2662.418223 | 0.986 |
| F7 | chr13 | 113750307 | 31 | 29 | 2714.71407 | 2541.153585 | 0.98622 |
| TERT | chr5 | 1298276 | 32 | 32 | 3013.892444 | 3012.808492 | 0.98624 |
| LDLR | chr19 | 11216112 | 28 | 31 | 2612.984482 | 2891.778499 | 0.9863 |
| LPA | chr6 | 160952714 | 38 | 33 | 1940.614558 | 1686.663419 | 0.9864 |
| CDKN1A | chr6 | 36651783 | 33 | 33 | 2752.788424 | 2751.237226 | 0.98642 |
| ABCA1 | chr9 | 107693980 | 5 | 10 | 469.047619 | 937.1428571 | 0.98648 |
| AMOTL2 | chr3 | 134102249 | 29 | 30 | 2574.651239 | 2664.758297 | 0.98686 |
| LCK | chr1 | 32713140 | 32 | 32 | 3063.702687 | 3062.696232 | 0.98702 |
| COL15A1 | chr9 | 101700190 | 35 | 33 | 2433.651797 | 2293.145557 | 0.98742 |
| F7 | chr13 | 113759087 | 31 | 33 | 1756.989921 | 1871.978741 | 0.9875 |
| APOA1 | chr11 | 116661101 | 35 | 32 | 3323.903739 | 3038.06303 | 0.988 |
| ICAM2 | chr17 | 62085412 | 33 | 32 | 3082.688531 | 2990.086805 | 0.98816 |
| TMPO | chr12 | 98907491 | 37 | 32 | 3497.837876 | 3025.650126 | 0.9887 |
| F7 | chr13 | 113757227 | 19 | 21 | 1092.34127 | 1208.984349 | 0.98872 |
| TIMP3 | chr22 | 33192087 | 33 | 32 | 1311.005708 | 1269.727947 | 0.98876 |
| CDKN1A | chr6 | 36652995 | 34 | 32 | 3179.275495 | 2991.732972 | 0.98882 |
| BAX | chr19 | 49453826 | 21 | 25 | 1857.82967 | 2209.962815 | 0.98882 |
| TERT | chr5 | 1296443 | 35 | 32 | 3196.399451 | 2921.646733 | 0.98894 |
| ARHGEF18 | chr19 | 7461331 | 22 | 22 | 2035.283883 | 2034.420996 | 0.98896 |
| TIMP3 | chr22 | 33195655 | 33 | 32 | 3014.085572 | 2921.776642 | 0.9892 |
| P2RY12 | chr3 | 151112407 | 37 | 33 | 2937.158239 | 2620.826899 | 0.9892 |
| F7 | chr13 | 113753844 | 35 | 32 | 2965.517934 | 2712.583339 | 0.98924 |
| LCK | chr1 | 32741158 | 20 | 26 | 797.7306027 | 1038.706359 | 0.98934 |
| ABCA1 | chr9 | 107690730 | 9 | 12 | 341.6666667 | 454.6536796 | 0.98946 |
| KLF2 | chr19 | 16427233 | 34 | 29 | 1284.055323 | 1096.281566 | 0.9897 |
| APOA1 | chr11 | 116672016 | 30 | 32 | 2509.97658 | 2678.776379 | 0.9898 |
| CDKN2B-AS1 | chr9 | 21981411 | 33 | 32 | 2697.693829 | 2614.9767 | 0.9898 |
| ALOX15 | chr17 | 4551739 | 24 | 23 | 2140.064103 | 2049.7114 | 0.98988 |
| PECAM1 | chr17 | 62409585 | 27 | 23 | 987.5320422 | 842.2849034 | 0.98994 |
| CDKN2B-AS1 | chr9 | 22000824 | 2 | 3 | 53.57142857 | 80 | 0.98996 |
| APOA1 | chr11 | 116668675 | 32 | 31 | 3003.295119 | 2908.698637 | 0.99016 |
| F2RL3 | chr19 | 16994209 | 32 | 32 | 3055.230149 | 3055.719303 | 0.9904 |
| LOXL1 | chr15 | 74244340 | 30 | 32 | 1445.703747 | 1543.663349 | 0.99066 |

| | | | | | | | |
|---|---|---|---|---|---|---|---|
| LOXL1 | chr15 | 74212301 | 33 | 32 | 3149.962109 | 3053.980803 | 0.99074 |
| APOA1 | chr11 | 116682995 | 6 | 2 | 473.3333333 | 157.1428571 | 0.9908 |
| F2RL3 | chr19 | 16990444 | 31 | 30 | 2904.513875 | 2810.050244 | 0.99096 |
| LDLR | chr19 | 11195083 | 25 | 21 | 2226.897547 | 1871.349206 | 0.99104 |
| CDKN2B-AS1 | chr9 | 21970976 | 15 | 12 | 496.8426018 | 396.7568543 | 0.99108 |
| CDKN2B-AS1 | chr9 | 21973422 | 32 | 31 | 1088.048678 | 1054.787508 | 0.99114 |
| LOXL1 | chr15 | 74218921 | 28 | 25 | 950.508003 | 849.6411808 | 0.99116 |
| BCL2 | chr18 | 60994697 | 38 | 32 | 3595.353239 | 3027.325847 | 0.99124 |
| CDKN1A | chr6 | 36643461 | 13 | 10 | 1255 | 965.7142857 | 0.99138 |
| LOXL1 | chr15 | 74219232 | 28 | 21 | 1122.402946 | 842.7473648 | 0.99156 |
| F7 | chr13 | 113753600 | 38 | 32 | 3240.643679 | 2729.612662 | 0.9917 |
| CTGF | chr6 | 132278725 | 39 | 32 | 3559.914163 | 2921.398343 | 0.9917 |
| LCK | chr1 | 32741970 | 29 | 26 | 2799.014042 | 2509.920635 | 0.99208 |
| NR3C1 | chr5 | 142789490 | 38 | 33 | 2965.125694 | 2574.165052 | 0.99232 |
| CDKN1A | chr6 | 36645608 | 9 | 7 | 238.1746032 | 185.497076 | 0.99256 |
| APOA1 | chr11 | 116702630 | 33 | 31 | 3136.017316 | 2945.231647 | 0.99272 |
| P2RY12 | chr3 | 151107811 | 35 | 32 | 3289.162695 | 3006.805943 | 0.99276 |
| CDKN2B-AS1 | chr9 | 21970986 | 10 | 10 | 323.4271284 | 322.9204129 | 0.99294 |
| GALNT2 | chr1 | 230187005 | 14 | 17 | 1136.520147 | 1380.750361 | 0.99308 |
| LDLR | chr19 | 11192547 | 31 | 26 | 2939.456018 | 2464.656234 | 0.99318 |
| LPA | chr6 | 11222780 | 21 | 27 | 748.8047099 | 961.9644496 | 0.9934 |
| ADRA2C | chr4 | 3759533 | 32 | 32 | 2602.748998 | 2603.437458 | 0.99352 |
| LOXL1 | chr15 | 74214969 | 29 | 29 | 1391.298598 | 1390.350607 | 0.99358 |
| ADRA2C | chr4 | 3761473 | 28 | 29 | 2275.971353 | 2356.379095 | 0.99358 |
| LPA | chr6 | 160952838 | 32 | 31 | 2281.627049 | 2208.970462 | 0.9937 |
| APOA1 | chr11 | 116679555 | 35 | 32 | 3064.049611 | 2801.934747 | 0.99384 |
| ADRA2C | chr4 | 3760461 | 34 | 30 | 2421.888434 | 2138.506218 | 0.9939 |
| NOS3 | chr7 | 150687599 | 32 | 32 | 2934.838287 | 2935.396784 | 0.9939 |
| ICAM2 | chr17 | 62087470 | 35 | 32 | 3344.280395 | 3057.897179 | 0.99464 |
| APOA1 | chr11 | 116677515 | 33 | 32 | 2878.00383 | 2791.391068 | 0.99476 |
| APOA1 | chr11 | 116679366 | 27 | 26 | 2519.052075 | 2425.259104 | 0.99482 |
| LCK | chr1 | 32740957 | 25 | 30 | 1332.565601 | 1600.028583 | 0.99484 |
| TMPO | chr12 | 98944098 | 37 | 32 | 3510.154702 | 3035.983877 | 0.99492 |
| ARHGEF18 | chr19 | 7469755 | 33 | 32 | 3175.579231 | 3079.612102 | 0.995 |
| APOA1 | chr11 | 116689654 | 25 | 29 | 2321.246074 | 2692.142857 | 0.99504 |
| PECAM1 | chr17 | 62397533 | 32 | 33 | 2347.191363 | 2419.88382 | 0.99554 |
| DHFR | chr5 | 79957277 | 35 | 32 | 3367.120877 | 3078.334638 | 0.99568 |
| ARHGEF18 | chr19 | 7469773 | 36 | 32 | 3374.202693 | 2999.555191 | 0.99574 |
| F2RL3 | chr19 | 16990512 | 29 | 31 | 2752.045911 | 2941.20671 | 0.99598 |
| TIMP3 | chr22 | 33187640 | 35 | 29 | 3014.67305 | 2497.289397 | 0.99668 |
| APOA1 | chr11 | 116681243 | 24 | 24 | 2359.775698 | 2360.012626 | 0.99674 |
| ALDOA | chr16 | 30057644 | 22 | 26 | 1858.482504 | 2196.008191 | 0.99684 |
| KLF2 | chr19 | 16430191 | 20 | 19 | 1728.809524 | 1642.65873 | 0.99722 |
| NOS3 | chr7 | 150706545 | 11 | 11 | 1053.333333 | 1053.214286 | 0.99736 |
| F7 | chr13 | 113752444 | 33 | 32 | 3058.765702 | 2965.91088 | 0.99738 |
| HMGCR | chr5 | 74631296 | 33 | 32 | 3119.031589 | 3024.683928 | 0.9974 |
| TIMP3 | chr22 | 33195657 | 33 | 32 | 3086.213048 | 2992.977882 | 0.99744 |

| | | | | | | | |
|---|---|---|---|---|---|---|---|
| APOA1 | chr11 | 116679835 | 33 | 32 | 2886.726742 | 2799.081198 | 0.99756 |
| ESR2 | chr14 | 64808703 | 40 | 32 | 3383.853981 | 2707.248669 | 0.99786 |
| LPA | chr6 | 160952893 | 33 | 33 | 2743.210215 | 2742.967225 | 0.99786 |
| F7 | chr13 | 113759592 | 32 | 32 | 2176.14144 | 2176.400972 | 0.99806 |
| FOXP3 | chrX | 49126999 | 20 | 13 | 1540.396825 | 1001.428571 | 0.99834 |
| TERT | chr5 | 1295656 | 4 | 5 | 103.3333333 | 129.1269841 | 0.9984 |
| ESR1 | chr6 | 152006733 | 36 | 32 | 3345.30508 | 2973.690806 | 0.9987 |
| FGF2 | chr4 | 123747533 | 29 | 29 | 1311.467015 | 1311.593615 | 0.99898 |
| BAX | chr19 | 49465134 | 32 | 32 | 2961.768399 | 2961.871382 | 0.9991 |
| TERT | chr5 | 1301070 | 33 | 32 | 3112.633925 | 3018.247368 | 0.99922 |
| ATF3 | chr1 | 212745305 | 8 | 9 | 780 | 877.5 | 1 |
| APOA1 | chr11 | 116712726 | 4 | 2 | 280 | 140 | 1 |
| TIMP3 | chr22 | 33190506 | 2 | 3 | 80 | 120 | 1 |
| TFPI | chr2 | 188425120 | 3 | 2 | 275 | 183.3333333 | 1 |
| TFPI | chr2 | 188424633 | 7 | 3 | 653.3333333 | 280 | 1 |
