## Supplementary Table 4 for "Investigating Coronary Artery Disease methylome through targeted bisulfite sequencing"

| Supplementary table 4 is a list of all the significantly differentially methylated regions identified from our study between controls and CAD subjects | | | | | | | |
|---|---|---|---|---|---|---|---|
| Gene | Chromosome | coordinate | No. of controls | No of cases | Control methylation count | Case methylation count | p value |
| APOA1 | chr11 | 116661248 | 31 | 32 | 2660.90125 | 3028.46628 | 0.000135 |
| PLA2G7 | chr6 | 46711700 | 35 | 32 | 2907.244716 | 2835.20577 | 0.000762 |
| FOXP3 | chrX | 49121289 | 21 | 18 | 1679.958791 | 1132.435065 | 0.001339 |
| CTGF | chr6 | 132279198 | 30 | 27 | 2813.331448 | 2168.752184 | 0.001641 |
| ICAM2 | chr17 | 62086376 | 30 | 33 | 1700.235569 | 1504.472239 | 0.001644 |
| CDKN1A | chr6 | 36651933 | 31 | 31 | 2987.672346 | 2805.204288 | 0.001662 |
| F7 | chr13 | 113756669 | 34 | 32 | 2508.381418 | 2579.3089 | 0.001924 |
| ABCG1 | chr21 | 43629820 | 37 | 32 | 3522.277779 | 2874.058612 | 0.002175 |
| PHACTR1 | chr6 | 12717192 | 2 | 4 | 200 | 331.4285714 | 0.002279 |
| NOS3 | chr7 | 150689700 | 26 | 30 | 1769.570013 | 2434.986921 | 0.002443 |
| GCK | chr7 | 44229968 | 10 | 19 | 419.4540708 | 1207.30464 | 0.002561 |
| KLF2 | chr19 | 16438034 | 29 | 30 | 972.958874 | 1288.309009 | 0.002597 |
| TFPI | chr2 | 188424480 | 22 | 22 | 2168.293651 | 1981.067821 | 0.002662 |
| LDLR | chr19 | 11221008 | 27 | 25 | 2269.989557 | 2338.599179 | 0.002838 |
| F2RL3 | chr19 | 16998127 | 17 | 15 | 1429.184982 | 996.0714286 | 0.002919 |
| APOA1 | chr11 | 116687627 | 32 | 32 | 2963.553796 | 3090.157093 | 0.003036 |
| KLF2 | chr19 | 16433304 | 35 | 32 | 3457.780687 | 2977.12297 | 0.003063 |
| TFPI | chr2 | 188420267 | 39 | 32 | 3004.300012 | 2689.91039 | 0.00308 |
| F7 | chr13 | 113755287 | 32 | 31 | 2949.859307 | 3033.300547 | 0.003337 |
| ADRA2C | chr4 | 3765853 | 18 | 17 | 1330.079365 | 1490.386003 | 0.003343 |
| ADRA2C | chr4 | 3766721 | 25 | 25 | 1350.293651 | 1672.112724 | 0.003424 |
| LDLR | chr19 | 11192639 | 22 | 20 | 2151.742424 | 1763.095238 | 0.003563 |
| GCK | chr7 | 44231646 | 9 | 19 | 572.1753247 | 1612.608225 | 0.003605 |
| HMGCR | chr5 | 74628717 | 31 | 30 | 2588.22911 | 2730.822788 | 0.003819 |
| ABCG1 | chr21 | 43717591 | 37 | 32 | 3104.243264 | 2890.112966 | 0.004575 |
| LDLR | chr19 | 11221692 | 31 | 29 | 2851.185835 | 2812.255157 | 0.00458 |
| CDKN2B-AS | chr9 | 22005889 | 25 | 27 | 920.4695867 | 1313.69129 | 0.004857 |
| CDKN2B-AS | chr9 | 21989468 | 24 | 26 | 809.5659048 | 1140.979321 | 0.004939 |
| LPA | chr6 | 160946112 | 32 | 32 | 3145.479996 | 2953.917403 | 0.005015 |
| APOA1 | chr11 | 116723854 | 25 | 21 | 739.7567362 | 814.7053852 | 0.0059 |
| ALOX15 | chr17 | 4553160 | 33 | 33 | 1574.973078 | 1308.38947 | 0.006322 |
| APOA1 | chr11 | 116697880 | 31 | 30 | 2920.575064 | 2676.844966 | 0.006435 |
| CDKN2B-AS | chr9 | 22009886 | 32 | 30 | 1756.675522 | 1369.526065 | 0.006622 |
| APOA1 | chr11 | 116701833 | 31 | 32 | 3050.047796 | 2739.742382 | 0.006639 |
| F7 | chr13 | 113750128 | 33 | 31 | 3204.001114 | 2826.589638 | 0.00664 |
| KLF2 | chr19 | 16438050 | 33 | 32 | 1347.652719 | 1591.862541 | 0.006742 |
| LCK | chr1 | 32739493 | 12 | 14 | 782.9691877 | 592.3412698 | 0.006759 |
| NOS3 | chr7 | 150694867 | 32 | 31 | 2749.596914 | 2861.04145 | 0.007161 |
| E2F6 | chr2 | 11613547 | 35 | 32 | 3322.792507 | 2931.948827 | 0.008323 |
| APOA1 | chr11 | 116696636 | 33 | 32 | 2604.42204 | 2694.135944 | 0.008741 |
| P2RY12 | chr3 | 151111174 | 19 | 11 | 1541.666667 | 672.0634921 | 0.008903 |
| ADRA2C | chr4 | 3761998 | 26 | 27 | 2479.91009 | 2369.89956 | 0.009325 |
| TIMP3 | chr22 | 33190664 | 32 | 33 | 2696.616007 | 2495.694229 | 0.009764 |
| TMPO | chr12 | 98900752 | 39 | 32 | 3833.655242 | 3079.942353 | 0.009896 |
| ABCA1 | chr9 | 107691120 | 15 | 11 | 1402.261905 | 1100 | 0.009979 |
| PLA2G7 | chr6 | 46713047 | 40 | 32 | 3390.057951 | 2841.381957 | 0.010035 |

| | | | | | | | |
|---|---|---|---|---|---|---|---|
| LDLR | chr19 | 11219442 | 34 | 32 | 3208.279783 | 3129.765784 | 0.010242 |
| CDKN2B-A | chr9 | 21989845 | 20 | 19 | 709.6549529 | 943.1349206 | 0.010339 |
| CDKN2B-A | chr9 | 22001063 | 33 | 33 | 2125.792617 | 1811.839638 | 0.010476 |
| NOS3 | chr7 | 150686444 | 30 | 33 | 1385.473035 | 1764.71109 | 0.011356 |
| LDLR | chr19 | 11197536 | 16 | 16 | 1533.055556 | 1417.380952 | 0.011657 |
| ESR1 | chr6 | 152010481 | 38 | 32 | 3517.55283 | 3046.173451 | 0.011904 |
| NOS3 | chr7 | 150683826 | 33 | 32 | 2733.188019 | 2912.199969 | 0.011959 |
| AMOTL2 | chr3 | 134098914 | 35 | 33 | 2860.117139 | 2481.069369 | 0.012019 |
| TERT | chr5 | 1301170 | 34 | 32 | 3139.173343 | 3076.933869 | 0.012343 |
| F7 | chr13 | 113756188 | 34 | 33 | 2418.248227 | 2133.7321 | 0.012345 |
| PECAM1 | chr17 | 62399742 | 14 | 22 | 1342.853535 | 1919.920635 | 0.012423 |
| GCK | chr7 | 44228831 | 36 | 32 | 3098.395854 | 2896.82119 | 0.012544 |
| ARHGEF18 | chr19 | 7461921 | 33 | 32 | 3189.298143 | 2770.217195 | 0.01274 |
| LDLR | chr19 | 11217939 | 30 | 29 | 2826.561129 | 2862.804516 | 0.012781 |
| KLF2 | chr19 | 16437980 | 31 | 28 | 1202.112436 | 1340.635794 | 0.012916 |
| CDKN2B-A | chr9 | 21968470 | 7 | 6 | 226.9047619 | 141.3071895 | 0.013315 |
| F2RL3 | chr19 | 16991169 | 27 | 26 | 2226.719033 | 2388.613609 | 0.013404 |
| ATF3 | chr1 | 212793952 | 37 | 32 | 3039.401717 | 2766.558684 | 0.013437 |
| LDLR | chr19 | 11221697 | 30 | 29 | 2759.691503 | 2816.666667 | 0.013457 |
| CDKN1A | chr6 | 36651959 | 30 | 31 | 2987.487513 | 2964.535595 | 0.0138 |
| AMOTL2 | chr3 | 134095282 | 32 | 32 | 2722.255284 | 2891.898575 | 0.014318 |
| FOXP3 | chrX | 49121784 | 16 | 9 | 1416.944444 | 885.7142857 | 0.014345 |
| F7 | chr13 | 113760587 | 8 | 2 | 510 | 200 | 0.014441 |
| APOA1 | chr11 | 116708414 | 32 | 30 | 2663.34869 | 2064.316386 | 0.014458 |
| LDLR | chr19 | 11199718 | 34 | 32 | 1827.30144 | 1506.978573 | 0.014459 |
| F7 | chr13 | 113750505 | 25 | 24 | 2339.484127 | 2373.333333 | 0.014679 |
| ICAM2 | chr17 | 62087479 | 34 | 32 | 3246.325922 | 3129.406038 | 0.014861 |
| TERT | chr5 | 1299860 | 12 | 17 | 731.0436788 | 772.6679879 | 0.014901 |
| CTGF | chr6 | 132274760 | 34 | 32 | 3138.349677 | 3068.303241 | 0.015203 |
| ESR1 | chr6 | 152002099 | 30 | 33 | 2880.262853 | 2940.172458 | 0.015664 |
| F2RL3 | chr19 | 16998196 | 17 | 9 | 752.2763348 | 253.7301587 | 0.015737 |
| LDLR | chr19 | 11224018 | 23 | 22 | 2089.569597 | 2133.360071 | 0.016217 |
| TERT | chr5 | 1299447 | 24 | 27 | 2068.101343 | 2548.134921 | 0.016643 |
| F2RL3 | chr19 | 16994756 | 32 | 31 | 2798.203341 | 2907.305539 | 0.016859 |
| ADRA2C | chr4 | 3760991 | 32 | 29 | 2853.639971 | 2766.175691 | 0.017136 |
| MCPH1 | chr8 | 6470832 | 34 | 31 | 3126.432337 | 2985.708171 | 0.017237 |
| COL15A1 | chr9 | 101698014 | 34 | 32 | 2895.740938 | 2866.269512 | 0.01724 |
| APOA1 | chr11 | 116687933 | 32 | 31 | 3033.537113 | 3034.001823 | 0.017259 |
| CDKN2B-A | chr9 | 21977554 | 13 | 10 | 429.1666667 | 515.4761905 | 0.017263 |
| ICAM2 | chr17 | 62080221 | 28 | 29 | 2693.326118 | 2879.79798 | 0.017375 |
| CDKN2B-A | chr9 | 22000964 | 33 | 33 | 2475.752006 | 2130.642638 | 0.017596 |
| TERT | chr5 | 1296463 | 34 | 33 | 2793.653158 | 2503.444008 | 0.01782 |
| APOA1 | chr11 | 116691928 | 32 | 32 | 2909.239159 | 3049.794166 | 0.01786 |
| LDLR | chr19 | 11216032 | 33 | 31 | 3082.045533 | 3012.098074 | 0.018164 |
| APOA1 | chr11 | 116725861 | 34 | 33 | 1838.142246 | 1550.56936 | 0.018762 |
| NOS3 | chr7 | 150679361 | 38 | 33 | 2086.237291 | 2000.576065 | 0.018976 |
| KLF2 | chr19 | 16433445 | 34 | 32 | 3281.516637 | 3004.594136 | 0.019297 |
| ATF3 | chr1 | 212739706 | 27 | 28 | 2472.885496 | 2265.885226 | 0.019381 |
| DHFR | chr5 | 79959441 | 20 | 19 | 1825.119048 | 1853.888889 | 0.019419 |
| ADRA2C | chr4 | 3764755 | 31 | 31 | 1899.559484 | 1592.950393 | 0.019441 |
| CDKN2B-A | chr9 | 21977451 | 29 | 25 | 1657.143918 | 1171.013709 | 0.019576 |

| | | | | | | | |
|---|---|---|---|---|---|---|---|
| APOA1 | chr11 | 116661216 | 31 | 32 | 2840.256015 | 3067.811663 | 0.019582 |
| ARHGEF18 | chr19 | 7464099 | 33 | 32 | 3039.025467 | 2763.302993 | 0.019779 |
| COL15A1 | chr9 | 101704337 | 35 | 32 | 3146.366722 | 2995.574006 | 0.019798 |
| PHACTR1 | chr6 | 12715861 | 25 | 28 | 1865.986455 | 2344.940476 | 0.02086 |
| ADRA2C | chr4 | 3766521 | 27 | 29 | 2386.282096 | 2337.967178 | 0.021116 |
| F2RL3 | chr19 | 16990296 | 30 | 31 | 2825.721092 | 3038.696581 | 0.021201 |
| APOA1 | chr11 | 116721465 | 33 | 32 | 3258.827229 | 2864.249855 | 0.021241 |
| F2RL3 | chr19 | 16998949 | 32 | 33 | 2733.366386 | 2608.380942 | 0.021284 |
| BCL2 | chr18 | 60995889 | 22 | 16 | 1932.832168 | 1531.309524 | 0.021624 |
| CDKN2B-AS | chr9 | 21978376 | 36 | 33 | 1317.189357 | 1388.007684 | 0.021637 |
| F7 | chr13 | 113750103 | 32 | 31 | 3067.717763 | 2854.622719 | 0.021658 |
| APOA1 | chr11 | 116725636 | 12 | 9 | 1117.575758 | 900 | 0.021702 |
| TMPO | chr12 | 98906185 | 33 | 32 | 3142.253761 | 2900.953057 | 0.021716 |
| LDLR | chr19 | 11192553 | 31 | 26 | 3034.931041 | 2375.40293 | 0.021903 |
| TMPO | chr12 | 98900798 | 41 | 32 | 3947.5111 | 3017.745819 | 0.022145 |
| CDKN2B-AS | chr9 | 21974079 | 5 | 3 | 130.7936508 | 130 | 0.022356 |
| F7 | chr13 | 113750985 | 30 | 23 | 2697.478355 | 2198.94958 | 0.022797 |
| APOA1 | chr11 | 116704450 | 33 | 33 | 2969.798386 | 2758.633113 | 0.023039 |
| MIA3 | chr1 | 222808651 | 37 | 32 | 3370.595736 | 3006.529588 | 0.023619 |
| LCK | chr1 | 32740607 | 15 | 14 | 1333.607504 | 1357.777778 | 0.023838 |
| FOXP3 | chrX | 49116085 | 24 | 30 | 2105.413059 | 2857.893218 | 0.023857 |
| TERT | chr5 | 1295754 | 27 | 21 | 888.1714169 | 856.9523768 | 0.02406 |
| APOA1 | chr11 | 116690546 | 31 | 33 | 2078.259868 | 2488.141553 | 0.024383 |
| CTGF | chr6 | 132273127 | 30 | 32 | 2591.790324 | 2497.287102 | 0.024401 |
| APOA1 | chr11 | 116714317 | 37 | 32 | 3526.825451 | 2985.376127 | 0.024481 |
| MCPH1 | chr8 | 6470178 | 19 | 23 | 1888.649684 | 2058.441558 | 0.024619 |
| ICAM2 | chr17 | 62091363 | 32 | 32 | 2970.289663 | 3124.852941 | 0.024756 |
| NR3C1 | chr5 | 142788918 | 36 | 32 | 3378.233411 | 2922.490197 | 0.024763 |
| ESR2 | chr14 | 64806245 | 37 | 33 | 3177.031669 | 2643.560749 | 0.025124 |
| ABCB1 | chr7 | 87343538 | 26 | 22 | 2420.627706 | 2179.807692 | 0.025417 |
| FOXP3 | chrX | 49115016 | 13 | 5 | 1055.238095 | 293.3333333 | 0.025441 |
| CDKN2B-AS | chr9 | 22005101 | 20 | 13 | 516.2954177 | 466.8903319 | 0.025643 |
| LDLR | chr19 | 11221389 | 34 | 32 | 3224.642383 | 3127.543572 | 0.026705 |
| APOA1 | chr11 | 116661335 | 30 | 32 | 2601.99794 | 2940.67472 | 0.026981 |
| ATF3 | chr1 | 212745495 | 32 | 32 | 3077.167157 | 2967.421722 | 0.027035 |
| LOXL1 | chr15 | 74210831 | 36 | 32 | 3271.388545 | 2998.377202 | 0.027319 |
| LOXL1 | chr15 | 74209559 | 25 | 23 | 2365.436508 | 2273.214286 | 0.027377 |
| CTGF | chr6 | 132280299 | 35 | 31 | 3200.148177 | 2708.951833 | 0.028042 |
| APOA1 | chr11 | 116678334 | 17 | 18 | 1571.190476 | 1757.142857 | 0.028059 |
| LDLR | chr19 | 11224057 | 24 | 18 | 2221.309524 | 1783.333333 | 0.028181 |
| CDKN2B-AS | chr9 | 21991667 | 37 | 33 | 2792.539068 | 2318.549655 | 0.028361 |
| FOXP3 | chrX | 49121276 | 21 | 21 | 1745.436785 | 1524.576012 | 0.028478 |
| APOA1 | chr11 | 116708556 | 29 | 30 | 2772.811772 | 2599.416997 | 0.028623 |
| TFPI | chr2 | 188426841 | 24 | 21 | 2076.395467 | 1970.093158 | 0.028664 |
| LCK | chr1 | 32741484 | 25 | 25 | 2211.498779 | 1961.381674 | 0.028778 |
| CDKN2B-AS | chr9 | 21990105 | 13 | 18 | 632.089169 | 642.1184371 | 0.028899 |
| LDLR | chr19 | 11221486 | 34 | 32 | 3259.867609 | 3137.146419 | 0.029659 |
| APOA1 | chr11 | 116691961 | 32 | 32 | 2951.202466 | 3088.256473 | 0.030179 |
| APOA1 | chr11 | 116708518 | 31 | 31 | 3021.752315 | 2924.063649 | 0.030359 |
| PECAM1 | chr17 | 62398226 | 36 | 33 | 3089.381442 | 2656.854054 | 0.030424 |
| ABCA1 | chr9 | 107542842 | 34 | 32 | 2828.844758 | 2451.085611 | 0.030518 |

| | | | | | | | |
|---|---|---|---|---|---|---|---|
| FGF2 | chr4 | 123741393 | 35 | 32 | 3285.559922 | 2916.456135 | 0.030539 |
| LOXL1 | chr15 | 74218620 | 15 | 19 | 494.8276723 | 890.0968476 | 0.030599 |
| TIMP3 | chr22 | 33189151 | 30 | 30 | 2039.994054 | 1806.386559 | 0.030895 |
| AKT2 | chr19 | 40794901 | 33 | 32 | 3230.521136 | 3041.45447 | 0.030959 |
| APOA1 | chr11 | 116691951 | 32 | 32 | 2959.625112 | 3106.468431 | 0.031043 |
| LOXL1 | chr15 | 74211401 | 35 | 32 | 3174.697564 | 2997.191084 | 0.031402 |
| CTGF | chr6 | 132272983 | 35 | 33 | 2325.489793 | 2037.538089 | 0.03144 |
| ALDOA | chr16 | 30061597 | 23 | 27 | 2184.629815 | 2399.73693 | 0.032084 |
| LOXL1 | chr15 | 74211612 | 14 | 11 | 1253.650794 | 1064.880952 | 0.0323 |
| CDKN1A | chr6 | 36654680 | 32 | 32 | 3071.608865 | 2959.694733 | 0.032456 |
| NOS3 | chr7 | 150695647 | 23 | 22 | 2249.047619 | 2046.226551 | 0.032844 |
| FOXP3 | chrX | 49130161 | 31 | 31 | 1785.234519 | 2004.693918 | 0.033135 |
| ESR1 | chr6 | 152007888 | 27 | 24 | 2148.986985 | 1585.078059 | 0.033343 |
| TERT | chr5 | 1253550 | 31 | 31 | 2679.064516 | 2879.627458 | 0.033582 |
| FOXP3 | chrX | 49126877 | 7 | 5 | 200.8333333 | 220 | 0.033758 |
| DHFR | chr5 | 79954125 | 6 | 4 | 440.7142857 | 385.7142857 | 0.033943 |
| AMOTL2 | chr3 | 134099761 | 31 | 31 | 3054.586623 | 2800.151876 | 0.034042 |
| ABCG1 | chr21 | 43637024 | 32 | 32 | 2661.951014 | 2836.541639 | 0.034298 |
| LDLR | chr19 | 11217263 | 27 | 31 | 2672.334055 | 2942.353449 | 0.034355 |
| ARHGEF18 | chr19 | 7468074 | 22 | 18 | 2014.761905 | 1759.047619 | 0.034624 |
| FOXP3 | chrX | 49130389 | 29 | 30 | 2507.239403 | 2329.236336 | 0.034699 |
| ADRA2C | chr4 | 3760184 | 35 | 32 | 2794.064131 | 2691.39899 | 0.034985 |
| APOA1 | chr11 | 116705516 | 21 | 24 | 1205.323843 | 1761.111111 | 0.035037 |
| APOA1 | chr11 | 116699075 | 19 | 23 | 1395.766595 | 1953.044733 | 0.035517 |
| APOA1 | chr11 | 116687953 | 32 | 32 | 3021.990188 | 3102.453819 | 0.035637 |
| CTGF | chr6 | 132280534 | 30 | 31 | 2032.789973 | 2348.484965 | 0.035764 |
| KLF2 | chr19 | 16426847 | 7 | 7 | 507.0238095 | 603.3333333 | 0.035802 |
| PHACTR1 | chr6 | 12717341 | 33 | 33 | 3016.100463 | 2839.434731 | 0.035836 |
| APOA1 | chr11 | 116674643 | 28 | 32 | 2778.825439 | 3021.018483 | 0.035998 |
| LPA | chr6 | 160950075 | 32 | 32 | 2930.8083 | 3040.222983 | 0.036044 |
| ABCG1 | chr21 | 43634598 | 28 | 31 | 2320.897034 | 2793.272283 | 0.036222 |
| GCK | chr7 | 44231431 | 14 | 14 | 1106.547619 | 1265 | 0.036557 |
| ARHGEF18 | chr19 | 7461329 | 24 | 22 | 2206.071429 | 2137.142857 | 0.037057 |
| LDLR | chr19 | 11199222 | 32 | 31 | 2670.389903 | 2745.749826 | 0.037064 |
| NOS3 | chr7 | 150705831 | 28 | 27 | 969.4632936 | 1092.170761 | 0.03708 |
| ABCB1 | chr7 | 87342828 | 32 | 31 | 2137.987643 | 2267.592598 | 0.037222 |
| ABCB1 | chr7 | 87350728 | 33 | 32 | 2823.381757 | 2503.065822 | 0.037437 |
| FGF2 | chr4 | 123746991 | 22 | 19 | 2000.125153 | 1834.040404 | 0.037899 |
| CDKN1A | chr6 | 36652302 | 33 | 32 | 3010.647734 | 3047.186878 | 0.038061 |
| APOA1 | chr11 | 116681284 | 23 | 23 | 2258.75 | 2163.883061 | 0.038299 |
| ABCA1 | chr9 | 107691631 | 38 | 32 | 3496.528734 | 2829.008842 | 0.038883 |
| FOXP3 | chrX | 49119927 | 32 | 33 | 2677.842477 | 2545.889789 | 0.038942 |
| LOXL1 | chr15 | 74213839 | 32 | 32 | 2613.61453 | 2448.185977 | 0.0391 |
| GCK | chr7 | 44183776 | 35 | 32 | 3230.062136 | 3035.631651 | 0.039704 |
| ALOX15 | chr17 | 4554325 | 8 | 6 | 676.0606061 | 587.5 | 0.039739 |
| APOA1 | chr11 | 116726424 | 17 | 21 | 528.6580087 | 802.3859963 | 0.039765 |
| ABCB1 | chr7 | 87352198 | 39 | 32 | 3370.126687 | 2857.536804 | 0.039805 |
| E2F6 | chr2 | 11613270 | 16 | 22 | 1471.825397 | 2131.388889 | 0.039821 |
| APOA1 | chr11 | 116692085 | 32 | 32 | 2801.374457 | 2927.095825 | 0.039844 |
| CTGF | chr6 | 132276533 | 33 | 32 | 2560.110867 | 2657.430024 | 0.04004 |
| LDLR | chr19 | 11222323 | 13 | 14 | 1181.428571 | 1359.722222 | 0.040136 |

| | | | | | | | |
|---|---|---|---|---|---|---|---|
| FGF2 | chr4 | 123746873 | 33 | 32 | 3115.965202 | 2903.292319 | 0.040155 |
| APOA1 | chr11 | 116661450 | 30 | 31 | 2305.965641 | 2597.882282 | 0.040162 |
| LDLR | chr19 | 11195038 | 33 | 31 | 3161.633989 | 2852.017377 | 0.040355 |
| NOS3 | chr7 | 150708045 | 30 | 33 | 1770.399749 | 2234.542125 | 0.040882 |
| APOA1 | chr11 | 116701560 | 31 | 28 | 3006.767677 | 2615.18759 | 0.041062 |
| LDLR | chr19 | 11223961 | 23 | 21 | 2172.711039 | 2074.603175 | 0.041117 |
| TERT | chr5 | 1296577 | 32 | 30 | 3125.173395 | 2846.152846 | 0.041422 |
| APOA1 | chr11 | 116708317 | 32 | 32 | 3089.397721 | 2989.079166 | 0.041896 |
| LDLR | chr19 | 11198964 | 23 | 25 | 2117.539683 | 2428.409091 | 0.041985 |
| FOXP3 | chrX | 49106474 | 21 | 23 | 1824.076063 | 1745.952381 | 0.042663 |
| ABCG1 | chr21 | 43635188 | 34 | 32 | 3059.219617 | 3042.813742 | 0.042721 |
| ADRA2C | chr4 | 3762695 | 36 | 32 | 2262.521825 | 2179.978249 | 0.042824 |
| LDLR | chr19 | 11223530 | 11 | 12 | 1100 | 1138.928571 | 0.042998 |
| BAX | chr19 | 49449696 | 31 | 32 | 2905.354111 | 3092.204905 | 0.043163 |
| ADRA2C | chr4 | 3762969 | 28 | 29 | 2392.273005 | 2279.694195 | 0.04326 |
| BAX | chr19 | 49453159 | 25 | 19 | 1886.262626 | 1243.001443 | 0.04328 |
| ABCG1 | chr21 | 43717781 | 37 | 32 | 3544.394002 | 2996.30796 | 0.043396 |
| P2RY12 | chr3 | 151105698 | 31 | 33 | 2771.802538 | 2694.418981 | 0.043457 |
| APOA1 | chr11 | 116692558 | 32 | 32 | 2901.019711 | 3019.623327 | 0.043698 |
| LDLR | chr19 | 11221653 | 32 | 32 | 2862.624706 | 2982.020371 | 0.044098 |
| F2RL3 | chr19 | 16998540 | 2 | 2 | 40 | 61.9047619 | 0.044156 |
| TERT | chr5 | 1295030 | 2 | 2 | 61.9047619 | 40 | 0.044156 |
| ARHGEF18 | chr19 | 7463987 | 32 | 32 | 2549.140648 | 2745.909699 | 0.0442 |
| ADRA2C | chr4 | 3760951 | 31 | 28 | 2495.931013 | 2453.137794 | 0.044382 |
| APOA1 | chr11 | 116715660 | 36 | 32 | 3353.911119 | 3056.259274 | 0.044464 |
| APOA1 | chr11 | 116725440 | 34 | 32 | 3336.79097 | 3011.21156 | 0.044464 |
| MCPH1 | chr8 | 6260206 | 12 | 22 | 1200 | 2152.103175 | 0.044521 |
| APOA1 | chr11 | 116708984 | 28 | 27 | 2650.445906 | 2657.478632 | 0.04508 |
| LCK | chr1 | 32740883 | 22 | 26 | 1440.770063 | 1980.06993 | 0.045196 |
| PECAM1 | chr17 | 62397478 | 32 | 32 | 3124.552596 | 3043.098589 | 0.045539 |
| LCK | chr1 | 32740399 | 26 | 28 | 2518.591921 | 2575.850641 | 0.045604 |
| LPA | chr6 | 11224714 | 2 | 4 | 146.6666667 | 142.1428571 | 0.045681 |
| MCPH1 | chr8 | 6470820 | 33 | 31 | 2985.514881 | 2946.734374 | 0.045939 |
| NOS3 | chr7 | 150695069 | 6 | 4 | 600 | 314.6031746 | 0.045977 |
| LOXL1 | chr15 | 74218777 | 16 | 12 | 406.1105561 | 449.6480643 | 0.046262 |
| MCPH1 | chr8 | 6470110 | 21 | 25 | 2063.214286 | 2337.234432 | 0.046356 |
| APOA1 | chr11 | 116663322 | 34 | 32 | 2985.450879 | 2936.043444 | 0.04636 |
| APOA1 | chr11 | 116724949 | 4 | 3 | 273.8095238 | 283.3333333 | 0.046401 |
| PHACTR1 | chr6 | 12711037 | 33 | 32 | 3027.076802 | 3052.293347 | 0.046483 |
| APOA1 | chr11 | 116676590 | 6 | 4 | 534.7619048 | 270.4761905 | 0.046518 |
| LOXL1 | chr15 | 74218715 | 16 | 17 | 472.056277 | 715.8899434 | 0.046519 |
| ADRA2C | chr4 | 3760295 | 36 | 32 | 3099.450658 | 2872.718022 | 0.04652 |
| AMOTL2 | chr3 | 134097086 | 34 | 32 | 2935.015198 | 2568.723451 | 0.046658 |
| GALNT2 | chr1 | 230188247 | 18 | 15 | 1594.642857 | 1448.333333 | 0.046755 |
| NOS3 | chr7 | 150710683 | 5 | 8 | 158.3333333 | 184.1269841 | 0.046857 |
| AKT2 | chr19 | 40795616 | 13 | 13 | 1300 | 1224.047619 | 0.04686 |
| ADRA2C | chr4 | 3761649 | 34 | 32 | 2961.127215 | 2911.275165 | 0.047017 |
| LDLR | chr19 | 11196237 | 35 | 32 | 3236.037618 | 2814.033387 | 0.047058 |
| GALNT2 | chr1 | 230190310 | 30 | 30 | 2675.920399 | 2858.187923 | 0.047156 |
| APOA1 | chr11 | 116724233 | 33 | 28 | 1905.424417 | 1345.588522 | 0.047204 |
| APOA1 | chr11 | 116661389 | 32 | 32 | 2913.801911 | 3040.75293 | 0.047318 |

| | | | | | | | |
|---|---|---|---|---|---|---|---|
| CDKN1A | chr6 | 36652031 | 29 | 30 | 2663.564214 | 2928.675214 | 0.047403 |
| TFPI | chr2 | 188425287 | 23 | 25 | 2192.247475 | 2245.898268 | 0.047463 |
| PHACTR1 | chr6 | 12717069 | 32 | 32 | 3020.009972 | 2907.285891 | 0.047678 |
| APOA1 | chr11 | 116726158 | 31 | 30 | 2947.512373 | 2744.647595 | 0.047958 |
| ADRA2C | chr4 | 3761372 | 32 | 30 | 2746.269954 | 2738.896761 | 0.048537 |
| LDLR | chr19 | 11221291 | 33 | 32 | 3186.732341 | 3004.793623 | 0.048744 |
| E2F6 | chr2 | 11613685 | 35 | 32 | 3312.092762 | 2937.045275 | 0.048802 |
| FOXP3 | chrX | 49117190 | 31 | 32 | 2571.896673 | 2880.247489 | 0.048879 |
| COL15A1 | chr9 | 101705919 | 7 | 5 | 190.8333333 | 102.2222222 | 0.049277 |
| GCK | chr7 | 44237017 | 38 | 32 | 3078.539383 | 2709.895073 | 0.049821 |
